# Integrative genomic reconstruction reveals heterogeneity in carbohydrate utilization across human gut bifidobacteria

**DOI:** 10.1101/2024.07.06.602360

**Authors:** Aleksandr A. Arzamasov, Dmitry A. Rodionov, Matthew C. Hibberd, Janaki L. Guruge, James E. Kent, Marat D. Kazanov, Semen A. Leyn, Marinela L. Elane, Kristija Sejane, Annalee Furst, Lars Bode, Michael J. Barratt, Jeffrey I. Gordon, Andrei L. Osterman

**Author notes:** Corresponding authors: Jeffrey I. Gordon Andrei L. Osterman.

## Abstract

Bifidobacteria are among the earliest colonizers of the human gut and are widely used as probiotics for their health-promoting properties. However, individual responses to probiotic supplementation may vary with strain type(s), microbiota composition, diet, or lifestyle conditions, highlighting the need for strain-level insights into the bifidobacterial metabolism of dietary and host glycans. Here, we systematically reconstructed 68 pathways involved in the utilization of mono-, di-, oligo-, and polysaccharides by analyzing the distribution of 589 curated metabolic functional roles (catabolic enzymes, transporters, transcriptional regulators) in 3083 non-redundant cultured *Bifidobacterium* isolates and metagenome-assembled genomes (MAGs) of human origin. Our analysis uncovered extensive inter- and intraspecies heterogeneity, including a distinct clade within the *Bifidobacterium longum* species capable of metabolizing α-glucans. We also identified isolates of Bangladeshi origin that harbor unique gene clusters implicated in the breakdown of xyloglucan and human milk oligosaccharides. Thirty-eight predicted carbohydrate utilization phenotypes were experimentally validated in 30 geographically diverse *Bifidobacterium* isolates *in vitro*. Our large-scale genomic compendium expands the knowledge of bifidobacterial carbohydrate metabolism and can inform the rational design of probiotic and synbiotic formulations tailored to strain-specific nutrient preferences.

---

Bifidobacteria are Gram-positive, saccharolytic microorganisms that predominantly inhabit animal gastrointestinal tracts^1^. Multiple *Bifidobacterium* species colonize the human gut throughout life, with dietary carbohydrate intake playing a key role in shaping this process^2^. Breastfeeding fosters the dominance of specific *Bifidobacterium* species within the neonatal gut microbiota due to their evolutionary adaptation to metabolize human milk oligosaccharides (HMOs)^3–5^. Weaning drives a gradual succession of bifidobacterial taxa from those tuned for HMO utilization to those more adapted to foraging dietary glycans (oligo- and polysaccharides) of plant origin^6–8^.

Geographic and cultural dietary differences also profoundly influence the bifidobacterial composition of neonatal microbiotas. For example, *Bifidobacterium longum* subsp. *infantis* (*Bl. infantis*), a specialist HMO utilizer, constitutes up to 90% of the gut microbial composition of healthy breastfed infants from non-Westernized populations^9–11^. Infants from Westernized populations often lack *Bl. infantis* and instead harbor less proficient HMO utilizers such as *Bifidobacterium longum* subsp. *longum* (*Bl. longum*), *Bifidobacterium breve*, and *Bifidobacterium pseudocatenulatum*; this appears to be due to a combination of lifestyle and cultural factors^9,12,13^.

Despite these differences, the predominance of bifidobacteria in gut communities is associated with multiple health benefits, particularly in infancy. Bifidobacterial fermentation products, lactate and acetate, can inhibit pathogen colonization^14,15^ and serve as substrates for cross-feeding among microbial community members^16,17^. Additionally, multiple *Bifidobacterium* species produce aromatic lactic acids modulating the immune system^18,19^. These beneficial traits underpin the widespread use of bifidobacterial strains as probiotics^20–23^, often supplemented with complementary prebiotics to facilitate engraftment^23,24^.

Further development is needed to rationally select probiotic strains and prebiotic glycans tailored for different populations. For instance, a probiotic *Bl. infantis* strain did not durably engraft in the microbiota of malnourished Bangladeshi infants whose diets were low in breastmilk compared to complementary foods^23^. Moreover, strains isolated from children in this population may harbor distinctive genomic adaptations for metabolizing glycans common in weaning diets^23^. Thus, strain-level insights into bifidobacterial carbohydrate metabolism, especially in understudied populations, may be instrumental in developing locally adapted pro- and synbiotics.

Genomics-based approaches, including the analysis of Carbohydrate Active Enzymes (CAZymes) repertoires^7,25–27^, genotype-to-phenotype matching^28–30^, and genome-scale metabolic models^31–33^, have been used to predict the carbohydrate utilization capabilities of bifidobacteria. However, functional gene annotations, particularly for glycan transporters, remain insufficiently curated and integrated into metabolic reconstructions^34^. In addition, current genomic resources may not fully capture the strain-level diversity within the *Bifidobacterium* genus.

We employed a combination of curated metabolic reconstruction and machine learning to map 68 carbohydrate utilization pathways across 263 reference and 2,820 additional *Bifidobacterium* genomes by analyzing the representation of 589 functional roles (catabolic enzymes, CAZymes, transporters, and transcriptional regulators). Thirty-eight predicted glycan utilization phenotypes were validated *in vitro* using 30 diverse strains. Our findings reveal remarkable inter- and intraspecies variability, including a distinct *Bifidobacterium longum* clade that metabolizes α-glucans and Bangladeshi strains carrying unique gene clusters for plant hemicellulose and HMO metabolism. These insights establish a genomic framework for predicting glycan utilization networks across *Bifidobacterium* lineages and support the development of targeted pre-, pro-, and synbiotics.

## RESULTS

### Curated metabolic reconstruction from reference genomes

The curated reconstruction encompassed a non-redundant reference set of 263 *Bifidobacterium* genomes of cultured isolates, including 19 newly sequenced genomes of Bangladeshi strains (**Supplementary Tables 1**,**2**). To refine taxonomic assignments, we first constructed a maximum-likelihood phylogenetic tree based on 487 core genes identified through a pangenome analysis (**Supplementary Fig. 1**). Additional pairwise average nucleotide identity (ANI) comparisons for 39 genomes allowed us to delineate the within-species structure of *Bifidobacterium longum* and *Bifidobacterium catenulatum* species (**Extended Data Fig. 1**; *Supplementary Note 1*). Overall, the reference set spanned 19 species and subspecies (**Supplementary Table 1**).

We leveraged a subsystem-based comparative genomics framework^35^ to reconstruct carbohydrate utilization pathways and predict associated metabolic phenotypes (**Fig. 1a**). We first mined 222 publications to identify 433 orthologous groups of carbohydrate utilization genes, whose encoded proteins were either experimentally characterized in bifidobacteria (210 metabolic functional roles), shared significant sequence similarity with proteins characterized in other microbial taxa (80 roles), or were previously assigned putative functions (143 roles). Genomic context analysis, including *in silico* transcriptional regulon reconstruction (*Supplementary Notes 2-3*), enabled tentative functional predictions for 156 additional gene groups involved in glycan metabolism. The resulting curated set of 589 roles — comprising 235 components of glycan-specific transporters, 197 catabolic CAZymes, 72 downstream catabolic enzymes, and 85 transcription factors — was used to functionally annotate 39,589 of 541,418 protein-coding genes across 263 reference genomes (**Supplementary Tables 4**,**8**). Manual curation improved 76.6% and 69% of annotations over Prokka and EggNOG-mapper, respectively, including more than 90% of annotations for transporters and transcriptional regulators (**Fig. 1b)**. The metabolic reconstruction also captured 82.2% of catabolic CAZymes identified by dbCAN (**Fig. 1c**).

**Fig. 1.**
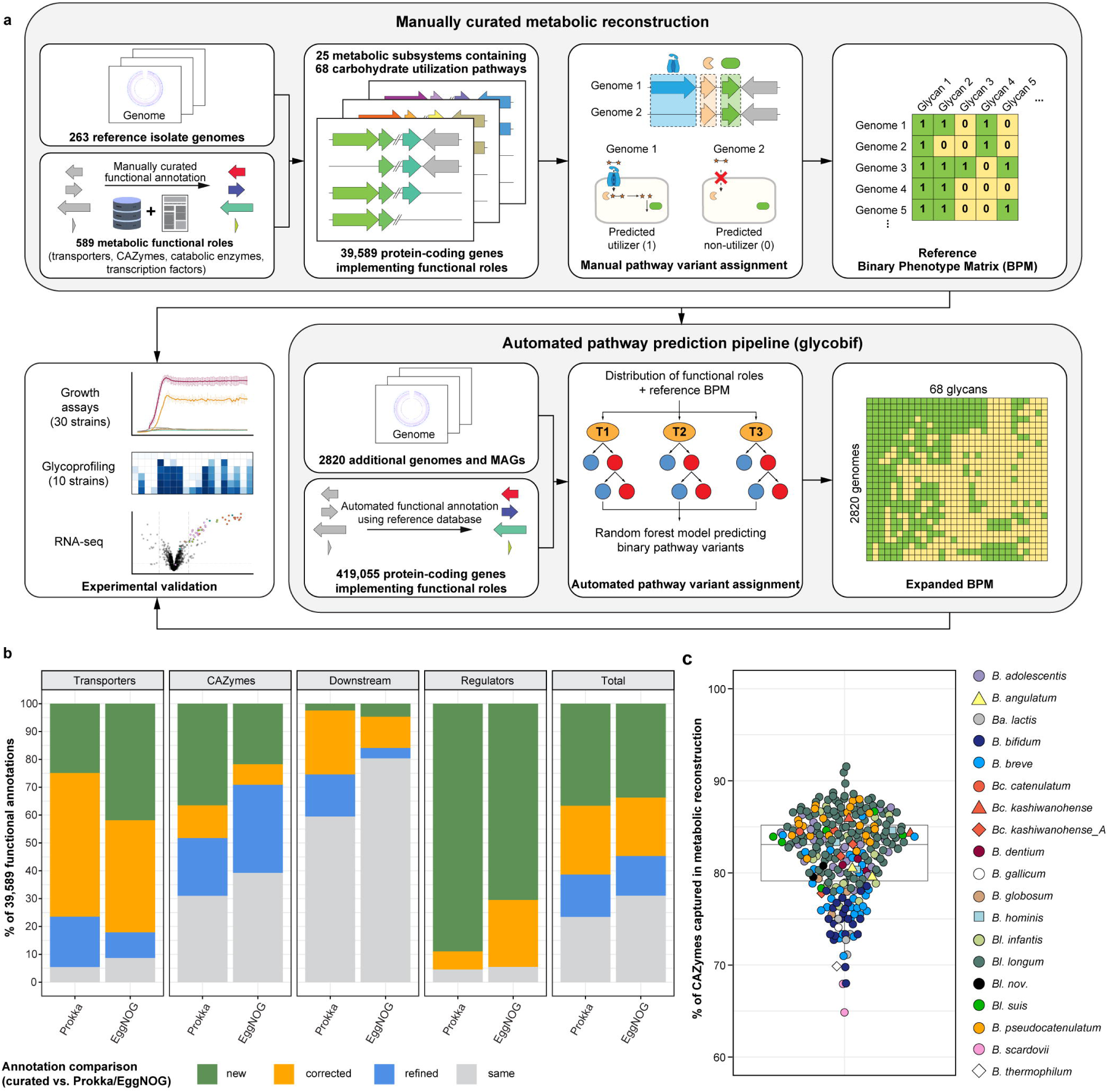
*In silico* reconstruction of carbohydrate metabolism in bifidobacteria. **a**, Overview of the computational framework. We analyzed the distribution of 589 metabolic functional roles (glycan transporters, CAZymes, downstream catabolic enzymes, transcription factors) across 263 reference *Bifidobacterium* genomes. Manual reconstruction of 68 carbohydrate utilization pathways enabled the assignment of binary pathway variants corresponding to predicted glycan utilization phenotypes based on specific genomic signatures (see **Extended Data Fig. 2**). This curated dataset was used to train an automated pathway prediction pipeline (glycobif), which outputs: (i) the distribution of functional roles across additional 2,820 *Bifidobacterium* genomes and (ii) a Binary Phenotype Matrix (BPM) reflecting the inferred presence or absence of pathways in each genome. Predicted phenotypes for 20 reference and 10 additional strains were compared with *in vitro* growth profiles to evaluate prediction accuracy. **b,** Comparison of functional gene annotations obtained via manual curation versus automated tools in the 263 reference *Bifidobacterium* genomes. Stacked bar plots show the distribution of 39,589 manually curated annotations across four categories: *new* = annotations that update non-specific predictions (e.g., hypothetical protein); *corrected* = annotations that replace specific but incorrect predictions; *refined* = annotations that add functional precision (e.g., substrate and linkage specificity for GHs); *same* = annotations that are essentially identical to automated output (**Supplementary Table 13**). **c**, Percentage of catabolic CAZymes (glycoside hydrolases, carbohydrate esterases, polysaccharide lyases) captured in the reconstructed metabolic pathways across the 263 reference *Bifidobacterium* genomes. Each point represents a genome

The genome-wide distribution of genes assigned to 589 functional roles was used to reconstruct 68 catabolic pathways: 18 for monosaccharides and their derivatives (sugar alcohols and acids), 39 for di- and oligosaccharides, and 11 for polysaccharides (**Supplementary Table 5**). To link gene conservation patterns with metabolic phenotypes, we first established rules that define genomic signatures distinguishing metabolic pathway variants (*Methods*; **Extended Data Fig. 2**; **Supplementary Table 6**). Each pathway variant was then converted into a binary phenotype, classifying strains as predicted utilizers (“1”) or non-utilizers (“0”) of specific carbohydrates. These assignments formed a Binary Phenotype Matrix (BPM), summarizing the predicted utilization profiles of 263 reference *Bifidobacterium* strains spanning 68 glycans (**Supplementary Table 7**). To evaluate the robustness of our reconstruction, we compared predicted phenotypes with *in vitro* growth data for 33 strains from previous studies^8,28,36^, yielding 94% accuracy (**Supplementary Table 17a–d**).

### Automated pathway prediction across large genomic datasets

We utilized the metabolic reconstruction for 263 reference *Bifidobacterium* genomes to analyze the representation of glycan utilization pathways in an additional set of 2,820 non-redundant genomes (**Supplementary Table 3**). This set included 364 isolate genomes and 2,456 high-quality metagenome-assembled genomes (MAGs) with completeness ≥97 % and contamination ≤3%. Twelve genomes represented four species absent from the reference collection, and nine were from Bangladeshi and Malawian strains isolated in previous studies^37–39^ and sequenced in this work (**Supplementary Table 1**). We used annotated protein sequences of reference strains to assign functions to a subset of 419,055 protein-coding genes in the 2,820 genomes. These functional annotations, combined with the reference BPM, were used to train a random forest model that predicted the presence of 68 reconstructed carbohydrate utilization pathways (*Methods*; **Fig. 1a**; **Supplementary Tables 9**,**10**).

### Interspecies diversity in glycan utilization

The BPMs for 263 reference and 2,820 additional genomes were merged to assess the distribution of glycan utilization pathways across the *Bifidobacterium* genus. Non-metric multidimensional scaling (NMDS) of the Hamming distance matrix derived from the BPM of 3,083 genomes revealed grouping by species/subspecies, with taxonomy explaining 91% of the variation (PERMANOVA R² = 0.91, P = 0.001; **Fig. 2a**). A significant dispersion effect (F = 81.06, P= 0.001) indicated that strain-level variability in pathway representation differed across taxa, likely influencing the separation.

**Fig. 2.**
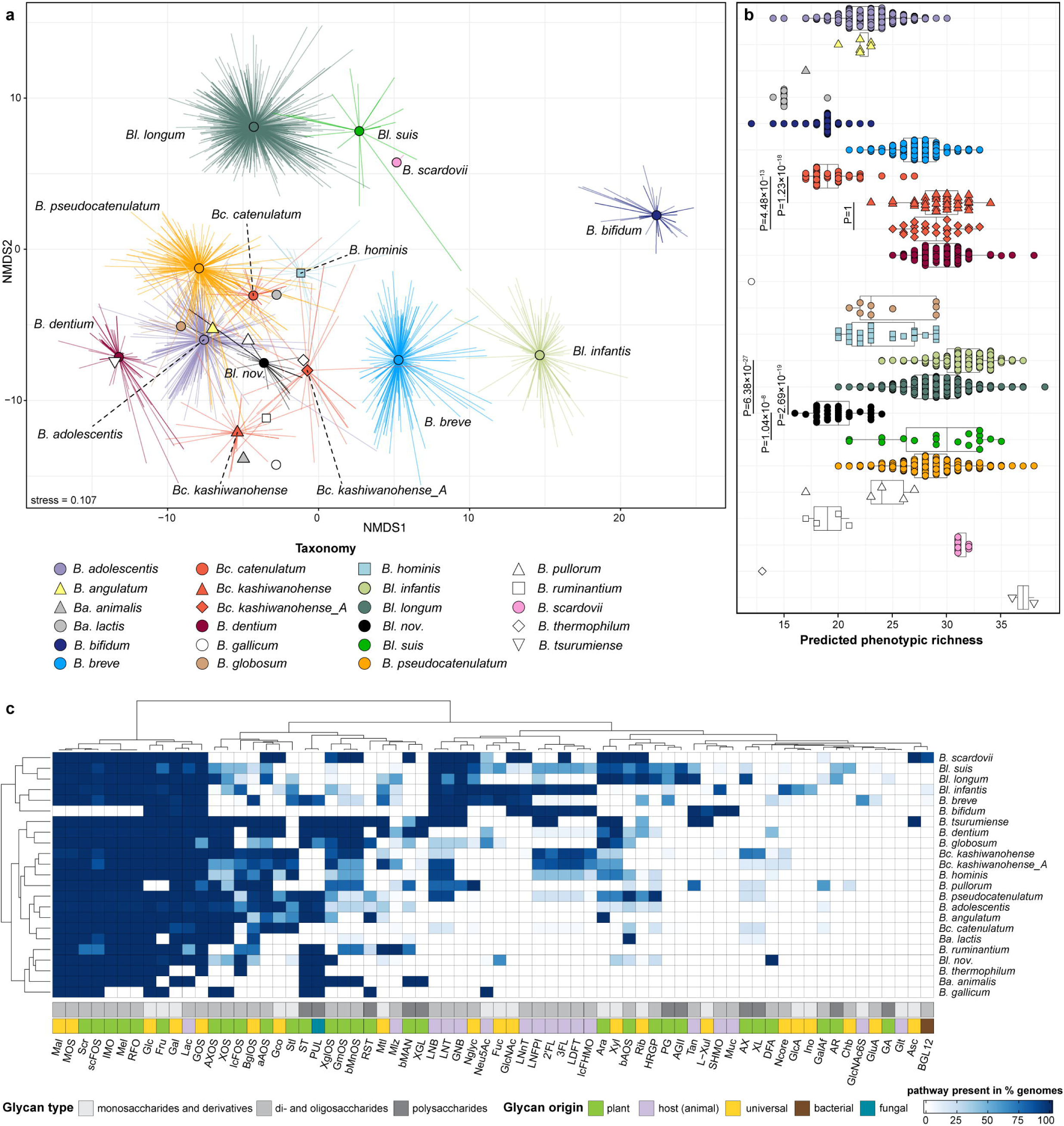
Representation of carbohydrate utilization pathways across 3,083 *Bifidobacterium* genomes. **a,** Non-metric multidimensional scaling (NMDS) of a Hamming distance matrix derived from the presence/absence patterns of 68 predicted carbohydrate utilization pathways across 627 isolate genomes plus 2,456 metagenome assembled genomes (MAGs). Points represent genomes; spider lines connect genomes to their group (taxon) centroid. Colors and shapes of centroids indicate taxonomic groups. **b**, Predicted phenotypic richness (the total number of carbohydrate utilization pathways) at species and strain levels. Each point represents a genome. Box plots show the median (center line), interquartile range (IQR; box bounds), and full data range excluding outliers (whiskers, defined as 1.5× IQR). Statistical comparisons were performed using a two-sided generalized linear model (GLM) with a Poisson distribution, followed by post hoc pairwise comparisons with Bonferroni correction. Sample size (n) per group corresponds to the number of genomes analyzed. **c,** Heatmap of the proportion of genomes within each taxon encoding the 68 predicted carbohydrate utilization pathways. Color intensity indicates the percentage of genomes that encode each pathway. Annotation rows at the bottom indicate pathway/phenotype classifications. Full names are provided in **Supplementary Table 5**.

We observed significant differences in predicted phenotypic richness (total number of predicted carbohydrate utilization pathways) between taxa, including phylogenetically close subspecies sharing over 95% ANI (Poisson generalized linear model, p < 2.2 × 10 ^16^; **Fig. 2b**). For example, *Bifidobacterium catenulatum* subsp. *catenulatum* (*Bc. catenulatum*) had significantly lower phenotypic richness than subspecies *kashiwanohense* (*Bc. kashiwanohense*) and *kashiwanohense_A* (*Bc. kashiwanohense_A*) due to the absence of utilization pathways for fucosylated HMOs (e.g., 2’- and 3-fucosyllactose, 2’FL/3FL), and certain plant oligosaccharides (e.g., beta-mannose oligosaccharides, bMnOS) in >95% of genomes (**Extended Data Fig. 3**; *Supplementary Note 4*).

Pathways for glucose (Glc), galactose (Gal), fructose (Fru), lactose (Lac), and galactooligosaccharide (GOS) utilization were identified in over 98% of analyzed genomes, defining the core catabolic potential of human-colonizing bifidobacteria (**Fig. 2c**). Pathways for sucrose (Scr), maltose (Mal), malto- and isomaltooligosaccharide (MOS and IMO), melibiose (Mel), raffinose-family oligosaccharide (RFO), and short-chain fructooligosaccharide (scFOS) metabolism were encoded in over 84% of genomes, indicating broad conservation across most species, except *Bifidobacterium bifidum*. Other pathways showed more sporadic distribution patterns, reflecting species-level adaptations for metabolizing dietary glycans of different origins and structures. For example, *B. bifidum* exhibited a notable specialization towards the metabolism of host mucin *O*-glycans, HMOs, and their degradation products lacto-*N*-biose (LNB), galacto-*N*-biose (GNB), and *N*-acetyl-D-glucosamine (GlcNAc) — consistent with prior reports^40^ (**Fig. 2c**). At the same time, most *B. bifidum* genomes lacked complete pathways for utilizing plant-derived mono-, di-/oligo- and polysaccharides. Other bifidobacteria, including the specialist HMO utilizer *Bl. infantis,* were more versatile in their predicted glycan utilization profiles, although the pathways for plant polysaccharide degradation were generally underrepresented compared to those for their mono- and oligosaccharide components (**Fig. 2c**).

Hierarchical clustering of the BPM for 263 reference genomes showed a moderate correlation with core-gene phylogeny (cophenetic correlation = 0.58, permutation test, p < 0.001; **Supplementary Fig. 2**), indicating incomplete concordance between predicted glycan utilization capabilities and phylogenetic relatedness. For example, the predicted phenotypic profiles of *Bl. infantis* and *Bl. longum*, two phylogenetically related subspecies within the *B. longum* species, were markedly different. The representation of glycan utilization pathways in *Bl. infantis* more closely resembled that of *B. breve* — a more distantly related species inhabiting the neonatal human gut. Given the importance of *B. longum* in infant microbiota development, we next conducted a more focused analysis of pathway variability within this heterogeneous species.

### Diversity of glycan metabolism within the *B. longum* species

The *B. longum* species comprises multiple subspecies distinguished by phylogeny and specific phenotypic traits^26,41–43^. Our phylogenomic and ANI analyses clustered *B. longum* genomes into three clades matching previously described subspecies — *infantis* (*Bl. infantis*), *longum* (*Bl. longum*), *suis* (*Bl. suis*) — and a distinct clade hereafter referred to as *Bl. nov.* (**Extended Data Fig. 1a**; **Supplementary Fig. 1**; *Supplementary Note 1*). *Bl. nov.* exhibited significantly lower predicted phenotypic richness than other subspecies (**Fig. 2b**), lacking pathways for LNB/GNB, *N*-glycan, HMO, and T-antigen (Tan) metabolism (**Fig. 2c**; *Supplementary Note 5*). Conversely, only *Bl. nov.* genomes encoded extracellular amylopullulanase ApuB (GH13_14_32), a bifunctional glycoside hydrolase (GH) cleaving both α-1,4 and α-1,6-glycosidic bonds in soluble starch (ST) and pullulan (PUL)^44^, and a pathway for difructose dianhydride (DFA) metabolism^45^. These findings suggest that *Bl. nov.* has a reduced capacity to metabolize host-derived glycans but can degrade α-glucans of plant and fungal origin.

Comparative analysis of carbohydrate utilization pathways revealed stark differences between *Bl. infantis* and *Bl. longum* (**Fig. 2c**). Consistent with previous studies, *Bl. infantis* genomes were distinguished by the presence of H1^46^ and FL1/2^4^ gene clusters, which enable the utilization of lacto-*N*-neotetraose (LNnT), 2’FL, 3FL, lactodifucotetraose (LDFT), lacto-*N*-fucopentaose I (LNFP I), and sialylated HMOs (SHMOs; **Supplementary Tables 7–10**). Predicted HMO utilization potential of *Bl. longum* was more limited: 35% of genomes encoded an extracellular lacto-*N*-biosidase (LnbX; GH136) that cleaves lacto-*N*-tetraose (LNT) and LNFP I^47^, and only 2.3% harbored a gene cluster for intracellular utilization of 2’/3FL, LDFT, and LNFP I^48,49^. Beyond HMO metabolism, *Bl. infantis* exclusively encoded utilization pathways for metabolizing glucuronate (GlcA), inositol (Ino), and gluconate (Gco) in 63%, 48%, and 23% of genomes, respectively (**Extended Data Fig. 4a,b,d,e**; *Supplementary Note 3*) *Bl. longum* genomes, by contrast, commonly encoded pathways for L-arabinose (Ara), α/β-arabinooligosaccharides (aAOS/bAOS), type II arabinogalactan (AGII), arabinan (AR), arabinoxylan (AX), and host- or plant-derived *O*-glycans (Tan, HRGP) (**Fig. 2c**). These findings illustrate an ecological divergence between *Bl. infantis* and *Bl. longum* shaped by their respective adaptations to thrive on milk glycans during breastfeeding versus plant-derived carbohydrates after weaning.

Predicted glycan utilization profiles within the *Bl. suis* group were highly heterogeneous. Most genomes encoded a set of pathways similar to that of *Bl. longum*, yet more frequently included pathways for *N*-acetylneuraminic acid (Neu5Ac; 44% vs. 2%), L-fucose (Fuc; 61% vs. 1.7%), and fucosylated HMOs (2’/3FL, LDFT, LNFP I; 56% vs. 2.3%), while lacking extracellular α-L-arabinofuranosidases required for AX degradation^50,51^ (**Fig. 2c**; **Supplementary Tables 8**,**10**). In contrast, the Bangladeshi strain Bg131.S11_17.F6 exhibited a *Bl. infantis*-like pathway profile (**Supplementary Figure 3**), carrying the H1 gene cluster for the utilization of multiple HMOs but lacking pathways for arabinose-containing glycans (e.g., Ara, aAOS, AGII). Consequently, this strain was predicted to utilize more distinct HMO structures (e.g., LNnT, SHMOs) than other Bangladeshi *Bl. suis* strains, while having a reduced capacity for plant glycan metabolism. Analysis of 10 additional *Bl. suis* genomes of animal origin confirmed the uniqueness of the Bg131.S11_17.F6 strain and further underscored the phenotypic heterogeneity within this group (**Extended Data Fig. 5**; *Supplementary Note 5*). Collectively, these findings reveal pronounced differences in carbohydrate utilization across *B. longum* subspecies and underscore pervasive strain-level heterogeneity within each clade, a phenomenon explored in detail below.

### Strain-level heterogeneity of glycan metabolism

Beyond interspecies differences, we observed extensive strain-level variability in predicted carbohydrate utilization capabilities. Of the 68 reconstructed pathways, 66 exhibited variability within at least one taxonomic group (**Fig. 2c**; **Supplementary Tables 7**,**9**). The genomic differences driving this heterogeneity ranged from individual genes encoding extracellular GHs enabling polysaccharide degradation to multi-gene clusters comprising up to 20 genes encoding complete metabolic pathways (**Supplementary Table 8**). In contrast, biosynthetic pathways for essential metabolites, such as amino acids and B vitamins, were largely conserved. A few exceptions included riboflavin (B2) biosynthesis, which varied across *Bl. suis* and *Bc. kashiwanohense_A*, and thiamine (B1) and niacin (B3) biosynthesis in *Bifidobacterium adolescentis* (**Extended Data Fig. 6**; **Supplementary Table 11**).

The observed heterogeneity suggests that while most *Bifidobacterium* taxa follow general ecological strategies centered on the utilization of specific core glycans, individual strains can exhibit substantial metabolic tuning. For instance, *Bl. infantis* Bg064.S07_13.C6 harbored pathways for xylooligosaccharide (XOS) and long-chain fructooligosaccharide (lcFOS) utilization, suggesting an enhanced capacity to metabolize dietary plant glycans compared to most other *Bl. infantis* strains (**Supplementary Fig. 3**). Conversely, several *B. adolescentis* isolates carried pathways for the utilization of LNB/GNB, *N*-glycans (strains AF96-10M2bTA, UN03-88), and fucosylated HMOs (M56B_1C3), highlighting the presence of traits characteristic of infant-adapted taxa in a species commonly associated with the adult gut (**Supplementary Tables 9,10**). Additional examples of notable strain-specific genomic features related to carbohydrate utilization were found in Bangladeshi isolates and are detailed below.

### Unique glycan utilization features of Bangladeshi strains

Our previous study identified distinctive genomic features of Bangladeshi *Bl. infantis* strains related to *N*-glycan and beta-glucoside catabolism^23^. Here, we identified a distinct gene cluster (*xgl*) in *Bc. kashiwanohense* Bg42221_1E1 and *Bc. kashiwanohense_A* Bg42221_1D3, two isolates from a Bangladeshi infant (**Fig. 3a**). This cluster encoded multiple GHs (families 3, 5_4, 29, 31, 42, 43_12), an unclassified carbohydrate esterase (CE), a β-glucoside kinase, and an ABC transport system, which together may catabolize plant hemicelluloses such as xyloglucans (XGL). The reconstructed XGL pathway involved the hydrolysis of the XGL backbone to oligosaccharides by extracellular endo-β-1,4-glucanase Xgl5A (GH5_4), which shares catalytic and glycan-binding residues with xyloglucanase PpXG5 from *Paenibacillus pabuli* XG5^52^ (**Extended Data Fig. 7a**). Released oligosaccharides would be imported into the cell and degraded sequentially to individual monosaccharides and cellobiose by the coordinated action of GHs and CEs via a mechanism similar to that described in *Ruminiclostridium cellulolyticum*^53^ (**Fig. 3b**). Regulon reconstruction of XglT, a putative TetR-family transcription factor, suggested potential co-regulation of the *xgl* cluster and *cbpA*, which encodes a GH94-family cellobiose phosphorylase (**Fig. 3a-c**; **Supplementary Table 19**). The *xgl* cluster was identified in only 3 of 110 studied *B. catenulatum* genomes but was conserved in *Bifidobacterium dentium* and *Bifidobacterium tsurumirense*.

**Fig. 3.**
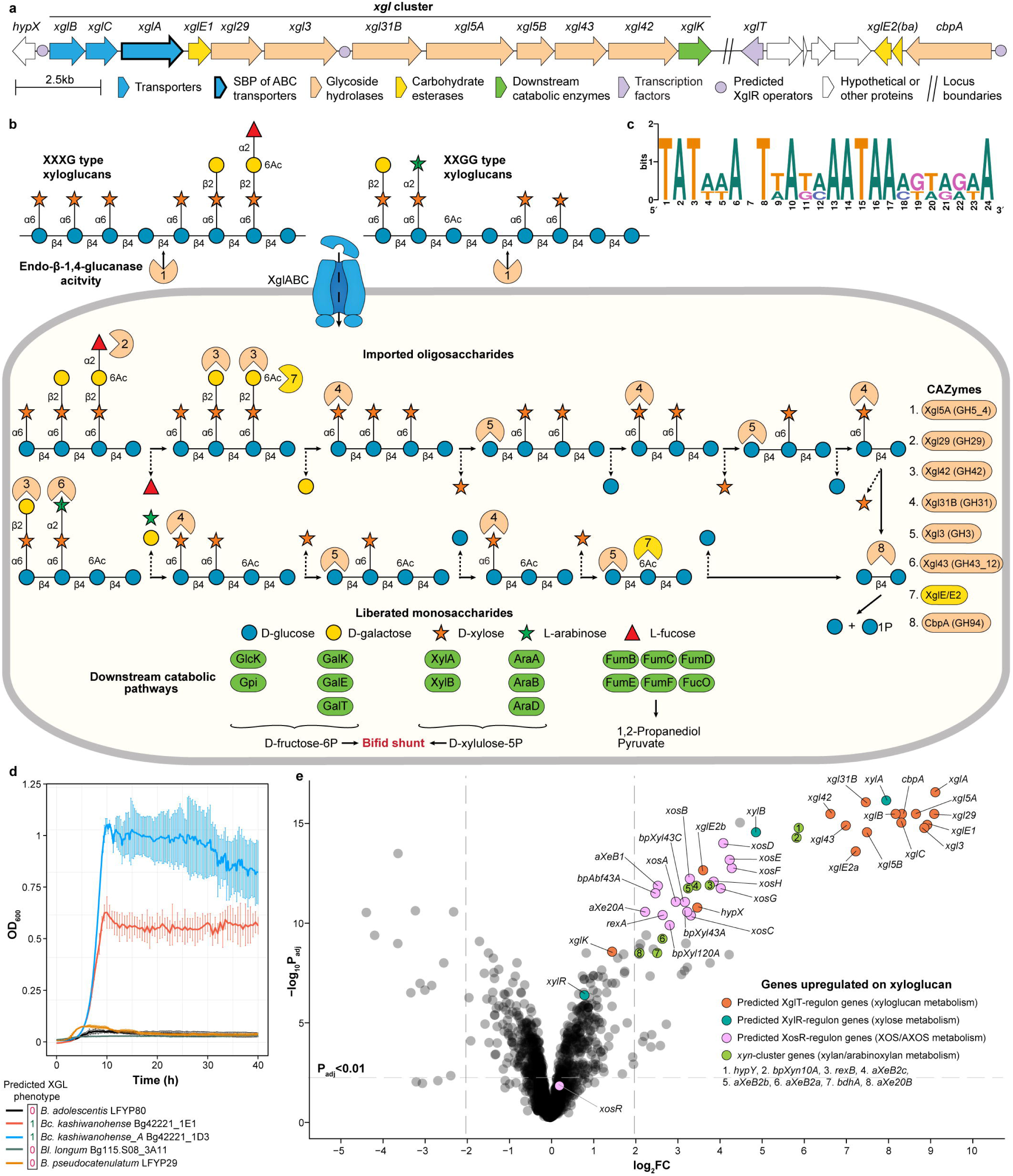
Integrated genomic and transcriptomic analysis of xyloglucan metabolism in *Bc. kashiwanohense* Bg42221_1E1. **a**, Gene clusters potentially driving xyloglucan degradation (phenotype XGL) in *Bc. kashiwanohense* Bg42221_1E1. **b,** Reconstructed xyloglucan degradation pathway in *Bc. kashiwanohense* Bg42221_1E1: (i) xyloglucan chains are potentially cleaved by extracellular endo-β-1,4-glucanase Xgl5A, (ii) released oligosaccharides are imported and metabolized inside the cell by a coordinated action of CAZymes and downstream catabolic enzymes. **c,** Predicted DNA-binding motif of the XglT transcriptional regulator potentially controlling xyloglucan metabolism genes. **d,** Growth curves of *Bifidobacterium* strains in MRS-AC supplemented with 0.5% tamarind xyloglucan. Data represent the mean ± s.d. of three biological replicates. **e,** Volcano plot showing differential gene expression (log_2_ fold change vs. −log_10_ adjusted P-value) in *Bc. kashiwanohense* Bg42221_1E1 grown in MRS-AC with tamarind xyloglucan (XGL) versus MRS-AC with lactose (Lac). Differential expression was assessed using moderated two-sided t-tests with empirical Bayes variance moderation. P-values were adjusted for multiple comparisons using the Benjamini–Hochberg procedure. Genes were considered differentially expressed at Padj < 0.01 and |log_2_FC| > 2. Genes belonging to the reconstructed XglT, XylR, and XosR regulons, as well as *xyn* cluster genes, are highlighted. Exact log FC values, test statistics, and adjusted P-values are provided in **Supplementary Table 21a**.

Another gene cluster, unique to *Bc. kashiwanohense* Bg42221_1E1, contained orthologs of H1 cluster HMO utilization genes from *Bl. infantis* and the “outlier” Bangladeshi *Bl. suis* Bg131.S11_17.F6 strain (**Fig. 4a**). The H1 variant identified in Bg42221_1E1 encoded two ABC transporters, one of which (HmoABC) is likely involved in the uptake of LNnT and other type II HMO structures^54^. Additionally, this cluster encoded orthologs of characterized β-*N*-acetylglucosaminidase (GH20), two α-fucosidases (GH29 and GH95), and α-sialidase (GH33), which mediate intracellular HMO hydrolysis^55–57^, along with downstream catabolic pathways for GlcNAc and Fuc. Regulon reconstruction suggested the potential transcriptional regulation of H1 cluster genes in *Bc. kashiwanohense* Bg42221_1E1 by NagR, a GlcNAc-responsive ROK-family repressor^58^ (**Fig. 4a**; **Supplementary Table 19**).

**Fig. 4.**
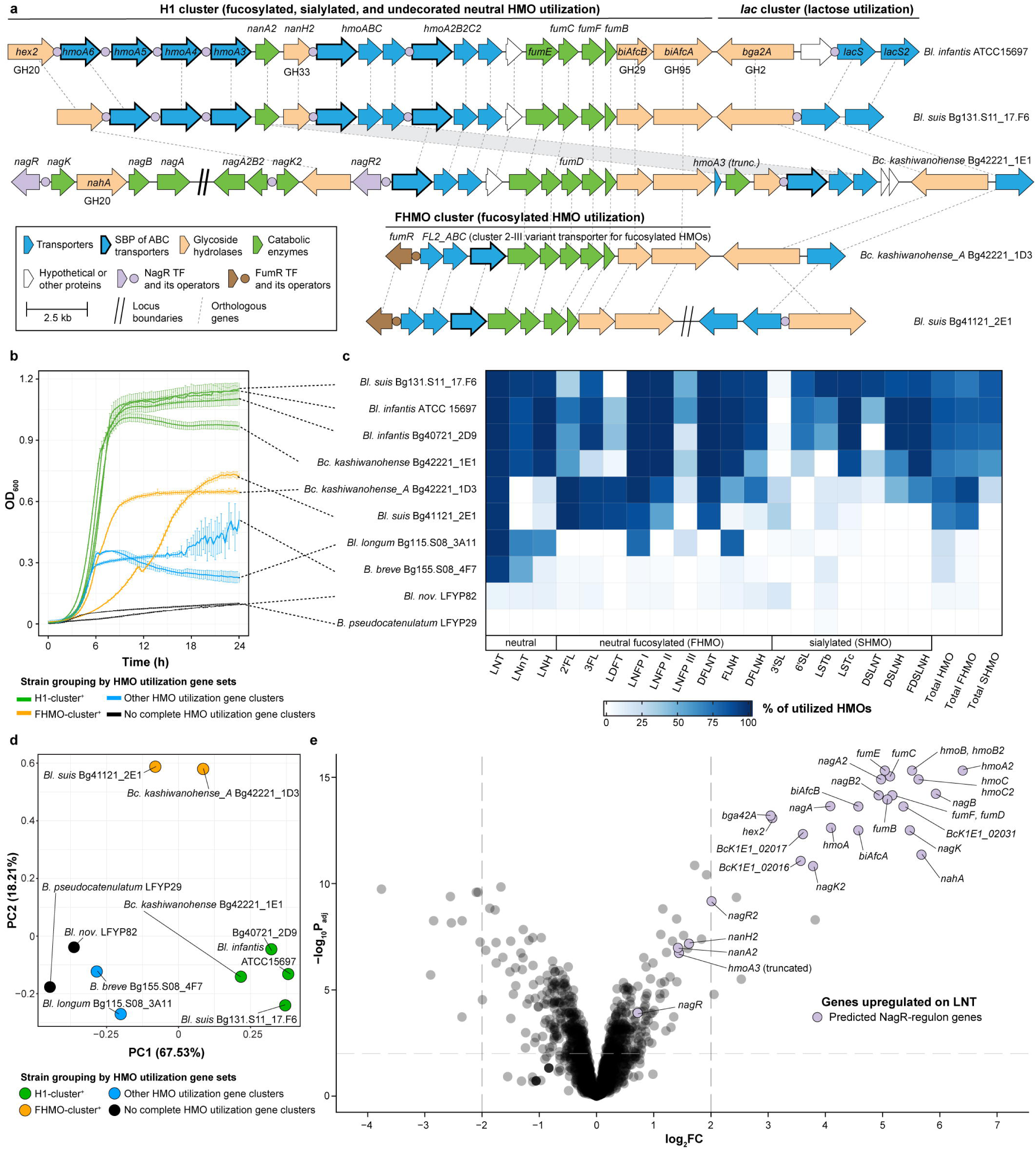
Comparative genomic and functional profiling of human milk oligosaccharide (HMO) utilization capabilities in *Bifidobacterium* strains. **a**, Schematic representation of HMO utilization genes in selected Bangladeshi *Bifidobacterium* strains and their homology to H1 cluster genes from *Bl. infantis* ATCC 15697. **b,** Growth curves of *Bifidobacterium* strains in MRS-AC supplemented with pooled HMOs. Data represent the mean ± s.d. of three biological replicates. Curves are color-coded based on the presence of specific HMO utilization gene clusters in the respective strains. **c,** HPLC-based quantification of HMO depletion from culture supernatants after 24 h. Data represent the percentage of utilized HMOs (mean of three biological replicates) relative to the medium control. Total HMO = total HMO utilized; total FHMO = total fucosylated HMO utilized; total SHMO = total sialylated HMO utilized. Concentrations of individual HMOs (nmol/mL) are provided in **Supplementary Table 18b**. **d**, Principal Component Analysis (PCA) based on combined growth metrics (maximum OD_600_, area under the curve, and maximum growth rate) and percent utilization of individual HMOs. Each point represents a strain, color-coded by the presence of specific HMO utilization gene clusters. **e,** Volcano plot showing differential gene expression (log_2_ fold change vs. −log_10_ adjusted P-value) in *Bc. kashiwanohense* Bg42221_1E1 grown in MRS-AC with lacto-*N*-tetraose (LNT) versus MRS-AC with lactose (Lac). Differential expression was assessed using moderated two-sided t-tests with empirical Bayes variance moderation. P-values were adjusted for multiple comparisons using the Benjamini–Hochberg procedure. Genes were considered differentially expressed at Padj < 0.01 and |log_2_FC| > 2. Genes belonging to the reconstructed NagR regulon are highlighted. Exact log FC values, test statistics, and adjusted P-values are provided in **Supplementary Table 21b**.

Other notable features of Bangladeshi *Bifidobacterium* isolates included a putative D-galactonate utilization pathway found exclusively in *B. breve* Bg41721_1C11 (**Extended Data Fig. 4d,e**; *Supplementary Note 3*). This strain, along with *B. breve* Bg131.S11_D6, also lacked the *nan* gene cluster, which encodes a well-characterized Neu5Ac catabolic pathway^59^ (**Extended Data Fig. 4f,g**). This absence of the *nan* cluster in these two strains was unexpected, given its previously reported broad conservation in *B. breve* genomes^29,59^ and its established role in utilizing Neu5Ac via cross-feeding on sialylated HMOs and mucin *O*-glycans degraded by *B. bifidum*^59,60^.

These results demonstrate that Bangladeshi *Bifidobacterium* strains carry unique adaptations for metabolizing dietary plant polysaccharides and HMO, while lacking pathways conserved in well-characterized strains. These traits may reflect strain-level adaptation to the diet and lifestyle of Bangladeshi children. We next investigated whether broader patterns of bifidobacterial carbohydrate utilization are associated with host age and lifestyle across diverse human populations.

### Associations between pathway profiles and lifestyle

We examined the enrichment of glycan utilization pathways in *Bifidobacterium* genomes from different populations based on: (i) host age/stage of gut microbiota maturation (< 3 years: infant/transitional; ≥ 3 years: adult-like), and (ii) host lifestyle (“Westernized” vs. “non-Westernized” as defined in Pasolli et al.^61^). Host glycan utilization pathways (e.g., for HMO) were enriched in genomes from the “age < 3” group, whereas plant glycan utilization pathways were more prevalent in adult-associated (≥ 3 years) genomes across both lifestyle groups (Fisher’s exact test, P_adj_ ≤ 0.01; **Extended Data Fig. 8a,b**). Within the “age < 3” group, 14 pathways (11 for plant glycans) were enriched in the Westernized group and 24, including 11 for HMOs and their constituent blocks LNB, Neu5Ac, and Fuc in the non-Westernized group (**Extended Data Fig. 8c**). These differences likely stem from the uneven taxonomic distribution of species and subspecies across Westernized and non-Westernized microbiota. For example, pathways enriched in the non-Westernized group were associated with *Bl. infantis*, a taxon more prevalent in that group (odds ratio = 4.98, 95% CI: 3.24–7.51, Fisher’s exact test, P = 1.82×10^−^11).

Within taxa, pathways for sorbitol, mannitol (Stl and Mtl), lcFOS, and pectic galactan (PG) metabolism were significantly enriched in Westernized *Bl. infantis* genomes (**Extended Data Fig. 8d**). The Stl and lcFOS utilization pathways were also enriched in Westernized *B. adolescentis* genomes, and the PG pathway was enriched in Westernized *Bl. longum* genomes. Conversely, non-Westernized *B. breve* genomes were enriched for melezitose (Mlz) and 1,2-β-oligoglucan utilization pathways but more frequently lacked the Neu5Ac pathway. These findings underscore how lifestyle-driven ecological pressures shape glycan utilization strategies among bifidobacteria. We next tested carbohydrate utilization phenotypes in representative isolates to validate predictions from the reconstruction framework.

### Growth-based validation of predicted glycan utilization

To experimentally test *in silico* carbohydrate utilization predictions, we conducted *in vitro* growth assays on 30 geographically diverse *Bifidobacterium* strains (15 Bangladeshi, 8 Malawian, 5 US, two European). Growth was tested on 43 substrates corresponding to 38 predicted glycan utilization phenotypes: 13 monosaccharides and derivatives, 18 di-/oligosaccharides, and 7 polysaccharides (**Fig. 5a**). Strains were cultured in a sugar-free De Man-Rogosa-Sharpe (MRS-AC) medium^4,58^ supplemented with the test substrate (5-10 mg/mL), and growth was defined using strain-specific OD_600_ thresholds (*Methods*; **Supplementary Fig. 4**). Growth outcomes were compared with predictions derived from manual curation (20 strains) or the automated pipeline (10 strains; **Fig. 5b**).

**Fig. 5.**
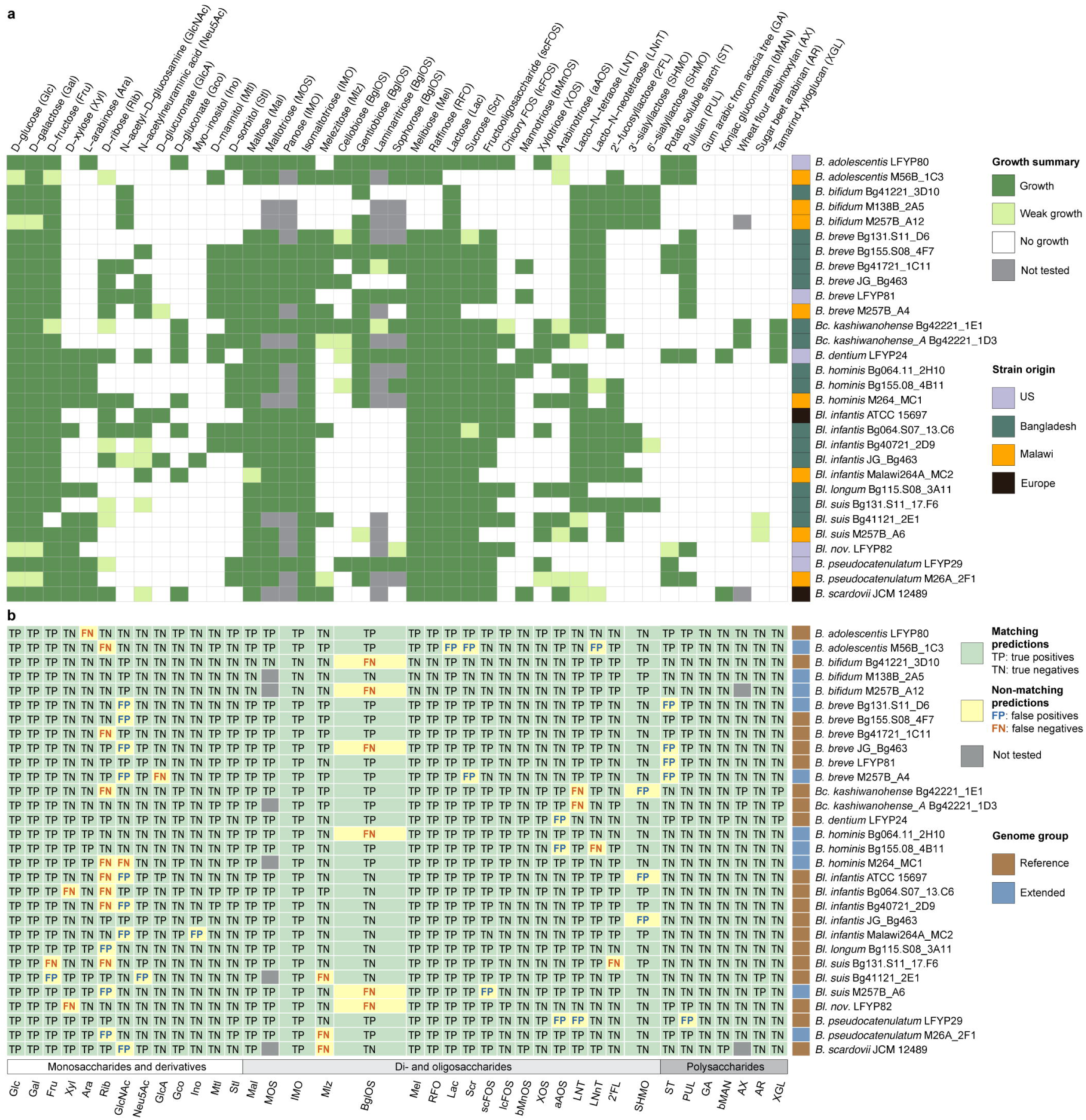
Comparison of predicted carbohydrate utilization phenotypes with *in vitro* growth data. **a**, Summary of *in vitro* growth profiles of 30 *Bifidobacterium* strains on 43 substrates. Growth, weak growth, and no growth were categorized based on strain-specific OD_600_ thresholds (*Methods*). The row annotation shows the geographical origin of each strain. The details about substrates are given in **Supplementary Table 15**; all growth curves are shown in **Supplementary Fig. 4**; raw OD_600_ data are provided in **Supplementary Table 16**. **b**, Comparison of 38 predicted carbohydrate utilization phenotypes with corresponding *in vitro* growth profiles for the same 30 strains. Predicted phenotypes were tested using a singular substrate, except IMO (panose and isomaltotriose), BglOS (cellobiose, gentiobiose, laminaritriose, and sophorose), and SHMO (3’-sialyllactose, 6’-sialyllactose). Prediction outcomes are color-coded to indicate agreement between predicted and observed phenotypes. The row annotation indicates whether genomes belong to the reference or extended dataset. The column annotation shows phenotype classification (glycan type). Summary data are provided in **Supplementary Table 17f**; full names of abbreviations are provided in **Supplementary Table 5**.

Prediction accuracy was similar for both approaches: 95 and 94%, respectively, with Matthews correlation coefficients of 0.9 and 0.89 (**Supplementary Table 17e,f**). False negative predictions (growth despite predicted non-utilization) likely stemmed from incomplete knowledge about monosaccharide and HMO transporters. For example, *Bl. suis* Bg131.S11_17.F6 grew on 2’FL despite lacking known 2’FL transporter genes^4,5^ (**Supplementary Table 8**). Some false positive predictions (predicted utilization but no growth) may have resulted from gene disruptions, such as a premature stop codon in the *gltA* gene in *B. pseudocatenulatum* LFYP29, which likely impaired the LNT transport function.

As predicted, *Bl. nov.* LFYP82 was the only *B. longum* strain to grow on ST and PUL (**Fig. 5a**). We also validated strain-specific utilization of short- and long-chain fructooligosaccharides (scFOS and lcFOS), XOS, and GlcA in *Bl. infantis* (**Fig. 5b; Extended Data Fig. 4c**). Similarly, while two *B. breve* strains grew on mannotriose (bMnOS), none grew on konjac glucomannan (bMAN), in contrast to *B. dentium* LFYP24 and *Bifidobacterium scardovii* JCM 12489, which harbored extracellular endo-β-1,4-mannanases (**Extended Data Fig. 7e-g**; *Supplementary Note 2*).

We further validated several unusual glycan utilization traits predicted in isolates from Bangladesh and Malawi. *Bc. kashiwanohense* Bg42221_1E1 and *Bc. kashiwanohense_A* Bg42221_1D3, both carrying the *xgl* cluster, grew in the medium supplemented with tamarind XGL (**Fig. 3d**). Bangladeshi *B. breve* isolates lacking the Neu5Ac utilization pathway (Bg41721_1C11 and Bg131.S11_D6) failed to grow on Neu5Ac, unlike other tested *B. breve* strains (**Extended Data Fig. 4h**). Finally, consistent with our prediction, *B. adolescentis* M56B_1C3 grew on 2’FL, highlighting the previously unrecognized capacity of this species (**Fig. 5a**).

### Glycoprofiling of HMO utilization

The H1 cluster in *Bl. infantis* is believed to enable the metabolism of multiple HMOs^46^, though the precise range of structures cannot be confidently predicted due to limited understanding of the functions of individual transporter genes. Given the presence of homologous H1 clusters in *Bl. suis* Bg131.S11_17.F6 and *Bc. kashiwanohense* Bg42221_1E1, we compared their HMO utilization with that of (i) two *Bl. infantis* strains, (ii) phylogenetically related strains carrying a fucosylated HMO utilization gene cluster (FHMO) instead of H1 (*Bl. suis* Bg41121_2E1 and *Bc. kashiwanohense_A* Bg42221_1D3; **Fig. 4a**), and (iii) distantly related strains predicted to have weak (*Bl. longum*, *B. breve*) or minimal (*Bl. nov.* and *B. pseudocatenulatum*) HMO utilization capacity.

Strains with the H1 cluster reached the highest optical densities when grown in MRS-AC supplemented with an HMO mixture isolated from pooled human milk (**Fig. 4b**). HPLC-based glycoprofiling of culture supernatants revealed that these strains consumed 72-86% of total HMOs by 24h, including multiple fucosylated and sialylated structures (**Fig. 4c**; **Extended Data Fig. 9**). However, *Bl. suis* Bg131.S11_17.F6 and *Bc. kashiwanohense* Bg42221_1E1 did not efficiently utilize 2’FL, the most abundant species in the mixture, in line with the absence of characterized 2’FL transporters. In contrast, strains carrying the FHMO cluster preferentially metabolized fucosylated HMOs, completely depleting 2′FL while exhibiting limited utilization of sialylated structures. Among H1- and FHMO-cluster negative strains, *Bl. longum* Bg115.S08_3A11 depleted LNT, LNnT, LNFP I, LNFP III, LNH, and FLNH likely via partial extracellular degradation by β-galactosidase and lacto-*N*-biosidase, whereas *B. breve* Bg155.S08_4F7 metabolized LNT and LNnT intracellularly (**Supplementary Fig. 5**). *Bl. nov.* LFYP82 failed to efficiently utilize any tested HMOs, consistent with lineage-specific loss of relevant genes. Principal component analysis of growth and HMO utilization data separated strains with H1 and FHMO clusters from each other and all others (**Fig. 4d**). These findings provide the experimental evidence that *Bc. kashiwanohense* and *Bl. suis* strains carrying the H1 cluster exhibit HMO utilization patterns comparable to *Bl. infantis*, and highlight how variation in HMO utilization gene clusters drives metabolic divergence among closely related *Bifidobacterium* strains.

### Transcriptional profiles of glycan utilization in *Bc. kashiwanohense*

We used RNA-seq to test regulon predictions and pathway assignments in *Bc. kashiwanohense* Bg42221_1E1, which harbors both *xgl* and H1 gene clusters. Transcriptomic comparison of cultures grown in MRS-AC-XGL vs. MRS-AC-Lac revealed strong induction (100-550 fold) of most *xgl* genes and *cbpA*, supporting their co-regulation by XglT and a shared role in XGL metabolism (**Fig. 3a,e**; **Supplementary Table 21a**). Genes involved in xylose, XOS/AXOS, xylan, and AX metabolism showed moderate upregulation (4-30 fold), likely in response to intracellular xylose release, which may serve as a transcriptional effector of predicted transcriptional repressors XylR (ROK family) and XosR (LacI family; **Extended Data Fig. 7b,c**; **Supplementary Table 19**; *Supplementary Note 2*)^62^. Next, we compared the transcriptomes of *Bc. kashiwanohense* Bg42221_1E1 grown in MRS-AC-LNT and MRS-AC-Lac. All H1 cluster genes except *nanA2* and *nanH2* were significantly upregulated in the presence of LNT (**Fig. 4e; Supplementary Table 21b**), consistent with the proposed regulation by NagR, a GlcNAc-responsive repressor implicated in the control of HMO utilization in bifidobacteria^58^. Overall, the observed transcriptomes suggest that the regulatory networks of this strain are adapted for foraging on mixtures of HMOs and plant oligo- and polysaccharides.

Our large-scale genomic and experimental analyses pinpoint ecological differences between glycan foraging strategies within *Bifidobacterium* that reflect species-level evolutionary adaptation to different habitats (e.g., infant gut vs. adult gut) and dietary carbohydrate composition. The results also underscore considerable strain-level variability, likely shaped by host lifestyle and local dietary exposures.

## DISCUSSION

The ability to metabolize dietary glycans is central to bifidobacterial fitness in the human gut, with 8–12% of their gene content dedicated to carbohydrate metabolism^63^. Understanding this process has fundamental and translational relevance, including for the development of pro- and prebiotics. While interspecies differences in glycan preferences within bifidobacteria are well documented^25,64,65^, within-species variability, particularly in populations from low- and middle-income countries, remains understudied. The growing availability of *Bifidobacterium* genomes, driven by culturomics^66–69^ and metagenomics^70–75^, provides an opportunity to address this gap. However, accurately predicting carbohydrate utilization phenotypes from genomic data remains challenging due to imprecise functional annotations (particularly of glycan transporters) generated by widely-used pipelines, as well as incomplete representation of metabolic pathways in public databases^34,76,77^. To address this gap, we reconstructed 68 carbohydrate utilization pathways encoded in 3,083 *Bifidobacterium* genomes and MAGs and validated 38 predicted phenotypes *in vitro*, with an overall accuracy exceeding 94%. Several false-negative predictions for specific mono- and oligosaccharides, including HMOs, suggest the existence of yet uncharacterized glycan transport mechanisms in bifidobacteria.

Our analysis revealed taxon-specific distribution of glycan utilization pathways, reflecting the adaptation of *Bifidobacterium* species and subspecies to distinct ecological niches, such as the infant and adult gut, and expanding on earlier observations^25,30,64,65^. For instance, we identified a distinct clade within the *B. longum* species (*Bl. nov*.) specialized in metabolizing plant and fungal α-glucans but incapable of utilizing host glycans such as LNB, GNB, and HMOs. As most *Bl. nov.* genomes originated from infants in Westernized populations, the unique glycan preferences of this clade might reflect an adaptation to differences in early-life diet and feeding practices. However, certain pathways, such as raffinose utilization, were widely conserved. A recent study demonstrated that raffinose metabolism supports *B. breve* colonization and persistence in *in vivo*^19^, suggesting that conservation of this pathway across bifidobacterial lineages may promote their persistence and transmission beyond infancy.

Another important observation was extensive strain-level variability driven by accessory genome differences ranging from single genes to gene cassettes encoding entire carbohydrate utilization pathways. Particularly notable was the heterogeneity of *Bl. suis* isolates of Bangladeshi origin. Similar strains were recently predicted to utilize HMOs and plant fibers based on the analysis of CAZyme repertoires and reclassified as *B. longum* subsp. *iuvenis*^7,43^. Our analysis revealed nuanced metabolic differences within this clade, suggesting the presence of two distinct “ecotypes”. The first, exemplified by strain Bg131.S11_17.F6, metabolizes a wide array of HMOs — including LNT, LNnT, long-chain fucosylated, and sialylated species — but not arabinose and plant polysaccharides, mimicking the glycan preferences of *Bl. infantis*. The second ecotype, represented by strain Bg41121_2E1, preferentially utilizes fucosylated HMO structures, including the abundant 2’FL, but can metabolize arabinose-containing oligo- and polysaccharides, phenotypically resembling previously characterized *Bl. longum* strains SC596 and MC10007^48,49^. These distinct and complementary strategies suggest ecological specialization within the *Bl. suis* group, potentially linked to diet and weaning status.

Further within-species variability is illustrated by Bangladeshi isolates carrying unique gene clusters. One notable example was the *xgl* cluster identified in *Bc. kashiwanohense* Bg42221_1E1 and *Bc. kashiwanohense_A* Bg42221_1D3 strains isolated from the same infant. This cluster enables xyloglucan degradation, a capability previously suggested only for *Bifidobacterium* species inhabiting captive marmosets^78^. Another unexpected discovery was the presence of an H1 cluster variant in *Bc. kashiwanohense* Bg42221_1E1, formerly considered to be exclusive for *Bl. infantis*^46^ and rare Bangladeshi *Bl. suis* group strains^7,23^. Although the precise substrate-specificities of transporters encoded by H1 cluster remain to be characterized, our data confirm that its presence enables the utilization of major HMOs found in human milk. *Bc. kashiwanohense* strains have been shown to metabolize fucosylated HMOs (2’FL and LNFP I) and plant hemicelluloses (xylan and arabinoxylan)^8^. Our findings expand the catabolic potential of *Bc. kashiwanohense* to xyloglucan and a broader range of HMOs (LNT, LNnT, and sialylated HMOs). These results indicate an evolutionary adaptation of Bangladeshi strains to thrive in the microbiota of weaning children, whose diets may combine breast milk with complementary foods rich in plant polysaccharides.

While we did not explicitly reconstruct cross-feeding interactions, the lower prevalence of complete catabolic pathways for polysaccharides (relative to those for their mono- and oligomeric building blocks) suggests that many *Bifidobacterium* strains may depend on syntrophic relationships, both within the genus and with keystone degraders such as *Bacteroides* and *Segatella* (formerly *Prevotella*)^79–81^. Recent evidence for widespread HMO-degrading capacity across gut microbes^82^ further supports the potential for cross-feeding involving bifidobacteria and other community members.

The genomic dataset analyzed in this study is skewed towards samples from high-income countries, highlighting the need for initiatives to expand collections of *Bifidobacterium* isolates and genomes from underrepresented populations in an ethical and culturally sensitive manner to capture global metabolic diversity^83^. Some observed within-species variability may reflect artifacts introduced by incomplete or contaminated MAGs and different assembly methods, underscoring the importance of expanding collections of high-quality MAGs and isolate genomes.

In summary, our study reveals how glycan foraging strategies vary across and within *Bifidobacterium* species, shaped by ecological factors, including host age, diet, and lifestyle. The comprehensive metabolic reconstruction and an automated pathway prediction pipeline provide a scalable framework for accurate functional annotation of bifidobacterial genomes and MAGs. This genomic compendium may inform the development of pro- and synbiotic formulations, including multi-strain consortia, whose members might efficiently colonize the gut across diverse lifestyle and dietary contexts.

## Acknowledgments

We thank Robert Olson (Hack Biology LLC) for helping with mcSEED maintenance, Kang Liu (SBP Medical Discovery Institute Genomics Core) for library preparation and sequencing, Swetha Nakshatri (Washington University) and Kennedy Spann (UC San Diego) for technical assistance, Glycom A/S and DSM for generously providing individual HMOs, Dr. Takane Katayama (Kyoto University) for generously sharing *Bifidobacterium scardovii* JCM 12489 and insightful discussions. We are grateful to Dr. Mark Manary (Washington University) and study personnel at the Malawi College of Medicine for the provision of fecal samples used to isolate the Malawian bifidobacterial strains described in this report. Bangladeshi isolates were provided through a long-standing collaborative research program between Washington University and members of the International Center for Diarrheal Disease Research, Bangladesh (icddr,b) led by Dr. Tahmeed Ahmed. An MTA was established to transfer these strains to SBP Medical Discovery Institute. This work was supported by grants from the National Institutes of Health (NIH) (DK30292 to A.L.O. and J.I.G.) and the Bill & Melinda Gates Foundation (INV-016367). L.B. is the UC San Diego Chair of Collaborative Human Milk Research, endowed by the Family Larsson-Rosenquist Foundation, Switzerland.

## Author contributions statement

A.A.A., D.A.R., M.J.B., J.I.G., and A.L.O. designed the study. A.A.A. did data curation and metabolic reconstruction. A.A.A., D.A.R., M.D.K., S.A.L., and A.L.O. developed the pathway prediction pipeline. J.L.G. isolated bacterial strains. M.C.H., J.E.K., and M.L.E. sequenced and assembled bacterial genomes. A.A.A. did growth and RNA-seq experiments. K.S., A.F., and L.B. isolated HMOs and did glycoprofiling analysis. A.A.A. wrote the initial draft with input from J.E.K. A.A.A., D.A.R., M.C.H., L.B., M.J.B., J.I.G., and A.L.O. edited the manuscript with invaluable assistance from co-authors.

## Competing interests statement

D.A.R. and A.L.O. are co-founders of Phenobiome Inc., a company pursuing the development of personalized nutritional solutions to balance the gut microbiome. L.B. is a coinventor on patent applications related to the use of HMOs in preventing necrotizing enterocolitis and other inflammatory diseases. The remaining authors of this paper declare no competing interests.

## METHODS

### Collection of *Bifidobacterium* genomes

Reference genomes for the manual *in silico* metabolic reconstruction were retrieved from Bacterial and Viral Bioinformatics Resource Center (BV-BRC)^84^ and Integrated Microbial Genomes (IMG)^85^ databases as of October 2020. We selected 335 genomes based on the following criteria: (i) human or probiotic product-derived *Bifidobacterium spp*. isolates, (ii) number of contigs ≤ 200, (iii) CheckM^86^ completeness ≥ 97 % and contamination ≤ 3%. Additionally, we sequenced 31 genomes of cultured isolates from fecal samples obtained from Bangladeshi children^37,38^. The resulting reference set of 366 genomes was clustered using dRep (v3.4.2)^87^ at a 99.95% ANI threshold (“dRep dereplicate -pa 0.9 -sa 0.9995 -nc 0.35 -- S_algorithm ANImf”), yielding 263 non-redundant genomes (**Supplementary Table 2**).

Additional human gut-derived *Bifidobacterium* genomes were collected from prior studies. These included isolate genomes from HBC^66^, BIO-ML^67^, and CGR^68^ collections, and MAGs from UHGG^70^, KIJ^71^, ELGG^72^, SPMP^73^, and IMGG^74^ collections and other datasets^75^. We used the same selection criteria for both isolate genomes and MAGs: (i) *Bifidobacterium* spp. based on Genome Taxonomy Database (GTDB) taxonomy^88^ excluding *Bifidobacterium leopoldii* and *Bifidobacterium vaginale*, (ii) number of contigs ≤ 200, (iii) CheckM completeness ≥ 97% and contamination ≤ 3%. The resulting 4,944 genomes were clustered via dRep using the following command “dRep dereplicate -pa 0.9 -sa 0.999 -nc 0.35 --S_algorithm fastANI”, yielding 2811 non-redundant genomes (**Supplementary Table 3**). We also sequenced nine genomes of strains isolated from Bangladeshi and Malawian infants previously^37–39^, resulting in 2,820 additional genomes.

### Isolation and sequencing of *Bifidobacterium* strains from Bangladeshi and Malawian donors

Fecal samples used for culturing Bangladeshi bifidobacterial strains were collected during studies conducted by the International Centre for Diarrhoeal Disease Research (icddr,b). These were (i) the MAL-ED birth cohort study of children aged 0-24 months (Interactions of Enteric Infections and Malnutrition and the Consequences for Child Health and Development; ClinicalTrials.gov identifier NCT02441426) and (ii) a cohort of healthy 12-24 month-old Bangladeshi children enrolled in parallel with children with acute malnutrition in a study of microbiota-directed complementary food (MDCF) prototypes (ClinicalTrials.gov identifier NCT03084731)^37,38^. Both studies were approved by the Ethical Review Committee of the icddr,b. Bifidobacterial strains were also isolated from fecal samples collected in a previously reported study of Malawian twins discordant for acute malnutrition^39^. The protocol was approved by the College of Medicine Research Ethics Committee of the University of Malawi and by the Human Research Protection Office of Washington University in St. Louis. Written informed consent, including provisions for future use of materials, was provided by the parents or guardians of participating children before enrolment. The details of genome sequencing and assembly are provided in *Supplementary Methods*.

### Taxonomy inference

Initial taxonomic assignments for the 263 reference *Bifidobacterium* genomes were retrieved from NCBI and BV-BRC for public genomes or inferred from 16S rRNA gene sequencing for Bangladeshi isolates. To refine these assignments, we built a pangenome via Panaroo (v1.3.2)^89^. Concatenated nucleotide sequences of 487 identified core genes (**Supplementary Table 12**) were aligned using MAFFT (v7.515)^90^. A maximum-likelihood phylogenetic tree was built in IQ-TREE (v2.2.0.3)^91^ and visualized via iTOL (v5)^92^. Pairwise ANI values (ANIb) were calculated using pyani (v0.2.12)^93^. The exact commands and parameters are provided in Supplementary Code File 1. The resulting tree topology was manually inspected to verify and correct taxonomic assignments of genomes based on their co-clustering with branches corresponding to the type or well-characterized strains of *Bifidobacterium* species and subspecies. ANI matrices were used to delineate the within-species structure for *Bifidobacterium longum* and *Bifidobacterium catenulatum* species. GTDB-based taxonomies of 2,820 additional genomes were retrieved from original publications and refined via ANI comparisons with 263 reference genomes. Species assignments were made if ANI to the closest reference genome was > 95%, and subspecies assignments if > 97%.

### Gene prediction and functional annotation

Protein-coding sequences were predicted and annotated with Prokka (v1.14.6)^94^ using default settings. The 263 reference genomes were additionally annotated via RASTtk (v1.073)^95^ and EggNOG-mapper (v2.1.12)^96^ with default settings (**Supplementary Table 13**). The representation of CAZymes in 263 genomes was analyzed using dbCAN (v4.0.0)^97^. HMMER, dbCAN-sub, and DIAMOND-based searches against the CAZy database were used for CAZyme identification. Only GHs, CEs, and polysaccharide lyases predicted by two or more methods were retained.

### Subsystems-based annotation and *in silico* metabolic reconstruction of glycan utilization pathways

We used a subsystems-based approach implemented in the SEED platform^35,95^ to reconstruct carbohydrate utilization pathways encoded in reference 263 *Bifidobacterium* strains. RASTtk-annotated genomes were uploaded to mcSEED (microbial community SEED), a clone of the SEED annotation environment. We created 25 subsystems that captured the representation of genes that implement *functional roles* (glycan transporters, CAZymes, downstream catabolic enzymes, and transcriptional regulators) involved in carbohydrate utilization (**Supplementary Table 4**). The list of functional roles was compiled via extensive literature search and using the information from Transporter Classification (TCDB)^98^, Carbohydrate Active Enzyme (CAZy)^99^, Kyoto Encyclopedia of Genes (KEGG)^100^, and RegPrecise^101^ databases. This knowledge was used to manually curate the automated functional annotations of protein-coding genes in a subset of reference *Bifidobacterium* genomes (e.g., type strains) where the respective functional roles had been experimentally characterized. Additional functional gene annotation to fill gaps in metabolic pathways was based on three genome context techniques: (i) clustering of genes on the chromosome (operons), (ii) co-regulation of genes by a common transcription factor, and (iii) co-occurrence of genes across related genomes. The approach used to reconstruct the regulons of transcription factors potentially regulating carbohydrate metabolism is outlined in *Supplementary methods*.

The propagation of curated annotations (corresponding to 589 functional roles) across all 263 reference genomes was performed using homology-based methods implemented in mcSEED. Orthologs were detected automatically using predefined protein family classifications such as PGFams (cross-genus protein families) and PLFams (genus-specific protein families)^84^. These assignments were manually refined by examining the gene neighborhood for each functional role. Genes with conserved gene neighborhoods were classified as orthologous, while paralogs were assigned distinct functional annotations. Overall, 39,589 out of 541,418 protein-coding gene annotations in 263 reference *Bifidobacterium* genomes were curated.

Next, we reconstructed 72 catabolic pathways spanning 25 subsystems (**Supplementary Table 5**). Many pathways included alternative biochemical modules (routes) driven by different sets of catabolic enzymes and diverse glycan transporters. For each pathway, we defined genomic signatures — sets of genes encoding functional roles that together represent the minimal gene complement required to form a complete pathway for the utilization or degradation of a specific carbohydrate (**Supplementary Table 6**). *Utilization* was defined as a process when a glycan molecule (mono- or oligosaccharide) is transported into the cell and then catabolized using a combination of CAZymes and downstream catabolic enzymes. *Degradation* referred to the partial hydrolysis of a glycan (e.g., a polysaccharide) by extracellular CAZymes, with the resulting mono- and oligosaccharides subsequently transported into the cell and metabolized.

We used the presence or absence of genes matching each genomic signature to assign a detailed pathway variant to each genome: (i) transporter + catabolic pathway (“U”), (ii) catabolic pathway without transporter (“P”), (iii) no catabolic pathway (“N”) (**Supplementary Tables 6,8**). For the purpose of automated phenotype profiling, these assignments were simplified into binary phenotypes: a complete pathway (“U”) indicated predicted utilization or degradation and was assigned phenotype “1”, while incomplete (“P”) or missing (“N”) pathway variants indicated no utilization and were assigned phenotype “0”. The resulting set of 72 predicted carbohydrate utilization phenotypes across 263 strains comprised the reference Binary Phenotype Matrix (BPM) (**Supplementary Table 7**). Four pathways (GalNAc, ManAc, Man, and GalA), for which all strains in the reference set were assigned variants “P” or “N” and phenotype “0”, respectively, were retained in the BPM but excluded from the pathway prediction pipeline.

### Pathway prediction pipeline (glycobif)

The workflow used to predict the carbohydrate utilization pathways encoded in additional 2,820 *Bifidobacterium* genomes is schematically illustrated in **Fig. 1a**. We first constructed a reference database containing (i) 39,589 functionally annotated protein sequences from 25 curated mcSEED subsystems across 263 reference genomes, and (ii) an additional set of 52,990 outgroup proteins (not captured in the subsystems) clustered at 95% amino acid identity and 95% coverage using MMSeqs2 (v14.7e284)^102^. Proteins encoded in the 2,820 query genomes were annotated by mapping their sequences to the reference database using DIAMOND (v2.1.4)^103^. To handle multidomain proteins, we first selected the top 50 hits for each query based on bitscore and clustered the alignment coordinates using DBSCAN (scikit-learn v1.2.1)^104^. Cluster centers were treated as potential domain boundaries, which were used to split query proteins into discrete domains, with database hits attributed to each individual domain. For each resulting domain with ≥ 35 amino acids, we applied the Gaussian kernel density modeling (*KernelDensity* function from *sklearn.neighbors*) to the distribution of sequence identity values, and used the highest local minimum (*argrelextrema* function from *sklearn.signal*) to filter out low-confidence hits. Annotations were assigned by majority vote from high-scoring, domain-specific reference hits. High-identity hits to outgroup proteins from the reference database were used as criteria to vote against applying annotation to a given query. This pipeline yielded 419,055 annotated protein sequences.

These annotations, together with the BPM for 263 reference genomes, were integrated into a machine learning-based pipeline to predict the presence (“1”) or absence (“0”) of carbohydrate utilization pathways in additional 2,820 genomes. We evaluated over 30 machine learning methods implemented in the *Caret* package (v6.0.86)^105^ using a leave-one-out cross-validation approach: for each reference genome, a model was trained on the remaining 262 genomes and then used to predict the binary variant for the held-out genome. This process was repeated for every pathway. Random forest was identified as the best-performing model based on prediction accuracy. Pathway-specific random forest models were then trained using the full reference set, excluding four pathways (GalNAc, ManAc, Man, and GalA) for which all genomes had predicted binary phenotype “0”. The list of functional roles used as model predictors for each remaining pathway was manually curated to match the genomic signatures delineating pathway variants (**Supplementary Table 6**). Model parameters were optimized using grid search, and mock genomes with custom genomic signatures were added to the training set to ensure that rare combinations from the reference collection were adequately learned and to improve model performance on incomplete MAGs. The resulting models were used to predict the presence or absence (1/0) of 68 carbohydrate utilization pathways in 2,820 genomes (**Supplementary Table 9**).

The representation of 29 additional metabolic pathways (e.g., the biosynthesis of B vitamins and amino acids) in both 263 reference and 2,820 additional genomes was inferred via a similar annotation pipeline as recently published^106^ (**Supplementary Table 11**).

### Visualization and statistical analysis

Non-metric multidimensional scaling (NMDS) was used for the ordination of the Hamming distance matrix calculated from the merged BPM of 3,083 genomes. PERMANOVA and a test for the homogeneity of multivariate dispersions were performed using the *adonis2* and *betadisper* functions in the *vegan* package (v2.6-8)^107^. Differences in predicted phenotypic richness were assessed via a generalized linear model (GLM) assuming Poisson distribution; post hoc comparisons with Bonferroni correction were conducted using the *emmeans* package (v1.10.6)^108^. Fisher’s exact test was used for the enrichment analysis of pathway representation. For each of the 68 carbohydrate utilization pathways, 2×2 contingency tables were constructed. Rows denoted the absence/presence of predicted pathways (“0” or “1”), whereas columns represented metadata categories: age group (age < 3 years vs. age ≥ 3) or host lifestyle (Westernized vs. non-Westernized)^61^. P-values were corrected for multiple testing using the Benjamini-Hochberg procedure, with Padj ≤ 0.01 considered significant (**Supplementary Table 14**). Odds ratios were calculated using the *fisher.test* function. Principal component analysis (PCA) was used for the ordination of growth and HMO consumption data. Data visualization was performed using *ggplot2* (v3.5.1)^109^, *ComplexHeatmap* (v2.18.0)^110^, and clinker (v0.0.27)^111^. Code for all analyses is provided in **Supplementary Code File 1**.

### *In vitro* growth of *Bifidobacterium* strains on selected carbohydrates

All strains except *B. scardovii* JCM 12489 were routinely grown in dextrose-free Lactobacilli de Man, Rogosa & Sharpe broth (Alpha Biosciences) supplemented with 0.34% (wt/vol) sodium ascorbate, 0.029% (wt/vol) L-cysteine-HCl monohydrate, and 10 mg/mL lactose (MRS-AC-Lac) in a chamber maintained with a gas mix of 10% H_2_, 10% CO_2_, and 80% N_2_ (Coy Laboratory Products). The MRS-C-Lac medium without sodium ascorbate was used for *B. scardovii* JCM 12489. The growth of 30 *Bifidobacterium* strains on 47 substrates (**Supplementary Table 15**) was measured using a custom carbohydrate array constructed in flat-bottom half-area 96-well plates (Costar). Wells were filled with 55 µl of a sterilized carbohydrate stock at 2× concentration (10 to 20 mg/mL) and transferred to the chamber 48 h before the experiment. Each substrate was tested in triplicate (or duplicate in a few cases). Sterile water and lactose served as negative and positive controls.

Cultures for assay inoculations were grown overnight at 37 °C in MRS-AC-Lac, diluted 1:100 into fresh medium, and incubated for 8 h. Cultures were then adjusted to OD = 0.6, and 100 μL aliquots were harvested at 5,000 × g for 5 min and resuspended in 6 mL of 2× MRS-AC (2× MRS-C for *B. scardovii*) without added carbohydrate. Each well in the carbohydrate array was loaded with 55 µL of inoculated 2× medium to make individual 110 µL cultures. Growth was monitored by measuring OD_600_ every 15 or 30 minutes for 40 h under constant linear shaking using a Synergy H1 Plate Reader (BioTek Instruments). Raw OD_600_ data were exported using Gen5 software (v2.05.5) and analyzed in R (v4.3.2). For all strains except *B. pseudocatenulatum* M26A_2F1 growth on each glycan was classified as follows: (i) no growth (“−”) if mean OD_600_ never exceeded 10% of the strain’s highest OD_600_ within a panel of carbon sources, (ii) weak growth (“w”) if mean OD_600_ did not exceed 25% of maximum OD_600_, (iii) growth (“+”) if mean OD_600_ was above 25% of maximum OD_600_ (**Supplementary Fig. 4**; **Supplementary Table 16**). For *B. pseudocatenulatum* M26A_2F1, thresholds were increased to 22% and 55%, respectively, due to some outgrowth in basal MRS-AC lacking an added substrate.

### Comparison of predicted phenotypes with growth data

Predicted carbohydrate utilization phenotypes were compared with growth data from this work or prior literature^8,28,36^. A predicted binary phenotype “1” was classified as a true positive (TP) if it matched with growth phenotypes “+/w” and as a false positive (FP) if it matched with “−”. A predicted binary phenotype “0” was considered a true negative (TN) if it matched with a growth phenotype “−” and a false negative (FN) if it matched with “+/w”. Most predicted utilization phenotypes were tested using individual substrates. Exceptions included phenotypes IMO, for which the growth was tested on panose and isomaltotriose, BglOS (cellobiose, gentiobiose, laminaritriose, and sophorose), and SHMO (3’-sialyllactose, 6’-sialyllactose). For these cases, growth (“+/w”) on at least one substrate was considered as a TP for a predicted binary phenotype “1”, and the absence of growth on all tested substrates was considered a TN for “0”. Standard binary classification metrics were calculated from TP, TN, FP, and FN counts (**Supplementary Table 17**).

### Glycoprofiling of HMO consumption

Overnight cultures of *Bifidobacterium* strains grown in MRS-AC-Lac were inoculated (0.5–1% [vol/vol]) into fresh medium and cultured to OD_600_=0.6. Cells were harvested at 5,000 × g for 5 min, washed with sugar-free MRS-AC, and used to inoculate 200 µL of MRS-AC supplemented with an HMO mixture (10 mg/mL, isolated from pooled human milk from ≥ 25 different donors) in 96-well plates (Costar) at OD_600_=0.005. Triplicate 100 µL samples of culture supernatant were collected at 8 or 24 h time points in separate experiments, filtered through Spin-X Centrifuge Tube Filter (0.22 µm, Costar), snap-frozen in liquid nitrogen, and stored at −80 °C. HMO concentrations in supernatants were analyzed by high-performance liquid chromatography (HPLC) with fluorescence detection^112^. Media samples (10 µL) were spiked with maltose as an internal standard, lyophilized, and labeled with the fluorophore 2-aminobenzamide (2AB). 2AB-labeled HMOs were separated on a TSKgel Amide-80 column (2.0 mm ID × 15 cm, 3 µm; Tosoh Bioscience) and detected at 360 nm excitation / 425 nm emission on Dionex Ultimate 3000 (Thermo Fisher Scientific). Nineteen HMO structures were annotated based on standard retention times and quantified relative to the internal standard: 2’-fucosyllactose (2’FL), 3-fucosyllactose (3FL), lactodifucotetraose (LDFT), 3’-sialyllactose (3’SL), 6’-sialyllactose (6’SL), lacto-*N*-tetraose (LNT), lacto-*N*-neotetraose (LNnT), lacto-*N*-fucopentaose (LNFP) I/II/III, sialyllacto-*N*-tetraose (LST) b and c, difucosyllacto-*N*-tetraose (DFLNT), lacto-*N*-hexaose (LNH), disialyllacto-*N*-tetraose (DSLNT), fucosyllacto-*N*-hexaose (FLNH), difucosyllacto-*N*-hexaose (DFLNH), fucodisialyllacto-*N*-hexaose (FDSLNH), disialyllacto-*N*-hexaose (DSLNH). Total HMO concentration was calculated as the sum of individual HMO concentrations (**Supplementary Table 18**). HMO utilization at 8 or 24 h was calculated relative to HMO concentrations in the medium controls.

### Transcriptional profiling (RNA-seq)

An overnight culture of *Bc. kashiwanohense* Bg42221_1E1 grown in MRS-AC-Glc was harvested at 5,000 × g for 5 min, washed in sugar-free MRS-AC, and used to inoculate MRS-AC supplemented with either 10 mg/mL of glucose, lactose, lacto-*N*-tetraose, or 5 mg/mL of tamarind xyloglucan at OD_600_=0.01. Samples (2 mL, biological triplicates) were collected at the early exponential phase (OD_600_=0.35) and pelleted in a prechilled centrifuge at 4,800 × g for 5 min. Cell pellets were snap-frozen in liquid nitrogen and stored at −80 °C until further use. The detailed RNA extraction protocol is described in *Supplementary methods*. Ribosomal RNA was depleted with the NEBNext rRNA depletion kit for bacteria (New England Biolabs) and a set of 20 pooled sequence-specific probes for *Bc. kashiwanohense* Bg42221_1E1 designed using the NEBNext Custom RNA Depletion Design Tool v1.0 (**Supplementary Table 20**). Barcoded libraries were made with the NEBNext Ultra II directional RNA library prep kit for Illumina (New England Biolabs). Libraries were pooled and sequenced (single-end 75-bp reads) on Illumina NextSeq 500 using the High Output V2 kit (Illumina). The details of sequencing data analysis are provided in *Supplementary methods*.

## Data availability

Genomes of *Bifidobacterium* isolates sequenced in this study have been deposited in GenBank (https://www.ncbi.nlm.nih.gov/bioproject/PRJNA1126848). Nucleotide FASTA and annotated protein FASTA files of 263 reference genomes are available on Figshare (https://doi.org/10.6084/m9.figshare.26053936). Additional *Bifidobacterium* genomes and MAGs were retrieved from the following publicly available databases and datasets: BV-BRC (https://www.bv-brc.org), IMG (https://img.jgi.doe.gov), BIO-ML (https://www.ncbi.nlm.nih.gov/bioproject/PRJNA544527), CGR (https://www.ncbi.nlm.nih.gov/bioproject/PRJNA482748), CGR2 (https://www.ncbi.nlm.nih.gov/bioproject/PRJNA903559), UHGG (http://ftp.ebi.ac.uk/pub/databases/metagenomics/mgnify_genomes), KIJ (https://www.decodebiome.org/HRGM1), ELGG (https://zenodo.org/records/6969520), SPMP (https://figshare.com/collections/SPMP/5993596), IMGG (https://www.ncbi.nlm.nih.gov/bioproject/PRJNA763692). The RNA-seq data set has been deposited in Gene Expression Omnibus (https://www.ncbi.nlm.nih.gov/geo/query/acc.cgi?acc=GSE239955). Source data are provided on GitHub (https://github.com/Arzamasov/glycobif).

## Code availability

Code detailing the data analysis steps is available on GitHub (https://github.com/Arzamasov/compendium_manuscript) and **Supplementary Code File 1.** The pipeline for analyzing the representation of carbohydrate utilization pathways encoded in bifidobacterial genomes (glycobif) is available on GitHub (https://github.com/Arzamasov/glycobif).

## Supplementary information

### Extended data figures

**Extended Data Fig. 1.**
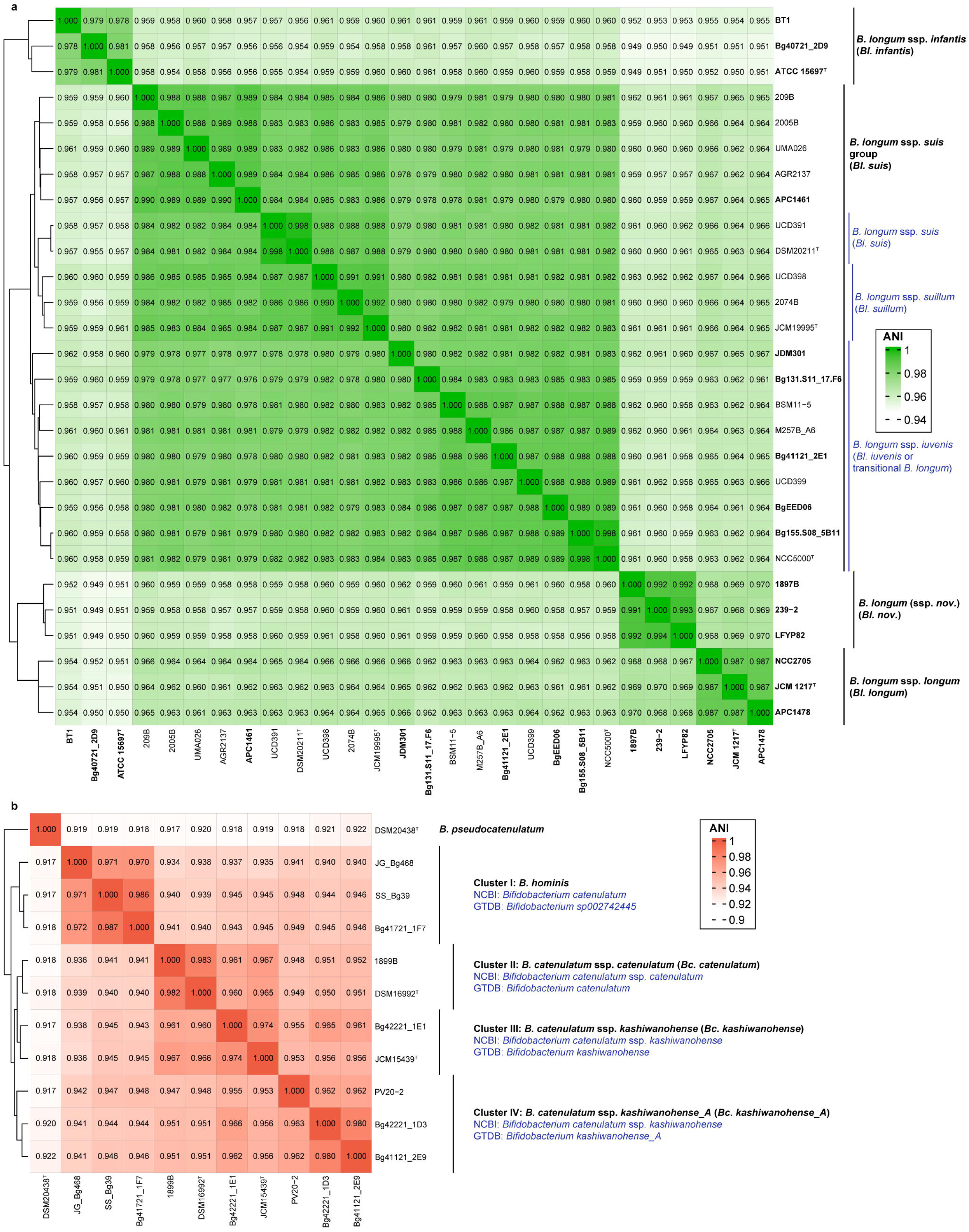
Pairwise average nucleotide identity (ANI) analysis of reference *B. longum* and *B. catenulatum* genomes. **a**, Pairwise ANI values for 28 *B. longum* genomes. For comparative purposes, this set contained 15 genomes from the reference collection (names are in bold) and 13 additional genomes, including the type strains of *Bl. suis* and *Bl. suillum* of non-human origin. Black lines indicate taxonomic groupings used in this study; blue lines denote the additional delineation proposed by Modesto et al.^43^. b, Pairwise ANI values for 11 reference *B. catenulatum*, *B. hominis*^69^, and *B. pseudocatenulatum* genomes. Black lines indicate taxonomic groupings used in this study; NCBI and GTDB taxonomy assignments are shown in blue.

**Extended Data Fig. 2.**
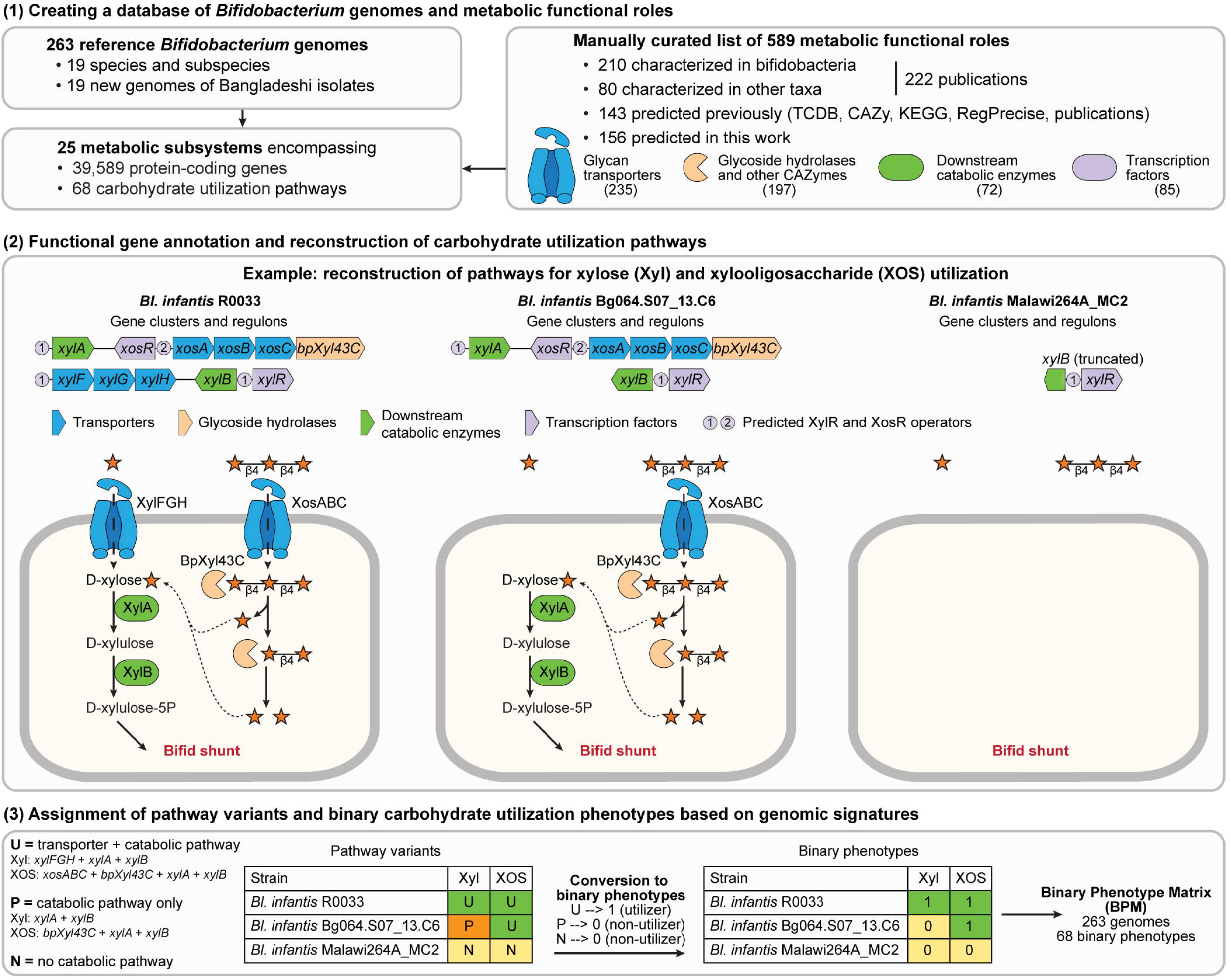
Reconstruction of carbohydrate utilization pathways and associated phenotypes. (1) A reference set of 263 Bifidobacterium genomes (**Supplementary Table 2**) and a list of 589 curated functional roles (**Supplementary Table 4**) were used to populate 25 metabolic subsystems capturing catabolic pathways for 68 glycans. (2) Functional gene annotation was performed using homology-based methods and three genome context techniques: (i) clustering of genes on the chromosome (operons), (ii) co-regulation of genes by a common transcription factor, and (iii) co-occurrence of genes across related genomes. Carbohydrate utilization pathways were reconstructed based on the distribution of functional roles. (3) Each genome was assigned a detailed pathway variant based on the presence of signature genes (**Supplementary Table 6**). These variants were then converted to predicted binary phenotypes (1 = utilizer, 0 = non-utilizer) to create a Binary Phenotype Matrix (BPM).

**Extended Data Fig. 3.**
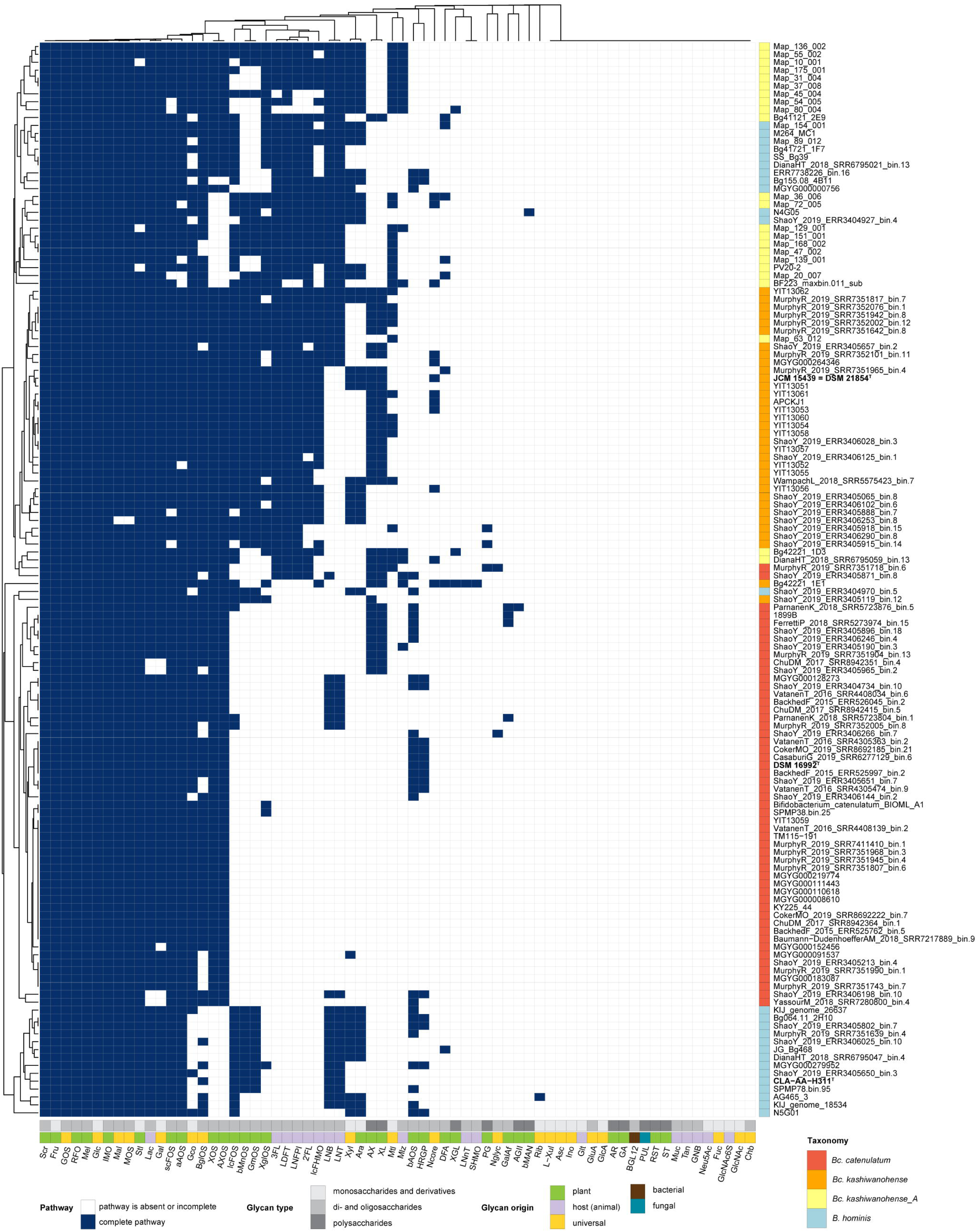
Representation of predicted carbohydrate utilization pathways in B. catenulatum and B. hominis genomes. The heatmap shows the hierarchical clustering of the Binary Phenotype Matrix (BPM) for 68 carbohydrate utilization pathways (columns) predicted in 110 *B. catenulatum* and 26 *B. hominis* genomes (rows). Pathway/phenotype classifications are shown in the bottom annotation tracks; taxonomic assignments are indicated on the right. Names of type strains are in bold. Full names pathway/phenotype names are provided in **Supplementary Table 5**.

**Extended Data Fig. 4.**
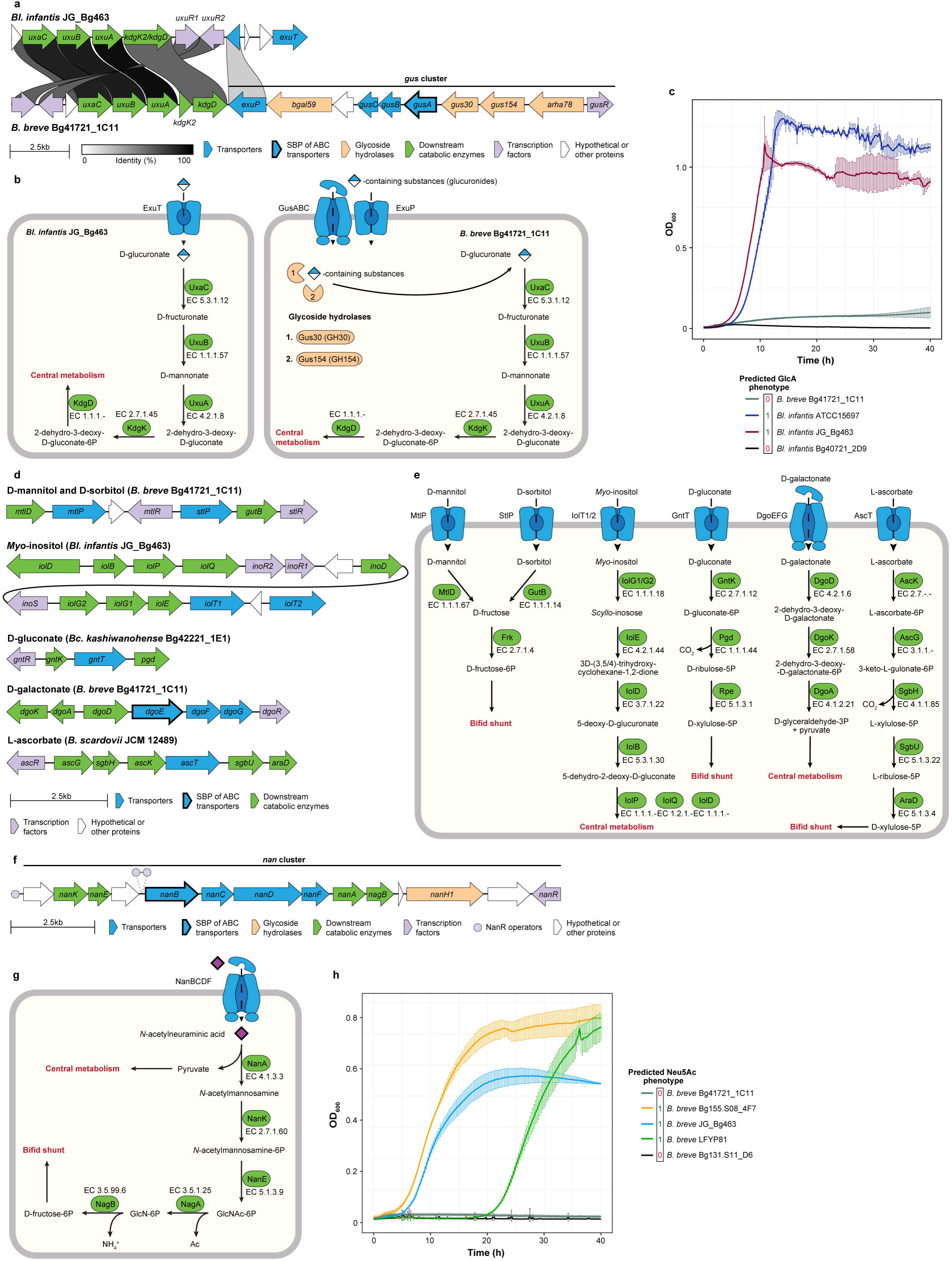
Genomics-based reconstruction of pathways involved in the utilization of monosaccharide derivatives. **a**, Gene clusters potentially driving D-glucuronate (GlcA) utilization by *Bl. infantis* JG_Bg 463 and glucuronide (GluA) utilization by *B. breve* Bg41721_1C11. Orthologous genes are linked, and the link color represents the sequence identity between corresponding protein products. **b**, Reconstructed D-glucuronate and glucuronide utilization pathways. **c**, Growth curves of selected *Bifidobacterium* strains in MRS-AC supplemented with 1% D-glucuronic acid. Data represent the mean ± s.d. of three biological replicates. **d**, Gene clusters potentially driving the utilization of (i) sugar alcohols D-mannitol (Mtl), D-sorbitol (Stl), and *myo*-inositol (Ino), (ii) sugar acids D-gluconate (Gco) and D-galactonate (Glt), (iii) L-ascorbate (Asc). **e**, Reconstructed utilization pathways for monosaccharide derivatives. **f**, Gene cluster (*nan*) driving sialic acid (Neu5Ac) utilization by *B. breve* strains. **g**, Neu5Ac utilization pathway in *B. breve*. **h**, Growth curves of selected *B. breve* strains in MRS-AC supplemented with 1% *N*-acetylneuraminic acid. Data represent the mean ± s.d. of three biological replicates.

**Extended Data Fig. 5.**
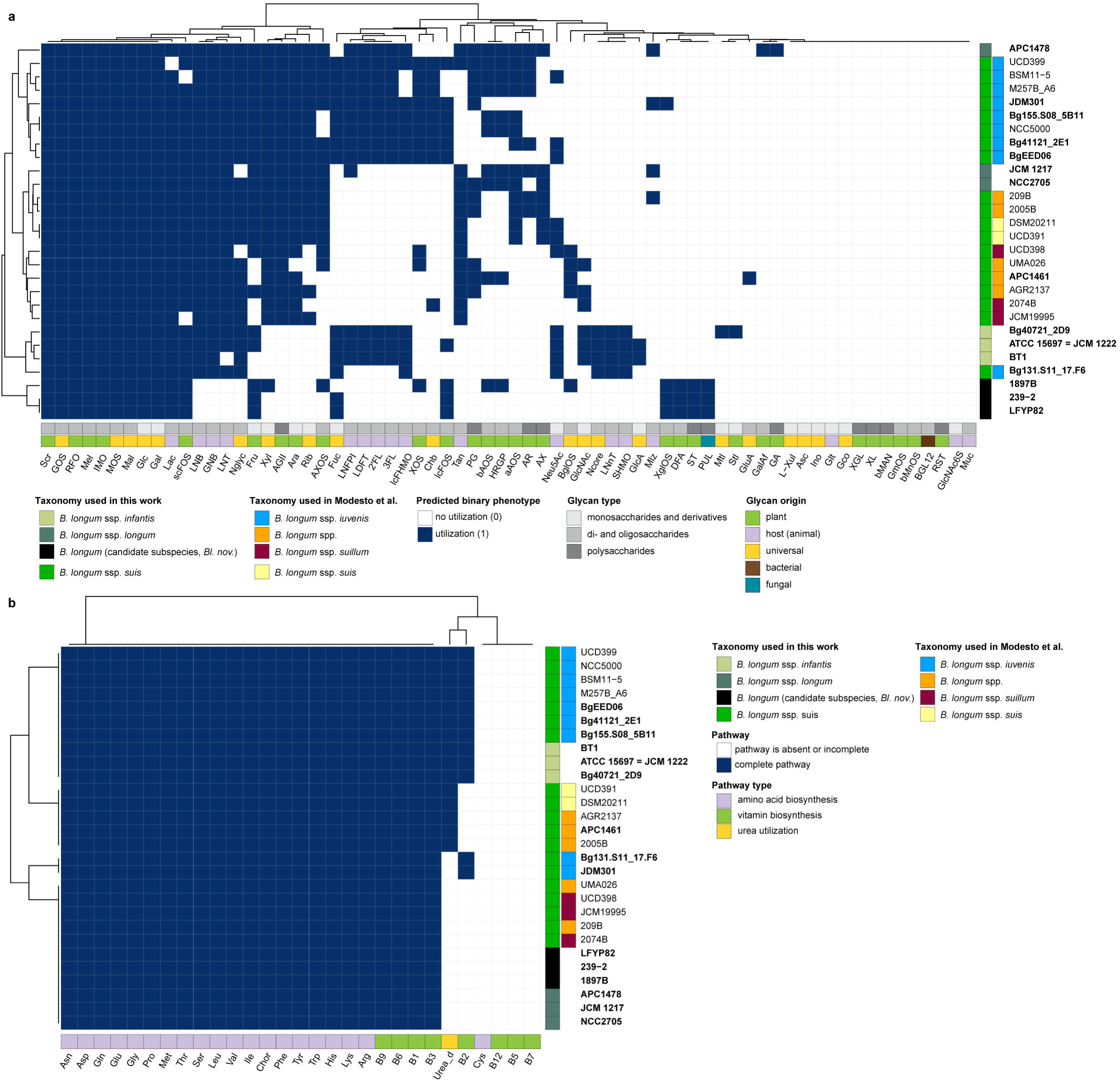
Representation of metabolic pathways in 28 selected B. longum isolate genomes. For comparative purposes, this dataset includes 15 genomes from the reference collection (names in bold) and 13 additional genomes, primarily of non-human origin, including type strains of *Bl. suis* and *Bl. suillum*. Annotation rows at the bottom represent pathway/phenotype classifications. Annotation columns on the right denote taxonomic groupings: the left column reflects assignments used in this study; the right column showing subspecies delineation proposed by Modesto et al.^43^. **a**, Representation of 68 carbohydrate utilization pathways and associated metabolic phenotypes. Full names are provided in **Supplementary Table 5**. **b**, Representation of 29 additional pathways, including the biosynthesis of amino acids and B vitamins, and urea utilization.

**Extended Data Fig. 6.**
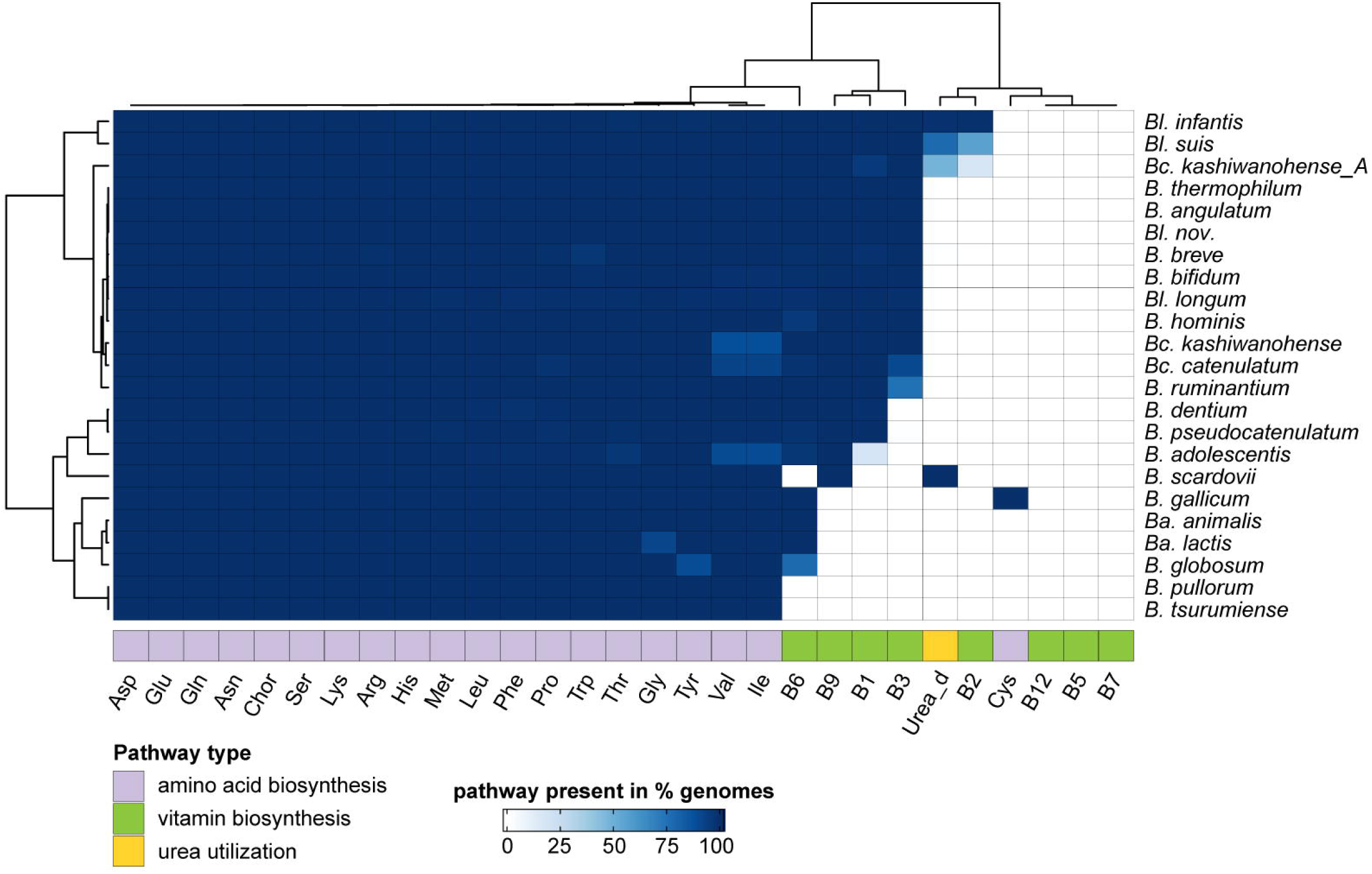
Representation of core biosynthetic pathways across 3,083 *Bifidobacterium genomes.* The heatmap depicts the prevalence of metabolic pathways corresponding to: (i) B vitamin and amino acid biosynthesis, (ii) urea utilization (Urea_d) within each taxon. Color intensity indicates the proportion of genomes that encode the respective metabolic pathway.

**Extended Data Fig. 7.**
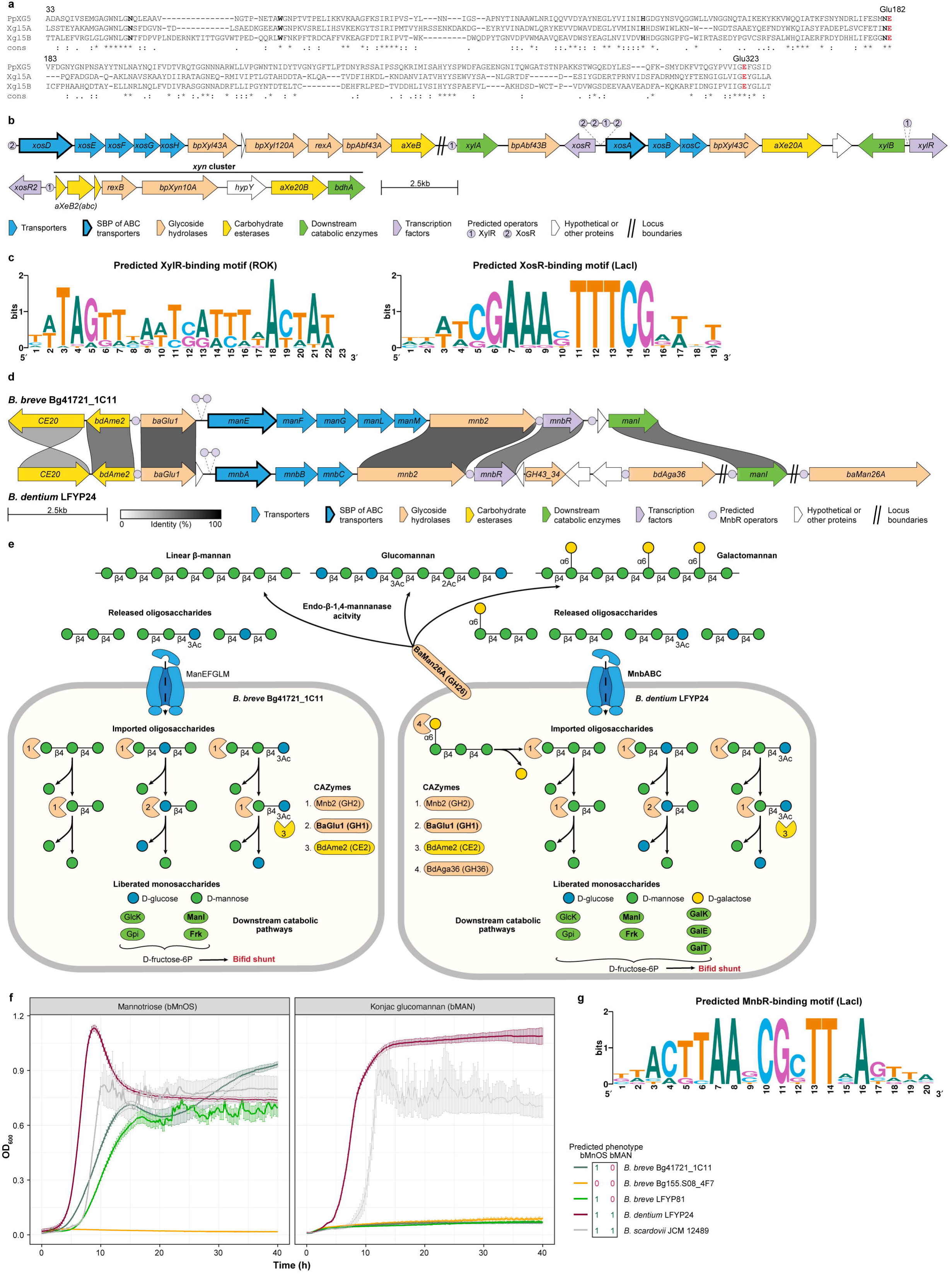
Genomics-based reconstruction of polysaccharide degradation and oligosaccharide utilization pathways. **a**, Alignment of the amino acid sequences of xyloglucan endo-β-1,4-glucanases (xyloglucanases): PpXG5 from *Paenibacillus pabuli* XG5 (characterized), Xgl5A and Xgl5B from *Bc*. *kashiwanohense* Bg42221_1E1 (putative). Conserved catalytic amino acid residues are highlighted in bold red, and conserved glycan-binding residues are in bold. **b**, Gene clusters potentially driving the degradation of (arabino)xylan and the utilization of released oligosaccharides in *Bc. kashiwanohense* Bg42221_1E1. **c**, Predicted DNA-binding motifs of transcription factors potentially controlling the expression of the gene clusters in **b**. Motifs were built based on operator sequences in the RegPrecise database and identified in this work (**Supplementary Table 19**). **d**, Gene clusters potentially driving β-mannose oligosaccharide utilization (phenotype bMnOS) in *B. breve* Bg41721_1C11 and β-mannan degradation (phenotype bMAN) in *B. dentium* LFYP24. Orthologous genes are linked, and the link color represents the sequence identity between corresponding protein products. **e**, Reconstructed β-mannose oligosaccharide utilization and β-mannan degradation pathways. Names of previously characterized enzymes and transporters are in bold. **f**, Growth curves of selected *Bifidobacterium* strains in the medium supplemented with 0.5% mannotriose or 0.5% konjac glucomannan. Data represent the mean ± s.d. of three biological replicates. **g**, Predicted MnbR DNA-binding motif based on operator sequences identified in this work (**Supplementary Table 19**).

**Extended Data Fig. 8.**
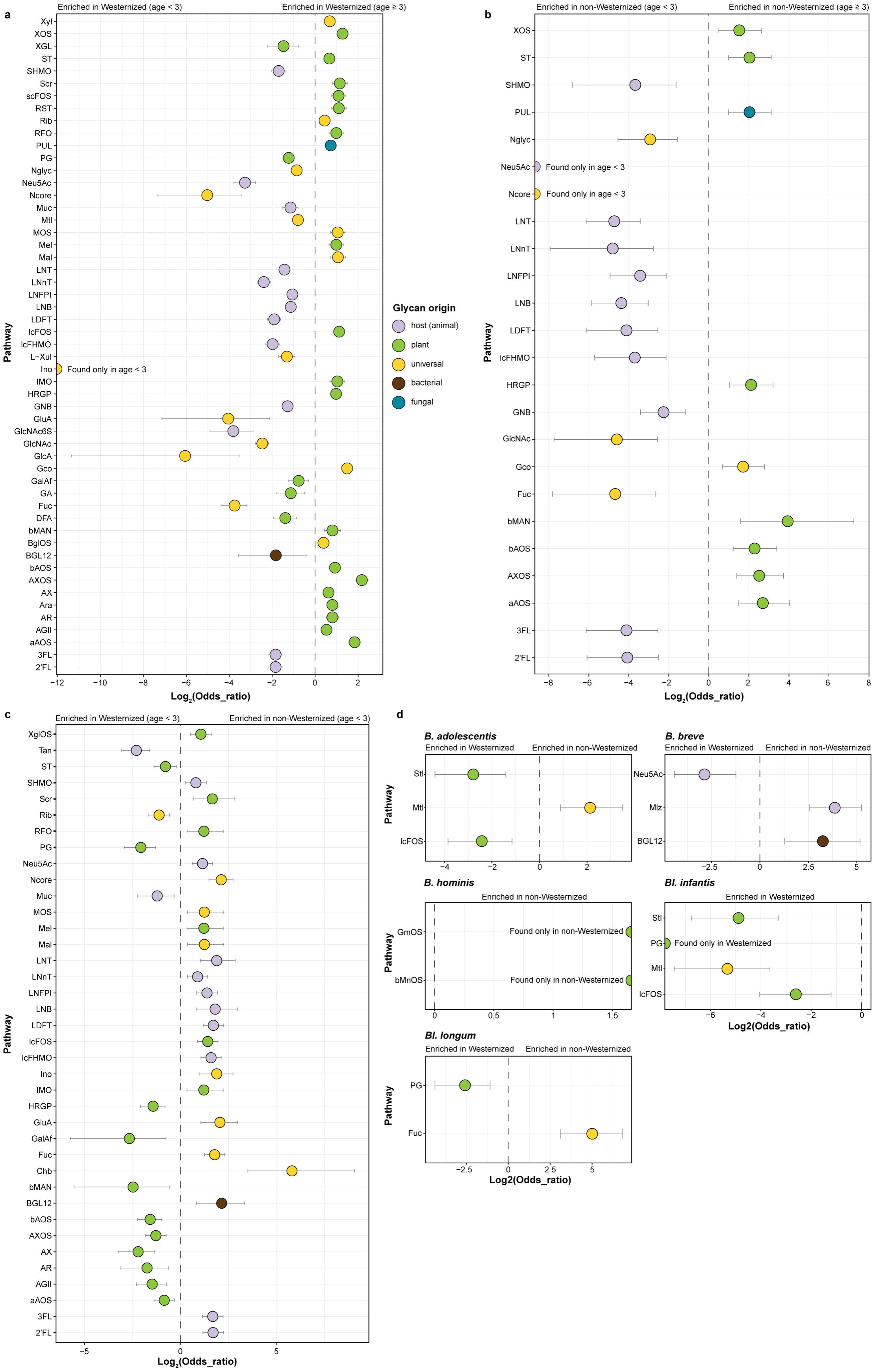
Pathway enrichment analysis across 3,083 *Bifidobacterium* genomes. Pathways significantly enriched in specific groups (Benjamini–Hochberg adjusted P ≤ 0.01; two-sided Fisher’s exact test) are shown. Points represent odds ratios, and horizontal lines indicate 95% confidence intervals. Pathways with infinite odds ratios are present exclusively in one group. Exact adjusted P-values are provided in **Supplementary Table 14**. **a**, Comparison between genomes from “Westernized (age < 3)” vs. “Westernized (age ≥ 3)” groups. **b**, Comparison between genomes from “non-Westernized (age < 3)” vs. “non-Westernized (age ≥ 3)” groups. **c**, Comparison between genomes from “Westernized (age < 3)” vs. “non-Westernized (age < 3)” groups. **d**, Comparison within individual taxa between genomes from “Westernized” vs. “non-Westernized” groups.

**Extended Data Fig. 9.**
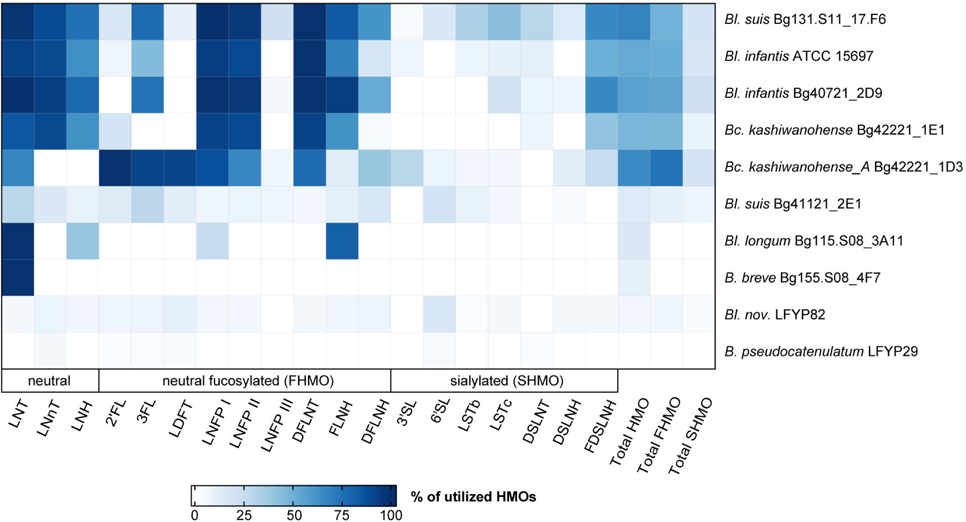
HPLC-based quantification of human milk oligosaccharide (HMO) depletion from culture supernatants after 8 h. Data represent the percentage of utilized HMOs (mean of three biological replicates) relative to the medium control. Total HMO = total HMO utilized; total FHMO = total fucosylated HMO utilized; total SHMO = total sialylated HMO utilized. Concentrations of individual HMOs (nmol/mL) are provided in **Supplementary Table 18a**.

### Supplementary figures

**Supplementary Fig. 1. Phylogenetic tree of 263 reference *Bifidobacterium* genomes.** The maximum-likelihood tree was constructed from a concatenated alignment of nucleotide sequences of 487 core genes. Node bootstrap support values are indicated by color.

**Supplementary Fig. 2. Concordance between phylogeny and predicted phenotypes.** The tanglegram compares the phylogenetic tree of 263 reference *Bifidobacterium* genomes with the Hamming-distance dendrogram derived from the Binary Phenotype Matrix (BPM). Lines connecting matching genomes are colored by taxonomic assignment.

**Supplementary Fig. 3. Variability in predicted carbohydrate utilization phenotypes across 263 reference *Bifidobacterium* strains.** The heatmap shows the hierarchical clustering (Hamming distance and average linkage) of the Binary Phenotype Matrix (BPM) for 68 carbohydrate utilization pathways (columns) reconstructed in 263 reference *Bifidobacterium* genomes (rows). Full names of abbreviations are provided in Supplementary Table 5. The annotation rows at the bottom represent pathway/phenotype classifications. The annotation column on the right denotes taxonomic groupings. Lines point to specific strains discussed in the text.

**Supplementary Fig. 4. Growth curves of 30 Bifidobacterium strains in MRS-AC medium supplemented with different substrates.** Data represent the mean and 95% confidence interval from three replicates (two for some substrates). Labels indicate the substrates used in each growth assay (see Supplementary Table 15 **for details**). “No substrate” refers to incubation in MRS-AC without added substrate (negative control). Red lines indicate the “no growth” threshold (10% of the maximum OD_600_ observed for a given strain across all substrates), while orange lines mark the “weak growth” threshold (25% of the maximum OD_600_).

**Supplementary Fig. 5**. **Reconstructed human milk oligosaccharide (HMO) utilization pathways in Bifidobacterium strains.** Pathways were inferred based on the integration of genomic predictions with growth phenotypes and HMO glycoprofiling data.

**Supplementary Table 1.** Distribution of 3,083 *Bifidobacterium* genomes by taxonomic group and geographic origin.

**Supplementary Table 2**. Metadata for 263 reference Bifidobacterium genomes used in curated metabolic reconstruction.

**Supplementary Table 3**. Metadata for 2,820 additional *Bifidobacterium* genomes used in the automated pathway prediction pipeline.

**Supplementary Table 4.** Curated list of functional roles comprising carbohydrate utilization pathways.

**Supplementary Table 5.** Glossary of reconstructed carbohydrate utilization pathways, associated substrates, and predicted metabolic phenotypes.

**Supplementary Table 6**. Genomic signatures used to assign pathway variants and predicted binary utilization phenotypes.

**Supplementary Table 7.** Binary Phenotype Matrix (BPM) depicting 68 carbohydrate utilization pathways reconstructed in 263 reference *Bifidobacterium* genomes.

**Supplementary Table 8.** Representation of functional roles comprising carbohydrate utilization pathways in 263 reference *Bifidobacterium* genomes.

**Supplementary Table 9.** Binary Phenotype Matrix (BPM) depicting 68 carbohydrate utilization pathways reconstructed in 2,820 additional *Bifidobacterium* genomes.

**Supplementary Table 10.** Representation of functional roles comprising carbohydrate utilization pathways in 2,820 additional *Bifidobacterium* genomes.

**Supplementary Table 11.** Binary Phenotype Matrix (BPM) depicting 30 additional metabolic pathways reconstructed in 3,083 *Bifidobacterium* genomes.

**Supplementary Table 12.** List of genes used to construct the phylogenetic tree of 263 reference *Bifidobacterium* genomes.

**Supplementary Table 13.** Comparison of manually curated and automated (Prokka, EggNOG-mapper) annotations for 263 reference *Bifidobacterium* genomes.

**Supplementary Table 14.** Pathway enrichment analysis.

**Supplementary Table 15.** List of substrates used in *in vitro* growth experiments.

**Supplementary Table 16.** Raw *in vitro* growth data for 30 *Bifidobacterium* strains.

**Supplementary Table 17.** Comparison between predicted carbohydrate utilization phenotypes and *in vitro* growth data.

**Supplementary Table 18.** Human milk oligosaccharide (HMO) utilization by *Bifidobacterium* strains.

**Supplementary Table 19.** Reconstructed regulons of transcription factors and their predicted operator sequences

**Supplementary Table 20.** Custom probes used to deplete *Bc. kashiwanohense* Bg42221_1E1 rRNA.

**Supplementary Table 21.** Differential gene expression in *Bc. kashiwanohense* Bg42221_1E1 grown on various carbon sources

## Notes

https://doi.org/10.6084/m9.figshare.26053936

https://github.com/Arzamasov/compendium_manuscript

https://github.com/Arzamasov/glycobif

### Summary of Updates

The manuscript has been substantially revised in response to the reviewers' comments.

## References

1. Alessandri, G., van Sinderen, D. & Ventura, M. The genus bifidobacterium: From genomics to functionality of an important component of the mammalian gut microbiota running title: Bifidobacterial adaptation to and interaction with the host. Comput Struct Biotechnol J 19, 1472–1487 (2021).

2. Arboleya, S., Watkins, C., Stanton, C. & Ross, R. P. Gut Bifidobacteria Populations in Human Health and Aging. Front Microbiol 7, 1204 (2016).

3. Stewart, C. J. et al. Temporal development of the gut microbiome in early childhood from the TEDDY study. Nature 562, 583–588 (2018).

4. Sakanaka, M. et al. Evolutionary adaptation in fucosyllactose uptake systems supports bifidobacteria-infant symbiosis. Sci Adv 5, eaaw7696 (2019).

5. Arzamasov, A. A. & Osterman, A. L. Milk glycan metabolism by intestinal bifidobacteria: insights from comparative genomics. Crit Rev Biochem Mol Biol 1–23 (2023) doi:10.1080/10409238.2023.2182272.

6. Kujawska, M. et al. Succession of Bifidobacterium longum Strains in Response to a Changing Early Life Nutritional Environment Reveals Dietary Substrate Adaptations. iScience 23, 101368 (2020).

7. Vatanen, T. et al. A distinct clade of Bifidobacterium longum in the gut of Bangladeshi children thrives during weaning. Cell (2022) doi:10.1016/j.cell.2022.10.011.

8. Orihara, K. et al. Characterization of Bifidobacterium kashiwanohense that utilizes both milk- and plant-derived oligosaccharides. Gut Microbes 15, 2207455 (2023).

9. Taft, D. H. et al. Bifidobacterium Species Colonization in Infancy: A Global Cross-Sectional Comparison by Population History of Breastfeeding. Nutrients 14, 1423 (2022).

10. Olm, M. R. et al. Robust variation in infant gut microbiome assembly across a spectrum of lifestyles. Science 376, 1220–1223 (2022).

11. Derrien, M. et al. Gut microbiome function and composition in infants from rural Kenya and association with human milk oligosaccharides. Gut Microbes 15, 2178793 (2023).

12. Tannock, G. W. et al. Comparison of the compositions of the stool microbiotas of infants fed goat milk formula, cow milk-based formula, or breast milk. Appl. Environ. Microbiol. 79, 3040–3048 (2013).

13. Casaburi, G. et al. Metagenomic insights of the infant microbiome community structure and function across multiple sites in the United States. Sci Rep 11, 1472 (2021).

14. Fukuda, S. et al. Bifidobacteria can protect from enteropathogenic infection through production of acetate. Nature 469, 543–547 (2011).

15. Hirano, R. et al. Next-generation prebiotic promotes selective growth of bifidobacteria, suppressing Clostridioides difficile. Gut Microbes 13, 1973835 (2021).

16. Belenguer, A. et al. Two routes of metabolic cross-feeding between Bifidobacterium adolescentis and butyrate-producing anaerobes from the human gut. Appl. Environ. Microbiol. 72, 3593–3599 (2006).

17. Rios-Covian, D., Gueimonde, M., Duncan, S. H., Flint, H. J. & de los Reyes-Gavilan, C. G. Enhanced butyrate formation by cross-feeding between Faecalibacterium prausnitzii and Bifidobacterium adolescentis. FEMS Microbiol. Lett. 362, (2015).

18. Laursen, M. F. et al. Bifidobacterium species associated with breastfeeding produce aromatic lactic acids in the infant gut. Nat Microbiol 6, 1367–1382 (2021).

19. Shiver, A. L. et al. Genome-scale resources in the infant gut symbiont Bifidobacterium breve reveal genetic determinants of colonization and host-microbe interactions. Cell S0092-8674(25)00195–3 (2025) doi:10.1016/j.cell.2025.02.010.

20. Maldonado-Gómez, M. X. et al. Stable Engraftment of Bifidobacterium longum AH1206 in the Human Gut Depends on Individualized Features of the Resident Microbiome. Cell Host Microbe 20, 515–526 (2016).

21. Frese, S. A. et al. Persistence of Supplemented Bifidobacterium longum subsp. infantis EVC001 in Breastfed Infants. mSphere 2, e00501–17 (2017).

22. Beck, L. C. et al. Strain-specific impacts of probiotics are a significant driver of gut microbiome development in very preterm infants. Nat Microbiol 7, 1525–1535 (2022).

23. Barratt, M. J. et al. Bifidobacterium infantis treatment promotes weight gain in Bangladeshi infants with severe acute malnutrition. Sci Transl Med 14, eabk1107 (2022).

24. Button, J. E. et al. Dosing a synbiotic of human milk oligosaccharides and B. infantis leads to reversible engraftment in healthy adult microbiomes without antibiotics. Cell Host Microbe 30, 712–725.e7 (2022).

25. Milani, C. et al. Bifidobacteria exhibit social behavior through carbohydrate resource sharing in the gut. Sci Rep 5, 15782 (2015).

26. Albert, K., Rani, A. & Sela, D. A. Comparative Pangenomics of the Mammalian Gut Commensal Bifidobacterium longum. Microorganisms 8, (2019).

27. Liu, J., Li, W., Yao, C., Yu, J. & Zhang, H. Comparative genomic analysis revealed genetic divergence between Bifidobacterium catenulatum subspecies present in infant versus adult guts. BMC Microbiol 22, 158 (2022).

28. Arboleya, S. et al. Gene-trait matching across the Bifidobacterium longum pan-genome reveals considerable diversity in carbohydrate catabolism among human infant strains. BMC Genomics 19, 33 (2018).

29. Bottacini, F. et al. Comparative genomics and genotype-phenotype associations in Bifidobacterium breve. Sci Rep 8, (2018).

30. Liu, S. et al. Gene-Phenotype Associations Involving Human-Residential Bifidobacteria (HRB) Reveal Significant Species- and Strain-Specificity in Carbohydrate Catabolism. Microorganisms 9, 883 (2021).

31. Magnúsdóttir, S. et al. Generation of genome-scale metabolic reconstructions for 773 members of the human gut microbiota. Nat Biotechnol 35, 81–89 (2017).

32. Devika, N. T. & Raman, K. Deciphering the metabolic capabilities of Bifidobacteria using genome-scale metabolic models. Sci Rep 9, 18222 (2019).

33. Schöpping, M., Gaspar, P., Neves, A. R., Franzén, C. J. & Zeidan, A. A. Identifying the essential nutritional requirements of the probiotic bacteria Bifidobacterium animalis and Bifidobacterium longum through genome-scale modeling. NPJ Syst Biol Appl 7, 47 (2021).

34. Casey, J. et al. Transporter annotations are holding up progress in metabolic modeling. *Front*. Syst. Biol. 4, (2024).

35. Overbeek, R. et al. The subsystems approach to genome annotation and its use in the project to annotate 1000 genomes. Nucleic Acids Research 33, 5691–5702 (2005).

36. Bottacini, F. et al. Comparative genomics of the Bifidobacterium breve taxon. BMC Genomics 15, 170 (2014).

37. Raman, A. S. et al. A sparse covarying unit that describes healthy and impaired human gut microbiota development. Science 365, (2019).

38. Gehrig, J. L. et al. Effects of microbiota-directed foods in gnotobiotic animals and undernourished children. Science 365, (2019).

39. Smith, M. I. et al. Gut microbiomes of Malawian twin pairs discordant for kwashiorkor. Science 339, 548–554 (2013).

40. Turroni, F. et al. Genome analysis of Bifidobacterium bifidum PRL2010 reveals metabolic pathways for host-derived glycan foraging. Proc Natl Acad Sci U S A 107, 19514–19519 (2010).

41. Mattarelli, P., Bonaparte, C., Pot, B. & Biavati, B. Proposal to reclassify the three biotypes of Bifidobacterium longum as three subspecies: Bifidobacterium longum subsp. longum subsp. nov., Bifidobacterium longum subsp. infantis comb. nov. and Bifidobacterium longum subsp. suis comb. nov. Int J Syst Evol Microbiol 58, 767–772 (2008).

42. Chaplin, A. V. et al. Intraspecies Genomic Diversity and Long-Term Persistence of Bifidobacterium longum. PLoS One 10, (2015).

43. Modesto, M. et al. Bifidobacterium longum subsp. iuvenis subsp. nov., a novel subspecies isolated from the faeces of weaning infants. Int J Syst Evol Microbiol 73, (2023).

44. O’Connell Motherway, M., et al. Characterization of ApuB, an extracellular type II amylopullulanase from Bifidobacterium breve UCC2003. Appl. Environ. Microbiol. 74, 6271–6279 (2008).

45. Kashima, T. et al. Identification of difructose dianhydride I synthase/hydrolase from an oral bacterium establishes a novel glycoside hydrolase family. J Biol Chem 297, 101324 (2021).

46. LoCascio, R. G., Desai, P., Sela, D. A., Weimer, B. & Mills, D. A. Broad conservation of milk utilization genes in Bifidobacterium longum subsp. infantis as revealed by comparative genomic hybridization. Appl Environ Microbiol 76, 7373–7381 (2010).

47. Sakurama, H. et al. Lacto-N-biosidase encoded by a novel gene of Bifidobacterium longum subspecies longum shows unique substrate specificity and requires a designated chaperone for its active expression. J. Biol. Chem. 288, 25194–25206 (2013).

48. Garrido, D. et al. A novel gene cluster allows preferential utilization of fucosylated milk oligosaccharides in Bifidobacterium longum subsp. longum SC596. Sci Rep 6, (2016).

49. Ojima, M. N. et al. Priority effects shape the structure of infant-type Bifidobacterium communities on human milk oligosaccharides. ISME J (2022) doi:10.1038/s41396-022-01270-3.

50. Komeno, M. et al. Two α-L-arabinofuranosidases from Bifidobacterium longum subsp. longum are involved in arabinoxylan utilization. Appl Microbiol Biotechnol 106, 1957– 1965 (2022).

51. Friess, L. et al. Two extracellular α-arabinofuranosidases are required for cereal-derived arabinoxylan metabolism by Bifidobacterium longum subsp. longum. Gut Microbes 16, 2353229 (2024).

52. Tm, G. et al. Characterization and three-dimensional structures of two distinct bacterial xyloglucanases from families GH5 and GH12. The Journal of biological chemistry 282, (2007).

53. J, R., et al. Mechanisms involved in xyloglucan catabolism by the cellulosome-producing bacterium Ruminiclostridium cellulolyticum. Scientific reports 6, (2016).

54. Garrido, D., Kim, J. H., German, J. B., Raybould, H. E. & Mills, D. A. Oligosaccharide Binding Proteins from Bifidobacterium longum subsp. infantis Reveal a Preference for Host Glycans. PLoS One 6, (2011).

55. Sela, D. A. et al. An infant-associated bacterial commensal utilizes breast milk sialyloligosaccharides. J. Biol. Chem. 286, 11909–11918 (2011).

56. Sela, D. A. et al. Bifidobacterium longum subsp. infantis ATCC 15697 α-Fucosidases Are Active on Fucosylated Human Milk Oligosaccharides. Appl Environ Microbiol 78, 795–803 (2012).

57. Garrido, D., Ruiz-Moyano, S. & Mills, D. A. Release and utilization of N-acetyl-d-glucosamine from human milk oligosaccharides by Bifidobacterium longum subsp. infantis. Anaerobe 18, 430–435 (2012).

58. Arzamasov, A. A. et al. Human Milk Oligosaccharide Utilization in Intestinal Bifidobacteria Is Governed by Global Transcriptional Regulator NagR. mSystems e0034322 (2022) doi:10.1128/msystems.00343-22.

59. Egan, M., O’Connell Motherway, M., Ventura, M. & van Sinderen, D. Metabolism of Sialic Acid by Bifidobacterium breve UCC2003. Appl Environ Microbiol 80, 4414–4426 (2014).

60. Egan, M. et al. Cross-feeding by Bifidobacterium breve UCC2003 during co-cultivation with Bifidobacterium bifidum PRL2010 in a mucin-based medium. BMC Microbiol 14, (2014).

61. Pasolli, E. et al. Extensive Unexplored Human Microbiome Diversity Revealed by Over 150,000 Genomes from Metagenomes Spanning Age, Geography, and Lifestyle. Cell 176, 649–662.e20 (2019).

62. Khoroshkin, M. S., Leyn, S. A., Van Sinderen, D. & Rodionov, D. A. Transcriptional Regulation of Carbohydrate Utilization Pathways in the Bifidobacterium Genus. Front Microbiol 7, (2016).

63. Milani, C. et al. Genomic encyclopedia of type strains of the genus Bifidobacterium. Appl Environ Microbiol 80, 6290–6302 (2014).

64. Odamaki, T. et al. Comparative Genomics Revealed Genetic Diversity and Species/Strain-Level Differences in Carbohydrate Metabolism of Three Probiotic Bifidobacterial Species. Int J Genomics 2015, 567809 (2015).

65. Crociani, F., Alessandrini, A., Mucci, M. M. & Biavati, B. Degradation of complex carbohydrates by Bifidobacterium spp. Int J Food Microbiol 24, 199–210 (1994).

66. Forster, S. C. et al. A human gut bacterial genome and culture collection for improved metagenomic analyses. Nat Biotechnol 37, 186–192 (2019).

67. Poyet, M. et al. A library of human gut bacterial isolates paired with longitudinal multiomics data enables mechanistic microbiome research. Nat Med 25, 1442–1452 (2019).

68. Lin, X. et al. The genomic landscape of reference genomes of cultivated human gut bacteria. Nat Commun 14, 1663 (2023).

69. Hitch, T. C. A. et al. HiBC: a publicly available collection of bacterial strains isolated from the human gut. Nat Commun 16, 4203 (2025).

70. Almeida, A. et al. A unified catalog of 204,938 reference genomes from the human gut microbiome. Nat Biotechnol 39, 105–114 (2021).

71. Kim, C. Y. et al. Human reference gut microbiome catalog including newly assembled genomes from under-represented Asian metagenomes. Genome Med 13, 134 (2021).

72. Zeng, S. et al. A compendium of 32,277 metagenome-assembled genomes and over 80 million genes from the early-life human gut microbiome. Nat Commun 13, 5139 (2022).

73. Gounot, J.-S. et al. Genome-centric analysis of short and long read metagenomes reveals uncharacterized microbiome diversity in Southeast Asians. Nat Commun 13, 6044 (2022).

74. Jin, H. et al. A high-quality genome compendium of the human gut microbiome of Inner Mongolians. Nat Microbiol 8, 150–161 (2023).

75. Kim, M. et al. Higher pathogen load in children from Mozambique vs. USA revealed by comparative fecal microbiome profiling. ISME Commun 2, 74 (2022).

76. de Crécy-Lagard, V. et al. A roadmap for the functional annotation of protein families: a community perspective. Database (Oxford) 2022, baac062 (2022).

77. Price, M. N. & Arkin, A. P. Interactive tools for functional annotation of bacterial genomes. Database 2024, baae089 (2024).

78. Zhu, L. et al. Captive Common Marmosets (Callithrix jacchus) Are Colonized throughout Their Lives by a Community of Bifidobacterium Species with Species-Specific Genomic Content That Can Support Adaptation to Distinct Metabolic Niches. mBio e0115321 (2021) doi:10.1128/mBio.01153-21.

79. Chang, H.-W. et al. Prevotella copri and microbiota members mediate the beneficial effects of a therapeutic food for malnutrition. Nat Microbiol 9, 922–937 (2024).

80. Munoz, J., James, K., Bottacini, F. & Van Sinderen, D. Biochemical analysis of cross-feeding behaviour between two common gut commensals when cultivated on plant-derived arabinogalactan. Microb Biotechnol (2020) doi:10.1111/1751-7915.13577.

81. Fernandez-Julia, P., Black, G. W., Cheung, W., Van Sinderen, D. & Munoz-Munoz, J. Fungal β-glucan-facilitated cross-feeding activities between Bacteroides and Bifidobacterium species. Commun Biol 6, 576 (2023).

82. Renwick, S. et al. Modulating the developing gut microbiota with 2’-fucosyllactose and pooled human milk oligosaccharides. Microbiome 13, 44 (2025).

83. Browne, H. P. et al. Boosting microbiome science worldwide could save millions of children’s lives. Nature 625, 237–240 (2024).

84. Davis, J. J. et al. The PATRIC Bioinformatics Resource Center: expanding data and analysis capabilities. Nucleic Acids Res 48, D606–D612 (2020).

85. Markowitz, V. M. et al. IMG: the Integrated Microbial Genomes database and comparative analysis system. Nucleic Acids Res 40, D115–122 (2012).

86. Parks, D. H., Imelfort, M., Skennerton, C. T., Hugenholtz, P. & Tyson, G. W. CheckM: assessing the quality of microbial genomes recovered from isolates, single cells, and metagenomes. Genome Res 25, 1043–1055 (2015).

87. Olm, M. R., Brown, C. T., Brooks, B. & Banfield, J. F. dRep: a tool for fast and accurate genomic comparisons that enables improved genome recovery from metagenomes through de-replication. ISME J 11, 2864–2868 (2017).

88. Parks, D. H. et al. GTDB: an ongoing census of bacterial and archaeal diversity through a phylogenetically consistent, rank normalized and complete genome-based taxonomy. Nucleic Acids Res 50, D785–D794 (2022).

89. Tonkin-Hill, G. et al. Producing polished prokaryotic pangenomes with the Panaroo pipeline. Genome Biol 21, 180 (2020).

90. Katoh, K. & Standley, D. M. MAFFT Multiple Sequence Alignment Software Version 7: Improvements in Performance and Usability. Mol Biol Evol 30, 772–780 (2013).

91. Minh, B. Q. et al. IQ-TREE 2: New Models and Efficient Methods for Phylogenetic Inference in the Genomic Era. Mol. Biol. Evol. 37, 1530–1534 (2020).

92. Letunic, I. & Bork, P. Interactive Tree Of Life (iTOL) v5: an online tool for phylogenetic tree display and annotation. Nucleic Acids Research 49, W293–W296 (2021).

93. Pritchard, L., Glover, R. H., Humphris, S., Elphinstone, J. G. & Toth, I. K. Genomics and taxonomy in diagnostics for food security: soft-rotting enterobacterial plant pathogens. Anal. Methods 8, 12–24 (2015).

94. Seemann, T. Prokka: rapid prokaryotic genome annotation. Bioinformatics 30, 2068–2069 (2014).

95. Overbeek, R. et al. The SEED and the Rapid Annotation of microbial genomes using Subsystems Technology (RAST). Nucleic Acids Res. 42, D206–214 (2014).

96. Cantalapiedra, C. P., Hernández-Plaza, A., Letunic, I., Bork, P. & Huerta-Cepas, J. eggNOG-mapper v2: Functional Annotation, Orthology Assignments, and Domain Prediction at the Metagenomic Scale. Mol Biol Evol 38, 5825–5829 (2021).

97. Zheng, J. et al. dbCAN3: automated carbohydrate-active enzyme and substrate annotation. Nucleic Acids Res gkad328 (2023) doi:10.1093/nar/gkad328.

98. Saier, M. H. et al. The Transporter Classification Database (TCDB): 2021 update. Nucleic Acids Res 49, D461–D467 (2021).

99. Drula, E. et al. The carbohydrate-active enzyme database: functions and literature. Nucleic Acids Res 50, D571–D577 (2022).

100. Kanehisa, M., Sato, Y., Kawashima, M., Furumichi, M. & Tanabe, M. KEGG as a reference resource for gene and protein annotation. Nucleic Acids Res. 44, D457–462 (2016).

101. Novichkov, P. S. et al. RegPrecise: a database of curated genomic inferences of transcriptional regulatory interactions in prokaryotes. Nucleic Acids Res 38, D111–D118 (2010).

102. Steinegger, M. & Söding, J. MMseqs2 enables sensitive protein sequence searching for the analysis of massive data sets. Nat Biotechnol 35, 1026–1028 (2017).

103. Buchfink, B., Xie, C. & Huson, D. H. Fast and sensitive protein alignment using DIAMOND. Nat. Methods 12, 59–60 (2015).

104. Abraham, A. et al. Machine learning for neuroimaging with scikit-learn. Front Neuroinform 8, 14 (2014).

105. Kuhn, M. Building Predictive Models in R Using the caret Package. Journal of Statistical Software 028, (2008).

106. Hibberd, M. C. et al. Bioactive glycans in a microbiome-directed food for children with malnutrition. Nature 625, 157–165 (2024).

107. Oksanen, J. et al. vegan: Community Ecology Package. (2022).

108. Lenth, R. V., et al. emmeans: Estimated Marginal Means, aka Least-Squares Means. (2022).

109. Wickham, Hadley. *Ggplot2: Elegant Graphics for Data Analysis*. (Springer-Verlag, New York, 2016).

110. Gu, Z., Eils, R. & Schlesner, M. Complex heatmaps reveal patterns and correlations in multidimensional genomic data. Bioinformatics 32, 2847–2849 (2016).

111. Gilchrist, C. L. M. & Chooi, Y.-H. clinker & clustermap.js: automatic generation of gene cluster comparison figures. Bioinformatics 37, 2473–2475 (2021).

112. Berger, P. K. et al. Stability of Human-Milk Oligosaccharide Concentrations Over 1 Week of Lactation and Over 6 Hours Following a Standard Meal. J Nutr 152, 2727–2733 (2023).

