## Supplementary Notes 1-5; Supplementary Methods; Supplementary Figs. 1-5 for "Integrative genomic reconstruction reveals heterogeneity in carbohydrate utilization across human gut bifidobacteria"

**Supplementary Note 1: Taxonomic assignments of reference *Bifidobacterium* genomes**

Previous studies have demonstrated that *Bifidobacterium* genomes can be misclassified in public databases[1–5]. Therefore, we used a maximum-likelihood phylogenetic tree based on the alignment of 487 core genes to check the taxonomic assignments of 263 reference *Bifidobacterium* genomes (**Supplementary Fig. 1**; **Supplementary Table 2**). This phylogenomic analysis was complemented with pairwise comparisons of Average Nucleotide Identity (ANI) values for select genomes from *Bifidobacterium longum* and *Bifidobacterium catenulatum* species **(Extended Data Fig. 1**).

The *B. longum* species consists of several distinct subspecies delineated based on the combination of genetic and phenotypic features (i.e., polyphasic taxonomy approach). Most *B. longum* strains of human origin belong to two subspecies: subsp. *longum* (*Bl. longum*) and subsp. *infantis* (*Bl. infantis*)[6]. The type strains of other subspecies, *B. longum* subsp. *suis* (*Bl. suis*)[6] and *B. longum* subsp. *suillium* (*Bl. suillium*)[7], were originally isolated from piglet feces; however, multiple reports[2,8,9,4,5,10,11] have described the genomic similarity of particular human-derived *B. longum* isolates to *Bl. suis*. Furthermore, a recent study proposed reclassifying a subgroup of *Bl. suis* strains, mainly isolated from Bangladeshi children, as *B. longum* subsp. *iuvenis* (*Bl. iuvenis*)[12].

We reproduced the observation that *B. longum* strains 157F, CCUG 52486, and CECT 7210 were incorrectly classified as *Bl. infantis* and instead belong to subsp. *longum*[1–5] based on their position within the phylogenetic tree (**Supplementary Fig. 1**). We also reclassified *B. longum* 981_BLON as *Bifidobacterium scardovii*, and *Bifidobacterium* sp. 12_1_47BFAA as *Bl. longum*. Three *B. longum* strains (1897B, 239-2, and LFYP82) formed a clade separate from *Bl. longum* strains (**Supplementary Fig. 1**), consistent with findings from previous studies[5,13]. *B. longum* JDM301 and APC1461 were classified as *Bl. suis* similarly to previous reports[2,4,5,9]. The *Bl. suis* phylogenetic clade also included four Bangladeshi isolates (BgEED06, Bg131.S11_17.F6, Bg155.S08_5B11, and Bg41121_2E1).

To better delineate the within-species structure of the *B. longum* complex, we compared pairwise ANI values of 15 reference and 13 additional *B. longum* strains, including the type strains of *Bl. suis*, *Bl. suillium*, and *Bl. iuvenis*. Hierarchical clustering of ANI values revealed four clusters corresponding to *Bl. infantis*, *Bl. longum*, a group of *Bl. suis* genomes, and a distinct clade (termed hereafter *Bl. nov.*) comprised strains 1897B, 239-2, and LFYP82 (**Extended Data Fig. 1a**). *Bl. nov* genomes shared 96.8-96.9% ANI with *Bl. longum* JCM 1217$^{T}$, 95.8-95.9% ANI with *Bl. suis* DSM 20211$^{T}$, and 95-95.1% ANI with *Bl. infantis* ATCC

15697$^T$. These results, coupled with the unique distribution of glycan utilization pathways in *Bl. nov.* strains (see **Supplementary Note 5**), suggest that they may represent a candidate subspecies within *B. longum.* We did not observe a clear resolution between *Bl. suis*, *Bl. suillium*, and *Bl. iuvenis* based on ANI values. The minimum pairwise ANI of 97.8% between the genomes across these three subspecies was comparable to strain-level ANI differences within *Bl. infantis*, for example, between strains BT1 and ATCC 15697$^T$ (**Extended Data Fig. 1a**). While these results support the separation of *Bl.* suis group genomes from other *B. longum* subspecies based on ANI, we argue that ANI values alone cannot robustly delineate *Bl. suis, Bl. suillium,* and *Bl. iuvenis*, consistent with similar observations by Albert et al[4].

Previous studies indicated that the *B. catenulatum* species complex comprises two subspecies: subsp. *catenulatum* (*Bc. catenulatum*) and subsp. *kashiwanohense* (*Bc. kashiwanohense*)[14,15]. Our initial phylogenomic analysis did not support a clear separation of 10 reference genomes classified as *B. catenulatum* by NCBI taxonomy[16] into these two subspecies (**Supplementary Fig. 1**). A follow-up hierarchical clustering of pairwise ANI values split the genomes into four clusters (**Extended Data Fig. 1b**). Cluster II contained conventional *Bc. catenulatum* strains (DSM 16992$^T$ and 1899B), whereas cluster III included the type strain of *Bc. kashiwanohense* (JCM 15439$^T$) and a new Bangladeshi isolate (Bg42221_1E1). Cluster I contained three Bangladeshi isolates (SS_Bg39, JG_Bg468, and Bg41721_1F7). Their genomes shared 93.8-94.5% ANI with *Bc. catenulatum* DSM 16992$^T$ and *Bc. kashiwanohense* JCM 15439$^T$, suggesting that these isolates may represent a putative novel *Bifidobacterium* species (*Bifidobacterium sp002742445* in GTDB[17]). Consistent with this notion, this species was recently formalized as *Bifidobacterium hominis* based on a 95% ANI threshold[18].

Cluster IV comprised the PV20-2 strain, previously isolated from an anemic Kenyan infant[19], along with two Bangladeshi isolates (Bg41121_2F9 and Bg42221_1D3) from this work. PV20-2 was previously classified as *Bifidobacterium kashiwanohense* in NCBI Taxonomy and *Bifidobacterium kashiwanohense_A* in GTDB. Genomes within cluster IV shared 94.8-95.6% ANI with *Bc. catenulatum* DSM 16992$^T$ and *Bc. kashiwanohense* JCM 15439$^T$, suggesting that the strains in this cluster might represent a candidate subspecies within *B. catenulatum*. Overall, our phylogenetic analysis suggests the existence of a novel *B. catenulatum* subspecies (*Bc. kashiwanohense_A*) composed of strains isolated from children in understudied non-Westernized populations (e.g., from Kenya and Bangladesh). However, additional studies are needed to test this hypothesis, given the extensive within-species variability of predicted carbohydrate utilization capabilities within *B. catenulatum* and *B. hominis* (see **Supplementary Note 4**).

#### Supplementary Note 2: Reconstruction of transcriptional regulons for carbohydrate metabolism genes

Regulon reconstruction offers a robust framework for predicting novel metabolic functional roles, such as glycan transporters, by leveraging the principle that genes co-regulated by the same transcription factor often encode functional roles involved in the same metabolic pathway[20,21]. Utilizing a previously established regulon reconstruction approach[22–26], we aimed to predict binding sites (operators) of various transcription factors to expand our understanding of transcriptional regulatory networks governing carbohydrate utilization in bifidobacteria.

We first reconstructed the regulon of a putative LacI-family transcription factor, MnbR, in *Bifidobacterium breve* Bg41721_1C11 and *Bifidobacterium dentium* LFYP24. In *B. dentium*, putative MnbR operators were identified in the promoter regions of (i) the *mnbABC-mnb2* operon encoding an ortholog of a biochemically characterized ATP-binding cassette (ABC) transporter for β-mannose and glucomannan oligosaccharides (bMnOS and GmOS, respectively)[27] and a putative β-mannosidase (GH2), (ii) *baMan26A* encoding an ortholog of a characterized extracellular endo-β-1,4-mannanase (GH26)[28], (iii) *manI* encoding a homolog of a characterized D-mannose isomerase (EC 5.3.1.7)[29], (iv) divergently transcribed genes *bdAme2* and *baGlu1* encoding a predicted carbohydrate esterase (CE2) and an ortholog of β-glucosidase (GH1) with a broad substrate-specificity[30], (v) *bdAga36* encoding a putative α-galactosidase (GH36), and (vi) *mnbR* encoding the transcription factor itself (**Extended Data Fig. 7d,g**; **Supplementary Tables 4,19**). The reconstructed MnbR regulon in *B. breve* Bg41721_1C11 contained fewer genes, namely *manI*, *bdAme2*, *baGlu1*, and the *manEFGLM-mnb2* operon. We hypothesize that *manEFGLM* is a non-orthologous replacement for *mnbABC* and encodes a previously uncharacterized five-component ABC transporter involved in the uptake of bMnOS and GmOS into the cell.

The structure of the reconstructed regulon implicates MnbR as a global regulator of bMnOS/GmOS and β-mannan (bMAN) metabolism in bifidobacteria. While the transcriptional effector(s) of MnbR is unknown, we surmise that mannose or short bMnOS may represent likely candidate effectors. The structure of the MnbR regulon allowed us to refine the previous model[27] of bMnOS, GmOS, and bMAN metabolism in bifidobacteria (**Extended Data Fig. 7e**). According to the updated model, strains possessing extracellular endo-β-1,4-mannanases (e.g., BaMan26A) cleave the backbone of various bMAN chains, including gluco- and galactomannan. The released bMnOS and GmOS are imported into the cell by either MnbABC or ManEFGLM transporters and are then hydrolyzed in a step-by-step fashion via a coordinated action of exo-acting β-mannosidase (Mnb2), β-glucosidase (BaGlu1), α-galactosidase (BdAga36), and carbohydrate esterase (BdAme2). The released glucose, galactose, and mannose residues are converted via their respective downstream pathways to fructose-6-phosphate, which enters the bifid shunt. Consistent with the metabolic reconstruction and predicted carbohydrate utilization phenotypes, *B. dentium* LFYP24 grew in MRS-AC supplemented with mannotriose or konjac glucomannan, whereas *B. breve* Bg41721_1C11 grew only in MRS-AC with mannotriose (**Fig. 5**; **Extended Data Fig. 7f**).

We expanded our previous regulon reconstruction for XosR, a LacI-family transcription factor potentially repressing a gene cluster involved in xylooligosaccharide (XOS) and arabinoxylooligosaccharide (AXOS) utilization in *Bifidobacterium adolescentis*, *Bifidobacterium angulatum*, *B. dentium*, and *Bifidobacterium gallicum*[23]. We identified additional candidate XosR-binding sites in the promoter region of an operon starting with the *xosD* gene in *Bc. kashiwanohense* Bg42221_1E1 and *Bifidobacterium pseudocatenulatum* JCM 1200 (**Extended Data Fig. 7b,c**; **Supplementary Table 19**). This operon encoded a biochemically characterized ABC transporter for XOS/AXOS (XosDEFGH) along with several cytoplasmic glycoside hydrolases (GHs) involved in the hydrolysis of XOS and AXOS[31] (**Supplementary Table 4**). The genes constituting the reconstructed XosR regulon were upregulated in *Bc. kashiwanohense* Bg42221_1E1 grown on tamarind xyloglucan (**Fig. 3e**) and in *B. pseudocatenulatum* JCM 1200 grown on xylose, XOS, or AXOS[31], suggesting that xylose may function as the candidate transcriptional effector of XosR. We did not identify putative XosR operators in the promoter regions of *xyn* cluster genes (**Extended Data Fig. 7b**), including *bpXyn10A,* a gene encoding an extracellular endo-β-1,4-xylanase (GH10) required for xylan and arabinoxylan depolymerization[32,33]. Therefore, a different transcription factor (XylR or XosR2) likely regulates the expression of *xyn* genes. Overall, these findings suggest that XosR is a negative regulator of XOS/AXOS utilization in bifidobacteria, whereas the expression of the key enzyme for (arabino)xylan degradation is controlled by a yet unknown mechanism.

We expanded the regulons of two LacI-family transcription factors from *Bl. longum* JCM 1217. The first, BgaR, had been predicted to control a gene cluster involved in beta-galactoside utilization in *Bl. longum*[23,34]. We identified an additional candidate BgaR-binding site in the promoter region of the *bl16Gal-bl13Gal* operon in *Bl. longum* JCM 1217 (**Supplementary Table 19**). These two genes encode extracellular GHs participating in the depolymerization of type II (e.g., larch wood) arabinogalactan (AGII) chains: exo-β-1,6-galactobiohydrolase (GH30_5) and exo-β-1,3-galactanase (GH43_24), respectively[35,36] (**Supplementary Table 4**). Based on the updated regulon structure and improved functional gene annotation, we propose that BgaR regulates AGII metabolism in *Bl. longum* by potentially controlling the expression of genes required for (i) the release of β-1,6-linked (arabino)galactooligosaccharides (*bl16Gal*, *bl13Gal*) from AGII, (ii) transport of the released oligosaccharides into the cell (*gosFGH*, *gosX*)[37,38], and (iii) their subsequent hydrolysis to monosaccharides by cytoplasmic exo-acting β-galactosidase (*bga42A*)[39,40] and α-L-arabinofuranosidase (*abf51B*)[41].

The second LacI-family transcription factor, AraQ, is a bifunctional global regulator that represses arabinose catabolism genes (*araBDA*) and, at the same time, activates multiple central carbohydrate metabolism genes (*gap*, *tkt*, *tal*, *pyk*, *eno*, *ldh*) in various *Bifidobacterium* species[23,24,42]. We identified additional potential AraQ operators in the promoter regions of the following genes in *Bl. longum* JCM 1217: *bll4HypBA1* and *bll3HypBA1* encoding β-L-arabinofuranosidases (GH146)[43,44], *blArafA* encoding α-1,3-L-arabinofuranosidase (GH43_22)[36], and *blArafD* encoding α-1,2/5-L-arabinofuranosidase (GH43_UC_26)[45] (**Supplementary Tables 4,19**). The results indicate that AraQ may also regulate the expression of multiple extracellular GHs involved in the removal of arabinofuranosyl residues from arabinose-containing polysaccharides (AGII, arabinan, and arabinoxylan) and hydroxyproline-rich glycoproteins.

#### Supplementary Note 3: Genomic reconstruction of monosaccharide derivative utilization pathways

We reconstructed metabolic pathways involved in the catabolism of monosaccharide derivatives such as sugar alcohols, sugar acids, and uronic acids. One such derivative, glucuronic acid (GlcA), is a structural component of plant polysaccharides, animal proteoglycans, and various glucuronides. We identified a gene cluster potentially involved in GlcA catabolism in nine *Bl. infantis*, six *B. breve,* one *Bl. longum*, and one *Bl. suis* genome from the reference dataset (**Supplementary Table 8**). This cluster encoded homologs of characterized GlcA pathway enzymes in *E. coli*[46], specifically uronate isomerase (UxaC), D-mannonate oxidoreductase (UxuB), and mannonate dehydratase (UxuA). It also encoded two putative functional roles — a kinase (KdgK) and a dehydrogenase (KdgD) — implemented by two separate genes in *B. breve*, *Bl. longum* and *Bl. suis,* or by a single gene fusion in *Bl. infantis* strains (**Extended Data Fig. 4a**). In *Bl. infantis*, the GlcA cluster contained genes for two MFS permeases, *exuT* and a truncated *exuP*. Other taxa lacked the *exuT* gene but harbored a large adjacent gene cluster termed *gus*. The *gus* cluster encoded multiple GHs from various families (30, 59, 78, and 154), an ABC transporter, and a full-sized ExuP (**Extended Data Fig. 4a**).

Based on the gene distribution pattern, we predicted that *Bl. infantis* strains could utilize GlcA by transporting it via the ExuT permease, whereas *B. breve*, *Bl. longum* and *Bl. suis* strains could utilize glucuronides (phenotype GluA), from which cytoplasmic β-glucuronidases (GH30[47] and GH154) would cleave terminal GlcA residues (**Extended Data Fig. 4b**). The freed GlcA would then be converted to 2-dehydro-3-deoxy-D-gluconate (KDG) by UxaC, UxuB, and UxuA. The exact metabolic fate of KDG is unclear since all reference strains lacked genes encoding homologs of 2-dehydro-3-deoxygluconate kinase and 2-dehydro-3-deoxy-6-phosphogluconate aldolase characterized in *E. coli*[46] and identified in certain non-human bifidobacteria[48]. We surmise that KdgK may phosphorylate KDG to 2-dehydro-3-deoxy-6-phosphogluconate, which is then likely shunted to central metabolism by KdgD via a yet unknown mechanism. Consistent with our genomic predictions, *Bl. infantis* strains carrying GlcA utilization genes grew in MRS-AC supplemented with GlcA (**Fig. 5**; **Extended Data Fig. 4c**), aligning with a prior study that reported GlcA fermentation only by *Bl. infantis*[49]. By extending the analysis to 3,083 *Bifidobacterium* genomes, we identified putative GlcA and GluA utilization pathways in 107 (predominantly *Bl. infantis*) and 63 (predominantly *B. breve*) genomes, respectively (**Fig. 2c**; **Supplementary Tables 7-10**). These findings suggest that GlcA and GluA metabolism is associated with infant-colonizing *Bifidobacterium* species; however, further studies are needed to clarify the structure of the underlying catabolic pathways.

Sugar alcohols (polyols) are naturally present in small quantities in fruits and vegetables and some are widely used as food additives (sweeteners). Bifidobacteria can ferment specific polyols, D-mannitol (Mtl) and D-sorbitol (Stl)[50,51], and the gene clusters involved in their utilization have previously been previously identified via genotype-to-phenotype matching[51,52] (**Extended Data Fig. 4d**). The structures of reconstructed Mtl and Stl utilization pathways are similar; both include the uptake of respective polyol molecules by major facilitator superfamily (MFS) transporters (MtlP or StlP) followed by oxidation to D-fructose by respective dehydrogenases (MtlD or GutB; **Extended Data Fig. 4e**). D-fructose is then phosphorylated to fructose-6-phosphate by fructokinase (Frk)[53]. In agreement with the predicted utilization phenotypes, all tested strains with a complete set of Mtl or Stl utilization genes grew in MRS-AC supplemented with Mtl or Stl, respectively (**Fig. 5**). Mtl and Stl utilization pathways were identified in 1,082 and 1,545 out of 3,083 *Bifidobacterium* genomes, respectively, indicating that the predicted capacity to metabolize these sugar alcohols is relatively widespread yet varies at the strain level (**Fig. 2c**; **Supplementary Tables 7-10**).

Inositol (Ino) is a polyol found in various mammalian and plant tissues in free (e.g., *myo*-inositol) and various conjugated (e.g., phytic acid) forms. We identified a gene cluster encoding homologs of characterized Ino catabolism enzymes from *Bacillus subtilis* and *Thermotoga maritima* in seven reference *Bl. infantis* genomes. These enzymes included two distinct *myo*-inositol 2-dehydrogenases (IolG1/IolG2)[54,55], *scyllo*-inosose dehydratase (IolE)[56], 3D-(3,5/4)-trihydroxycyclohexane-1,2-dione hydrolase (IolD)[57], and 5-deoxy-glucuronate isomerase (IolB) (**Extended Data Fig. 4d**; **Supplementary Table 8**). The identified enzymes allow the sequential catabolism of *myo*-inositol to 5-dehydro-2-deoxy-D-gluconate (**Extended Data Fig. 4e**). We, however, did not find the homologs of enzymes that catalyze the last two steps in the conventional Ino catabolism pathway from *Ba. subtilis*[57], 5-dehydro-2-deoxy-D-gluconate kinase (IolC) and 5-dehydro-2-deoxyphosphogluconate aldolase (IolJ), in *Bifidobacterium* genomes. This observation suggests that *Bl. infantis* either possesses non-orthologous gene displacements encoding these two enzymatic activities (yet to be identified) or metabolizes 5-dehydro-2-deoxy-D-gluconate via a yet unknown pathway that may involve putative dehydrogenases (IolP, IolQ, InoD), kinase (InoK) and transketolase (InoXn, InoXc) encoded in the same gene cluster (**Supplementary Tables 4,8**). Among the two tested *Bl. infantis* strains carrying Ino utilization genes, one exhibited weak growth in MRS-AC supplemented with Ino (**Fig. 5**). This result suggests that Ino does not support high biomass accumulation when used as a substrate, possibly because its catabolic pathway is not connected to the bifid shunt. Extending the analysis to 3,083 *Bifidobacterium* genomes, we found the putative Ino utilization pathway in 76 *Bl. infantis* and one *Bifidobacterium bifidum* genome (**Fig. 2c**; **Supplementary Tables 7-10**). These results suggest that the ability to catabolize Ino is confined to infant-colonizing *Bifidobacterium* species.

D-gluconate (Gco) is a sugar acid that naturally occurs in many foods (fruits, honey, and wine) and is used as a food additive (acidity regulator). We identified a gene cluster encoding homologs of characterized Gco utilization enzymes from *Ba. subtilis* and *Gluconobacter oxydans* in 56 reference *Bifidobacterium* genomes (**Extended Data Fig. 4d**; **Supplementary Table 8**). These elements included D-gluconate MFS permease (GntT)[58], gluconokinase (GntK)[59], and 6-phosphogluconate dehydrogenase (Gnd)[59]. Based on the reconstructed pathway, Gco is transported into the cell by GntT, phosphorylated to 6-phosphogluconate by GntK, and decarboxylated to ribulose-5-phosphate by Gnd (**Extended Data Fig. 4e**). The latter compound is isomerized to xylulose-5-phosphate, which enters the bifid shunt. Supporting the reconstruction, all tested strains with a complete set of Gco utilization genes grew in MRS-AC supplemented with Gco (**Fig. 5**). We identified the putative Gco utilization pathway in 1,016 out of 3,083 *Bifidobacterium* genomes, suggesting that while the capacity to metabolize this sugar acid is relatively common, it remains strain-dependent (**Fig. 2c**; **Supplementary Tables 7-10**).

D-galactonate (Glt) is a sugar acid found in mammalian tissues and body secretions and produced by various bacteria as an intermediate of galactose metabolism[60]. We identified a putative Glt utilization gene cluster in *B. breve* Bg41721_1C11 (**Supplementary Table 8**). This cluster encoded homologs of galactonate dehydratase (DgoD), 2-dehydro-3-deoxygalactonokinase (DgoK), 2-dehydro-3-deoxyphosphogalactonate aldolase (DgoA) from *E. coli*[61], and a predicted ABC transport system (DgoEFG; **Extended Data Fig. 4d**). We hypothesize that Glt is transported into the cytoplasm by DgoEFG and then dehydrated to 2-dehydro-3-deoxy-D-galactonate by DgoD. The latter intermediate is phosphorylated to 2-dehydro-3-deoxy-D-galactonate-6-phosphate by DgoK and then further cleaved by DgoA to D-glyceraldehyde 3-phosphate and pyruvate (**Extended Data Fig. 4e**). The putative Glt utilization pathway was identified in a single reference strain (*B. breve* Bg41721_1C11) and three additional genomes from the extended dataset (**Supplementary Tables 7-10**), making this pathway the rarest among 68 studied. Nevertheless, further research is required to characterize the Glt pathway and its potential importance in bifidobacteria.

L-ascorbate (Asc) is a conjugate base of L-ascorbic acid (vitamin C), an essential cofactor involved in multiple metabolic processes. We identified a putative Asc utilization gene cluster in two reference *Bifidobacterium scardovii* genomes (**Supplementary Table 8**). This cluster encoded homologs of downstream Asc utilization enzymes from *E. coli,* namely 3-keto-L-gulonate 6-phosphate decarboxylase (SgbH), L-xylulose 5-phosphate 3-epimerase (SgbU), and L-ribulose-5-phosphate 4-epimerase (AraD)[62,63], as well as a putative kinase (AscK), lactonase (AscG), and a permease (AscT) (**Extended Data Fig. 4d**). We surmise that AscT transports Asc into the cell, while AscK functions as an ascorbate kinase, phosphorylating Asc to ascorbate-6-phosphate, which AscG then converts to 3-keto-L-gulonate 6-phosphate (**Extended Data Fig. 4e**). The latter compound would be converted to D-xylulose-5-phosphate by a coordinated action of SgbH, SgbU, and AraD (**Extended Data Fig. 4e**). Overall, the proposed bifidobacterial pathway resembles the anaerobic Asc utilization pathway in *E. coli*, except that Asc transport and phosphorylation in *E. coli* are mediated by a phosphoenolpyruvate:carbohydrate phosphotransferase system (PTS)[64]. Among 3,083 genomes, the putative Asc utilization pathway was identified only in *B. scardovii* and

*Bifidobacterium tsurumiense* (**Fig. 2c**; **Supplementary Tables 7-10**). Additional studies are required to assess its functional relevance and confirm its enzymatic architecture.

**Supplementary Note 4: Analysis of metabolic pathway representation in *Bifidobacterium catenulatum* and *Bifidobacterium hominis* genomes**

Phylogenomic and ANI analyses of reference genomes revealed the presence of three subspecies (*Bc. catenulatum*, *Bc. kashiwanohense*, *Bc. kashiwanohense_A*) within the *B. catenulatum* species and indicated their phylogenetic relatedness with recently proposed *Bifidobacterium hominis*[18]. To explore potential phenotypic differences among these taxa, we examined the representation of various metabolic pathways in 110 *B. catenulatum* and 26 *B. hominis* genomes (**Supplementary Tables 2,3**).

Hierarchical clustering of the binary representation of predicted glycan utilization pathways revealed two distinct clusters among the *B. hominis* genomes (**Extended Data Fig. 3**). One cluster, comprising 14 genomes and including the type strain CLA-AA-H311[18], diverged from the *B. catenulatum* species complex due to the absence of pathways for XOS, AXOS, and several fucosylated HMOs (FHMOs) — 2'-fucosyllactose (2'FL), 3-fucosyllactose (3FL), lactodifucotetraose (LDFT), lacto-*N*-fucopentaose I (LNFP I). In contrast, the remaining 12 *B. hominis* genomes harbored pathways for XOS/AXOS, FHMOs, or both, showing a profile more similar to *Bc. kashiwanohense_A*. These results suggest the existence of two phenotypically distinct "ecotypes"[65] within *B. hominis*: one seemingly adapted to thrive in children still receiving breast milk through the capacity to utilize FHMOs and another more likely specialized for the adult gut. The marked within-species heterogeneity also complicates efforts to delineate *B. hominis* from *B. catenulatum* based solely on glycan utilization capabilities.

*Bc. kashiwanohense* had significantly higher predicted phenotypic richness than *Bc. catenulatum*, but not *Bc. kashiwanohense_A* (Poisson generalized linear model, $P = 1.23 \times 10^{-18}$ and $P = 1$, Bonferroni-corrected post-hoc test, **Fig. 2b**). Specifically, pathways involved in the utilization of FHMOs and certain plant-derived oligosaccharides (bMnOS, GmOS, and lcFOS) were present in over 94% of *Bc. kashiwanohense* genomes yet absent in more than 95% of *Bc. catenulatum* genomes (**Extended Data Fig. 3**). We also observed substantial strain-level heterogeneity within each *B. catenulatum* subspecies. For instance, the predicted ability to depolymerize plant hemicelluloses xylan (XL) and arabinoxylan (AX) into XOS/AXOS via endo-β-1,4-xylanase BpXyn10[32,33] varied among *Bc. catenulatum, Bc. kashiwanohense,* and *Bc. kashiwanohense*_A. Among other notable examples, *Bc. kashiwanohense* Bg42221_1E1 was the only genome with the H1 gene cluster, that conveys the capability to metabolize multiple HMOs, including lacto-*N*-neotetraose or LNnT, long-chain fucosylated and sialylated HMOs (**Fig. 4**). It was also one of only three genomes encoding the xyloglucan (XGL) degradation pathway (**Fig. 3**).

Most *B. hominis* and *B. catenulatum* genomes encoded similar conserved sets of B vitamin and amino acid biosynthesis pathways (**Extended Data Fig. 6**), suggesting minimal differences in essential micronutrient production. However, a notable exception was the presence of a complete riboflavin (vitamin B2) pathway in 4 of 23 *Bc. kashiwanohense_A* genomes (**Supplementary Table 11**). Additionally, 11 of 23 *Bc. kashiwanohense_A* genomes encoded a urea utilization pathway homologous to one characterized in *Bl. infantis*[66] and previously unreported in the *B. catenulatum* species complex. These observations highlight the presence of unique metabolic pathways, extending beyond carbohydrate metabolism, in genomes recovered from non-Westernized populations.

Our findings highlight notable within-species heterogeneity in carbohydrate metabolism among *B. hominis* and *B. catenulatum* complexes. In particular, *Bc. kashiwanohense*, *Bc. kashiwanohense_A*, and a subset of *B. hominis* strains exhibit a more expansive repertoire of glycan utilization pathways than *Bc. catenulatum* and appear adapted to thrive in the gut of weaning children. However, these contrasts are partly obscured by considerable strain-level variability, making it difficult to delineate subspecies boundaries based solely on glycan utilization profiles.

#### Supplementary Note 5: Analysis of metabolic pathway representation in *Bifidobacterium longum* genomes

Phylogenomic and ANI analyses suggested that three reference *B. longum* genomes (1897B, 239-2, and LFYP82) form a distinct clade within the *Bifidobacterium longum* species (*Bl. nov.*). To assess potential phenotypic differences between *Bl. nov.* and other *B. longum* subspecies, we examined the distribution of various metabolic pathways across 34 *Bl. nov.* genomes.

All 34 genomes encoded the same set of complete biosynthetic pathways for B vitamins (B1, B3, B6, and B9) and amino acids as genomes of their closest phylogenetic relative, *Bl. longum* (**Extended Data Fig. 6**). In contrast, we observed stark differences in the representation of predicted glycan utilization pathways between *Bl. nov*. and other *B. longum* subspecies. The total number of carbohydrate utilization pathways in *Bl. nov*. was significantly lower compared to *Bl. infantis*, *Bl. longum*, and *Bl. suis* (Poisson generalized linear model,

Bonferroni-corrected post-hoc test, $P = 6.38 \times 10^{-27}$, $P = 2.69 \times 10^{-19}$, $P = 1.04 \times 10^{-8}$, respectively; **Fig. 2b**). One of the striking observations was the absence of the lacto-*N*-biose/galacto-*N*-biose (LNB/GNB) utilization pathway in all 34 *Bl. nov*. genomes (**Fig. 2c**; **Supplementary Tables 7-10**). This pathway, which enables the metabolism of LNB and GNB[67,68], structural components of type I HMOs and mucin-*O*-glycans, respectively, was conserved in all other *B. longum* genomes (**Fig. 2c**). All *Bl. nov*. genomes also lacked complete catabolic pathways for *N*-glycans[29,69], simple *O*-glycans like T-antigen (Tan)[70], and various HMOs (lacto-*N*-tetraose or LNT, LNnT, 2'FL, 3FL, LDFT, LNFP I)[69,71]. These findings indicate that *Bl. nov*. has a reduced capacity to catabolize host-derived glycans, such as those found in human milk.

At the same time, *Bl. nov*. had glycan utilization capabilities absent in other *B. longum* subspecies. All 34 genomes encoded the orthologs of the characterized extracellular amylopullulanase ApuB (GH13_14_32), a bifunctional enzyme that hydrolyzes both α-1,4 and α-1,6-glycosidic bonds in soluble starch (ST) and pullulan (PUL)[72,73] (**Supplementary Tables 8,10**). This suggests that, unlike other *B. longum* subspecies, *Bl. nov.* can degrade α-glucans of plant (ST) and fungal (PUL) origin (**Fig. 2c**), a prediction experimentally validated for the strain LFYP82 (**Fig. 5**). Additionally, all *Bl. nov.* genomes encoded a putative pathway for difructose anhydride (DFA) metabolism previously characterized in *Bifidobacterium dentium*[74]. These findings indicate that *Bl. nov.* harbors distinct genomic signatures related to glycan metabolism, providing a clear basis for differentiating this clade from other *B. longum* subspecies.

Our phylogenetic analysis of *Bl. suis* group genomes suggested that *Bl. suis, Bl. suillium,* and *Bl. iuvenis* could not be robustly delineated based on ANI values. Previous studies used the distribution of specific gene clusters and phenotypic traits to facilitate characterizing these subspecies. For instance, *Bl. suillium* was distinguished from *Bl. suis* based on the absence of urease activity[7], whereas *Bl. iuvenis* strains were separated based on the presence of a vitamin B2 biosynthesis cluster and the ability to metabolize 3FL[12]. We therefore analyzed the representation of pathways involved in carbohydrate and urea utilization, as well as B vitamin and amino acid biosynthesis, across 19 *Bl. suis* group genomes of cultured isolates, including the non-human type strains of *Bl. suis* and *Bl. suillium*. All 19 genomes encoded biosynthetic pathways for vitamins B1, B3, B6, and B9 and all amino acids except Cys (**Extended Data Fig. 5b**). Consistent with the previous study[12], all *Bl. iuvenis* strains additionally harbored the B2 biosynthesis pathway similarly to *Bl. infantis*. The urea utilization pathway was sporadically represented across the *Bl. suis* group. Specifically, it was absent in two out of nine *Bl. iuvenis* strains. These results suggest that the urease activity is not a reliable marker for subspecies delineation, as previously noted[4].

We observed substantial variation in the distribution of predicted glycan utilization pathways in 19 *Bl. suis* group strains. The Bg131.S11_17.F6 strain, isolated from a Bangladeshi infant, exhibited a *Bl. infantis*-like pattern of pathway representation (**Extended Data Fig. 5a**; **Supplementary Fig. 3**), most notably characterized by the presence of the H1 gene cluster implicated in the utilization of multiple neutral, neutral fucosylated and sialylated HMO species[75] (**Fig. 4a**; **Supplementary Table 8**). In contrast, this strain lacked catabolic pathways for plant-derived sugars, such as xylose (Xyl), arabinose (Ara), and type II arabinogalactan (AGII), which were characteristic of *Bl. longum* and other *Bl. suis* group strains (**Extended Data Fig. 5a**; **Supplementary Fig. 3**). However, unlike *Bl. infantis*, *Bl. suis* Bg131.S11_17.F6 lacked genomic loci FL1 and FL2 that encode characterized[71] ABC transport systems for FHMOs (2'FL, 3FL, LDFT, and LNFP I; **Supplementary Table 8**).

The predicted carbohydrate utilization capabilities of the three other *Bl. suis* isolates of Bangladeshi origin (Bg41121_2E1, BgEED06, Bg155.S08_5B11) were notably different. These strains lacked a complete H1 cluster and instead possessed an FHMO utilization locus, which orthologs have been previously described in *B. breve*, *Bc. kashiwanohense*, *Bl. longum,* and *B. pseudocatenulatum*[76–80] (**Fig. 4a**). While both clusters encoded α-fucosidases (BiAfcA and BiAfcB) and downstream fucose catabolism enzymes, they differed in the representation of transporters and other GHs. The H1 cluster in Bg131.S11_17.F6 encoded five putative ABC transporters for HMOs and α-sialidase NanH2[81,82], whereas the FHMO cluster harbored the 2-III variant of the ABC transporter for fucosylated HMOs (FL2_ABC), which facilitates the uptake of 2'FL, 3FL, LDFT, LNFP I, lacto-*N*-fucopentaose II (LNFP II), and lacto-*N*-difucohexaoses I/II (LNDFH I/II)[79] (**Supplementary Fig. 5**). Consistent with these genomic differences, HMO glycoprofiling revealed distinct utilization profiles among Bangladeshi *Bl. suis* group strains. Strain Bg131.S11_17.F6 depleted over 80% of LNT, LNnT, lacto-*N*-hexaose (LNH), 3FL, LNFP I/II, 6'-sialyllactose (6'SL), sialyllacto-N-tetraoses b/c (LST b/c), and longer fucosylated and sialylated HMO chains, but did not efficiently metabolize 2'FL and LDFT (**Fig. 4c**). In contrast, strain Bg41121_2E1 efficiently consumed LNT and FHMOs (2'FL, 3FL, LDFT, LNFP I/II, LNDFH) but not LNnT, LNH, and sialylated structures.

Additionally, *Bl. suis* Bg41121_2E1, BgEED06, and Bg155.S08_5B11 possessed conserved pathways for the utilization of Xyl, Ara, XOS, long-chain fructooligosaccharides (lcFOS), and AGII, which were absent in Bg131.S11_17.F6 (**Extended Data Fig. 5a**). This enhanced capacity to metabolize plant-derived mono-, oligo-, and polysaccharides was validated for *Bl. suis* Bg41121_2E1*,* which, unlike Bg131.S11_17.F6, demonstrated growth on xylose, arabinose, xylotriose, arabinotriose, chicory fructooligosaccharides, and sugar beet arabinan (**Fig. 5**). The overall representation of glycan utilization pathways in *Bl. suis* Bg41121_2E1, BgEED06, and Bg155.S08_5B11 closely resembled that of *Bl. longum* strains (e.g., APC1478 and SC596; **Extended Data Fig. 5a**; **Supplementary Fig. 3**). Some subtle distinctions of these Bangladeshi strains included the presence of the fucose (Fuc) utilization pathway[78], and the lack of specific extracellular α-L-arabinofuranosidases (BlArafD, BlArafE, AxuA, AxuB) required for the degradation of AX[83,84] (**Supplementary Tables 4,8**).

Our results highlight that the phenotypic differences among *Bl. suis* group strains are comparable to those observed between distinct *B. longum* subspecies, such as *Bl. longum* and *Bl. infantis*, and likely reflect adaptation to different ecological niches. For example, strain Bg131.S11_17.F6 appears to share a niche with *Bl. infantis* in breastfed infants, as it can utilize a broad range of HMOs, including long-chain fucosylated and sialylated structures, but cannot metabolize Xyl- and Ara-containing plant glycans (ecotype I). In contrast, strains Bg41121_2E1, BgEED06, and Bg155.S08_5B11 can efficiently forage FHMOs and a broader set of plant mono-, oligo-, and polysaccharides, demonstrating an adaptation to the gut of weaning children consuming a diet rich in both milk and plant glycans (ecotype II). Bangladeshi isolates with similar traits were recently proposed as a separate subspecies, *Bl. iuvenis*, distinct from other *Bl. suis* group strains of human origin[12]. However, the substantial strain-level phenotypic heterogeneity within Bangladeshi isolates demonstrated in this study challenges the previous delineation of *Bl. iuvenis* as a distinct subspecies within *B. longum*. Instead, our results support grouping *Bl. iuvenis*, *Bl. suis*, and *Bl. suillium* into a single subspecies (*Bl. suis*), characterized by high genomic heterogeneity and plasticity, as previously noted by Albert et al[4].

**Supplementary methods**

**Isolation and sequencing of *Bifidobacterium* strains from Bangladeshi and Malawian donors**

*Culturing.* Fecal samples were collected from 6-24-month-old Bangladeshi and Malawian children from the cohorts described in the previous studies[85–87]. Samples were pulverized in liquid nitrogen, and a ~0.1 g aliquot of each sample was transferred to a chamber maintained with a gas mix of 5% $H_2$, 20% $CO_2$, and 75% $N_2$ (Coy Laboratory Products). Samples were diluted 1:10 (wt/vol) with PBS supplemented with 0.05% L-cysteine-HCl in 50 mL conical plastic tubes containing 5 mL of 2 mm-diameter glass beads (VWR International). Tubes were gently vortexed, and the resulting slurry was passed through a 100 μm-pore diameter nylon cell strainer (BD Falcon). A 500 μL aliquot of each processed fecal sample was added to 4.5 mL of PBS, and a 1:10, 1:100, 1:1000, and 1:10,000 dilution series was prepared. LYBHI (brain-heart infusion medium supplemented with 0.5% yeast extract) agar plates were streaked with 100 μL of each dilution. Plates were incubated for 2-3 days at 37 °C. Colonies were picked into 96 deep-well plates (Thermo Fisher Scientific) containing 600 μL of Wilkins-Chalgren broth and incubated overnight at 37°C. Isolate stocks were prepared by combining 50 μL of culture with 50 μL of 30% glycerol in PBS in shallow 96-well plates. Stocks were then frozen at −80 °C for future use. A 500 μL aliquot of each culture was transferred to 2 mL screw cap tubes and pelleted by centrifugation. The resulting supernatant was discarded, and DNA was extracted from pellets with phenol:chloroform. V4-16S rRNA gene amplicons were generated by PCR and sequenced (paired-end 250-bp reads) on the Illumina MiSeq (Illumina). Clonal isolates whose V4-16S rRNA gene amplicon sequences shared ≥ 97% sequence identity with *Bifidobacterium* spp. were subjected to full-length 16S rRNA gene sequencing using primers 8F (AGAGTTTGATCCTGGCTCAG) and 1391R (GACGGGCGGTGTGTRCA).

*Illumina sequencing*. Cryopreserved stocks were streaked onto De Man, Rogosa, and Sharpe (MRS) agar plates and incubated at 37 °C under anoxic conditions twice. Single colonies were picked, inoculated into 6 mL of MRS medium, and cultured to the late log phase. Cell pellets were recovered by centrifugation, and genomic DNA was isolated using phenol:chloroform extraction. Libraries for shotgun sequencing were constructed using the TruSeq Nano DNA Library Prep Kit (Illumina), pooled, and sequenced (paired-end 150-bp reads) on the Illumina NextSeq (Illumina).

*PacBio* and *Nanopore sequencing*. Cryopreserved stocks were streaked onto Brain Heart Infusion (BHI) agar plates and incubated at 37 °C under anoxic conditions overnight. Single colonies were picked, inoculated into 6 mL of MRS medium, and cultured for two days.

Cell pellets were recovered by centrifugation, and genomic DNA was recovered using the MagAttract HMW DNA Extraction Kit (Qiagen). Libraries for PacBio long-read sequencing were prepared using the SMRTbell Template Prep Kit v1.0 (PacBio) + Barcoded Adapter Kit v8A (PacBio) and sequenced on the PacBio Sequel System (PacBio) with a mean ± SD read length of 3,681 ± 861 bp. Libraries for Nanopore long-read sequencing were prepared using the SQK-NBD114.24 native barcoding kit (Nanopore) in accordance with the manufacturer's protocol. Sequencing was carried out on the MinION Mk1B using a FLO-MIN114 flow cell (Nanopore) with a Q-score threshold of 8.

*Read processing and genome assembly.* Raw Illumina reads were demultiplexed via bcl2fastq (v1.3.0) and pre-processed to remove low-quality bases and reads using Trim Galore (v0.4.5)[88] or bbtools (v38.26)[89]. Quality-controlled reads were then subsampled to a depth of ~100-fold coverage using bbtools (v38.26)[89]. Raw PacBio reads were demultiplexed and converted to FASTQ format (SMRT Tools software, v5.1.0 or 6.0.0). Raw Nanopore reads were basecalled using Dorado (v0.3.0) employing the high accuracy calling model (dna_r10.4.1_e8.2_400bps_HAC@v4.2.0). The basecalled reads were demultiplexed with Guppy (v6.5.7) and subsequently filtered using Filtlong (v0.2.1) (top 90% retained, minimum length = 1000 bp) or Nanofilt (v2.8.0)[90] (q=10, minimum length = 1000 bp).

For short-read assembly, paired-end Illumina reads corresponding to each genome were assembled using SPAdes (v3.13.0) with the "—careful" option[91]. Hybrid Illumina/PacBio assemblies were constructed via Unicycler (v0.4.7)[92]. Hybrid Illumina/Nanopore assemblies were constructed using Unicycler (v0.5.0) with SPAdes (v3.15.2) and Racon (v1.5.0)[93]. For *Bc. kashiwanohense* Bg42221_1E1, long-read-only assemblies were constructed using Trycycler (v0.5.0)[94]. Briefly, trimmed reads were subset into 16 non-identical sets and assembled using four separate assemblers: Flye (v2.9.3)[95], Minimap and Miniasm (v0.1.3)[96], Canu (v2.2)[97], and Raven (v1.8.3)[98]. Assemblies with more than one contig were discarded, and the remaining were clustered, checked for circularity, and then aligned to form a consensus long-read assembly. The resulting assembly was polished using Medaka (v1.11.2) (long reads) followed by Polypolish (v0.5.0)[99] (short reads). Assembly quality statistics were generated using Quast (v4.5 or 5.2)[100]. Open reading frames were identified and annotated using Prokka (v1.12 or 1.14.6)[101].

##### HMO isolation from pooled donor human milk

Pooled human milk oligosaccharides (pHMOs) were isolated from donor milk collected from multiple individuals. After centrifugation, the lipid layer was removed, and proteins were precipitated from the aqueous phase by the addition of ice-cold ethanol and subsequent centrifugation. Ethanol was removed from the HMO-containing supernatant by roto-evaporation. Lactose and salts were removed by gel filtration chromatography over a BioRad P2 column (100 cm x 316 mm, Bio-Rad) using a semi-automated fast protein liquid chromatography (FPLC) system. Only pHMO preparations containing <2% lactose were used in bacterial growth assays.

**Reconstruction of transcriptional regulons**

We used a comparative genomics approach previously applied for the *de novo* prediction of operator sequences and regulon reconstruction in bifidobacteria. For transcription factors with known regulons, existing positional weight matrices (PWMs) were retrieved from prior studies[23,24,26] and the RegPrecise database[34]. For *de novo* inference of novel PWMs, we first identified candidate transcription factors, which genes clustered alongside carbohydrate utilization genes. For each regulator and its putative targets, upstream intergenic regions were extracted and aligned using Pro-Coffee[102]. Conserved regions within these alignments were analyzed with SignalX (v1.0)[103], a motif discovery tool based on the expectation-maximization algorithm. We focused on palindromic motifs (inverted repeats) ranging from 16 to 24 bp in length. Identified motifs were used to construct PWMs, which were then applied to scan upstream regions (up to −350 bp from the translation start site) of all genes in genomes encoding the corresponding TF orthologs, using Genome Explorer (v1.0)[103]. Site detection thresholds were set based on the lowest PWM score among the training set sequences. Candidate operator sequences were further validated by the phylogenetic footprinting approach. Consensus transcription factor binding motifs were visualized using WebLogo (v2.8.2)[104].

**RNA isolation**

Frozen cell pellets from 2 mL of early exponential phase cultures ($OD_{600}$=0.35) were resuspended in a solution containing 710 µL of the extraction mixture (200 mM NaCl, 20 mM EDTA, 20% SDS), 500 µL of phenol:chloroform:isoamyl alcohol (125:24:1, pH 4.5), and 250 µL of acid-washed glass beads (212-300 µm; Sigma). Cells were disrupted using the Bead Ruptor 12 (Omni International) by alternating 2 min of homogenizing at 6 m/s and 2 min of cooling on ice. Cellular debris was removed by centrifugation (16,000 × g for 10 min), and the aqueous phase was collected. RNA was precipitated with isopropanol and sodium acetate (pH 5.5) at −20 °C overnight, washed twice in ice-cold 70% ethanol, and resuspended in nucleasefree water. Total RNA was subjected to two consecutive DNase treatments (30 min each) with the Turbo DNase (Ambion) and the Baseline-ZERO DNase (Lucigen), respectively. After each treatment, RNA was purified using the MEGAclear Transcription Clean-Up Kit (Ambion). RNA was quantified via the Qubit Broad Range RNA Assay Kit (Invitrogen). RNA integrity was assessed by the 4200 TapeStation System (Agilent). All samples had RIN values exceeding 8.0.

**RNA-seq data processing**

Raw reads were demultiplexed and quality-checked via FastQC (v0.11.9)[105]. Low-quality bases, Illumina sequencing adapters, and short reads (< 20 bp) were trimmed by Cutadapt (v4.1)[106]. Filtered reads were aligned to rRNA gene sequences extracted from the *Bc. kashiwanohense* Bg42221_1E1 genome using Bowtie2 (v2.4.5)[107]. Reads not mapping to rRNA genes were pseudoaligned to the *Bc. kashiwanohense* Bg42221_1E1 transcriptome using Kallisto (v0.48)[108]. All downstream analyses were carried out using R (v4.3.2) and Bioconductor (v3.15)[109]. Transcript abundance estimates from Kallisto were imported into R via the *tximport* package (v1.30.0)[110] and normalized using the trimmed mean of M-values (TMM) method implemented in the *edgeR* package (v4.0.16)[111]. Genes were filtered to retain those with count per million (CPM) > 1 in at least three samples. Principal component analysis (PCA) was performed on TMM-normalized count data to assess potential batch effects. Normalized and filtered count data were variance-stabilized using the *voom* function in the *limma* package (v3.58.1)[112], and differential expression analysis was carried out using linear modeling in *limma*. Genes were considered differentially expressed at Padj < 0.01 and $|\log_2 FC| \geq 2$, with multiple testing correction applied using the Benjamini-Hochberg procedure[113]. Data visualization was performed using the *ggplot2* package (v3.5.1)[114]. The code used for the analysis is available in **Supplementary Code File 1**.

**References**


1. LoCascio, R. G., Desai, P., Sela, D. A., Weimer, B. & Mills, D. A. Broad conservation of milk utilization genes in Bifidobacterium longum subsp. infantis as revealed by comparative genomic hybridization. *Appl Environ Microbiol* **76**, 7373–7381 (2010).
2. Chaplin, A. V. *et al.* Intraspecies Genomic Diversity and Long-Term Persistence of Bifidobacterium longum. *PLoS One* **10**, (2015).
3. O'Callaghan, A., Bottacini, F., O'Connell Motherway, M. & van Sinderen, D. Pangenome analysis of Bifidobacterium longum and site-directed mutagenesis through by-pass of restriction-modification systems. *BMC Genomics* **16**, (2015).
4. Albert, K., Rani, A. & Sela, D. A. Comparative Pangenomics of the Mammalian Gut Commensal Bifidobacterium longum. *Microorganisms* **8**, (2019).
5. Tarracchini, C. *et al.* Phylogenomic disentangling of the Bifidobacterium longum subsp. infantis taxon. *Microb Genom* **7**, (2021).
6. Mattarelli, P., Bonaparte, C., Pot, B. & Biavati, B. Proposal to reclassify the three biotypes of Bifidobacterium longum as three subspecies: Bifidobacterium longum subsp. longum subsp. nov., Bifidobacterium longum subsp. infantis comb. nov. and Bifidobacterium longum subsp. suis comb. nov. *Int J Syst Evol Microbiol* **58**, 767–772 (2008).
7. Yanokura, E. *et al.* Subspeciation of Bifidobacterium longum by multilocus approaches and amplified fragment length polymorphism: Description of B. longum subsp. suillum subsp. nov., isolated from the faeces of piglets. *Syst Appl Microbiol* **38**, 305–314 (2015).
8. Bunesova, V., Lacroix, C. & Schwab, C. Fucosyllactose and L-fucose utilization of infant Bifidobacterium longum and Bifidobacterium kashiwanohense. *BMC Microbiol* **16**, (2016).
9. Arboleya, S. *et al.* Gene-trait matching across the Bifidobacterium longum pan-genome reveals considerable diversity in carbohydrate catabolism among human infant strains. *BMC Genomics* **19**, 33 (2018).
10. Barratt, M. J. *et al.* Bifidobacterium infantis treatment promotes weight gain in Bangladeshi infants with severe acute malnutrition. *Sci Transl Med* **14**, eabk1107 (2022).
11. Vatanen, T. *et al.* A distinct clade of Bifidobacterium longum in the gut of Bangladeshi children thrives during weaning. *Cell* (2022) doi:10.1016/j.cell.2022.10.011.
12. Modesto, M. *et al.* Bifidobacterium longum subsp. iuvenis subsp. nov., a novel subspecies isolated from the faeces of weaning infants. *Int J Syst Evol Microbiol* **73**, (2023).
13. Freitas, A. C. & Hill, J. E. Bifidobacteria isolated from vaginal and gut microbiomes are indistinguishable by comparative genomics. *PLoS One* **13**, e0196290 (2018).
14. Nouioui, I. *et al.* Genome-Based Taxonomic Classification of the Phylum Actinobacteria. *Front Microbiol* **9**, 2007 (2018).
15. Liu, J., Li, W., Yao, C., Yu, J. & Zhang, H. Comparative genomic analysis revealed genetic divergence between Bifidobacterium catenulatum subspecies present in infant versus adult guts. *BMC Microbiology* **22**, 158 (2022).
16. Schoch, C. L. *et al.* NCBI Taxonomy: a comprehensive update on curation, resources and tools. *Database (Oxford)* **2020**, baaa062 (2020).
17. Parks, D. H. *et al.* GTDB: an ongoing census of bacterial and archaeal diversity through a phylogenetically consistent, rank normalized and complete genome-based taxonomy. *Nucleic Acids Res* **50**, D785–D794 (2022).
18. Hitch, T. C. A. *et al.* HiBC: a publicly available collection of bacterial strains isolated from the human gut. *Nat Commun* **16**, 4203 (2025).

19. Vazquez-Gutierrez, P. *et al.* Complete and Assembled Genome Sequence of Bifidobacterium kashiwanohense PV20-2, Isolated from the Feces of an Anemic Kenyan Infant. *Genome Announc* **3**, e01467-14 (2015).
20. Rodionov, D. A. Comparative Genomic Reconstruction of Transcriptional Regulatory Networks in Bacteria. *Chem Rev* **107**, 3467–3497 (2007).
21. Gelfand, M. S. & Rodionov, D. A. Comparative genomics and functional annotation of bacterial transporters. *Physics of Life Reviews* **5**, 22–49 (2008).
22. Rodionov, D. A. *et al.* Transcriptional regulation of the carbohydrate utilization network in Thermotoga maritima. *Front Microbiol* **4**, (2013).
23. Khoroshkin, M. S., Leyn, S. A., Van Sinderen, D. & Rodionov, D. A. Transcriptional Regulation of Carbohydrate Utilization Pathways in the Bifidobacterium Genus. *Front Microbiol* **7**, (2016).
24. Arzamasov, A. A., van Sinderen, D. & Rodionov, D. A. Comparative Genomics Reveals the Regulatory Complexity of Bifidobacterial Arabinose and Arabino-Oligosaccharide Utilization. *Front Microbiol* **9**, 776 (2018).
25. Rodionov, D. A. *et al.* Transcriptional Regulation of Plant Biomass Degradation and Carbohydrate Utilization Genes in the Extreme Thermophile Caldicellulosiruptor bescii. *mSystems* e0134520 (2021) doi:10.1128/mSystems.01345-20.
26. Arzamasov, A. A. *et al.* Human Milk Oligosaccharide Utilization in Intestinal Bifidobacteria Is Governed by Global Transcriptional Regulator NagR. *mSystems* e0034322 (2022) doi:10.1128/msystems.00343-22.
27. Ejby, M. *et al.* Two binding proteins of the ABC transporter that confers growth of Bifidobacterium animalis subsp. lactis ATCC27673 on β-mannan possess distinct manno-oligosaccharide-binding profiles. *Mol. Microbiol.* **112**, 114–130 (2019).
28. Kulcinskaja, E., Rosengren, A., Ibrahim, R., Kolenová, K. & Stålbrand, H. Expression and characterization of a Bifidobacterium adolescentis beta-mannanase carrying mannan-binding and cell association motifs. *Appl. Environ. Microbiol.* **79**, 133–140 (2013).
29. Cordeiro, R. L. *et al.* Mechanism of high-mannose N-glycan breakdown and metabolism by Bifidobacterium longum. *Nat Chem Biol* (2022) doi:10.1038/s41589-022-01202-4.
30. Youn, S. Y., Park, M. S. & Ji, G. E. Identification of the beta-glucosidase gene from Bifidobacterium animalis subsp. lactis and its expression in B. bifidum BGN4. *J Microbiol Biotechnol* **22**, 1714–1723 (2012).
31. Saito, Y. *et al.* Multiple Transporters and Glycoside Hydrolases Are Involved in Arabinoxylan-Derived Oligosaccharide Utilization in Bifidobacterium pseudocatenulatum. *Appl Environ Microbiol* **86**, (2020).
32. Watanabe, Y. *et al.* Xylan utilisation promotes adaptation of Bifidobacterium pseudocatenulatum to the human gastrointestinal tract. *ISME COMMUN.* **1**, 1–11 (2021).
33. Orihara, K. *et al.* Characterization of Bifidobacterium kashiwanohense that utilizes both milk- and plant-derived oligosaccharides. *Gut Microbes* **15**, 2207455 (2023).
34. Novichkov, P. S. *et al.* RegPrecise: a database of curated genomic inferences of transcriptional regulatory interactions in prokaryotes. *Nucleic Acids Res* **38**, D111–D118 (2010).
35. Fujita, K., Sakaguchi, T., Sakamoto, A., Shimokawa, M. & Kitahara, K. Bifidobacterium longum subsp. longum Exo-β-1,3-Galactanase, an enzyme for the degradation of type II arabinogalactan. *Appl. Environ. Microbiol.* **80**, 4577–4584 (2014).
36. Fujita, K. *et al.* Degradative enzymes for type II arabinogalactan side chains in Bifidobacterium longum subsp. longum. *Appl. Microbiol. Biotechnol.* **103**, 1299–1310 (2019).

37. Sotoya, H. *et al.* Identification of genes involved in galactooligosaccharide utilization in Bifidobacterium breve strain YIT 4014T. *Microbiology (Reading, Engl.)* **163**, 1420–1428 (2017).
38. Theilmann, M. C., Fredslund, F., Svensson, B., Lo Leggio, L. & Abou Hachem, M. Substrate preference of an ABC importer corresponds to selective growth on β-(1,6)-galactosides in Bifidobacterium animalis subsp. lactis. *J Biol Chem* **294**, 11701–11711 (2019).
39. Viborg, A. H. *et al.* A β1-6/β1-3 galactosidase from Bifidobacterium animalis subsp. lactis Bl-04 gives insight into sub-specificities of β-galactoside catabolism within Bifidobacterium. *Mol. Microbiol.* (2014) doi:10.1111/mmi.12815.
40. Ambrogi, V. *et al.* Characterization of GH2 and GH42 β-galactosidases derived from bifidobacterial infant isolates. *AMB Express* **9**, (2019).
41. Margolles, A. & de los Reyes-Gavilán, C. G. Purification and Functional Characterization of a Novel α-l-Arabinofuranosidase from Bifidobacterium longum B667. *Appl Environ Microbiol* **69**, 5096–5103 (2003).
42. Lanigan, N. *et al.* Transcriptional control of central carbon metabolic flux in Bifidobacteria by two functionally similar, yet distinct LacI-type regulators. *Sci Rep* **9**, 17851 (2019).
43. Ishiwata, A. *et al.* Bifidobacterial GH family 146 β-L-arabinofuranosidase (Bll4HypBA1) as the last enzyme for the complete removal of oligoarabinofuranosides from HRGP. *Chembiochem* (2022) doi:10.1002/cbic.202200637.
44. Fujita, K. *et al.* Bifidobacterial GH146 β-L-arabinofuranosidase for the removal of β1,3-L-arabinofuranosides on plant glycans. *Appl Microbiol Biotechnol* **108**, 199 (2024).
45. Komeno, M., Hayamizu, H., Fujita, K. & Ashida, H. Two Novel α-l-Arabinofuranosidases from Bifidobacterium longum subsp. longum Belonging to Glycoside Hydrolase Family 43 Cooperatively Degrade Arabinan. *Appl. Environ. Microbiol.* (2019) doi:10.1128/AEM.02582-18.
46. Wichelecki, D. J. *et al.* Discovery of function in the enolase superfamily: D-mannonate and d-gluconate dehydratases in the D-mannonate dehydratase subgroup. *Biochemistry* **53**, 2722–2731 (2014).
47. Sakurama, H. *et al.* β-Glucuronidase from Lactobacillus brevis useful for baicalin hydrolysis belongs to glycoside hydrolase family 30. *Appl Microbiol Biotechnol* **98**, 4021–4032 (2014).
48. Zhu, L. *et al.* Captive Common Marmosets (Callithrix jacchus) Are Colonized throughout Their Lives by a Community of Bifidobacterium Species with Species-Specific Genomic Content That Can Support Adaptation to Distinct Metabolic Niches. *mBio* e0115321 (2021) doi:10.1128/mBio.01153-21.
49. Crociani, F., Alessandrini, A., Mucci, M. M. & Biavati, B. Degradation of complex carbohydrates by Bifidobacterium spp. *Int J Food Microbiol* **24**, 199–210 (1994).
50. Ventura, M. *et al.* The Bifidobacterium dentium Bd1 genome sequence reflects its genetic adaptation to the human oral cavity. *PLoS Genet* **5**, e1000785 (2009).
51. Lee, J.-H. & O'Sullivan, D. J. Genomic insights into bifidobacteria. *Microbiol. Mol. Biol. Rev.* **74**, 378–416 (2010).
52. Bottacini, F. *et al.* Comparative genomics and genotype-phenotype associations in Bifidobacterium breve. *Sci Rep* **8**, (2018).
53. Caescu, C. I., Vidal, O., Krzewinski, F., Artenie, V. & Bouquelet, S. Bifidobacterium longum requires a fructokinase (Frk; ATP:D-fructose 6-phosphotransferase, EC 2.7.1.4) for fructose catabolism. *J. Bacteriol.* **186**, 6515–6525 (2004).

54. Fujita, Y., Shindo, K., Miwa, Y. & Yoshida, K. Bacillus subtilis inositol dehydrogenase-encoding gene (idh): sequence and expression in Escherichia coli. *Gene* **108**, 121–125 (1991).
55. Rodionova, I. A. *et al.* Novel inositol catabolic pathway in Thermotoga maritima. *Environ Microbiol* **15**, 2254–2266 (2013).
56. Yoshida, K.-I. *et al.* The fifth gene of the iol operon of Bacillus subtilis, iolE, encodes 2-keto-myo-inositol dehydratase. *Microbiology (Reading)* **150**, 571–580 (2004).
57. Yoshida, K. *et al.* myo-Inositol catabolism in Bacillus subtilis. *J Biol Chem* **283**, 10415–10424 (2008).
58. Fujita, Y., Fujita, T., Miwa, Y., Nihashi, J. & Aratani, Y. Organization and transcription of the gluconate operon, gnt, of Bacillus subtilis. *J Biol Chem* **261**, 13744–13753 (1986).
59. Rauch, B., Pahlke, J., Schweiger, P. & Deppenmeier, U. Characterization of enzymes involved in the central metabolism of Gluconobacter oxydans. *Appl Microbiol Biotechnol* **88**, 711–718 (2010).
60. Singh, B. *et al.* Molecular and Functional Insights into the Regulation of d-Galactonate Metabolism by the Transcriptional Regulator DgoR in Escherichia coli. *J Bacteriol* **201**, e00281-18 (2019).
61. Deacon, J. & Cooper, R. A. D-Galactonate utilisation by enteric bacteria. The catabolic pathway in Escherichia coli. *FEBS Lett* **77**, 201–205 (1977).
62. Ibañez, E. *et al.* Role of the yiaR and yiaS genes of Escherichia coli in metabolism of endogenously formed L-xylulose. *J Bacteriol* **182**, 4625–4627 (2000).
63. Yew, W. S. & Gerlt, J. A. Utilization of L-ascorbate by Escherichia coli K-12: assignments of functions to products of the yjf-sga and yia-sgb operons. *J Bacteriol* **184**, 302–306 (2002).
64. Zhang, Z., Aboulwafa, M., Smith, M. H. & Saier, M. H. The ascorbate transporter of Escherichia coli. *J Bacteriol* **185**, 2243–2250 (2003).
65. Van Rossum, T., Ferretti, P., Maistrenko, O. M. & Bork, P. Diversity within species: interpreting strains in microbiomes. *Nat Rev Microbiol* **18**, 491–506 (2020).
66. You, X., Rani, A., Özcan, E., Lyu, Y. & Sela, D. A. Bifidobacterium longum subsp. infantis utilizes human milk urea to recycle nitrogen within the infant gut microbiome. *Gut Microbes* **15**, 2192546 (2023).
67. Kitaoka, M., Tian, J. & Nishimoto, M. Novel Putative Galactose Operon Involving Lacto-N-Biose Phosphorylase in Bifidobacterium longum. *Appl Environ Microbiol* **71**, 3158–3162 (2005).
68. Nishimoto, M. & Kitaoka, M. Identification of N-Acetylhexosamine 1-Kinase in the Complete Lacto-N-Biose I/Galacto-N-Biose Metabolic Pathway in Bifidobacterium longum. *Appl Environ Microbiol* **73**, 6444–6449 (2007).
69. Arzamasov, A. A. & Osterman, A. L. Milk glycan metabolism by intestinal bifidobacteria: insights from comparative genomics. *Crit Rev Biochem Mol Biol* 1–23 (2023) doi:10.1080/10409238.2023.2182272.
70. Fujita, K. *et al.* Identification and molecular cloning of a novel glycoside hydrolase family of core 1 type O-glycan-specific endo-alpha-N-acetylgalactosaminidase from Bifidobacterium longum. *J. Biol. Chem.* **280**, 37415–37422 (2005).
71. Sakanaka, M. *et al.* Varied Pathways of Infant Gut-Associated Bifidobacterium to Assimilate Human Milk Oligosaccharides: Prevalence of the Gene Set and Its Correlation with Bifidobacteria-Rich Microbiota Formation. *Nutrients* **12**, (2019).
72. O'Connell Motherway, M. *et al.* Characterization of ApuB, an extracellular type II amylopullulanase from Bifidobacterium breve UCC2003. *Appl. Environ. Microbiol.* **74**, 6271–6279 (2008).

73. Kim, S.-Y. *et al.* Enzymatic analysis of truncation mutants of a type II pullulanase from Bifidobacterium adolescentis P2P3, a resistant starch-degrading gut bacterium. *Int J Biol Macromol* **193**, 1340–1349 (2021).
74. Kashima, T. *et al.* Identification of difructose dianhydride I synthase/hydrolase from an oral bacterium establishes a novel glycoside hydrolase family. *J Biol Chem* **297**, 101324 (2021).
75. Sela, D. A. *et al.* The genome sequence of Bifidobacterium longum subsp. infantis reveals adaptations for milk utilization within the infant microbiome. *Proc Natl Acad Sci U S A* **105**, 18964–18969 (2008).
76. Matsuki, T. *et al.* A key genetic factor for fucosyllactose utilization affects infant gut microbiota development. *Nat Commun* **7**, 11939 (2016).
77. Garrido, D. *et al.* A novel gene cluster allows preferential utilization of fucosylated milk oligosaccharides in Bifidobacterium longum subsp. longum SC596. *Sci Rep* **6**, (2016).
78. James, K. *et al.* Metabolism of the predominant human milk oligosaccharide fucosyllactose by an infant gut commensal. *Sci Rep* **9**, 15427 (2019).
79. Ojima, M. N. *et al.* Diversification of a fucosyllactose transporter within the genus Bifidobacterium. *Appl Environ Microbiol* AEM0143721 (2021) doi:10.1128/AEM.01437-21.
80. Sanchez-Gallardo, R. *et al.* Selective human milk oligosaccharide utilization by members of the Bifidobacterium pseudocatenulatum taxon. *Appl Environ Microbiol* e0064824 (2024) doi:10.1128/aem.00648-24.
81. Sela, D. A. *et al.* An infant-associated bacterial commensal utilizes breast milk sialyloligosaccharides. *J. Biol. Chem.* **286**, 11909–11918 (2011).
82. Garrido, D., Kim, J. H., German, J. B., Raybould, H. E. & Mills, D. A. Oligosaccharide Binding Proteins from Bifidobacterium longum subsp. infantis Reveal a Preference for Host Glycans. *PLoS One* **6**, (2011).
83. Komeno, M. *et al.* Two α-L-arabinofuranosidases from Bifidobacterium longum subsp. longum are involved in arabinoxylan utilization. *Appl Microbiol Biotechnol* **106**, 1957–1965 (2022).
84. Friess, L. *et al.* Two extracellular α-arabinofuranosidases are required for cereal-derived arabinoxylan metabolism by Bifidobacterium longum subsp. longum. *Gut Microbes* **16**, 2353229 (2024).
85. Gehrig, J. L. *et al.* Effects of microbiota-directed foods in gnotobiotic animals and undernourished children. *Science* **365**, (2019).
86. Raman, A. S. *et al.* A sparse covarying unit that describes healthy and impaired human gut microbiota development. *Science* **365**, (2019).
87. Smith, M. I. *et al.* Gut microbiomes of Malawian twin pairs discordant for kwashiorkor. *Science* **339**, 548–554 (2013).
88. Krueger, F. *et al.* TrimGalore. Zenodo https://doi.org/10.5281/zenodo.5127898 (2019).
89. Bushnell, B. BBTools software package. (2014).
90. De Coster, W., D'Hert, S., Schultz, D. T., Cruts, M. & Van Broeckhoven, C. NanoPack: visualizing and processing long-read sequencing data. *Bioinformatics* **34**, 2666–2669 (2018).
91. Bankevich, A. *et al.* SPAdes: A New Genome Assembly Algorithm and Its Applications to Single-Cell Sequencing. *J Comput Biol* **19**, 455–477 (2012).
92. Wick, R. R., Judd, L. M., Gorrie, C. L. & Holt, K. E. Unicycler: Resolving bacterial genome assemblies from short and long sequencing reads. *PLoS Comput Biol* **13**, e1005595 (2017).
93. Vaser, R., Sović, I., Nagarajan, N. & Šikić, M. Fast and accurate de novo genome assembly from long uncorrected reads. *Genome Res* **27**, 737–746 (2017).

94. Wick, R. R. *et al.* Trycycler: consensus long-read assemblies for bacterial genomes. *Genome Biol* **22**, 266 (2021).
95. Kolmogorov, M., Yuan, J., Lin, Y. & Pevzner, P. A. Assembly of long, error-prone reads using repeat graphs. *Nat Biotechnol* **37**, 540–546 (2019).
96. Li, H. Minimap and miniasm: fast mapping and de novo assembly for noisy long sequences. *Bioinformatics* **32**, 2103–2110 (2016).
97. Koren, S. *et al.* Canu: scalable and accurate long-read assembly via adaptive k-mer weighting and repeat separation. *Genome Res* **27**, 722–736 (2017).
98. Vaser, R. & Šikić, M. Time- and memory-efficient genome assembly with Raven. *Nat Comput Sci* **1**, 332–336 (2021).
99. Wick, R. R. & Holt, K. E. Polypolish: Short-read polishing of long-read bacterial genome assemblies. *PLoS Comput Biol* **18**, e1009802 (2022).
100. Gurevich, A., Saveliev, V., Vyahhi, N. & Tesler, G. QUAST: quality assessment tool for genome assemblies. *Bioinformatics* **29**, 1072–1075 (2013).
101. Seemann, T. Prokka: rapid prokaryotic genome annotation. *Bioinformatics* **30**, 2068–2069 (2014).
102. Di Tommaso, P. *et al.* T-Coffee: a web server for the multiple sequence alignment of protein and RNA sequences using structural information and homology extension. *Nucleic Acids Res* **39**, W13–W17 (2011).
103. Mironov, A. A., Vinokurova, N. P. & Gelfand, M. S. Software for analysis of bacterial genomes. *Mol Biol* **34**, 222–231 (2000).
104. Crooks, G. E., Hon, G., Chandonia, J.-M. & Brenner, S. E. WebLogo: A Sequence Logo Generator. *Genome Res* **14**, 1188–1190 (2004).
105. Andrews, S. FastQC: A Quality Control Tool for High Throughput Sequence Data. http://www.bioinformatics.babraham.ac.uk/projects/fastqc/ (2010).
106. Martin, M. Cutadapt removes adapter sequences from high-throughput sequencing reads. *EMBnet.journal* **17**, 10–12 (2011).
107. Langmead, B. & Salzberg, S. L. Fast gapped-read alignment with Bowtie 2. *Nat Methods* **9**, 357–359 (2012).
108. Bray, N. L., Pimentel, H., Melsted, P. & Pachter, L. Near-optimal probabilistic RNA-seq quantification. *Nat Biotechnol* **34**, 525–527 (2016).
109. Huber, W. *et al.* Orchestrating high-throughput genomic analysis with Bioconductor. *Nat Methods* **12**, 115–121 (2015).
110. Soneson, C., Love, M. I. & Robinson, M. D. Differential analyses for RNA-seq: transcript-level estimates improve gene-level inferences. *F1000Res* **4**, 1521 (2015).
111. Robinson, M. D., McCarthy, D. J. & Smyth, G. K. edgeR: a Bioconductor package for differential expression analysis of digital gene expression data. *Bioinformatics* **26**, 139–140 (2010).
112. Ritchie, M. E. *et al.* limma powers differential expression analyses for RNA-sequencing and microarray studies. *Nucleic Acids Res* **43**, e47 (2015).
113. Benjamini, Y. & Hochberg, Y. Controlling the False Discovery Rate: A Practical and Powerful Approach to Multiple Testing. *Journal of the Royal Statistical Society. Series B (Methodological)* **57**, 289–300 (1995).
114. Wickham, Hadley. *Ggplot2: Elegant Graphics for Data Analysis*. (Springer-Verlag, New York, 2016).

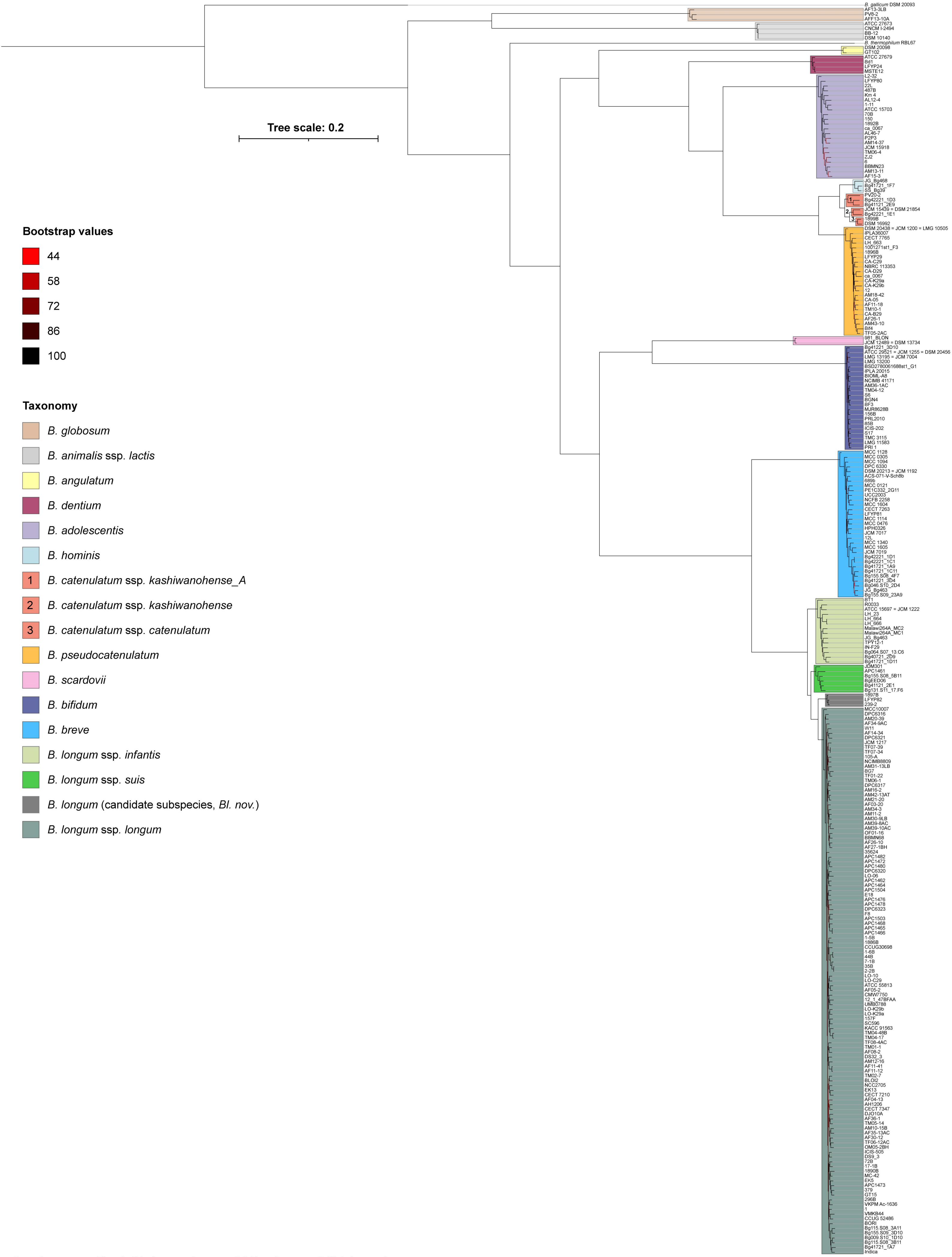


**Supplementary Fig. 1. Phylogenetic tree of 263 reference *Bifidobacterium* genomes.**
The maximum-likelihood tree was constructed from an alignment of concatenated nucleotide sequences of 487 core genes. Node bootstrap support values are indicated by color.

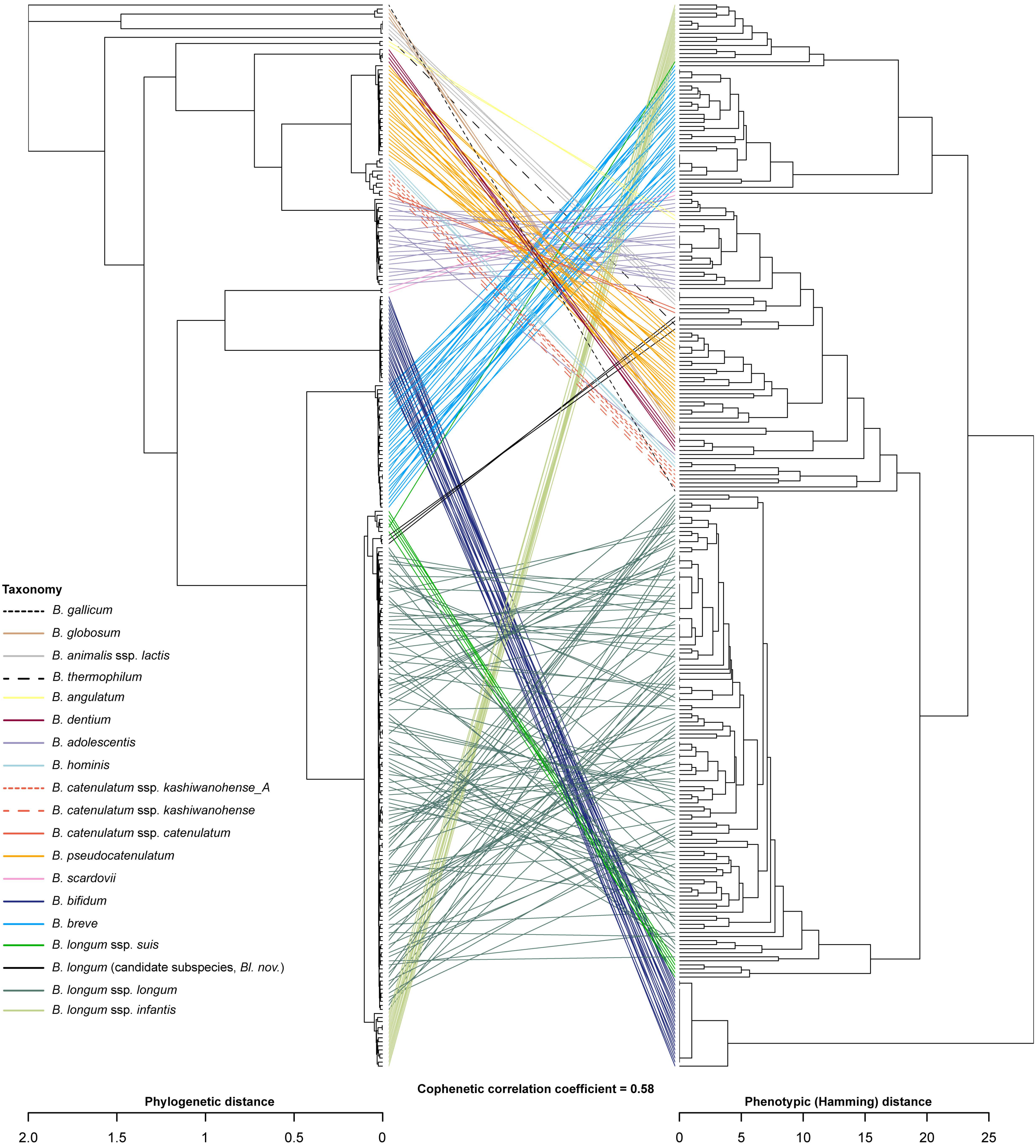


**Supplementary Fig. 2. Concordance between phylogeny and predicted phenotypes.**
The tanglegram compares the phylogenetic tree of 263 reference *Bifidobacterium* genomes with the Hamming-distance dendrogram derived from the Binary Phenotype Matrix (BPM). Lines connecting matching genomes are colored by taxonomic assignment.

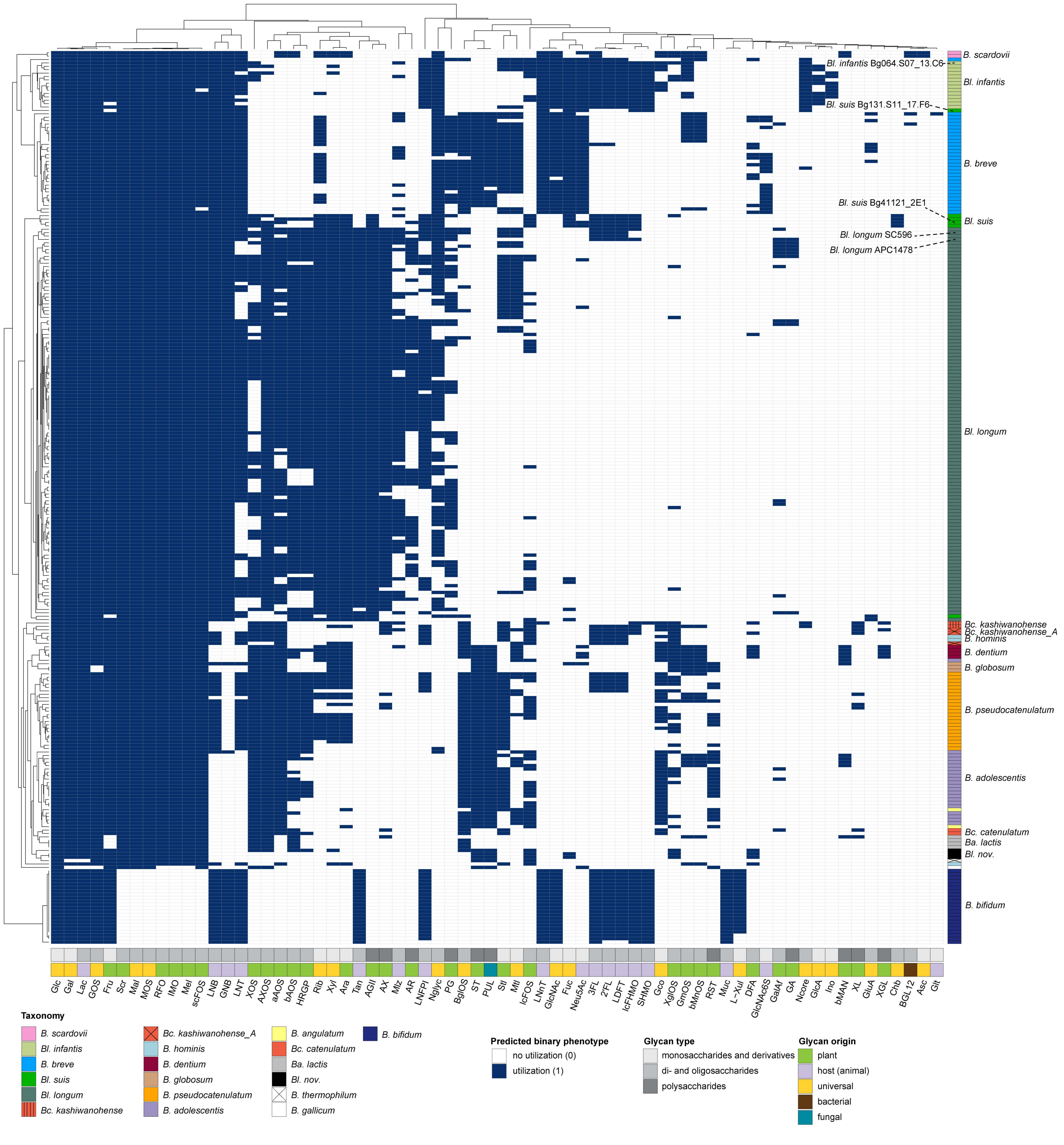


**Supplementary Fig. 3. Variability in predicted carbohydrate utilization phenotypes across 263 reference *Bifidobacterium* strains.**
The heatmap shows the hierarchical clustering (Hamming distance and average linkage) of the Binary Phenotype Matrix (BPM) for 68 carbohydrate utilization pathways (columns) reconstructed in 263 reference *Bifidobacterium* genomes (rows). Full names of abbreviations are provided in Supplementary Table 5.
The annotation rows at the bottom represent pathway/phenotype classifications. The annotation column on the right denotes taxonomic groupings.
Lines point to specific strains discussed in the text.

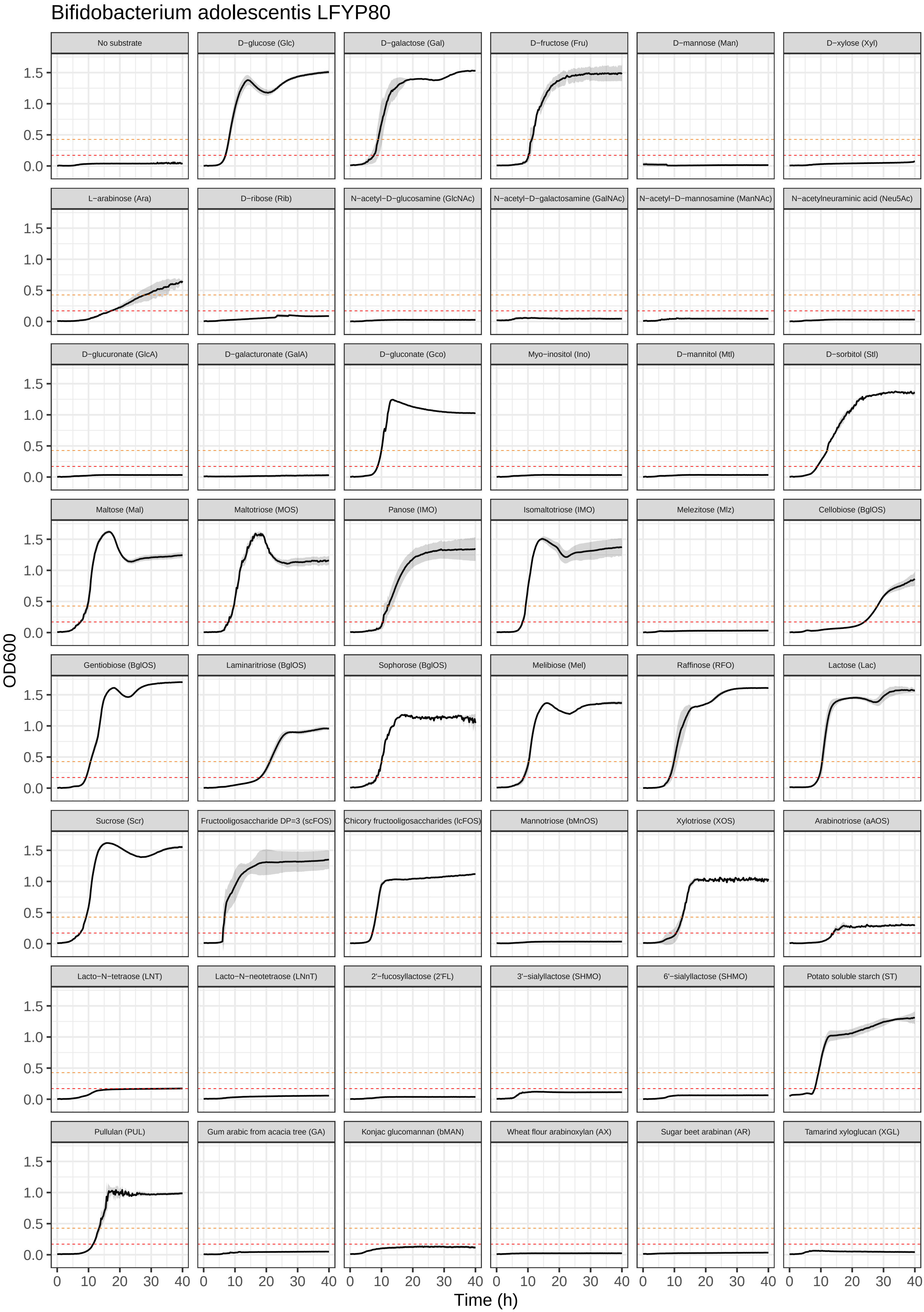
Bifidobacterium adolescentis LFYP80
No substrate
D−glucose (Glc)
D−galactose (Gal)
D−fructose (Fru)
D−mannose (Man)
D−xylose (Xyl)
L−arabinose (Ara)
D−ribose (Rib)
N−acetyl−D−glucosamine (GlcNAc)
N−acetyl−D−galactosamine (GalNAc)
N−acetyl−D−mannosamine (ManNAc)
N−acetylneuraminic acid (Neu5Ac)
D−glucuronate (GlcA)
D−galacturonate (GalA)
D−gluconate (Gco)
Myo−inositol (Ino)
D−mannitol (Mtl)
D−sorbitol (Stl)
Maltose (Mal)
Maltotriose (MOS)
Panose (IMO)
Isomaltotriose (IMO)
Melezitose (Mlz)
Cellobiose (BglOS)
Gentiobiose (BglOS)
Laminaritriose (BglOS)
Sophorose (BglOS)
Melibiose (Mel)
Raffinose (RFO)
Lactose (Lac)
Sucrose (Scr)
Fructooligosaccharide DP=3 (scFOS)
Chicory fructooligosaccharides (lcFOS)
Mannotriose (bMnOS)
Xylotriose (XOS)
Arabinotriose (aAOS)
Lacto−N−tetraose (LNT)
Lacto−N−neotetraose (LNnT)
2'−fucosyllactose (2'FL)
3'−sialyllactose (SHMO)
6'−sialyllactose (SHMO)
Potato soluble starch (ST)
Pullulan (PUL)
Gum arabic from acacia tree (GA)
Konjac glucomannan (bMAN)
Wheat flour arabinoxylan (AX)
Sugar beet arabinan (AR)
Tamarind xyloglucan (XGL)
OD600
0.0
0.5
1.0
1.5
0
10
20
30
40
Time (h)

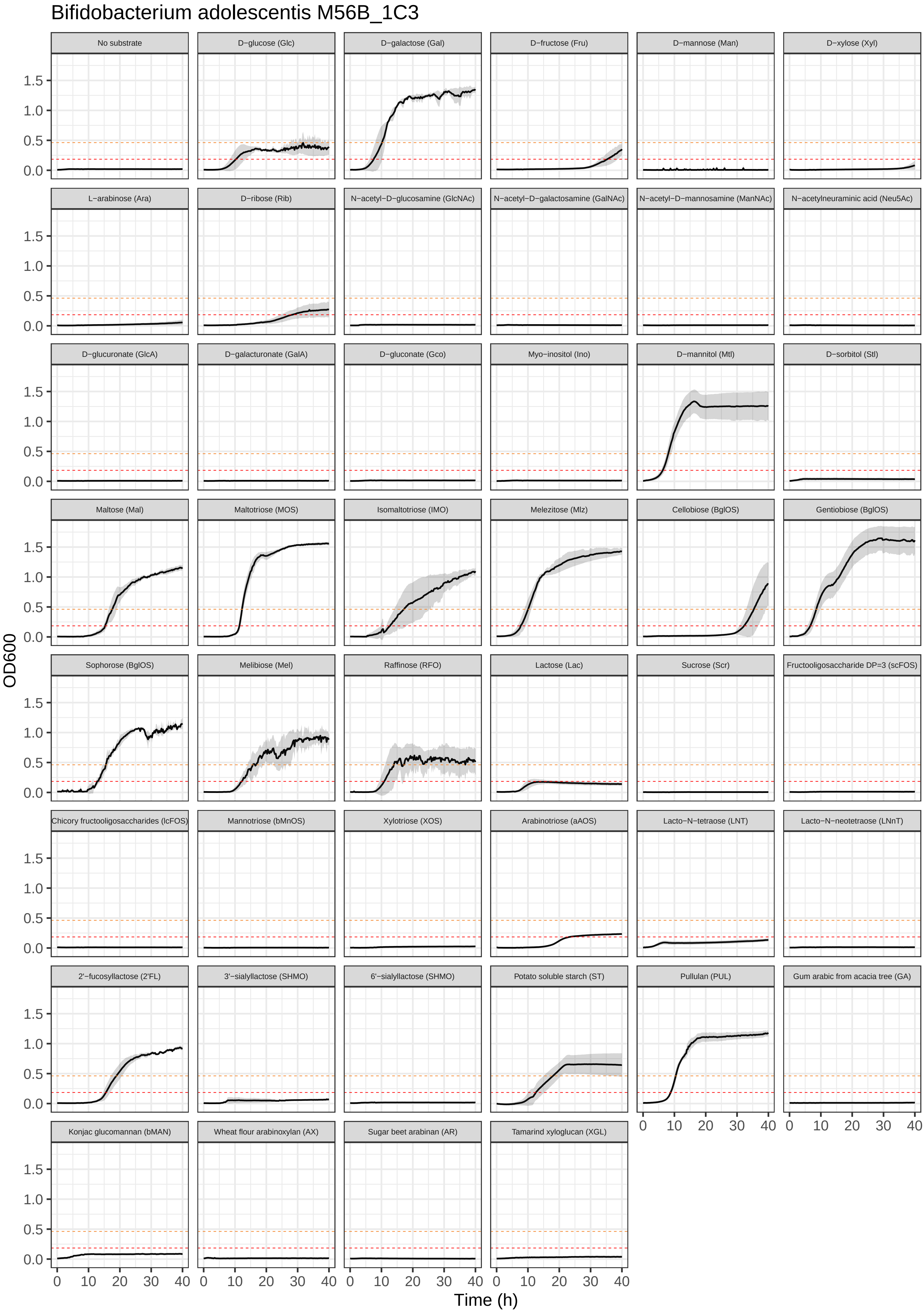

Bifidobacterium adolescentis M56B_1C3
No substrate
D−glucose (Glc)
D−galactose (Gal)
D−fructose (Fru)
D−mannose (Man)
D−xylose (Xyl)
L−arabinose (Ara)
D−ribose (Rib)
N−acetyl−D−glucosamine (GlcNAc)
N−acetyl−D−galactosamine (GalNAc)
N−acetyl−D−mannosamine (ManNAc)
N−acetylneuraminic acid (Neu5Ac)
D−glucuronate (GlcA)
D−galacturonate (GalA)
D−gluconate (Gco)
Myo−inositol (Ino)
D−mannitol (Mtl)
D−sorbitol (Stl)
Maltose (Mal)
Maltotriose (MOS)
Isomaltotriose (IMO)
Melezitose (Mlz)
Cellobiose (BglOS)
Gentiobiose (BglOS)
Sophorose (BglOS)
Melibiose (Mel)
Raffinose (RFO)
Lactose (Lac)
Sucrose (Scr)
Fructooligosaccharide DP=3 (scFOS)
Chicory fructooligosaccharides (lcFOS)
Mannotriose (bMnOS)
Xylotriose (XOS)
Arabinotriose (aAOS)
Lacto−N−tetraose (LNT)
Lacto−N−neotetraose (LNnT)
2'−fucosyllactose (2'FL)
3'−sialyllactose (SHMO)
6'−sialyllactose (SHMO)
Potato soluble starch (ST)
Pullulan (PUL)
Gum arabic from acacia tree (GA)
Konjac glucomannan (bMAN)
Wheat flour arabinoxylan (AX)
Sugar beet arabinan (AR)
Tamarind xyloglucan (XGL)
OD600
0.0
0.5
1.0
1.5
0
10
20
30
40
Time (h)

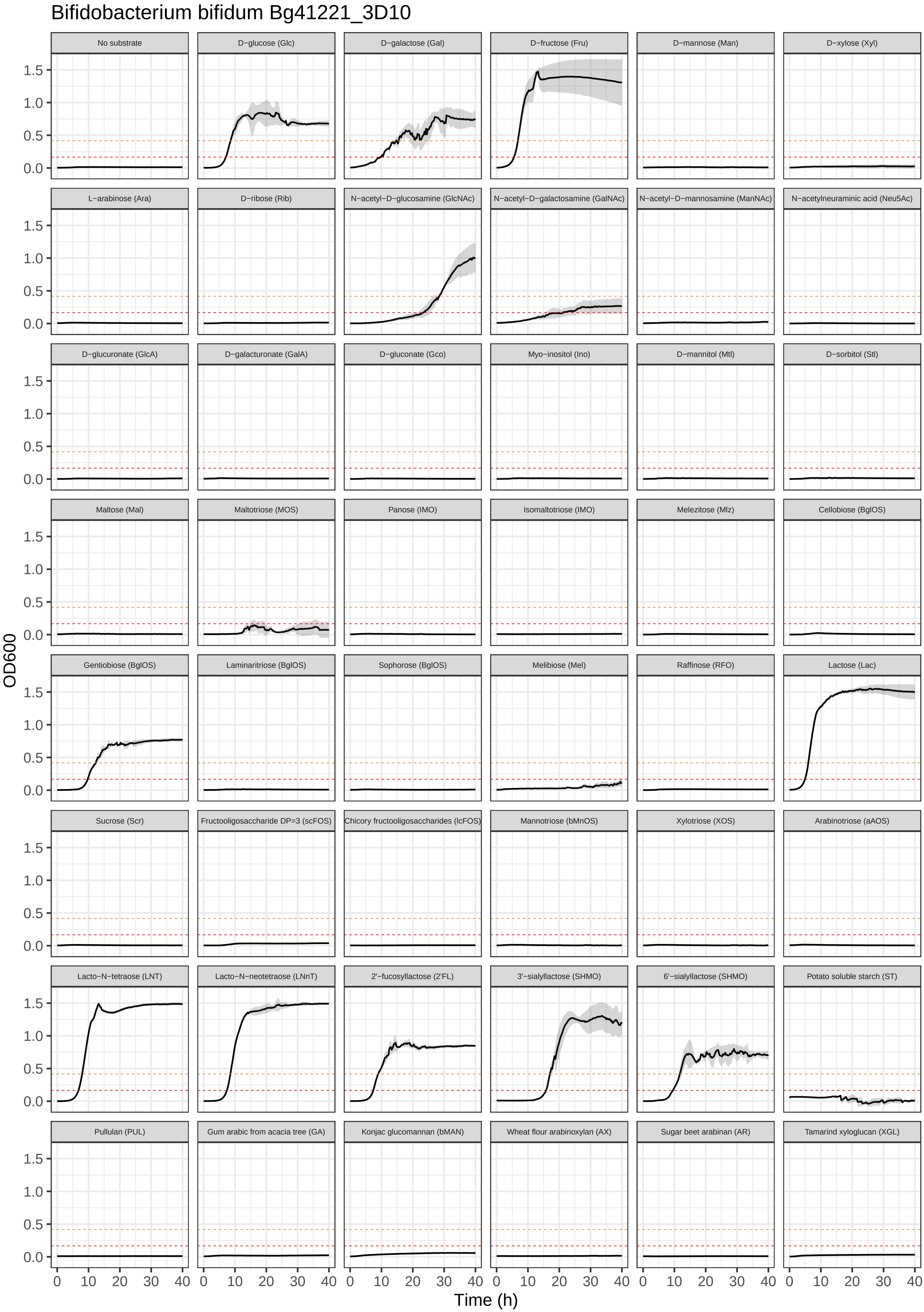

Bifidobacterium bifidum Bg41221_3D10
No substrate
D−glucose (Glc)
D−galactose (Gal)
D−fructose (Fru)
D−mannose (Man)
D−xylose (Xyl)
L−arabinose (Ara)
D−ribose (Rib)
N−acetyl−D−glucosamine (GlcNAc)
N−acetyl−D−galactosamine (GalNAc)
N−acetyl−D−mannosamine (ManNAc)
N−acetylneuraminic acid (Neu5Ac)
D−glucuronate (GlcA)
D−galacturonate (GalA)
D−gluconate (Gco)
Myo−inositol (Ino)
D−mannitol (Mtl)
D−sorbitol (Stl)
Maltose (Mal)
Maltotriose (MOS)
Panose (IMO)
Isomaltotriose (IMO)
Melezitose (Mlz)
Cellobiose (BglOS)
Gentiobiose (BglOS)
Laminaritriose (BglOS)
Sophorose (BglOS)
Melibiose (Mel)
Raffinose (RFO)
Lactose (Lac)
Sucrose (Scr)
Fructooligosaccharide DP=3 (scFOS)
Chicory fructooligosaccharides (lcFOS)
Mannotriose (bMnOS)
Xylotriose (XOS)
Arabinotriose (aAOS)
Lacto−N−tetraose (LNT)
Lacto−N−neotetraose (LNnT)
2'−fucosyllactose (2'FL)
3'−sialyllactose (SHMO)
6'−sialyllactose (SHMO)
Potato soluble starch (ST)
Pullulan (PUL)
Gum arabic from acacia tree (GA)
Konjac glucomannan (bMAN)
Wheat flour arabinoxylan (AX)
Sugar beet arabinan (AR)
Tamarind xyloglucan (XGL)
OD600
0.0
0.5
1.0
1.5
0
10
20
30
40
Time (h)

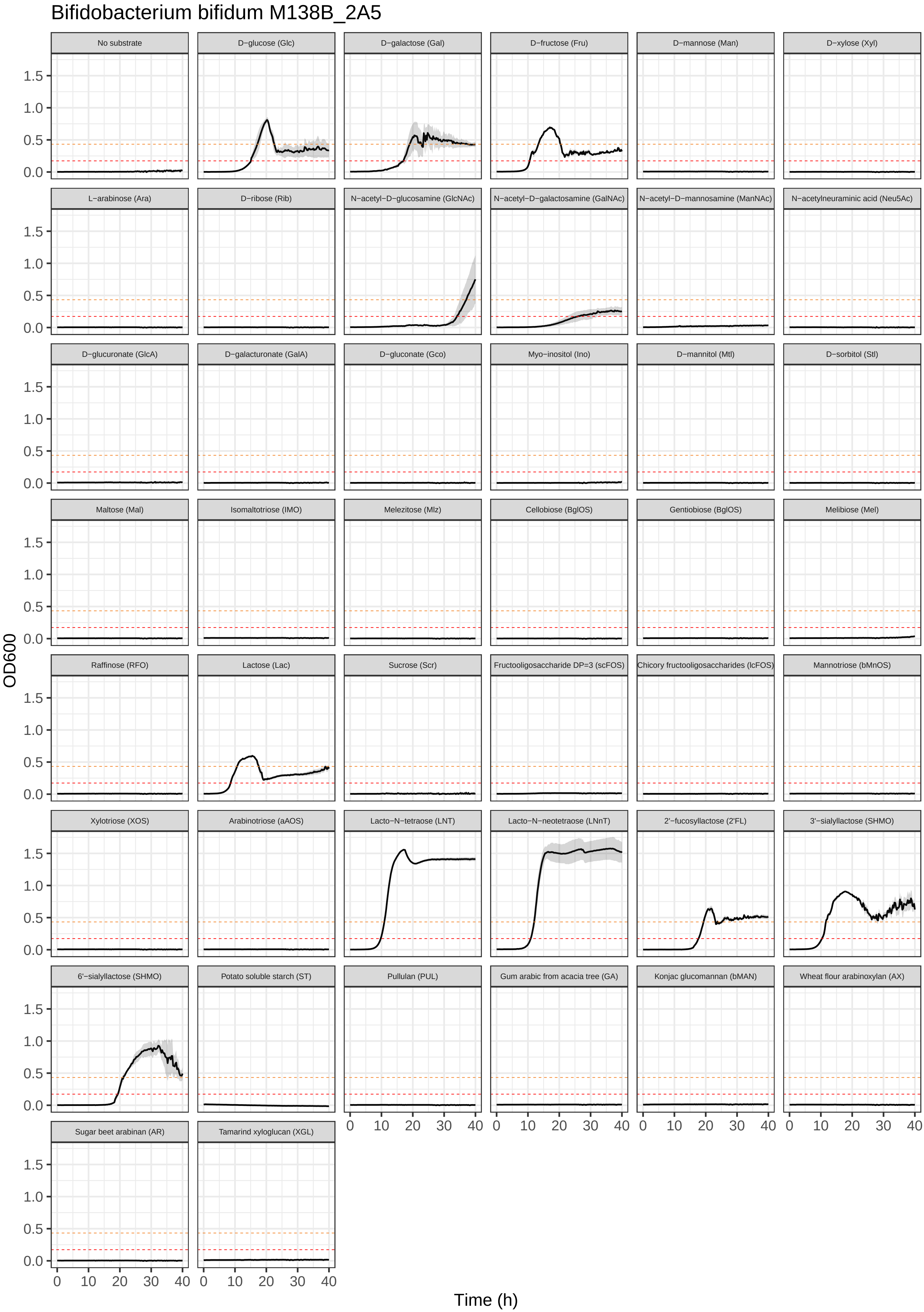
Bifidobacterium bifidum M138B_2A5
No substrate
D−glucose (Glc)
D−galactose (Gal)
D−fructose (Fru)
D−mannose (Man)
D−xylose (Xyl)
L−arabinose (Ara)
D−ribose (Rib)
N−acetyl−D−glucosamine (GlcNAc)
N−acetyl−D−galactosamine (GalNAc)
N−acetyl−D−mannosamine (ManNAc)
N−acetylneuraminic acid (Neu5Ac)
D−glucuronate (GlcA)
D−galacturonate (GalA)
D−gluconate (Gco)
Myo−inositol (Ino)
D−mannitol (Mtl)
D−sorbitol (Stl)
Maltose (Mal)
Isomaltotriose (IMO)
Melezitose (Mlz)
Cellobiose (BglOS)
Gentiobiose (BglOS)
Melibiose (Mel)
Raffinose (RFO)
Lactose (Lac)
Sucrose (Scr)
Fructooligosaccharide DP=3 (scFOS)
Chicory fructooligosaccharides (lcFOS)
Mannotriose (bMnOS)
Xylotriose (XOS)
Arabinotriose (aAOS)
Lacto−N−tetraose (LNT)
Lacto−N−neotetraose (LNnT)
2'−fucosyllactose (2'FL)
3'−sialyllactose (SHMO)
6'−sialyllactose (SHMO)
Potato soluble starch (ST)
Pullulan (PUL)
Gum arabic from acacia tree (GA)
Konjac glucomannan (bMAN)
Wheat flour arabinoxylan (AX)
Sugar beet arabinan (AR)
Tamarind xyloglucan (XGL)
OD600
0.0
0.5
1.0
1.5
0
10
20
30
40
Time (h)

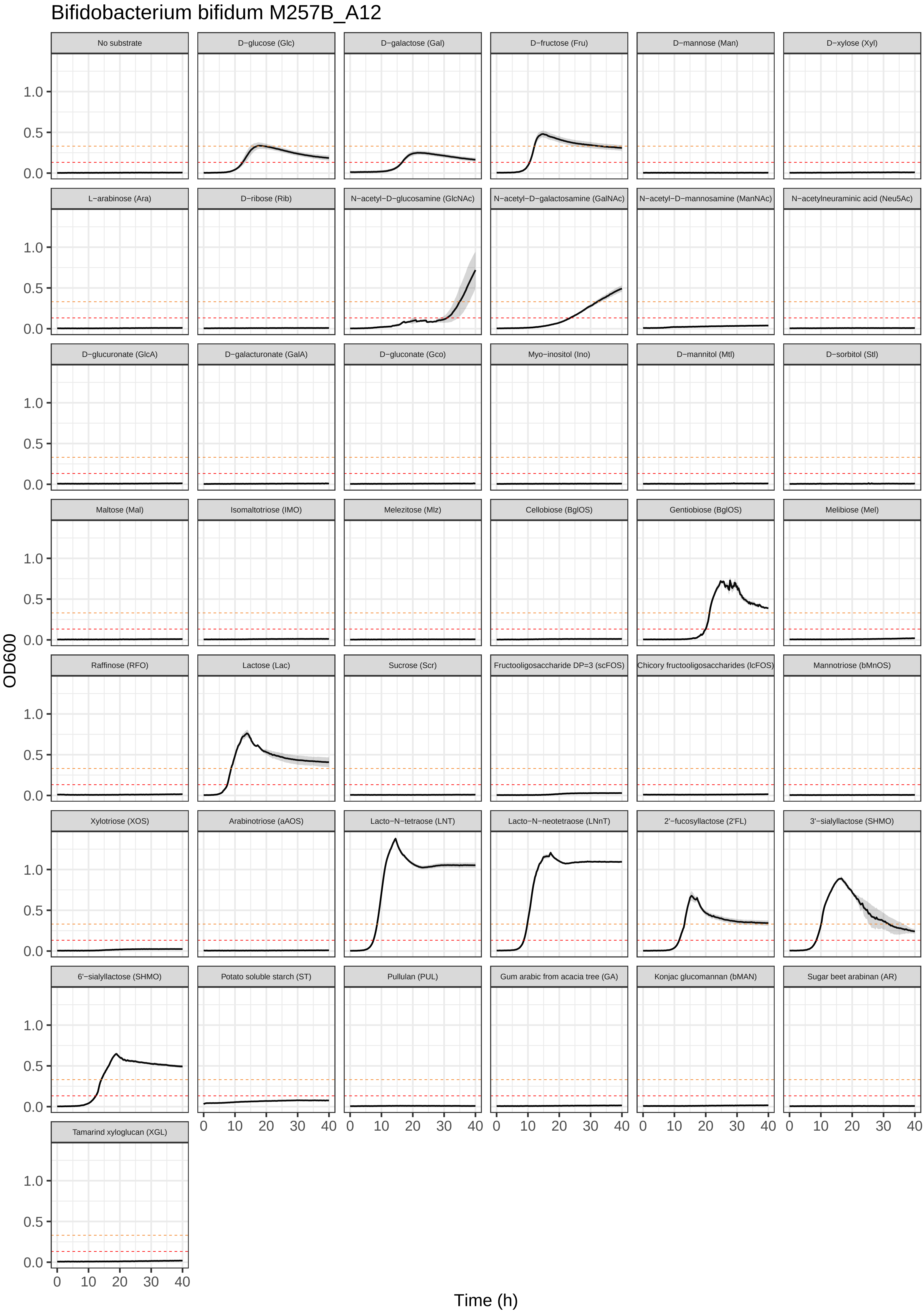

Bifidobacterium bifidum M257B_A12
No substrate
D−glucose (Glc)
D−galactose (Gal)
D−fructose (Fru)
D−mannose (Man)
D−xylose (Xyl)
L−arabinose (Ara)
D−ribose (Rib)
N−acetyl−D−glucosamine (GlcNAc)
N−acetyl−D−galactosamine (GalNAc)
N−acetyl−D−mannosamine (ManNAc)
N−acetylneuraminic acid (Neu5Ac)
D−glucuronate (GlcA)
D−galacturonate (GalA)
D−gluconate (Gco)
Myo−inositol (Ino)
D−mannitol (Mtl)
D−sorbitol (Stl)
Maltose (Mal)
Isomaltotriose (IMO)
Melezitose (Mlz)
Cellobiose (BglOS)
Gentiobiose (BglOS)
Melibiose (Mel)
Raffinose (RFO)
Lactose (Lac)
Sucrose (Scr)
Fructooligosaccharide DP=3 (scFOS)
Chicory fructooligosaccharides (lcFOS)
Mannotriose (bMnOS)
Xylotriose (XOS)
Arabinotriose (aAOS)
Lacto−N−tetraose (LNT)
Lacto−N−neotetraose (LNnT)
2'−fucosyllactose (2'FL)
3'−sialyllactose (SHMO)
6'−sialyllactose (SHMO)
Potato soluble starch (ST)
Pullulan (PUL)
Gum arabic from acacia tree (GA)
Konjac glucomannan (bMAN)
Sugar beet arabinan (AR)
Tamarind xyloglucan (XGL)
OD600
0.0
0.5
1.0
0
10
20
30
40
Time (h)

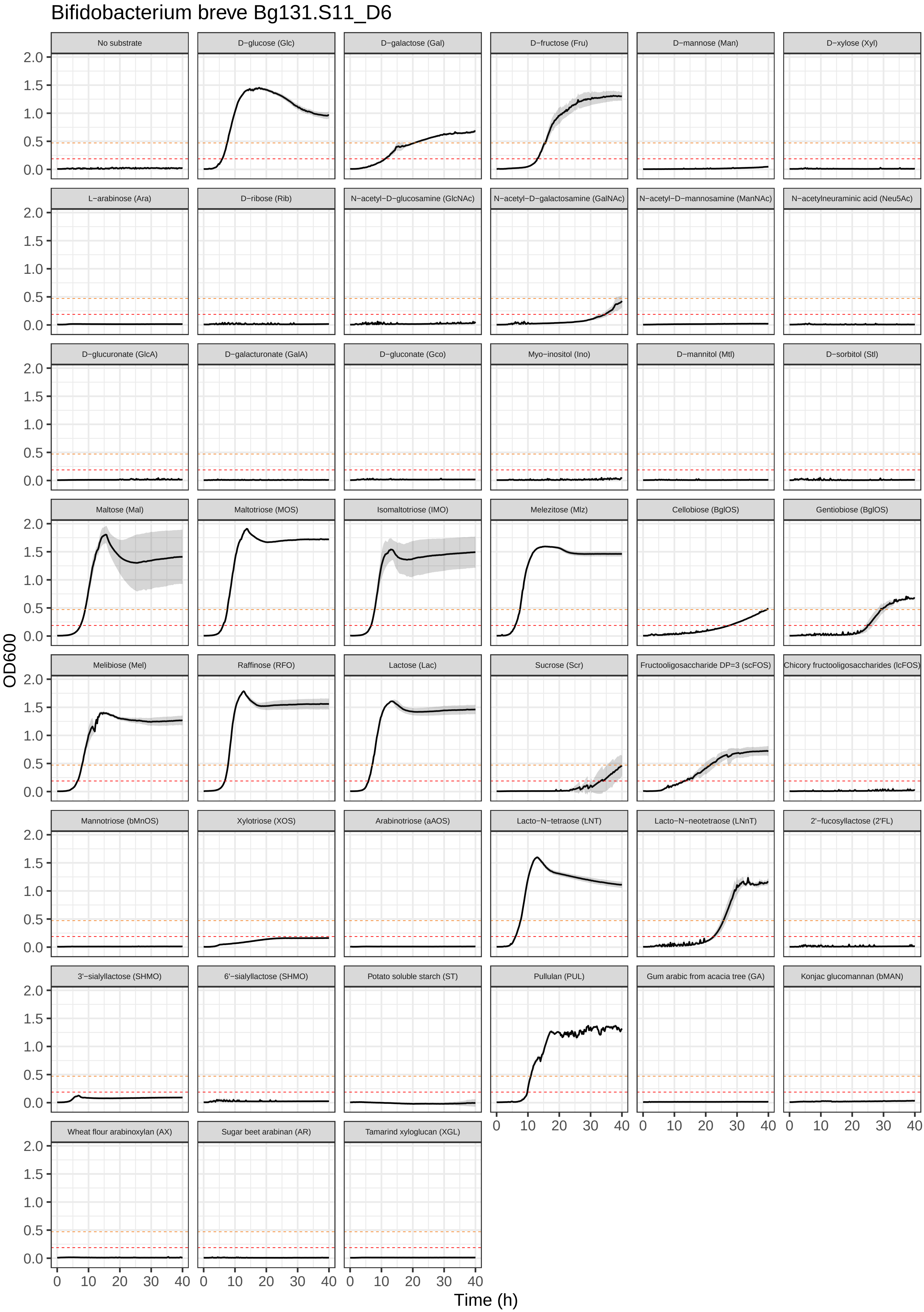

Bifidobacterium breve Bg131.S11_D6
No substrate
D−glucose (Glc)
D−galactose (Gal)
D−fructose (Fru)
D−mannose (Man)
D−xylose (Xyl)
L−arabinose (Ara)
D−ribose (Rib)
N−acetyl−D−glucosamine (GlcNAc)
N−acetyl−D−galactosamine (GalNAc)
N−acetyl−D−mannosamine (ManNAc)
N−acetylneuraminic acid (Neu5Ac)
D−glucuronate (GlcA)
D−galacturonate (GalA)
D−gluconate (Gco)
Myo−inositol (Ino)
D−mannitol (Mtl)
D−sorbitol (Stl)
Maltose (Mal)
Maltotriose (MOS)
Isomaltotriose (IMO)
Melezitose (Mlz)
Cellobiose (BglOS)
Gentiobiose (BglOS)
Melibiose (Mel)
Raffinose (RFO)
Lactose (Lac)
Sucrose (Scr)
Fructooligosaccharide DP=3 (scFOS)
Chicory fructooligosaccharides (lcFOS)
Mannotriose (bMnOS)
Xylotriose (XOS)
Arabinotriose (aAOS)
Lacto−N−tetraose (LNT)
Lacto−N−neotetraose (LNnT)
2'−fucosyllactose (2'FL)
3'−sialyllactose (SHMO)
6'−sialyllactose (SHMO)
Potato soluble starch (ST)
Pullulan (PUL)
Gum arabic from acacia tree (GA)
Konjac glucomannan (bMAN)
Wheat flour arabinoxylan (AX)
Sugar beet arabinan (AR)
Tamarind xyloglucan (XGL)
OD600
Time (h)
0.0
0.5
1.0
1.5
2.0
0
10
20
30
40

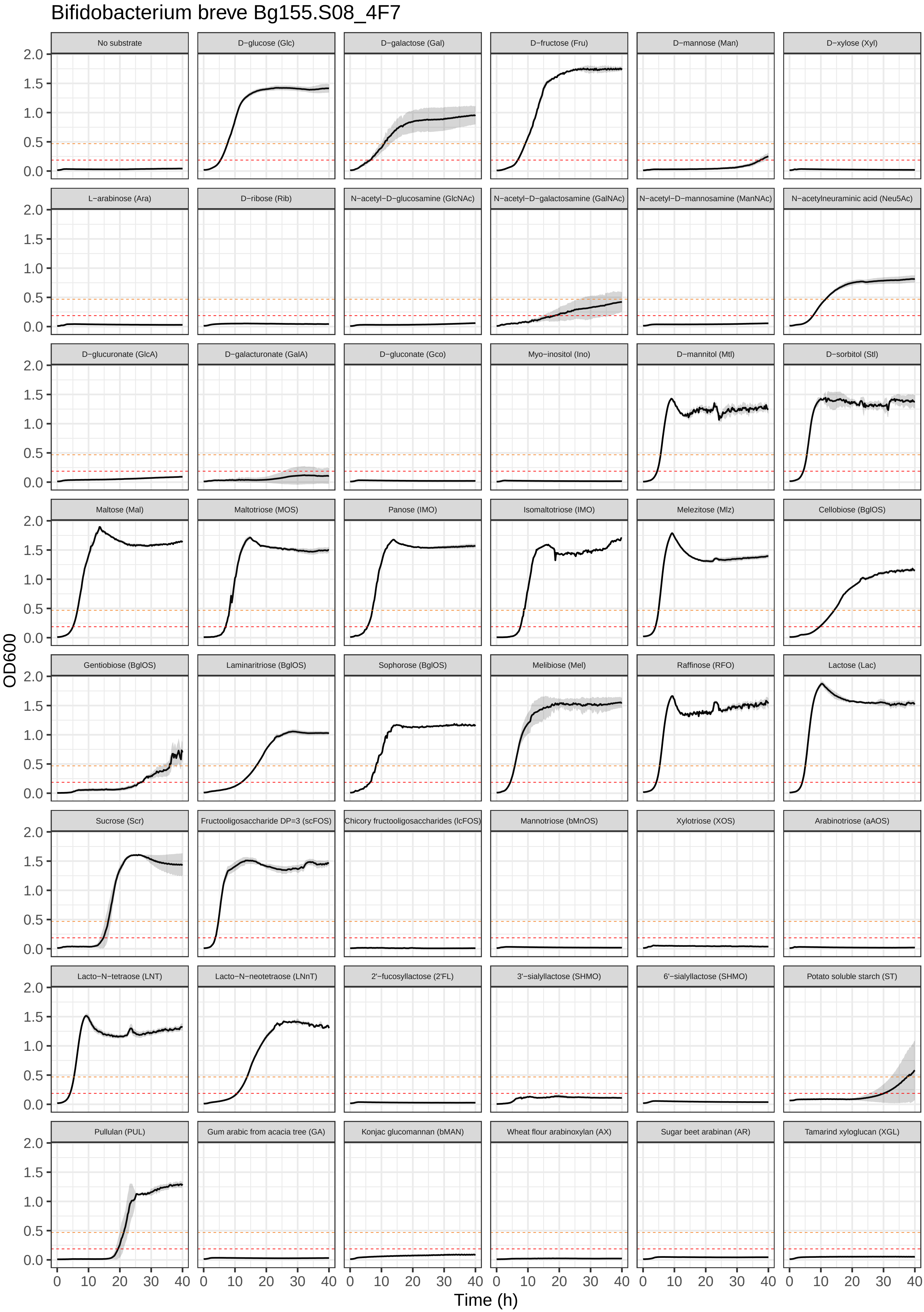
Bifidobacterium breve Bg155.S08_4F7
No substrate
D−glucose (Glc)
D−galactose (Gal)
D−fructose (Fru)
D−mannose (Man)
D−xylose (Xyl)
L−arabinose (Ara)
D−ribose (Rib)
N−acetyl−D−glucosamine (GlcNAc)
N−acetyl−D−galactosamine (GalNAc)
N−acetyl−D−mannosamine (ManNAc)
N−acetylneuraminic acid (Neu5Ac)
D−glucuronate (GlcA)
D−galacturonate (GalA)
D−gluconate (Gco)
Myo−inositol (Ino)
D−mannitol (Mtl)
D−sorbitol (Stl)
Maltose (Mal)
Maltotriose (MOS)
Panose (IMO)
Isomaltotriose (IMO)
Melezitose (Mlz)
Cellobiose (BglOS)
Gentiobiose (BglOS)
Laminaritriose (BglOS)
Sophorose (BglOS)
Melibiose (Mel)
Raffinose (RFO)
Lactose (Lac)
Sucrose (Scr)
Fructooligosaccharide DP=3 (scFOS)
Chicory fructooligosaccharides (lcFOS)
Mannotriose (bMnOS)
Xylotriose (XOS)
Arabinotriose (aAOS)
Lacto−N−tetraose (LNT)
Lacto−N−neotetraose (LNnT)
2'−fucosyllactose (2'FL)
3'−sialyllactose (SHMO)
6'−sialyllactose (SHMO)
Potato soluble starch (ST)
Pullulan (PUL)
Gum arabic from acacia tree (GA)
Konjac glucomannan (bMAN)
Wheat flour arabinoxylan (AX)
Sugar beet arabinan (AR)
Tamarind xyloglucan (XGL)
OD600
Time (h)

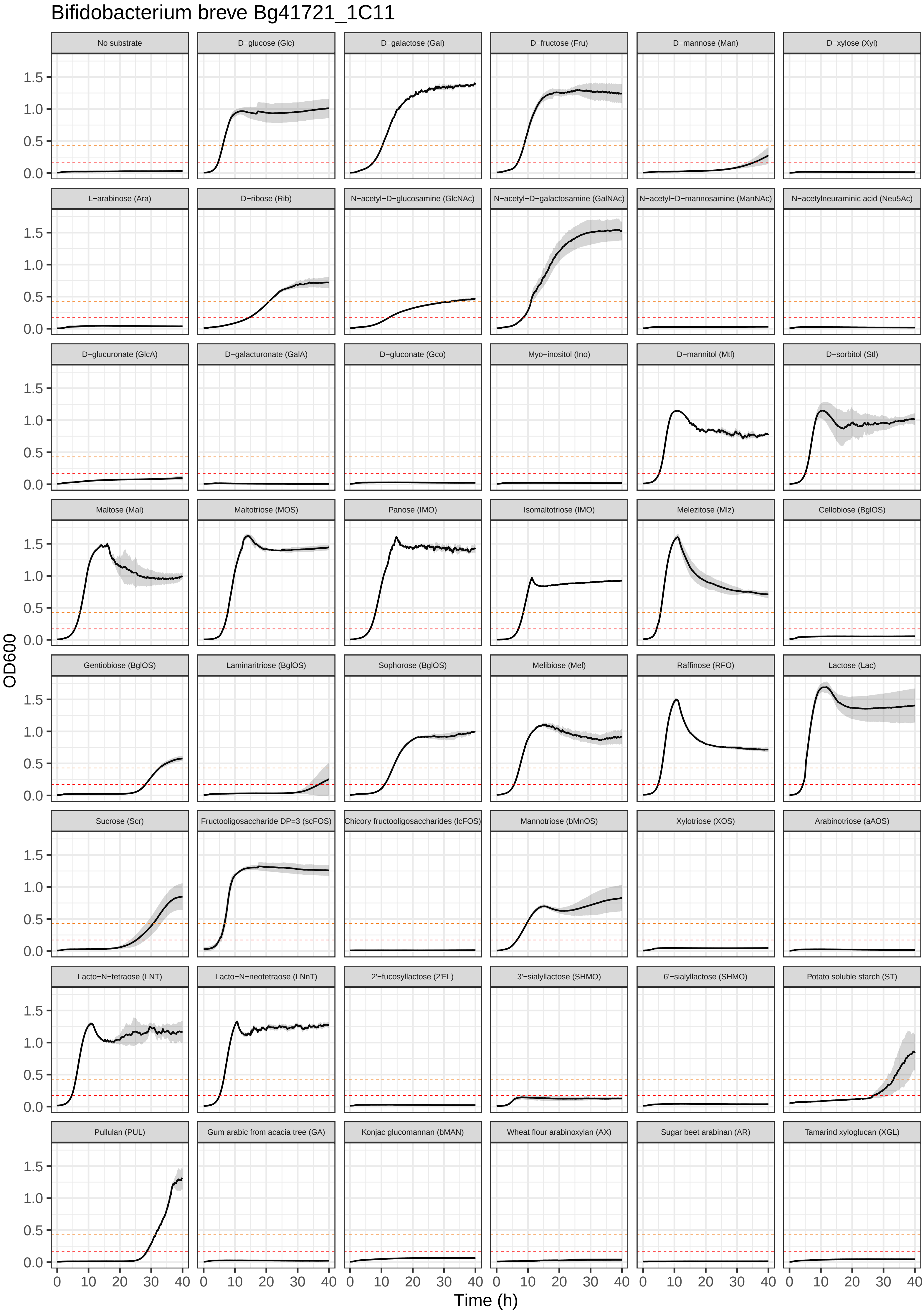

Bifidobacterium breve Bg41721_1C11
No substrate
D−glucose (Glc)
D−galactose (Gal)
D−fructose (Fru)
D−mannose (Man)
D−xylose (Xyl)
L−arabinose (Ara)
D−ribose (Rib)
N−acetyl−D−glucosamine (GlcNAc)
N−acetyl−D−galactosamine (GalNAc)
N−acetyl−D−mannosamine (ManNAc)
N−acetylneuraminic acid (Neu5Ac)
D−glucuronate (GlcA)
D−galacturonate (GalA)
D−gluconate (Gco)
Myo−inositol (Ino)
D−mannitol (Mtl)
D−sorbitol (Stl)
Maltose (Mal)
Maltotriose (MOS)
Panose (IMO)
Isomaltotriose (IMO)
Melezitose (Mlz)
Cellobiose (BglOS)
Gentiobiose (BglOS)
Laminaritriose (BglOS)
Sophorose (BglOS)
Melibiose (Mel)
Raffinose (RFO)
Lactose (Lac)
Sucrose (Scr)
Fructooligosaccharide DP=3 (scFOS)
Chicory fructooligosaccharides (lcFOS)
Mannotriose (bMnOS)
Xylotriose (XOS)
Arabinotriose (aAOS)
Lacto−N−tetraose (LNT)
Lacto−N−neotetraose (LNnT)
2'−fucosyllactose (2'FL)
3'−sialyllactose (SHMO)
6'−sialyllactose (SHMO)
Potato soluble starch (ST)
Pullulan (PUL)
Gum arabic from acacia tree (GA)
Konjac glucomannan (bMAN)
Wheat flour arabinoxylan (AX)
Sugar beet arabinan (AR)
Tamarind xyloglucan (XGL)
OD600
Time (h)
0.0
0.5
1.0
1.5
0
10
20
30
40

#### Bifidobacterium breve JG_Bg463

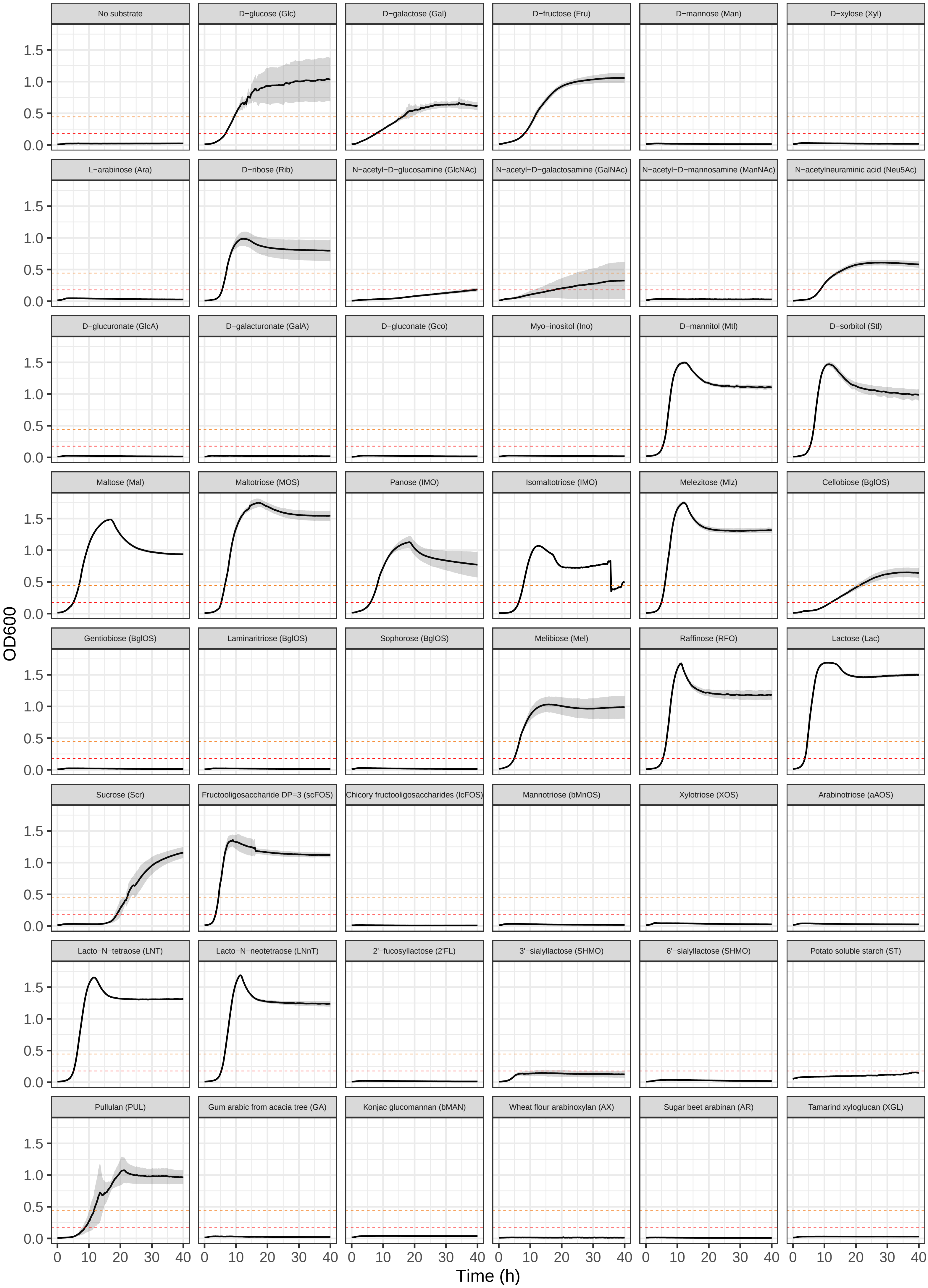

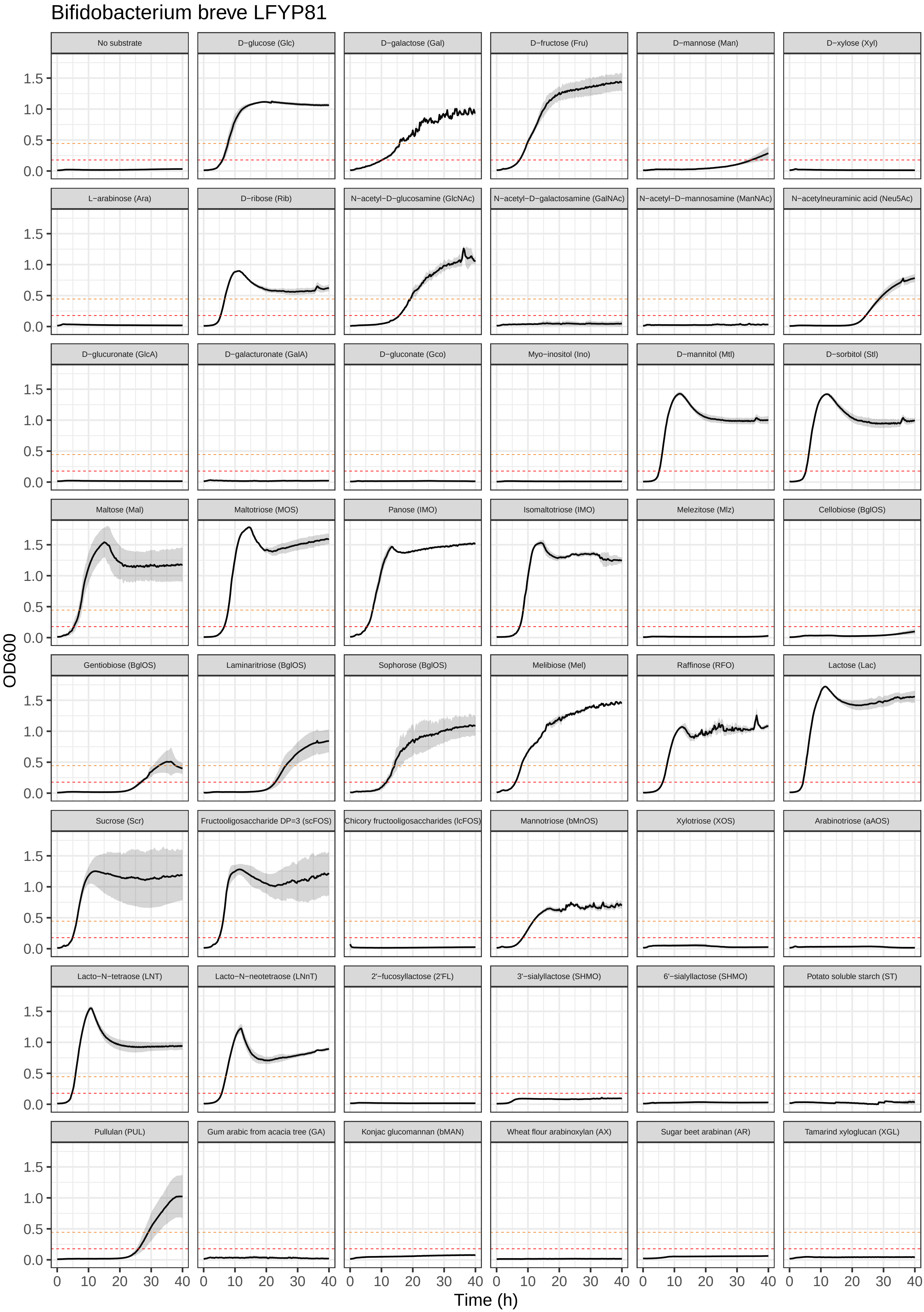

Bifidobacterium breve LFYP81
No substrate
D−glucose (Glc)
D−galactose (Gal)
D−fructose (Fru)
D−mannose (Man)
D−xylose (Xyl)
L−arabinose (Ara)
D−ribose (Rib)
N−acetyl−D−glucosamine (GlcNAc)
N−acetyl−D−galactosamine (GalNAc)
N−acetyl−D−mannosamine (ManNAc)
N−acetylneuraminic acid (Neu5Ac)
D−glucuronate (GlcA)
D−galacturonate (GalA)
D−gluconate (Gco)
Myo−inositol (Ino)
D−mannitol (Mtl)
D−sorbitol (Stl)
Maltose (Mal)
Maltotriose (MOS)
Panose (IMO)
Isomaltotriose (IMO)
Melezitose (Mlz)
Cellobiose (BglOS)
Gentiobiose (BglOS)
Laminaritriose (BglOS)
Sophorose (BglOS)
Melibiose (Mel)
Raffinose (RFO)
Lactose (Lac)
Sucrose (Scr)
Fructooligosaccharide DP=3 (scFOS)
Chicory fructooligosaccharides (lcFOS)
Mannotriose (bMnOS)
Xylotriose (XOS)
Arabinotriose (aAOS)
Lacto−N−tetraose (LNT)
Lacto−N−neotetraose (LNnT)
2'−fucosyllactose (2'FL)
3'−sialyllactose (SHMO)
6'−sialyllactose (SHMO)
Potato soluble starch (ST)
Pullulan (PUL)
Gum arabic from acacia tree (GA)
Konjac glucomannan (bMAN)
Wheat flour arabinoxylan (AX)
Sugar beet arabinan (AR)
Tamarind xyloglucan (XGL)
OD600
0.0
0.5
1.0
1.5
Time (h)
0
10
20
30
40

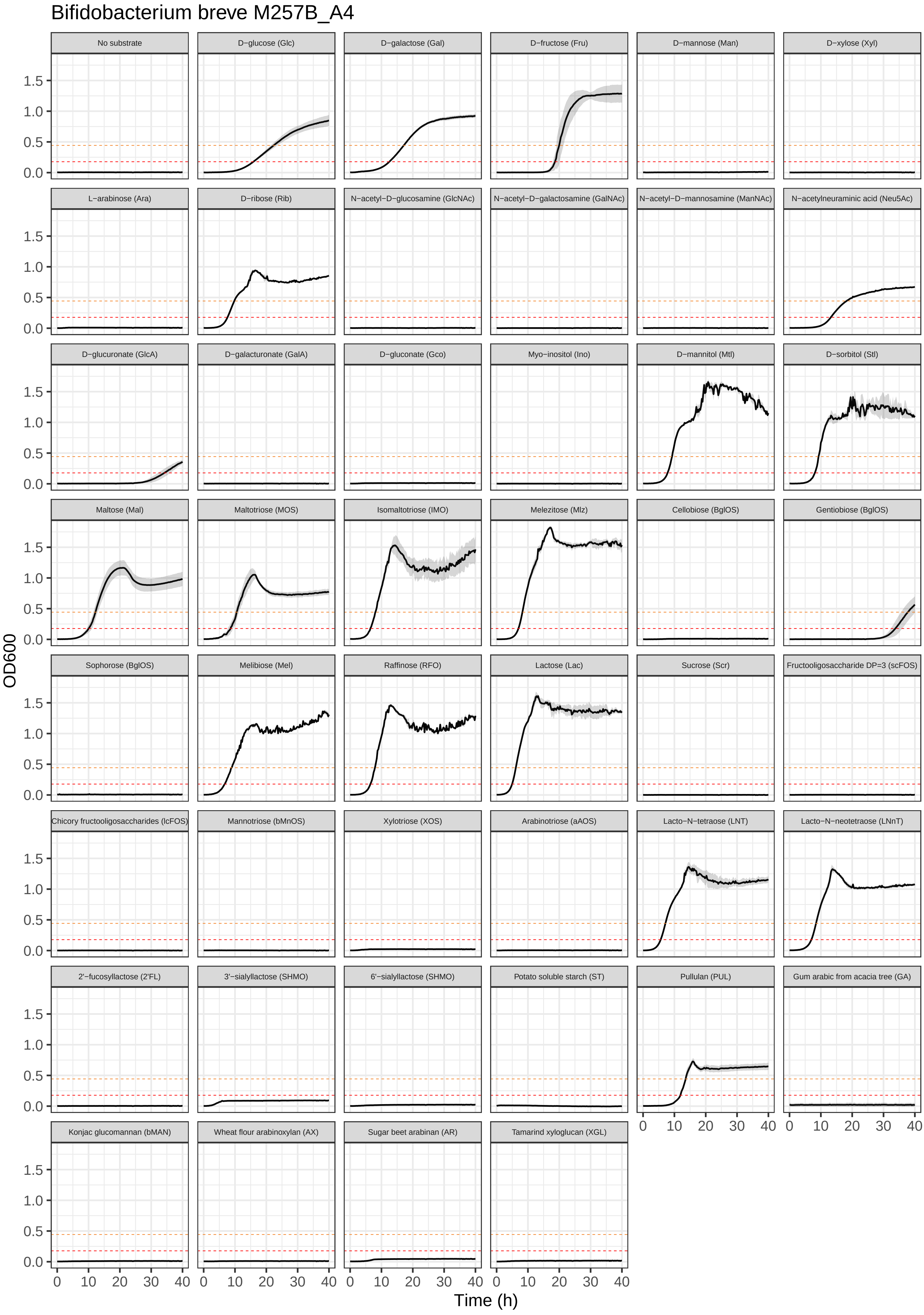
Bifidobacterium breve M257B_A4
No substrate
D−glucose (Glc)
D−galactose (Gal)
D−fructose (Fru)
D−mannose (Man)
D−xylose (Xyl)
L−arabinose (Ara)
D−ribose (Rib)
N−acetyl−D−glucosamine (GlcNAc)
N−acetyl−D−galactosamine (GalNAc)
N−acetyl−D−mannosamine (ManNAc)
N−acetylneuraminic acid (Neu5Ac)
D−glucuronate (GlcA)
D−galacturonate (GalA)
D−gluconate (Gco)
Myo−inositol (Ino)
D−mannitol (Mtl)
D−sorbitol (Stl)
Maltose (Mal)
Maltotriose (MOS)
Isomaltotriose (IMO)
Melezitose (Mlz)
Cellobiose (BglOS)
Gentiobiose (BglOS)
Sophorose (BglOS)
Melibiose (Mel)
Raffinose (RFO)
Lactose (Lac)
Sucrose (Scr)
Fructooligosaccharide DP=3 (scFOS)
Chicory fructooligosaccharides (lcFOS)
Mannotriose (bMnOS)
Xylotriose (XOS)
Arabinotriose (aAOS)
Lacto−N−tetraose (LNT)
Lacto−N−neotetraose (LNnT)
2'−fucosyllactose (2'FL)
3'−sialyllactose (SHMO)
6'−sialyllactose (SHMO)
Potato soluble starch (ST)
Pullulan (PUL)
Gum arabic from acacia tree (GA)
Konjac glucomannan (bMAN)
Wheat flour arabinoxylan (AX)
Sugar beet arabinan (AR)
Tamarind xyloglucan (XGL)
OD600
0.0
0.5
1.0
1.5
0
10
20
30
40
Time (h)

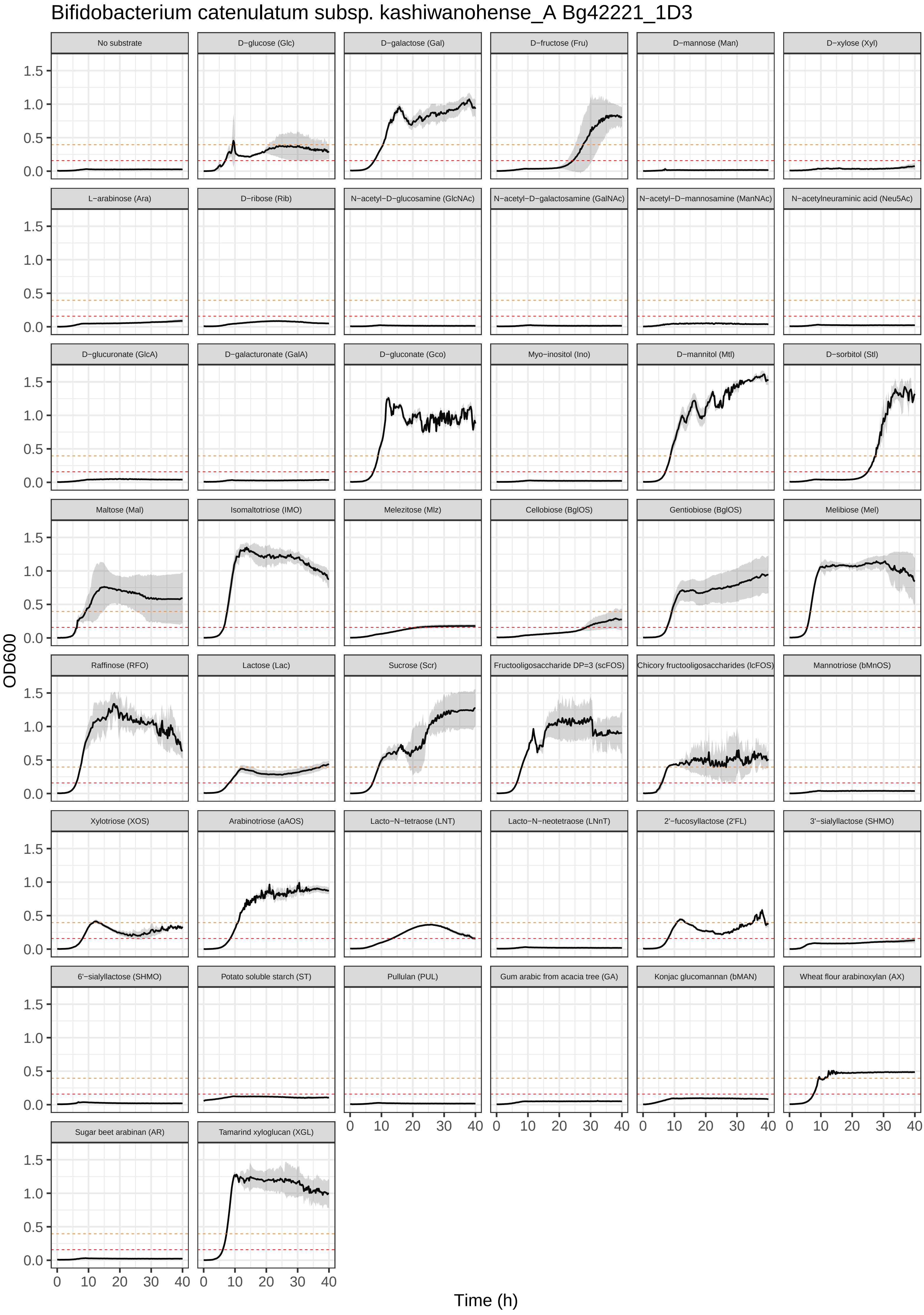

Bifidobacterium catenulatum subsp. kashiwanohense_A Bg42221_1D3
No substrate
D−glucose (Glc)
D−galactose (Gal)
D−fructose (Fru)
D−mannose (Man)
D−xylose (Xyl)
L−arabinose (Ara)
D−ribose (Rib)
N−acetyl−D−glucosamine (GlcNAc)
N−acetyl−D−galactosamine (GalNAc)
N−acetyl−D−mannosamine (ManNAc)
N−acetylneuraminic acid (Neu5Ac)
D−glucuronate (GlcA)
D−galacturonate (GalA)
D−gluconate (Gco)
Myo−inositol (Ino)
D−mannitol (Mtl)
D−sorbitol (Stl)
Maltose (Mal)
Isomaltotriose (IMO)
Melezitose (Mlz)
Cellobiose (BglOS)
Gentiobiose (BglOS)
Melibiose (Mel)
Raffinose (RFO)
Lactose (Lac)
Sucrose (Scr)
Fructooligosaccharide DP=3 (scFOS)
Chicory fructooligosaccharides (lcFOS)
Mannotriose (bMnOS)
Xylotriose (XOS)
Arabinotriose (aAOS)
Lacto−N−tetraose (LNT)
Lacto−N−neotetraose (LNnT)
2'−fucosyllactose (2'FL)
3'−sialyllactose (SHMO)
6'−sialyllactose (SHMO)
Potato soluble starch (ST)
Pullulan (PUL)
Gum arabic from acacia tree (GA)
Konjac glucomannan (bMAN)
Wheat flour arabinoxylan (AX)
Sugar beet arabinan (AR)
Tamarind xyloglucan (XGL)
OD600
0.0
0.5
1.0
1.5
0
10
20
30
40
Time (h)

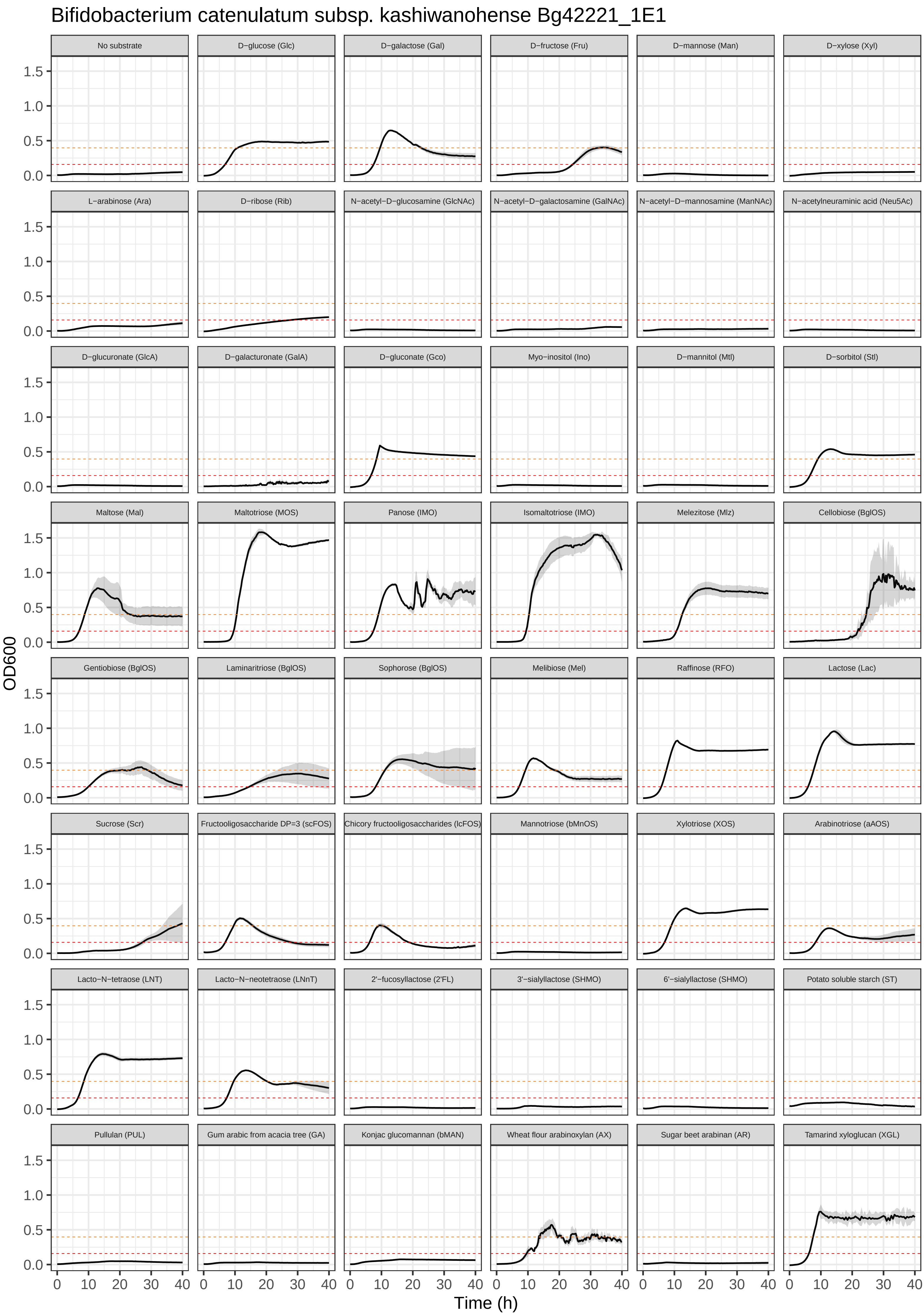
Bifidobacterium catenulatum subsp. kashiwanohense Bg42221_1E1
No substrate
D−glucose (Glc)
D−galactose (Gal)
D−fructose (Fru)
D−mannose (Man)
D−xylose (Xyl)
L−arabinose (Ara)
D−ribose (Rib)
N−acetyl−D−glucosamine (GlcNAc)
N−acetyl−D−galactosamine (GalNAc)
N−acetyl−D−mannosamine (ManNAc)
N−acetylneuraminic acid (Neu5Ac)
D−glucuronate (GlcA)
D−galacturonate (GalA)
D−gluconate (Gco)
Myo−inositol (Ino)
D−mannitol (Mtl)
D−sorbitol (Stl)
Maltose (Mal)
Maltotriose (MOS)
Panose (IMO)
Isomaltotriose (IMO)
Melezitose (Mlz)
Cellobiose (BglOS)
Gentiobiose (BglOS)
Laminaritriose (BglOS)
Sophorose (BglOS)
Melibiose (Mel)
Raffinose (RFO)
Lactose (Lac)
Sucrose (Scr)
Fructooligosaccharide DP=3 (scFOS)
Chicory fructooligosaccharides (lcFOS)
Mannotriose (bMnOS)
Xylotriose (XOS)
Arabinotriose (aAOS)
Lacto−N−tetraose (LNT)
Lacto−N−neotetraose (LNnT)
2'−fucosyllactose (2'FL)
3'−sialyllactose (SHMO)
6'−sialyllactose (SHMO)
Potato soluble starch (ST)
Pullulan (PUL)
Gum arabic from acacia tree (GA)
Konjac glucomannan (bMAN)
Wheat flour arabinoxylan (AX)
Sugar beet arabinan (AR)
Tamarind xyloglucan (XGL)
OD600
0.0
0.5
1.0
1.5
Time (h)
0
10
20
30
40

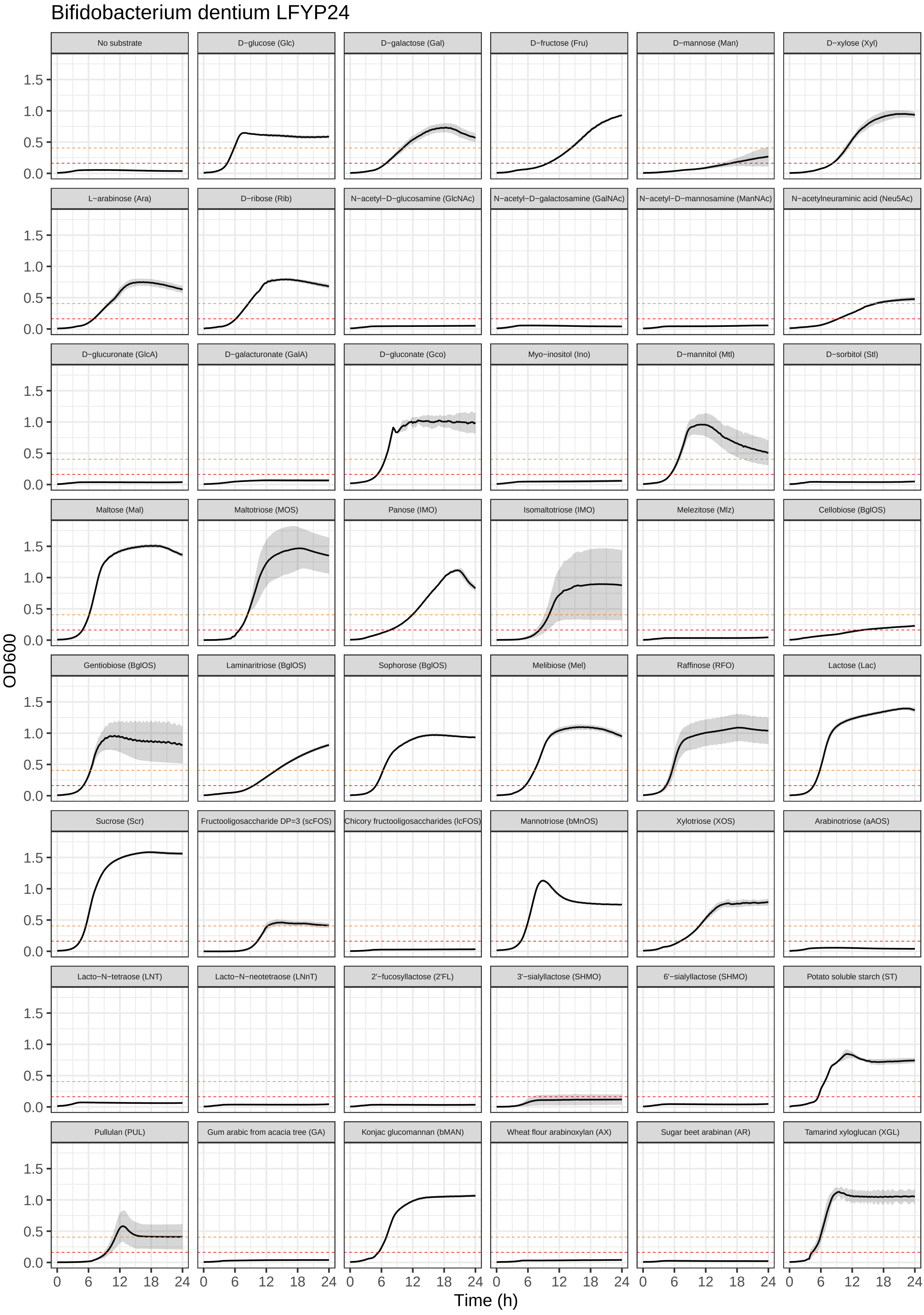
Bifidobacterium dentium LFYP24
No substrate
D−glucose (Glc)
D−galactose (Gal)
D−fructose (Fru)
D−mannose (Man)
D−xylose (Xyl)
L−arabinose (Ara)
D−ribose (Rib)
N−acetyl−D−glucosamine (GlcNAc)
N−acetyl−D−galactosamine (GalNAc)
N−acetyl−D−mannosamine (ManNAc)
N−acetylneuraminic acid (Neu5Ac)
D−glucuronate (GlcA)
D−galacturonate (GalA)
D−gluconate (Gco)
Myo−inositol (Ino)
D−mannitol (Mtl)
D−sorbitol (Stl)
Maltose (Mal)
Maltotriose (MOS)
Panose (IMO)
Isomaltotriose (IMO)
Melezitose (Mlz)
Cellobiose (BglOS)
Gentiobiose (BglOS)
Laminaritriose (BglOS)
Sophorose (BglOS)
Melibiose (Mel)
Raffinose (RFO)
Lactose (Lac)
Sucrose (Scr)
Fructooligosaccharide DP=3 (scFOS)
Chicory fructooligosaccharides (lcFOS)
Mannotriose (bMnOS)
Xylotriose (XOS)
Arabinotriose (aAOS)
Lacto−N−tetraose (LNT)
Lacto−N−neotetraose (LNnT)
2'−fucosyllactose (2'FL)
3'−sialyllactose (SHMO)
6'−sialyllactose (SHMO)
Potato soluble starch (ST)
Pullulan (PUL)
Gum arabic from acacia tree (GA)
Konjac glucomannan (bMAN)
Wheat flour arabinoxylan (AX)
Sugar beet arabinan (AR)
Tamarind xyloglucan (XGL)
OD600
0.0
0.5
1.0
1.5
0
6
12
18
24
Time (h)

Bifidobacterium hominis Bg064.11_2H10

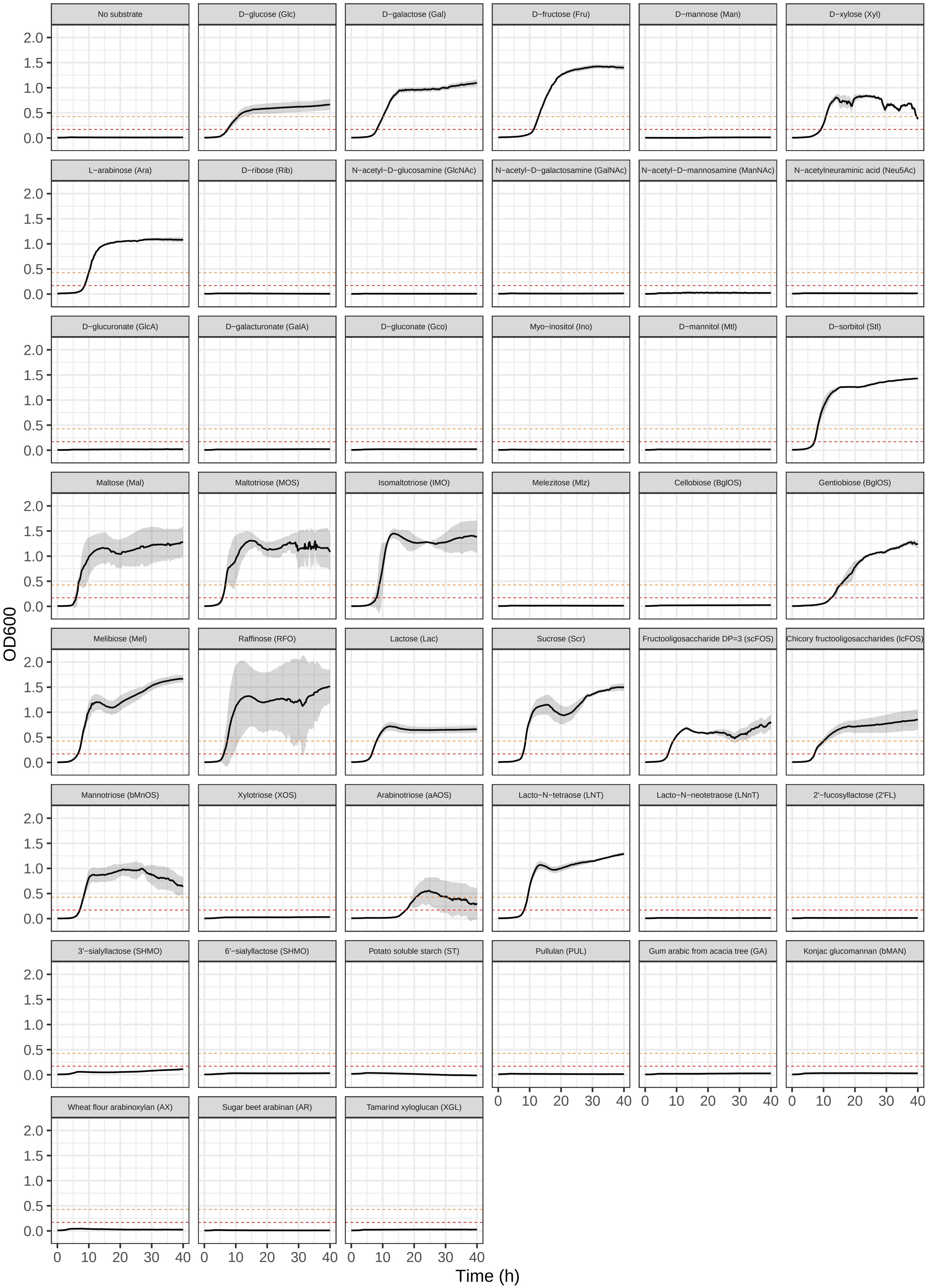

Bifidobacterium hominis Bg155.08_4B11

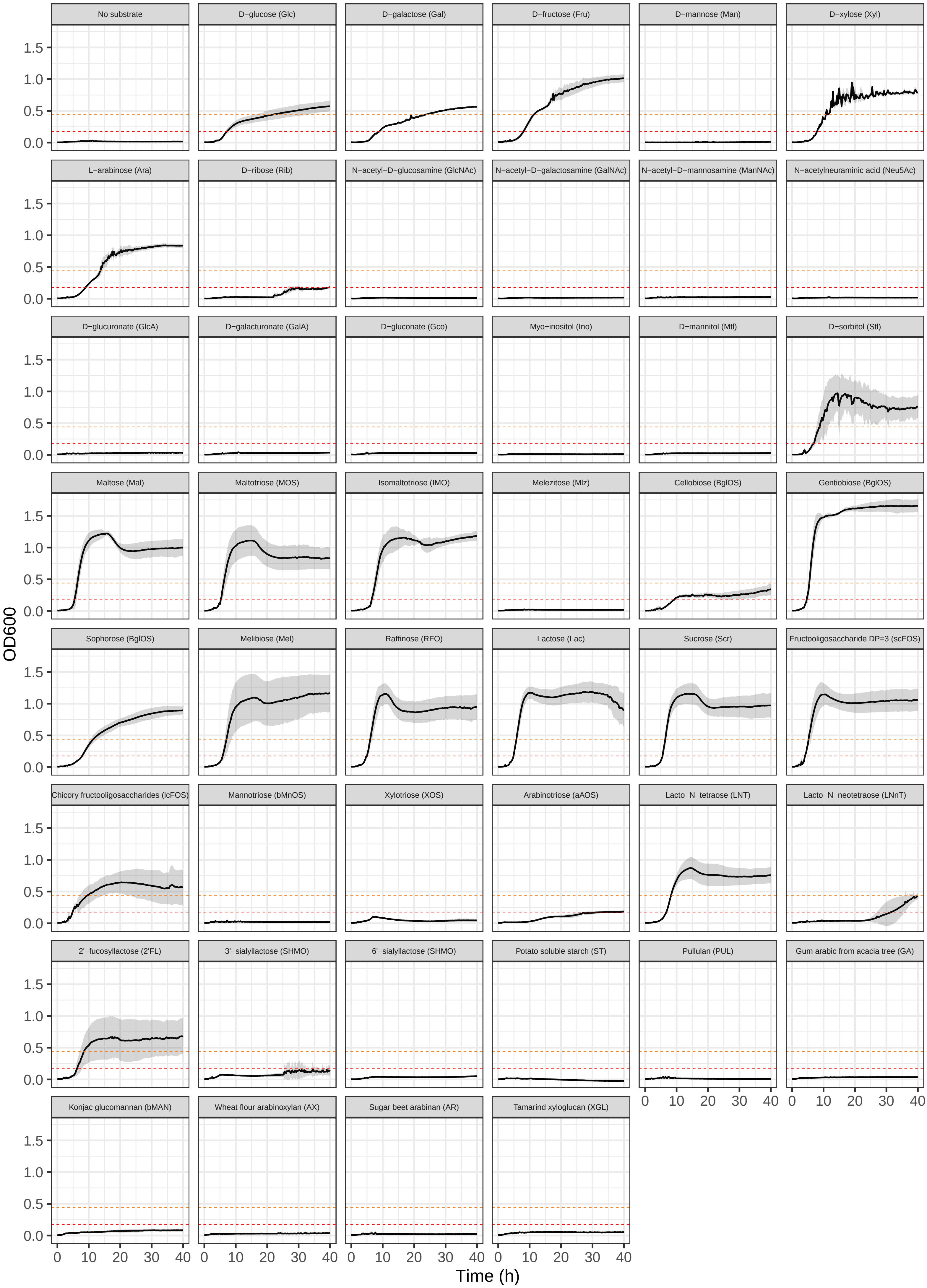

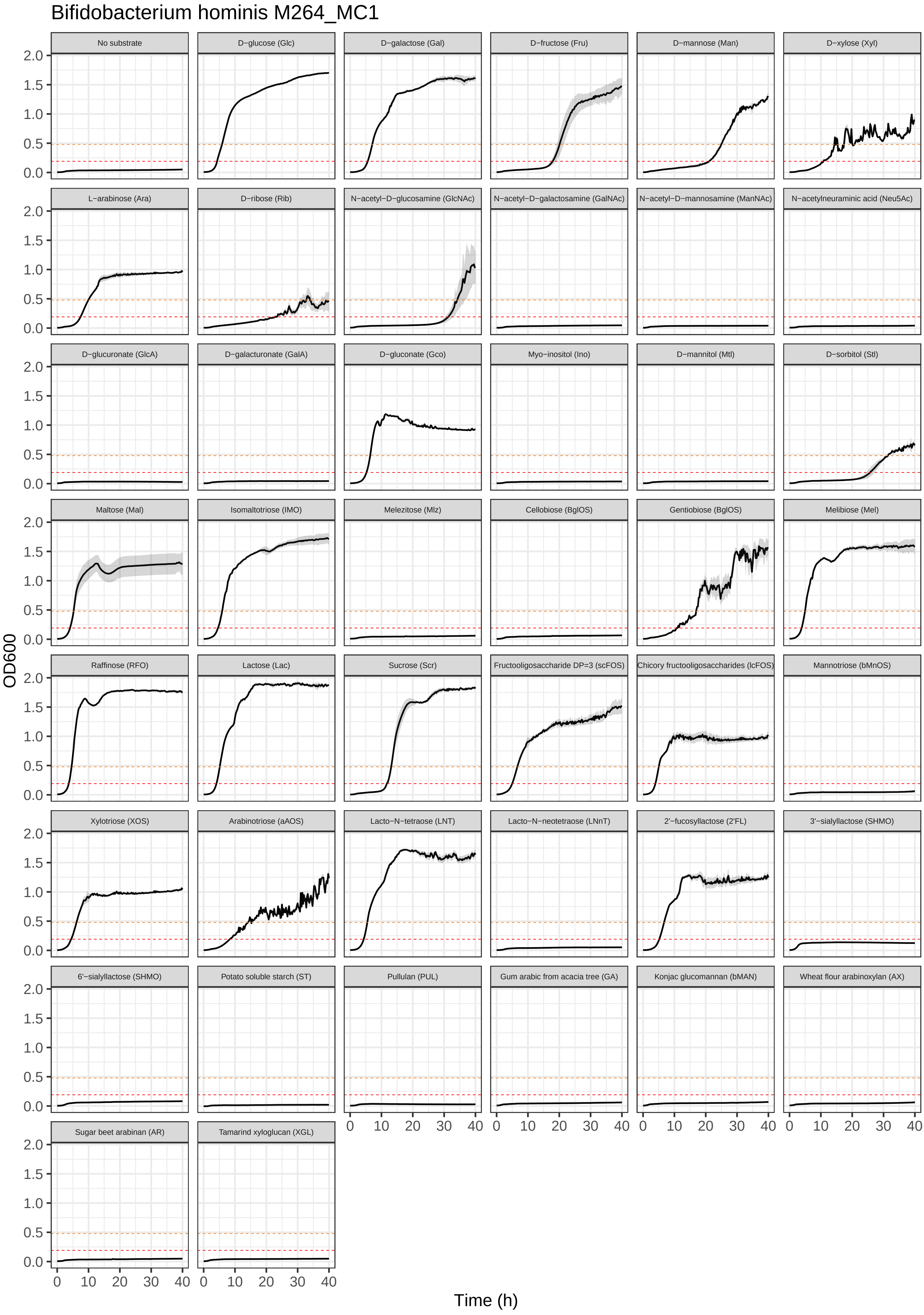
Bifidobacterium hominis M264_MC1
No substrate
D−glucose (Glc)
D−galactose (Gal)
D−fructose (Fru)
D−mannose (Man)
D−xylose (Xyl)
L−arabinose (Ara)
D−ribose (Rib)
N−acetyl−D−glucosamine (GlcNAc)
N−acetyl−D−galactosamine (GalNAc)
N−acetyl−D−mannosamine (ManNAc)
N−acetylneuraminic acid (Neu5Ac)
D−glucuronate (GlcA)
D−galacturonate (GalA)
D−gluconate (Gco)
Myo−inositol (Ino)
D−mannitol (Mtl)
D−sorbitol (Stl)
Maltose (Mal)
Isomaltotriose (IMO)
Melezitose (Mlz)
Cellobiose (BglOS)
Gentiobiose (BglOS)
Melibiose (Mel)
Raffinose (RFO)
Lactose (Lac)
Sucrose (Scr)
Fructooligosaccharide DP=3 (scFOS)
Chicory fructooligosaccharides (lcFOS)
Mannotriose (bMnOS)
Xylotriose (XOS)
Arabinotriose (aAOS)
Lacto−N−tetraose (LNT)
Lacto−N−neotetraose (LNnT)
2'−fucosyllactose (2'FL)
3'−sialyllactose (SHMO)
6'−sialyllactose (SHMO)
Potato soluble starch (ST)
Pullulan (PUL)
Gum arabic from acacia tree (GA)
Konjac glucomannan (bMAN)
Wheat flour arabinoxylan (AX)
Sugar beet arabinan (AR)
Tamarind xyloglucan (XGL)
OD600
0.0
0.5
1.0
1.5
2.0
Time (h)
0
10
20
30
40

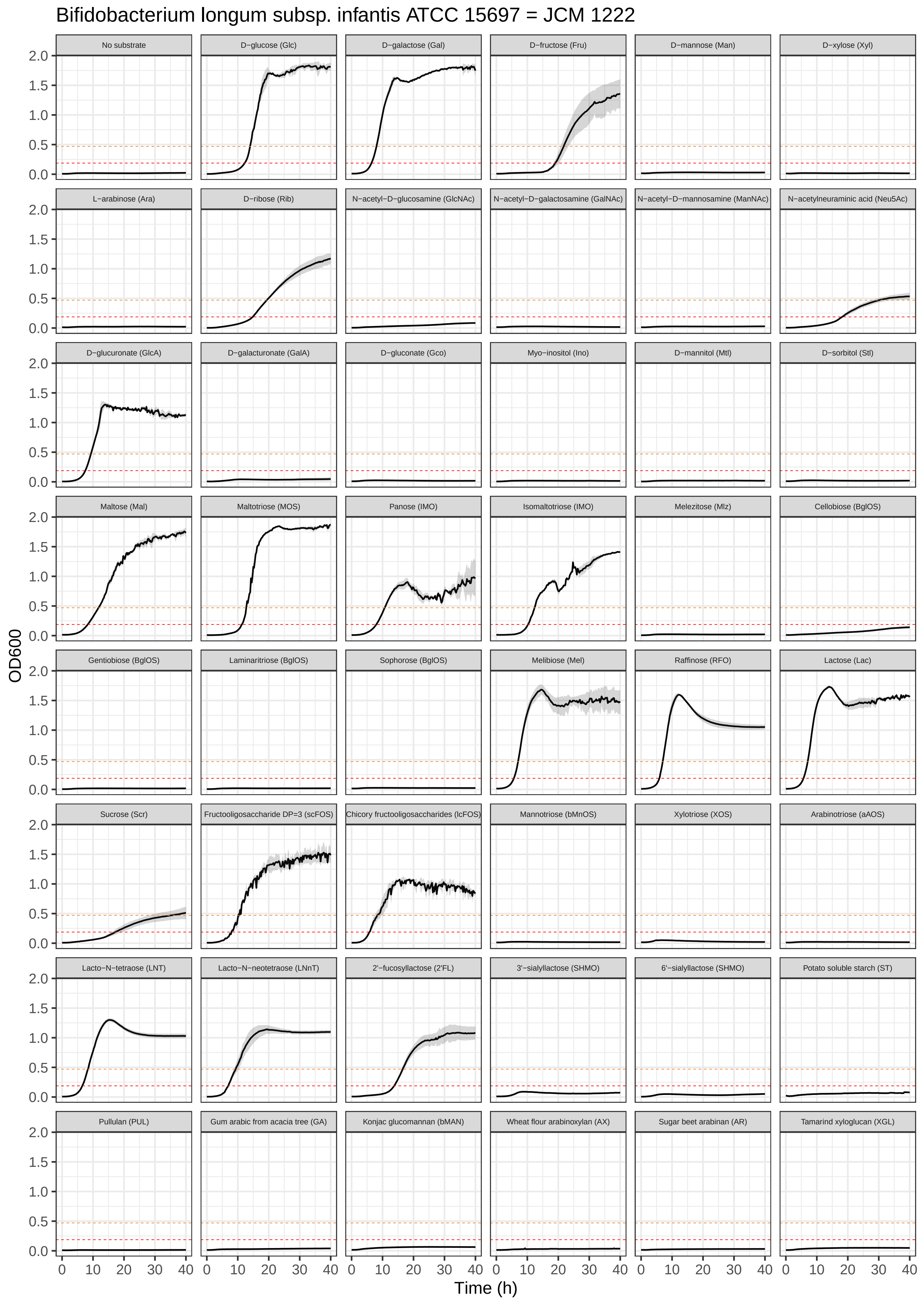
Bifidobacterium longum subsp. infantis ATCC 15697 = JCM 1222
No substrate
D−glucose (Glc)
D−galactose (Gal)
D−fructose (Fru)
D−mannose (Man)
D−xylose (Xyl)
L−arabinose (Ara)
D−ribose (Rib)
N−acetyl−D−glucosamine (GlcNAc)
N−acetyl−D−galactosamine (GalNAc)
N−acetyl−D−mannosamine (ManNAc)
N−acetylneuraminic acid (Neu5Ac)
D−glucuronate (GlcA)
D−galacturonate (GalA)
D−gluconate (Gco)
Myo−inositol (Ino)
D−mannitol (Mtl)
D−sorbitol (Stl)
Maltose (Mal)
Maltotriose (MOS)
Panose (IMO)
Isomaltotriose (IMO)
Melezitose (Mlz)
Cellobiose (BglOS)
Gentiobiose (BglOS)
Laminaritriose (BglOS)
Sophorose (BglOS)
Melibiose (Mel)
Raffinose (RFO)
Lactose (Lac)
Sucrose (Scr)
Fructooligosaccharide DP=3 (scFOS)
Chicory fructooligosaccharides (lcFOS)
Mannotriose (bMnOS)
Xylotriose (XOS)
Arabinotriose (aAOS)
Lacto−N−tetraose (LNT)
Lacto−N−neotetraose (LNnT)
2'−fucosyllactose (2'FL)
3'−sialyllactose (SHMO)
6'−sialyllactose (SHMO)
Potato soluble starch (ST)
Pullulan (PUL)
Gum arabic from acacia tree (GA)
Konjac glucomannan (bMAN)
Wheat flour arabinoxylan (AX)
Sugar beet arabinan (AR)
Tamarind xyloglucan (XGL)
OD600
Time (h)
0.0
0.5
1.0
1.5
2.0
0
10
20
30
40

Bifidobacterium longum subsp. infantis Bg064.S07_13.C6

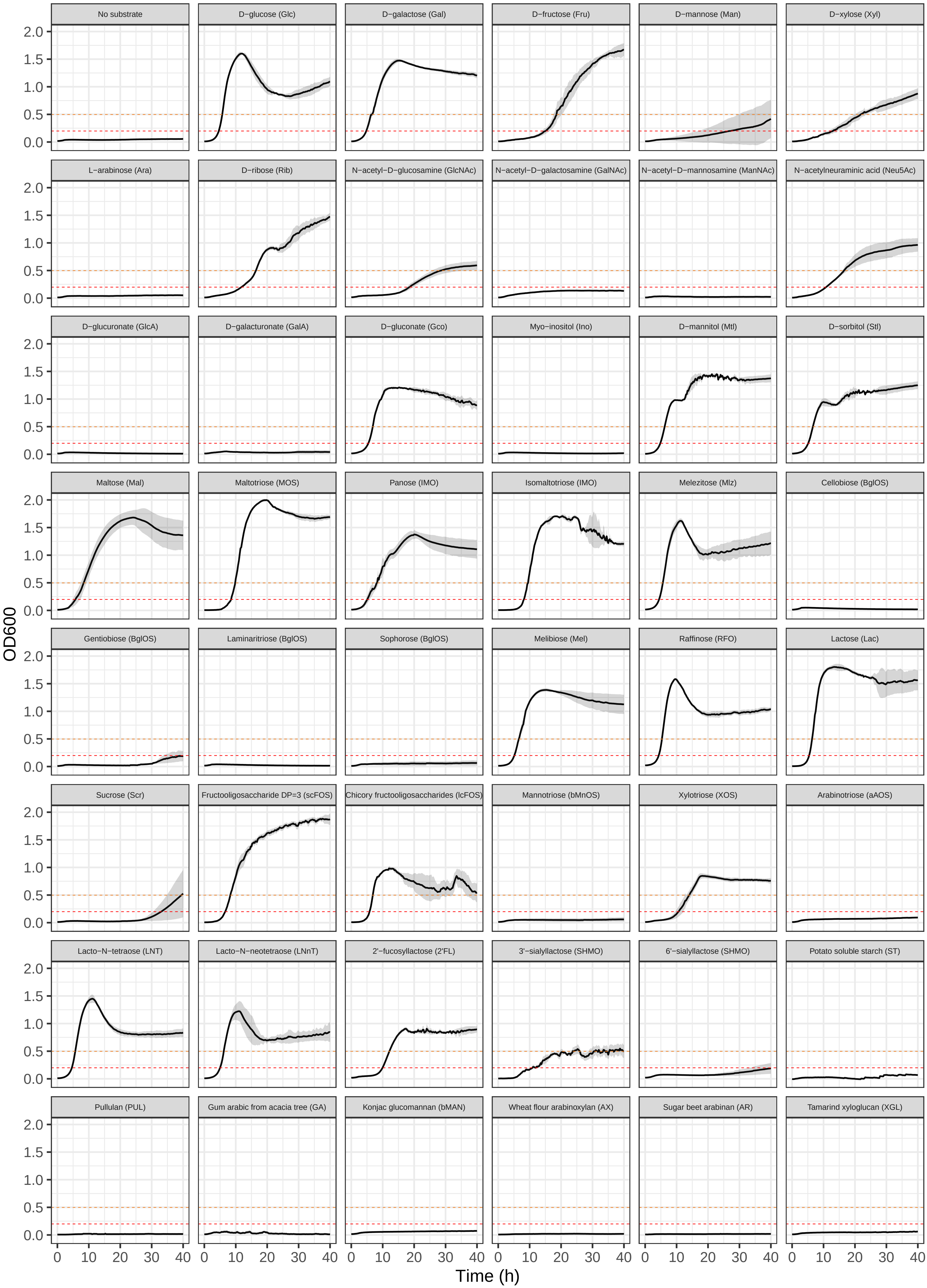

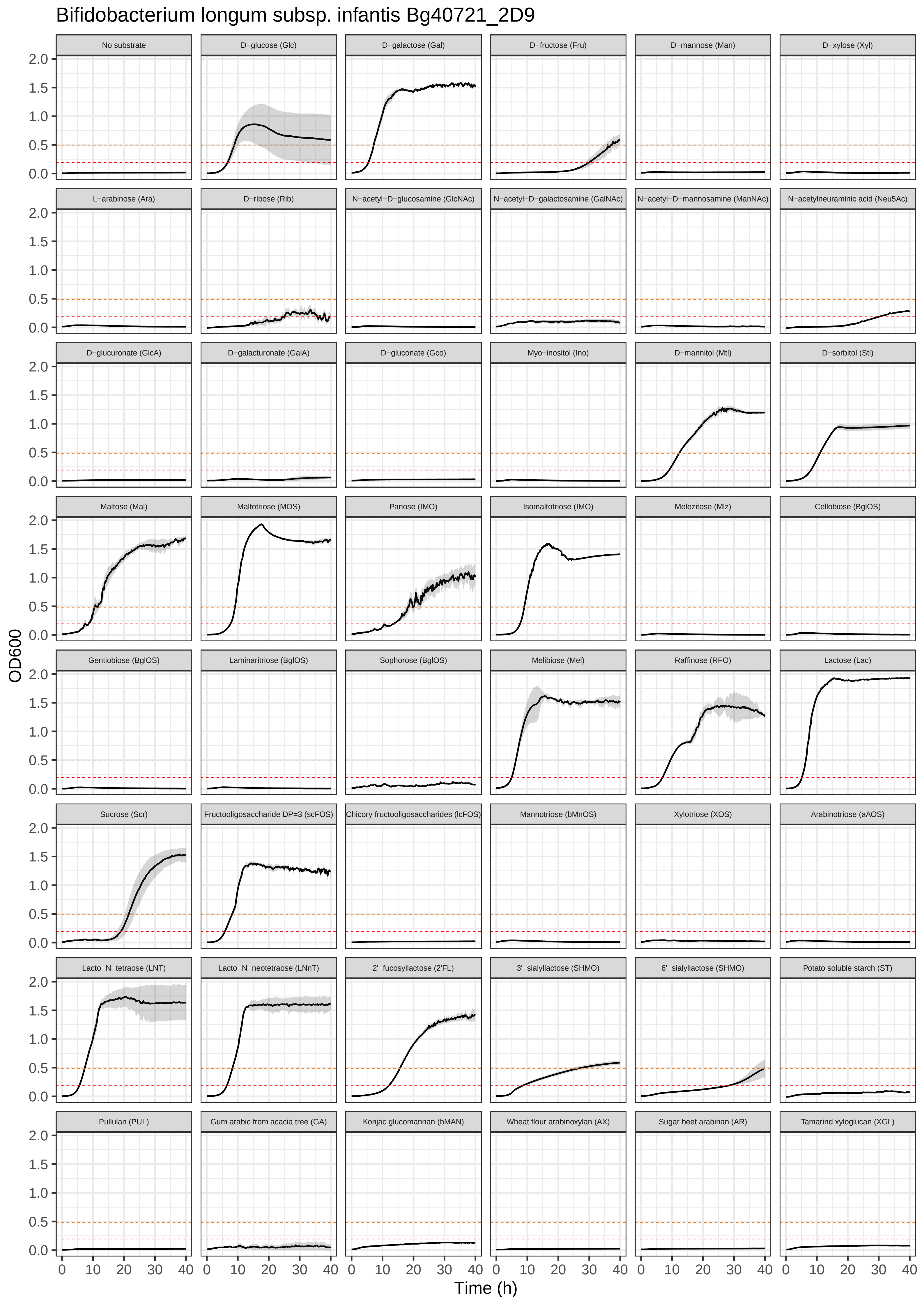
Bifidobacterium longum subsp. infantis Bg40721_2D9
No substrate
D−glucose (Glc)
D−galactose (Gal)
D−fructose (Fru)
D−mannose (Man)
D−xylose (Xyl)
L−arabinose (Ara)
D−ribose (Rib)
N−acetyl−D−glucosamine (GlcNAc)
N−acetyl−D−galactosamine (GalNAc)
N−acetyl−D−mannosamine (ManNAc)
N−acetylneuraminic acid (Neu5Ac)
D−glucuronate (GlcA)
D−galacturonate (GalA)
D−gluconate (Gco)
Myo−inositol (Ino)
D−mannitol (Mtl)
D−sorbitol (Stl)
Maltose (Mal)
Maltotriose (MOS)
Panose (IMO)
Isomaltotriose (IMO)
Melezitose (Mlz)
Cellobiose (BglOS)
Gentiobiose (BglOS)
Laminaritriose (BglOS)
Sophorose (BglOS)
Melibiose (Mel)
Raffinose (RFO)
Lactose (Lac)
Sucrose (Scr)
Fructooligosaccharide DP=3 (scFOS)
Chicory fructooligosaccharides (lcFOS)
Mannotriose (bMnOS)
Xylotriose (XOS)
Arabinotriose (aAOS)
Lacto−N−tetraose (LNT)
Lacto−N−neotetraose (LNnT)
2'−fucosyllactose (2'FL)
3'−sialyllactose (SHMO)
6'−sialyllactose (SHMO)
Potato soluble starch (ST)
Pullulan (PUL)
Gum arabic from acacia tree (GA)
Konjac glucomannan (bMAN)
Wheat flour arabinoxylan (AX)
Sugar beet arabinan (AR)
Tamarind xyloglucan (XGL)
OD600
Time (h)
0.0
0.5
1.0
1.5
2.0
0
10
20
30
40

Bifidobacterium longum subsp. infantis JG_Bg463

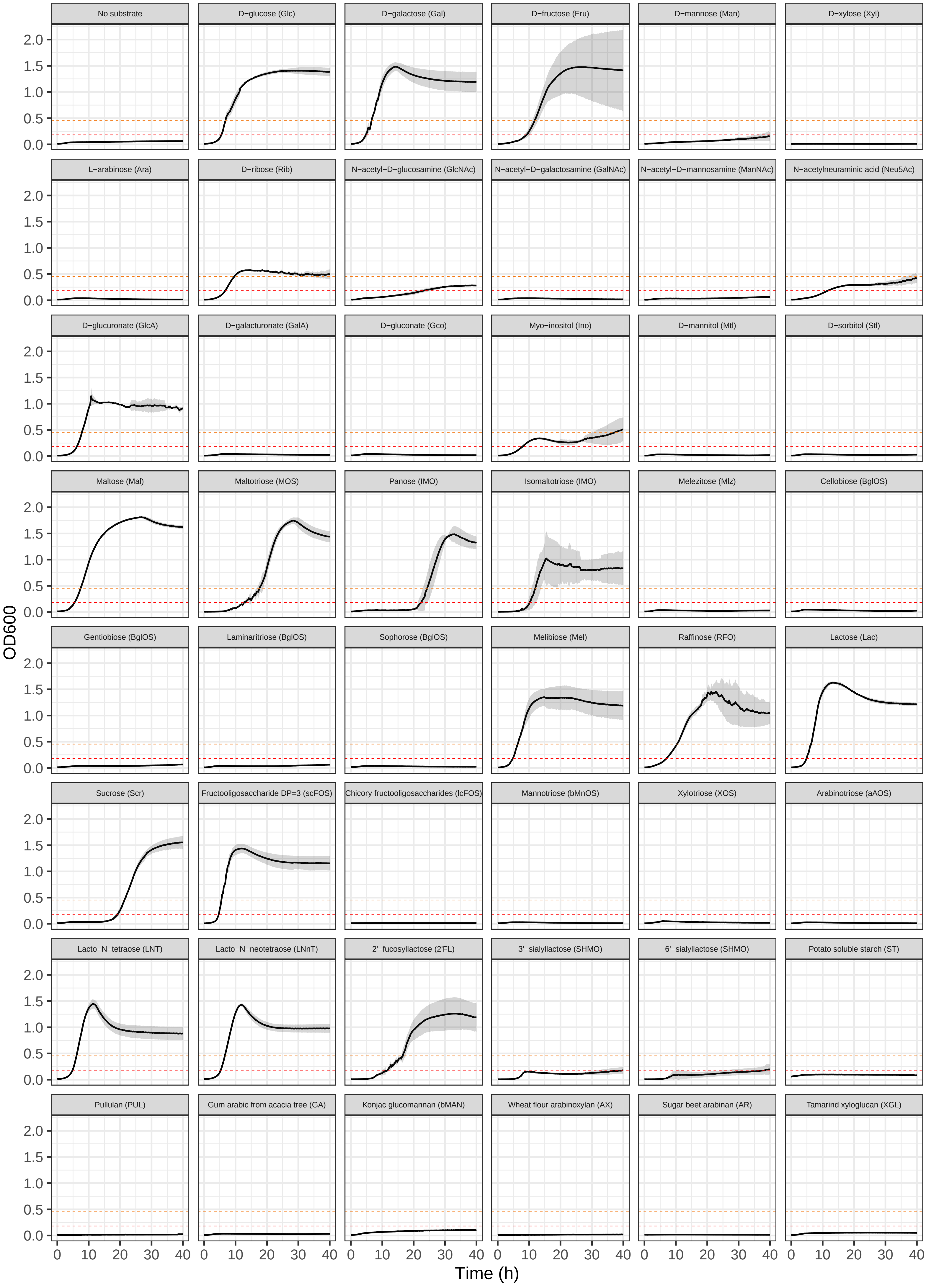

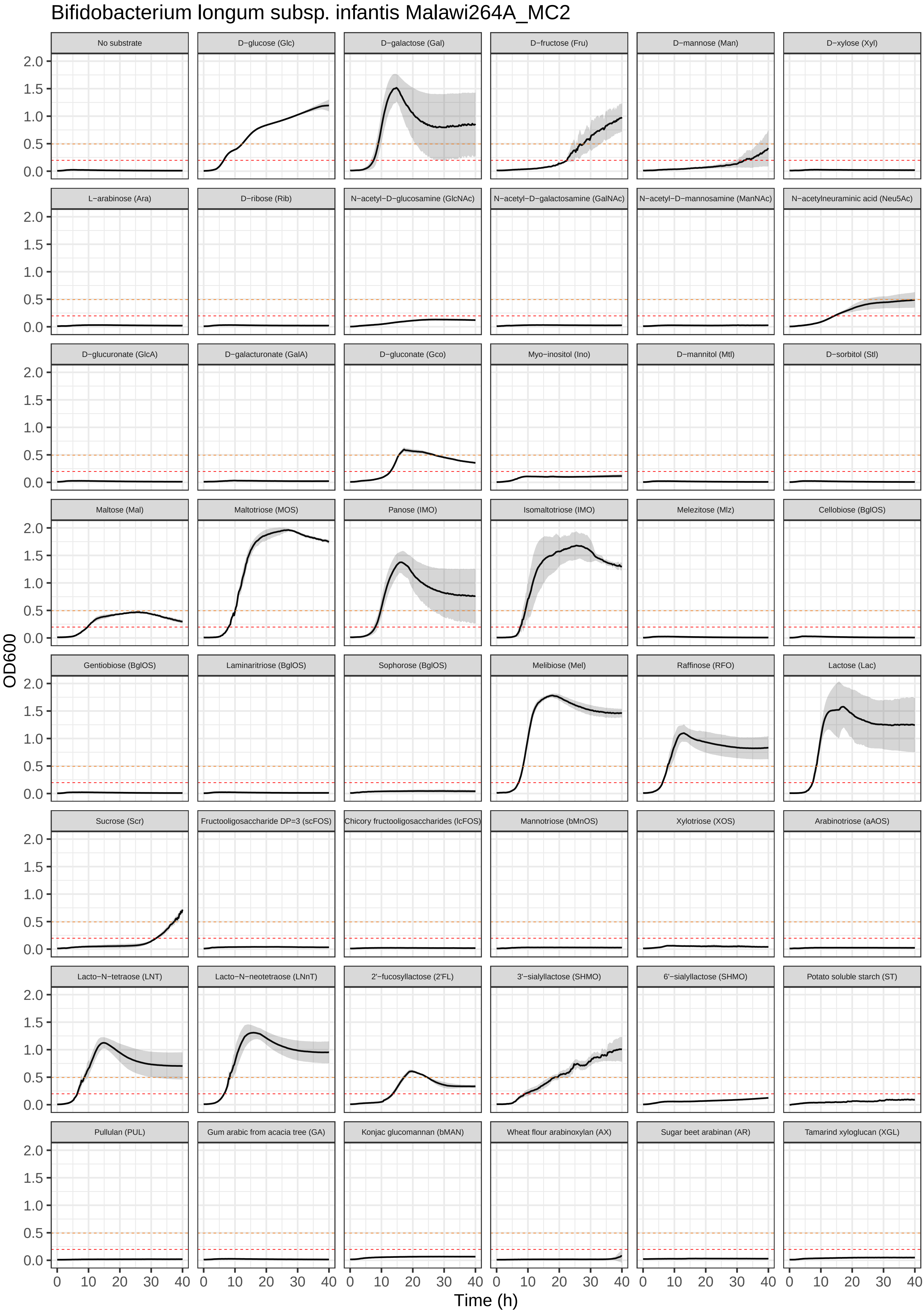

Bifidobacterium longum subsp. infantis Malawi264A_MC2
No substrate
D−glucose (Glc)
D−galactose (Gal)
D−fructose (Fru)
D−mannose (Man)
D−xylose (Xyl)
L−arabinose (Ara)
D−ribose (Rib)
N−acetyl−D−glucosamine (GlcNAc)
N−acetyl−D−galactosamine (GalNAc)
N−acetyl−D−mannosamine (ManNAc)
N−acetylneuraminic acid (Neu5Ac)
D−glucuronate (GlcA)
D−galacturonate (GalA)
D−gluconate (Gco)
Myo−inositol (Ino)
D−mannitol (Mtl)
D−sorbitol (Stl)
Maltose (Mal)
Maltotriose (MOS)
Panose (IMO)
Isomaltotriose (IMO)
Melezitose (Mlz)
Cellobiose (BglOS)
Gentiobiose (BglOS)
Laminaritriose (BglOS)
Sophorose (BglOS)
Melibiose (Mel)
Raffinose (RFO)
Lactose (Lac)
Sucrose (Scr)
Fructooligosaccharide DP=3 (scFOS)
Chicory fructooligosaccharides (lcFOS)
Mannotriose (bMnOS)
Xylotriose (XOS)
Arabinotriose (aAOS)
Lacto−N−tetraose (LNT)
Lacto−N−neotetraose (LNnT)
2'−fucosyllactose (2'FL)
3'−sialyllactose (SHMO)
6'−sialyllactose (SHMO)
Potato soluble starch (ST)
Pullulan (PUL)
Gum arabic from acacia tree (GA)
Konjac glucomannan (bMAN)
Wheat flour arabinoxylan (AX)
Sugar beet arabinan (AR)
Tamarind xyloglucan (XGL)
OD600
0.0
0.5
1.0
1.5
2.0
0
10
20
30
40
Time (h)

Bifidobacterium longum subsp. longum Bg115.S08_3A11

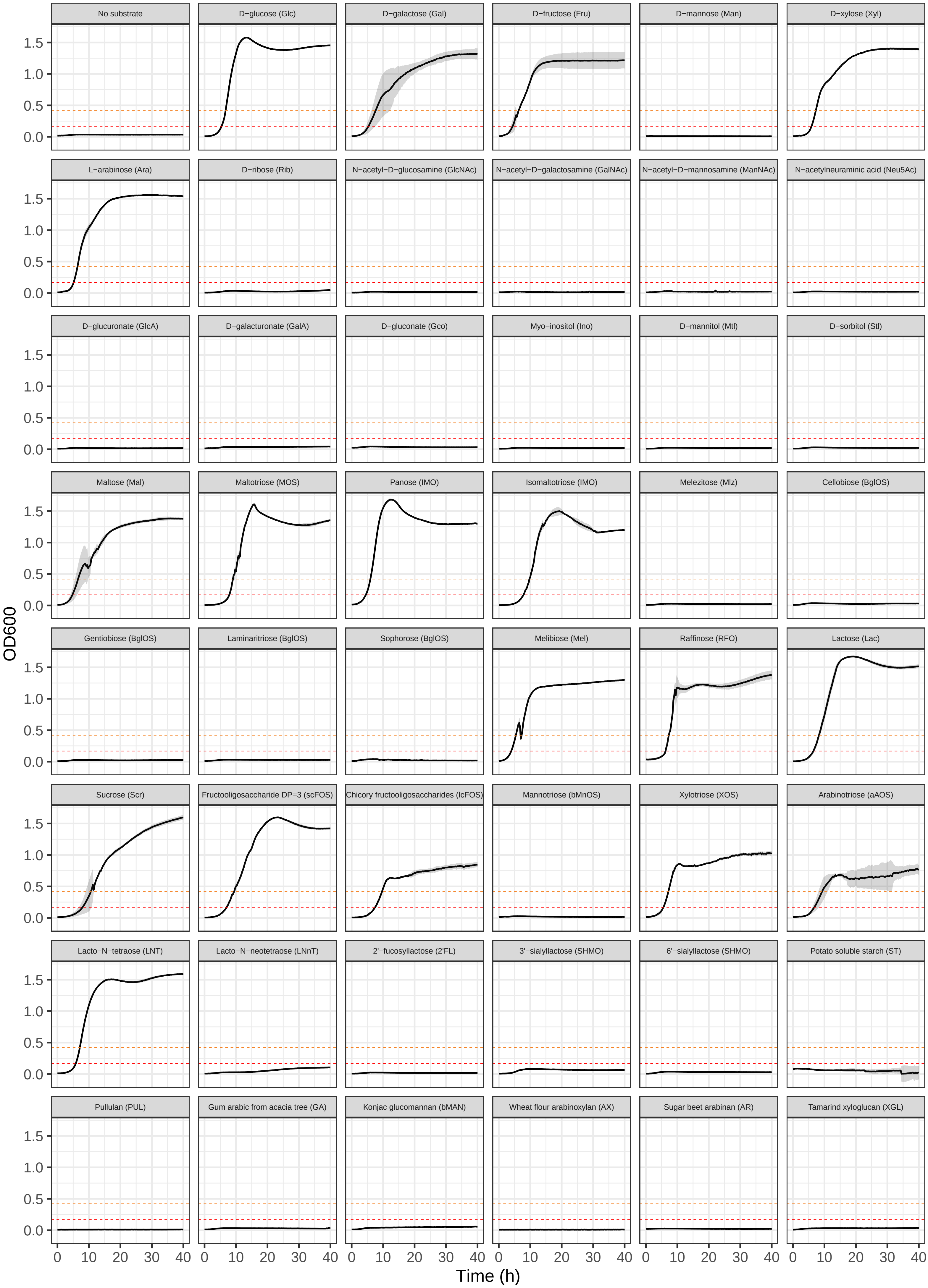

### Bifidobacterium longum subsp. suis Bg131.S11_17.F6

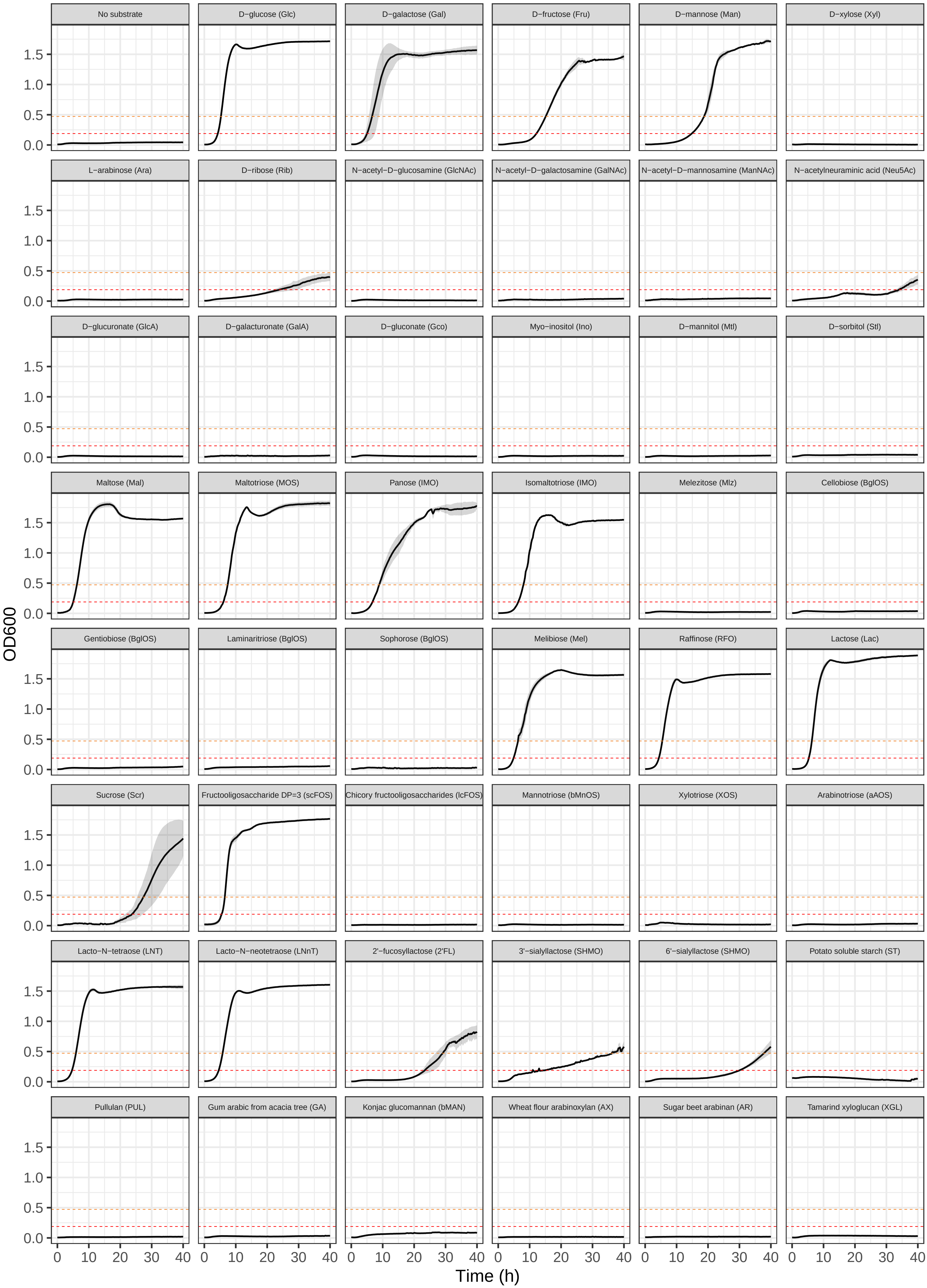

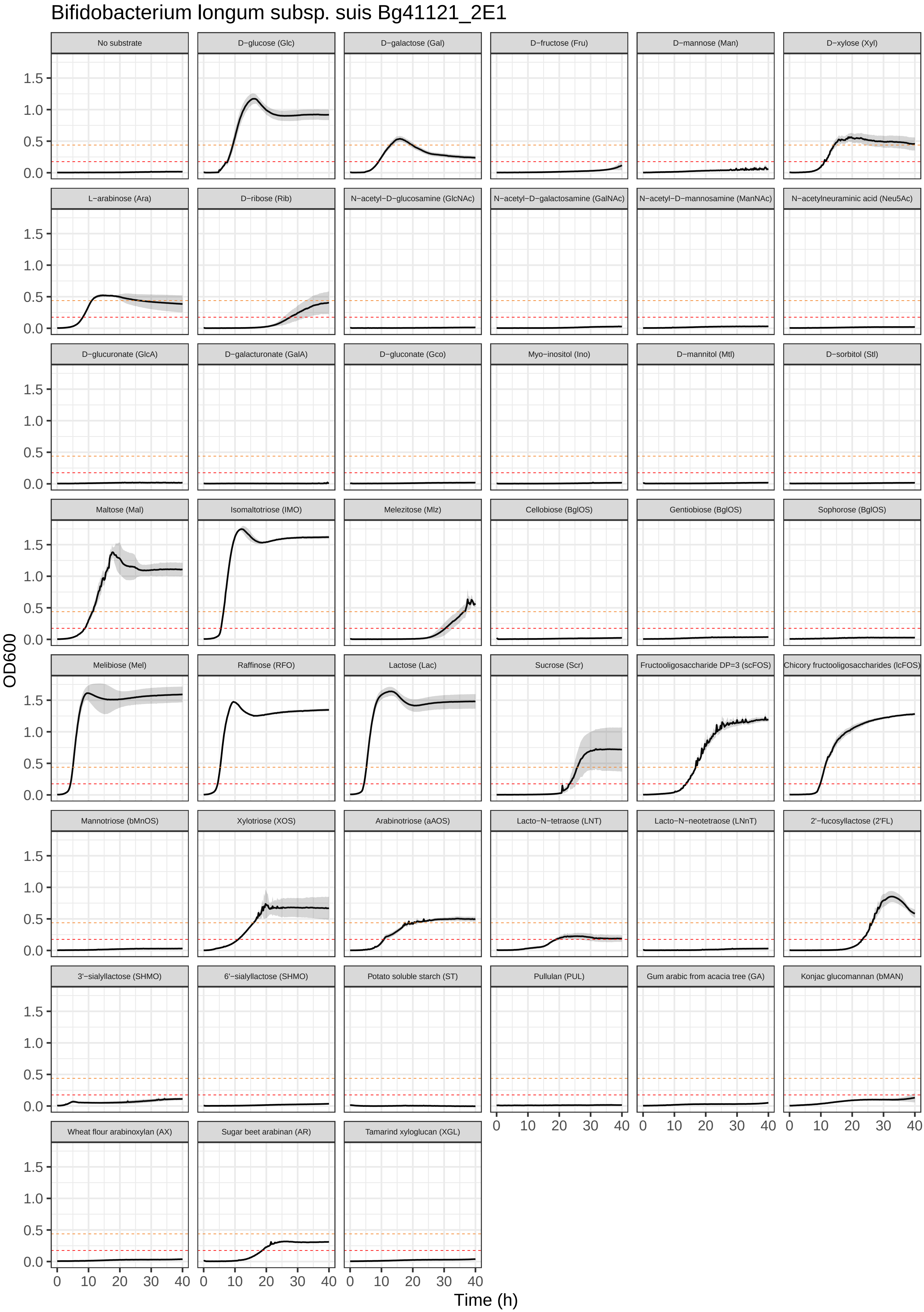

Bifidobacterium longum subsp. suis Bg41121_2E1
No substrate
D−glucose (Glc)
D−galactose (Gal)
D−fructose (Fru)
D−mannose (Man)
D−xylose (Xyl)
L−arabinose (Ara)
D−ribose (Rib)
N−acetyl−D−glucosamine (GlcNAc)
N−acetyl−D−galactosamine (GalNAc)
N−acetyl−D−mannosamine (ManNAc)
N−acetylneuraminic acid (Neu5Ac)
D−glucuronate (GlcA)
D−galacturonate (GalA)
D−gluconate (Gco)
Myo−inositol (Ino)
D−mannitol (Mtl)
D−sorbitol (Stl)
Maltose (Mal)
Isomaltotriose (IMO)
Melezitose (Mlz)
Cellobiose (BglOS)
Gentiobiose (BglOS)
Sophorose (BglOS)
Melibiose (Mel)
Raffinose (RFO)
Lactose (Lac)
Sucrose (Scr)
Fructooligosaccharide DP=3 (scFOS)
Chicory fructooligosaccharides (lcFOS)
Mannotriose (bMnOS)
Xylotriose (XOS)
Arabinotriose (aAOS)
Lacto−N−tetraose (LNT)
Lacto−N−neotetraose (LNnT)
2'−fucosyllactose (2'FL)
3'−sialyllactose (SHMO)
6'−sialyllactose (SHMO)
Potato soluble starch (ST)
Pullulan (PUL)
Gum arabic from acacia tree (GA)
Konjac glucomannan (bMAN)
Wheat flour arabinoxylan (AX)
Sugar beet arabinan (AR)
Tamarind xyloglucan (XGL)
OD600
0.0
0.5
1.0
1.5
0
10
20
30
40
Time (h)

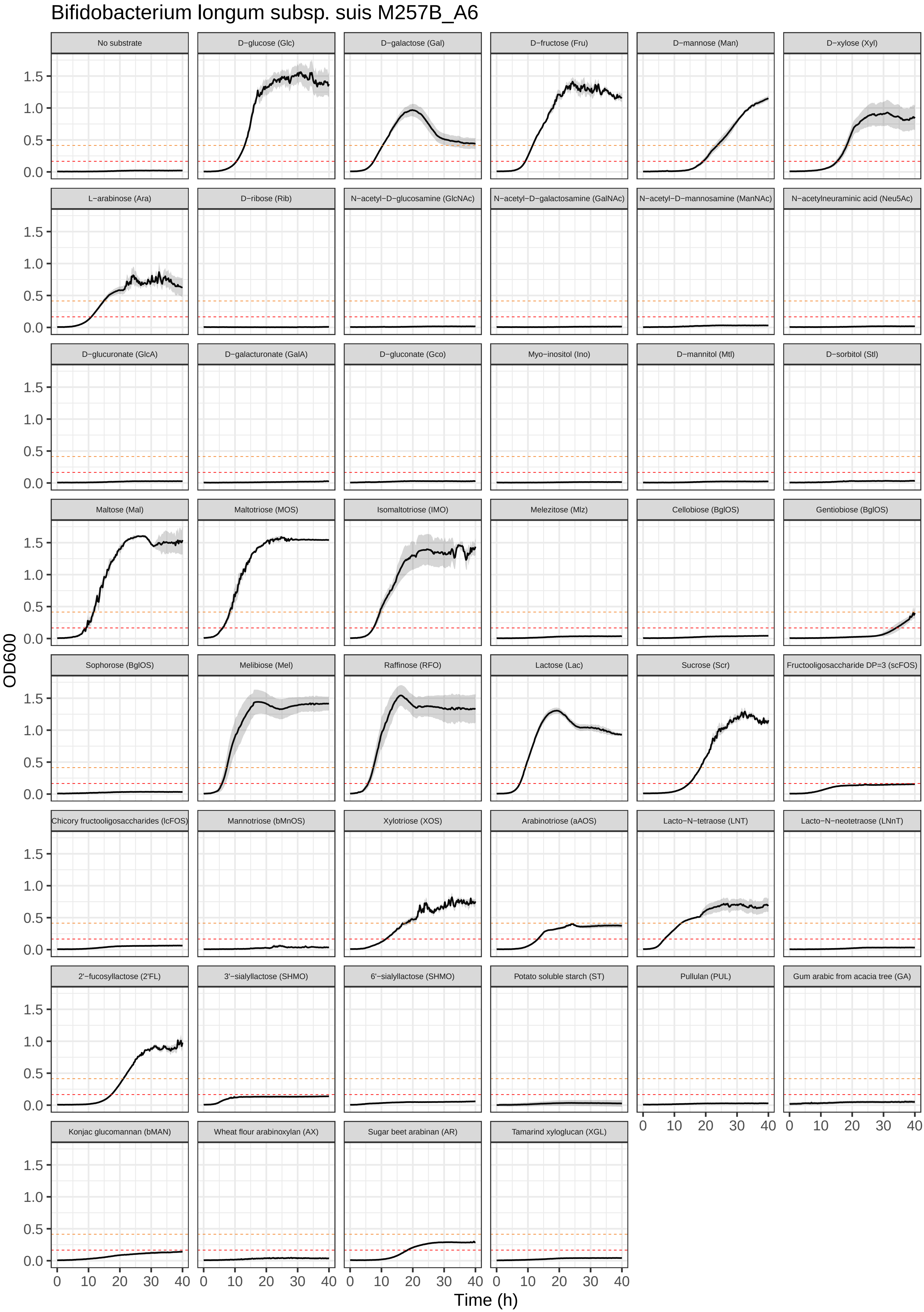
Bifidobacterium longum subsp. suis M257B_A6
No substrate
D−glucose (Glc)
D−galactose (Gal)
D−fructose (Fru)
D−mannose (Man)
D−xylose (Xyl)
L−arabinose (Ara)
D−ribose (Rib)
N−acetyl−D−glucosamine (GlcNAc)
N−acetyl−D−galactosamine (GalNAc)
N−acetyl−D−mannosamine (ManNAc)
N−acetylneuraminic acid (Neu5Ac)
D−glucuronate (GlcA)
D−galacturonate (GalA)
D−gluconate (Gco)
Myo−inositol (Ino)
D−mannitol (Mtl)
D−sorbitol (Stl)
Maltose (Mal)
Maltotriose (MOS)
Isomaltotriose (IMO)
Melezitose (Mlz)
Cellobiose (BglOS)
Gentiobiose (BglOS)
Sophorose (BglOS)
Melibiose (Mel)
Raffinose (RFO)
Lactose (Lac)
Sucrose (Scr)
Fructooligosaccharide DP=3 (scFOS)
Chicory fructooligosaccharides (lcFOS)
Mannotriose (bMnOS)
Xylotriose (XOS)
Arabinotriose (aAOS)
Lacto−N−tetraose (LNT)
Lacto−N−neotetraose (LNnT)
2'−fucosyllactose (2'FL)
3'−sialyllactose (SHMO)
6'−sialyllactose (SHMO)
Potato soluble starch (ST)
Pullulan (PUL)
Gum arabic from acacia tree (GA)
Konjac glucomannan (bMAN)
Wheat flour arabinoxylan (AX)
Sugar beet arabinan (AR)
Tamarind xyloglucan (XGL)
OD600
Time (h)
0.0
0.5
1.0
1.5
0
10
20
30
40

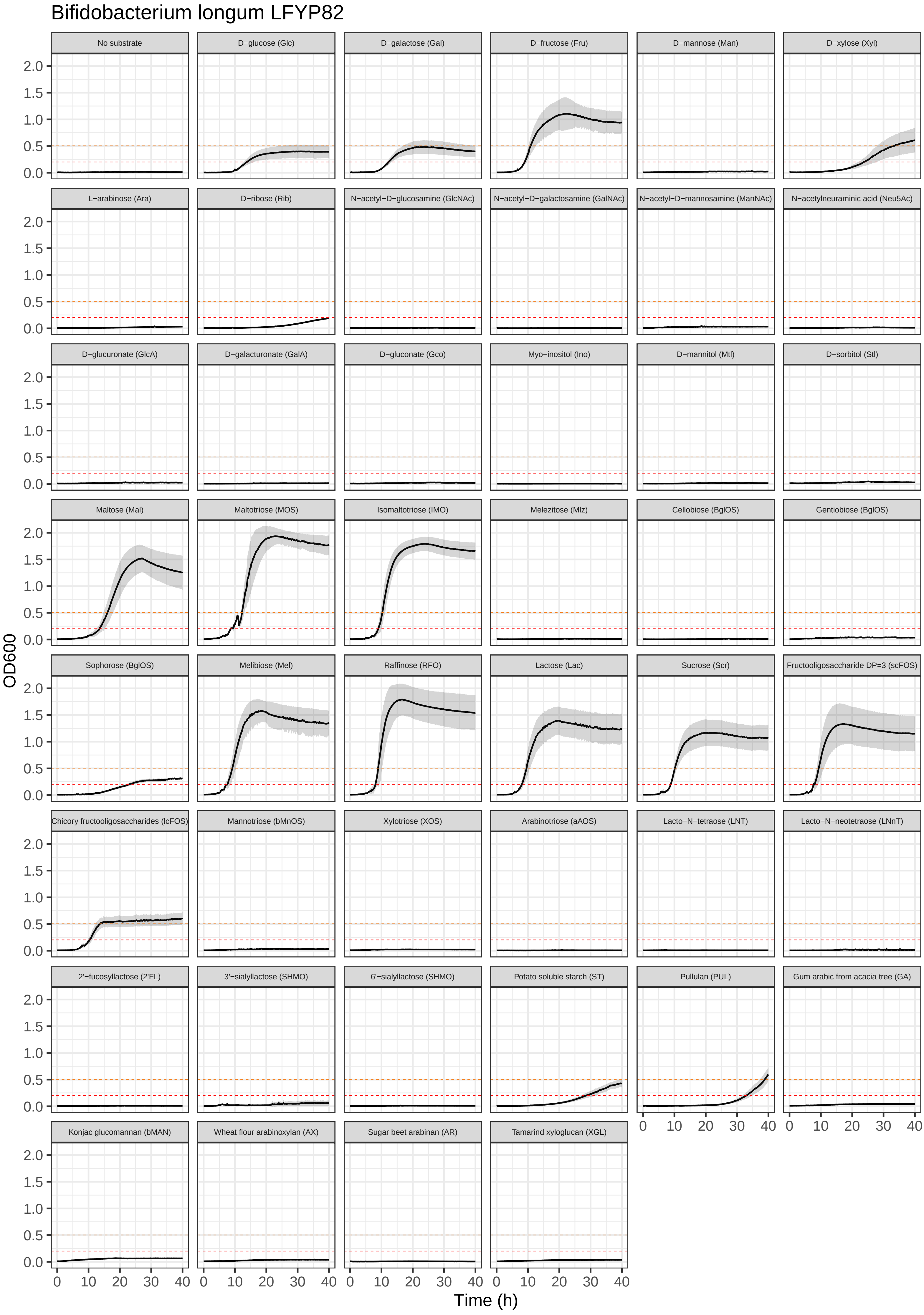

Bifidobacterium longum LFYP82
No substrate
D−glucose (Glc)
D−galactose (Gal)
D−fructose (Fru)
D−mannose (Man)
D−xylose (Xyl)
L−arabinose (Ara)
D−ribose (Rib)
N−acetyl−D−glucosamine (GlcNAc)
N−acetyl−D−galactosamine (GalNAc)
N−acetyl−D−mannosamine (ManNAc)
N−acetylneuraminic acid (Neu5Ac)
D−glucuronate (GlcA)
D−galacturonate (GalA)
D−gluconate (Gco)
Myo−inositol (Ino)
D−mannitol (Mtl)
D−sorbitol (Stl)
Maltose (Mal)
Maltotriose (MOS)
Isomaltotriose (IMO)
Melezitose (Mlz)
Cellobiose (BglOS)
Gentiobiose (BglOS)
Sophorose (BglOS)
Melibiose (Mel)
Raffinose (RFO)
Lactose (Lac)
Sucrose (Scr)
Fructooligosaccharide DP=3 (scFOS)
Chicory fructooligosaccharides (lcFOS)
Mannotriose (bMnOS)
Xylotriose (XOS)
Arabinotriose (aAOS)
Lacto−N−tetraose (LNT)
Lacto−N−neotetraose (LNnT)
2'−fucosyllactose (2'FL)
3'−sialyllactose (SHMO)
6'−sialyllactose (SHMO)
Potato soluble starch (ST)
Pullulan (PUL)
Gum arabic from acacia tree (GA)
Konjac glucomannan (bMAN)
Wheat flour arabinoxylan (AX)
Sugar beet arabinan (AR)
Tamarind xyloglucan (XGL)
OD600
0.0
0.5
1.0
1.5
2.0
0
10
20
30
40
Time (h)

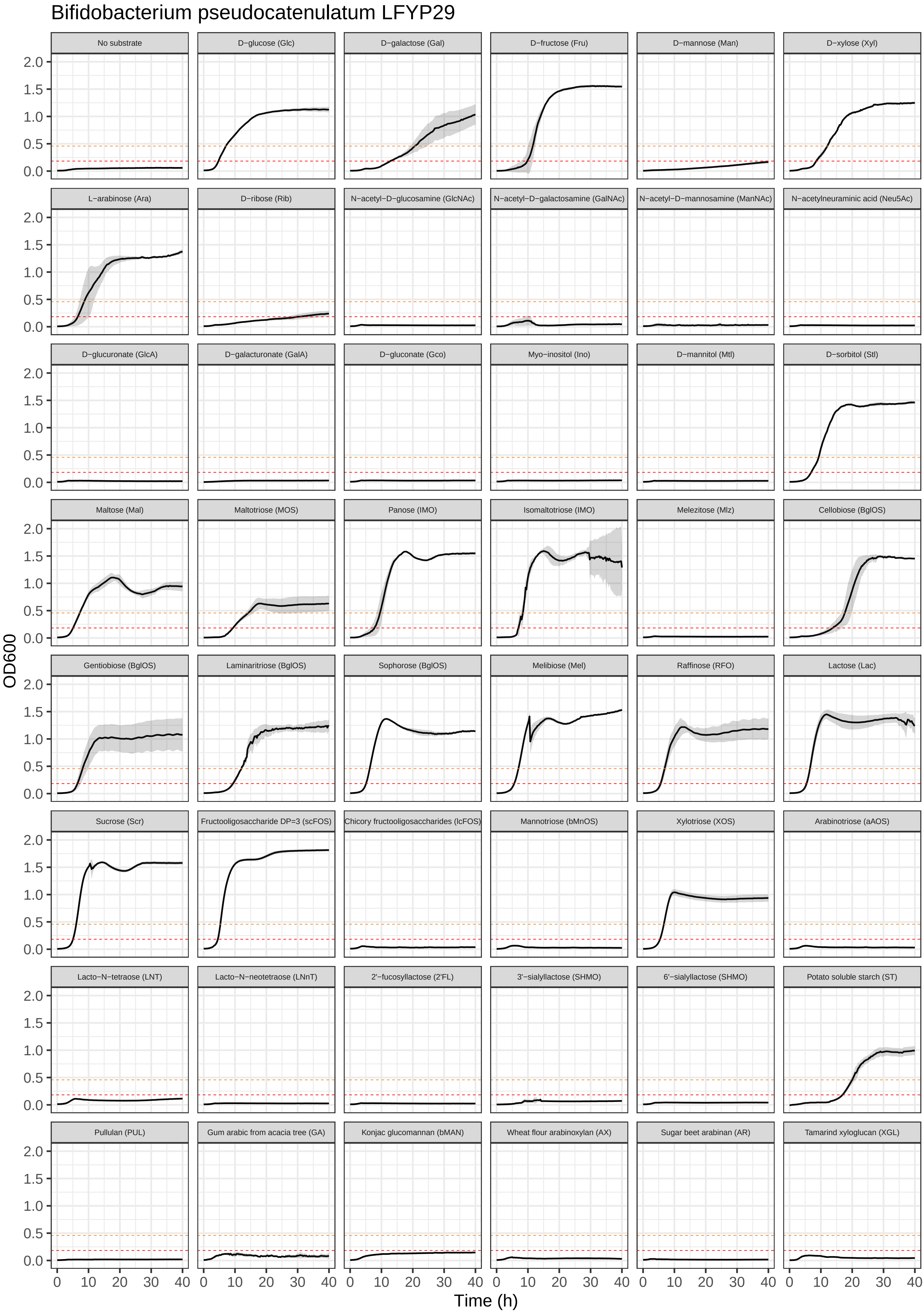
Bifidobacterium pseudocatenulatum LFYP29
No substrate
D−glucose (Glc)
D−galactose (Gal)
D−fructose (Fru)
D−mannose (Man)
D−xylose (Xyl)
L−arabinose (Ara)
D−ribose (Rib)
N−acetyl−D−glucosamine (GlcNAc)
N−acetyl−D−galactosamine (GalNAc)
N−acetyl−D−mannosamine (ManNAc)
N−acetylneuraminic acid (Neu5Ac)
D−glucuronate (GlcA)
D−galacturonate (GalA)
D−gluconate (Gco)
Myo−inositol (Ino)
D−mannitol (Mtl)
D−sorbitol (Stl)
Maltose (Mal)
Maltotriose (MOS)
Panose (IMO)
Isomaltotriose (IMO)
Melezitose (Mlz)
Cellobiose (BglOS)
Gentiobiose (BglOS)
Laminaritriose (BglOS)
Sophorose (BglOS)
Melibiose (Mel)
Raffinose (RFO)
Lactose (Lac)
Sucrose (Scr)
Fructooligosaccharide DP=3 (scFOS)
Chicory fructooligosaccharides (lcFOS)
Mannotriose (bMnOS)
Xylotriose (XOS)
Arabinotriose (aAOS)
Lacto−N−tetraose (LNT)
Lacto−N−neotetraose (LNnT)
2'−fucosyllactose (2'FL)
3'−sialyllactose (SHMO)
6'−sialyllactose (SHMO)
Potato soluble starch (ST)
Pullulan (PUL)
Gum arabic from acacia tree (GA)
Konjac glucomannan (bMAN)
Wheat flour arabinoxylan (AX)
Sugar beet arabinan (AR)
Tamarind xyloglucan (XGL)
OD600
0.0
0.5
1.0
1.5
2.0
Time (h)
0
10
20
30
40

Bifidobacterium pseudocatenulatum M26A_2F1

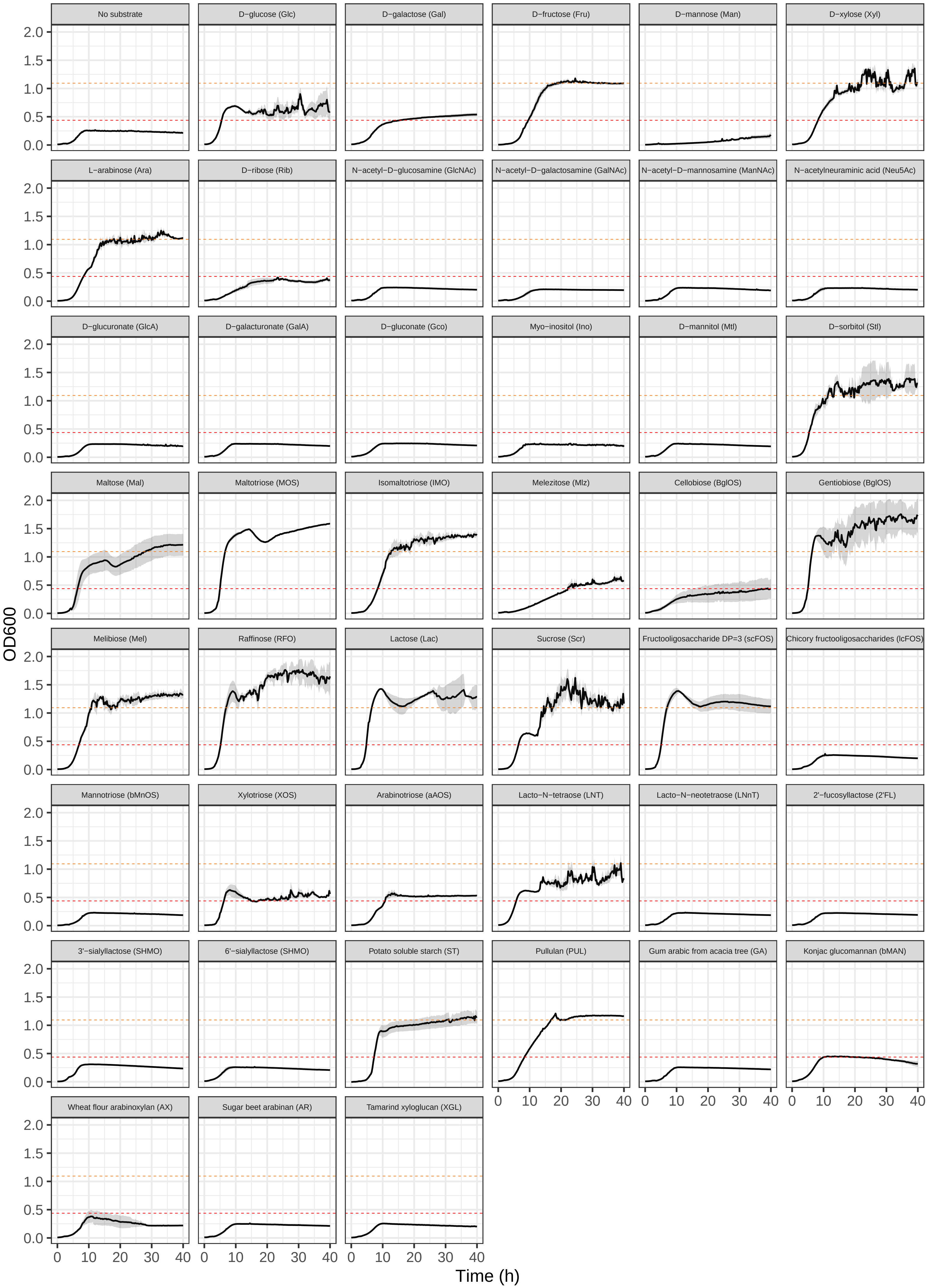

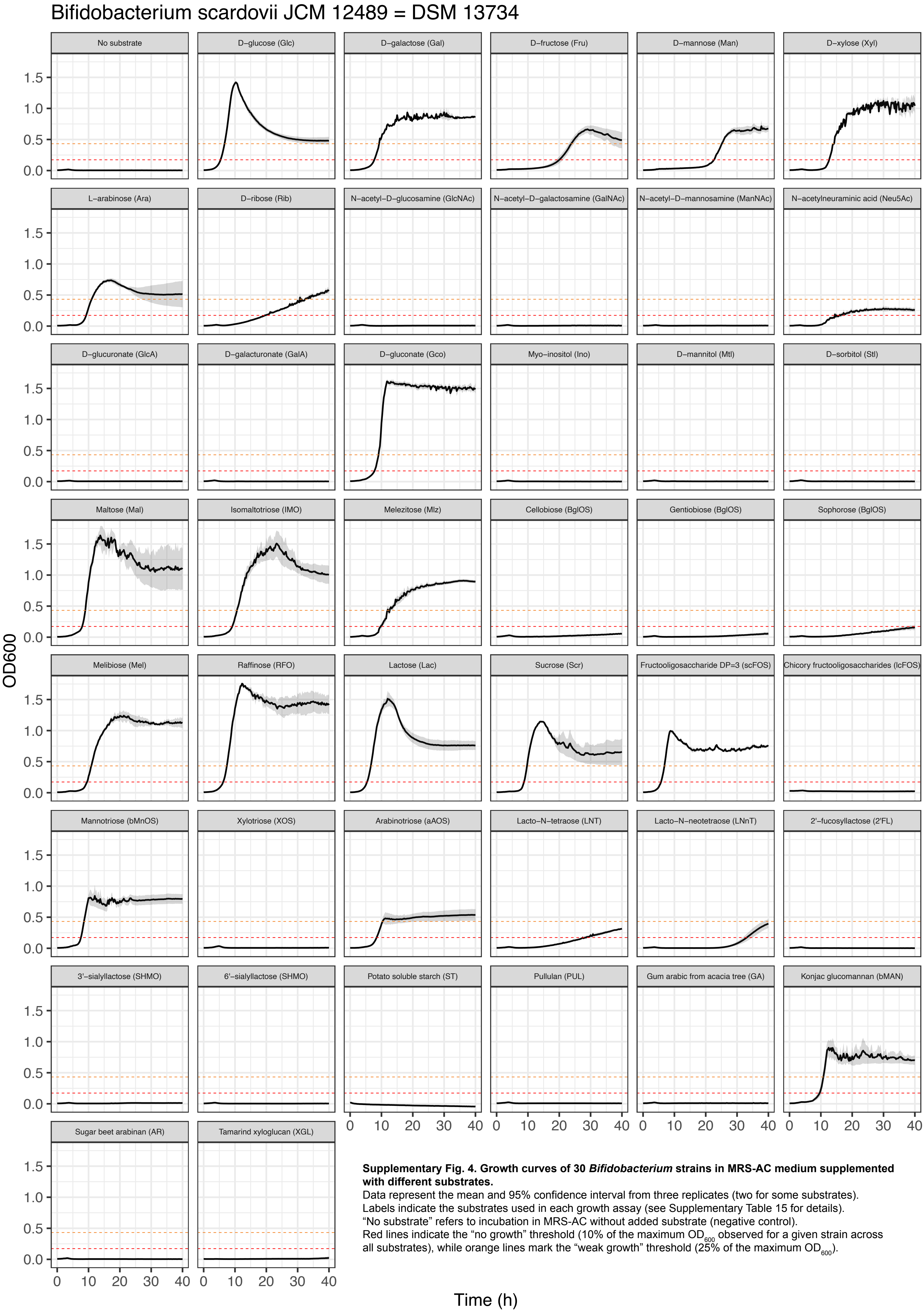


**Supplementary Fig. 4. Growth curves of 30 *Bifidobacterium* strains in MRS-AC medium supplemented with different substrates.**
Data represent the mean and 95% confidence interval from three replicates (two for some substrates).
Labels indicate the substrates used in each growth assay (see Supplementary Table 15 for details).
"No substrate" refers to incubation in MRS-AC without added substrate (negative control).
Red lines indicate the "no growth" threshold (10% of the maximum $OD_{600}$ observed for a given strain across all substrates), while orange lines mark the "weak growth" threshold (25% of the maximum $OD_{600}$).

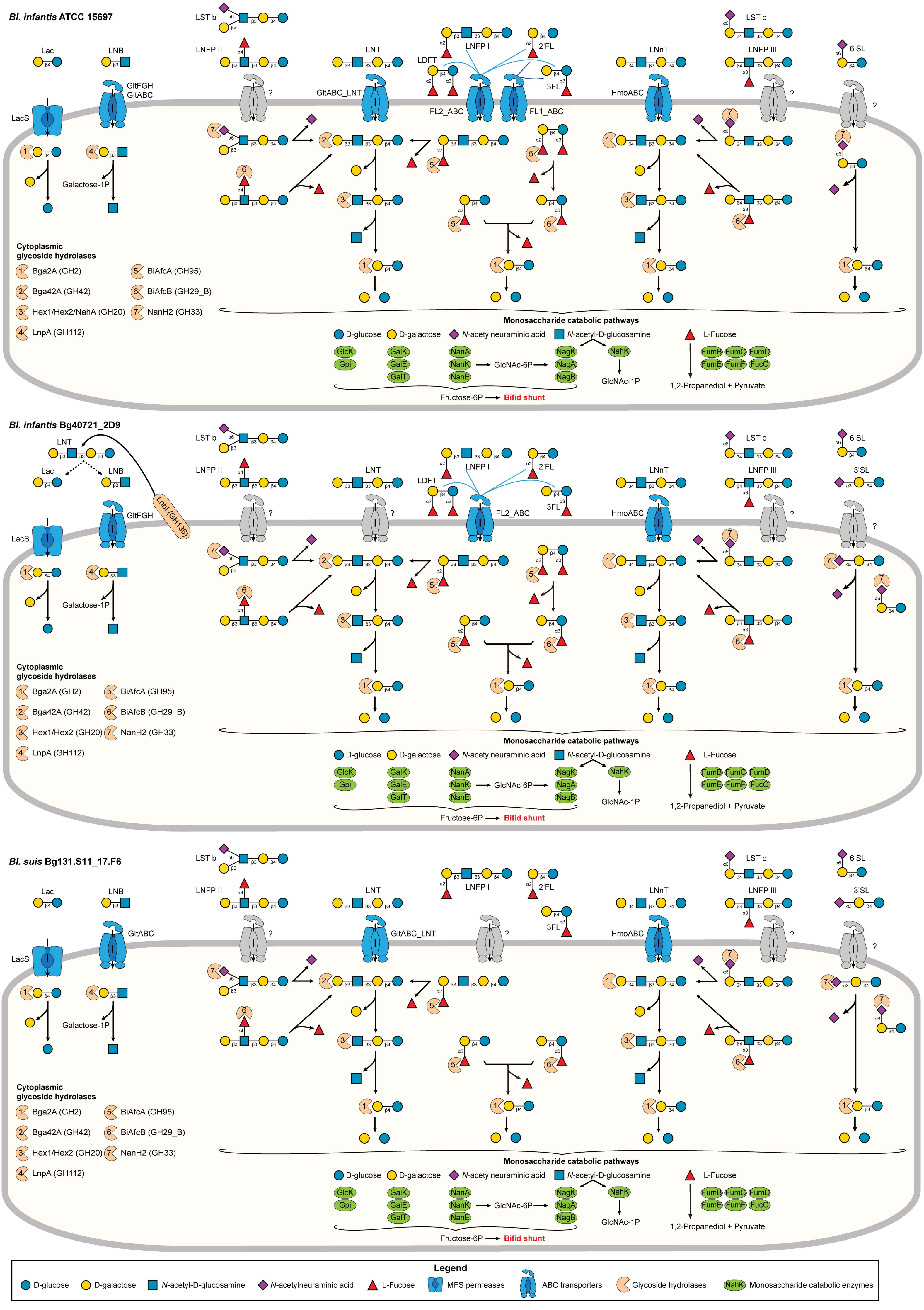

Bl. infantis ATCC 15697
LST b
LNFP II
Lac
LNB
LNT
LNFP I
2'FL
LDFT
3FL
LNnT
LST c
LNFP III
6'SL
GltFGH
GltABC
LacS
GltABC_LNT
FL2_ABC
FL1_ABC
HmoABC
?
Galactose-1P
Cytoplasmic glycoside hydrolases
1 Bga2A (GH2)
2 Bga42A (GH42)
3 Hex1/Hex2/NahA (GH20)
4 LnpA (GH112)
5 BiAfcA (GH95)
6 BiAfcB (GH29_B)
7 NanH2 (GH33)
Monosaccharide catabolic pathways
D-glucose
D-galactose
N-acetylneuraminic acid
N-acetyl-D-glucosamine
L-Fucose
GlcK
Gpi
GalK
GalE
GalT
NanA
NanK
NanE
GlcNAc-6P
NagK
NagA
NagB
NahK
GlcNAc-1P
FumB
FumC
FumD
FumE
FumF
FucO
1,2-Propanediol + Pyruvate
Fructose-6P
Bifid shunt
Bl. infantis Bg40721_2D9
LNT
Lac
LNB
LnbI (GH136)
LST b
LNFP II
LNFP I
2'FL
LDFT
3FL
LNnT
LST c
LNFP III
6'SL
3'SL
LacS
GltFGH
FL2_ABC
HmoABC
?
Galactose-1P
Cytoplasmic glycoside hydrolases
1 Bga2A (GH2)
2 Bga42A (GH42)
3 Hex1/Hex2 (GH20)
4 LnpA (GH112)
5 BiAfcA (GH95)
6 BiAfcB (GH29_B)
7 NanH2 (GH33)
Monosaccharide catabolic pathways
D-glucose
D-galactose
N-acetylneuraminic acid
N-acetyl-D-glucosamine
L-Fucose
GlcK
Gpi
GalK
GalE
GalT
NanA
NanK
NanE
GlcNAc-6P
NagK
NagA
NagB
NahK
GlcNAc-1P
FumB
FumC
FumD
FumE
FumF
FucO
1,2-Propanediol + Pyruvate
Fructose-6P
Bifid shunt
Bl. suis Bg131.S11_17.F6
LST b
LNFP II
Lac
LNB
LNT
LNFP I
2'FL
3FL
LNnT
LST c
LNFP III
6'SL
3'SL
LacS
GltABC
GltABC_LNT
HmoABC
?
Galactose-1P
Cytoplasmic glycoside hydrolases
1 Bga2A (GH2)
2 Bga42A (GH42)
3 Hex1/Hex2 (GH20)
4 LnpA (GH112)
5 BiAfcA (GH95)
6 BiAfcB (GH29_B)
7 NanH2 (GH33)
Monosaccharide catabolic pathways
D-glucose
D-galactose
N-acetylneuraminic acid
N-acetyl-D-glucosamine
L-Fucose
GlcK
Gpi
GalK
GalE
GalT
NanA
NanK
NanE
GlcNAc-6P
NagK
NagA
NagB
NahK
GlcNAc-1P
FumB
FumC
FumD
FumE
FumF
FucO
1,2-Propanediol + Pyruvate
Fructose-6P
Bifid shunt
Legend
D-glucose
D-galactose
N-acetyl-D-glucosamine
N-acetylneuraminic acid
L-Fucose
MFS permeases
ABC transporters
Glycoside hydrolases
NahK
Monosaccharide catabolic enzymes

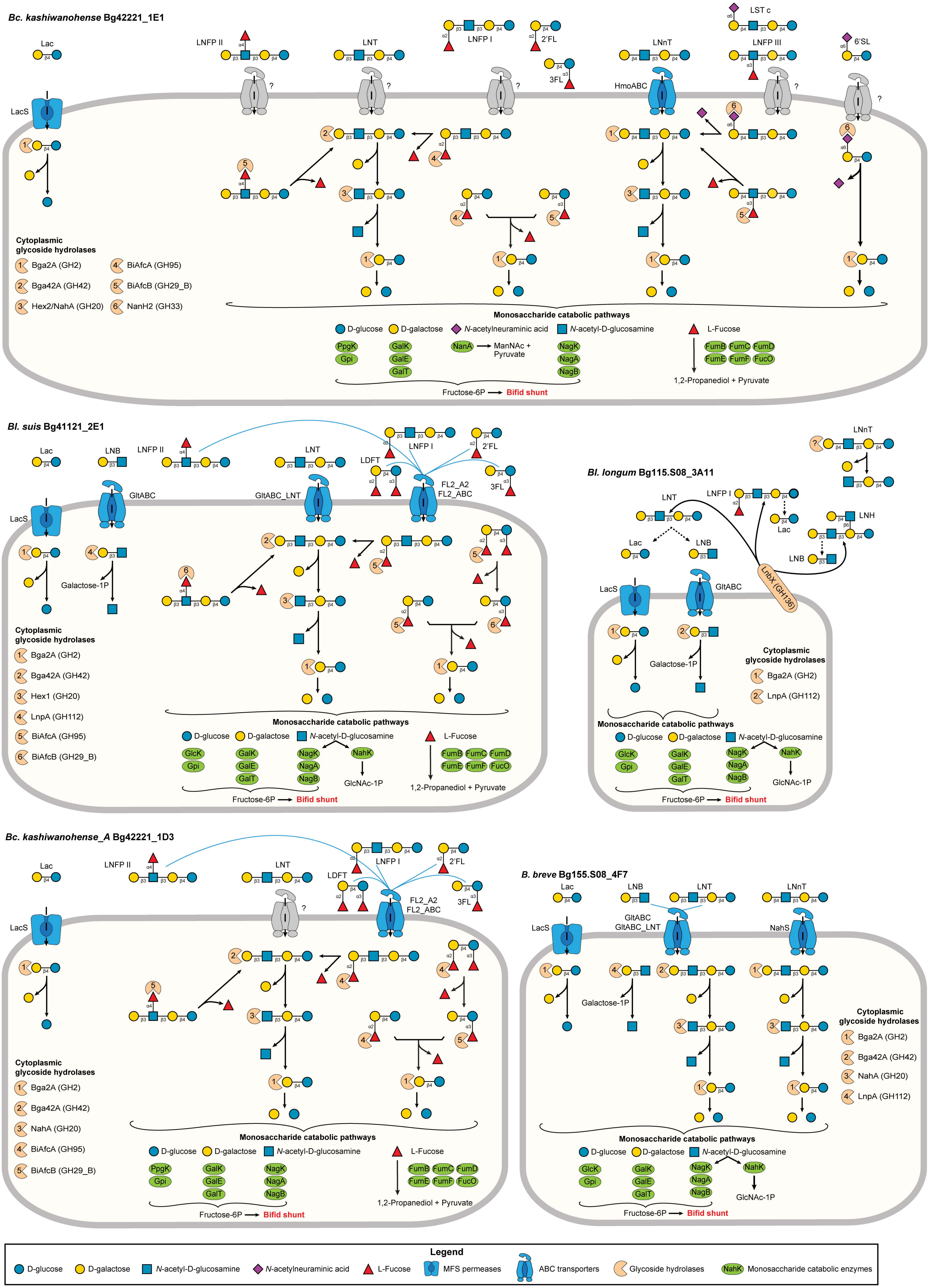


**Supplementary Fig. 5. Reconstructed human milk oligosaccharide (HMO) utilization pathways in *Bifidobacterium* strains.**
Pathways were inferred based on the integration of genomic predictions with growth phenotypes and HMO glycoprofiling data.
