## Supplementary Code File 1 for "Integrative genomic reconstruction reveals heterogeneity in carbohydrate utilization across human gut bifidobacteria"

#### Contents

**1 Background** **4**

**2 Reproducibility and accessibility** **5**

**3 R packages and external functions** **5**

**4 Summary of genomes and functional roles** **6**

9.4 Hierarchical clustering of the BPM (other pathways) of 28 *Bifidobacterium longum* genomes . 39

**10 Representation of predicted metabolic pathways in 3083 *Bifidobacterium* genomes** **41**

11.31.3 Growth in MRS-AC supplemented with 0.5% mannotriose or 0.5% konjac glucomannan 149
11.31.4 Growth in MRS-AC supplemented with 1% D-glucuronic acid sodium salt . . . . . 151
11.31.5 Growth in MRS-AC supplemented with 1% HMOs from pooled human milk (pHMOs) 152

**14 Session info 181**

### 1 Background

**This supplementary code file describes**:

1. Summary of reference genomes and functional roles
2. Phylogenetic analysis of reference *Bifidobacterium* genomes
3. Comparison of gene annotations
4. Analysis of CAZyme representation in 263 *Bifidobacterium* strains
5. Representation of predicted carbohydrate utilization phenotypes in 263 reference *Bifidobacterium* strains
6. Representation of predicted metabolic pathways in 28 *Bifidobacterium longum* genomes

7. Representation of predicted metabolic pathways in 3083 *Bifidobacterium* genomes
8. Analysis of *in vitro* growth data
9. Analysis of human milk oligosaccharide (HMO) consumption data
10. Analysis of RNA-seq data

---

#### 2 Reproducibility and accessibility

All code used in this analysis (including the Rmarkdown document used to compile this supplementary code file) is available on GitHub **here**. Once the GitHub repo has been downloaded, navigate to `compendium_manuscript/` to find the Rmarkdown document as well as the RProject file. This should be your working directory for executing code.

1. To fully reproduce the phylogenetic analysis of 263 reference *Bifidobacterium* genomes, you will need to download FNA files from **Figshare**. Downloaded FNA files should be placed in `data/genomes/263_NR_ref_genomes/fna/`
2. To fully reproduce the analysis of the CAZyme representation in 263 reference *Bifidobacterium* genomes, you will need to predict CDS using tools like **Prodigal** or **Prokka**. The resulting FAA files should be placed in `data/genomes/263_NR_ref_genomes/faa/`
3. To fully reproduce the RNA-seq data analysis, you will need to download raw FASTQ files from Gene Expression Omnibus under accession **GSE239955**. Downloaded FASTQ files should be placed in `data/rnaseq/fastq/`. Otherwise, `data/rnaseq/kallisto/` already contains Kallisto mapping outputs
4. The pathway prediction pipeline **glycobif** used for additional 2820 isolate genomes and MAGs is available in another GitHub **repository**

---

#### 3 R packages and external functions

A set of R packages was used for this analysis. The **pacman** package was used to simplify downloading and loading the required packages. All graphics and data wrangling were handled using the **tidyverse suite of packages**.

```
# install or load the pacman package for the rapid installation of required packages
if (!require("pacman")) install.packages("pacman")
# use pacman to install or load all required packages
pacman::p_load("tidyverse", "readxl", "patchwork", "ComplexHeatmap", "ggbeeswarm", "ggrepel",
               "circlize", "ggpubr", "tximport", "rhdf5", "gt", "edgeR", "cowplot", "limma",
               "vegan", "car", "AER", "emmeans", "gcplyr", "ape", "phytools", "dendextend",
               "ggfortify", "ggordiplots")
set.seed(1992)
```

A set of external R functions was used to keep the code tidy. All used R scripts with functions can be found in `compendium_manuscript/code/`.

```r
source("code/growth_curve_plotter.R") # plots growth curves
source("code/calculate_statistics.R") # calculates the number of true positives (TP),
# true negatives (TN), false positives (FP), false negatives (FN)
# by comparing predicted binary carbohydrate utilization phenotypes ("1" and "0")
# with growth phenotypes ("+/w" and "-")
source("code/profile.R") # calculates counts per million (CPM) for each gene and
# plots the distribution of CPM values for each sample
source("code/deg_list.R") # selects differentially expressed genes (DEGs) based on input cut-offs
# outputs an annotated table with DEGs to a file
```

---

### 4 Summary of genomes and functional roles

#### 4.1 Introduction

This block contains code used for summarizing information about (i) 3083 non-redundant *Bifidobacterium* genomes and (ii) 589 functional metabolic roles.

#### 4.2 Load data

```r
# read table with data on reference genomes
info_ref.df <- read_tsv("data/tables/BPM_263_NR_genomes_carbs.txt") %>%
  dplyr::select(genome_ID, curated_taxonomy, continent, country)
# read table with data on additional genomes
info_add.df <- read_tsv("data/tables/BPM_2820_NR_genomes_carbs.txt") %>%
  dplyr::select(genome_ID, curated_taxonomy, continent, country)
# read table with functional roles
fr.df <- read_tsv("data/tables/functional_roles.txt")
```

#### 4.3 Table with summary of 3083 *Bifidobacterium* genomes

Here, we summarize information about 3083 genomes (isolate and assembled from metagenomes) of human-colonizing bifidobacteria.

First, the number of genomes per species/subspecies.

```r
# summarize genome counts per curated_taxonomy
ref_summary_tax <- info_ref.df %>%
  group_by(curated_taxonomy) %>%
  summarise(ref_genomes = n(), .groups = "drop")
add_summary_tax <- info_add.df %>%
  group_by(curated_taxonomy) %>%
  summarise(add_genomes = n(), .groups = "drop")

# merge summaries
summary_table_tax <- full_join(ref_summary_tax, add_summary_tax, by = "curated_taxonomy") %>%
  replace_na(list(ref_genomes = 0, add_genomes = 0)) %>%
  mutate(total_genomes = ref_genomes + add_genomes)
```

| curated_taxonomy | ref_genomes | add_genomes | total_genomes |
|---|---|---|---|
| Bifidobacterium adolescentis | 22 | 480 | 502 |
| Bifidobacterium angulatum | 2 | 4 | 6 |
| Bifidobacterium animalis subsp. lactis | 4 | 8 | 12 |
| Bifidobacterium bifidum | 22 | 391 | 413 |
| Bifidobacterium breve | 31 | 276 | 307 |
| Bifidobacterium catenulatum subsp. catenulatum | 2 | 51 | 53 |
| Bifidobacterium catenulatum subsp. kashiwanohense | 2 | 32 | 34 |
| Bifidobacterium catenulatum subsp. kashiwanohense_A | 3 | 20 | 23 |
| Bifidobacterium dentium | 4 | 102 | 106 |
| Bifidobacterium gallicum | 1 | 0 | 1 |
| Bifidobacterium globosum | 3 | 6 | 9 |
| Bifidobacterium hominis | 3 | 23 | 26 |
| Bifidobacterium longum subsp. infantis | 14 | 144 | 158 |
| Bifidobacterium longum subsp. longum | 115 | 846 | 961 |
| Bifidobacterium longum subsp. nov. | 3 | 31 | 34 |
| Bifidobacterium longum subsp. suis | 6 | 12 | 18 |
| Bifidobacterium pseudocatenulatum | 23 | 374 | 397 |
| Bifidobacterium scardovii | 2 | 8 | 10 |
| Bifidobacterium thermophilum | 1 | 0 | 1 |
| Bifidobacterium animalis subsp. animalis | 0 | 1 | 1 |
| Bifidobacterium pullorum | 0 | 5 | 5 |
| Bifidobacterium ruminantium | 0 | 4 | 4 |
| Bifidobacterium tsurumiense | 0 | 2 | 2 |
| TOTAL | 263 | 2820 | 3083 |

```r
# add a summary row at the bottom
summary_row_tax <- tibble(
  curated_taxonomy = "TOTAL",
  ref_genomes = sum(summary_table_tax$ref_genomes, na.rm = TRUE),
  add_genomes = sum(summary_table_tax$add_genomes, na.rm = TRUE),
  total_genomes = sum(summary_table_tax$total_genomes, na.rm = TRUE)
)

# combine the main summary table with the summary row
summary_table_tax <- bind_rows(summary_table_tax, summary_row_tax)

# print the summary table
gt(summary_table_tax)
```

Second, the number of genomes per region.

```r
# summarize genome counts per continent
ref_summary_cont <- info_ref.df %>%
  group_by(continent) %>%
  summarise(ref_genomes = n(), .groups = "drop")
```

| continent | ref_genomes | add_genomes | total_genomes |
|---|---|---|---|
| Africa | 5 | 94 | 99 |
| Asia | 130 | 798 | 928 |
| Europe | 95 | 1217 | 1312 |
| North_America | 32 | 367 | 399 |
| South_America | 1 | 13 | 14 |
| Oceania | 0 | 330 | 330 |
| NA | 0 | 1 | 1 |
| TOTAL | 263 | 2820 | 3083 |

```
add_summary_cont <- info_add.df %>%
  group_by(continent) %>%
  summarise(add_genomes = n(), .groups = "drop")

# merge summaries
summary_table_cont <- full_join(ref_summary_cont, add_summary_cont, by = "continent") %>%
  replace_na(list(ref_genomes = 0, add_genomes = 0)) %>%
  mutate(total_genomes = ref_genomes + add_genomes)

# add a summary row at the bottom
summary_row_cont <- tibble(
  continent = "TOTAL",
  ref_genomes = sum(summary_table_cont$ref_genomes, na.rm = TRUE),
  add_genomes = sum(summary_table_cont$add_genomes, na.rm = TRUE),
  total_genomes = sum(summary_table_cont$total_genomes, na.rm = TRUE)
)

# combine the main summary table with the summary row
summary_table_cont <- bind_rows(summary_table_cont, summary_row_cont)

# print the summary table
gt(summary_table_cont)
```

#### 4.4 Summary of functional roles

Here we calculate:

- The number of unique functional roles (total and stratified by (i) experimental evidence and (ii) type)
  - Total number of roles
  - Roles stratified by (i) experimental evidence and (ii) type
- The number of publications from which the data about functional roles were collected

```
# calculate the total number of unique functional roles
num_roles <- fr.df %>%
  distinct(functional_role_description) %>%
  nrow()
```

```r
# calculate the total number of unique functional roles characterized in bifidobacteria
num_characterized_roles <- fr.df %>%
  filter(evidence == "characterized_bifido") %>%
  distinct(functional_role_description) %>%
  nrow()
# calculate the total number of unique functional roles that have homology
# with roles characterized in other microbial taxa
num_homology_roles <- fr.df %>%
  filter(evidence == "homology_with_characterized") %>%
  distinct(functional_role_description) %>%
  nrow()
# calculate the total number of unique previosly predicted functional roles
num_predicted_before_roles <- fr.df %>%
  filter(evidence == "predicted_previously") %>%
  distinct(functional_role_description) %>%
  nrow()
# calculate the total number of unique novel predicted functional roles
num_predicted_new_roles <- fr.df %>%
  filter(evidence == "predicted_new") %>%
  distinct(functional_role_description) %>%
  nrow()

# stratify the functional roles by type
num_transporters <- fr.df %>%
  filter(functional_role_type == "transporter") %>%
  distinct(functional_role_description) %>%
  nrow()
num_donwstream <- fr.df %>%
  filter(functional_role_type == "downstream_catabolism") %>%
  distinct(functional_role_description) %>%
  nrow()
num_cazy <- fr.df %>%
  filter(functional_role_type == "CAZyme") %>%
  distinct(functional_role_description) %>%
  nrow()
num_reg <- fr.df %>%
  filter(functional_role_type == "regulator") %>%
  distinct(functional_role_description) %>%
  nrow()

# calculate the total number of publications
# split PMID values and extract unique values
unique_pmids <- unique(unlist(strsplit(fr.df$reference, ";")))
num_unique_pmids <- length(unique_pmids)

# print the counts
fr_output <- paste("Total number of functional roles:", num_roles, "\n",
                "Total number of roles characterized in bifidobacteria:",
                num_characterized_roles, "\n",
                "Total number of roles homologous that characterized in other taxa:",
                num_homology_roles, "\n",
                "Total number of roles predicted previosly:",
                num_predicted_before_roles - 1, "\n",
```

```
                 "Total number of functional roles predicted in this work:",
                 num_predicted_new_roles, "\n",
                 "Transporters and their components:", num_transporters, "\n",
                 "Downstream catabolic enzymes:", num_donwstream, "\n",
                 "CAZymes:", num_cazy, "\n",
                 "Transcriptional regulators:", num_reg, "\n",
                 "Total number of publications:", num_unique_pmids - 2, "\n")
cat(fr_output)
```

```
## Total number of functional roles: 589
##  Total number of roles characterized in bifidobacteria: 210
##  Total number of roles homologous that characterized in other taxa: 80
##  Total number of roles predicted previosly: 143
##  Total number of functional roles predicted in this work: 156
##  Transporters and their components: 235
##  Downstream catabolic enzymes: 72
##  CAZymes: 197
##  Transcriptional regulators: 85
##  Total number of publications: 222
```

---

### 5 Phylogenetic analysis of reference *Bifidobacterium* genomes

#### 5.1 Introduction

This block contains the code for building:

1. Phylogenetic tree of 263 reference *Bifidobacterium* genomes based on 487 core genes. The topology of the resulting tree was manually inspected to check (and correct if needed) taxonomic assignments of genomes based on their co-clustering with branches corresponding to the type or well-characterized strains of various *Bifidobacterium* taxa
2. Average Nucleotide Identity (ANI) matrices for select strains belonging to *Bifidobacterium longum* and *Bifidobacterium catenulatum* species

The following software is required:

1. **Prokka (v1.14.6)**
2. **Panaroo (v1.3.2)**
3. **CD-HIT (v4.8.1)**
4. **MAFFT (v7.515)**
5. **IQ-TREE (v2.2.0.3)**
6. **pyani (v0.2.12)**

To install these tools, you can use **mamba** and yml files in `envs` to create respective environments.

**Note**: installation via `mamba` was tested on macOS and may not always work on Linux-based operating systems. In the latter case, you may need to install the required software manually.

1. `mamba env create -f envs/prokka.yml` # Prokka (v1.14.6)

2. `mamba env create -f envs/panaroo.yml` # Panaroo (v1.3.2); CD-HIT (v4.8.1); MAFFT (v7.515)
3. `mamba env create -f envs/iqtree.yml` # IQ-TREE (v2.2.0.3)
4. `mamba env create -f envs/pyani.yml` # pyani (v0.2.12)

**Note**: FNA files could not be stored in the GitHub repo due to size limitations. Thus, you will need to download them from Figshare under **accession**. Put downloaded FNA files to `data/genomes/263_NR_ref_genomes/fna/`.

#### 5.2 Annotating genomes using Prokka

We used **Prokka** for annotating genomes. Prokka takes contig nucleotide fasta (FNA) and outputs annotated genomes in a standardized format (GFF3), which is recognized by many tools used to calculate pangenomes. The script that performs annotation can be found in `code/run_prokka.sh/`. A simple bash operation at the end of the script collects all created GFF3 files and puts them in the `data/genomes/263_NR_ref_genomes/gff/` folder.

```
source ~/.bash_profile
source code/run_prokka.sh
```

#### 5.3 Calculating pangenome using Panaroo

We used **Panaroo** to identify core genes shared among 263 reference *Bifidobacterium* genomes. Prokka-annotated GFF3 files were used as input. Since genomes of multiple different *Bifidobacterium* species were used, we relaxed the sequence identity threshold (`--threshold 0.8`) and length difference cutoff (`--len_dif_percent 0.9`). As part of the Panaroo pipeline, concatenated nucleotide sequences of 487 identified core genes were aligned via MAFFT.

```
source ~/.bash_profile
#set -ex

### SOFTWARE SETUP ##
####################
# required tools: panaroo=1.3.2; cd-hit=4.8.1; mafft=7.515
# set the name of the environment with installed tools
environment_name="panaroo"
# activate selected conda environment
eval "$(command conda 'shell.bash' 'hook' 2> /dev/null)" # initializes conda in sub-shell
conda activate ${environment_name}
conda info|egrep "conda version|active environment"

# run panaroo
mkdir -p data/pangenome
panaroo -i data/genomes/263_NR_ref_genomes/gff/*.gff \
-o data/pangenome/panaroo_strict_i80_l90 \
--clean-mode strict \
-a core --aligner mafft \
--threshold 0.8 \
--len_dif_percent 0.9 \
-t 16
```

#### 5.4 Building phylogenetic tree using IQ-TREE

We used a maximum-likelihood-based algorithm with ultrafast bootstrap approximation (UFBoot) implemented in **IQ-TREE** to build a phylogenetic tree based on the alignment of the core genes. Depending on your computing power, this process may take multiple days; you can use the prebuilt tree (`data/phylogeny/tree_263_NR_ref_genomes/tree_263_NR_genomes.treefile`) as an alternative.

```
source ~/.bash_profile
#set -ex

### SOFTWARE SETUP ##
####################
# required tools: iqtree=2.2.0.3
# set the name of the environment with installed tools
environment_name="iqtree"
# activate selected conda environment
eval "$(command conda 'shell.bash' 'hook' 2> /dev/null)" # initializes conda in sub-shell
conda activate ${environment_name}
conda info|egrep "conda version|active environment"

# copy the filtered alignment file to a new folder
mkdir -p data/phylogeny/tree_263_NR_ref_genomes
cp data/pangenome/panaroo_strict_i80_l90/core_gene_alignment.aln \
data/phylogeny/tree_263_NR_ref_genomes/core_gene_alignment.aln
cd data/phylogeny/tree_263_NR_ref_genome

# build the tree
iqtree -s core_gene_alignment.aln -o 561180.4 -m GTR+F+R10 -B 1000 -T 16
# if you do not have much time or computing power, turn on the fast tree search mode
#iqtree -s core_gene_alignment.aln -o 561180.4 -m GTR+F+R10 -T 16 -fast
```

#### 5.5 Visualizing phylogenetic tree

The phylogenetic tree of 263 *Bifidobacterium* genomes was manually visualized in **iTOL**.

#### 5.6 Calculating ANI of *Bifidobacterium longum* genomes

The phylogenomic analysis indicated that the *Bifidobacterium longum* species might have a more complex subspecies structure than previously described. To investigate it further, we computed pairwise ANI indices of 15 reference and 13 additional *Bifidobacterium longum* genomes using the ANIb algorithm implemented in **pyani**. For comparative purposes, the 13 additional genomes included isolates of non-human origin, such as type strains of *Bifidobacterium longum* subsp. *suis* and *Bifidobacterium longum* subsp. *suillum*.

To run the analysis, put the 28 corresponding FNA files to `data/genomes/Blongum_genomes/`.

```
source ~/.bash_profile
#set -ex

### SOFTWARE SETUP ##
####################
# required tools: pyani=0.2.12
# set the name of the environment with installed tools
```

```
environment_name="pyani"
# activate selected conda environment
eval "$(command conda 'shell.bash' 'hook' 2> /dev/null)" # initializes conda in sub-shell
conda activate ${environment_name}
conda info|egrep "conda version|active environment"

# run pyani
average_nucleotide_identity.py -i data/genomes/Blongum_genomes/ \
-o data/phylogeny/ANIb_28_Blongum_genomes \
-m ANIb
```

Load data.

```
# read the table with calculated ANI values for 28 Bifidobacterium longum genomes
ANIb_Blon <- read_tsv("data/phylogeny/ANIb_28_Blongum_genomes/ANIb_percentage_identity.tab") %>%
  # use the first column as row names
  column_to_rownames(var="...1")
```

The code chunk below creates a heatmap based on calculated ANI values.

Hierarchical clustering options:

- Maximum distance
- Average method

```
# convert the tibble with ANI values to a matrix
ANIb_Blon_mat <- as.matrix(ANIb_Blon)
ANIb_Blon_mat <- round(ANIb_Blon_mat, digits = 3)
old_to_new_names <- c("ATCC15697" = "ATCC 15697 = JCM 1222",
                      "JCM1217" = "JCM 1217")

# update row names only for specified ones, preserving the rest
rownames(ANIb_Blon_mat) <- ifelse(rownames(ANIb_Blon_mat) %in% names(old_to_new_names),
                                  old_to_new_names[rownames(ANIb_Blon_mat)],
                                  rownames(ANIb_Blon_mat))

# update column names only for specified ones, preserving the rest
colnames(ANIb_Blon_mat) <- ifelse(colnames(ANIb_Blon_mat) %in% names(old_to_new_names),
                                  old_to_new_names[colnames(ANIb_Blon_mat)],
                                  colnames(ANIb_Blon_mat))

# set colors
ANI_col_fun <- circlize::colorRamp2(c(0.945, 1), c("white", "#00b900"))
# create a function that will add ANI values to each cell
ANI_cell_fun = function(j, i, x, y, w, h, fill){
  grid.rect(x, y, w, h, gp = gpar(fill = fill, col = fill))
  # add ANI values to each cell
   if(ANIb_Blon_mat[j, i] <= 1){
    grid.text(sprintf("%.3f", ANIb_Blon_mat[j, i]), x, y, gp = gpar(fontsize = 5))
  }
}
```

```r
# define the distance measure and clustering method
dist_method = "maximum"
clust_method = "average"

# compute hierarchical clustering for rows
row_dist <- dist(t(ANIb_Blon_mat), method = dist_method)
row_clust <- hclust(row_dist, method = clust_method)

# compute hierarchical clustering for columns
col_dist <- dist(ANIb_Blon_mat, method = dist_method)
col_clust <- hclust(col_dist, method = clust_method)

# specify the name of the output
pdf("results/phylogeny/ANIb_28_Blongum.pdf", width=8.3, height=8)
# plot the heatmap
ANIb_Blon_ht <- ComplexHeatmap::Heatmap(ANIb_Blon_mat,
                             rect_gp = gpar(type = "none"),
                             column_dend_side = "bottom",
                             column_title = "ANI of Bifidobacterium longum genomes",
                             name = "ANI",
                             col = ANI_col_fun,
                             cell_fun = ANI_cell_fun,
                             cluster_rows = row_clust,
                             cluster_columns = col_clust,
                             row_names_side = "right")

draw(ANIb_Blon_ht)
# save the heatmap to a file
dev.off()
```

```
## pdf
##   2
```

```r
# draw the heatmap again to show it in the compiled markdown file
draw(ANIb_Blon_ht)
```

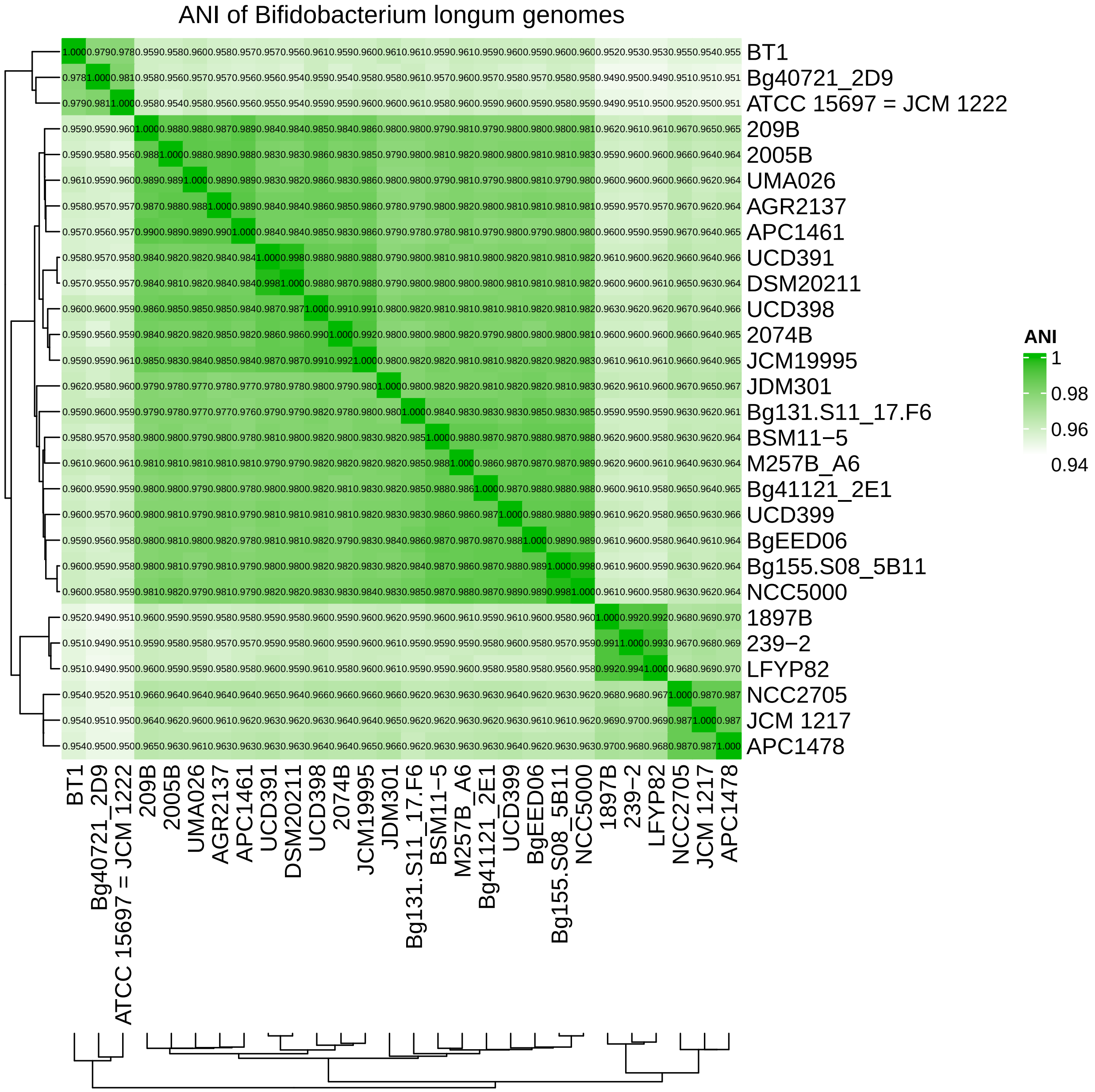

#### 5.7 Calculating ANI of *Bifidobacterium catenulatum* genomes

The phylogenetic analysis indicated that the *Bifidobacterium catenulatum* species might have a more complex subspecies structure than previously described. To investigate it further, we computed pairwise ANI indices of 11 reference genomes using the ANIb algorithm implemented in pyani. To run the analysis, put the 11 corresponding FNA files to `data/genomes/Bcatenulatum_group_genomes/`.

```
source ~/.bash_profile
#set -ex

### SOFTWARE SETUP ##
####################
# required tools: pyani=0.2.12
```

```
# set the name of the environment with installed tools
environment_name="pyani"
# activate selected conda environment
eval "$(command conda 'shell.bash' 'hook' 2> /dev/null)" # initializes conda in sub-shell
conda activate ${environment_name}
conda info|egrep "conda version|active environment"

# run pyani
average_nucleotide_identity.py -i data/genomes/Bcatenulatum_group_genomes/ \
-o data/phylogeny/ANIb_11_Bcatenulatum_genomes \
-m ANIb
```

Load data.

```
# read the file with calculated ANI values for 11 Bifidobacterium catenulatum-like genomes
ANIb_Bcat <- read_tsv("data/phylogeny/ANIb_11_Bcatenulatum_genomes/ANIb_percentage_identity.tab") %>%
  # use the first column as row names
  column_to_rownames(var="...1")
```

The code chunk below creates a heatmap based on calculated ANI values.

Hierarchical clustering options:

- Maximum distance
- Average method

```
# convert the tibble with ANI values to a matrix
ANIb_Bcat_mat <- as.matrix(ANIb_Bcat)
ANIb_Bcat_mat <- round(ANIb_Bcat_mat, digits = 3)

# set colors
ANI_col_fun <- circlize::colorRamp2(c(0.935, 1), c("white", "tomato2"))
# create a function that will add ANI values to each cell
ANI_cell_fun = function(j, i, x, y, w, h, fill){
  grid.rect(x, y, w, h, gp = gpar(fill = fill, col = fill))
  # add ANI values to each cell
   if(ANIb_Bcat_mat[j, i] <= 1){
    grid.text(sprintf("%.3f", ANIb_Bcat_mat[j, i]), x, y, gp = gpar(fontsize = 8))
  }
}

# define the distance measure and clustering method
dist_method = "maximum"
clust_method = "average"

# compute hierarchical clustering for rows
row_dist <- dist(t(ANIb_Bcat_mat), method = dist_method)
row_clust <- hclust(row_dist, method = clust_method)

# compute hierarchical clustering for columns
col_dist <- dist(ANIb_Bcat_mat, method = dist_method)
col_clust <- hclust(col_dist, method = clust_method)
```

```r
# specify the name of the output
pdf("results/phylogeny/ANb_11_Bcatenulatum.pdf", width=8, height=8)
# plot the heatmap
ANIb_Bcat_ht <- ComplexHeatmap::Heatmap(ANIb_Bcat_mat,
                             rect_gp = gpar(type = "none"),
                             column_dend_side = "bottom",
                             column_title = "ANI of Bifidobacterium catenulatum group genomes",
                             name = "ANI",
                             col = ANI_col_fun,
                             cell_fun = ANI_cell_fun,
                             cluster_rows = row_clust,
                             cluster_columns = col_clust,
                             row_names_side = "left")

draw(ANIb_Bcat_ht)
# save the heatmap to a file
dev.off()
```

```
## pdf
##   2
```

```r
# draw the heatmap again to show it in the compiled markdown file
draw(ANIb_Bcat_ht)
```

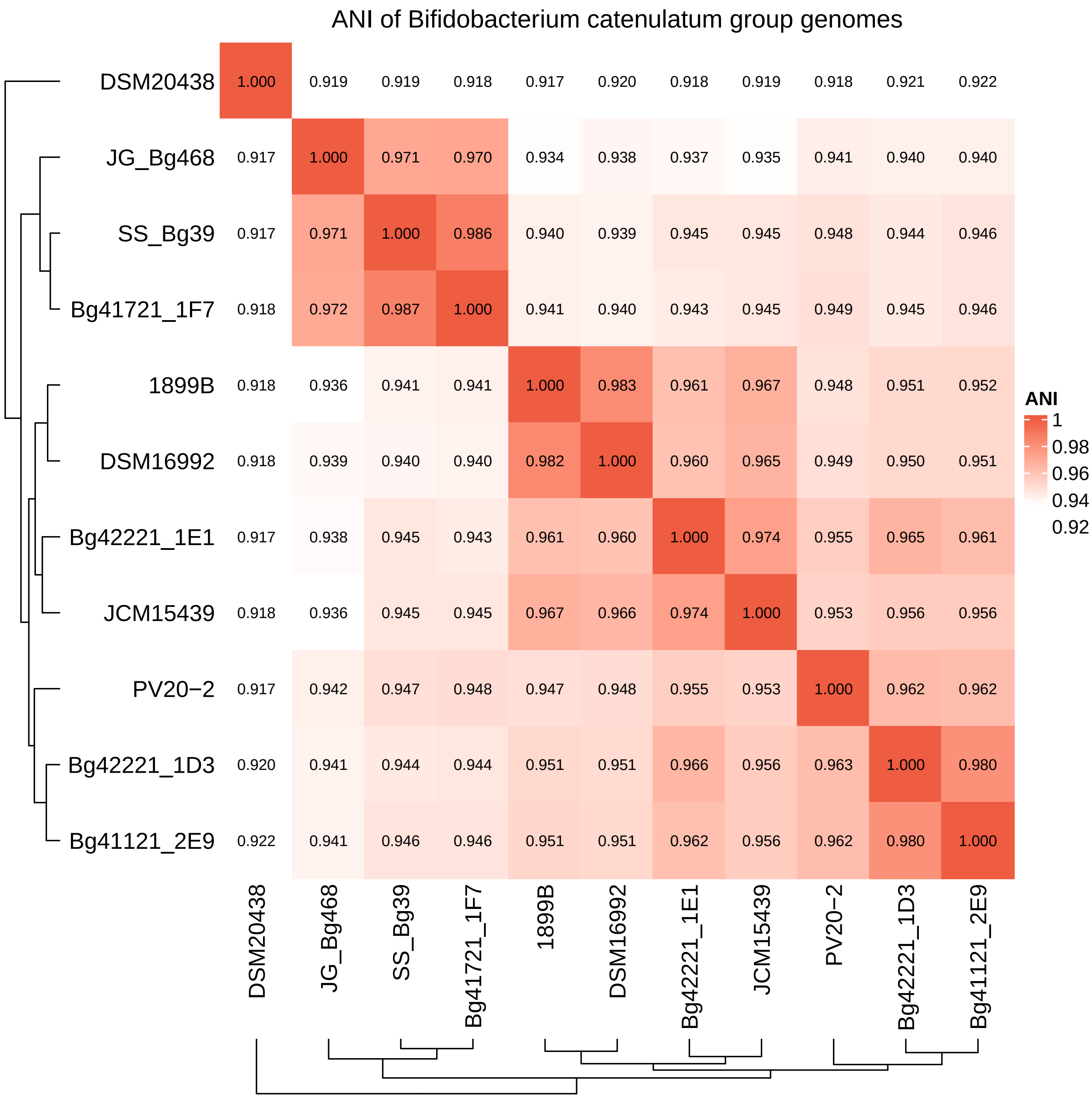

### 6 Comparison of gene annotations

#### 6.1 Introduction

This block contains code used for comparing curated gene annotations used in this work with annotations produced by commonly used automated tools **Prokka (v1.14.5)** and **EggNOG-mapper (v2.1.12)** for 263 reference *Bifidobacterium* genomes.

#### 6.2 Load data

```
# read table with annotations
annotations.df <- read_xlsx("data/tables/annotation_comparison.xlsx")
```

#### 6.3 Gene annotations: manual curation vs. Prokka/EggNOG

We compared gene annotations obtained via manual curation with annotations produced by Prokka and EggNOG-mapper. EggNOG-mapper outputs multiple annotation fields (description, EC number, KEGG_ko numbers). We aggregated information from these fields, prioritizing the descriptions of EC and KEGG_ko numbers.

This comparison has four categories:

- new = manually curated annotations update **general (non-specific)** annotations produced by Prokka/EggNOG-mapper. Example:
  - Putative fructose MFS permease (TC 2.A.1.7) vs. hypothetical protein (Prokka) or Major Facilitator Superfamily (EggNOG)
  - 2'FL, 3FL ABC transporter substrate-binding component (TC 3.A.1.1) vs. hypothetical protein (Prokka) or multiple sugar transport system substrate-binding protein (EggNOG)
- corrected = manually curated annotations update **incorrect specific** annotations produced by Prokka/EggNOG-mapper. Examples:
  - D-mannose isomerase (EC 5.3.1.7) vs. Sulfoquinovose isomerase (Prokka) or N-acylglucosamine 2-epimerase (EggNOG)
  - Arabinoxylooligosaccharide ABC transporter permease component 1 (TC 3.A.1.1) vs. Melibiose/raffinose/stachyose import permease protein MelD (Prokka) or xylobiose transport system permease protein (EggNOG)
- refined = annotations are similar in principle, but manually curated ones are more precise. Examples:
  - Adding linkage specificity to glycoside hydrolases: alpha-1,6-L-fucosidase (GH29) vs. alpha-L-fucosidase [EC:3.2.1.51] (EggNOG)
  - Specific annotation vs. list with many activities including the correct one: D-mannonate oxidoreductase (EC 1.1.1.57) vs. fructuronate reductase; mannonate oxidoreductase; mannonic dehydrogenase; D-mannonate dehydrogenase; D-mannonate:NAD+ oxidoreductase; tagaturonate reductase; altronic oxidoreductase; altronate oxidoreductase; TagUAR; altronate dehydrogenase; D-tagaturonate reductase (EggNOG)
- same = annotations are identical

Plot the results of the comparison.

First, what percentage of 589 functional roles is constituted by each comparison group.

```
# convert prokka_comparison and eggnog_comparison
# into a single column "comparison" with a source column
comparison_by_role <- annotations.df %>%
  pivot_longer(cols = c(prokka_comparison, eggnog_comparison),
               names_to = "source", values_to = "comparison") %>%
```

```r
  # recode source before grouping
  mutate(source = dplyr::recode(source,
                         "prokka_comparison" = "Prokka",
                         "eggnog_comparison" = "EggNOG")) %>%
  group_by(functional_role_type, source, comparison) %>%
  summarise(n = n(), .groups = "drop")

# add a "total" category by summing counts across all functional_role_type
comparison_by_role <- comparison_by_role %>%
  bind_rows(comparison_by_role %>%
            group_by(source, comparison) %>%
            summarise(n = sum(n), .groups = "drop") %>%
            mutate(functional_role_type = "total"))  # add a new category total

# ensure factor ordering
comparison_by_role <- comparison_by_role %>%
  mutate(comparison = factor(comparison, levels = c("new", "corrected", "refined", "same")),
                             source = factor(source, levels = c("Prokka", "EggNOG")),
                            functional_role_type = factor(functional_role_type,
                             levels = c("transporter", "CAZyme", "downstream_catabolism",
                                        "regulator", "total")))

# create a stacked barplot
ggplot(comparison_by_role,
       aes(x = source, y = n, fill = comparison)) +
  geom_bar(stat = "identity", position = "fill") +
  scale_fill_manual(values = c("new" = "#5e9155", "corrected" = "#ffa600",
                               "refined" = "#4f8fe6", "same" = "#d1d3d4")) +
  theme_bw() +
  facet_grid(~ functional_role_type) +
  labs(title = "total",
       x = "%  of 589 functional roles", y = "") %>%
  scale_y_continuous(breaks = seq(0, 1, 0.1),
                     labels = scales::percent_format(accuracy = 1)) +
 theme(axis.text.x = element_text(angle = 60, hjust = 1))
```

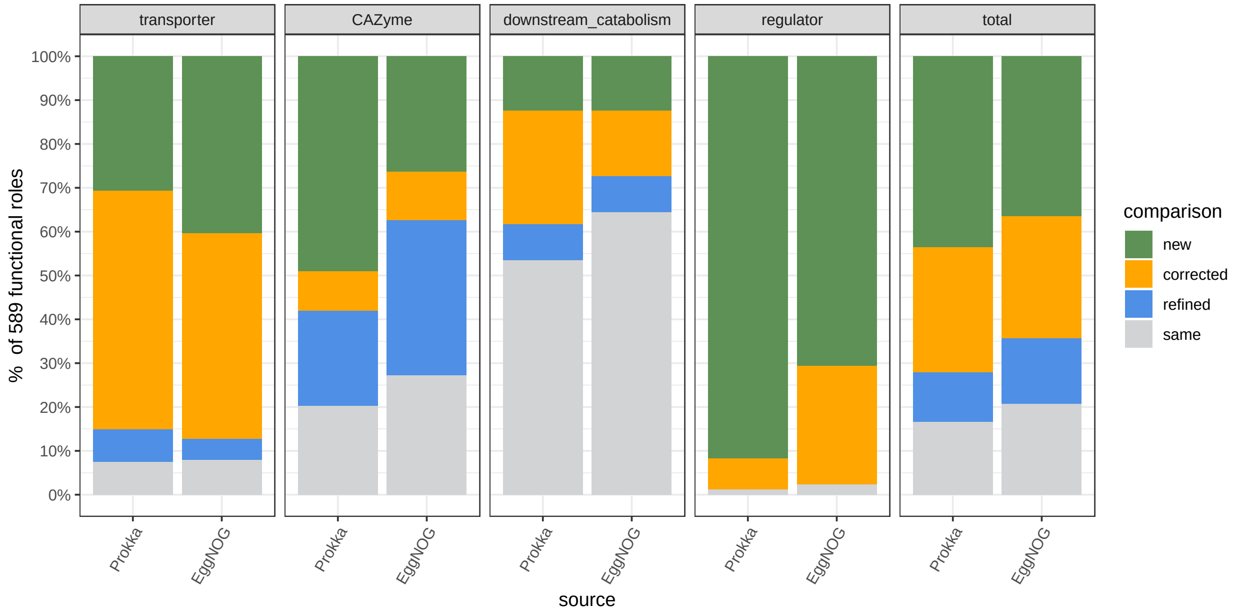

```
# save the figure to a file
ggsave("results/annotations/percent_fr.pdf", device = "pdf", width = 10, height = 5)
```

Second, what percentage of 39589 annotated genes is constituted by each comparsion group.

```
# split "mcseed_IDs" into multiple rows
annotations.df_expanded <- annotations.df %>%
  separate_rows(mcseed_IDs, sep = "; ")

# convert prokka_comparison and eggnog_comparison
# into a single column "comparison" with a source column
comparison_by_role_expanded <- annotations.df_expanded %>%
  pivot_longer(cols = c(prokka_comparison, eggnog_comparison),
               names_to = "source", values_to = "comparison") %>%
  mutate(source = dplyr::recode(source,
                          "prokka_comparison" = "Prokka",
                          "eggnog_comparison" = "EggNOG")) %>%
  group_by(functional_role_type, source, comparison) %>%
  summarise(n = n(), .groups = "drop")

# add a "total" category by summing counts across all functional_role_type
comparison_by_role_expanded <- comparison_by_role_expanded %>%
  bind_rows(comparison_by_role_expanded %>%
              group_by(source, comparison) %>%
              summarise(n = sum(n), .groups = "drop") %>%
              mutate(functional_role_type = "total"))

# ensure factor ordering
comparison_by_role_expanded <- comparison_by_role_expanded %>%
  mutate(comparison = factor(comparison, levels = c("new", "corrected", "refined", "same")),
         source = factor(source, levels = c("Prokka", "EggNOG")),
         functional_role_type = factor(functional_role_type,
                                       levels = c("transporter", "CAZyme",
```

```
                                                "downstream_catabolism", "regulator",
                                                "total")))

# create a stacked barplot
ggplot(comparison_by_role_expanded,
       aes(x = source, y = n, fill = comparison)) +
  geom_bar(stat = "identity", position = "fill") +
  scale_fill_manual(values = c("new" = "#5e9155", "corrected" = "#ffa600",
                               "refined" = "#4f8fe6", "same" = "#d1d3d4")) +
  theme_bw() +
  facet_grid(~ functional_role_type) +
  labs(title = "total",
       x = "%  of annotations of 39589 genes", y = "") %>%
  scale_y_continuous(breaks = seq(0, 1, 0.1),
                     labels = scales::percent_format(accuracy = 1)) +
 theme(axis.text.x = element_text(angle = 60, hjust = 1))
```

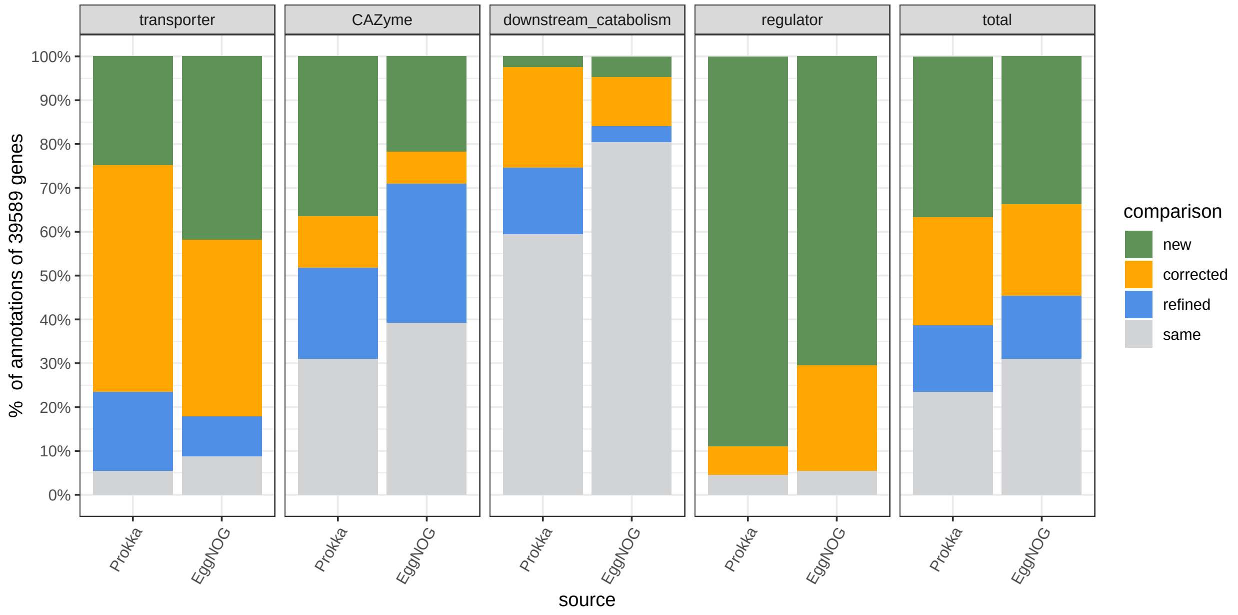

```
# save the figure to a file
ggsave("results/annotations/percent_genes.pdf", device = "pdf", width = 10, height = 5)
```

Calculate the percentage of updated annotations for each tool:

```
# sum counts for new + corrected + refined + same
subset_counts <- comparison_by_role_expanded %>%
  filter(comparison %in% c("new", "corrected", "refined")) %>%
  group_by(source, functional_role_type) %>%
  summarise(sum_n = sum(n), .groups = "drop")

# sum all counts
total_counts <- comparison_by_role_expanded %>%
  filter(comparison %in% c("new", "corrected", "refined", "same")) %>%
```

| source | functional_role_type | sum_n | sum_all | percentage |
|---|---|---|---|---|
| Prokka | transporter | 11260 | 11905 | 94.6 |
| Prokka | CAZyme | 9398 | 13618 | 69.0 |
| Prokka | downstream_catabolism | 2782 | 6858 | 40.6 |
| Prokka | regulator | 6881 | 7208 | 95.5 |
| Prokka | total | 30321 | 39589 | 76.6 |
| EggNOG | transporter | 10871 | 11905 | 91.3 |
| EggNOG | CAZyme | 8276 | 13618 | 60.8 |
| EggNOG | downstream_catabolism | 1347 | 6858 | 19.6 |
| EggNOG | regulator | 6815 | 7208 | 94.5 |
| EggNOG | total | 27309 | 39589 | 69.0 |

```
  group_by(source, functional_role_type) %>%
  summarise(sum_all = sum(n), .groups = "drop") %>%
  dplyr::select(-source, -functional_role_type)

# join with subset_counts and calculate percentages
percentage_results <- cbind(subset_counts, total_counts) %>%
  mutate(percentage = round((sum_n / sum_all) * 100, 1))

# plot the table:
gt(percentage_results)
```

---

### 7 CAZyme representation in 263 *Bifidobacterium* genomes

#### 7.1 Introduction

This block contains the code used to analyze the representation of genes encoding Carbohydrate Active Enzymes (CAZymes, specifically glycoside hydrolases (GHs), carbohydrate esterases (CEs), and polysaccharide lyases (PLs)) in 263 *Bifidobacterium* genomes. Specifically, we checked how many of these CAZymes were captured in our metabolic reconstruction.

The following software is required:

1. **dbCAN (v4.0.0)**

**Note**: Given the potential challenges with installing and running dbCAN, we provide processed dbCAN outputs in `data/CAZyme`:

1. `GH_output_subfamilies` # concatenated dbCAN output
2. `CAZyme_families` # representation of GH/CE/PL families in 263 *Bifidobacterium* genomes. GH subfamilies are collapsed (e.g., GH43_22 and GH_43_24 are treated as GH43)
3. `CAZyme_subfamilies` # representation of GH/CE/PL families and subfamilies in 263 *Bifidobacterium* genomes. GH subfamilies are treated as distinct columns

#### 7.2 Load data

```
# read the table with metadata for 263 genomes
bpm_263_join <- read_tsv("data/tables/BPM_263_NR_genomes_carbs.txt",
                         col_types = cols(.default = "c")) %>%
                arrange(genome_ID) %>%
                dplyr::select(genome_ID, genome_name, curated_taxonomy)
# read the table with CAZyme representation
cazy_subfam_df <- read_tsv("data/CAZyme/CAZyme_subfamilies.txt",
                           col_types = cols(.default = "c")) %>%
                  mutate_at(c(3:111), as.numeric) %>%
                  left_join(bpm_263_join, by = c("Organism" = "genome_name")) %>%
                  dplyr::select(-seed_id, -CE0, -GH0) %>%
                  arrange(genome_ID)
# read the processed dbCAN output
cazy_df <- read_tsv("data/CAZyme/GH_output_subfamilies.txt",
                    col_types = cols(.default = "c"))
```

#### 7.3 Set colors

Define point colors and shapes used throughout this block.

```
# arguments for scale_shape_manual
genomes_263_shapes <- c(21,24,21,21,21,
                        21,24,23,21,21,
                        21,22,21,21,21,
                        21,21,21,23)
# arguments for scale_fill_manual
# breaks (how groups are encoded in the table)
genomes_263_breaks <- c("Bifidobacterium adolescentis",
                        "Bifidobacterium angulatum",
                        "Bifidobacterium animalis subsp. lactis",
                        "Bifidobacterium bifidum",
                        "Bifidobacterium breve",
                        "Bifidobacterium catenulatum subsp. catenulatum",
                        "Bifidobacterium catenulatum subsp. kashiwanohense",
                        "Bifidobacterium catenulatum subsp. kashiwanohense_A",
                        "Bifidobacterium dentium",
                        "Bifidobacterium gallicum",
                        "Bifidobacterium globosum",
                        "Bifidobacterium hominis",
                        "Bifidobacterium longum subsp. infantis",
                        "Bifidobacterium longum subsp. longum",
                        "Bifidobacterium longum subsp. nov.",
                        "Bifidobacterium longum subsp. suis",
                        "Bifidobacterium pseudocatenulatum",
                        "Bifidobacterium scardovii",
                        "Bifidobacterium thermophilum")
# values (color codes)
genomes_263_colors <- c("#9b91c1", # Bifidobacterium adolescentis
                        "#ffff7f", # Bifidobacterium angulatum
                        "grey", # Bifidobacterium animalis subsp. lactis
```

```
                        "#222c80", # Bifidobacterium bifidum
                        "#00a2ff", # Bifidobacterium breve
                        "tomato2", # Bifidobacterium catenulatum subsp. catenulatum
                        "tomato2", # Bifidobacterium catenulatum subsp. kashiwanohense
                        "tomato2", # Bifidobacterium catenulatum subsp. kashiwanohense_A
                        "#8e063c", # Bifidobacterium dentium
                        "#ffffff", # Bifidobacterium gallicum
                        "#d0a079", # Bifidobacterium globosum
                        "#a2d2de", # Bifidobacterium hominis
                        "#c0d28b", # Bifidobacterium longum subsp. infantis
                        "#51796f", # Bifidobacterium longum subsp. longum
                        "black", # Bifidobacterium longum subsp. nov.
                        "#00b400", # Bifidobacterium longum subsp. suis
                        "#ffa600", # Bifidobacterium pseudocatenulatum
                        "#f79cd4", # Bifidobacterium scardovii
                        "#ffffff") # Bifidobacterium thermophilum)

# labels (what will appear in the legend)
genomes_263_species <- c("B. adolescentis",
                         "B. angulatum",
                         "B. animalis ssp. lactis",
                         "B. bifidum",
                         "B. breve",
                         "B. catenulatum ssp. catenulatum",
                         "B. catenulatum ssp. kashiwanohense",
                         "B. catenulatum ssp. kashiwanohense_A",
                         "B. dentium",
                         "B. gallicum",
                         "B. globosum",
                         "B. hominis",
                         "B. longum ssp. infantis",
                         "B. longum ssp. longum",
                         "B. longum ssp. nov.",
                         "B. longum ssp. suis",
                         "B. pseudocatenulatum",
                         "B. scardovii",
                         "B. thermophilum")
```

#### 7.4 Ordination

We used Principal Component Analysis (PCA) for the ordination of a table containing the representation of GH/CE/PL (sub)families in 263 genomes.

```
# extract the matrix from the CAZyme tibble
cazy_subfam_mat <- as.matrix((cazy_subfam_df[, 2:108]))
# do PCA
cazy.pca.res <- prcomp(cazy_subfam_mat, scale.=F, retx=T)
# sdev^2 captures eigenvalues from the PCA result
cazy.pc.var <- cazy.pca.res$sdev^2
# calculate the percentage of the total variance explained by each PC
cazy.pc.per <- round(cazy.pc.var/sum(cazy.pc.var)*100, 1)
# extract PCA results to a tibble
```

```r
cazy.pca.res.df <- as_tibble(cazy.pca.res$x)

# plot the PCA results
# create a vector with taxonomy (group)
cazy.species <- cazy_subfam_df$curated_taxonomy
# create a vector with genome names
cazy.genomes <- cazy_subfam_df$Organism
# select points (genomes) that will be labeled
cazy.genomes_short <- ifelse(cazy.genomes ==
                        "Bifidobacterium longum subsp. suis Bg131.S11_17.F6",
                        "Bg131.S11_17.F6",
                        ifelse(cazy.genomes ==
                        "Bifidobacterium catenulatum subsp. kashiwanohense Bg42221_1E1",
                        "Bg42221_1E1", ""))
# plot
ggplot(cazy.pca.res.df) +
  aes(x=PC1, y=PC2, fill=cazy.species, shape=cazy.species, stroke = 0.15) +
  geom_point(size=2) +
  scale_shape_manual(values=genomes_263_shapes) +
  guides(shape="none") +
  scale_fill_manual(name = "Taxonomy",
                    breaks=genomes_263_breaks,
                    values=genomes_263_colors,
                    labels=genomes_263_species) +
  guides(fill = guide_legend(override.aes=list(shape=genomes_263_shapes))) +
  # add text labels for selected genomes
  geom_text_repel(aes(label = cazy.genomes_short), size = 3, fontface=1, color="black",
                  min.segment.length = 0,
                  seed = 42, box.padding = 1, max.overlaps = 100) +
  labs(title= "PCA of GH/CE/PL (sub)families representation") +
  xlab(paste0("PC1 (",cazy.pc.per[1],"%",")")) +
  ylab(paste0("PC2 (",cazy.pc.per[2],"%",")")) +
  coord_fixed(1) +
  theme_bw() +
  theme(plot.title = element_text(face="bold"),
        axis.title = element_text(color = "black"),
        axis.text = element_text(color = "black"))
```

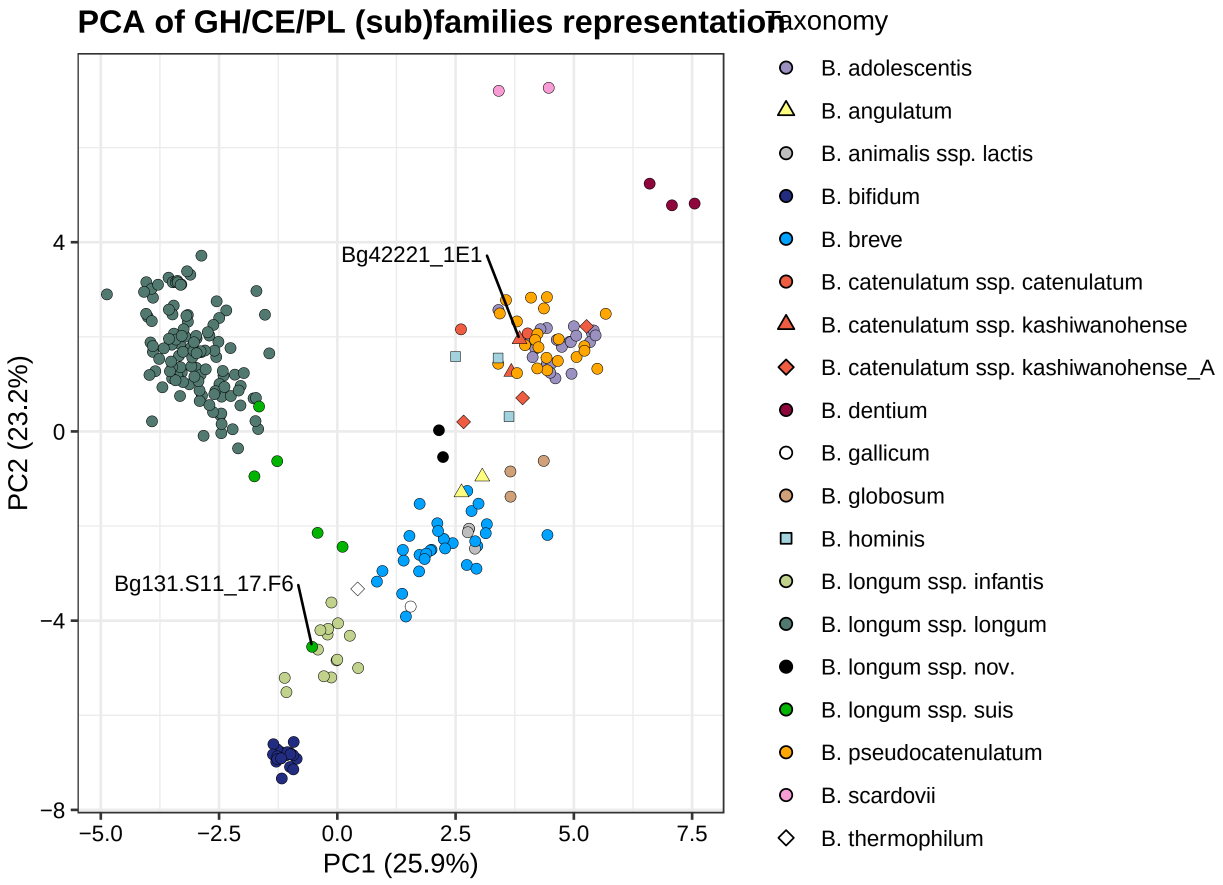

```
# save the plot to a file
ggsave("results/CAZyme/CAZyme_263_PCA.pdf", width = 7, height = 5)
```

#### 7.5 Percent of CAZymes captured in the metabolic reconstruction

For each genome, we calculated the percentage of CAZymes (GHs/CEs/PLs) captured in the metabolic reconstruction (i.e., ratio: number of GHs/CEs/PLs captured in mcSEED subsystems / total number of GHs/CEs/PLs identified by dbCAN).

```
# calculate the ratio of CAZymes (GHs/CEs/PLs) captured in mcSEED subsystems
cazy_captured_mcseed <- cazy_df %>%
  group_by(Organism) %>%
```

```
  summarize(ratio = 100*(1 - (sum(Subsystem == "-") / n()))) %>%
  left_join(bpm_263_join, by = c("Organism" = "genome_name")) %>%
  arrange(genome_ID)

# plot a swarmplot + boxplot
# create a vector with curated taxonomy
cazy_species <- cazy_captured_mcseed$curated_taxonomy
# plot
ggplot() +
  geom_boxplot(data = cazy_captured_mcseed,
               mapping = aes(x="", y=ratio),
               outlier.shape = NA, width=0.9, lwd = 0.3) +
  ggbeeswarm::geom_quasirandom(data = cazy_captured_mcseed,
                               aes(x="", y=ratio, fill=cazy_species, shape=cazy_species),
                               color = "black", stroke = 0.15, size = 4) +
  scale_shape_manual(values=genomes_263_shapes) +
  guides(shape="none") +
  scale_fill_manual(name = "Taxonomy",
                    breaks=genomes_263_breaks,
                    values=genomes_263_colors,
                    labels=genomes_263_species) +
  guides(fill = guide_legend(override.aes=list(shape=genomes_263_shapes))) +
  theme_bw() +
  theme(plot.title = element_text(face="bold"),
        axis.title = element_text(color = "black"),
        axis.text = element_text(color = "black")) +
  coord_cartesian(ylim = c(60, 100)) +
  labs(x = "", y = "% CAZymes captured in metabolic reconstruction")
```

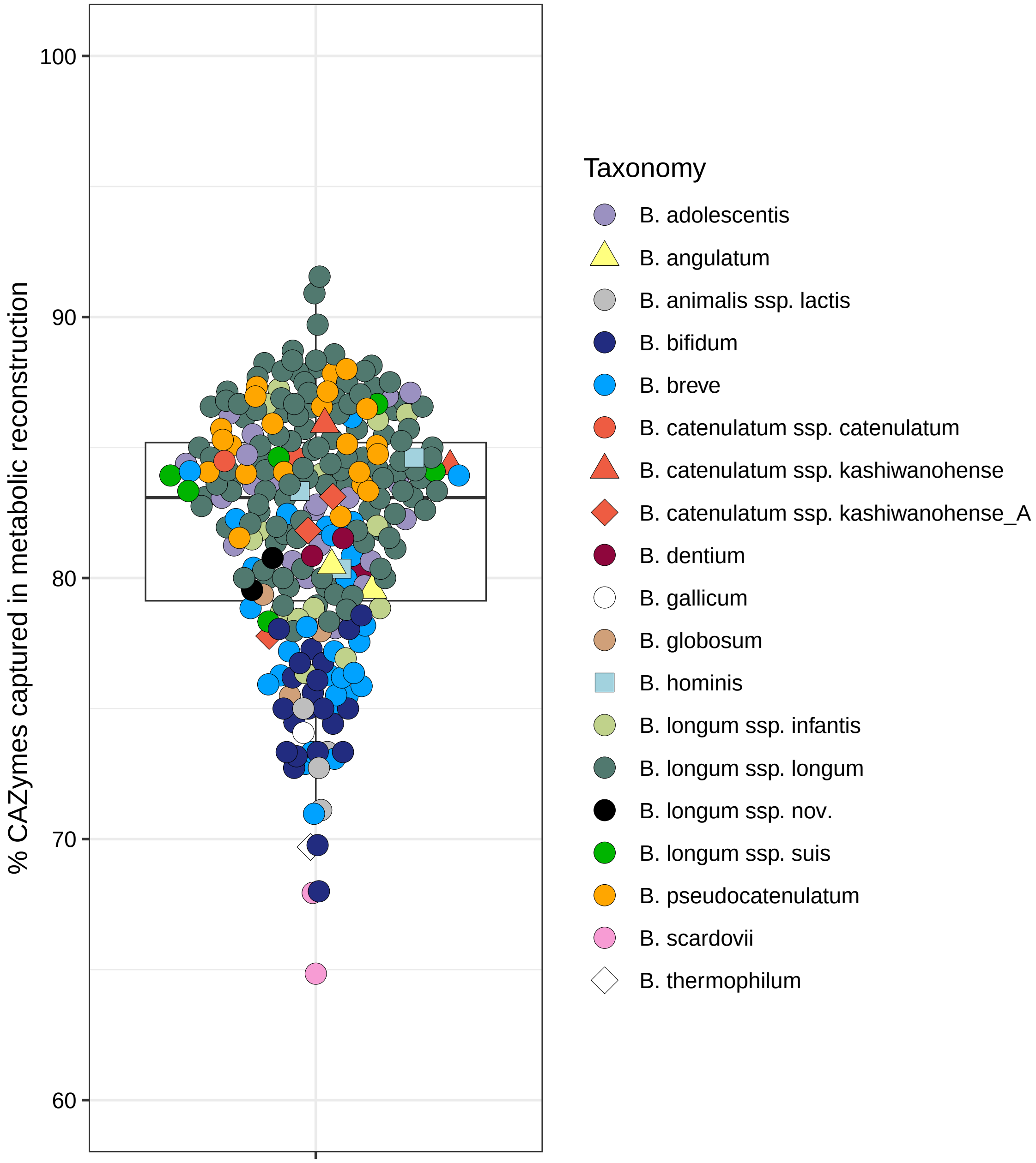

```
# save the plot to a file
ggsave("results/CAZyme/percent_captured_in_mcSEED.pdf", width = 7, height = 7)
```

The total and average (across all 263 genomes) percentages of captured CAZymes:

```
# calculate the total % of captured CAZymes
not_captured <- cazy_df %>%
  filter(Subsystem == "-") %>%
  nrow()
captured <- round(1 - not_captured / length(cazy_df$Subsystem), digits = 3) * 100
```

```r
# calculate the average % of captured CAZymes
ratio_mean <- round(mean(cazy_captured_mcseed$ratio), digits = 1)
ratio_sd <- round(sd(cazy_captured_mcseed$ratio), digits = 1)
# print calculated values
print(paste0("Total percentage of captured CAZymes:"," ",captured, "%"))
```

```
## [1] "Total percentage of captured CAZymes: 82.2%"
```

```r
print(paste0("Mean ± SD percentage of captured CAZymes:"," ",ratio_mean," ","±"," ",ratio_sd))
```

```
## [1] "Mean ± SD percentage of captured CAZymes: 81.9 ± 4.6"
```

---

### 8 Representation of predicted carbohydrate utilization phenotypes in 263 reference *Bifidobacterium* strains

#### 8.1 Introduction

The block describes the analysis of Binary Phenotype Matrix (BPM) containing 68 carbohydrate utilization phenotypes predicted in 263 reference *Bifidobacterium* strains. Four additional phenotypes (GalNAc, ManNAc, Man, GalA) were excluded from the analysis since, for these glycans, all strains had **predicted binary phenotype 0**.

#### 8.2 Load data

```r
# read the BPM (carbs)
bpm_263_carb_df <- read_tsv("data/tables/BPM_263_NR_genomes_carbs.txt",
                            col_types = cols(.default = "c")) %>%
                            mutate_at(c(4:75), as.numeric) %>%
                            dplyr::select(-c(ManNAc, GalNAc, Man, GalA)) %>%
                            arrange(genome_ID)
# read the table with metadata for predicted carbohydrate utilization phenotypes
phenotype_metadata <- read_tsv("data/tables/phenotype_metadata_carbs.txt",
                      col_types = cols(.default = "c")) %>%
                      filter(!(phenotype %in% c("ManNAc", "GalNAc", "Man", "GalA")))
```

#### 8.3 Hierarchical clustering of the BPM of 263 *Bifidobacterium* genomes

The following heatmap shows the hierarchical clustering of the BPM for 263 *Bifidobacterium* genomes.

Hierarchical clustering options:

- **Distance metric**: Hamming distance (equivalent to Manhattan distance for binary data)
- **Linkage method**: Average

```r
# extract the binary matrix
bpm_263_mat <- as.matrix((bpm_263_carb_df[, 4:71]))
# add rownames to the matrix
rownames(bpm_263_mat) <- bpm_263_carb_df$genome_ID

# create a vector with taxonomy (group)
taxonomy <- bpm_263_carb_df$curated_taxonomy
# extract vectors containing data about glycan type and origin
glycan_type <- phenotype_metadata$type_group
glycan_origin <- phenotype_metadata$origin
# create a coloring function
col_fun <- structure(c("white", "#08306b"), names = c("0", "1"))
# create a row annotation specifying taxonomy
ha_263_1 <- HeatmapAnnotation(
  which = c("row"),
  Taxonomy = taxonomy,
  col = list(Taxonomy = c("Bifidobacterium adolescentis" = "#9b91c1",
                          "Bifidobacterium angulatum" = "#ffff7f",
                          "Bifidobacterium animalis subsp. lactis" = "#bebebe",
                          "Bifidobacterium bifidum" = "#222c80",
                          "Bifidobacterium breve" = "#00a2ff",
                          "Bifidobacterium catenulatum subsp. catenulatum" = "#ee5c42",
                          "Bifidobacterium catenulatum subsp. kashiwanohense" = "#ee5c42",
                          "Bifidobacterium catenulatum subsp. kashiwanohense_A" = "#ee5c42",
                          "Bifidobacterium dentium" = "#8e063c",
                          "Bifidobacterium gallicum" = "#ffffff",
                          "Bifidobacterium globosum" = "#d0a079",
                          "Bifidobacterium hominis" = "#a2d2de",
                          "Bifidobacterium longum subsp. infantis" = "#c0d28b",
                          "Bifidobacterium longum subsp. longum" = "#51796f",
                          "Bifidobacterium longum subsp. suis" = "#00b400",
                          "Bifidobacterium longum subsp. nov." = "black",
                          "Bifidobacterium pseudocatenulatum" = "#ffa600",
                          "Bifidobacterium scardovii" = "#f79cd4",
                          "Bifidobacterium thermophilum" = "#ffffff")),
  show_annotation_name = FALSE,
  simple_anno_size = unit(3, "mm"))
# create two column annotations specifying glycan type and origin
ha_263_2 <- HeatmapAnnotation(
  type = glycan_type,
  origin = glycan_origin,
  col = list(type = c("monosaccharides_and_derivatives" = "#E6E7E8",
                      "di_and_oligosaccharides" = "#BCBEC0",
                      "polysaccharides" = "#808285"),
             origin = c("universal" = "#ffd22d",
                        "host (animal)" = "#cbbedd",
                        "plant" = "#8dc63f",
                        "bacterial" = "#603913",
                        "fungal" = "#008aa1" )),
  show_annotation_name = FALSE,
  simple_anno_size = unit(3, "mm"))

# plot the heatmap
```

```r
pdf("results/phenotypes/BPM_263_heatmap.pdf", width=15, height=10)
ht_263_all <- ComplexHeatmap::Heatmap(bpm_263_mat,
                              name = "Predicted phenotype",
                              right_annotation = ha_263_1,
                              bottom_annotation = ha_263_2,
                              col = col_fun,
                              clustering_distance_rows = function(m)
                                dist(m, method = "manhattan"),
                              clustering_distance_columns = function(m)
                                dist(m, method = "manhattan"),
                              clustering_method_rows = "average",
                              clustering_method_columns = "average",
                              rect_gp = gpar(col = "grey", lwd = 0.05),
                              # do not show row names
                              show_row_names = TRUE,
                              row_names_gp = gpar(fontsize = 3),
                              column_names_gp = gpar(fontsize = 5),
                              column_names_rot = 70,
                              width = unit(200, "mm"),
                              height = unit(200, "mm"))
draw(ht_263_all)
# save the heatmap to a file
dev.off()
```

```
## pdf
##   2
```

```r
# draw the heatmap again to show it in the compiled markdown file
draw(ht_263_all)
```

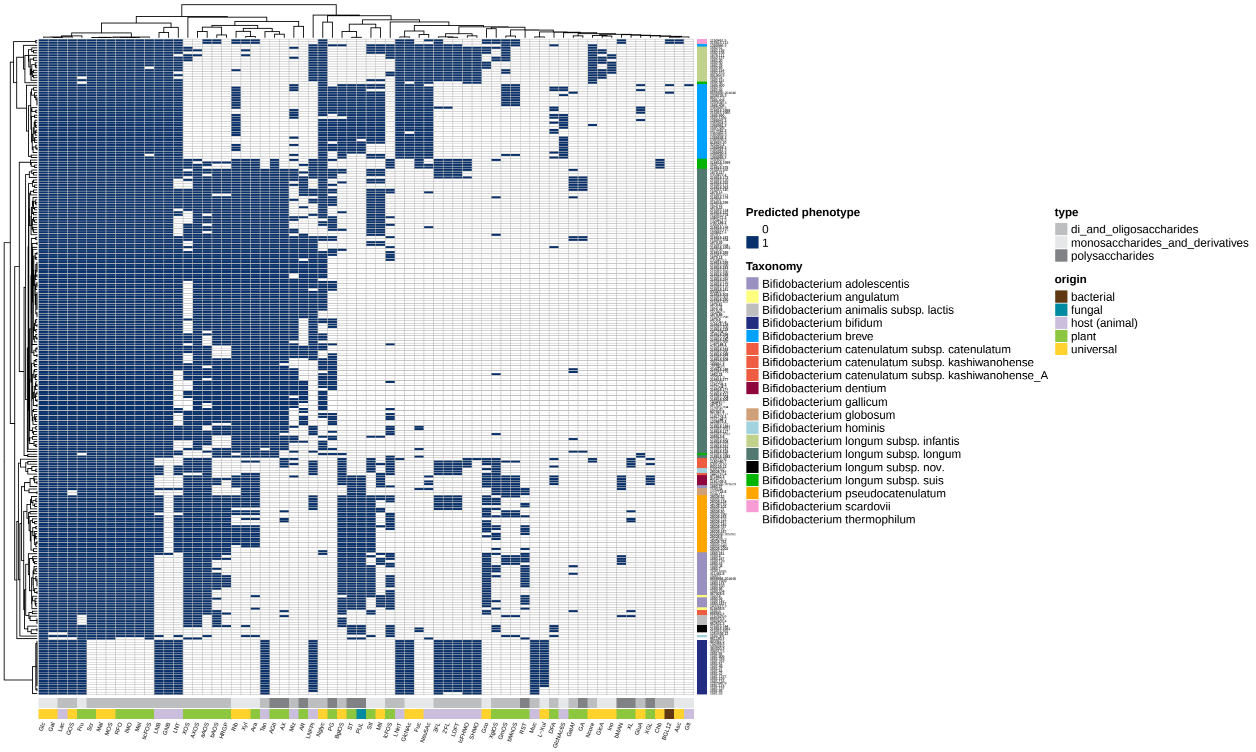

#### 8.4 Conservation of predicted carbohydrate utilization phenotypes

```
# calcualte "average" phenotypes for each taxon
bpm_263_mean <- bpm_263_carb_df %>%
  group_by(curated_taxonomy) %>%
  summarise_at(vars(Glc:`L-Xul`), list(mean))

# count columns with only 0 and 1
# i.e., binary phenotypes that are always conserved within taxonomic groups
binary_columns <- sum(apply(bpm_263_mean , 2, function(col) all(col %in% c(0, 1))))

# count columns with intermediate values in the [0,1] range
# i.e., binary phenotypes that vary within taxonomic groups
range_columns <- sum(apply(bpm_263_mean, 2, function(col) any(col > 0 & col < 1)))

# print the counts
print(paste0("Number of predicted phenotypes conserved within taxonomic groups: ", binary_columns))
```

```
## [1] "Number of predicted phenotypes conserved within taxonomic groups: 11"
```

```
print(paste0("Number of variable predicted phenotypes: ", range_columns))
```

```
## [1] "Number of variable predicted phenotypes: 57"
```

#### 8.5 Comparison between phylogenetic and metabolic distances

Here, we generate a tanglegram to compare the **p**hylogenetic relationships and "metabolic distance" among *Bifidobacterium* genomes. The phylogenetic tree is constructed from 263 genomes and rooted using an outgroup. Metabolic distance is calculated using a Hamming distance matrix based on the presence or absence of carbohydrate utilization pathways.

```
# read the file with the phylogenetic tree
tree_263 <- read.tree("data/phylogeny/tree_263_NR_ref_genomes/tree_263_NR_genomes.treefile")
# force the tree to be ultrametric using penalized likelihood
tree_263_ultra <- chronos(tree_263, lambda = 1)
```

```
##
## Setting initial dates...
## Fitting in progress... get a first set of estimates
##          (Penalised) log-lik = -17.87337
## Optimising rates... dates... -17.87337
## Optimising rates... dates... -17.87333
##
## log-Lik = -17.81193
## PHIIC = 1601.67
```

```
# define an outgroup
outgroup <- "561180.4"
# check if the outgroup exists in the tree
if (outgroup %in% tree_263_ultra$tip.label) {
  tree_263_ultra_rooted <- root(tree_263_ultra, outgroup = outgroup, resolve.root = TRUE)
} else {
  stop("Outgroup not found in the tree. Choose a different outgroup.")
}
# convert hclust to dendrogram
dend1 <- as.dendrogram(as.hclust(tree_263_ultra_rooted))

# extract the binary matrix
bpm_263_mat <- as.matrix((bpm_263_carb_df[, 4:71]))
# add rownames to the matrix
rownames(bpm_263_mat) <- bpm_263_carb_df$genome_ID
# compute the hamming distance matrix
hamming_dist <- as.matrix(dist(bpm_263_mat, method = "manhattan"))
# convert the distance matrix into an hclust object
hclust_hamming <- hclust(as.dist(hamming_dist), method = "average")
# convert hclust to dendrogram
dend2 <- as.dendrogram(hclust_hamming)

# plot the tanglegram
# set tanglegram link colors
taxonomy_colors <- c(
  "Bifidobacterium adolescentis" = "#9b91c1",
  "Bifidobacterium angulatum" = "#ffff7f",
  "Bifidobacterium animalis subsp. lactis" = "#bebebe",
  "Bifidobacterium bifidum" = "#222c80",
  "Bifidobacterium breve" = "#00a2ff",
  "Bifidobacterium catenulatum subsp. catenulatum" = "#ee5c42",
```

```r
  "Bifidobacterium catenulatum subsp. kashiwanohense" = "#ee5c42",
  "Bifidobacterium catenulatum subsp. kashiwanohense_A" = "#ee5c42",
  "Bifidobacterium dentium" = "#8e063c",
  "Bifidobacterium gallicum" = "#ffffff",
  "Bifidobacterium globosum" = "#d0a079",
  "Bifidobacterium hominis" = "#a2d2de",
  "Bifidobacterium longum subsp. infantis" = "#c0d28b",
  "Bifidobacterium longum subsp. longum" = "#51796f",
  "Bifidobacterium longum subsp. suis" = "#00b400",
  "Bifidobacterium longum subsp. nov." = "black",
  "Bifidobacterium pseudocatenulatum" = "#ffa600",
  "Bifidobacterium scardovii" = "#f79cd4",
  "Bifidobacterium thermophilum" = "#ffffff"
)

labels <- dend1 %>%
  set("labels_to_char") %>%
  labels

metadata <- bpm_263_carb_df %>%
  dplyr::select(genome_ID, curated_taxonomy) %>%
  mutate(taxonomy_color = taxonomy_colors[curated_taxonomy]) %>%
  mutate(genome_ID = factor(genome_ID, levels = labels)) %>%
  arrange(genome_ID)

# create a coloring vector
cols <- as.character(metadata$taxonomy_color)

# save the tanglegram
pdf("results/phenotypes/tanglegram.pdf", width = 10, height = 10)
tanglegram(dend1, dend2,
           highlight_distinct_edges = TRUE,
           common_subtrees_color_lines = TRUE,
           margin_inner = 2,
           fast = TRUE,
           lwd = 1,
           dLeaf = 0.1,
           type = "r",
           color_lines = cols,
           lab.cex = 0.1)

# save the tanglegram to a file
dev.off()
```

```
## pdf
##   2
```

We next calculate the **cophenetic correlation coefficient (CCC)** between two dendrograms and tests its statistical significance using a permutation test. The CCC measures how well a hierarchical clustering preserves the original pairwise distances between observations.

```r
# function to perform permutation test
permutation_test_cophenetic <- function(dend1, dend2, n_perm = 1000) {

  # step 1: compute observed cophenetic correlation
  observed_cor <- cor_cophenetic(dend1, dend2)

  # step 2: initialize vector to store permuted correlations
  permuted_cor <- numeric(n_perm)

  # step 3: perform permutations
  for (i in seq_len(n_perm)) {
    # shuffle labels in one dendrogram
    dend2_shuffled <- dend2
    labels(dend2_shuffled) <- sample(labels(dend2))

    # compute cophenetic correlation for permuted tree
    permuted_cor[i] <- cor_cophenetic(dend1, dend2_shuffled)
  }

  # step 4: compute p-value
  p_value <- mean(permuted_cor >= observed_cor)

  # step 5: return results
  list(
    observed_correlation = observed_cor,
    permuted_correlations = permuted_cor,
    p_value = p_value
  )
}

# run the permutation test with 1000 permutations
result <- permutation_test_cophenetic(dend1, dend2, n_perm = 1000)

# print the results
print(paste0("Cophenetic correlation coefficient: ", round((result$observed_correlation), digits = 3)))
```

```
## [1] "Cophenetic correlation coefficient: 0.584"
```

```r
print(paste0("p-value: ", result$p_value))
```

```
## [1] "p-value: 0"
```

---

### 9 Representation of reconstructed metabolic pathways in 28 *Bifidobacterium longum* genomes

#### 9.1 Introduction

This block describes the analysis of the representation of various metabolic pathways in 28 *Bifidobacterium longum* genomes. This genomic dataset included 15 genomes from the reference set + 13 additional *Bifidobacterium longum* genomes. For comparative purposes, the 13 additional genomes included isolates of non-human origin, such as type strains of *Bifidobacterium longum* subsp. *suis* and *Bifidobacterium longum* subsp. *suillum*.

##### 9.2 Load data

```
# read the table with BPM (carbohydrate utilization) for 28 B. longum genomes
bpm_28_Blon_df <- read_tsv("data/tables/BPM_28_Blongum_genomes_carbs.txt",
                           col_types = cols(.default = "c")) %>%
                            mutate_at(c(4:71), as.numeric)
# read the table with BPM (other pathways) for 26 B.longum genomes
bpm_28_Blon_df2 <- read_tsv("data/tables/BPM_28_Blongum_genomes_other.txt",
                            col_types = cols(.default = "c")) %>%
                            mutate_at(c(4:32), as.numeric)
# read the table with metadata for predicted carbohydrate utilization phenotypes
phenotype_metadata <- read_tsv("data/tables/phenotype_metadata_carbs.txt",
                               col_types = cols(.default = "c")) %>%
                               filter(!(phenotype %in% c("ManNAc", "GalNAc", "Man", "GalA")))
# read the table with metadata for other pathways
phenotype_metadata_other <- read_tsv("data/tables/phenotype_metadata_other.txt",
                               col_types = cols(.default = "c"))
```

##### 9.3 Hierarchical clustering of the BPM (carbohydrate utilization) of 28 *Bifidobacterium longum* genomes

The following heatmap shows the hierarchical clustering of BPM with the representation of carbohydrate utilization phenotypes predicted for 28 *Bifidobacterium longum* strains.

Hierarchical clustering options:

- **Distance metric**: Hamming distance (equivalent to Manhattan distance for binary data)
- **Linkage method**: Average

```
# extract the binary matrix
bpm_Blon_mat <- as.matrix((bpm_28_Blon_df[, 4:71]))
# add rownames to the matrix
genome_id_Blon <- bpm_28_Blon_df$genome_ID
rownames(bpm_Blon_mat) <- genome_id_Blon

# create vectors with taxonomy
Blon_tax1 <- bpm_28_Blon_df$group
Blon_tax2 <- bpm_28_Blon_df$add_tax
# extract vectors containing data about glycan type and origin
glycan_type <- phenotype_metadata$type_group
glycan_origin <- phenotype_metadata$origin

# create a coloring function
col_fun <- structure(c("white", "#08306b"), names = c("0", "1"))
# create two row annotations specifying taxonomy
ha_Blon1 <- HeatmapAnnotation(
  which = c("row"),
```

```
  Taxonomy1 = Blon_tax1,
  Taxonomy2 = Blon_tax2,
  col = list(Taxonomy1 = c("longum_infantis" = "#c0d28b",
                          "longum_longum" = "#51796f",
                          "longum_suis" = "#00b400",
                          "longum_nov" = "black"),
             Taxonomy2 = c("suis" = "#ffff91",
                          "spp" = "#ffa600",
                          "suillum" = "#8e063c",
                          "iuvenis" = "#00a2ff",
                          "longum" = "white",
                          "infantis" = "white",
                          "nov" = "white")),
             show_annotation_name = FALSE,
             simple_anno_size = unit(3, "mm"))
# create two column annotations specifying glycan type and origin
ha_Blon2 <- HeatmapAnnotation(
  type = glycan_type,
  origin = glycan_origin,
  col = list(type = c("monosaccharides_and_derivatives" = "#E6E7E8",
                      "di_and_oligosaccharides" = "#BCBEC0",
                      "polysaccharides" = "#808285"),
             origin = c("universal" = "#ffd22d",
                        "host (animal)" = "#cbbedd",
                        "plant" = "#8dc63f",
                        "fungal" = "#008aa1",
                        "bacterial" = "#603913")),
             show_annotation_name = FALSE,
             simple_anno_size = unit(3, "mm"))

# plot the heatmap
pdf("results/phenotypes/Blon_carb_heatmap.pdf", width=15, height=10)
ht_Blon_carb <- ComplexHeatmap::Heatmap(bpm_Blon_mat,
                              name = "Predicted phenotype",
                              right_annotation = ha_Blon1,
                              bottom_annotation = ha_Blon2,
                              col = col_fun,
                              clustering_distance_rows = function(m)
                                dist(m, method = "manhattan"),
                              clustering_distance_columns = function(m)
                                dist(m, method = "manhattan"),
                              clustering_method_rows = "average",
                              clustering_method_columns = "average",
                              rect_gp = gpar(col = "grey", lwd = 0.05),
                              show_row_names = TRUE,
                              row_names_gp = gpar(fontsize = 3),
                              column_names_gp = gpar(fontsize = 5),
                              column_names_rot = 60,
                              width = unit(250, "mm"),
                              height = unit(100, "mm"))
draw(ht_Blon_carb)
# save the heatmap to a file
dev.off()
```

```
## pdf
##   2
```

```r
# draw the heatmap again to show it in the compiled markdown file
draw(ht_Blon_carb)
```

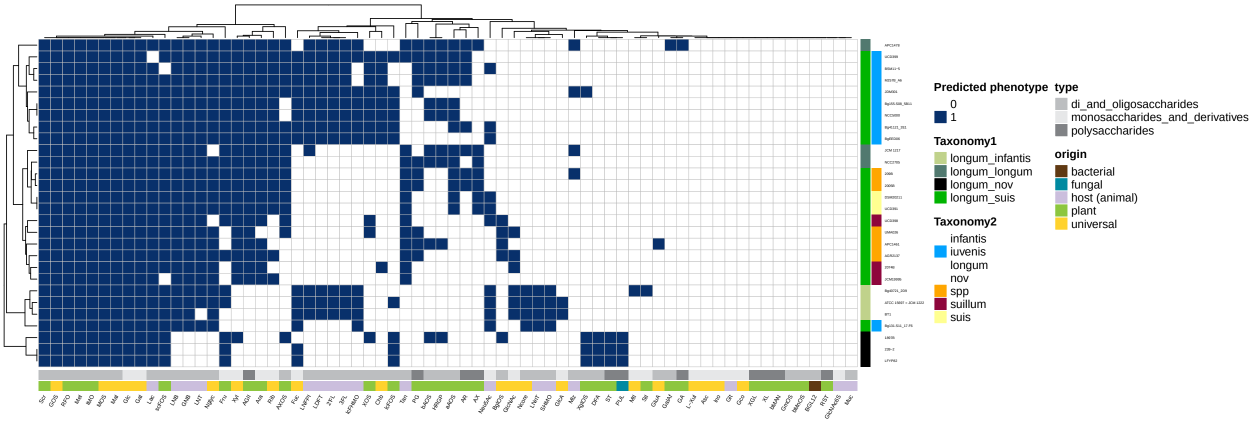

#### 9.4 Hierarchical clustering of the BPM (other pathways) of 28 *Bifidobacterium longum* genomes

The following heatmap shows the hierarchical clustering of BPM with the representation of select metabolic pathways (amino acid/vitamin biosynthesis, urea utilization) predicted in 28 *Bifidobacterium longum* genomes.

Hierarchical clustering options:

- **Distance metric**: Hamming distance (equivalent to Manhattan distance for binary data)
- **Linkage method**: Average

```r
# extract the binary matrix
bpm_Blon_mat2 <- as.matrix((bpm_28_Blon_df2[, 4:32]))
# add rownames to the matrix
genome_id_Blon <- bpm_28_Blon_df2$genome_ID
rownames(bpm_Blon_mat2) <- genome_id_Blon
```

```r
# create a vector with taxonomy (group)
Blon_tax1 <- bpm_28_Blon_df2$group
Blon_tax2 <- bpm_28_Blon_df2$add_tax
# extract vectors containing data about glycan type and origin
pathway_type <- phenotype_metadata_other$type
# create a coloring function
col_fun <- structure(c("white", "#08306b"), names = c("0", "1"))
# create two row annotations specifying taxonomy
ha_Blon1 <- HeatmapAnnotation(
  which = c("row"),
  Taxonomy1 = Blon_tax1,
  Taxonomy2 = Blon_tax2,
  col = list( Taxonomy1 = c("longum_infantis" = "#c0d28b",
                            "longum_longum" = "#51796f",
                            "longum_suis" = "#00b400",
                            "longum_nov" = "black"),
              Taxonomy2 = c("suis" = "#ffff91",
                            "spp" = "#ffa600",
                            "suillum" = "#8e063c",
                            "iuvenis" = "#00a2ff",
                            "longum" = "white",
                            "infantis" = "white",
                            "nov" = "white")),
              show_annotation_name = FALSE,
              simple_anno_size = unit(3, "mm"))
# create a column annotation specifying the pathway type
ha_Blon3 <- HeatmapAnnotation(
  type = pathway_type,
  col = list(type = c("vitamin" = "#8dc63f",
                      "amino_acid" = "#cbbedd",
                      "other" = "#808285")),
  show_annotation_name = FALSE,
  simple_anno_size = unit(3, "mm"))

# plot the heatmap
pdf("results/phenotypes/Blon_vit_heatmap.pdf", width=15, height=10)
ht_Blon_vit <- ComplexHeatmap::Heatmap(bpm_Blon_mat2,
                              name = "Predicted phenotype",
                              bottom_annotation = ha_Blon3,
                              right_annotation = ha_Blon1,
                              col = col_fun,
                              clustering_distance_rows = function(m)
                                dist(m, method = "manhattan"),
                              clustering_distance_columns = function(m)
                                dist(m, method = "manhattan"),
                              clustering_method_rows = "average",
                              clustering_method_columns = "average",
                              rect_gp = gpar(col = "grey", lwd = 0.05),
                              show_row_names = TRUE,
                              row_names_gp = gpar(fontsize = 3),
                              column_names_gp = gpar(fontsize = 5),
                              column_names_rot = 60,
                              width = unit(100, "mm"),
```

```r
                                   height = unit(80, "mm"))
draw(ht_Blon_vit)
# save the heatmap to a file
dev.off()
```

```
## pdf
##   2
```

```r
# draw the heatmap again to show it in the compiled markdown file
draw(ht_Blon_vit)
```

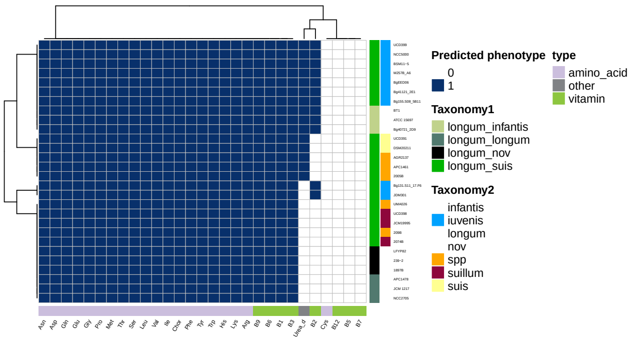

### 10 Representation of predicted metabolic pathways in 3083 *Bifidobacterium* genomes

#### 10.1 Introduction

This section describes various analyses of two binary BPM for 3083 non-redundant *Bifidobacterium* genomes. This genomic dataset was assembled by merging 263 reference and 2820 additional genomes.

1. The first BPM depicts the presence/absence of carbohydrate utilization pathways
2. The second BPM depicts the presence/absence of biosynthetic pathways + urea utilization

#### 10.2 Load data

```r
# read BPM (carbs) for 263 reference genomes
bpm_263_df <- read_tsv("data/tables/BPM_263_NR_genomes_carbs.txt",
                       col_types = cols(.default = "c")) %>%
                       mutate_at(c(4:75), as.numeric) %>%
                       dplyr::select(-c(ManNAc, GalNAc, Man, GalA)) %>%
                       arrange(genome_ID)
# read BPM (carbs) for additional genomes
bpm_2820_df <- read_tsv("data/tables/BPM_2820_NR_genomes_carbs.txt",
                        col_types = cols(.default = "c")) %>%
                        mutate_at(c(3:70), as.numeric) %>%
                        arrange(genome_ID)
# read the table with metadata for carbohydrate utilization pathways
phenotype_metadata <- read_tsv("data/tables/phenotype_metadata_carbs.txt",
                      col_types = cols(.default = "c")) %>%
                      filter(!(phenotype %in% c("ManNAc", "GalNAc", "Man", "GalA")))
# extract BPM for 263 genomes for merging
bpm_263_carb_df <- bpm_263_df %>%
  dplyr::select(genome_ID, curated_taxonomy, c(Glc:`L-Xul`))
# extract BPM for 2820 genomes for merging
bpm_2820_carb_df <- bpm_2820_df %>%
  dplyr::select(genome_ID, curated_taxonomy, c(Glc:`L-Xul`))
# check that column names in both BPMs are identical
colnames(bpm_263_carb_df) == colnames(bpm_2820_carb_df)
```

```
##  [1] TRUE TRUE TRUE TRUE TRUE TRUE TRUE TRUE TRUE TRUE TRUE TRUE TRUE TRUE TRUE
## [16] TRUE TRUE TRUE TRUE TRUE TRUE TRUE TRUE TRUE TRUE TRUE TRUE TRUE TRUE TRUE
## [31] TRUE TRUE TRUE TRUE TRUE TRUE TRUE TRUE TRUE TRUE TRUE TRUE TRUE TRUE TRUE
## [46] TRUE TRUE TRUE TRUE TRUE TRUE TRUE TRUE TRUE TRUE TRUE TRUE TRUE TRUE TRUE
## [61] TRUE TRUE TRUE TRUE TRUE TRUE TRUE TRUE TRUE TRUE
```

```r
# merge the BPMs
bpm_3083_carb_df <- as_tibble(merge(bpm_263_carb_df, bpm_2820_carb_df, all=TRUE) %>%
  arrange(genome_ID))
# read the table with BPM for 3083 genomes with predicted presence/absence of biosynthetic pathways
bpm_3083_other_df <- read_xlsx("data/tables/BPM_3083_genomes_other.xlsx") %>%
                     mutate_at(c(3:31), as.numeric) %>%
                     arrange(genome_ID)
# read the table with metadata for biosynthetic pathways
phenotype_metadata_other <- read_tsv("data/tables/phenotype_metadata_other.txt",
                                     col_types = cols(.default = "c"))
```

#### 10.3 Set colors

Defines point colors and shapes used throughout this block.

```r
# arguments for scale_shape_manual
genomes_3083_shapes <- c(21,24,24,21,21,
                         21,21,24,23,21,
                         21,21,22,21,21,
                         21,21,21,24,22,
                         21,23,25)
# arguments for scale_fill_manual
```

```
# breaks (how groups are encoded in the table)
genomes_3083_breaks <- c("Bifidobacterium adolescentis",
                         "Bifidobacterium angulatum",
                         "Bifidobacterium animalis subsp. animalis",
                         "Bifidobacterium animalis subsp. lactis",
                         "Bifidobacterium bifidum",
                         "Bifidobacterium breve",
                         "Bifidobacterium catenulatum subsp. catenulatum",
                         "Bifidobacterium catenulatum subsp. kashiwanohense",
                         "Bifidobacterium catenulatum subsp. kashiwanohense_A",
                         "Bifidobacterium dentium",
                         "Bifidobacterium gallicum",
                         "Bifidobacterium globosum",
                         "Bifidobacterium hominis",
                         "Bifidobacterium longum subsp. infantis",
                         "Bifidobacterium longum subsp. longum",
                         "Bifidobacterium longum subsp. nov.",
                         "Bifidobacterium longum subsp. suis",
                         "Bifidobacterium pseudocatenulatum",
                         "Bifidobacterium pullorum",
                         "Bifidobacterium ruminantium",
                         "Bifidobacterium scardovii",
                         "Bifidobacterium thermophilum",
                         "Bifidobacterium tsurumiense")
# values (color codes)
genomes_3083_colors <- c("#9b91c1", # Bifidobacterium adolescentis
                         "#ffff7f", # Bifidobacterium angulatum
                         "grey" , # Bifidobacterium animalis subsp. animalis
                         "grey", # Bifidobacterium animalis subsp. lactis
                         "#222c80", # Bifidobacterium bifidum
                         "#00a2ff", # Bifidobacterium breve
                         "tomato2", # Bifidobacterium catenulatum subsp. catenulatum
                         "tomato2", # Bifidobacterium catenulatum subsp. kashiwanohense
                         "tomato2", # Bifidobacterium catenulatum subsp. kashiwanohense_A
                         "#8e063c", # Bifidobacterium dentium
                         "#ffffff", # Bifidobacterium gallicum
                         "#d0a079", # Bifidobacterium globosum
                         "#a2d2de", # Bifidobacterium hominis
                         "#c0d28b", # Bifidobacterium longum subsp. infantis
                         "#51796f", # Bifidobacterium longum subsp. longum
                         "black", # Bifidobacterium longum subsp. nov.
                         "#00b400", # Bifidobacterium longum subsp. suis
                         "#ffa600", # Bifidobacterium pseudocatenulatum
                         "#ffffff", # Bifidobacterium pullorum
                         "#ffffff", # Bifidobacterium ruminantium
                         "#f79cd4", # Bifidobacterium scardovii
                         "#ffffff", # Bifidobacterium thermophilum
                         "#ffffff") # Bifidobacterium tsurumiense
# labels (what will appear in the legend)
genomes_3083_species <- c("B. adolescentis",
                          "B. angulatum",
                          "Ba. animalis",
                          "Ba. lactis",
```

```
                          "B. bifidum",
                          "B. breve",
                          "Bc. catenulatum",
                          "Bc. kashiwanohense",
                          "Bc. kashiwanohense_A",
                          "B. dentium",
                          "B. gallicum",
                          "B. globosum",
                          "B. hominis",
                          "Bl. infantis",
                          "Bl. longum",
                          "Bl. nov.",
                          "Bl. suis",
                          "B. pseudocatenulatum",
                          "B. pullorum",
                          "B. ruminantium",
                          "B. scardovii",
                          "B. thermophilum",
                          "B. tsurumiense")
```

#### 10.4 Statistical analysis

The code chunks below describe various statistical analyses.

##### 10.4.1 PERMANOVA

We used the Hamming distance (equivalent to Manhattan distance for binary data) to create a distance matrix from the BPM. We then performed a PERMANOVA (using the adonis2 function) to assess the effect of taxonomy on the constructed dissimilarity matrix (differences in group centroids).

```
# extract the binary matrix
bpm_3083_mat <- as.matrix((bpm_3083_carb_df[, 3:70]))
# calculate Hamming distance
bpm_3083_mat_dist <- vegdist(bpm_3083_mat, method ="manhattan")
# create a vector with taxonomy
taxonomy <- as.factor(bpm_3083_carb_df$curated_taxonomy)
# perform PERMANOVA
permanova_result <- adonis2(bpm_3083_mat_dist ~ taxonomy, permutations = 999, parallel = 8)
permanova_result
```

```
## Permutation test for adonis under reduced model
## Permutation: free
## Number of permutations: 999
##
## adonis2(formula = bpm_3083_mat_dist ~ taxonomy, permutations = 999, parallel = 8)
##            Df SumOfSqs      R2      F Pr(>F)
## Model      22   636433 0.90932 1394.7  0.001 ***
## Residual 3060    63470 0.09068
## Total    3082   699903 1.00000
## ---
## Signif. codes:  0 '***' 0.001 '**' 0.01 '*' 0.05 '.' 0.1 ' ' 1
```

The PERMANOVA results show a significant effect of taxonomy on the dissimilarity matrix (bpm_3083_mat_dist), explaining 91% of the variation ($R^2 = 0.91$, $F = 1394.7$, $p = 0.001$), indicating that taxonomy significantly influences the differences observed in the data.

##### 10.4.2 Homogeneity of multivariate dispersions

We assessed the homogeneity of multivariate dispersions using the betadisper function followed by a permutation test with permutest.

```
betadisper_result <- betadisper(bpm_3083_mat_dist, taxonomy)
permutest(betadisper_result)
```

```
##
## Permutation test for homogeneity of multivariate dispersions
## Permutation: free
## Number of permutations: 999
##
## Response: Distances
##             Df  Sum Sq Mean Sq      F N.Perm Pr(>F)
## Groups      22  7339.3  333.60 81.056    999  0.001 ***
## Residuals 3060 12594.1    4.12
## ---
## Signif. codes:  0 '***' 0.001 '**' 0.01 '*' 0.05 '.' 0.1 ' ' 1
```

The test for homogeneity of multivariate dispersions shows a significant difference between groups ($F = 81.056$, $p = 0.001$), indicating that the variation in dispersion among taxonomic groups is not equal. This result, combined with the significant effect of taxonomy in the PERMANOVA analysis, suggests that the observed differences in the dissimilarity matrix are influenced by both the central tendency and dispersion of the groups.

#### 10.5 Ordination

Ordination techniques summarize the data in a reduced number of dimensions while accounting for as much of the variability in the original data set as possible. Here, we use Non-metric MultiDimensional Scaling (NMDS) to visualize the level of similarity or dissimilarity between genomes based on a Hamming distance matrix.

```
# extract the binary matrix
bpm_3083_mat <- as.matrix((bpm_3083_carb_df[, 3:70]))
# do NMDS
nmds <- metaMDS(bpm_3083_mat,
             autotransform = FALSE,
             distance = "manhattan",
             engine = "monoMDS",
             k = 2,
             weakties = TRUE,
             model = "global",
             maxit = 300,
             try = 40,
             trymax = 100,
             parallel = 8)
```

```
## Run 0 stress 0.1068156
## Run 1 stress 0.1067765
## ... New best solution
## ... Procrustes: rmse 0.000345819  max resid 0.01242866
## Run 2 stress 0.1070227
## ... Procrustes: rmse 0.0007324424  max resid 0.01472983
## Run 3 stress 0.10717
## ... Procrustes: rmse 0.0009343673  max resid 0.02535644
## Run 4 stress 0.1069523
## ... Procrustes: rmse 0.0006104684  max resid 0.02527891
## Run 5 stress 0.1070437
## ... Procrustes: rmse 0.0007570445  max resid 0.02530916
## Run 6 stress 0.1068285
## ... Procrustes: rmse 0.0007031646  max resid 0.02816589
## Run 7 stress 0.1067965
## ... Procrustes: rmse 0.0008025511  max resid 0.03279926
## Run 8 stress 0.1069717
## ... Procrustes: rmse 0.0006504716  max resid 0.02525668
## Run 9 stress 0.1071304
## ... Procrustes: rmse 0.0006842554  max resid 0.02649743
## Run 10 stress 0.1070664
## ... Procrustes: rmse 0.0006744269  max resid 0.0252449
## Run 11 stress 0.1067813
## ... Procrustes: rmse 0.0007798291  max resid 0.03280343
## Run 12 stress 0.1072481
## ... Procrustes: rmse 0.0008822679  max resid 0.02647825
## Run 13 stress 0.1067719
## ... New best solution
## ... Procrustes: rmse 0.0007506487  max resid 0.03281488
## Run 14 stress 0.1069222
## ... Procrustes: rmse 0.0008841395  max resid 0.02525274
## Run 15 stress 0.1067673
## ... New best solution
## ... Procrustes: rmse 0.0006452657  max resid 0.02513746
## Run 16 stress 0.1069514
## ... Procrustes: rmse 0.0007230837  max resid 0.03275046
## Run 17 stress 0.1067092
## ... New best solution
## ... Procrustes: rmse 0.000588934  max resid 0.02518269
## Run 18 stress 0.1068028
## ... Procrustes: rmse 0.0006770257  max resid 0.03274255
## Run 19 stress 0.1077001
## Run 20 stress 0.1079722
## Run 21 stress 0.1068383
## ... Procrustes: rmse 0.0007039633  max resid 0.02523024
## Run 22 stress 0.1070097
## ... Procrustes: rmse 0.0007514134  max resid 0.02520312
## Run 23 stress 0.1069948
## ... Procrustes: rmse 0.0008476902  max resid 0.02529314
## Run 24 stress 0.1068961
## ... Procrustes: rmse 0.0006458196  max resid 0.02526049
## Run 25 stress 0.1068927
## ... Procrustes: rmse 0.0007790491  max resid 0.02521741
## Run 26 stress 0.1071495
```

```
## ... Procrustes: rmse 0.0008940186  max resid 0.03260698
## Run 27 stress 0.1068757
## ... Procrustes: rmse 0.0008298531  max resid 0.03255685
## Run 28 stress 0.1071274
## ... Procrustes: rmse 0.001044566  max resid 0.0323554
## Run 29 stress 0.1072105
## Run 30 stress 0.1069192
## ... Procrustes: rmse 0.0008333777  max resid 0.03296947
## Run 31 stress 0.1069658
## ... Procrustes: rmse 0.0007397569  max resid 0.02529256
## Run 32 stress 0.1071389
## ... Procrustes: rmse 0.0006726747  max resid 0.02380715
## Run 33 stress 0.1070565
## ... Procrustes: rmse 0.001024646  max resid 0.03272374
## Run 34 stress 0.1068863
## ... Procrustes: rmse 0.0008467185  max resid 0.03258288
## Run 35 stress 0.1067885
## ... Procrustes: rmse 0.000637455  max resid 0.02516002
## Run 36 stress 0.1068952
## ... Procrustes: rmse 0.0007915687  max resid 0.03252945
## Run 37 stress 0.1071111
## ... Procrustes: rmse 0.0008945946  max resid 0.02846407
## Run 38 stress 0.106981
## ... Procrustes: rmse 0.0008387633  max resid 0.0252669
## Run 39 stress 0.1068978
## ... Procrustes: rmse 0.0007797103  max resid 0.03277927
## Run 40 stress 0.1070159
## ... Procrustes: rmse 0.0008025884  max resid 0.02529039
## Run 41 stress 0.1068518
## ... Procrustes: rmse 0.0006398479  max resid 0.02522513
## Run 42 stress 0.1067993
## ... Procrustes: rmse 0.0007550324  max resid 0.03295501
## Run 43 stress 0.1070936
## ... Procrustes: rmse 0.001092521  max resid 0.0324759
## Run 44 stress 0.1070023
## ... Procrustes: rmse 0.0009254978  max resid 0.03263762
## Run 45 stress 0.1070092
## ... Procrustes: rmse 0.0009898692  max resid 0.03250282
## Run 46 stress 0.1068999
## ... Procrustes: rmse 0.0008627861  max resid 0.03255762
## Run 47 stress 0.1068145
## ... Procrustes: rmse 0.0006379109  max resid 0.02521268
## Run 48 stress 0.1070278
## ... Procrustes: rmse 0.0009069553  max resid 0.03250932
## Run 49 stress 0.1069662
## ... Procrustes: rmse 0.0007415256  max resid 0.02528757
## Run 50 stress 0.1071767
## ... Procrustes: rmse 0.001051709  max resid 0.03275517
## Run 51 stress 0.10692
## ... Procrustes: rmse 0.0008543258  max resid 0.03300534
## Run 52 stress 0.1067962
## ... Procrustes: rmse 0.0005511742  max resid 0.02520728
## Run 53 stress 0.1070895
## ... Procrustes: rmse 0.000962858  max resid 0.02532108
```

```
## Run 54 stress 0.1068755
## ... Procrustes: rmse 0.000830897  max resid 0.0327425
## Run 55 stress 0.1068895
## ... Procrustes: rmse 0.0007826196  max resid 0.02521194
## Run 56 stress 0.1072326
## Run 57 stress 0.1069032
## ... Procrustes: rmse 0.0006527357  max resid 0.02522734
## Run 58 stress 0.1068327
## ... Procrustes: rmse 0.0007396511  max resid 0.03303277
## Run 59 stress 0.1068801
## ... Procrustes: rmse 0.0006609969  max resid 0.02522835
## Run 60 stress 0.1068181
## ... Procrustes: rmse 0.000705348  max resid 0.032891
## Run 61 stress 0.1067931
## ... Procrustes: rmse 0.0007318363  max resid 0.03274713
## Run 62 stress 0.1071157
## ... Procrustes: rmse 0.000898514  max resid 0.02534741
## Run 63 stress 0.1070093
## ... Procrustes: rmse 0.0008801927  max resid 0.03260692
## Run 64 stress 0.1069993
## ... Procrustes: rmse 0.0009229578  max resid 0.03243356
## Run 65 stress 0.1076212
## Run 66 stress 0.1068626
## ... Procrustes: rmse 0.0007406359  max resid 0.0252171
## Run 67 stress 0.1071141
## ... Procrustes: rmse 0.001003329  max resid 0.03232227
## Run 68 stress 0.1070032
## ... Procrustes: rmse 0.000743672  max resid 0.02533762
## Run 69 stress 0.1071254
## ... Procrustes: rmse 0.001079363  max resid 0.03239105
## Run 70 stress 0.106993
## ... Procrustes: rmse 0.0009108261  max resid 0.03255979
## Run 71 stress 0.1070954
## ... Procrustes: rmse 0.00103295  max resid 0.03254009
## Run 72 stress 0.106793
## ... Procrustes: rmse 0.000639171  max resid 0.02513301
## Run 73 stress 0.1071278
## ... Procrustes: rmse 0.000895946  max resid 0.03255381
## Run 74 stress 0.106773
## ... Procrustes: rmse 0.0006011248  max resid 0.02517211
## Run 75 stress 0.1070667
## ... Procrustes: rmse 0.001050155  max resid 0.03266149
## Run 76 stress 0.1068791
## ... Procrustes: rmse 0.0008282676  max resid 0.03285743
## Run 77 stress 0.1069022
## ... Procrustes: rmse 0.0007184167  max resid 0.02523543
## Run 78 stress 0.1082185
## Run 79 stress 0.1067042
## ... New best solution
## ... Procrustes: rmse 0.0004874937  max resid 0.01418561
## Run 80 stress 0.1068991
## ... Procrustes: rmse 0.000784628  max resid 0.02544023
## Run 81 stress 0.1072625
## Run 82 stress 0.1070689
```

```
## ... Procrustes: rmse 0.0009622589  max resid 0.02555319
## Run 83 stress 0.1069394
## ... Procrustes: rmse 0.0007871437  max resid 0.02548093
## Run 84 stress 0.1067904
## ... Procrustes: rmse 0.0005629986  max resid 0.02540402
## Run 85 stress 0.1069181
## ... Procrustes: rmse 0.0008341064  max resid 0.032625
## Run 86 stress 0.1071369
## ... Procrustes: rmse 0.001116186  max resid 0.0323682
## Run 87 stress 0.1068506
## ... Procrustes: rmse 0.0006310524  max resid 0.0254377
## Run 88 stress 0.1068966
## ... Procrustes: rmse 0.0007757285  max resid 0.03266259
## Run 89 stress 0.1070438
## ... Procrustes: rmse 0.0009628582  max resid 0.03241514
## Run 90 stress 0.1070195
## ... Procrustes: rmse 0.001000309  max resid 0.03240402
## Run 91 stress 0.1069218
## ... Procrustes: rmse 0.0008218559  max resid 0.02549739
## Run 92 stress 0.1069364
## ... Procrustes: rmse 0.0007727137  max resid 0.02545627
## Run 93 stress 0.1070447
## ... Procrustes: rmse 0.0009450014  max resid 0.02553963
## Run 94 stress 0.1068966
## ... Procrustes: rmse 0.0008279078  max resid 0.03270984
## Run 95 stress 0.1069
## ... Procrustes: rmse 0.0008573983  max resid 0.03261177
## Run 96 stress 0.1068209
## ... Procrustes: rmse 0.0007037664  max resid 0.03281889
## Run 97 stress 0.1071417
## ... Procrustes: rmse 0.0009186363  max resid 0.02647573
## Run 98 stress 0.1068998
## ... Procrustes: rmse 0.0008279388  max resid 0.03273516
## Run 99 stress 0.1070131
## ... Procrustes: rmse 0.0009140001  max resid 0.02551837
## Run 100 stress 0.1067981
## ... Procrustes: rmse 0.0006686262  max resid 0.03278236
## Run 101 stress 0.1069905
## ... Procrustes: rmse 0.0009526386  max resid 0.03241314
## Run 102 stress 0.1072098
## Run 103 stress 0.1068557
## ... Procrustes: rmse 0.0006728743  max resid 0.02545099
## Run 104 stress 0.1069312
## ... Procrustes: rmse 0.0009192531  max resid 0.03248516
## *** Best solution was not repeated -- monoMDS stopping criteria:
##    104: scale factor of the gradient < sfgrmin
```

```r
# extract NMDS results to a tibble
nmds.df <- as_tibble(nmds$points)

# plot the NMDS results
# create a vector with taxonomy
taxonomy <- as.factor(bpm_3083_carb_df$curated_taxonomy)
# create a vector with genome names
```

```r
IDs <- bpm_3083_carb_df$genome_ID
# select genomes that will be marked by text
genomes_short <- dplyr::case_when(
  IDs == "1695.38" ~ "Bg131.S11_17.F6",
  IDs == "630129.38" ~ "Bg42221_1E1",
  IDs == "M56B_1C3" ~ "M56B_1C3",
  IDs == "Map_139_001" ~ "Map_139_001",
  TRUE ~ NA_character_
)
# plot
ggplot(nmds.df) +
  aes(x=MDS1, y=MDS2, fill=taxonomy, shape = taxonomy) +
  geom_point(size=2.5) +
  scale_shape_manual(values=genomes_3083_shapes) +
  guides(shape="none") +
  scale_fill_manual(name = "Taxonomy",
                    breaks=genomes_3083_breaks,
                    values=genomes_3083_colors,
                    labels=genomes_3083_species) +
  guides(fill = guide_legend(override.aes=list(shape=genomes_3083_shapes))) +
  geom_text_repel(aes(label = genomes_short), size = 3, fontface=1, color="black",
                  min.segment.length = 0, seed = 20, box.padding = 1,
                  max.overlaps = 100) +
  labs(title= "NMDS") +
  coord_fixed(1) +
  theme_bw() +
  theme(plot.title = element_text(face="bold"))
```

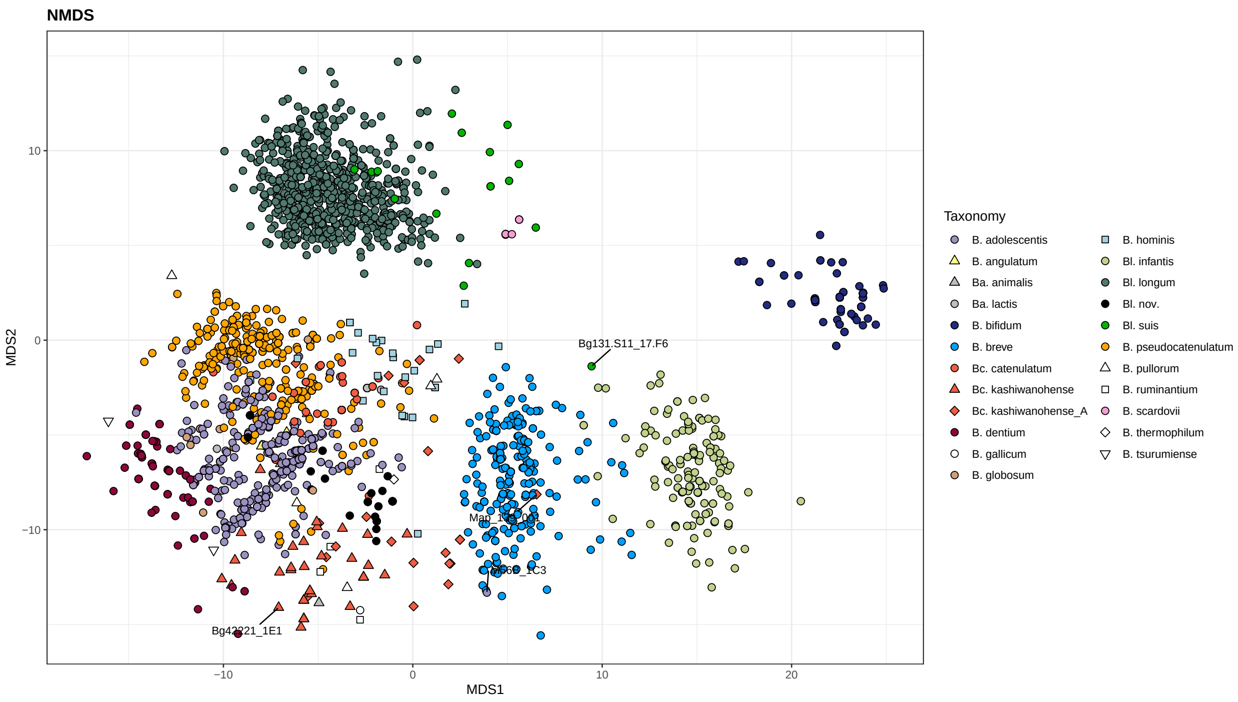

```r
# save the file
ggsave("results/phenotypes/bpm_3083_NMDS_Hamming_weak_TRUE.pdf", width = 14, height = 10)
```

The stress value achieved was 0.107, indicating a good fit of the 2-dimensional model to the data, with stress type 1 and weak ties applied.

Alternative visualization: show centroids + lines connecting centroids with individual points ("spiders").

```r
# plot ordination using gg_ordiplot
# this function creates several data.frames
nmds_ord <- gg_ordiplot(ord = nmds,
                        groups = taxonomy,
                        ellipse = FALSE,
                        hull = FALSE,
                        spiders = TRUE,
                        plot = FALSE)
# save the data frame with spiders
nmds_spiders.df <- nmds_ord$df_spiders

# plot
ggplot(nmds_spiders.df) +
  geom_segment(aes(x=cntr.x, y=cntr.y,
                   xend=x, yend=y,
                   color = Group),
                   size = 0.4,
```

```r
                    alpha = 0.5) +
  geom_point(aes(x=cntr.x, cntr.y,
                 fill = Group,
                 shape = Group),
                 size = 4) +
  scale_color_manual(breaks=genomes_3083_breaks,
                     values=genomes_3083_colors,
                     labels=genomes_3083_species) +
  scale_shape_manual(values=genomes_3083_shapes) +
  guides(shape="none") +
  scale_fill_manual(name = "Taxonomy",
                    breaks=genomes_3083_breaks,
                    values=genomes_3083_colors,
                    labels=genomes_3083_species) +
  guides(fill = guide_legend(override.aes=list(shape=genomes_3083_shapes))) +
  labs(title= "NMDS") +
  coord_fixed(1) +
  theme_bw() +
  theme(plot.title = element_text(face="bold"))
```

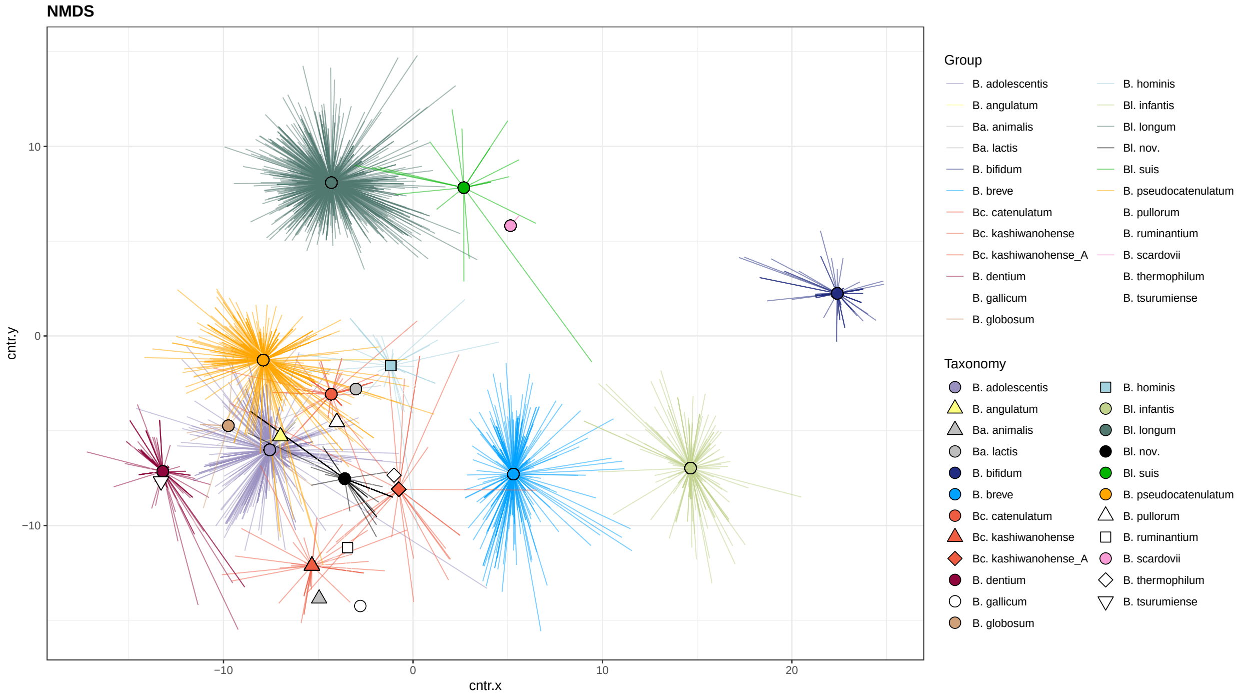

```r
# save the file
ggsave("results/phenotypes/bpm_3083_NMDS_Hamming_spiders.pdf", width = 14, height = 10)
```

#### 10.6 Predicted phenotypic richness

Here, we calculate predicted phenotypic richness, i.e., the total number of different carbohydrate utilization pathways predicted in each genome.

```
# create a new tibble
phenotype_richness <- bpm_3083_carb_df %>%
  mutate(curated_taxonomy = factor(curated_taxonomy, levels = rev(genomes_3083_breaks))) %>%
  group_by(genome_ID, curated_taxonomy) %>%
  summarize(total_phenotypes = sum(across(Glc:`L-Xul`)))

# plot boxplot + swarmplot for each species/subspecies
ggplot() +
  geom_boxplot(data = phenotype_richness,
               mapping = aes(y = curated_taxonomy, x = total_phenotypes),
               outlier.shape = NA, width=0.9, lwd = 0.3) +
  ggbeeswarm::geom_quasirandom(data = phenotype_richness,
                               aes(y=curated_taxonomy, x=total_phenotypes,
                                   fill=curated_taxonomy, shape=curated_taxonomy),
                               color = "black", stroke = 0.1, size = 4) +
  scale_shape_manual(values=rev(genomes_3083_shapes)) +
  guides(shape="none") +
  scale_fill_manual(name = "Taxonomy",
                    breaks=rev(c(genomes_3083_breaks)),
                    values=rev(c(genomes_3083_colors)),
                    labels=rev(c(genomes_3083_species))) +
  guides(fill = guide_legend(override.aes=list(shape=rev(genomes_3083_shapes)))) +
  theme_bw() +
  theme(plot.title = element_text(face="bold"),
        axis.title = element_text(color = "black"),
        axis.text = element_text(color = "black"),
        axis.text.y = element_blank(),
        axis.ticks.y = element_blank()) +
  labs(y = "", x = "Predicted phenotypic richness") +
  scale_x_continuous(breaks = seq(0, max(phenotype_richness$total_phenotypes), by = 5))
```

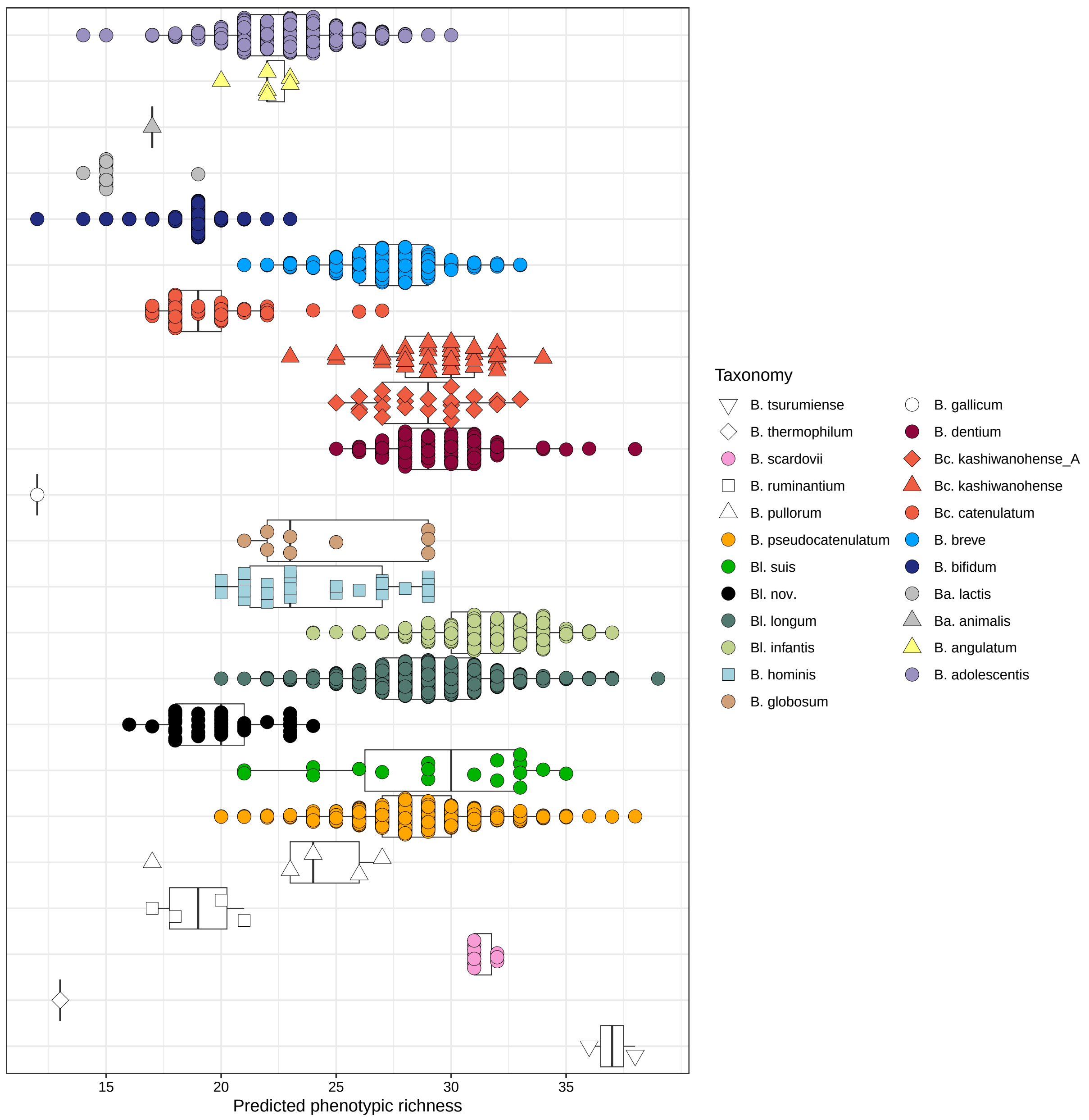

```
# save the plot to a file
ggsave("results/phenotypes/bpm_3083_phenotypic_richness.pdf", width = 10, height = 12)
```

##### 10.6.1 Statistics for predicted phenotypic richness: generalized linear model (GLM)

Here, we compare the predicted phenotypic richness of different *Bifidobacterum* clades using a generalized linear model (GLM) that assumes Poisson distribution. The phenotypic richness of *B. longum* subsp. *nov.* serves as a reference group.

```
# build a glm model
phenotype_richness$curated_taxonomy <- as.factor(phenotype_richness$curated_taxonomy)
phenotype_richness$curated_taxonomy <- relevel(phenotype_richness$curated_taxonomy,
                                               ref = "Bifidobacterium longum subsp. nov.")
```

```r
glm_result <- glm(total_phenotypes ~ curated_taxonomy, data = phenotype_richness, family = poisson)
# check the model
par(mfrow = c(2, 2))
plot(glm_result)
```

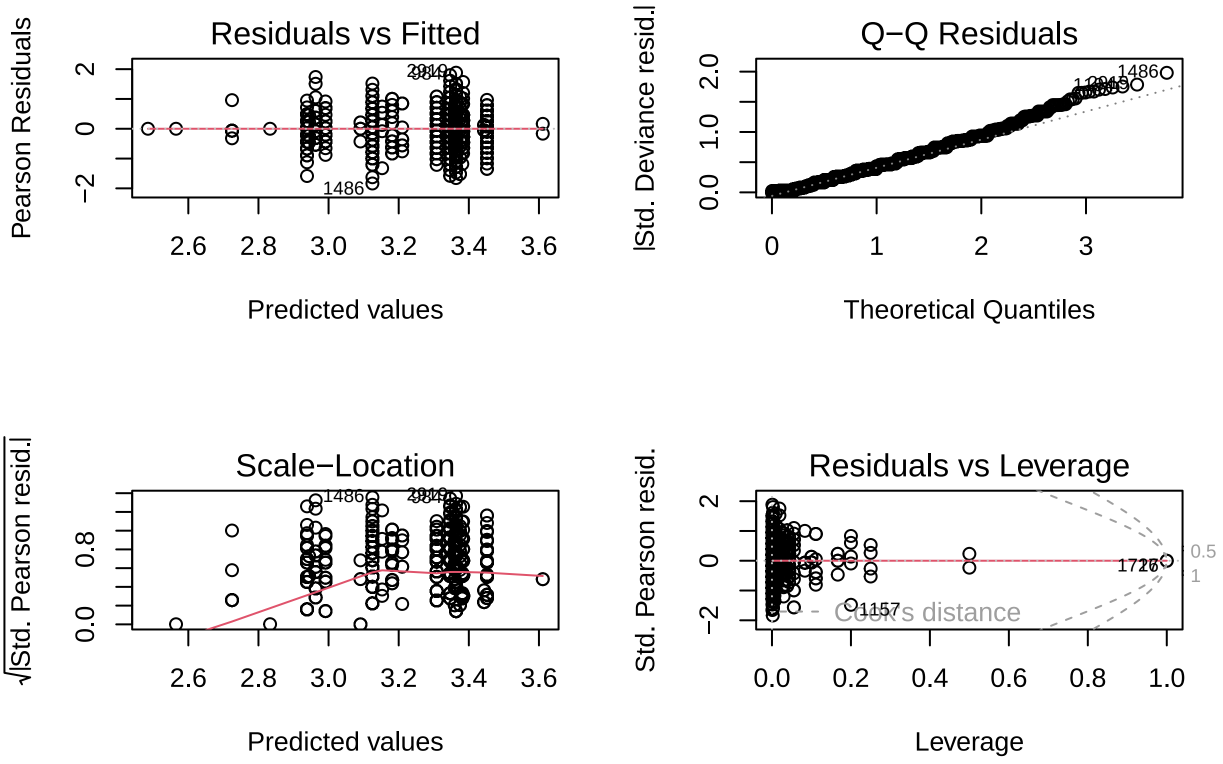

1. The residuals are randomly scattered around zero (but slightly shifted to the right)
2. The residuals are normally distributed
3. No extreme heteroscedasticity
4. One strong outlier based on Cook's distance (but kept for the analysis)

Overall, the Poisson regression model seems good; however, a check for overdispersion is required:

```r
dispersion_test <- dispersiontest(glm_result)
print(dispersion_test)
```

```
##
##  Overdispersion test
##
## data:  glm_result
## z = -122.62, p-value = 1
## alternative hypothesis: true dispersion is greater than 1
## sample estimates:
## dispersion
##  0.2044217
```

The Poisson regression model does not exhibit overdispersion. In fact, it shows underdispersion with a dispersion parameter significantly less than 1. This means that the Poisson model is appropriate for our data, and we do not need to consider alternative models like the quasi-Poisson or negative binomial models to account for overdispersion. Multicollinearity is not an issue here, so we conclude that the model is appropriate for our data.

```r
summary(glm_result)
```

```
##
## Call:
## glm(formula = total_phenotypes ~ curated_taxonomy, family = poisson,
##     data = phenotype_richness)
##
## Coefficients:
##                                                                   Estimate
## (Intercept)                                                        2.99131
## curated_taxonomyBifidobacterium tsurumiense                        0.61961
## curated_taxonomyBifidobacterium thermophilum                      -0.42636
## curated_taxonomyBifidobacterium scardovii                          0.45231
## curated_taxonomyBifidobacterium ruminantium                       -0.04687
## curated_taxonomyBifidobacterium pullorum                           0.16143
## curated_taxonomyBifidobacterium pseudocatenulatum                  0.35545
## curated_taxonomyBifidobacterium longum subsp. suis                 0.38362
## curated_taxonomyBifidobacterium longum subsp. longum               0.37228
## curated_taxonomyBifidobacterium longum subsp. infantis             0.46048
## curated_taxonomyBifidobacterium hominis                            0.18994
## curated_taxonomyBifidobacterium globosum                           0.21864
## curated_taxonomyBifidobacterium gallicum                          -0.50640
## curated_taxonomyBifidobacterium dentium                            0.39244
## curated_taxonomyBifidobacterium catenulatum subsp. kashiwanohense_A  0.36846
## curated_taxonomyBifidobacterium catenulatum subsp. kashiwanohense    0.39008
## curated_taxonomyBifidobacterium catenulatum subsp. catenulatum    -0.02818
## curated_taxonomyBifidobacterium breve                              0.31699
## curated_taxonomyBifidobacterium bifidum                           -0.05275
## curated_taxonomyBifidobacterium animalis subsp. lactis            -0.26673
## curated_taxonomyBifidobacterium animalis subsp. animalis          -0.15810
## curated_taxonomyBifidobacterium angulatum                          0.09973
## curated_taxonomyBifidobacterium adolescentis                       0.13374
##                                                                   Std. Error
## (Intercept)                                                          0.03843
## curated_taxonomyBifidobacterium tsurumiense                          0.12244
## curated_taxonomyBifidobacterium thermophilum                         0.28000
## curated_taxonomyBifidobacterium scardovii                            0.06835
## curated_taxonomyBifidobacterium ruminantium                          0.12098
## curated_taxonomyBifidobacterium pullorum                             0.10012
## curated_taxonomyBifidobacterium pseudocatenulatum                    0.03957
## curated_taxonomyBifidobacterium longum subsp. suis                   0.05812
## curated_taxonomyBifidobacterium longum subsp. longum                 0.03890
## curated_taxonomyBifidobacterium longum subsp. infantis               0.04096
## curated_taxonomyBifidobacterium hominis                              0.05545
## curated_taxonomyBifidobacterium globosum                             0.07721
## curated_taxonomyBifidobacterium gallicum                             0.29122
## curated_taxonomyBifidobacterium dentium                              0.04239
## curated_taxonomyBifidobacterium catenulatum subsp. kashiwanohense_A    0.05466
```

```
## curated_taxonomyBifidobacterium catenulatum subsp. kashiwanohense      0.04977
## curated_taxonomyBifidobacterium catenulatum subsp. catenulatum         0.04952
## curated_taxonomyBifidobacterium breve                                  0.03995
## curated_taxonomyBifidobacterium bifidum                                0.04007
## curated_taxonomyBifidobacterium animalis subsp. lactis                 0.08332
## curated_taxonomyBifidobacterium animalis subsp. animalis               0.24556
## curated_taxonomyBifidobacterium angulatum                              0.09515
## curated_taxonomyBifidobacterium adolescentis                           0.03956
##                                                                     z value
## (Intercept)                                                          77.832
## curated_taxonomyBifidobacterium tsurumiense                           5.061
## curated_taxonomyBifidobacterium thermophilum                         -1.523
## curated_taxonomyBifidobacterium scardovii                             6.617
## curated_taxonomyBifidobacterium ruminantium                          -0.387
## curated_taxonomyBifidobacterium pullorum                              1.612
## curated_taxonomyBifidobacterium pseudocatenulatum                     8.983
## curated_taxonomyBifidobacterium longum subsp. suis                    6.600
## curated_taxonomyBifidobacterium longum subsp. longum                  9.571
## curated_taxonomyBifidobacterium longum subsp. infantis               11.242
## curated_taxonomyBifidobacterium hominis                               3.426
## curated_taxonomyBifidobacterium globosum                              2.832
## curated_taxonomyBifidobacterium gallicum                             -1.739
## curated_taxonomyBifidobacterium dentium                               9.257
## curated_taxonomyBifidobacterium catenulatum subsp. kashiwanohense_A   6.741
## curated_taxonomyBifidobacterium catenulatum subsp. kashiwanohense     7.838
## curated_taxonomyBifidobacterium catenulatum subsp. catenulatum       -0.569
## curated_taxonomyBifidobacterium breve                                 7.934
## curated_taxonomyBifidobacterium bifidum                              -1.317
## curated_taxonomyBifidobacterium animalis subsp. lactis               -3.201
## curated_taxonomyBifidobacterium animalis subsp. animalis             -0.644
## curated_taxonomyBifidobacterium angulatum                             1.048
## curated_taxonomyBifidobacterium adolescentis                          3.381
##                                                                     Pr(>|z|)
## (Intercept)                                                          < 2e-16
## curated_taxonomyBifidobacterium tsurumiense                         4.18e-07
## curated_taxonomyBifidobacterium thermophilum                        0.127829
## curated_taxonomyBifidobacterium scardovii                           3.66e-11
## curated_taxonomyBifidobacterium ruminantium                         0.698424
## curated_taxonomyBifidobacterium pullorum                            0.106894
## curated_taxonomyBifidobacterium pseudocatenulatum                    < 2e-16
## curated_taxonomyBifidobacterium longum subsp. suis                  4.11e-11
## curated_taxonomyBifidobacterium longum subsp. longum                 < 2e-16
## curated_taxonomyBifidobacterium longum subsp. infantis               < 2e-16
## curated_taxonomyBifidobacterium hominis                             0.000614
## curated_taxonomyBifidobacterium globosum                            0.004630
## curated_taxonomyBifidobacterium gallicum                            0.082054
## curated_taxonomyBifidobacterium dentium                              < 2e-16
## curated_taxonomyBifidobacterium catenulatum subsp. kashiwanohense_A 1.57e-11
## curated_taxonomyBifidobacterium catenulatum subsp. kashiwanohense   4.59e-15
## curated_taxonomyBifidobacterium catenulatum subsp. catenulatum      0.569282
## curated_taxonomyBifidobacterium breve                               2.12e-15
## curated_taxonomyBifidobacterium bifidum                             0.187972
## curated_taxonomyBifidobacterium animalis subsp. lactis              0.001367
## curated_taxonomyBifidobacterium animalis subsp. animalis            0.519693
```

```
## curated_taxonomyBifidobacterium angulatum                               0.294551
## curated_taxonomyBifidobacterium adolescentis                            0.000722
##
## (Intercept)                                                             ***
## curated_taxonomyBifidobacterium tsurumiense                             ***
## curated_taxonomyBifidobacterium thermophilum
## curated_taxonomyBifidobacterium scardovii                               ***
## curated_taxonomyBifidobacterium ruminantium
## curated_taxonomyBifidobacterium pullorum
## curated_taxonomyBifidobacterium pseudocatenulatum                       ***
## curated_taxonomyBifidobacterium longum subsp. suis                      ***
## curated_taxonomyBifidobacterium longum subsp. longum                    ***
## curated_taxonomyBifidobacterium longum subsp. infantis                  ***
## curated_taxonomyBifidobacterium hominis                                 ***
## curated_taxonomyBifidobacterium globosum                                **
## curated_taxonomyBifidobacterium gallicum                                .
## curated_taxonomyBifidobacterium dentium                                 ***
## curated_taxonomyBifidobacterium catenulatum subsp. kashiwanohense_A ***
## curated_taxonomyBifidobacterium catenulatum subsp. kashiwanohense   ***
## curated_taxonomyBifidobacterium catenulatum subsp. catenulatum
## curated_taxonomyBifidobacterium breve                                   ***
## curated_taxonomyBifidobacterium bifidum
## curated_taxonomyBifidobacterium animalis subsp. lactis                  **
## curated_taxonomyBifidobacterium animalis subsp. animalis
## curated_taxonomyBifidobacterium angulatum
## curated_taxonomyBifidobacterium adolescentis                            ***
## ---
## Signif. codes:  0 '***' 0.001 '**' 0.01 '*' 0.05 '.' 0.1 ' ' 1
##
## (Dispersion parameter for poisson family taken to be 1)
##
##     Null deviance: 2660.61  on 3082  degrees of freedom
## Residual deviance:  633.75  on 3060  degrees of freedom
## AIC: 16369
##
## Number of Fisher Scoring iterations: 4
```

A Poisson regression model was fitted to evaluate the relationship between taxonomic groups and phenotypic richness. The overall model fit was assessed using a likelihood ratio test, comparing the full model to a null model containing only the intercept.

```r
# fit the null model (intercept-only model)
null_model <- glm(total_phenotypes ~ 1, data = phenotype_richness, family = poisson)
lr_test <- anova(null_model, glm_result, test = "Chisq")
print(lr_test)
```

```
## Analysis of Deviance Table
##
## Model 1: total_phenotypes ~ 1
## Model 2: total_phenotypes ~ curated_taxonomy
##   Resid. Df Resid. Dev Df Deviance  Pr(>Chi)
## 1      3082    2660.61
## 2      3060     633.75 22   2026.9 < 2.2e-16 ***
```

```
## ---
## Signif. codes:  0 '***' 0.001 '**' 0.01 '*' 0.05 '.' 0.1 ' ' 1
```

The likelihood ratio test comparing the full model to the null model indicated a significant improvement in model fit when including the taxonomic groups as predictors. The full model showed a significant reduction in residual deviance (from 2660.61 to 633.75) with a chi-squared value of 2026.9 and 22 degrees of freedom, resulting in a p-value less than 2.2e-16.

Run post-hoc tests and save the result as a table:

```
posthoc_glm <- emmeans(glm_result, ~ curated_taxonomy)
pairwise_comparisons <- as_tibble(contrast(posthoc_glm, method = "pairwise",
                                          adjust = "bonferroni")) %>%
  write_tsv("results/phenotypes/phenotypic_richness_stats.txt")
#gt(pairwise_comparisons)
```

#### 10.7 Conservation of predicted carbohydrate utilization pathways

The following heatmap shows the percent of genomes harboring a specific carbohydrate utilization pathway in each species/subspecies.

Hierarchical clustering options:

- **Distance metric**: Euclidean distance
- **Linkage method**: Ward's D2

```
# create a tibble where pathways are averaged at species level
bpm_3083_mean <- bpm_3083_carb_df %>%
  group_by(curated_taxonomy) %>%
  summarise_at(vars(Glc:`L-Xul`), list(mean))
# extract the binary matrix
bpm_3083_mean_mat <- as.matrix(bpm_3083_mean[, 2:69])
# add rownames to the matrix
rownames(bpm_3083_mean_mat) <- bpm_3083_mean$curated_taxonomy

# extract vectors containing data about glycan type and origin
glycan_type <- phenotype_metadata$type_group
glycan_origin <- phenotype_metadata$origin
# create two column annotations specifying glycan type and origin
ha_3083_mean <- HeatmapAnnotation(
  type = glycan_type,
  origin = glycan_origin,
  col = list(type = c("monosaccharides_and_derivatives" = "#E6E7E8",
                      "di_and_oligosaccharides" = "#BCBEC0",
                      "polysaccharides" = "#808285"),
             origin = c("universal" = "#ffd22d",
                        "host (animal)" = "#cbbedd",
                        "plant" = "#8dc63f",
                        "bacterial" = "#754c29",
                        "fungal" = "#008aa1")),
  show_annotation_name = FALSE,
  simple_anno_size = unit(3, "mm"))
```

```r
# add a coloring function
col_fun <- colorRamp2(c(0, 0.125, 0.25, 0.375, 0.5, 0.625, 0.75, 0.875, 1),
                      c("#ffffff", "#DEEBF7", "#C6DBEF", "#9ECAE1", "#6BAED6",
                        "#4292C6", "#2171B5", "#08519C", "#08306B"))

# specify the name of the output file
pdf("results/phenotypes/bpm_3083_collapsed_heatmap.pdf", width=15, height=8)
# plot the heatmap
ht_3083 <- ComplexHeatmap::Heatmap(bpm_3083_mean_mat,
        col = col_fun,
        bottom_annotation = ha_3083_mean,
        clustering_distance_rows = function(m) dist(m, method = "euclidean"),
        clustering_distance_columns = function(m) dist(m, method = "euclidean"),
        clustering_method_rows = "ward.D2",
        clustering_method_columns = "ward.D2",
        column_names_side = "bottom",
        rect_gp = gpar(col = "black", lwd = 0.1),
        row_names_gp = gpar(fontsize = 6),
        column_names_gp = gpar(fontsize = 6),
        column_names_rot = 75,
        width = unit(180, "mm"),
        height = unit(50, "mm"),
        heatmap_legend_param = list(
          col_fun = col_fun,
          at = c(0, 0.25, 0.5, 0.75, 1),
          title = "% of genomes with pathway",
          direction = "horizontal",
          title_position = "topcenter",
          border = "black",
          legend_width = unit(40, "mm"))
        )
draw(ht_3083)
# save the heatmap to a file
dev.off()
```

```
## pdf
##   2
```

```r
# draw the heatmap again to show it in the compiled markdown file
draw(ht_3083)
```

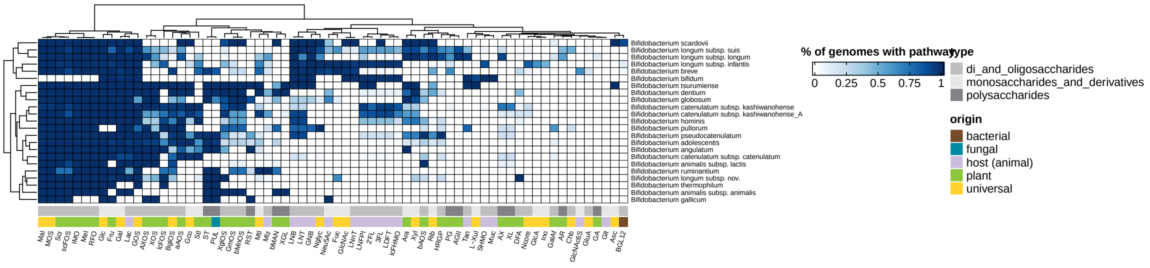

Number of predicted pathways with variability within species/subspecies.

```r
# count columns with only 0s and 1s
binary_columns <- sum(apply(bpm_3083_mean_mat, 2, function(col) all(col %in% c(0, 1))))

# count columns with intermediate values in the [0,1] range
range_columns <- sum(apply(bpm_3083_mean_mat, 2, function(col) any(col > 0 & col < 1)))

# print the counts
print(paste0("Number of predicted patwhays conserved within taxonomic groups: ", binary_columns))
```

```
## [1] "Number of predicted patwhays conserved within taxonomic groups: 2"
```

```r
print(paste0("Number of variable pathway: ", range_columns))
```

```
## [1] "Number of variable pathway: 66"
```

#### 10.8 Conservation of predicted B vitamin and amino acid biosynthesis pathways

The following heatmap demonstrates the percent of genomes harboring a specific pathway in each species/subspecies. These pathways include:

- Biosynthesis of B vitamins
- Biosynthesis of amino acids
- Urea utilization

Hierarchical clustering options:

- **Distance metric**: Euclidean distance
- **Linkage method**: Ward's D2

```
# create a tibble where pathways are averaged at species level
bpm_3083_mean2 <- bpm_3083_other_df %>%
  group_by(curated_taxonomy) %>%
  summarise_at(vars(B1:Urea_d), list(mean))
# extract the binary matrix
bpm_3083_mean2_mat <- as.matrix(bpm_3083_mean2[, 2:30])
# add rownames to the matrix
rownames(bpm_3083_mean2_mat) <- bpm_3083_mean2$curated_taxonomy

# extract vectors containing data about pathway type
pathway_type <- phenotype_metadata_other$type
# create a column annotations specifying pathway type
ha_3083_2_mean <- HeatmapAnnotation(
  type = pathway_type,
  col = list(type = c("vitamin" = "#8dc63f",
                      "amino_acid" = "#cbbedd",
                      "other" = "#ffd22d")),
  show_annotation_name = FALSE,
  simple_anno_size = unit(3, "mm"))
# add a coloring function
col_fun <- colorRamp2(c(0, 0.125, 0.25, 0.375, 0.5, 0.625, 0.75, 0.875, 1),
                      c("#ffffff", "#DEEBF7", "#C6DBEF", "#9ECAE1", "#6BAED6",
                        "#4292C6", "#2171B5", "#08519C", "#08306B"))

# specify the name of the output file
pdf("results/phenotypes/bpm_3083_collapsed_heatmap_other.pdf", width=15, height=8)
# plot the heatmap
ht_3083_other <- ComplexHeatmap::Heatmap(bpm_3083_mean2_mat,
        col = col_fun,
        bottom_annotation = ha_3083_2_mean,
        clustering_distance_rows = function(m) dist(m, method = "euclidean"),
        clustering_distance_columns = function(m) dist(m, method = "euclidean"),
        clustering_method_rows = "ward.D2",
        clustering_method_columns = "ward.D2",
        column_names_side = "bottom",
        rect_gp = gpar(col = "black", lwd = 0.1),
        row_names_gp = gpar(fontsize = 6),
        column_names_gp = gpar(fontsize = 6),
        column_names_rot = 60,
        width = unit(100, "mm"),
        height = unit(50, "mm"),
        heatmap_legend_param = list(
          col_fun = col_fun,
          at = c(0, 0.25, 0.5, 0.75, 1),
          title = "% of predicted producers",
          direction = "horizontal",
          title_position = "topcenter",
          border = "black",
          legend_width = unit(40, "mm"))
        )
```

```r
draw(ht_3083_other)
# save the heatmap to a file
dev.off()
```

```
## pdf
##   2
```

```r
# draw the heatmap again to show it in the compiled markdown file
draw(ht_3083_other)
```

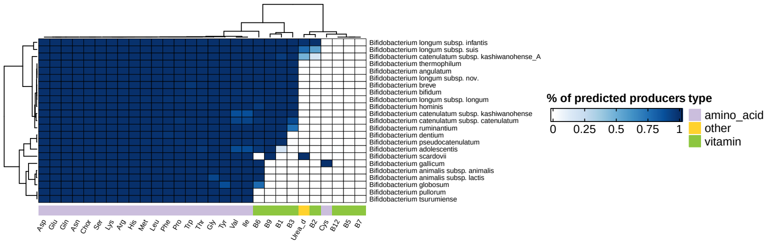

#### 10.9 Hierarchical clustering of the BPM (carbohydrate utilization) of 136 *Bifidobacterium catenulatum* and *Bifidobacterium hominis* genomes

The following heatmap shows the hierarchical clustering of BPM with the representation of carbohydrate utilization pathways predicted in 110 *Bifidobacterium catenulatum* and 26 *B. hominis* genomes.

Hierarchical clustering options:

- **Distance metric**: Hamming distance (equivalent to Manhattan distance for binary data)
- **Linkage method**: Average

```r
# filter rows based on taxonomy
bpm_cat_carb_df <- bpm_3083_carb_df %>%
  filter(curated_taxonomy %in% c("Bifidobacterium catenulatum subsp. catenulatum",
                                 "Bifidobacterium catenulatum subsp. kashiwanohense",
                                 "Bifidobacterium catenulatum subsp. kashiwanohense_A",
                                 "Bifidobacterium hominis")) %>%
  dplyr::select(genome_ID, curated_taxonomy, c(Glc:`L-Xul`))
```

```r
# extract the binary matrix
bpm_cat_mat <- as.matrix((bpm_cat_carb_df[, 3:70]))
# add rownames to the matrix
genome_id_cat <- bpm_cat_carb_df$genome_ID
rownames(bpm_cat_mat) <- genome_id_cat

# create a vector with taxonomy (group)
subsp_cat <- bpm_cat_carb_df$curated_taxonomy
# extract vectors containing data about glycan type and origin
phenotype_metadata <- phenotype_metadata %>%
  filter(!(phenotype %in% c("ManNAc", "GalNAc", "Man", "GalA")))
glycan_type <- phenotype_metadata$type_group
glycan_origin <- phenotype_metadata$origin
# create a coloring function
col_fun <- structure(c("white", "#08306b"), names = c("0", "1"))
# create a row annotation specifying taxonomy
ha_cat1 <- HeatmapAnnotation(
  which = c("row"),
  Taxonomy = subsp_cat,
  col = list(Taxonomy = c("Bifidobacterium catenulatum subsp. catenulatum" = "tomato2",
                          "Bifidobacterium catenulatum subsp. kashiwanohense" = "#ffa600",
                          "Bifidobacterium catenulatum subsp. kashiwanohense_A" = "#ffff7f",
                          "Bifidobacterium hominis" = "#a2d2de")),
  show_annotation_name = FALSE,
  simple_anno_size = unit(3, "mm"))
# create two column annotations specifying glycan type and origin
ha_cat2 <- HeatmapAnnotation(
  type = glycan_type,
  origin = glycan_origin,
  col = list(type = c("monosaccharides_and_derivatives" = "#E6E7E8",
                      "di_and_oligosaccharides" = "#BCBEC0",
                      "polysaccharides" = "#808285"),
             origin = c("universal" = "#ffd22d",
                        "host (animal)" = "#cbbedd",
                        "plant" = "#8dc63f",
                        "bacterial" = "#603913",
                        "fungal" = "#008aa1")),
  show_annotation_name = FALSE,
  simple_anno_size = unit(3, "mm"))

# plot the heatmap
pdf("results/phenotypes/Bcatenulatum_heatmap.pdf", width=18, height=18)
ht_bcatenulatum <- ComplexHeatmap::Heatmap(bpm_cat_mat,
                              name = "Predicted phenotype",
                              right_annotation = ha_cat1,
                              bottom_annotation = ha_cat2,
                              col = col_fun,
                              clustering_distance_rows = function(m)
                              dist(m, method = "binary"),
                              clustering_distance_columns = function(m)
                              dist(m, method = "binary"),
                              clustering_method_rows = "average",
                              clustering_method_columns = "average",
```

```r
                              rect_gp = gpar(col = "grey", lwd = 0.05),
                              show_row_names = TRUE,
                              row_names_gp = gpar(fontsize = 3),
                              column_names_gp = gpar(fontsize = 5),
                              column_names_rot = 60,
                              width = unit(200, "mm"),
                              height = unit(300, "mm"))
draw(ht_bcatenulatum)
# save the heatmap to a file
dev.off()
```

```
## pdf
##   2
```

```r
# draw the heatmap again to show it in the compiled markdown file
draw(ht_bcatenulatum)
```

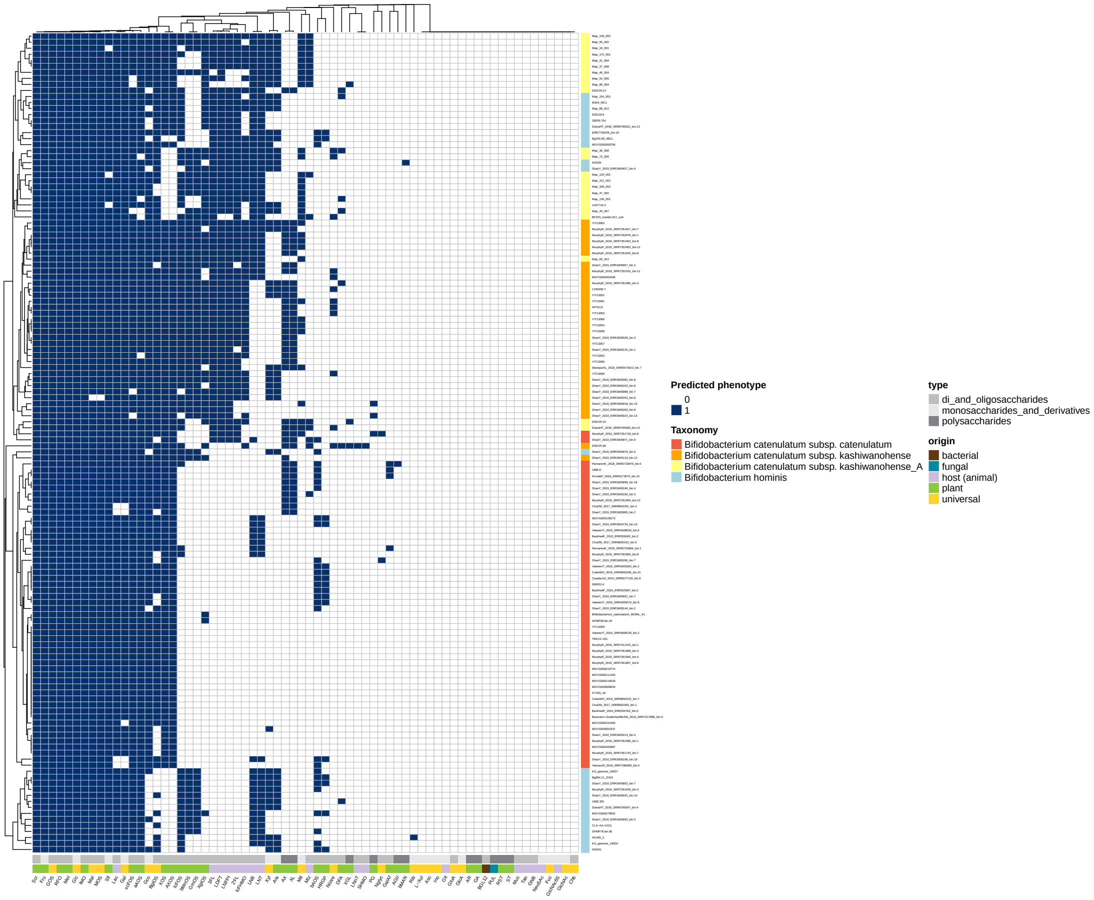

#### 10.10 Pathway enrichment analysis

Here we do a pathway enrichment analysis using Fisher's exact test. Pathways significantly enriched ($P_{adj}$ 0.01) in specified groups are shown.

```
# create a table for Fisher exact test
# extract data for 263 genomes
bpm_263_fish_df <- bpm_263_df %>%
  dplyr::select(genome_ID, curated_taxonomy, c(Glc:`L-Xul`), non_westernized, microbiota_age_group)
# extract data for 2820 genomes
bpm_2820_fish_df <- bpm_2820_df %>%
  dplyr::select(genome_ID, curated_taxonomy, c(Glc:`L-Xul`), non_westernized, microbiota_age_group)
# reorder columns in `bpm_263_fish_df` based on `bpm_2820_fish_df`
```

```
bpm_263_fish_df <- bpm_263_fish_df[, colnames(bpm_2820_fish_df)]
# merge the two tables
bpm_3083_fish_df <- merge(bpm_2820_fish_df, bpm_263_fish_df, all=TRUE)
```

###### 10.10.1 Westernized (age < 3) vs. non-Westernized (age < 3)

```
bpm_west_inf_df <- bpm_3083_fish_df %>%
  filter(microbiota_age_group == "child") %>%
  dplyr::select(genome_ID, c(Glc:`L-Xul`), non_westernized)
# create a contingency table
# i.e., calculate the number of occurrences of "yes" and "no" in the column `non_westernized`
# for each binary phenotype
contingency_table_west <- bpm_west_inf_df  %>%
  pivot_longer(cols = -c(genome_ID, non_westernized), names_to = "phenotype", values_to = "value") %>%
  group_by(phenotype, non_westernized, value) %>%
  summarise(count = n()) %>%
  pivot_wider(names_from = non_westernized, values_from = count) %>%
  replace(is.na(.), 0)
# keep only rows where neither all values are 0 nor all values are 1 within each group
contingency_table_west <- contingency_table_west %>%
  group_by(phenotype) %>%
  filter(!all(value == 0) & !all(value == 1))
# perform the Fisher's exact test, calculate odds ratios and adjusted p-values
result_west <- contingency_table_west %>%
  group_by(phenotype) %>%
  summarise(p_value = fisher.test(matrix(c(no, yes), nrow = 2))$p.value,
            odds_ratio = fisher.test(matrix(c(no, yes), nrow = 2))$estimate,
            conf_int_low = fisher.test(matrix(c(no, yes), nrow = 2))$conf.int[1],
            conf_int_high = fisher.test(matrix(c(no, yes), nrow = 2))$conf.int[2]) %>%
  mutate(p_adj = p.adjust(p_value, method = "fdr"))
# add information about phenotypes
results_west_ann <- left_join(result_west, phenotype_metadata, by = "phenotype")
# create a data frame with the log2-transformed odds ratios
odds_west_df <- results_west_ann %>%
  mutate(log2_odds_ratio = log2(odds_ratio)) %>%
  arrange(desc(log2_odds_ratio)) %>%
  dplyr::select(phenotype, origin, odds_ratio,
              conf_int_low, conf_int_high, log2_odds_ratio, p_value, p_adj)
# save the table
write_tsv(odds_west_df, "results/pathway_enrichment/tables/Westernized_age_less_3_vs_non_Westernized_ag

# plot odds ratios with 95% CI
ggplot(data = subset(odds_west_df, p_adj <= 0.01)) +
  aes(x = log2_odds_ratio, y = phenotype, fill = origin) +
  geom_errorbarh(aes(xmin = log2(conf_int_low),
                     xmax = log2(conf_int_high)),
                     height = 0.2,
                     position = position_dodge(width = 0.25),
                     linewidth = 1,
                     color = "grey50") +
  geom_point(position = position_dodge(width = 0.25) , shape = 21, size = 5) +
```

```r
  geom_vline(xintercept = 0, color = "black", linetype = "dashed") +
  scale_fill_manual(values = c("universal" = "#ffd22d",
                               "host (animal)" = "#cbbedd",
                               "plant" = "#8dc63f",
                               "fungal" = "#008aa1",
                               "bacterial" = "#603913")) +
  labs(x = "Log2(Odds_ratio)", y = "Pathway") +
  theme_bw() +
  theme(
    legend.title = element_blank(),
    plot.title = element_text(face = "bold"),
    axis.title = element_text(color = "black"),
    axis.text = element_text(color = "black")
  )
```

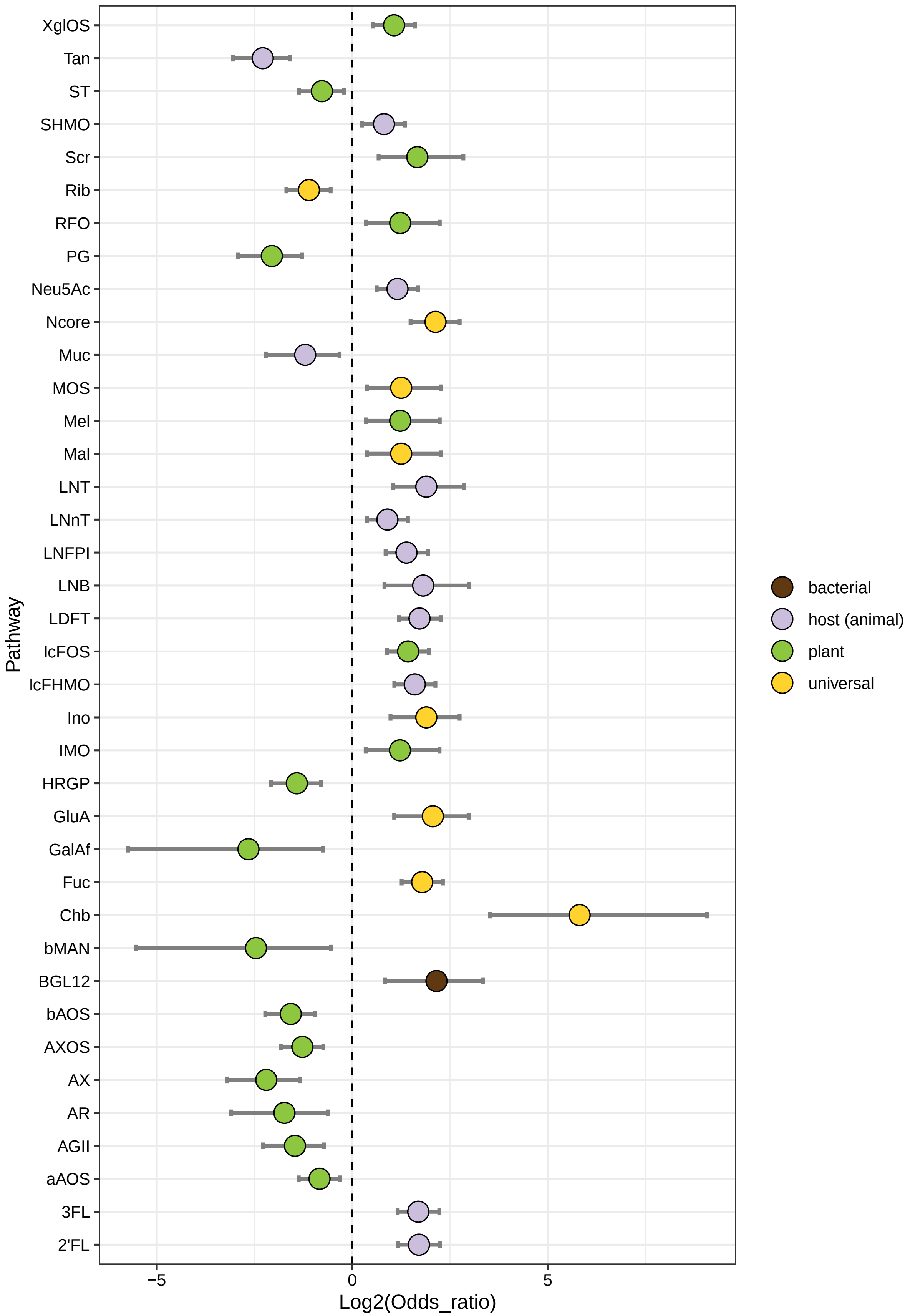

XglOS
Tan
ST
SHMO
Scr
Rib
RFO
PG
Neu5Ac
Ncore
Muc
MOS
Mel
Mal
LNT
LNnT
LNFPI
LNB
LDFT
lcFOS
lcFHMO
Ino
IMO
HRGP
GluA
GalAf
Fuc
Chb
bMAN
BGL12
bAOS
AXOS
AX
AR
AGII
aAOS
3FL
2'FL
Pathway
−5
0
5
Log2(Odds_ratio)
bacterial
host (animal)
plant
universal

```r
# save the figure to a file
ggsave("results/pathway_enrichment/child_westernized_vs_child_non_westernized.pdf",
       device = "pdf", width = 7, height = 10)
```

### 10.10.2 Westernized (age   3) vs. non-Westernized (age   3)

```r
bpm_west_ad_df <- bpm_3083_fish_df %>%
  filter(microbiota_age_group == "adult") %>%
  dplyr::select(genome_ID, c(Glc:`L-Xul`), non_westernized)
# create a contingency table
# i.e., calculate the number of occurrences of "yes" and "no" in the column `non_westernized`
# for each binary phenotype
contingency_table_west <- bpm_west_ad_df   %>%
  pivot_longer(cols = -c(genome_ID, non_westernized), names_to = "phenotype", values_to = "value") %>%
  group_by(phenotype, non_westernized, value) %>%
  summarise(count = n()) %>%
  pivot_wider(names_from = non_westernized, values_from = count) %>%
  replace(is.na(.), 0)
# keep only rows where neither all values are 0 nor all values are 1 within each group
contingency_table_west <- contingency_table_west %>%
  group_by(phenotype) %>%
  filter(!all(value == 0) & !all(value == 1))
# perform the Fisher's exact test, calculate odds ratios and adjusted p-values
result_west <- contingency_table_west %>%
  group_by(phenotype) %>%
  summarise(p_value = fisher.test(matrix(c(no, yes), nrow = 2))$p.value,
            odds_ratio = fisher.test(matrix(c(no, yes), nrow = 2))$estimate,
            conf_int_low = fisher.test(matrix(c(no, yes), nrow = 2))$conf.int[1],
            conf_int_high = fisher.test(matrix(c(no, yes), nrow = 2))$conf.int[2]) %>%
  mutate(p_adj = p.adjust(p_value, method = "fdr"))
# add information about phenotypes
results_west_ann <- left_join(result_west, phenotype_metadata, by = "phenotype")
# create a data frame with the log2-transformed odds ratios
odds_west_df <- results_west_ann %>%
  mutate(log2_odds_ratio = log2(odds_ratio)) %>%
  arrange(desc(log2_odds_ratio)) %>%
  dplyr::select(phenotype, origin, odds_ratio,
                conf_int_low, conf_int_high, log2_odds_ratio, p_value, p_adj)

# plot odds ratios with 95% CI
ggplot(data = subset(odds_west_df, p_adj <= 0.01)) +
  aes(x = log2_odds_ratio, y = phenotype, fill = origin) +
  geom_errorbarh(aes(xmin = log2(conf_int_low),
                     xmax = log2(conf_int_high)),
                     height = 0.2,
                     position = position_dodge(width = 0.25),
                     linewidth = 1,
                     color = "grey50") +
  geom_point(position = position_dodge(width = 0.25) , shape = 21, size = 5) +
  geom_vline(xintercept = 0, color = "black", linetype = "dashed") +
  scale_fill_manual(values = c("universal" = "#ffd22d",
                               "host (animal)" = "#cbbedd",
```

```
                                "plant" = "#8dc63f",
                                "fungal" = "#008aa1",
                                "bacterial" = "#603913")) +
  labs(x = "Log2(Odds_ratio)", y = "Pathway") +
  theme_bw() +
  theme(
    legend.title = element_blank(),
    plot.title = element_text(face = "bold"),
    axis.title = element_text(color = "black"),
    axis.text = element_text(color = "black")
  )
```

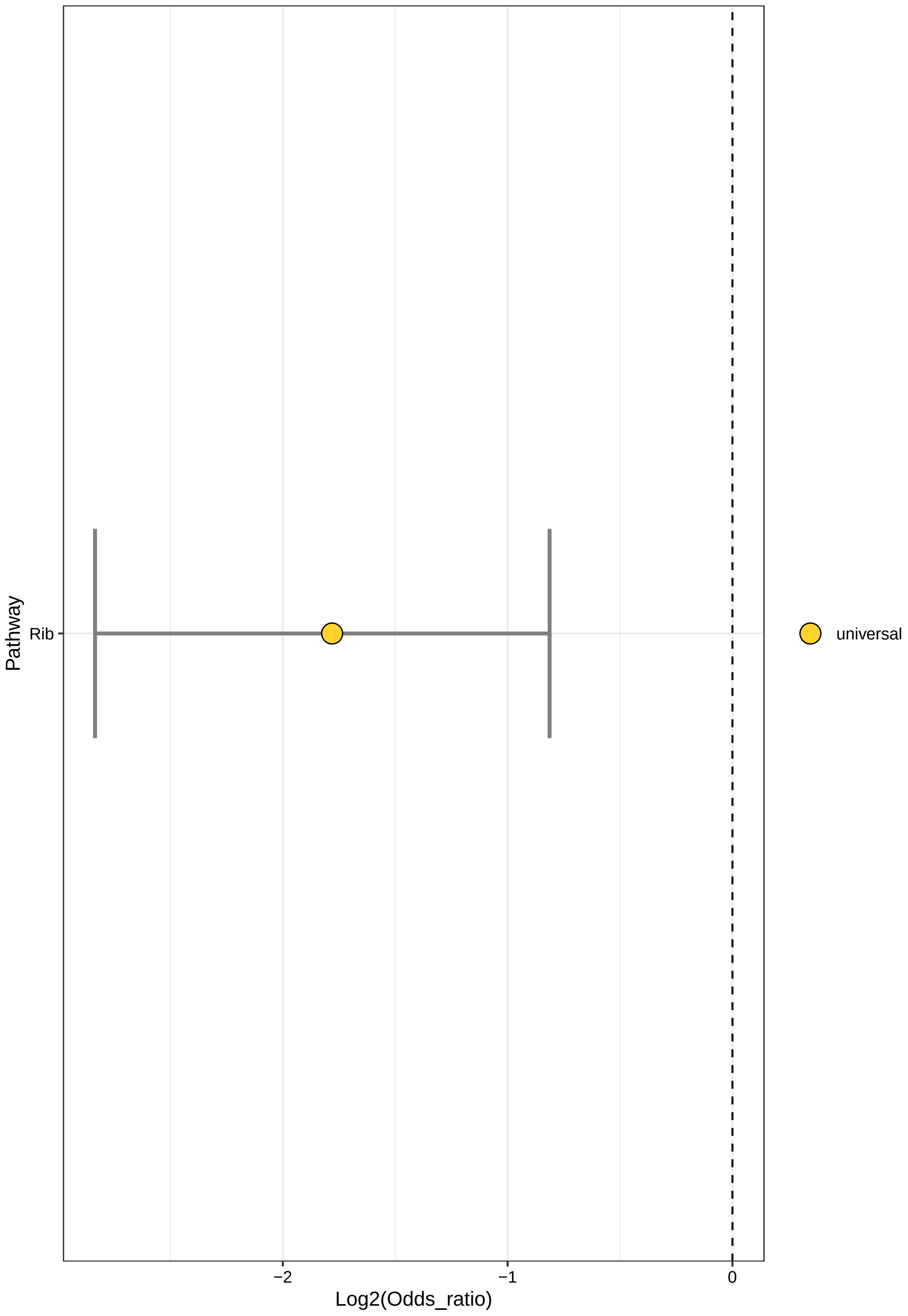
Pathway
Rib
−2
−1
0
Log2(Odds_ratio)
universal

```r
# save the figure to a file
ggsave("results/pathway_enrichment/adult_westernized_vs_adult_non_westernized.pdf",
       device = "pdf", width = 7, height = 10)
```

##### 10.10.3 Westernized (age < 3) vs. Westernized (age 3)

```r
bpm_age_west_df <- bpm_3083_fish_df %>%
  filter(non_westernized == "no") %>%
  dplyr::select(genome_ID, c(Glc:`L-Xul`), microbiota_age_group)
# create a contingency table
# i.e., calculate the number of occurrences of "yes" and "no" in the column
# `microbiota_age_group` for each binary phenotype
contingency_table_age <- bpm_age_west_df %>%
  pivot_longer(cols = -c(genome_ID, microbiota_age_group), names_to = "phenotype", values_to = "value")
  group_by(phenotype, microbiota_age_group, value) %>%
  summarise(count = n()) %>%
  pivot_wider(names_from = microbiota_age_group, values_from = count) %>%
  dplyr::select(phenotype, value, child, adult) %>%
  replace(is.na(.), 0)
# keep only rows where neither all values are 0 nor all values are 1 within each group
contingency_table_age <- contingency_table_age %>%
  group_by(phenotype) %>%
  filter(!all(value == 0) & !all(value == 1))
# perform the Fisher's exact test, calculate odds ratios and adjusted p-values
result_age <- contingency_table_age %>%
  group_by(phenotype) %>%
  summarise(p_value = fisher.test(matrix(c(child, adult), nrow = 2))$p.value,
            odds_ratio = fisher.test(matrix(c(child, adult), nrow = 2))$estimate,
            conf_int_low = fisher.test(matrix(c(child, adult), nrow = 2))$conf.int[1],
            conf_int_high = fisher.test(matrix(c(child, adult), nrow = 2))$conf.int[2]) %>%
  mutate(p_adj = p.adjust(p_value, method = "fdr"))
# add information about phenotypes
results_age_ann <- left_join(result_age, phenotype_metadata, by = "phenotype")
# create a data frame with the log2-transformed odds ratios
odds_age_df <- results_age_ann %>%
  mutate(log2_odds_ratio = log2(odds_ratio)) %>%
  arrange(desc(log2_odds_ratio)) %>%
  dplyr::select(phenotype, origin, odds_ratio,
                conf_int_low, conf_int_high, log2_odds_ratio, p_value, p_adj)
# save the table
write_tsv(odds_age_df, "results/pathway_enrichment/tables/Westernized_age_less_3_vs_Westernized_age_mor

# plot odds ratios with 95% CI
ggplot(data = subset(odds_age_df, p_adj <= 0.01)) +
  aes(x = log2_odds_ratio, y = phenotype, fill = origin) +
  geom_errorbarh(aes(xmin = log2(conf_int_low),
                     xmax = log2(conf_int_high)),
                     height = 0.2,
                     position = position_dodge(width = 0.25),
                     linewidth = 1,
                     color = "grey50") +
  geom_point(position = position_dodge(width = 0.25) , shape = 21, size = 5) +
```

```r
geom_vline(xintercept = 0, color = "black", linetype = "dashed") +
scale_fill_manual(values = c("universal" = "#ffd22d",
                             "host (animal)" = "#cbbedd",
                             "plant" = "#8dc63f",
                             "fungal" = "#008aa1",
                             "bacterial" = "#603913")) +
scale_x_continuous(breaks = seq(-12,4, 2)) +
labs(x = "Log2(Odds_ratio)", y = "Pathway") +
theme_bw() +
theme(
  legend.title = element_blank(),
  plot.title = element_text(face = "bold"),
  axis.title = element_text(color = "black"),
  axis.text = element_text(color = "black")
)
```

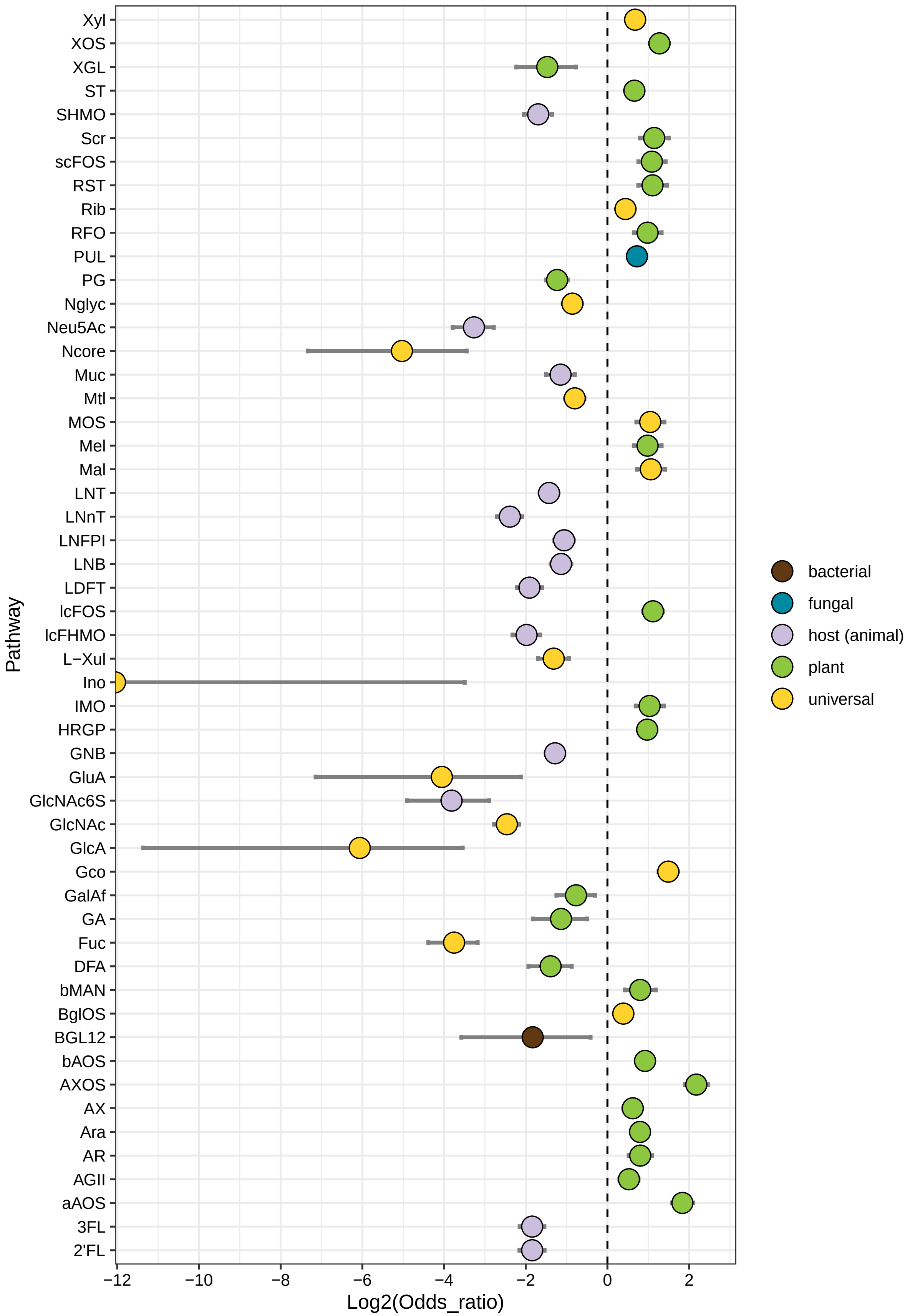
Xyl
XOS
XGL
ST
SHMO
Scr
scFOS
RST
Rib
RFO
PUL
PG
Nglyc
Neu5Ac
Ncore
Muc
Mtl
MOS
Mel
Mal
LNT
LNnT
LNFPI
LNB
LDFT
lcFOS
lcFHMO
L−Xul
Ino
IMO
HRGP
GNB
GluA
GlcNAc6S
GlcNAc
GlcA
Gco
GalAf
GA
Fuc
DFA
bMAN
BglOS
BGL12
bAOS
AXOS
AX
Ara
AR
AGII
aAOS
3FL
2'FL
Pathway
−12
−10
−8
−6
−4
−2
0
2
Log2(Odds_ratio)
bacterial
fungal
host (animal)
plant
universal

```r
# save the figure to a file
ggsave("results/pathway_enrichment/child_westernized_vs_adult_westernized.pdf",
       device = "pdf", width = 7, height = 10)
```

### 10.10.4 non-Westernized (age < 3) vs. non-Westernized (age   3)

```r
bpm_age_west_df <- bpm_3083_fish_df %>%
  filter(non_westerenized == "yes") %>%
  dplyr::select(genome_ID, c(Glc:`L-Xul`), microbiota_age_group)
# create a contingency table
# i.e., calculate the number of occurrences of "yes" and "no" in the column
# `microbiota_age_group` for each binary phenotype
contingency_table_age <- bpm_age_west_df %>%
  pivot_longer(cols = -c(genome_ID, microbiota_age_group), names_to = "phenotype", values_to = "value")
  group_by(phenotype, microbiota_age_group, value) %>%
  summarise(count = n()) %>%
  pivot_wider(names_from = microbiota_age_group, values_from = count) %>%
  dplyr::select(phenotype, value, child, adult) %>%
  replace(is.na(.), 0)
# keep only rows where neither all values are 0 nor all values are 1 within each group
contingency_table_age <- contingency_table_age %>%
  group_by(phenotype) %>%
  filter(!all(value == 0) & !all(value == 1))
# perform the Fisher's exact test, calculate odds ratios and adjusted p-values
result_age <- contingency_table_age %>%
  group_by(phenotype) %>%
  summarise(p_value = fisher.test(matrix(c(child, adult), nrow = 2))$p.value,
            odds_ratio = fisher.test(matrix(c(child, adult), nrow = 2))$estimate,
            conf_int_low = fisher.test(matrix(c(child, adult), nrow = 2))$conf.int[1],
            conf_int_high = fisher.test(matrix(c(child, adult), nrow = 2))$conf.int[2]) %>%
  mutate(p_adj = p.adjust(p_value, method = "fdr"))
# add information about phenotypes
results_age_ann <- left_join(result_age, phenotype_metadata, by = "phenotype")
# create a data frame with the log2-transformed odds ratios
odds_age_df <- results_age_ann %>%
  mutate(log2_odds_ratio = log2(odds_ratio)) %>%
  arrange(desc(log2_odds_ratio)) %>%
  dplyr::select(phenotype, origin, odds_ratio,
                conf_int_low, conf_int_high, log2_odds_ratio, p_value, p_adj) %>%
  filter((phenotype != "Lac"))
# save the table
write_tsv(odds_age_df, "results/pathway_enrichment/tables/non_Westernized_age_less_3_vs_non_Westernized

# plot odds ratios with 95% CI
ggplot(data = subset(odds_age_df, p_adj <= 0.01)) +
  aes(x = log2_odds_ratio, y = phenotype, fill = origin) +
  geom_errorbarh(aes(xmin = log2(conf_int_low),
                     xmax = log2(conf_int_high)),
                     height = 0.2,
                     position = position_dodge(width = 0.25),
                     linewidth = 1,
                     color = "grey50") +
```

```r
  geom_point(position = position_dodge(width = 0.25) , shape = 21, size = 5) +
  geom_vline(xintercept = 0, color = "black", linetype = "dashed") +
  scale_fill_manual(values = c("universal" = "#ffd22d",
                               "host (animal)" = "#cbbedd",
                               "plant" = "#8dc63f",
                               "fungal" = "#008aa1",
                               "bacterial" = "#603913")) +
  labs(x = "Log2(Odds_ratio)", y = "Pathway") +
  theme_bw() +
  theme(
    legend.title = element_blank(),
    plot.title = element_text(face = "bold"),
    axis.title = element_text(color = "black"),
    axis.text = element_text(color = "black")
  )
```

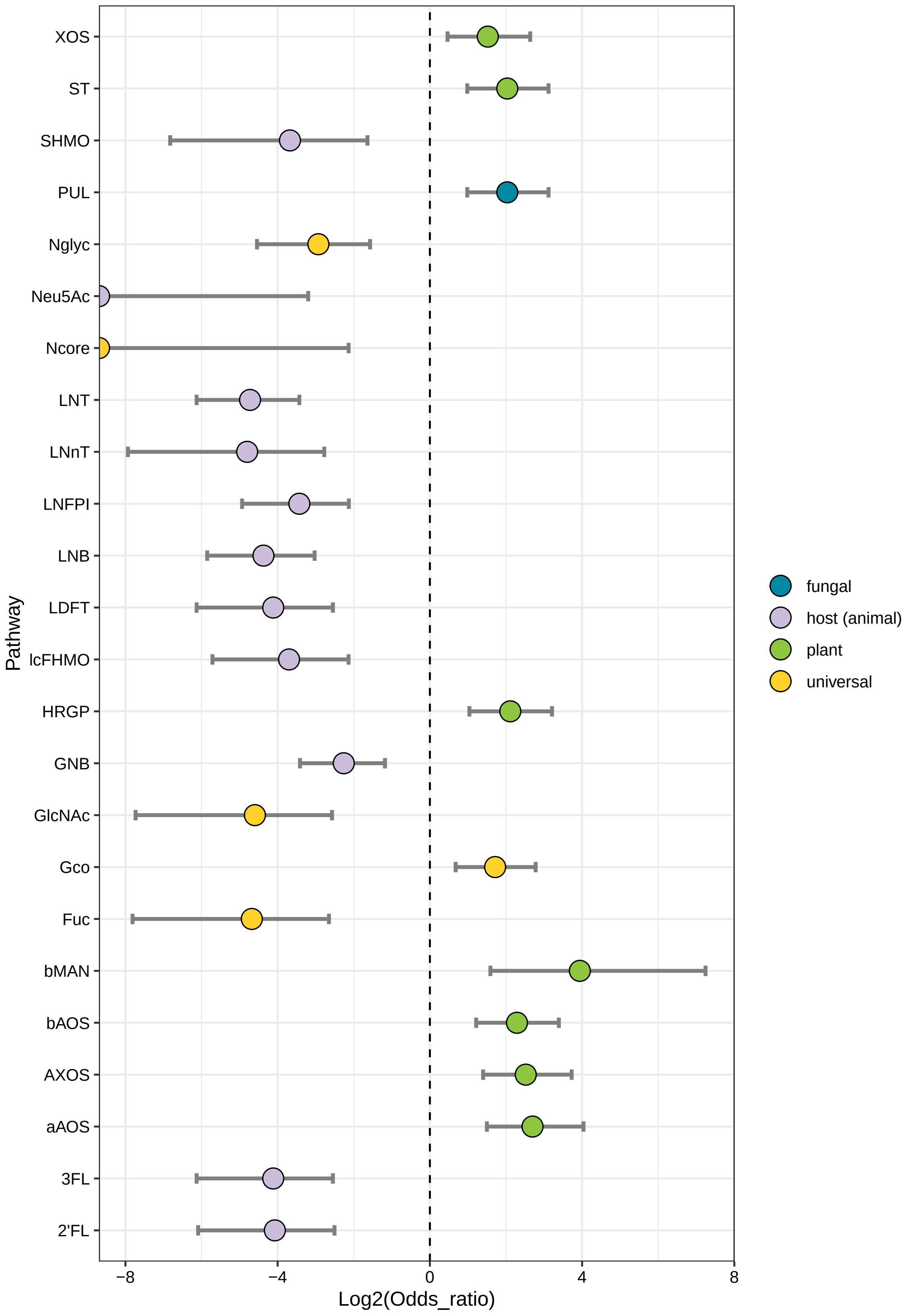
XOS
ST
SHMO
PUL
Nglyc
Neu5Ac
Ncore
LNT
LNnT
LNFPI
LNB
LDFT
lcFHMO
HRGP
GNB
GlcNAc
Gco
Fuc
bMAN
bAOS
AXOS
aAOS
3FL
2'FL
Pathway
−8
−4
0
4
8
Log2(Odds_ratio)
fungal
host (animal)
plant
universal

```
# save the figure to a file
ggsave("results/pathway_enrichment/child_non_westernized_vs_adult_non_westernized.pdf",
       device = "pdf", width = 7, height = 10)
```

###### 10.10.5 Westernized vs. non-Westernized within each taxon

```
# prepare data
bpm_3083_ado_df <- bpm_3083_fish_df %>%
  dplyr::select(curated_taxonomy, genome_ID, c(Glc:`L-Xul`), non_westernized)

# create a contingency table
contingency_table_west <- bpm_3083_ado_df %>%
  pivot_longer(cols = -c(genome_ID, curated_taxonomy, non_westernized),
               names_to = "phenotype", values_to = "value") %>%
  group_by(curated_taxonomy, phenotype, non_westernized, value) %>%
  summarise(count = n(), .groups = 'drop') %>%
  pivot_wider(names_from = non_westernized, values_from = count) %>%
  replace(is.na(.), 0)

# remove any taxon × pathway groups where all genomes either all encode a pathway
# or all do not encode a pathway
contingency_table_west <- contingency_table_west %>%
  group_by(curated_taxonomy, phenotype) %>%
  filter(n_distinct(value) == 2,      # must have both 0 and 1
         any(value == 0 & (`no` + `yes`) > 0),
         any(value == 1 & (`no` + `yes`) > 0)) %>%
  ungroup()

# perform the Fisher's exact test, calculate odds ratios and adjusted p-values
result_west <- contingency_table_west %>%
  group_by(curated_taxonomy, phenotype) %>%
  summarise(
    p_value = fisher.test(matrix(c(no, yes), nrow = 2))$p.value,
    odds_ratio = fisher.test(matrix(c(no, yes), nrow = 2))$estimate,
    conf_int_low = fisher.test(matrix(c(no, yes), nrow = 2))$conf.int[1],
    conf_int_high = fisher.test(matrix(c(no, yes), nrow = 2))$conf.int[2],
    .groups = 'drop'
  ) %>%
  group_by(curated_taxonomy) %>%
  mutate(p_adj = p.adjust(p_value, method = "fdr"))

# add information about phenotypes
results_west_ann <- left_join(result_west, phenotype_metadata, by = "phenotype")

# create a data frame with the log2-transformed odds ratios
odds_west_df <- results_west_ann %>%
  mutate(log2_odds_ratio = log2(odds_ratio)) %>%
  arrange(desc(log2_odds_ratio)) %>%
  dplyr::select(curated_taxonomy, phenotype, origin, odds_ratio,
                conf_int_low, conf_int_high, log2_odds_ratio, p_value, p_adj)

# save the table
```

```r
write_tsv(odds_west_df, "results/pathway_enrichment/tables/non_Westernized_vs_non_Westernized_taxon_spe

# create individual plots
plots <- odds_west_df %>%
  filter(p_adj <= 0.01) %>%
  split(.$curated_taxonomy) %>%
  purrr::map(~ ggplot(data = .x) +
          aes(x = log2_odds_ratio, y = phenotype, fill = origin) +
          geom_errorbarh(aes(xmin = log2(conf_int_low),
                             xmax = log2(conf_int_high)),
                             height = 0.2,
                             position = position_dodge(width = 0.25),
                             linewidth = 1,
                             color = "grey50") +
          geom_point(position = position_dodge(width = 0.25) , shape = 21, size = 5) +
          geom_vline(xintercept = 0, color = "black", linetype = "dashed") +
          scale_fill_manual(values = c("universal" = "#ffd22d",
                               "host (animal)" = "#cbbedd",
                               "plant" = "#8dc63f",
                               "fungal" = "#008aa1",
                               "bacterial" = "#603913")) +
          labs(x = "Log2(Odds_ratio)", y = "Phenotype", title = .x$curated_taxonomy[1]) +
          theme_bw() +
          theme(legend.title = element_blank(),
                plot.title = element_text(face = "bold"),
                axis.title = element_text(color = "black"),
                axis.text = element_text(color = "black")))

# combine plots using patchwork
combined_plot <- wrap_plots(plots, ncol = 2)
combined_plot
```

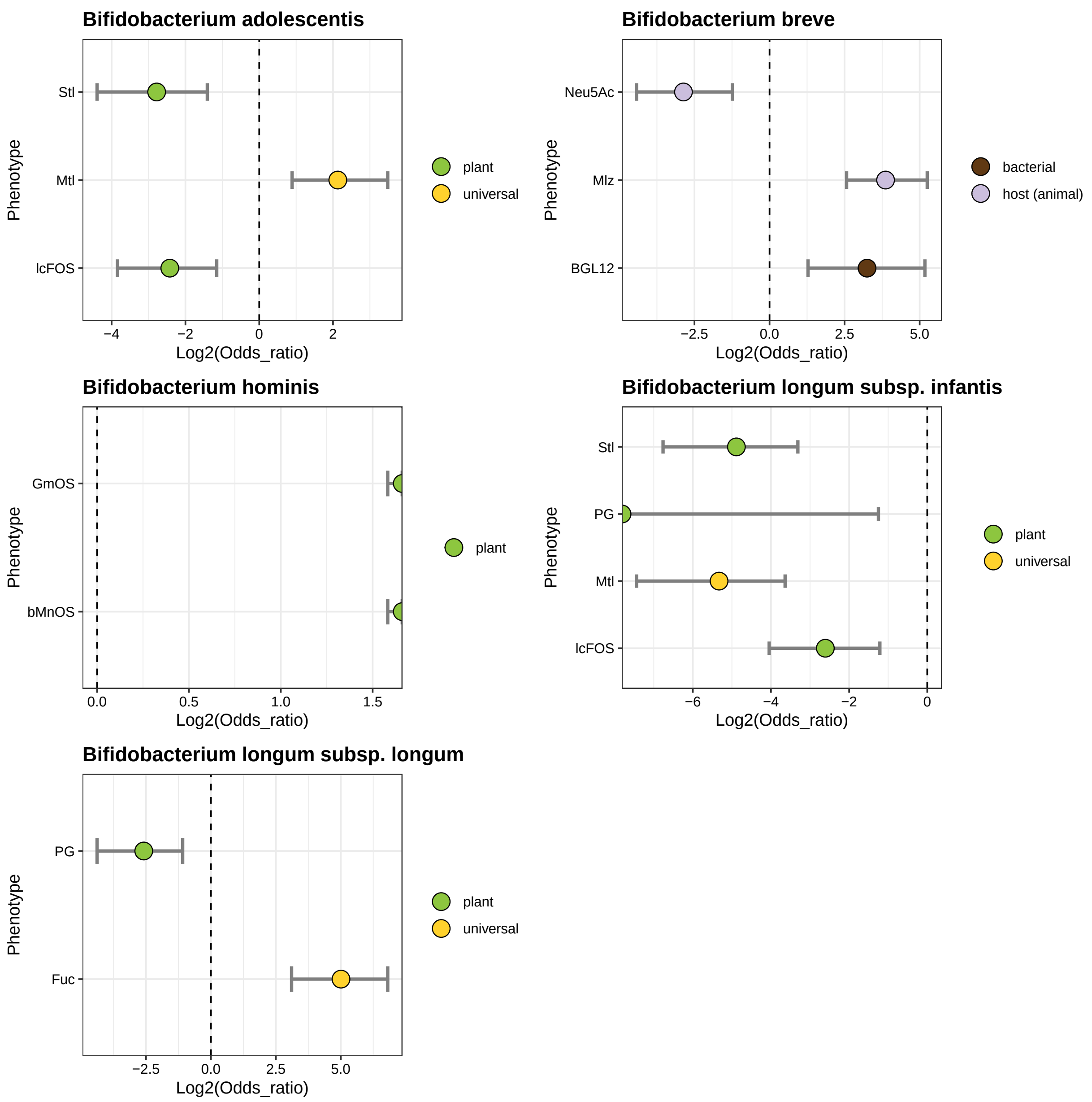

```
# save the combined plot
ggsave("results/pathway_enrichment/taxon_west_nonwest_combined.pdf", combined_plot,
       device = "pdf", width = 14, height = 10)
```

##### 10.10.6 Taxononomy differences: Westernized vs. non-Westernized

```
tax_3083_west_df <- bpm_3083_fish_df %>%
  select(genome_ID, curated_taxonomy, non_westernized)
# create a contingency table
west_tax <- tax_3083_west_df %>%
  pivot_longer(cols = -c(genome_ID, non_westernized), names_to = "group", values_to = "value") %>%
```

```r
  group_by(non_westernized, value) %>%
  summarise(count = n()) %>%
  pivot_wider(names_from = non_westernized, values_from = count) %>%
  replace(is.na(.), 0) %>%
  add_row(value = "total",
          `no` = sum(.[["no"]]),
          `yes` = sum(.[["yes"]])) %>%
  select(-"NA")

# calculate complement row for a given row
calculate_complement <- function(row, total_row) {
  new_row <- data.frame(
    value = paste0("NOT_", row$value),
    no = total_row$no - row$no,
    yes = total_row$yes - row$yes
  )
  return(new_row)
}
data <- west_tax
# filter out the "total" row
data_filtered <- data[data$value != "total", ]
# create a new data frame to store the results
new_data <- tibble()
# calculate complement rows
for (i in 1:nrow(data_filtered)) {
  original_row <- data_filtered[i, ]
  complement_row <- calculate_complement(original_row, data[data$value == "total", ])

  # append both original and complement rows to the new data frame
  new_data <- rbind(new_data, original_row, complement_row)
}
# add a column with taxonomy for grouping
west_tax_contingency <- new_data %>%
  mutate(curated_taxonomy = case_when(
    grepl("Bifidobacterium adolescentis", value) ~ "adolescentis",
    grepl("Bifidobacterium angulatum", value) ~ "angulatum",
    grepl("Bifidobacterium animalis subsp. animalis", value) ~ "animalis_animalis",
    grepl("Bifidobacterium animalis subsp. lactis", value) ~ "animalis_lactis",
    grepl("Bifidobacterium bifidum", value) ~ "bifidum",
    grepl("Bifidobacterium breve", value) ~ "breve",
    grepl("Bifidobacterium pseudocatenulatum", value) ~ "pseudocatenulatum",
    grepl("Bifidobacterium catenulatum subsp. catenulatum", value) ~ "catenulatum",
    grepl("Bifidobacterium catenulatum subsp. kashiwanohense_A", value) ~ "kashiwanohense_A",
    grepl("Bifidobacterium catenulatum subsp. kashiwanohense", value) ~ "kashiwanohense",
    grepl("Bifidobacterium dentium", value) ~ "dentium",
    grepl("Bifidobacterium gallicum", value) ~ "gallicum",
    grepl("Bifidobacterium globosum", value) ~ "globosum",
    grepl("Bifidobacterium hominis", value) ~ "hominis",
    grepl("Bifidobacterium longum subsp. infantis", value) ~ "longum_infantis",
    grepl("Bifidobacterium longum subsp. longum", value) ~ "longum_longum",
    grepl("Bifidobacterium longum subsp. nov.", value) ~ "longum_nov",
    grepl("Bifidobacterium longum subsp. suis", value) ~ "longum_suis",
    grepl("Bifidobacterium pullorum", value) ~ "pullorum",
```

| curated_taxonomy | odds_ratio | conf_int_low | conf_int_high | p_value | p_adj |
|---|---|---|---|---|---|
| adolescentis | 0.8324200 | 0.53501238 | 1.2554571 | 4.344427e-01 | 7.686294e-01 |
| angulatum | 0.0000000 | 0.00000000 | 15.2909643 | 1.000000e+00 | 1.000000e+00 |
| animalis_animalis | 0.0000000 | 0.00000000 | 539.9006224 | 1.000000e+00 | 1.000000e+00 |
| animalis_lactis | 1.2703990 | 0.02937557 | 8.8194537 | 5.646815e-01 | 8.886582e-01 |
| bifidum | 0.4890587 | 0.26561290 | 0.8382157 | 5.764201e-03 | 1.657208e-02 |
| breve | 1.5990267 | 1.02728344 | 2.4188134 | 2.939105e-02 | 7.511045e-02 |
| catenulatum | 0.5431719 | 0.06360423 | 2.0915075 | 5.795597e-01 | 8.886582e-01 |
| dentium | 0.2613798 | 0.03102419 | 0.9810688 | 4.519263e-02 | 1.039430e-01 |
| gallicum | 0.0000000 | 0.00000000 | 539.9006224 | 1.000000e+00 | 1.000000e+00 |
| globosum | 4.0153056 | 0.40448130 | 21.2674187 | 1.173490e-01 | 2.453661e-01 |
| hominis | 28.5465486 | 11.85582757 | 73.7609154 | 1.023854e-14 | 1.177433e-13 |
| kashiwanohense | 0.4203340 | 0.01028422 | 2.5380610 | 7.247770e-01 | 1.000000e+00 |
| kashiwanohense_A | Inf | 89.25587757 | Inf | 2.868127e-28 | 6.596692e-27 |
| longum_infantis | 4.9808076 | 3.24259630 | 7.5168964 | 2.379526e-12 | 1.824303e-11 |
| longum_longum | 0.2597595 | 0.15947490 | 0.4053167 | 4.970358e-12 | 2.857956e-11 |
| longum_nov | 0.0000000 | 0.00000000 | 1.5956268 | 1.665000e-01 | 3.191250e-01 |
| longum_suis | 14.5171809 | 5.04411930 | 41.8304220 | 6.431113e-07 | 2.958312e-06 |
| pseudocatenulatum | 0.4002985 | 0.20131097 | 0.7245575 | 1.118707e-03 | 3.679950e-03 |
| pullorum | 0.0000000 | 0.00000000 | 15.2909643 | 1.000000e+00 | 1.000000e+00 |
| ruminantium | 42.3237016 | 3.38219467 | 2196.8719880 | 1.119985e-03 | 3.679950e-03 |
| scardovii | 0.0000000 | 0.00000000 | 6.2545416 | 1.000000e+00 | 1.000000e+00 |
| thermophilum | 0.0000000 | 0.00000000 | 539.9006224 | 1.000000e+00 | 1.000000e+00 |
| tsurumiense | 0.0000000 | 0.00000000 | 74.3790348 | 1.000000e+00 | 1.000000e+00 |

```
    grepl("Bifidobacterium scardovii", value) ~ "scardovii",
    grepl("Bifidobacterium thermophilum", value) ~ "thermophilum",
    grepl("Bifidobacterium tsurumiense", value) ~ "tsurumiense",
    grepl("Bifidobacterium ruminantium", value) ~ "ruminantium",
    TRUE ~ NA_character_
  ))
# perform the Fisher's exact test, calculate odds ratios and adjusted p-values
result_west_tax <-  west_tax_contingency %>%
  select(curated_taxonomy , value, everything()) %>%
  group_by(curated_taxonomy ) %>%
  summarise(p_value = fisher.test(matrix(c(yes, no), nrow = 2))$p.value,
            odds_ratio = fisher.test(matrix(c(yes, no), nrow = 2))$estimate,
            conf_int_low = fisher.test(matrix(c(yes, no), nrow = 2))$conf.int[1],
            conf_int_high = fisher.test(matrix(c(yes, no), nrow = 2))$conf.int[2]) %>%
  mutate(p_adj = p.adjust(p_value, method = "fdr")) %>%
  select(curated_taxonomy, odds_ratio, conf_int_low, conf_int_high, p_value, p_adj)
# save the table
write_tsv(result_west_tax, "results/pathway_enrichment/tables/taxonomy_West_non_West.txt")
gt(result_west_tax)
```

### 11 Analysis of *in vitro* growth data

#### 11.1 *B. adolescentis* LFYP80

Growth curves of *B. adolescentis* LFYP80 in MRS-AC supplemented with various (n = 47) carbon sources.

```
growth_curves("Bifidobacterium adolescentis LFYP80",
              "data/growth/formatted/Badolescentis_LFYP80.txt",
              "results/growth/growth_curves/Badolescentis_LFYP80.pdf",
              40)
```

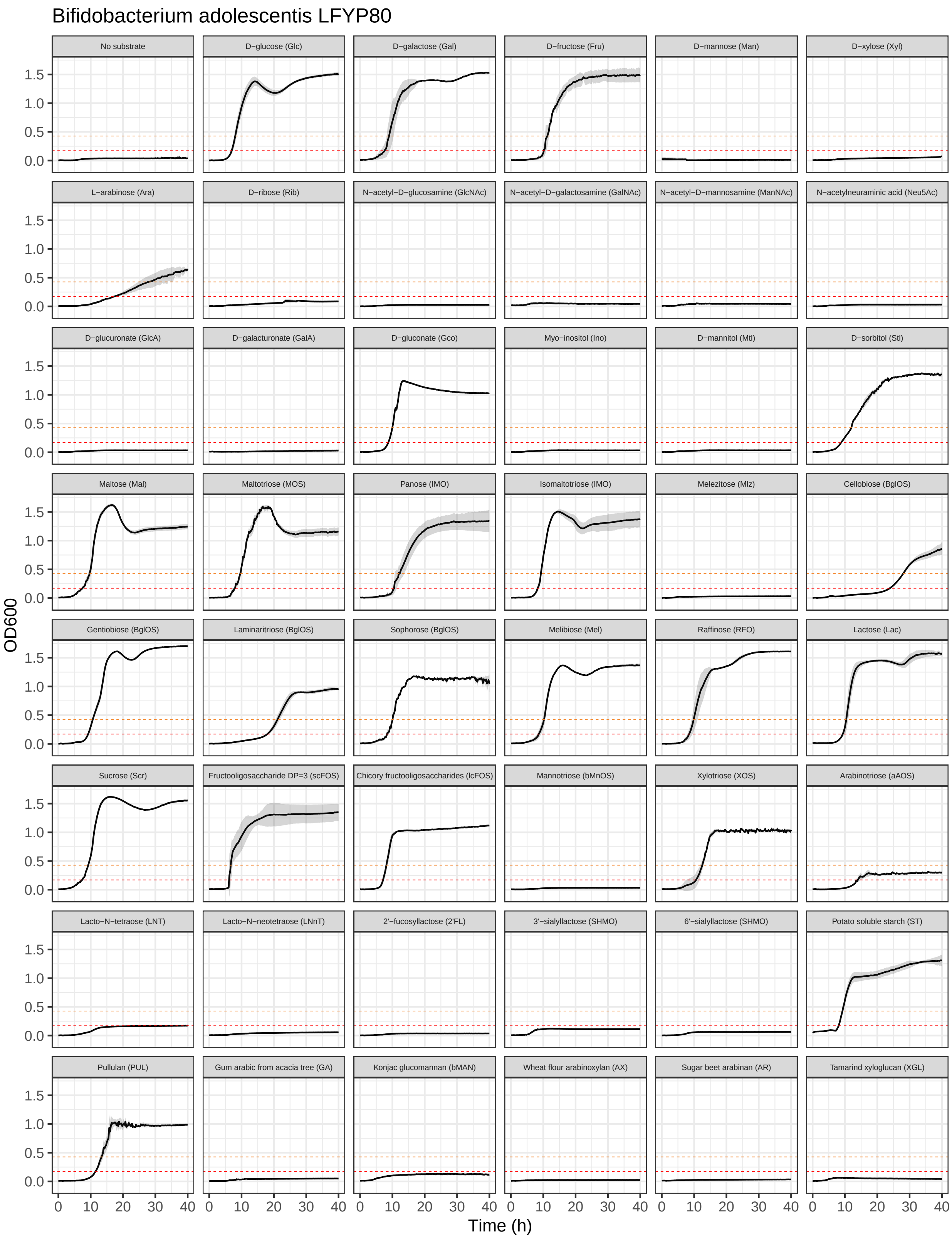

Bifidobacterium adolescentis LFYP80
No substrate
D−glucose (Glc)
D−galactose (Gal)
D−fructose (Fru)
D−mannose (Man)
D−xylose (Xyl)
L−arabinose (Ara)
D−ribose (Rib)
N−acetyl−D−glucosamine (GlcNAc)
N−acetyl−D−galactosamine (GalNAc)
N−acetyl−D−mannosamine (ManNAc)
N−acetylneuraminic acid (Neu5Ac)
D−glucuronate (GlcA)
D−galacturonate (GalA)
D−gluconate (Gco)
Myo−inositol (Ino)
D−mannitol (Mtl)
D−sorbitol (Stl)
Maltose (Mal)
Maltotriose (MOS)
Panose (IMO)
Isomaltotriose (IMO)
Melezitose (Mlz)
Cellobiose (BglOS)
Gentiobiose (BglOS)
Laminaritriose (BglOS)
Sophorose (BglOS)
Melibiose (Mel)
Raffinose (RFO)
Lactose (Lac)
Sucrose (Scr)
Fructooligosaccharide DP=3 (scFOS)
Chicory fructooligosaccharides (lcFOS)
Mannotriose (bMnOS)
Xylotriose (XOS)
Arabinotriose (aAOS)
Lacto−N−tetraose (LNT)
Lacto−N−neotetraose (LNnT)
2'−fucosyllactose (2'FL)
3'−sialyllactose (SHMO)
6'−sialyllactose (SHMO)
Potato soluble starch (ST)
Pullulan (PUL)
Gum arabic from acacia tree (GA)
Konjac glucomannan (bMAN)
Wheat flour arabinoxylan (AX)
Sugar beet arabinan (AR)
Tamarind xyloglucan (XGL)
OD600
0.0
0.5
1.0
1.5
Time (h)
0
10
20
30
40

#### 11.2 *B. adolescentis* M56B_1C3

Growth curves of *B. adolescentis* M56B_1C3 in MRS-AC supplemented with various (n = 45) carbon sources.

```
growth_curves("Bifidobacterium adolescentis M56B_1C3",
              "data/growth/formatted/Badolescentis_M56B_1C3.txt",
              "results/growth/growth_curves/Badolescentis_M56B_1C3.pdf",
              40)
```

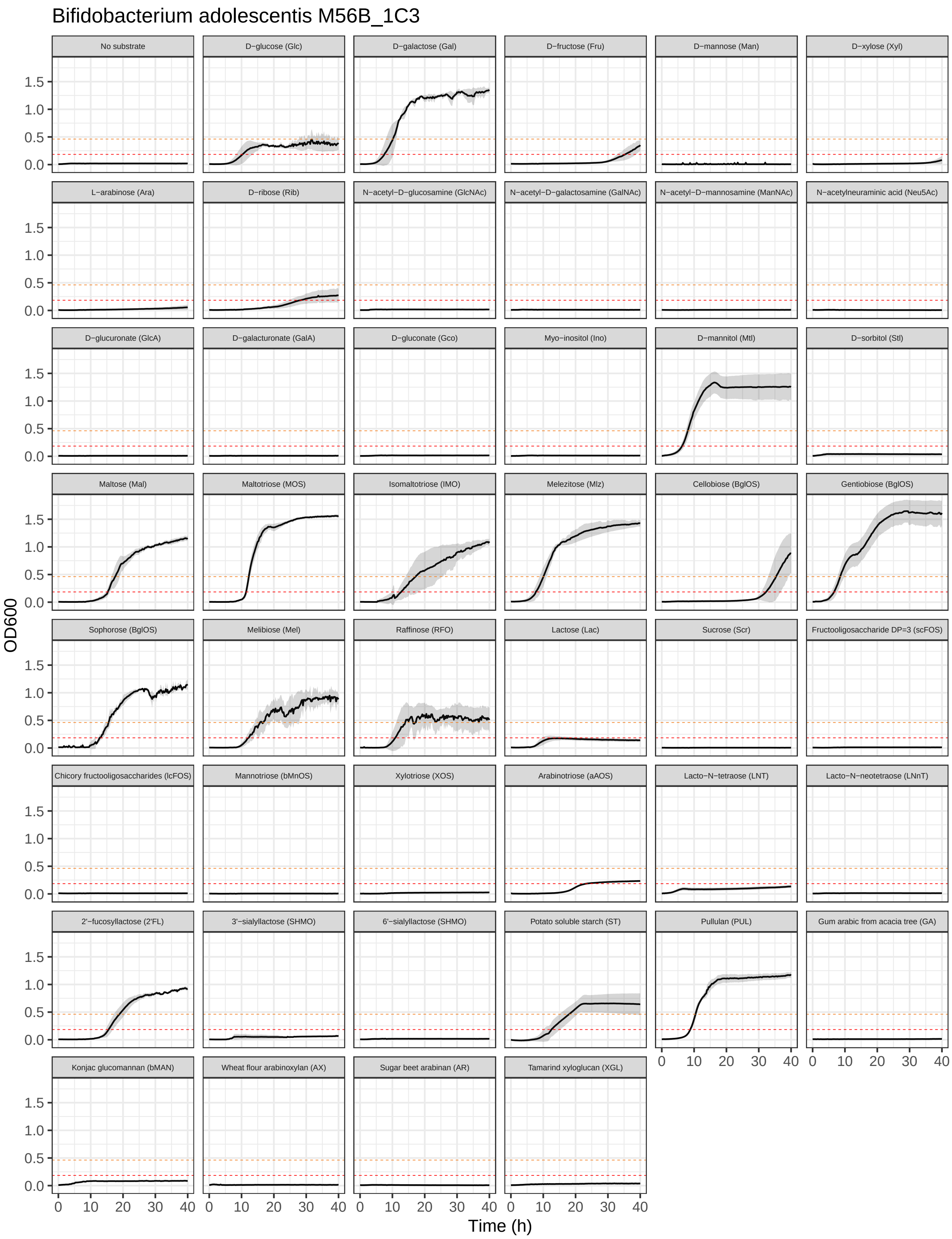

Bifidobacterium adolescentis M56B_1C3
No substrate
D−glucose (Glc)
D−galactose (Gal)
D−fructose (Fru)
D−mannose (Man)
D−xylose (Xyl)
L−arabinose (Ara)
D−ribose (Rib)
N−acetyl−D−glucosamine (GlcNAc)
N−acetyl−D−galactosamine (GalNAc)
N−acetyl−D−mannosamine (ManNAc)
N−acetylneuraminic acid (Neu5Ac)
D−glucuronate (GlcA)
D−galacturonate (GalA)
D−gluconate (Gco)
Myo−inositol (Ino)
D−mannitol (Mtl)
D−sorbitol (Stl)
Maltose (Mal)
Maltotriose (MOS)
Isomaltotriose (IMO)
Melezitose (Mlz)
Cellobiose (BglOS)
Gentiobiose (BglOS)
Sophorose (BglOS)
Melibiose (Mel)
Raffinose (RFO)
Lactose (Lac)
Sucrose (Scr)
Fructooligosaccharide DP=3 (scFOS)
Chicory fructooligosaccharides (lcFOS)
Mannotriose (bMnOS)
Xylotriose (XOS)
Arabinotriose (aAOS)
Lacto−N−tetraose (LNT)
Lacto−N−neotetraose (LNnT)
2'−fucosyllactose (2'FL)
3'−sialyllactose (SHMO)
6'−sialyllactose (SHMO)
Potato soluble starch (ST)
Pullulan (PUL)
Gum arabic from acacia tree (GA)
Konjac glucomannan (bMAN)
Wheat flour arabinoxylan (AX)
Sugar beet arabinan (AR)
Tamarind xyloglucan (XGL)
OD600
1.5
1.0
0.5
0.0
0
10
20
30
40
Time (h)

#### 11.3 *B. bifidum* Bg41221_3D10

Growth curves of *B. bifidum* Bg41221_3D10 in MRS-AC supplemented with various (n = 47) carbon sources.

```
growth_curves("Bifidobacterium bifidum Bg41221_3D10",
              "data/growth/formatted/Bbifidum_Bg41221_3D10.txt",
              "results/growth/growth_curves/Bbifidum_Bg41221_3D10.pdf",
              40)
```

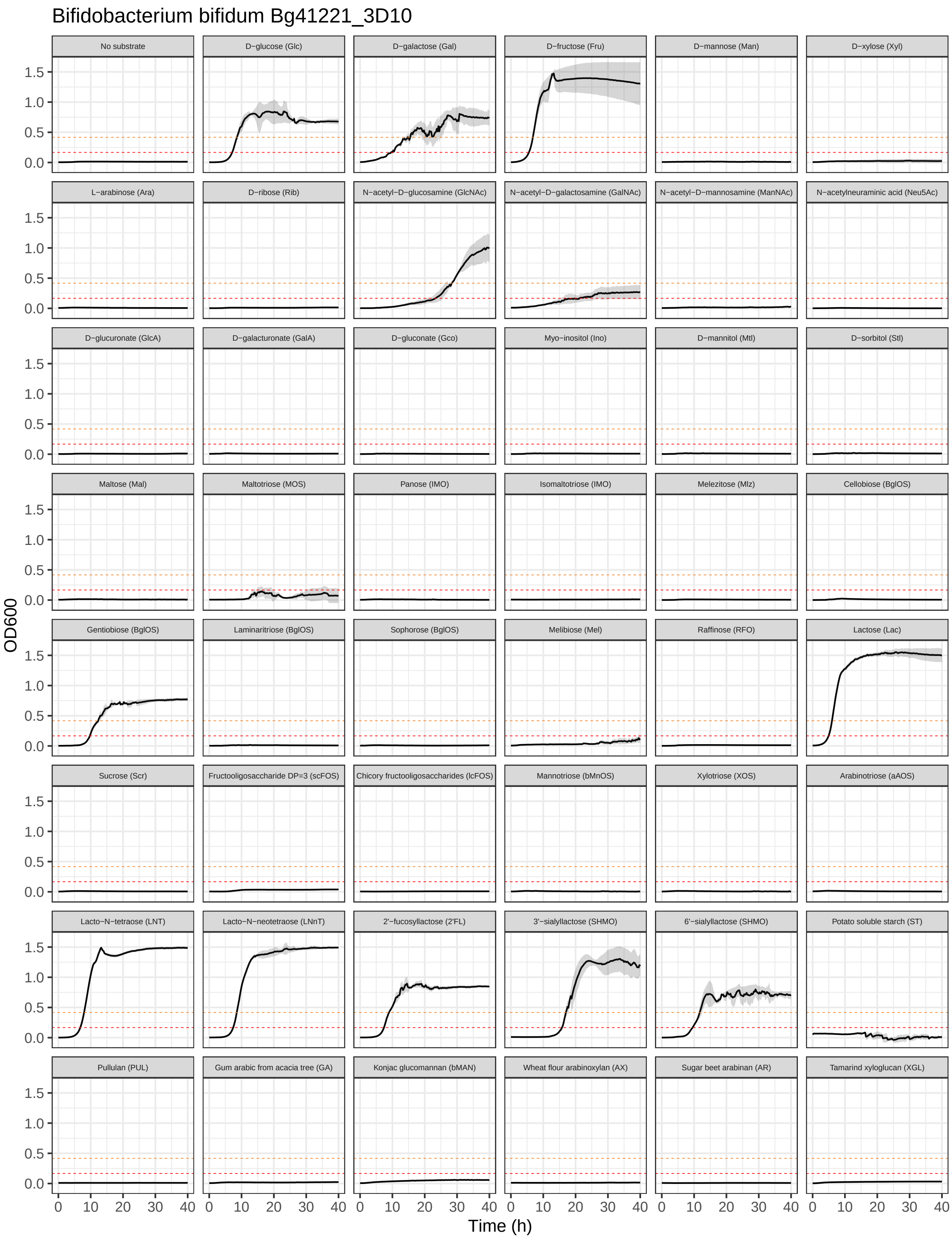
Bifidobacterium bifidum Bg41221_3D10
No substrate
D−glucose (Glc)
D−galactose (Gal)
D−fructose (Fru)
D−mannose (Man)
D−xylose (Xyl)
L−arabinose (Ara)
D−ribose (Rib)
N−acetyl−D−glucosamine (GlcNAc)
N−acetyl−D−galactosamine (GalNAc)
N−acetyl−D−mannosamine (ManNAc)
N−acetylneuraminic acid (Neu5Ac)
D−glucuronate (GlcA)
D−galacturonate (GalA)
D−gluconate (Gco)
Myo−inositol (Ino)
D−mannitol (Mtl)
D−sorbitol (Stl)
Maltose (Mal)
Maltotriose (MOS)
Panose (IMO)
Isomaltotriose (IMO)
Melezitose (Mlz)
Cellobiose (BglOS)
Gentiobiose (BglOS)
Laminaritriose (BglOS)
Sophorose (BglOS)
Melibiose (Mel)
Raffinose (RFO)
Lactose (Lac)
Sucrose (Scr)
Fructooligosaccharide DP=3 (scFOS)
Chicory fructooligosaccharides (lcFOS)
Mannotriose (bMnOS)
Xylotriose (XOS)
Arabinotriose (aAOS)
Lacto−N−tetraose (LNT)
Lacto−N−neotetraose (LNnT)
2'−fucosyllactose (2'FL)
3'−sialyllactose (SHMO)
6'−sialyllactose (SHMO)
Potato soluble starch (ST)
Pullulan (PUL)
Gum arabic from acacia tree (GA)
Konjac glucomannan (bMAN)
Wheat flour arabinoxylan (AX)
Sugar beet arabinan (AR)
Tamarind xyloglucan (XGL)
OD600
0.0
0.5
1.0
1.5
0
10
20
30
40
Time (h)

#### 11.4 *B. bifidum* M138B_2A5

Growth curves of *B. bifidum* M138B_2A5 in MRS-AC supplemented with various (n = 43) carbon sources.

```
growth_curves("Bifidobacterium bifidum M138B_2A5",
              "data/growth/formatted/Bbifidum_M138B_2A5.txt",
              "results/growth/growth_curves/Bbifidum_M138B_2A5.pdf",
              40)
```

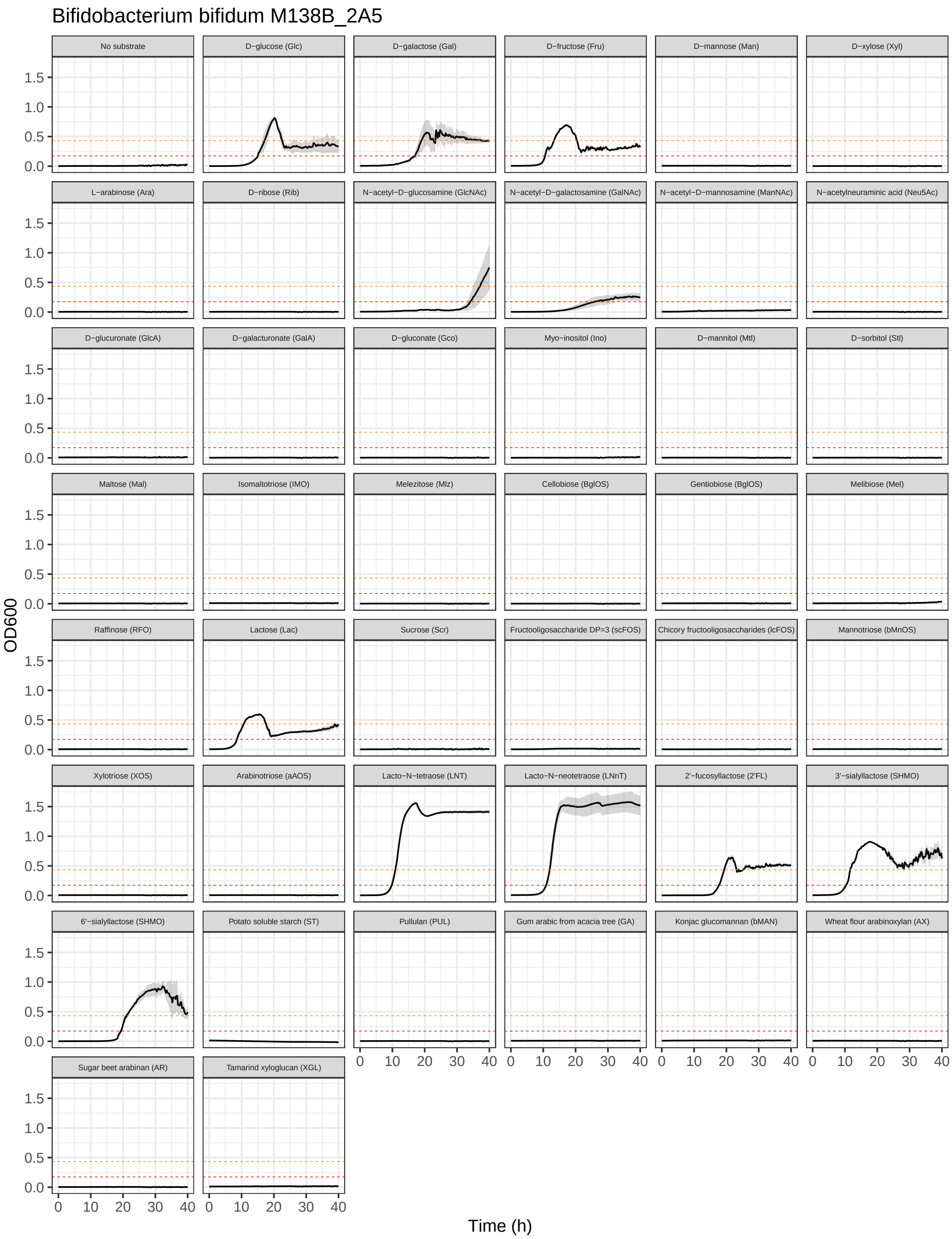
Bifidobacterium bifidum M138B_2A5
No substrate
D−glucose (Glc)
D−galactose (Gal)
D−fructose (Fru)
D−mannose (Man)
D−xylose (Xyl)
L−arabinose (Ara)
D−ribose (Rib)
N−acetyl−D−glucosamine (GlcNAc)
N−acetyl−D−galactosamine (GalNAc)
N−acetyl−D−mannosamine (ManNAc)
N−acetylneuraminic acid (Neu5Ac)
D−glucuronate (GlcA)
D−galacturonate (GalA)
D−gluconate (Gco)
Myo−inositol (Ino)
D−mannitol (Mtl)
D−sorbitol (Stl)
Maltose (Mal)
Isomaltotriose (IMO)
Melezitose (Mlz)
Cellobiose (BglOS)
Gentiobiose (BglOS)
Melibiose (Mel)
Raffinose (RFO)
Lactose (Lac)
Sucrose (Scr)
Fructooligosaccharide DP=3 (scFOS)
Chicory fructooligosaccharides (lcFOS)
Mannotriose (bMnOS)
Xylotriose (XOS)
Arabinotriose (aAOS)
Lacto−N−tetraose (LNT)
Lacto−N−neotetraose (LNnT)
2'−fucosyllactose (2'FL)
3'−sialyllactose (SHMO)
6'−sialyllactose (SHMO)
Potato soluble starch (ST)
Pullulan (PUL)
Gum arabic from acacia tree (GA)
Konjac glucomannan (bMAN)
Wheat flour arabinoxylan (AX)
Sugar beet arabinan (AR)
Tamarind xyloglucan (XGL)
OD600
0.0
0.5
1.0
1.5
0
10
20
30
40
Time (h)

#### 11.5 *B. bifidum* M257B_A12

Growth curves of *B. bifidum* M257B_A12 in MRS-AC supplemented with various (n = 42) carbon sources.

```
growth_curves("Bifidobacterium bifidum M257B_A12",
              "data/growth/formatted/Bbifidum_M257B_A12.txt",
              "results/growth/growth_curves/Bbifidum_M257B_A12.pdf",
              40)
```

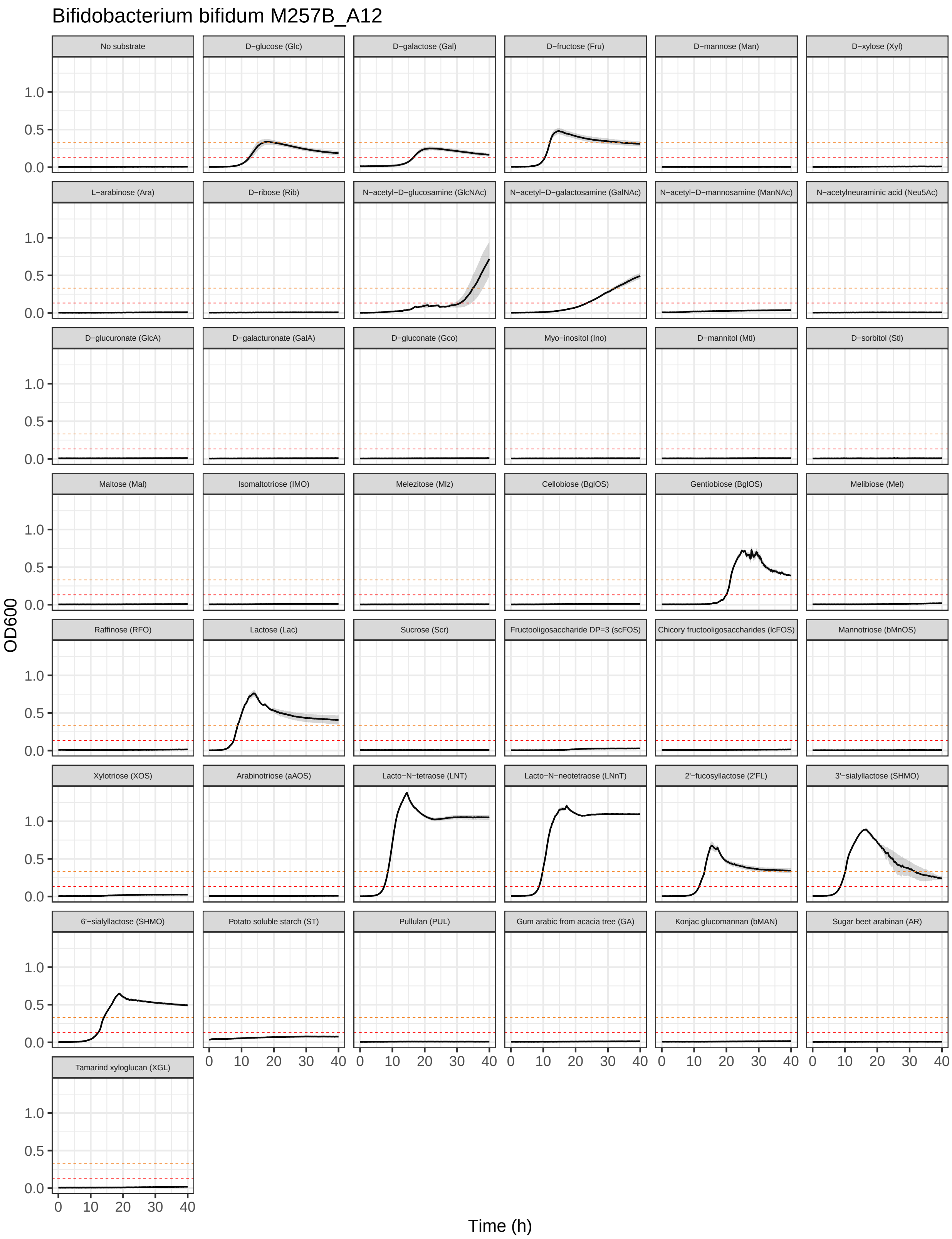
Bifidobacterium bifidum M257B_A12
No substrate
D−glucose (Glc)
D−galactose (Gal)
D−fructose (Fru)
D−mannose (Man)
D−xylose (Xyl)
L−arabinose (Ara)
D−ribose (Rib)
N−acetyl−D−glucosamine (GlcNAc)
N−acetyl−D−galactosamine (GalNAc)
N−acetyl−D−mannosamine (ManNAc)
N−acetylneuraminic acid (Neu5Ac)
D−glucuronate (GlcA)
D−galacturonate (GalA)
D−gluconate (Gco)
Myo−inositol (Ino)
D−mannitol (Mtl)
D−sorbitol (Stl)
Maltose (Mal)
Isomaltotriose (IMO)
Melezitose (Mlz)
Cellobiose (BglOS)
Gentiobiose (BglOS)
Melibiose (Mel)
Raffinose (RFO)
Lactose (Lac)
Sucrose (Scr)
Fructooligosaccharide DP=3 (scFOS)
Chicory fructooligosaccharides (lcFOS)
Mannotriose (bMnOS)
Xylotriose (XOS)
Arabinotriose (aAOS)
Lacto−N−tetraose (LNT)
Lacto−N−neotetraose (LNnT)
2'−fucosyllactose (2'FL)
3'−sialyllactose (SHMO)
6'−sialyllactose (SHMO)
Potato soluble starch (ST)
Pullulan (PUL)
Gum arabic from acacia tree (GA)
Konjac glucomannan (bMAN)
Sugar beet arabinan (AR)
Tamarind xyloglucan (XGL)
OD600
0.0
0.5
1.0
0 10 20 30 40
Time (h)

#### 11.6 *B. breve* Bg131.S11_D6

Growth curves of *B. breve* Bg131.S11_D6 in MRS-AC supplemented with various (n = 44) carbon sources.

```
growth_curves("Bifidobacterium breve Bg131.S11_D6",
              "data/growth/formatted/Bbreve_Bg131.S11_D6.txt",
              "results/growth/growth_curves/Bbreve_Bg131.S11_D6.pdf",
              40)
```

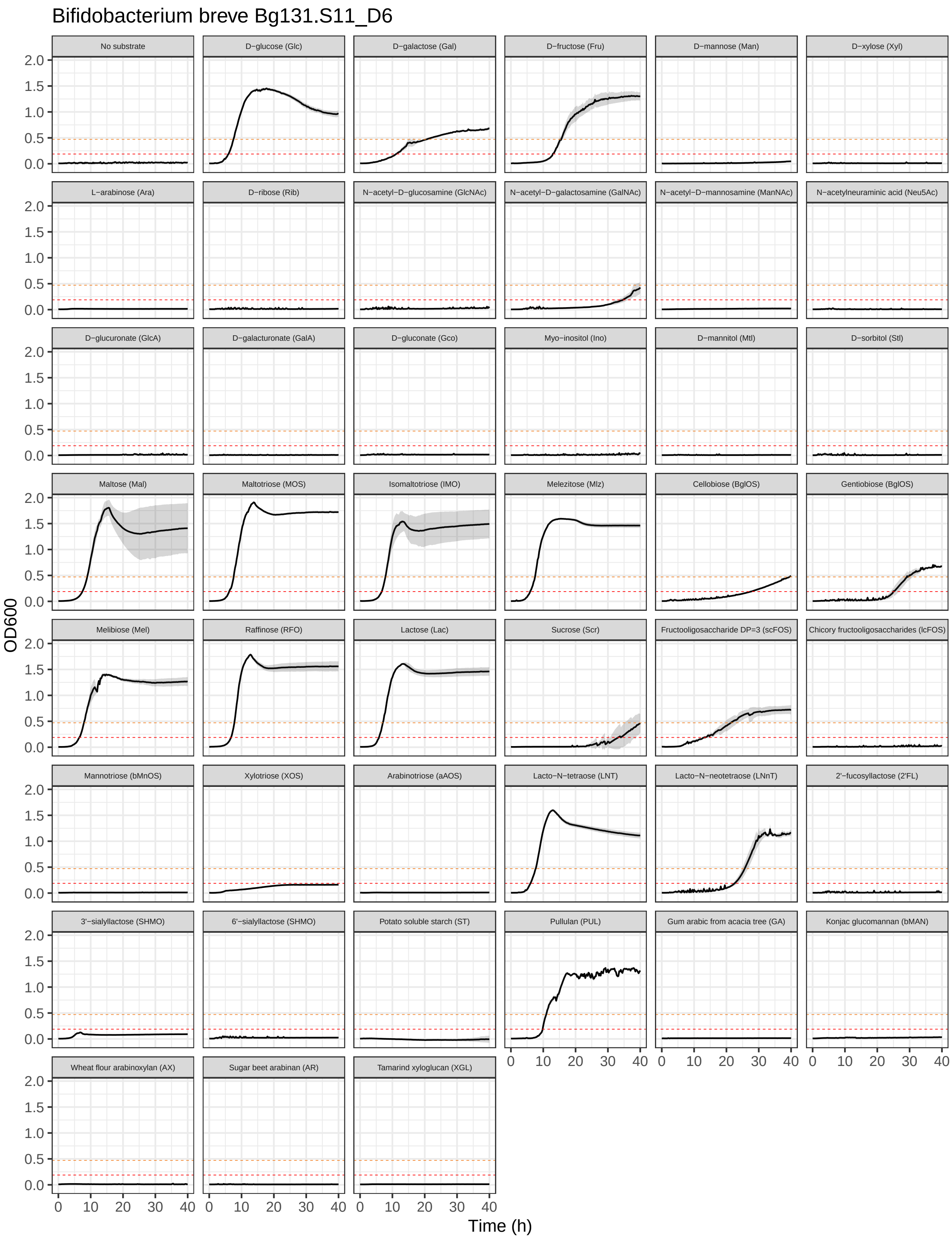

Bifidobacterium breve Bg131.S11_D6
No substrate
D−glucose (Glc)
D−galactose (Gal)
D−fructose (Fru)
D−mannose (Man)
D−xylose (Xyl)
L−arabinose (Ara)
D−ribose (Rib)
N−acetyl−D−glucosamine (GlcNAc)
N−acetyl−D−galactosamine (GalNAc)
N−acetyl−D−mannosamine (ManNAc)
N−acetylneuraminic acid (Neu5Ac)
D−glucuronate (GlcA)
D−galacturonate (GalA)
D−gluconate (Gco)
Myo−inositol (Ino)
D−mannitol (Mtl)
D−sorbitol (Stl)
Maltose (Mal)
Maltotriose (MOS)
Isomaltotriose (IMO)
Melezitose (Mlz)
Cellobiose (BglOS)
Gentiobiose (BglOS)
Melibiose (Mel)
Raffinose (RFO)
Lactose (Lac)
Sucrose (Scr)
Fructooligosaccharide DP=3 (scFOS)
Chicory fructooligosaccharides (lcFOS)
Mannotriose (bMnOS)
Xylotriose (XOS)
Arabinotriose (aAOS)
Lacto−N−tetraose (LNT)
Lacto−N−neotetraose (LNnT)
2'−fucosyllactose (2'FL)
3'−sialyllactose (SHMO)
6'−sialyllactose (SHMO)
Potato soluble starch (ST)
Pullulan (PUL)
Gum arabic from acacia tree (GA)
Konjac glucomannan (bMAN)
Wheat flour arabinoxylan (AX)
Sugar beet arabinan (AR)
Tamarind xyloglucan (XGL)
OD600
2.0
1.5
1.0
0.5
0.0
0
10
20
30
40
Time (h)

#### 11.7 *B. breve* Bg155.S08_4F7

Growth curves of *B. breve* Bg155.S08_4F7 in MRS-AC supplemented with various (n = 47) carbon sources.

```
growth_curves("Bifidobacterium breve Bg155.S08_4F7",
              "data/growth/formatted/Bbreve_Bg155.S08_4F7.txt",
              "results/growth/growth_curves/Bbreve_Bg155.S08_4F7.pdf",
              40)
```

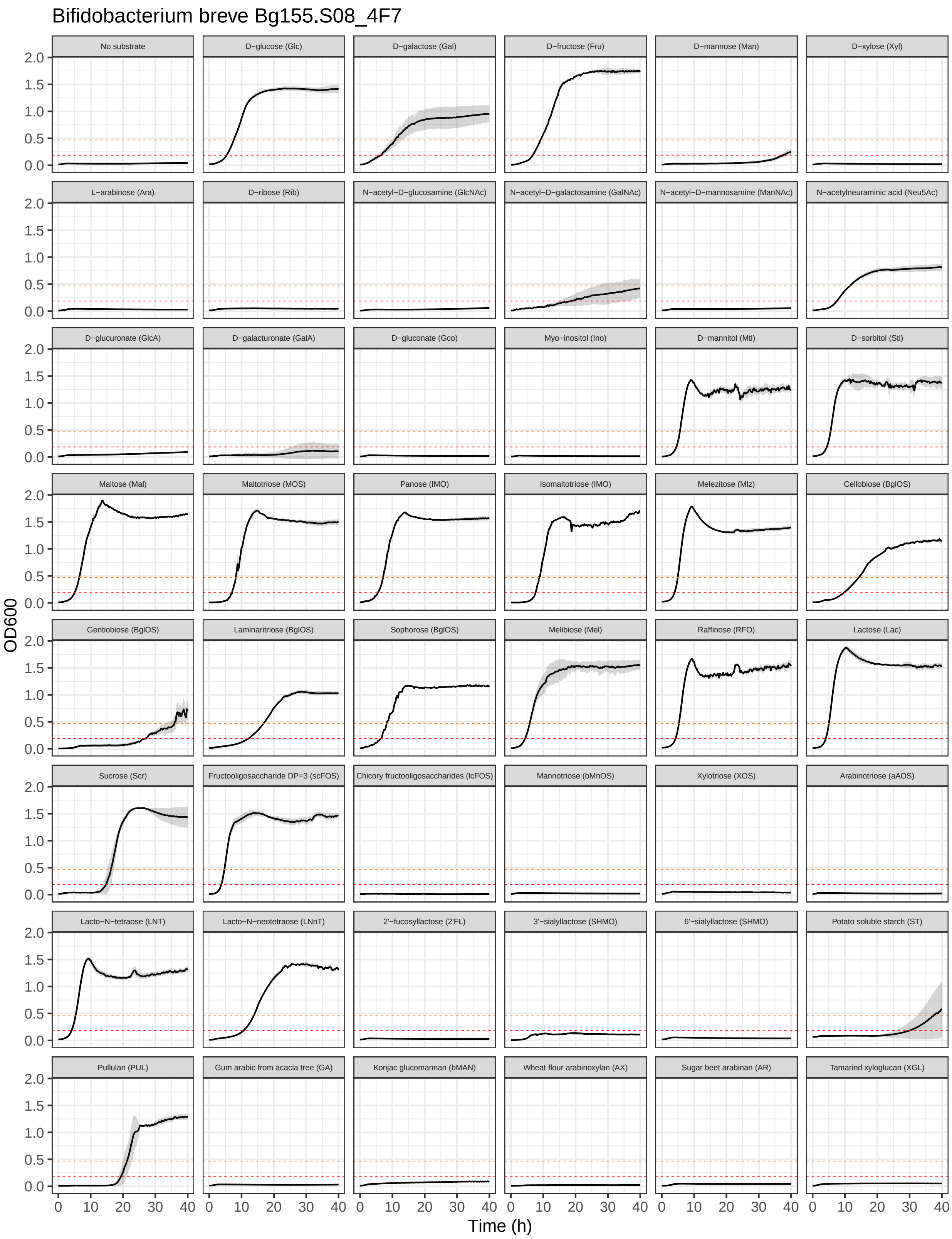
Bifidobacterium breve Bg155.S08_4F7
No substrate
D−glucose (Glc)
D−galactose (Gal)
D−fructose (Fru)
D−mannose (Man)
D−xylose (Xyl)
L−arabinose (Ara)
D−ribose (Rib)
N−acetyl−D−glucosamine (GlcNAc)
N−acetyl−D−galactosamine (GalNAc)
N−acetyl−D−mannosamine (ManNAc)
N−acetylneuraminic acid (Neu5Ac)
D−glucuronate (GlcA)
D−galacturonate (GalA)
D−gluconate (Gco)
Myo−inositol (Ino)
D−mannitol (Mtl)
D−sorbitol (Stl)
Maltose (Mal)
Maltotriose (MOS)
Panose (IMO)
Isomaltotriose (IMO)
Melezitose (Mlz)
Cellobiose (BglOS)
Gentiobiose (BglOS)
Laminaritriose (BglOS)
Sophorose (BglOS)
Melibiose (Mel)
Raffinose (RFO)
Lactose (Lac)
Sucrose (Scr)
Fructooligosaccharide DP=3 (scFOS)
Chicory fructooligosaccharides (lcFOS)
Mannotriose (bMnOS)
Xylotriose (XOS)
Arabinotriose (aAOS)
Lacto−N−tetraose (LNT)
Lacto−N−neotetraose (LNnT)
2'−fucosyllactose (2'FL)
3'−sialyllactose (SHMO)
6'−sialyllactose (SHMO)
Potato soluble starch (ST)
Pullulan (PUL)
Gum arabic from acacia tree (GA)
Konjac glucomannan (bMAN)
Wheat flour arabinoxylan (AX)
Sugar beet arabinan (AR)
Tamarind xyloglucan (XGL)
OD600
0.0
0.5
1.0
1.5
2.0
0
10
20
30
40
Time (h)

#### 11.8 *B. breve* Bg41721_1C11

Growth curves of *B. breve* Bg41721_1C11 in MRS-AC supplemented with various (n = 47) carbon sources.

```
growth_curves("Bifidobacterium breve Bg41721_1C11",
              "data/growth/formatted/Bbreve_Bg41721_1C11.txt",
              "results/growth/growth_curves/Bbreve_Bg41721_1C11.pdf",
              40)
```

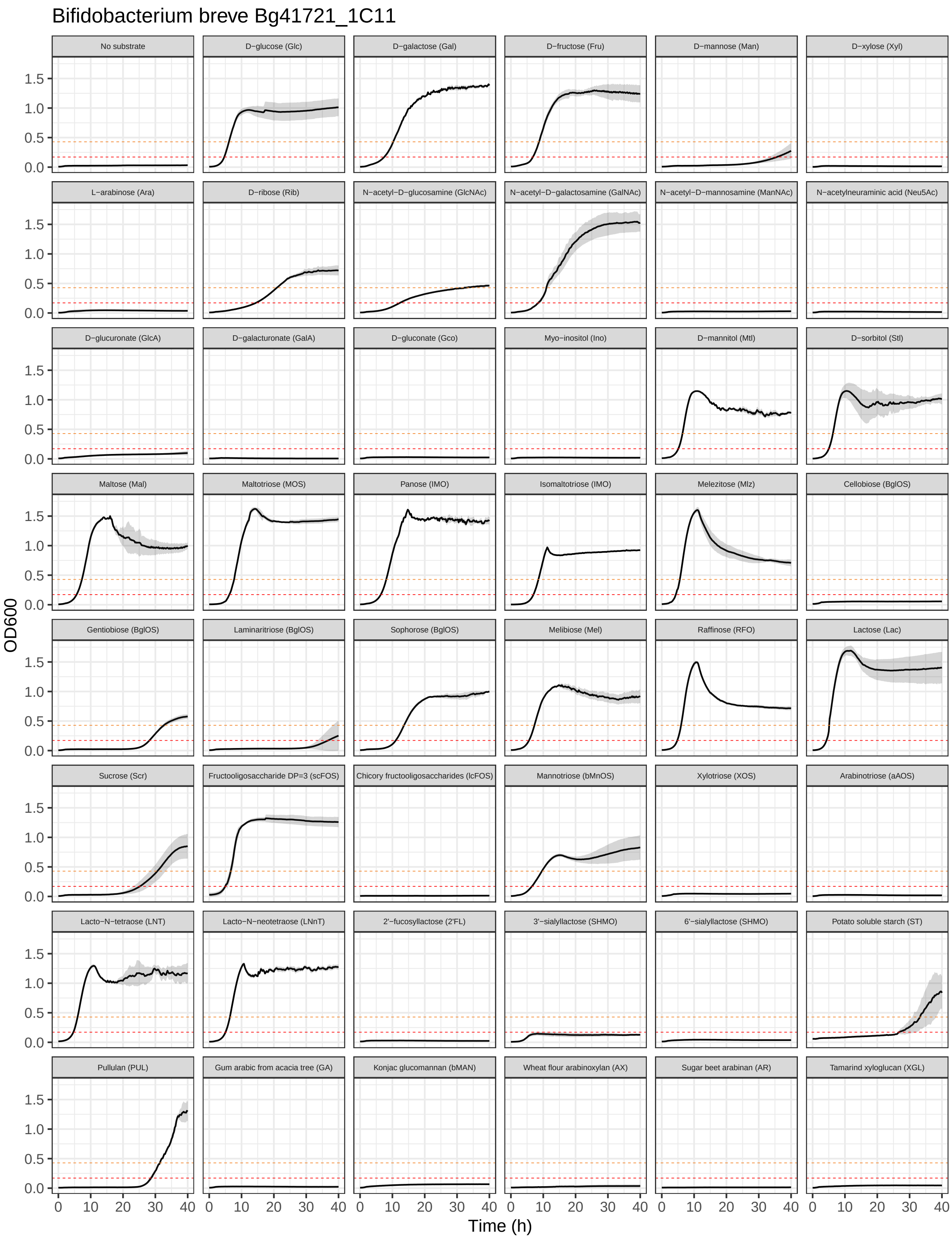
Bifidobacterium breve Bg41721_1C11
No substrate
D−glucose (Glc)
D−galactose (Gal)
D−fructose (Fru)
D−mannose (Man)
D−xylose (Xyl)
L−arabinose (Ara)
D−ribose (Rib)
N−acetyl−D−glucosamine (GlcNAc)
N−acetyl−D−galactosamine (GalNAc)
N−acetyl−D−mannosamine (ManNAc)
N−acetylneuraminic acid (Neu5Ac)
D−glucuronate (GlcA)
D−galacturonate (GalA)
D−gluconate (Gco)
Myo−inositol (Ino)
D−mannitol (Mtl)
D−sorbitol (Stl)
Maltose (Mal)
Maltotriose (MOS)
Panose (IMO)
Isomaltotriose (IMO)
Melezitose (Mlz)
Cellobiose (BglOS)
Gentiobiose (BglOS)
Laminaritriose (BglOS)
Sophorose (BglOS)
Melibiose (Mel)
Raffinose (RFO)
Lactose (Lac)
Sucrose (Scr)
Fructooligosaccharide DP=3 (scFOS)
Chicory fructooligosaccharides (lcFOS)
Mannotriose (bMnOS)
Xylotriose (XOS)
Arabinotriose (aAOS)
Lacto−N−tetraose (LNT)
Lacto−N−neotetraose (LNnT)
2'−fucosyllactose (2'FL)
3'−sialyllactose (SHMO)
6'−sialyllactose (SHMO)
Potato soluble starch (ST)
Pullulan (PUL)
Gum arabic from acacia tree (GA)
Konjac glucomannan (bMAN)
Wheat flour arabinoxylan (AX)
Sugar beet arabinan (AR)
Tamarind xyloglucan (XGL)
OD600
0.0
0.5
1.0
1.5
0
10
20
30
40
Time (h)

#### 11.9 *B. breve* JG_Bg463

Growth curves of *B. breve* JG_Bg463 in MRS-AC supplemented with various (n = 47) carbon sources.

```
growth_curves("Bifidobacterium breve JG_Bg463",
              "data/growth/formatted/Bbreve_JG_Bg463.txt",
              "results/growth/growth_curves/Bbreve_JG_Bg463.pdf",
              40)
```

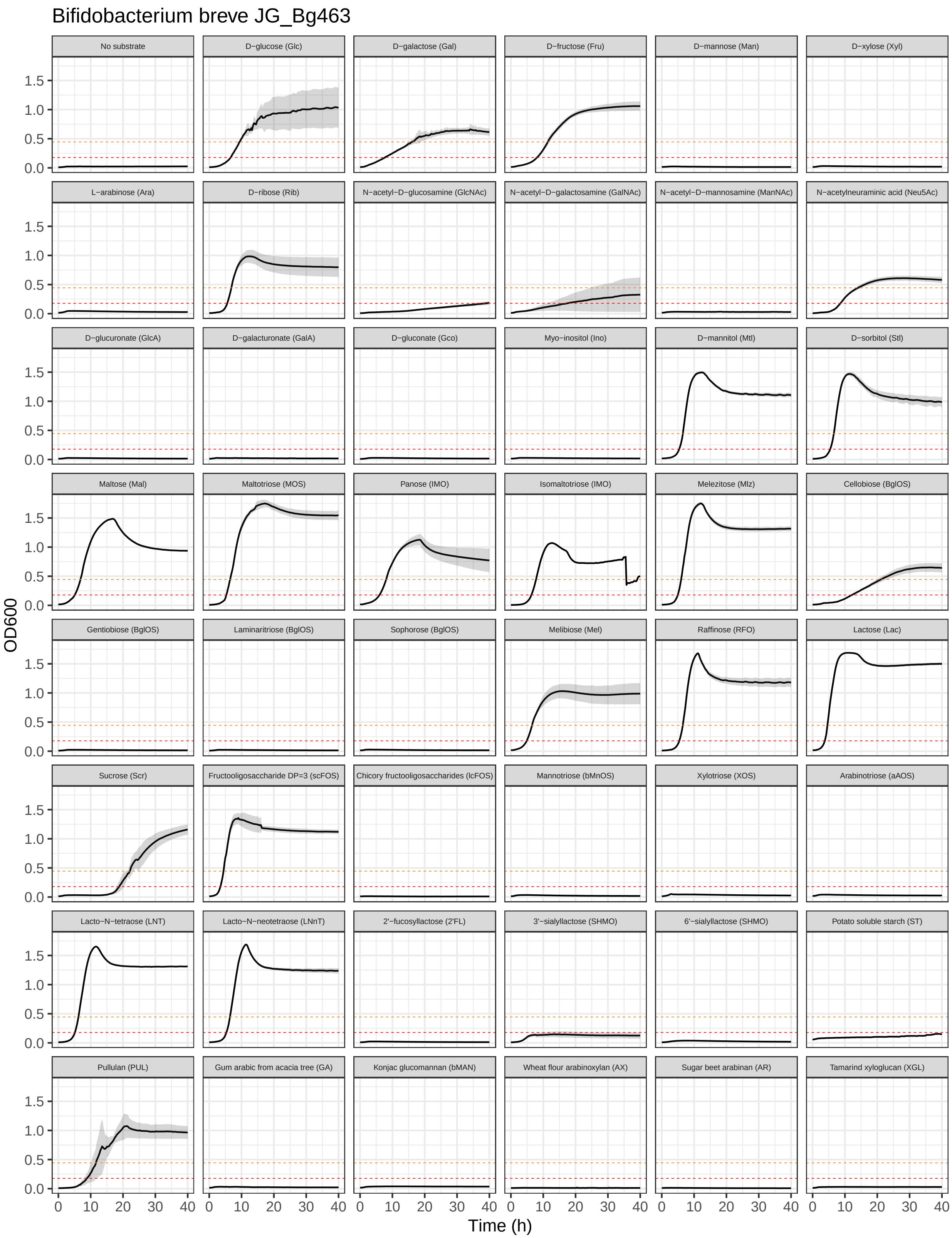
Bifidobacterium breve JG_Bg463
No substrate
D−glucose (Glc)
D−galactose (Gal)
D−fructose (Fru)
D−mannose (Man)
D−xylose (Xyl)
L−arabinose (Ara)
D−ribose (Rib)
N−acetyl−D−glucosamine (GlcNAc)
N−acetyl−D−galactosamine (GalNAc)
N−acetyl−D−mannosamine (ManNAc)
N−acetylneuraminic acid (Neu5Ac)
D−glucuronate (GlcA)
D−galacturonate (GalA)
D−gluconate (Gco)
Myo−inositol (Ino)
D−mannitol (Mtl)
D−sorbitol (Stl)
Maltose (Mal)
Maltotriose (MOS)
Panose (IMO)
Isomaltotriose (IMO)
Melezitose (Mlz)
Cellobiose (BglOS)
Gentiobiose (BglOS)
Laminaritriose (BglOS)
Sophorose (BglOS)
Melibiose (Mel)
Raffinose (RFO)
Lactose (Lac)
Sucrose (Scr)
Fructooligosaccharide DP=3 (scFOS)
Chicory fructooligosaccharides (lcFOS)
Mannotriose (bMnOS)
Xylotriose (XOS)
Arabinotriose (aAOS)
Lacto−N−tetraose (LNT)
Lacto−N−neotetraose (LNnT)
2'−fucosyllactose (2'FL)
3'−sialyllactose (SHMO)
6'−sialyllactose (SHMO)
Potato soluble starch (ST)
Pullulan (PUL)
Gum arabic from acacia tree (GA)
Konjac glucomannan (bMAN)
Wheat flour arabinoxylan (AX)
Sugar beet arabinan (AR)
Tamarind xyloglucan (XGL)
OD600
0.0
0.5
1.0
1.5
Time (h)
0
10
20
30
40

#### 11.10 *B. breve* LFYP81

Growth curves of *B. breve* LFYP81 in MRS-AC supplemented with various (n = 47) carbon sources.

```
growth_curves("Bifidobacterium breve LFYP81",
              "data/growth/formatted/Bbreve_LFYP81.txt",
              "results/growth/growth_curves/Bbreve_LFYP81.pdf",
              40)
```

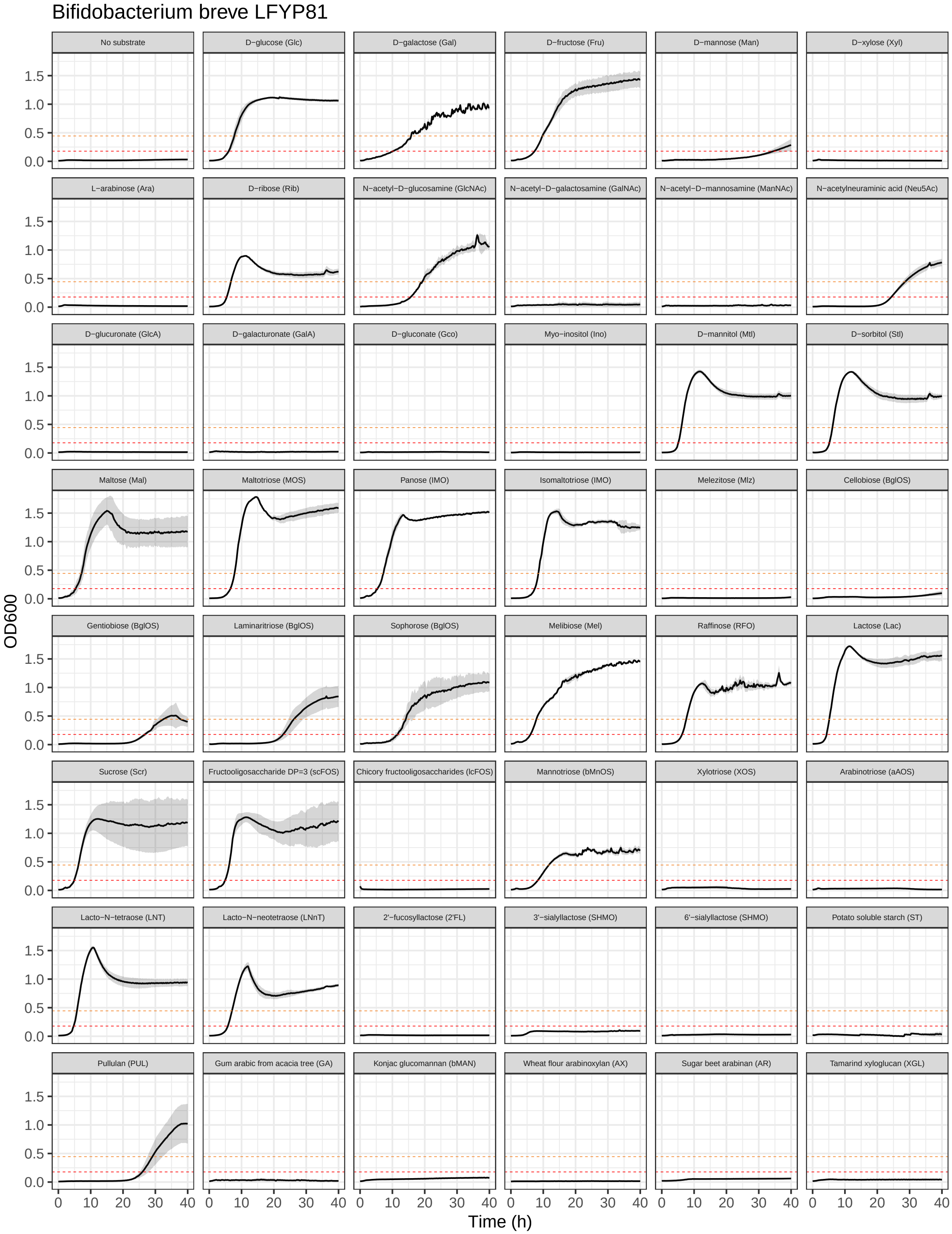

Bifidobacterium breve LFYP81
No substrate
D−glucose (Glc)
D−galactose (Gal)
D−fructose (Fru)
D−mannose (Man)
D−xylose (Xyl)
L−arabinose (Ara)
D−ribose (Rib)
N−acetyl−D−glucosamine (GlcNAc)
N−acetyl−D−galactosamine (GalNAc)
N−acetyl−D−mannosamine (ManNAc)
N−acetylneuraminic acid (Neu5Ac)
D−glucuronate (GlcA)
D−galacturonate (GalA)
D−gluconate (Gco)
Myo−inositol (Ino)
D−mannitol (Mtl)
D−sorbitol (Stl)
Maltose (Mal)
Maltotriose (MOS)
Panose (IMO)
Isomaltotriose (IMO)
Melezitose (Mlz)
Cellobiose (BglOS)
Gentiobiose (BglOS)
Laminaritriose (BglOS)
Sophorose (BglOS)
Melibiose (Mel)
Raffinose (RFO)
Lactose (Lac)
Sucrose (Scr)
Fructooligosaccharide DP=3 (scFOS)
Chicory fructooligosaccharides (lcFOS)
Mannotriose (bMnOS)
Xylotriose (XOS)
Arabinotriose (aAOS)
Lacto−N−tetraose (LNT)
Lacto−N−neotetraose (LNnT)
2'−fucosyllactose (2'FL)
3'−sialyllactose (SHMO)
6'−sialyllactose (SHMO)
Potato soluble starch (ST)
Pullulan (PUL)
Gum arabic from acacia tree (GA)
Konjac glucomannan (bMAN)
Wheat flour arabinoxylan (AX)
Sugar beet arabinan (AR)
Tamarind xyloglucan (XGL)
OD600
0.0
0.5
1.0
1.5
0
10
20
30
40
Time (h)

##### 11.11 *B. breve* M257B_A4

Growth curves of *B. breve* M257B_A4 in MRS-AC supplemented with various (n = 45) carbon sources.

```r
growth_curves("Bifidobacterium breve M257B_A4",
              "data/growth/formatted/Bbreve_M257B_A4.txt",
              "results/growth/growth_curves/Bbreve_M257B_A4.pdf",
              40)
```

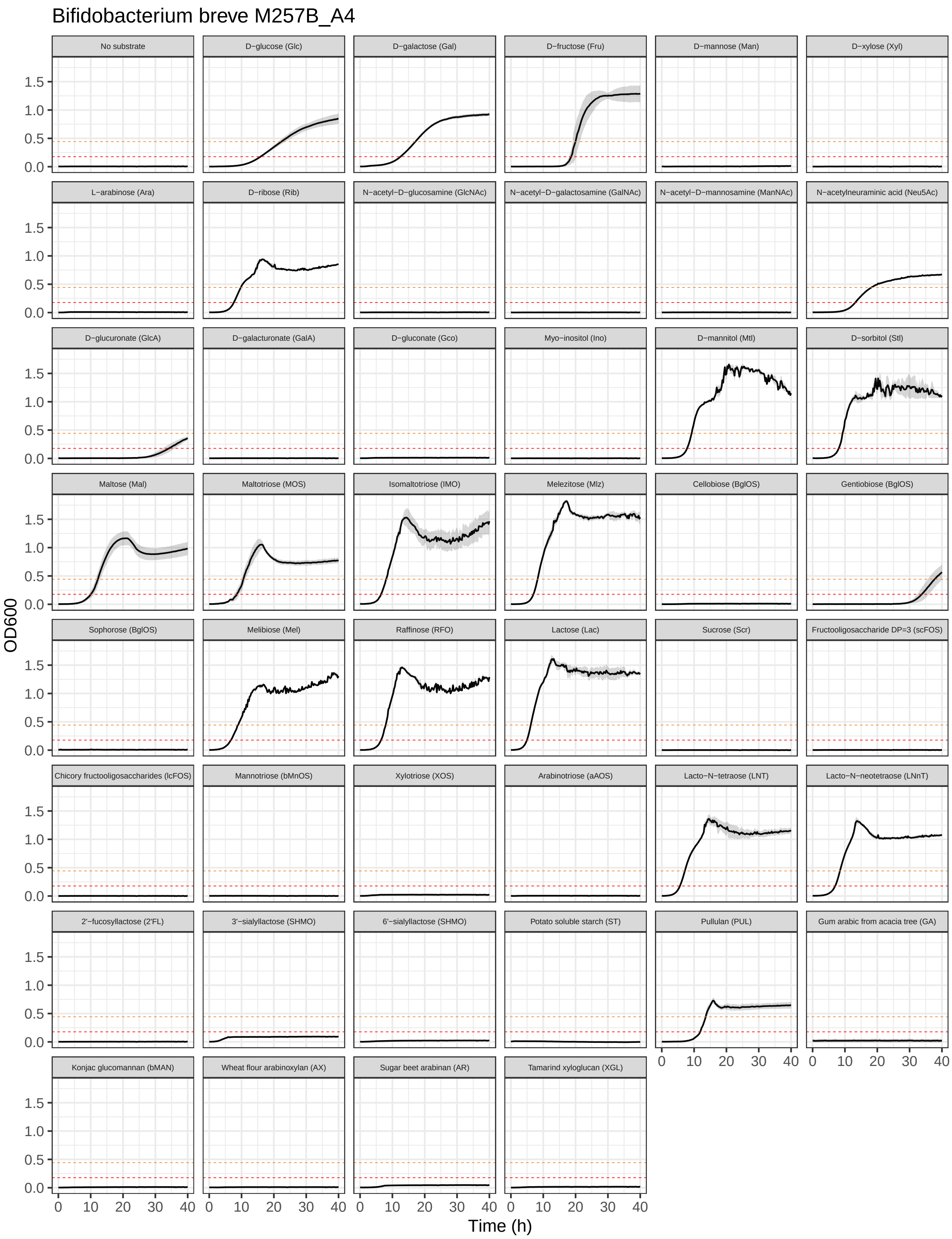

Bifidobacterium breve M257B_A4
No substrate
D−glucose (Glc)
D−galactose (Gal)
D−fructose (Fru)
D−mannose (Man)
D−xylose (Xyl)
L−arabinose (Ara)
D−ribose (Rib)
N−acetyl−D−glucosamine (GlcNAc)
N−acetyl−D−galactosamine (GalNAc)
N−acetyl−D−mannosamine (ManNAc)
N−acetylneuraminic acid (Neu5Ac)
D−glucuronate (GlcA)
D−galacturonate (GalA)
D−gluconate (Gco)
Myo−inositol (Ino)
D−mannitol (Mtl)
D−sorbitol (Stl)
Maltose (Mal)
Maltotriose (MOS)
Isomaltotriose (IMO)
Melezitose (Mlz)
Cellobiose (BglOS)
Gentiobiose (BglOS)
Sophorose (BglOS)
Melibiose (Mel)
Raffinose (RFO)
Lactose (Lac)
Sucrose (Scr)
Fructooligosaccharide DP=3 (scFOS)
Chicory fructooligosaccharides (lcFOS)
Mannotriose (bMnOS)
Xylotriose (XOS)
Arabinotriose (aAOS)
Lacto−N−tetraose (LNT)
Lacto−N−neotetraose (LNnT)
2'−fucosyllactose (2'FL)
3'−sialyllactose (SHMO)
6'−sialyllactose (SHMO)
Potato soluble starch (ST)
Pullulan (PUL)
Gum arabic from acacia tree (GA)
Konjac glucomannan (bMAN)
Wheat flour arabinoxylan (AX)
Sugar beet arabinan (AR)
Tamarind xyloglucan (XGL)
OD600
0.0
0.5
1.0
1.5
0
10
20
30
40
Time (h)

##### 11.12 *B. catenulatum* subsp. *kashiwanohense* Bg42221_1E1

Growth curves of *B. catenulatum* subsp. *kashiwanohense* Bg42221_1E1 in MRS-AC supplemented with various (n = 47) carbon sources.

```
growth_curves("Bifidobacterium catenulatum subsp. kashiwanohense Bg42221_1E1",
              "data/growth/formatted/Bc_kashiwanohense_Bg42221_1E1.txt",
              "results/growth/growth_curves/Bc_kashiwanohense_Bg42221_1E1.pdf",
              40)
```

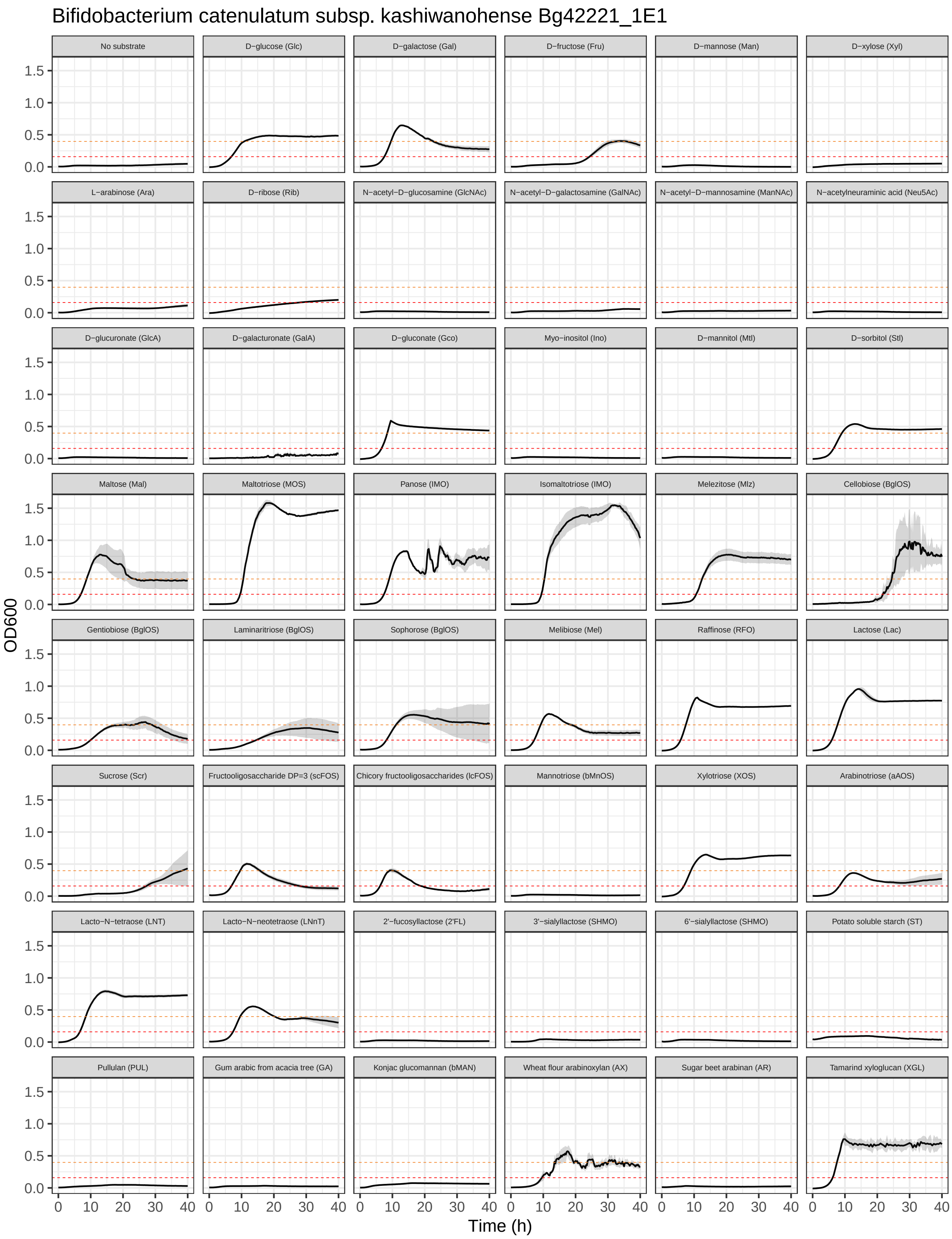

Bifidobacterium catenulatum subsp. kashiwanohense Bg42221_1E1
No substrate
D−glucose (Glc)
D−galactose (Gal)
D−fructose (Fru)
D−mannose (Man)
D−xylose (Xyl)
L−arabinose (Ara)
D−ribose (Rib)
N−acetyl−D−glucosamine (GlcNAc)
N−acetyl−D−galactosamine (GalNAc)
N−acetyl−D−mannosamine (ManNAc)
N−acetylneuraminic acid (Neu5Ac)
D−glucuronate (GlcA)
D−galacturonate (GalA)
D−gluconate (Gco)
Myo−inositol (Ino)
D−mannitol (Mtl)
D−sorbitol (Stl)
Maltose (Mal)
Maltotriose (MOS)
Panose (IMO)
Isomaltotriose (IMO)
Melezitose (Mlz)
Cellobiose (BglOS)
Gentiobiose (BglOS)
Laminaritriose (BglOS)
Sophorose (BglOS)
Melibiose (Mel)
Raffinose (RFO)
Lactose (Lac)
Sucrose (Scr)
Fructooligosaccharide DP=3 (scFOS)
Chicory fructooligosaccharides (lcFOS)
Mannotriose (bMnOS)
Xylotriose (XOS)
Arabinotriose (aAOS)
Lacto−N−tetraose (LNT)
Lacto−N−neotetraose (LNnT)
2'−fucosyllactose (2'FL)
3'−sialyllactose (SHMO)
6'−sialyllactose (SHMO)
Potato soluble starch (ST)
Pullulan (PUL)
Gum arabic from acacia tree (GA)
Konjac glucomannan (bMAN)
Wheat flour arabinoxylan (AX)
Sugar beet arabinan (AR)
Tamarind xyloglucan (XGL)
OD600
Time (h)
0.0
0.5
1.0
1.5
0
10
20
30
40

##### 11.13 *B. catenulatum* subsp. *kashiwanohense_A* Bg42221_1D3

Growth curves of *B. catenulatum* subsp. *kashiwanohense_A* Bg42221_1D3 in MRS-AC supplemented with various (n = 43) carbon sources.

```
growth_curves("Bifidobacterium catenulatum subsp. kashiwanohense_A Bg42221_1D3",
              "data/growth/formatted/Bc_kashiwanohense_A Bg42221_1D3.txt",
              "results/growth/growth_curves/Bc_kashiwanohense_A Bg42221_1D3.pdf",
              40)
```

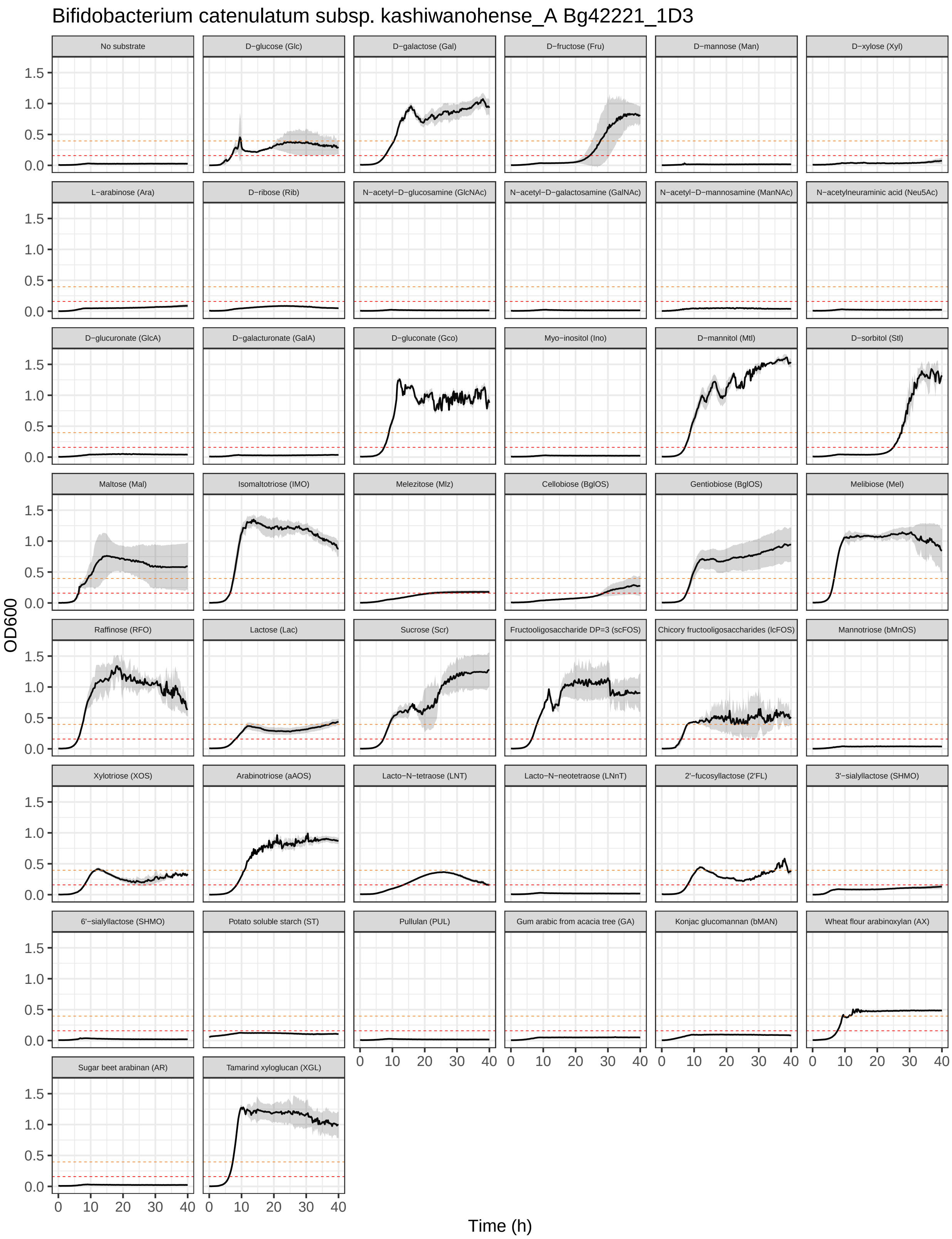

Bifidobacterium catenulatum subsp. kashiwanohense_A Bg42221_1D3
No substrate
D−glucose (Glc)
D−galactose (Gal)
D−fructose (Fru)
D−mannose (Man)
D−xylose (Xyl)
L−arabinose (Ara)
D−ribose (Rib)
N−acetyl−D−glucosamine (GlcNAc)
N−acetyl−D−galactosamine (GalNAc)
N−acetyl−D−mannosamine (ManNAc)
N−acetylneuraminic acid (Neu5Ac)
D−glucuronate (GlcA)
D−galacturonate (GalA)
D−gluconate (Gco)
Myo−inositol (Ino)
D−mannitol (Mtl)
D−sorbitol (Stl)
Maltose (Mal)
Isomaltotriose (IMO)
Melezitose (Mlz)
Cellobiose (BglOS)
Gentiobiose (BglOS)
Melibiose (Mel)
Raffinose (RFO)
Lactose (Lac)
Sucrose (Scr)
Fructooligosaccharide DP=3 (scFOS)
Chicory fructooligosaccharides (lcFOS)
Mannotriose (bMnOS)
Xylotriose (XOS)
Arabinotriose (aAOS)
Lacto−N−tetraose (LNT)
Lacto−N−neotetraose (LNnT)
2'−fucosyllactose (2'FL)
3'−sialyllactose (SHMO)
6'−sialyllactose (SHMO)
Potato soluble starch (ST)
Pullulan (PUL)
Gum arabic from acacia tree (GA)
Konjac glucomannan (bMAN)
Wheat flour arabinoxylan (AX)
Sugar beet arabinan (AR)
Tamarind xyloglucan (XGL)
OD600
0.0
0.5
1.0
1.5
0
10
20
30
40
Time (h)

#### 11.14 *B. dentium* LFYP24

Growth curves of *B. dentium* LFYP24 in MRS-AC supplemented with various (n = 47) carbon sources.

```
growth_curves("Bifidobacterium dentium LFYP24",
              "data/growth/formatted/Bdentium_LFYP24.txt",
              "results/growth/growth_curves/Bdentium_LFYP24.pdf",
              24)
```

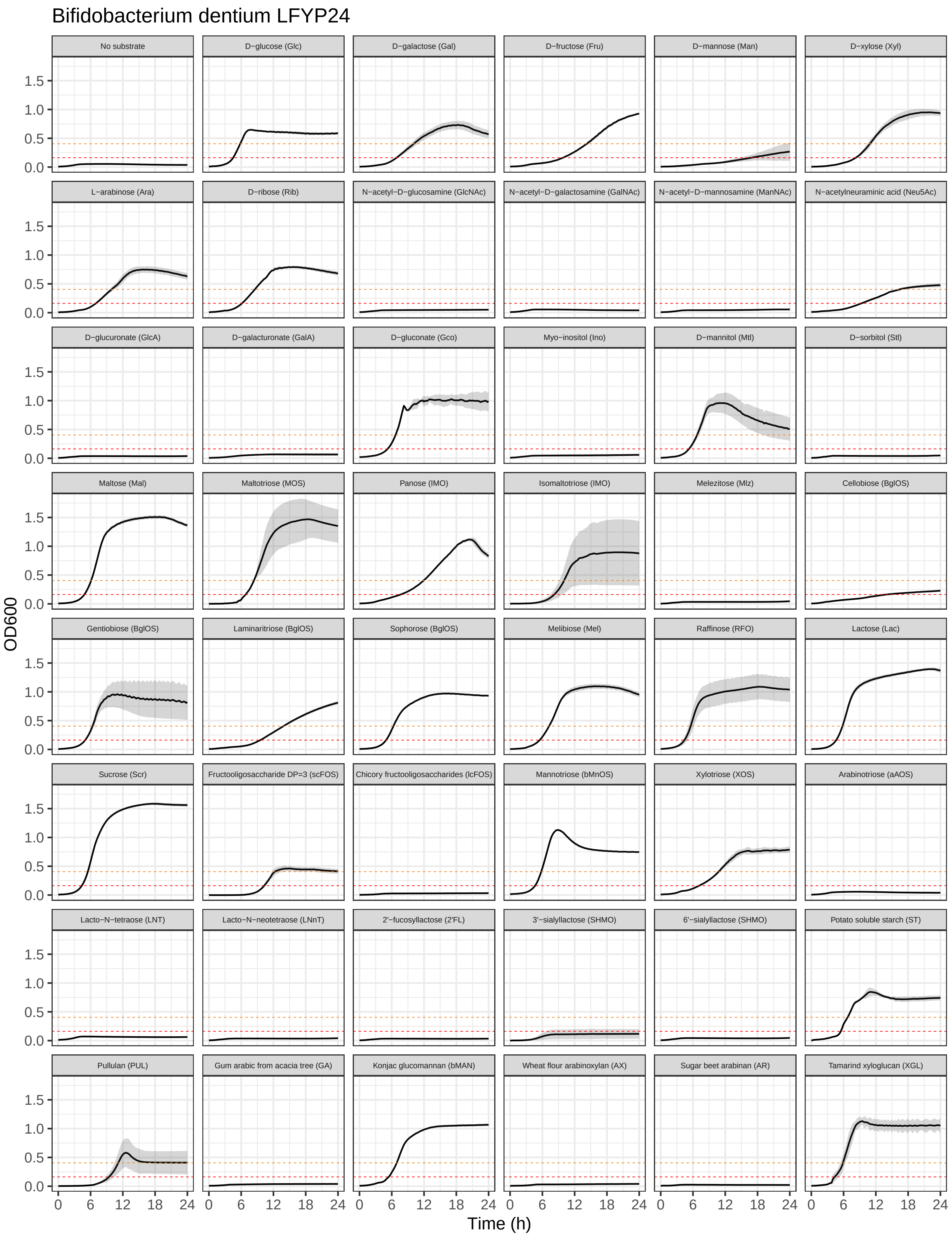
Bifidobacterium dentium LFYP24
No substrate
D−glucose (Glc)
D−galactose (Gal)
D−fructose (Fru)
D−mannose (Man)
D−xylose (Xyl)
L−arabinose (Ara)
D−ribose (Rib)
N−acetyl−D−glucosamine (GlcNAc)
N−acetyl−D−galactosamine (GalNAc)
N−acetyl−D−mannosamine (ManNAc)
N−acetylneuraminic acid (Neu5Ac)
D−glucuronate (GlcA)
D−galacturonate (GalA)
D−gluconate (Gco)
Myo−inositol (Ino)
D−mannitol (Mtl)
D−sorbitol (Stl)
Maltose (Mal)
Maltotriose (MOS)
Panose (IMO)
Isomaltotriose (IMO)
Melezitose (Mlz)
Cellobiose (BglOS)
Gentiobiose (BglOS)
Laminaritriose (BglOS)
Sophorose (BglOS)
Melibiose (Mel)
Raffinose (RFO)
Lactose (Lac)
Sucrose (Scr)
Fructooligosaccharide DP=3 (scFOS)
Chicory fructooligosaccharides (lcFOS)
Mannotriose (bMnOS)
Xylotriose (XOS)
Arabinotriose (aAOS)
Lacto−N−tetraose (LNT)
Lacto−N−neotetraose (LNnT)
2'−fucosyllactose (2'FL)
3'−sialyllactose (SHMO)
6'−sialyllactose (SHMO)
Potato soluble starch (ST)
Pullulan (PUL)
Gum arabic from acacia tree (GA)
Konjac glucomannan (bMAN)
Wheat flour arabinoxylan (AX)
Sugar beet arabinan (AR)
Tamarind xyloglucan (XGL)
OD600
Time (h)
0.0
0.5
1.0
1.5
0
6
12
18
24

#### 11.15 *B. hominis* Bg064.11__2H10

Growth curves of *B. hominis* Bg064.11__2H10 in MRS-AC supplemented with various (n = 44) carbon sources.

```
growth_curves("Bifidobacterium hominis Bg064.11_2H10",
              "data/growth/formatted/Bhominis_Bg064.11_2H10.txt",
              "results/growth/growth_curves/Bhominis_Bg064.11_2H10.pdf",
              40)
```

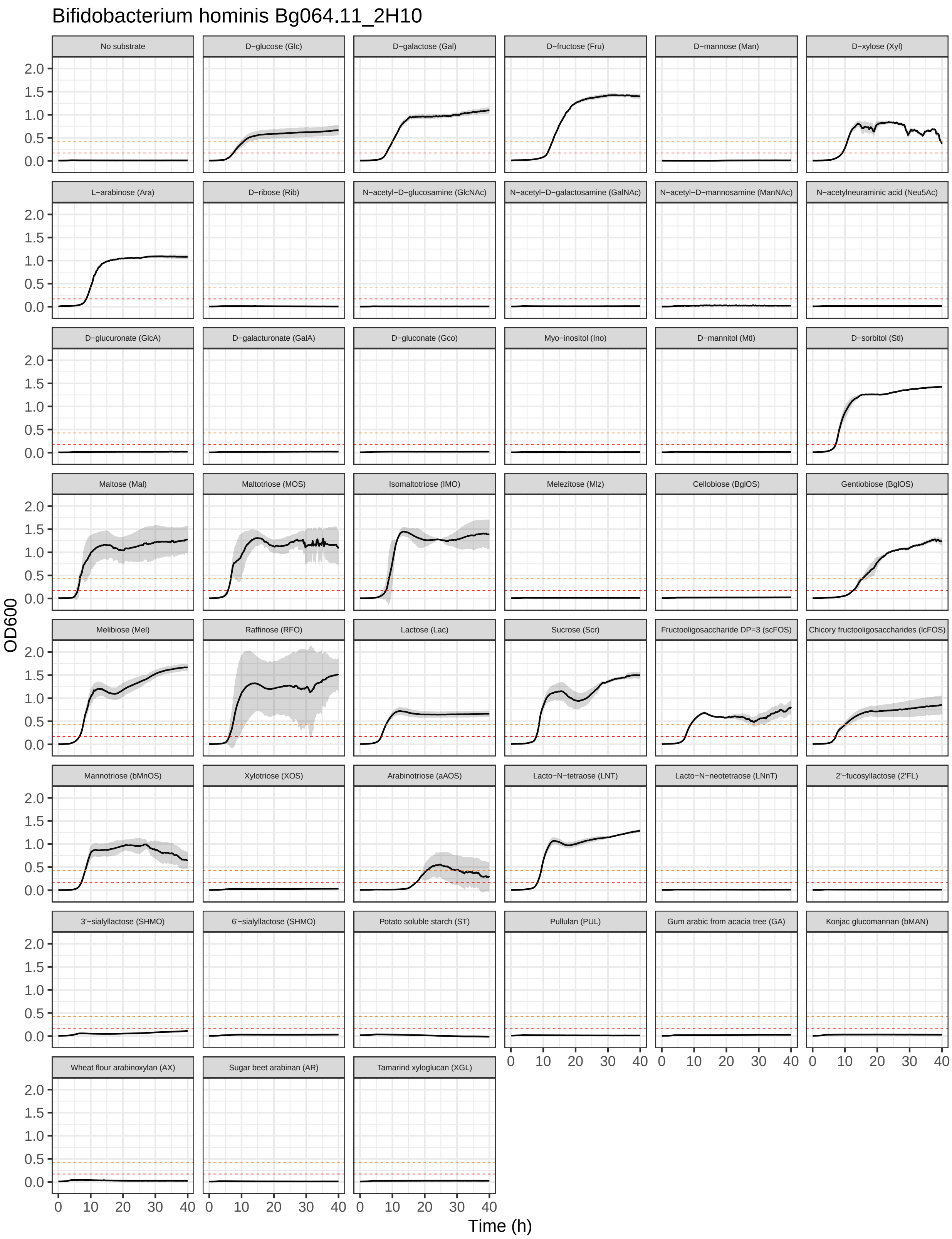

Bifidobacterium hominis Bg064.11_2H10
No substrate
D−glucose (Glc)
D−galactose (Gal)
D−fructose (Fru)
D−mannose (Man)
D−xylose (Xyl)
L−arabinose (Ara)
D−ribose (Rib)
N−acetyl−D−glucosamine (GlcNAc)
N−acetyl−D−galactosamine (GalNAc)
N−acetyl−D−mannosamine (ManNAc)
N−acetylneuraminic acid (Neu5Ac)
D−glucuronate (GlcA)
D−galacturonate (GalA)
D−gluconate (Gco)
Myo−inositol (Ino)
D−mannitol (Mtl)
D−sorbitol (Stl)
Maltose (Mal)
Maltotriose (MOS)
Isomaltotriose (IMO)
Melezitose (Mlz)
Cellobiose (BglOS)
Gentiobiose (BglOS)
Melibiose (Mel)
Raffinose (RFO)
Lactose (Lac)
Sucrose (Scr)
Fructooligosaccharide DP=3 (scFOS)
Chicory fructooligosaccharides (lcFOS)
Mannotriose (bMnOS)
Xylotriose (XOS)
Arabinotriose (aAOS)
Lacto−N−tetraose (LNT)
Lacto−N−neotetraose (LNnT)
2'−fucosyllactose (2'FL)
3'−sialyllactose (SHMO)
6'−sialyllactose (SHMO)
Potato soluble starch (ST)
Pullulan (PUL)
Gum arabic from acacia tree (GA)
Konjac glucomannan (bMAN)
Wheat flour arabinoxylan (AX)
Sugar beet arabinan (AR)
Tamarind xyloglucan (XGL)
OD600
0.0
0.5
1.0
1.5
2.0
0
10
20
30
40
Time (h)

#### 11.16 *B. hominis* Bg155.08_4B11

Growth curves of *B. hominis* Bg155.08_4B11 in MRS-AC supplemented with various (n = 45) carbon sources.

```
growth_curves("Bifidobacterium hominis Bg155.08_4B11",
              "data/growth/formatted/Bhominis_Bg155.08_4B11.txt",
              "results/growth/growth_curves/Bhominis_Bg155.08_4B11.pdf",
              40)
```

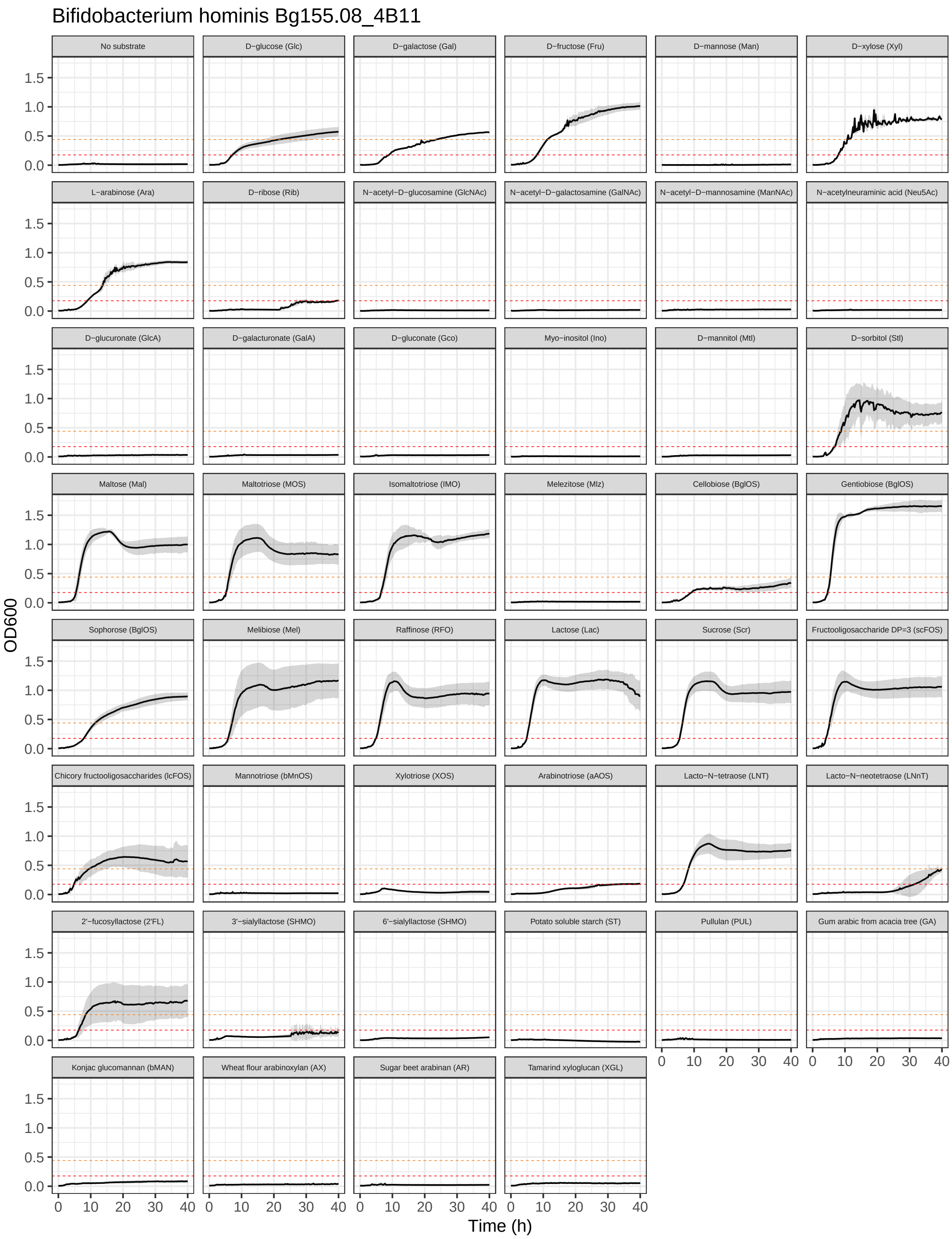
Bifidobacterium hominis Bg155.08_4B11
No substrate
D−glucose (Glc)
D−galactose (Gal)
D−fructose (Fru)
D−mannose (Man)
D−xylose (Xyl)
L−arabinose (Ara)
D−ribose (Rib)
N−acetyl−D−glucosamine (GlcNAc)
N−acetyl−D−galactosamine (GalNAc)
N−acetyl−D−mannosamine (ManNAc)
N−acetylneuraminic acid (Neu5Ac)
D−glucuronate (GlcA)
D−galacturonate (GalA)
D−gluconate (Gco)
Myo−inositol (Ino)
D−mannitol (Mtl)
D−sorbitol (Stl)
Maltose (Mal)
Maltotriose (MOS)
Isomaltotriose (IMO)
Melezitose (Mlz)
Cellobiose (BglOS)
Gentiobiose (BglOS)
Sophorose (BglOS)
Melibiose (Mel)
Raffinose (RFO)
Lactose (Lac)
Sucrose (Scr)
Fructooligosaccharide DP=3 (scFOS)
Chicory fructooligosaccharides (lcFOS)
Mannotriose (bMnOS)
Xylotriose (XOS)
Arabinotriose (aAOS)
Lacto−N−tetraose (LNT)
Lacto−N−neotetraose (LNnT)
2'−fucosyllactose (2'FL)
3'−sialyllactose (SHMO)
6'−sialyllactose (SHMO)
Potato soluble starch (ST)
Pullulan (PUL)
Gum arabic from acacia tree (GA)
Konjac glucomannan (bMAN)
Wheat flour arabinoxylan (AX)
Sugar beet arabinan (AR)
Tamarind xyloglucan (XGL)
OD600
0.0
0.5
1.0
1.5
0
10
20
30
40
Time (h)

##### 11.17 *B. hominis* M264_MC1

Growth curves of *B. hominis* M264_MC1 in MRS-AC supplemented with various (n = 43) carbon sources.

```
growth_curves("Bifidobacterium hominis M264_MC1",
              "data/growth/formatted/Bhominis_M264_MC1.txt",
              "results/growth/growth_curves/Bhominis_M264_MC1.pdf",
              40)
```

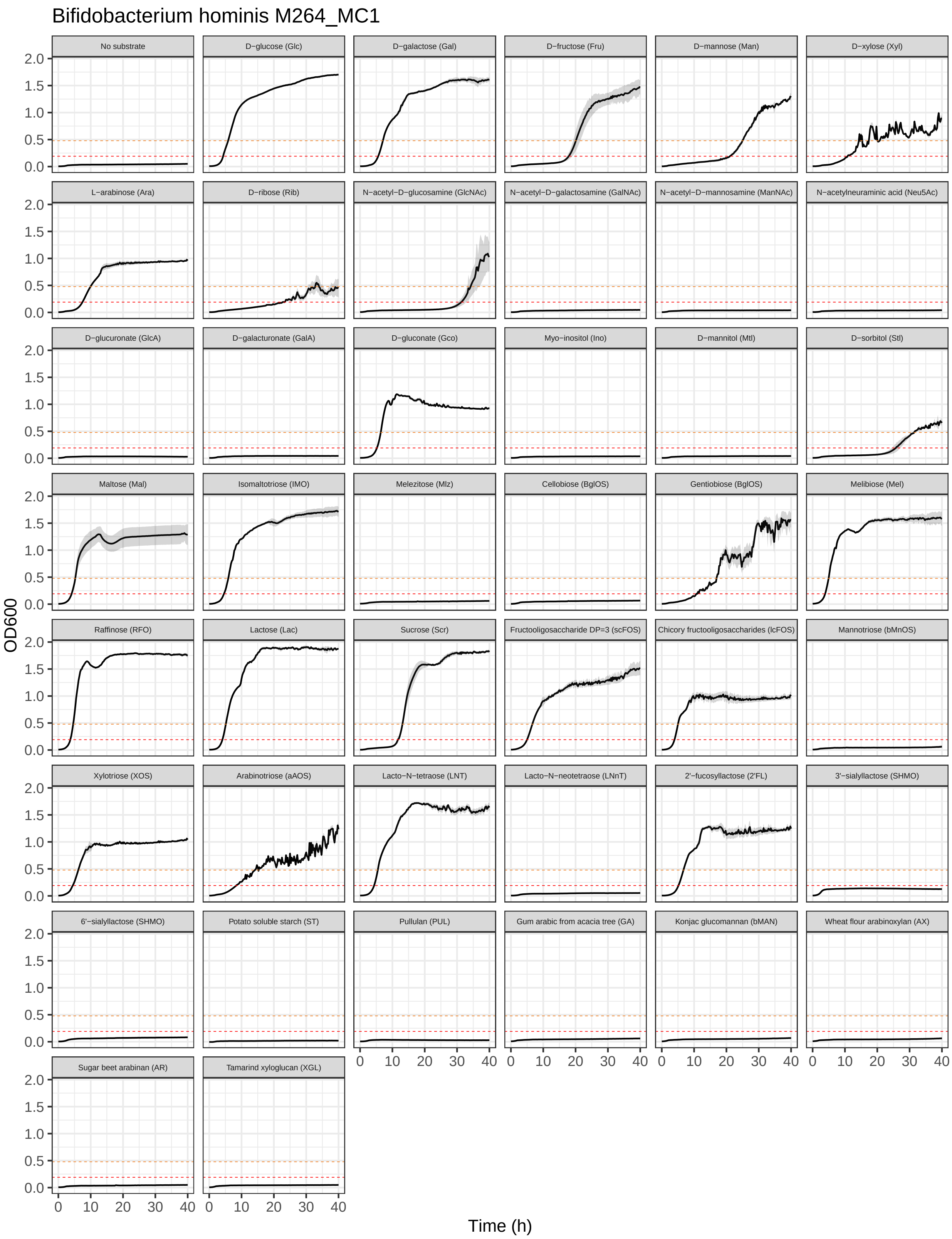
Bifidobacterium hominis M264_MC1
No substrate
D−glucose (Glc)
D−galactose (Gal)
D−fructose (Fru)
D−mannose (Man)
D−xylose (Xyl)
L−arabinose (Ara)
D−ribose (Rib)
N−acetyl−D−glucosamine (GlcNAc)
N−acetyl−D−galactosamine (GalNAc)
N−acetyl−D−mannosamine (ManNAc)
N−acetylneuraminic acid (Neu5Ac)
D−glucuronate (GlcA)
D−galacturonate (GalA)
D−gluconate (Gco)
Myo−inositol (Ino)
D−mannitol (Mtl)
D−sorbitol (Stl)
Maltose (Mal)
Isomaltotriose (IMO)
Melezitose (Mlz)
Cellobiose (BglOS)
Gentiobiose (BglOS)
Melibiose (Mel)
Raffinose (RFO)
Lactose (Lac)
Sucrose (Scr)
Fructooligosaccharide DP=3 (scFOS)
Chicory fructooligosaccharides (lcFOS)
Mannotriose (bMnOS)
Xylotriose (XOS)
Arabinotriose (aAOS)
Lacto−N−tetraose (LNT)
Lacto−N−neotetraose (LNnT)
2'−fucosyllactose (2'FL)
3'−sialyllactose (SHMO)
6'−sialyllactose (SHMO)
Potato soluble starch (ST)
Pullulan (PUL)
Gum arabic from acacia tree (GA)
Konjac glucomannan (bMAN)
Wheat flour arabinoxylan (AX)
Sugar beet arabinan (AR)
Tamarind xyloglucan (XGL)
OD600
Time (h)
0.0
0.5
1.0
1.5
2.0
0
10
20
30
40

#### 11.18 *B. longum* subsp. *infantis* ATCC 15697 = JCM 1222

Growth curves of *B. longum* subsp. *infantis* ATCC 15697 = JCM 1222 in MRS-AC supplemented with various (n = 47) carbon sources.

```r
growth_curves("Bifidobacterium longum subsp. infantis ATCC 15697 = JCM 1222",
              "data/growth/formatted/Bl_infantis_ATCC15697.txt",
              "results/growth/growth_curves/Bl_infantis_ATCC15697.pdf",
              40)
```

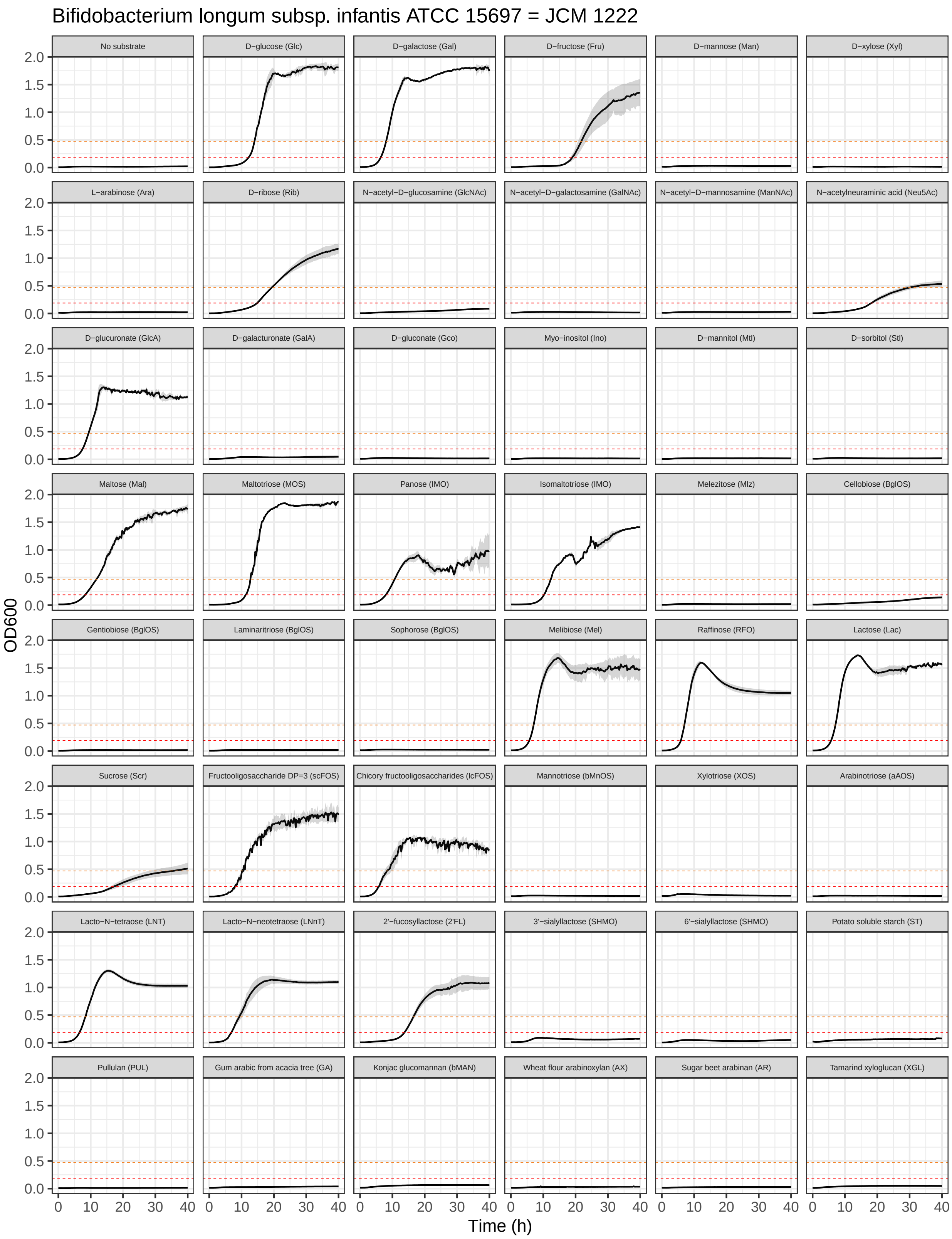
Bifidobacterium longum subsp. infantis ATCC 15697 = JCM 1222
No substrate
D−glucose (Glc)
D−galactose (Gal)
D−fructose (Fru)
D−mannose (Man)
D−xylose (Xyl)
L−arabinose (Ara)
D−ribose (Rib)
N−acetyl−D−glucosamine (GlcNAc)
N−acetyl−D−galactosamine (GalNAc)
N−acetyl−D−mannosamine (ManNAc)
N−acetylneuraminic acid (Neu5Ac)
D−glucuronate (GlcA)
D−galacturonate (GalA)
D−gluconate (Gco)
Myo−inositol (Ino)
D−mannitol (Mtl)
D−sorbitol (Stl)
Maltose (Mal)
Maltotriose (MOS)
Panose (IMO)
Isomaltotriose (IMO)
Melezitose (Mlz)
Cellobiose (BglOS)
Gentiobiose (BglOS)
Laminaritriose (BglOS)
Sophorose (BglOS)
Melibiose (Mel)
Raffinose (RFO)
Lactose (Lac)
Sucrose (Scr)
Fructooligosaccharide DP=3 (scFOS)
Chicory fructooligosaccharides (lcFOS)
Mannotriose (bMnOS)
Xylotriose (XOS)
Arabinotriose (aAOS)
Lacto−N−tetraose (LNT)
Lacto−N−neotetraose (LNnT)
2'−fucosyllactose (2'FL)
3'−sialyllactose (SHMO)
6'−sialyllactose (SHMO)
Potato soluble starch (ST)
Pullulan (PUL)
Gum arabic from acacia tree (GA)
Konjac glucomannan (bMAN)
Wheat flour arabinoxylan (AX)
Sugar beet arabinan (AR)
Tamarind xyloglucan (XGL)
OD600
0.0
0.5
1.0
1.5
2.0
Time (h)
0
10
20
30
40

##### 11.19 *B. longum* subsp. *infantis* Bg064.S07_13.C6

Growth curves of *B. longum* subsp. *infantis* Bg064.S07_13.C6 in MRS-AC supplemented with various (n = 47) carbon sources.

```
growth_curves("Bifidobacterium longum subsp. infantis Bg064.S07_13.C6",
              "data/growth/formatted/Bl_infantis_Bg064.S07_13.C6.txt",
              "results/growth/growth_curves/Bl_infantis_Bg064.S07_13.C6.pdf",
              40)
```

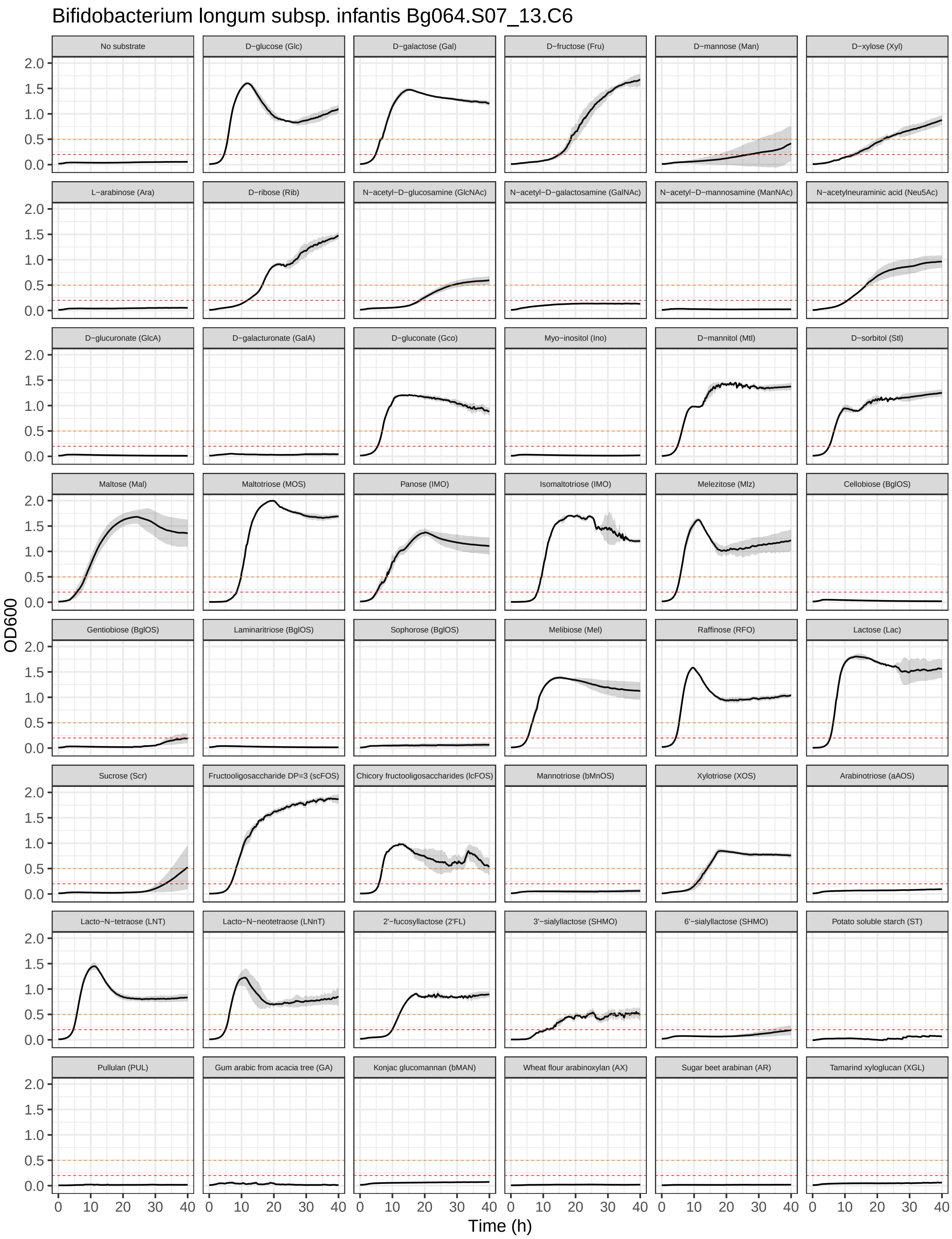
Bifidobacterium longum subsp. infantis Bg064.S07_13.C6
No substrate
D−glucose (Glc)
D−galactose (Gal)
D−fructose (Fru)
D−mannose (Man)
D−xylose (Xyl)
L−arabinose (Ara)
D−ribose (Rib)
N−acetyl−D−glucosamine (GlcNAc)
N−acetyl−D−galactosamine (GalNAc)
N−acetyl−D−mannosamine (ManNAc)
N−acetylneuraminic acid (Neu5Ac)
D−glucuronate (GlcA)
D−galacturonate (GalA)
D−gluconate (Gco)
Myo−inositol (Ino)
D−mannitol (Mtl)
D−sorbitol (Stl)
Maltose (Mal)
Maltotriose (MOS)
Panose (IMO)
Isomaltotriose (IMO)
Melezitose (Mlz)
Cellobiose (BglOS)
Gentiobiose (BglOS)
Laminaritriose (BglOS)
Sophorose (BglOS)
Melibiose (Mel)
Raffinose (RFO)
Lactose (Lac)
Sucrose (Scr)
Fructooligosaccharide DP=3 (scFOS)
Chicory fructooligosaccharides (lcFOS)
Mannotriose (bMnOS)
Xylotriose (XOS)
Arabinotriose (aAOS)
Lacto−N−tetraose (LNT)
Lacto−N−neotetraose (LNnT)
2'−fucosyllactose (2'FL)
3'−sialyllactose (SHMO)
6'−sialyllactose (SHMO)
Potato soluble starch (ST)
Pullulan (PUL)
Gum arabic from acacia tree (GA)
Konjac glucomannan (bMAN)
Wheat flour arabinoxylan (AX)
Sugar beet arabinan (AR)
Tamarind xyloglucan (XGL)
OD600
0.0
0.5
1.0
1.5
2.0
0
10
20
30
40
Time (h)

#### 11.20 *B. longum* subsp. *infantis* Bg40721_2D9

Growth curves of *B. longum* subsp. *infantis* Bg40721_2D9 in MRS-AC supplemented with various (n = 47) carbon sources.

```
growth_curves("Bifidobacterium longum subsp. infantis Bg40721_2D9",
              "data/growth/formatted/Bl_infantis_Bg40721_2D9.txt",
              "results/growth/growth_curves/Bl_infantis_Bg40721_2D9.pdf",
              40)
```

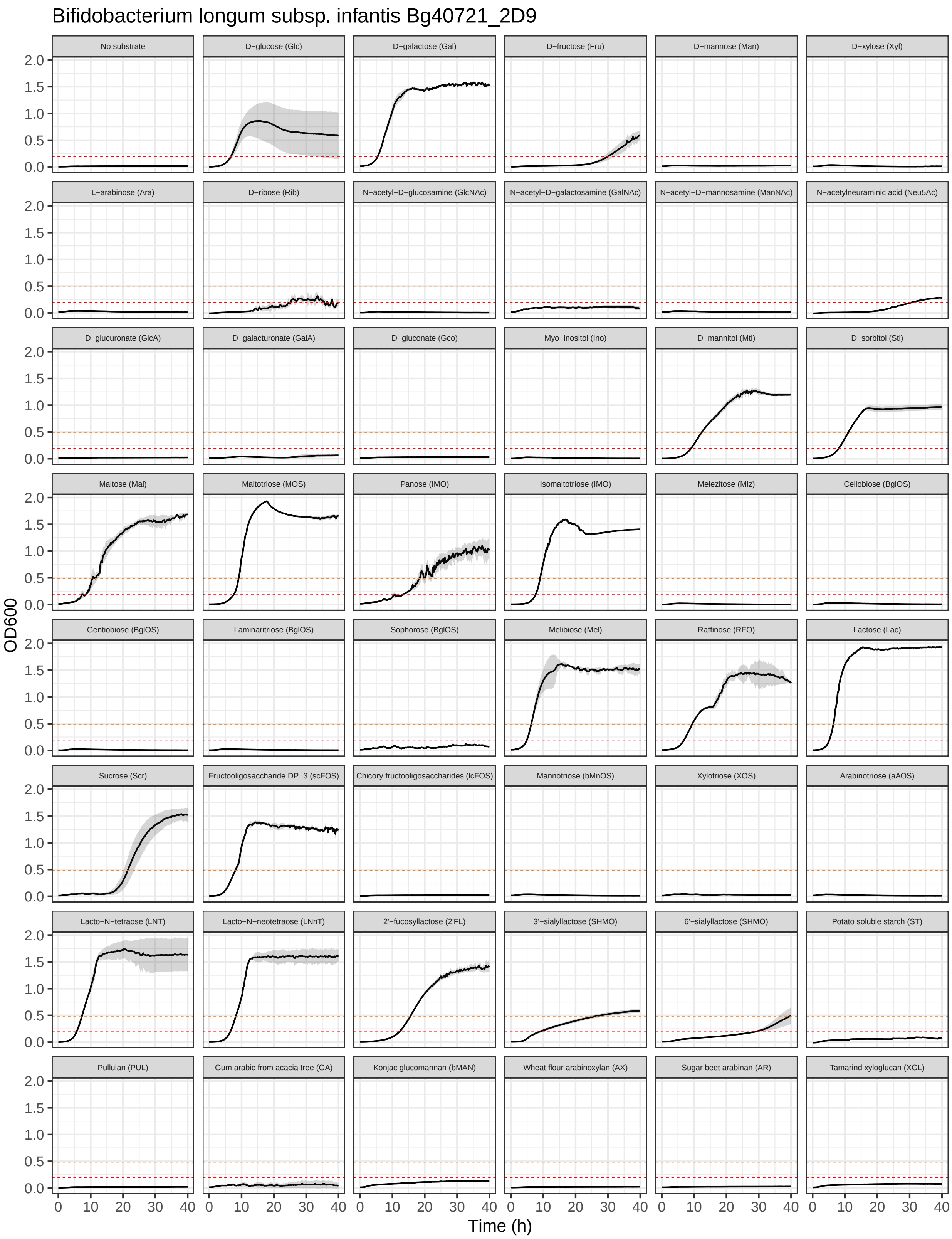
Bifidobacterium longum subsp. infantis Bg40721_2D9
No substrate
D−glucose (Glc)
D−galactose (Gal)
D−fructose (Fru)
D−mannose (Man)
D−xylose (Xyl)
L−arabinose (Ara)
D−ribose (Rib)
N−acetyl−D−glucosamine (GlcNAc)
N−acetyl−D−galactosamine (GalNAc)
N−acetyl−D−mannosamine (ManNAc)
N−acetylneuraminic acid (Neu5Ac)
D−glucuronate (GlcA)
D−galacturonate (GalA)
D−gluconate (Gco)
Myo−inositol (Ino)
D−mannitol (Mtl)
D−sorbitol (Stl)
Maltose (Mal)
Maltotriose (MOS)
Panose (IMO)
Isomaltotriose (IMO)
Melezitose (Mlz)
Cellobiose (BglOS)
Gentiobiose (BglOS)
Laminaritriose (BglOS)
Sophorose (BglOS)
Melibiose (Mel)
Raffinose (RFO)
Lactose (Lac)
Sucrose (Scr)
Fructooligosaccharide DP=3 (scFOS)
Chicory fructooligosaccharides (lcFOS)
Mannotriose (bMnOS)
Xylotriose (XOS)
Arabinotriose (aAOS)
Lacto−N−tetraose (LNT)
Lacto−N−neotetraose (LNnT)
2'−fucosyllactose (2'FL)
3'−sialyllactose (SHMO)
6'−sialyllactose (SHMO)
Potato soluble starch (ST)
Pullulan (PUL)
Gum arabic from acacia tree (GA)
Konjac glucomannan (bMAN)
Wheat flour arabinoxylan (AX)
Sugar beet arabinan (AR)
Tamarind xyloglucan (XGL)
OD600
Time (h)
0.0
0.5
1.0
1.5
2.0
0
10
20
30
40

#### 11.21 *B. longum* subsp. *infantis* JG_Bg463

Growth curves of *B. longum* subsp. *infantis* JG_Bg463 in MRS-AC supplemented with various (n = 47) carbon sources.

```
growth_curves("Bifidobacterium longum subsp. infantis JG_Bg463",
              "data/growth/formatted/Bl_infantis_JG_Bg463.txt",
              "results/growth/growth_curves/Bl_infantis_JG_Bg463.pdf",
              40)
```

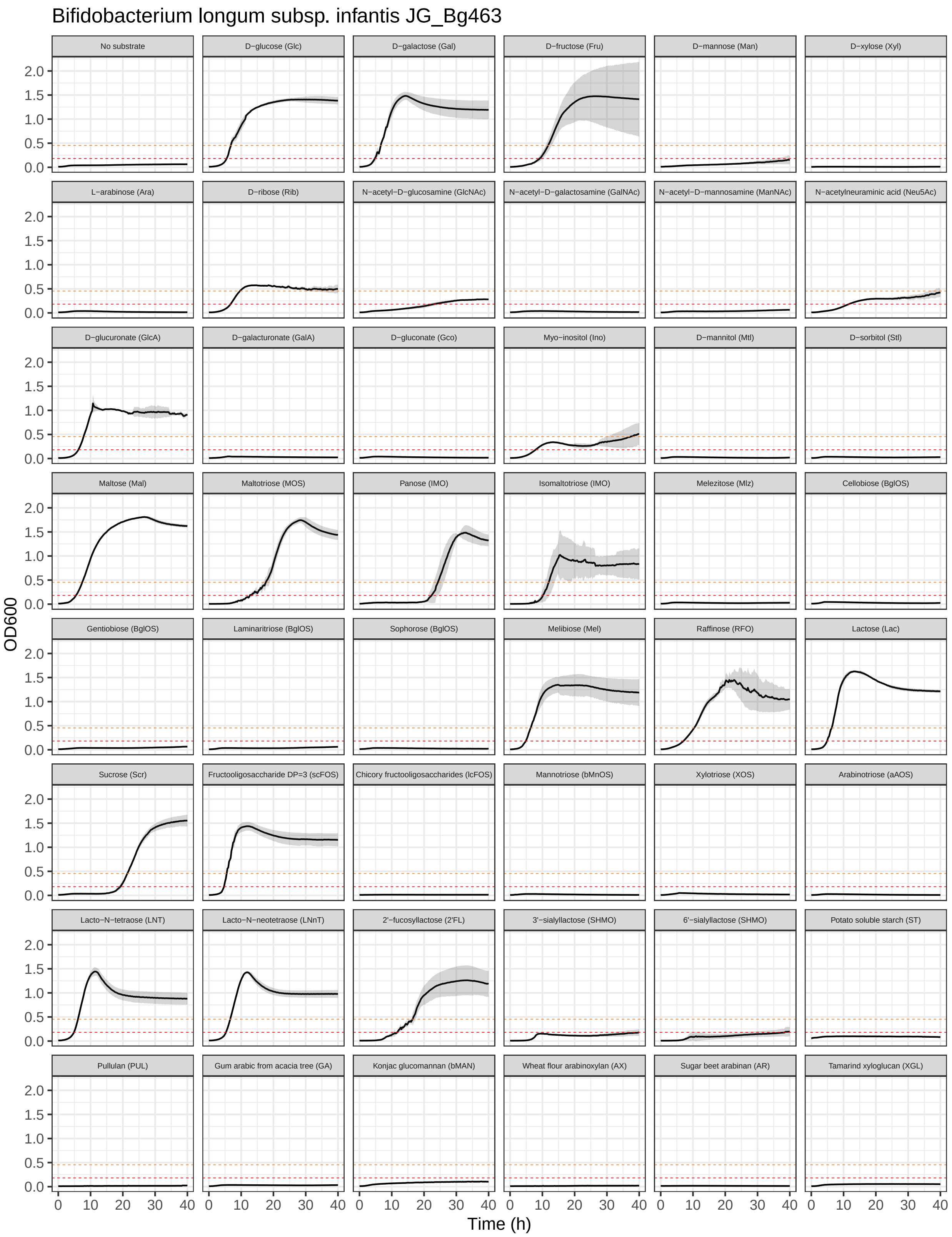
Bifidobacterium longum subsp. infantis JG_Bg463
No substrate
D−glucose (Glc)
D−galactose (Gal)
D−fructose (Fru)
D−mannose (Man)
D−xylose (Xyl)
L−arabinose (Ara)
D−ribose (Rib)
N−acetyl−D−glucosamine (GlcNAc)
N−acetyl−D−galactosamine (GalNAc)
N−acetyl−D−mannosamine (ManNAc)
N−acetylneuraminic acid (Neu5Ac)
D−glucuronate (GlcA)
D−galacturonate (GalA)
D−gluconate (Gco)
Myo−inositol (Ino)
D−mannitol (Mtl)
D−sorbitol (Stl)
Maltose (Mal)
Maltotriose (MOS)
Panose (IMO)
Isomaltotriose (IMO)
Melezitose (Mlz)
Cellobiose (BglOS)
Gentiobiose (BglOS)
Laminaritriose (BglOS)
Sophorose (BglOS)
Melibiose (Mel)
Raffinose (RFO)
Lactose (Lac)
Sucrose (Scr)
Fructooligosaccharide DP=3 (scFOS)
Chicory fructooligosaccharides (lcFOS)
Mannotriose (bMnOS)
Xylotriose (XOS)
Arabinotriose (aAOS)
Lacto−N−tetraose (LNT)
Lacto−N−neotetraose (LNnT)
2'−fucosyllactose (2'FL)
3'−sialyllactose (SHMO)
6'−sialyllactose (SHMO)
Potato soluble starch (ST)
Pullulan (PUL)
Gum arabic from acacia tree (GA)
Konjac glucomannan (bMAN)
Wheat flour arabinoxylan (AX)
Sugar beet arabinan (AR)
Tamarind xyloglucan (XGL)
OD600
0.0
0.5
1.0
1.5
2.0
0
10
20
30
40
Time (h)

#### 11.22 *B. longum* subsp. *infantis* Malawi264A_MC2

Growth curves of *B. longum* subsp. *infantis* Malawi264A_MC2 in MRS-AC supplemented with various (n = 47) carbon sources.

```
growth_curves("Bifidobacterium longum subsp. infantis Malawi264A_MC2",
              "data/growth/formatted/Bl_infantis_Malawi264A_MC2.txt",
              "results/growth/growth_curves/Bl_infantis_Malawi264A_MC2.pdf",
              40)
```

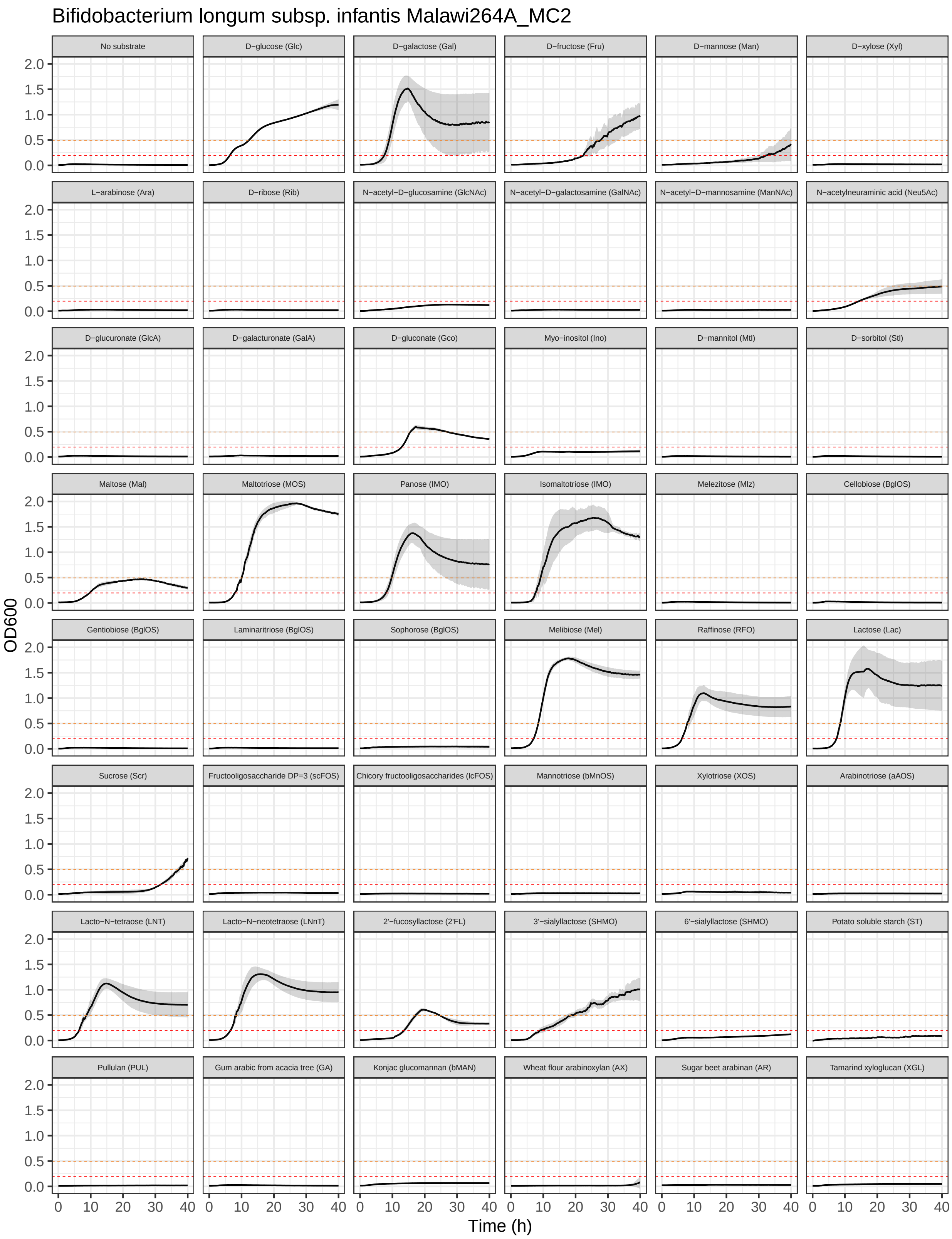

Bifidobacterium longum subsp. infantis Malawi264A_MC2
No substrate
D−glucose (Glc)
D−galactose (Gal)
D−fructose (Fru)
D−mannose (Man)
D−xylose (Xyl)
L−arabinose (Ara)
D−ribose (Rib)
N−acetyl−D−glucosamine (GlcNAc)
N−acetyl−D−galactosamine (GalNAc)
N−acetyl−D−mannosamine (ManNAc)
N−acetylneuraminic acid (Neu5Ac)
D−glucuronate (GlcA)
D−galacturonate (GalA)
D−gluconate (Gco)
Myo−inositol (Ino)
D−mannitol (Mtl)
D−sorbitol (Stl)
Maltose (Mal)
Maltotriose (MOS)
Panose (IMO)
Isomaltotriose (IMO)
Melezitose (Mlz)
Cellobiose (BglOS)
Gentiobiose (BglOS)
Laminaritriose (BglOS)
Sophorose (BglOS)
Melibiose (Mel)
Raffinose (RFO)
Lactose (Lac)
Sucrose (Scr)
Fructooligosaccharide DP=3 (scFOS)
Chicory fructooligosaccharides (lcFOS)
Mannotriose (bMnOS)
Xylotriose (XOS)
Arabinotriose (aAOS)
Lacto−N−tetraose (LNT)
Lacto−N−neotetraose (LNnT)
2'−fucosyllactose (2'FL)
3'−sialyllactose (SHMO)
6'−sialyllactose (SHMO)
Potato soluble starch (ST)
Pullulan (PUL)
Gum arabic from acacia tree (GA)
Konjac glucomannan (bMAN)
Wheat flour arabinoxylan (AX)
Sugar beet arabinan (AR)
Tamarind xyloglucan (XGL)
OD600
0.0
0.5
1.0
1.5
2.0
0
10
20
30
40
Time (h)

##### 11.23 *B. longum* subsp. *longum* Bg115.S08_3A11

Growth curves of *B. longum* subsp. *longum* Bg115.S08_3A11 in MRS-AC supplemented with various (n = 47) carbon sources.

```
growth_curves("Bifidobacterium longum subsp. longum Bg115.S08_3A11",
              "data/growth/formatted/Bl_longum_Bg115.S08_3A11.txt",
              "results/growth/growth_curves/Bl_longum_Bg115.S08_3A11.pdf",
              40)
```

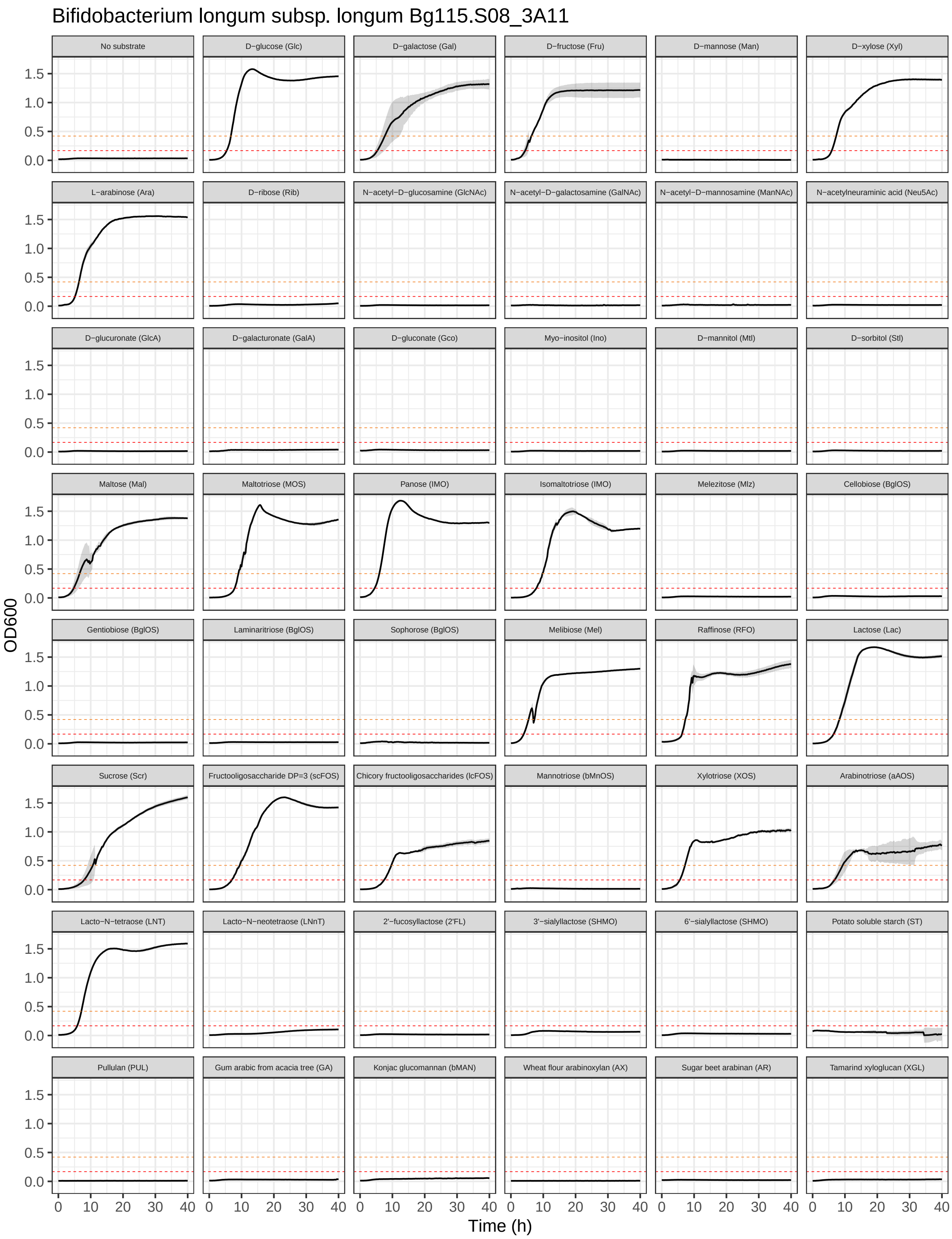
Bifidobacterium longum subsp. longum Bg115.S08_3A11
No substrate
D−glucose (Glc)
D−galactose (Gal)
D−fructose (Fru)
D−mannose (Man)
D−xylose (Xyl)
L−arabinose (Ara)
D−ribose (Rib)
N−acetyl−D−glucosamine (GlcNAc)
N−acetyl−D−galactosamine (GalNAc)
N−acetyl−D−mannosamine (ManNAc)
N−acetylneuraminic acid (Neu5Ac)
D−glucuronate (GlcA)
D−galacturonate (GalA)
D−gluconate (Gco)
Myo−inositol (Ino)
D−mannitol (Mtl)
D−sorbitol (Stl)
Maltose (Mal)
Maltotriose (MOS)
Panose (IMO)
Isomaltotriose (IMO)
Melezitose (Mlz)
Cellobiose (BglOS)
Gentiobiose (BglOS)
Laminaritriose (BglOS)
Sophorose (BglOS)
Melibiose (Mel)
Raffinose (RFO)
Lactose (Lac)
Sucrose (Scr)
Fructooligosaccharide DP=3 (scFOS)
Chicory fructooligosaccharides (lcFOS)
Mannotriose (bMnOS)
Xylotriose (XOS)
Arabinotriose (aAOS)
Lacto−N−tetraose (LNT)
Lacto−N−neotetraose (LNnT)
2'−fucosyllactose (2'FL)
3'−sialyllactose (SHMO)
6'−sialyllactose (SHMO)
Potato soluble starch (ST)
Pullulan (PUL)
Gum arabic from acacia tree (GA)
Konjac glucomannan (bMAN)
Wheat flour arabinoxylan (AX)
Sugar beet arabinan (AR)
Tamarind xyloglucan (XGL)
OD600
0.0
0.5
1.0
1.5
Time (h)
0
10
20
30
40

##### 11.24 *B. longum* subsp. *suis* Bg131.S11_17.F6

Growth curves of *B. longum* subsp. *suis* Bg131.S11_17.F6 in MRS-AC supplemented with various (n = 47) carbon sources.

```
growth_curves("Bifidobacterium longum subsp. suis Bg131.S11_17.F6",
              "data/growth/formatted/Bl_suis_Bg131.S11_17.F6.txt",
              "results/growth/growth_curves/Bl_suis_Bg131.S11_17.F6.pdf",
              40)
```

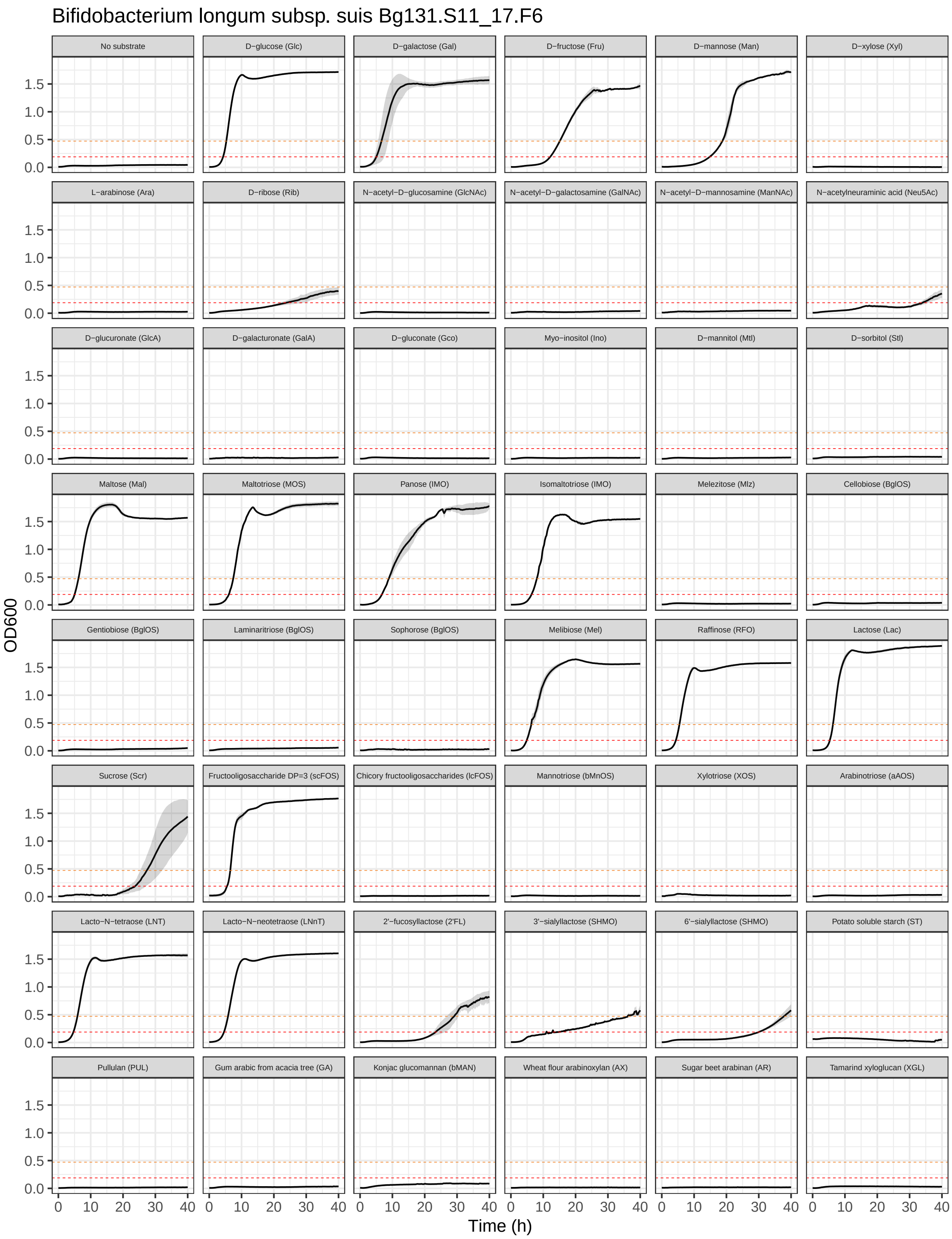

Bifidobacterium longum subsp. suis Bg131.S11_17.F6
No substrate
D−glucose (Glc)
D−galactose (Gal)
D−fructose (Fru)
D−mannose (Man)
D−xylose (Xyl)
L−arabinose (Ara)
D−ribose (Rib)
N−acetyl−D−glucosamine (GlcNAc)
N−acetyl−D−galactosamine (GalNAc)
N−acetyl−D−mannosamine (ManNAc)
N−acetylneuraminic acid (Neu5Ac)
D−glucuronate (GlcA)
D−galacturonate (GalA)
D−gluconate (Gco)
Myo−inositol (Ino)
D−mannitol (Mtl)
D−sorbitol (Stl)
Maltose (Mal)
Maltotriose (MOS)
Panose (IMO)
Isomaltotriose (IMO)
Melezitose (Mlz)
Cellobiose (BglOS)
Gentiobiose (BglOS)
Laminaritriose (BglOS)
Sophorose (BglOS)
Melibiose (Mel)
Raffinose (RFO)
Lactose (Lac)
Sucrose (Scr)
Fructooligosaccharide DP=3 (scFOS)
Chicory fructooligosaccharides (lcFOS)
Mannotriose (bMnOS)
Xylotriose (XOS)
Arabinotriose (aAOS)
Lacto−N−tetraose (LNT)
Lacto−N−neotetraose (LNnT)
2'−fucosyllactose (2'FL)
3'−sialyllactose (SHMO)
6'−sialyllactose (SHMO)
Potato soluble starch (ST)
Pullulan (PUL)
Gum arabic from acacia tree (GA)
Konjac glucomannan (bMAN)
Wheat flour arabinoxylan (AX)
Sugar beet arabinan (AR)
Tamarind xyloglucan (XGL)
OD600
0.0
0.5
1.0
1.5
0
10
20
30
40
Time (h)

##### 11.25 ***B. longum* subsp. *suis* Bg41121_2E1**

Growth curves of *B. longum* subsp. *suis* Bg41121_2E1 in MRS-AC supplemented with various (n = 44) carbon sources.

```
growth_curves("Bifidobacterium longum subsp. suis Bg41121_2E1",
              "data/growth/formatted/Bl_suis_Bg41121_2E1.txt",
              "results/growth/growth_curves/Bl_suis_Bg41121_2E1.pdf",
              40)
```

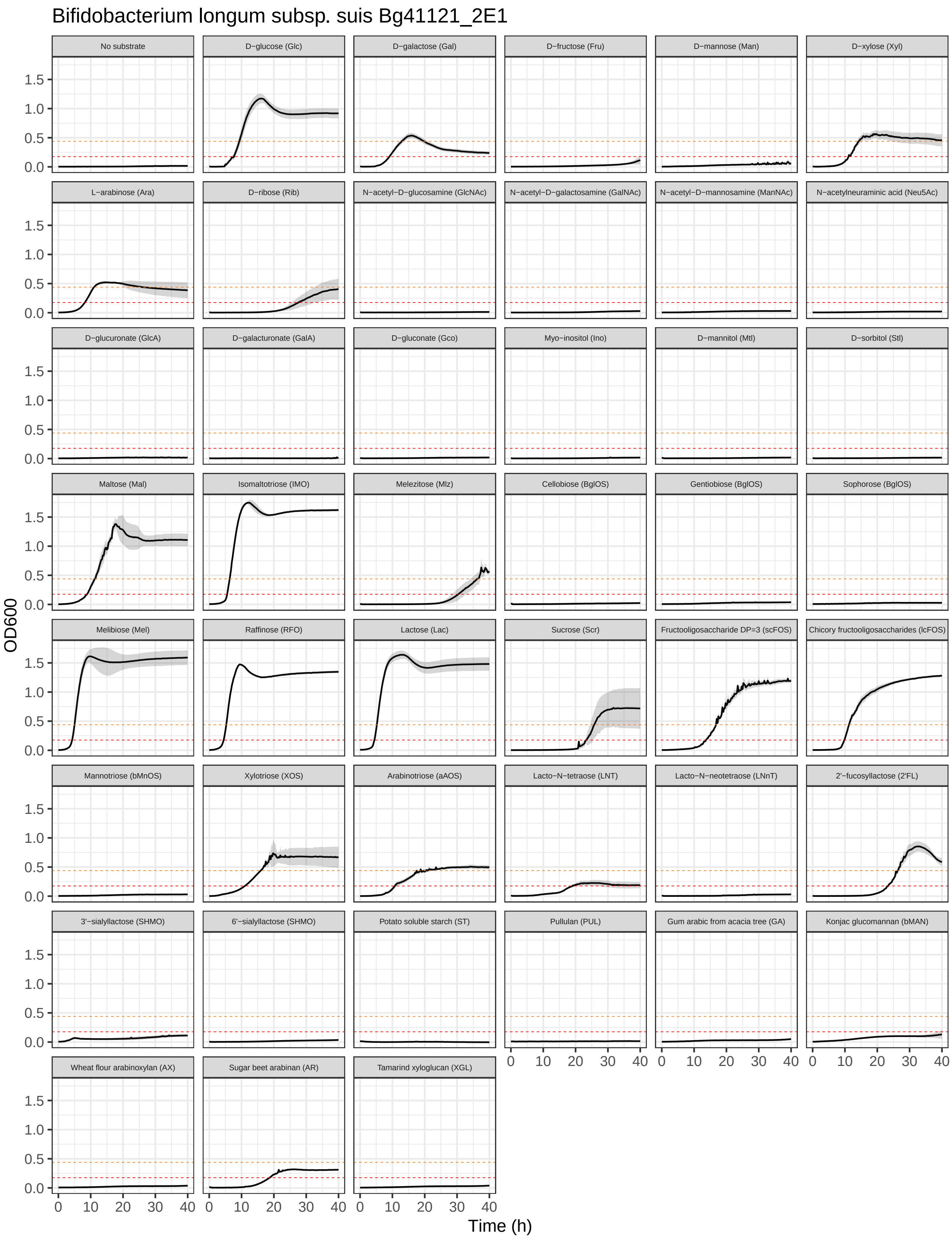
Bifidobacterium longum subsp. suis Bg41121_2E1
No substrate
D−glucose (Glc)
D−galactose (Gal)
D−fructose (Fru)
D−mannose (Man)
D−xylose (Xyl)
L−arabinose (Ara)
D−ribose (Rib)
N−acetyl−D−glucosamine (GlcNAc)
N−acetyl−D−galactosamine (GalNAc)
N−acetyl−D−mannosamine (ManNAc)
N−acetylneuraminic acid (Neu5Ac)
D−glucuronate (GlcA)
D−galacturonate (GalA)
D−gluconate (Gco)
Myo−inositol (Ino)
D−mannitol (Mtl)
D−sorbitol (Stl)
Maltose (Mal)
Isomaltotriose (IMO)
Melezitose (Mlz)
Cellobiose (BglOS)
Gentiobiose (BglOS)
Sophorose (BglOS)
Melibiose (Mel)
Raffinose (RFO)
Lactose (Lac)
Sucrose (Scr)
Fructooligosaccharide DP=3 (scFOS)
Chicory fructooligosaccharides (lcFOS)
Mannotriose (bMnOS)
Xylotriose (XOS)
Arabinotriose (aAOS)
Lacto−N−tetraose (LNT)
Lacto−N−neotetraose (LNnT)
2'−fucosyllactose (2'FL)
3'−sialyllactose (SHMO)
6'−sialyllactose (SHMO)
Potato soluble starch (ST)
Pullulan (PUL)
Gum arabic from acacia tree (GA)
Konjac glucomannan (bMAN)
Wheat flour arabinoxylan (AX)
Sugar beet arabinan (AR)
Tamarind xyloglucan (XGL)
OD600
0.0
0.5
1.0
1.5
0
10
20
30
40
Time (h)

#### 11.26 *B. longum* subsp. *suis* M257B_A6

Growth curves of *B. longum* subsp. *suis* M257B_A6 in MRS-AC supplemented with various (n = 45) carbon sources.

```
growth_curves("Bifidobacterium longum subsp. suis M257B_A6",
              "data/growth/formatted/Bl_suis_M257B_A6.txt",
              "results/growth/growth_curves/Bl_suis_M257B_A6.pdf",
              40)
```

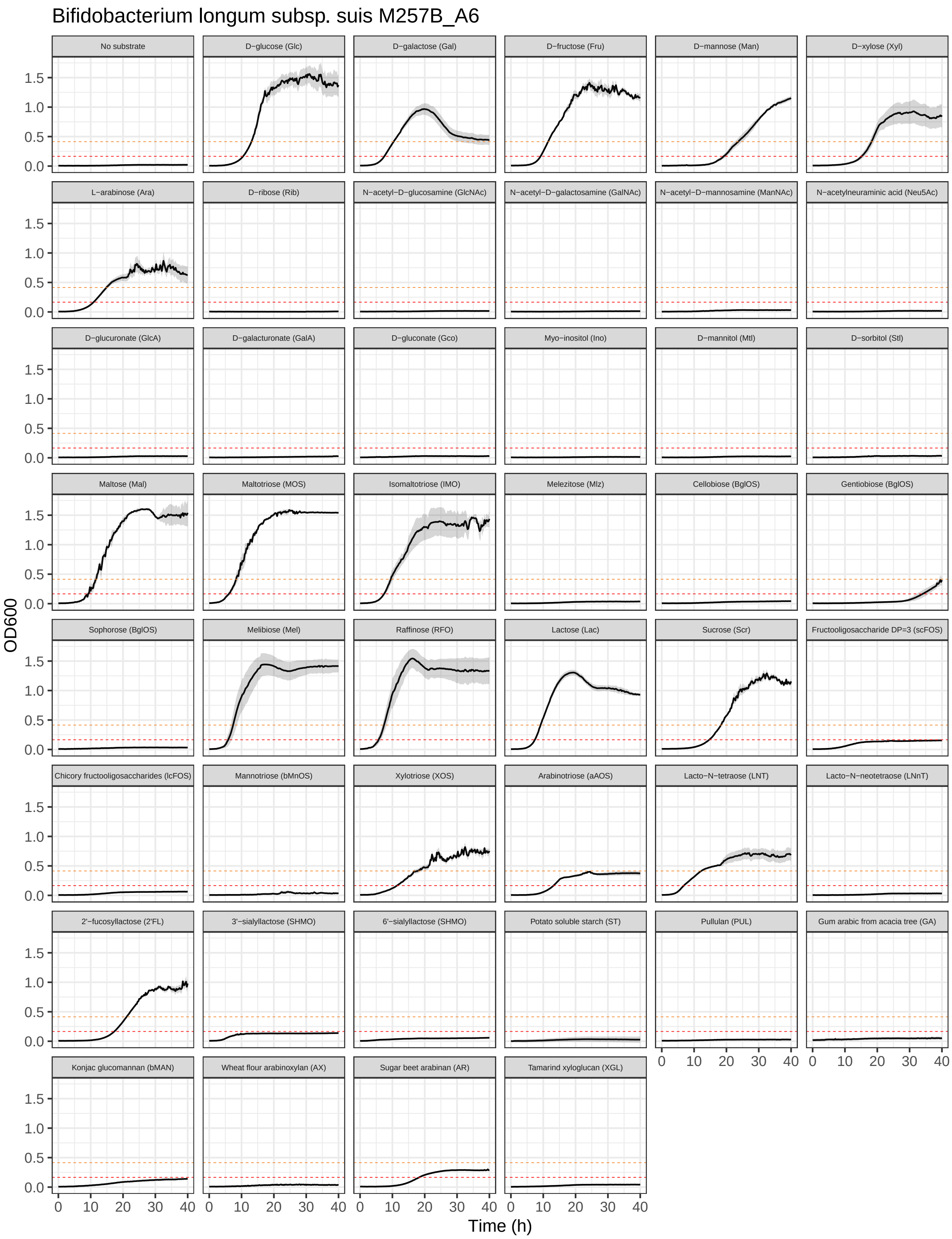

Bifidobacterium longum subsp. suis M257B_A6
No substrate
D−glucose (Glc)
D−galactose (Gal)
D−fructose (Fru)
D−mannose (Man)
D−xylose (Xyl)
L−arabinose (Ara)
D−ribose (Rib)
N−acetyl−D−glucosamine (GlcNAc)
N−acetyl−D−galactosamine (GalNAc)
N−acetyl−D−mannosamine (ManNAc)
N−acetylneuraminic acid (Neu5Ac)
D−glucuronate (GlcA)
D−galacturonate (GalA)
D−gluconate (Gco)
Myo−inositol (Ino)
D−mannitol (Mtl)
D−sorbitol (Stl)
Maltose (Mal)
Maltotriose (MOS)
Isomaltotriose (IMO)
Melezitose (Mlz)
Cellobiose (BglOS)
Gentiobiose (BglOS)
Sophorose (BglOS)
Melibiose (Mel)
Raffinose (RFO)
Lactose (Lac)
Sucrose (Scr)
Fructooligosaccharide DP=3 (scFOS)
Chicory fructooligosaccharides (lcFOS)
Mannotriose (bMnOS)
Xylotriose (XOS)
Arabinotriose (aAOS)
Lacto−N−tetraose (LNT)
Lacto−N−neotetraose (LNnT)
2'−fucosyllactose (2'FL)
3'−sialyllactose (SHMO)
6'−sialyllactose (SHMO)
Potato soluble starch (ST)
Pullulan (PUL)
Gum arabic from acacia tree (GA)
Konjac glucomannan (bMAN)
Wheat flour arabinoxylan (AX)
Sugar beet arabinan (AR)
Tamarind xyloglucan (XGL)
OD600
Time (h)
0.0
0.5
1.0
1.5
0
10
20
30
40

#### 11.27 *B. longum* LFYP82

Growth curves of *B. longum* LFYP82 in MRS-AC supplemented with various (n = 45) carbon sources.

```
growth_curves("Bifidobacterium longum LFYP82",
              "data/growth/formatted/Blongum_LFYP82.txt",
              "results/growth/growth_curves/Blongum_LFYP82.pdf",
              40)
```

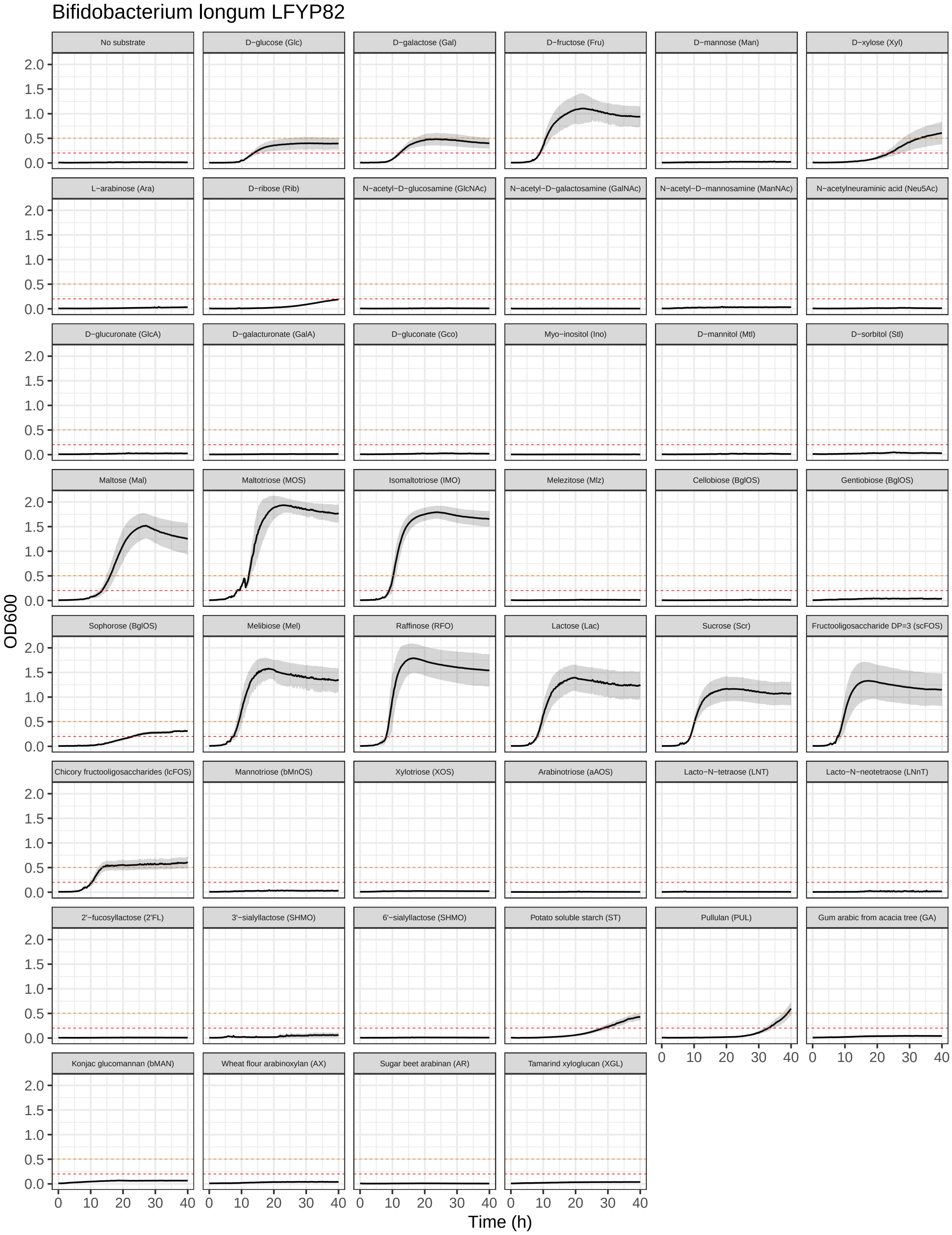

Bifidobacterium longum LFYP82
No substrate
D−glucose (Glc)
D−galactose (Gal)
D−fructose (Fru)
D−mannose (Man)
D−xylose (Xyl)
L−arabinose (Ara)
D−ribose (Rib)
N−acetyl−D−glucosamine (GlcNAc)
N−acetyl−D−galactosamine (GalNAc)
N−acetyl−D−mannosamine (ManNAc)
N−acetylneuraminic acid (Neu5Ac)
D−glucuronate (GlcA)
D−galacturonate (GalA)
D−gluconate (Gco)
Myo−inositol (Ino)
D−mannitol (Mtl)
D−sorbitol (Stl)
Maltose (Mal)
Maltotriose (MOS)
Isomaltotriose (IMO)
Melezitose (Mlz)
Cellobiose (BglOS)
Gentiobiose (BglOS)
Sophorose (BglOS)
Melibiose (Mel)
Raffinose (RFO)
Lactose (Lac)
Sucrose (Scr)
Fructooligosaccharide DP=3 (scFOS)
Chicory fructooligosaccharides (lcFOS)
Mannotriose (bMnOS)
Xylotriose (XOS)
Arabinotriose (aAOS)
Lacto−N−tetraose (LNT)
Lacto−N−neotetraose (LNnT)
2'−fucosyllactose (2'FL)
3'−sialyllactose (SHMO)
6'−sialyllactose (SHMO)
Potato soluble starch (ST)
Pullulan (PUL)
Gum arabic from acacia tree (GA)
Konjac glucomannan (bMAN)
Wheat flour arabinoxylan (AX)
Sugar beet arabinan (AR)
Tamarind xyloglucan (XGL)
OD600
Time (h)

##### 11.28 *B. pseudocatenulatum* LFYP29

Growth curves of *B. pseudocatenulatum* LFYP29 in MRS-AC supplemented with various (n = 47) carbon sources.

```
growth_curves("Bifidobacterium pseudocatenulatum LFYP29",
              "data/growth/formatted/Bpseudocatenulatum_LFYP29.txt",
              "results/growth/growth_curves/Bpseudocatenulatum_LFYP29.pdf",
              40)
```

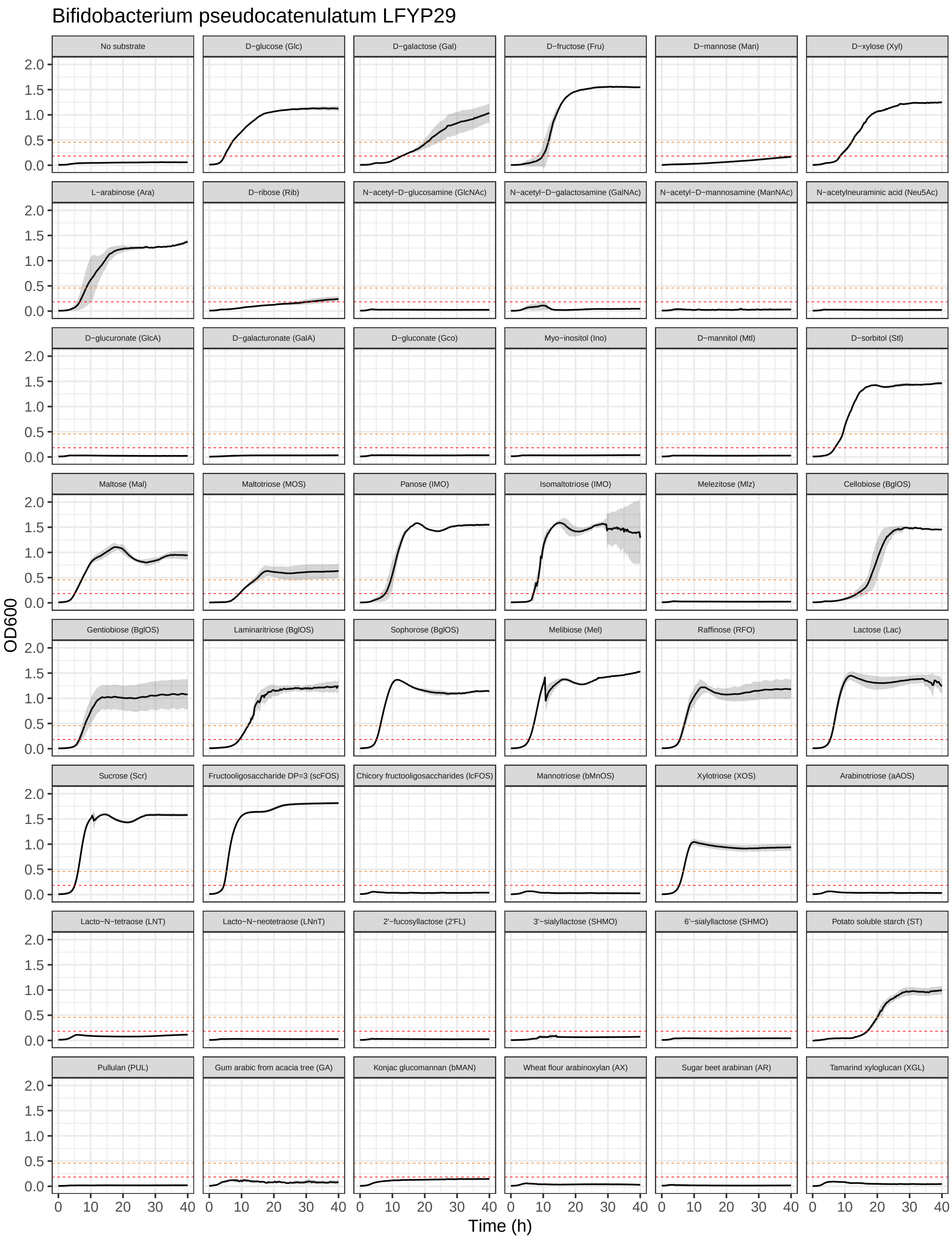
Bifidobacterium pseudocatenulatum LFYP29
No substrate
D−glucose (Glc)
D−galactose (Gal)
D−fructose (Fru)
D−mannose (Man)
D−xylose (Xyl)
L−arabinose (Ara)
D−ribose (Rib)
N−acetyl−D−glucosamine (GlcNAc)
N−acetyl−D−galactosamine (GalNAc)
N−acetyl−D−mannosamine (ManNAc)
N−acetylneuraminic acid (Neu5Ac)
D−glucuronate (GlcA)
D−galacturonate (GalA)
D−gluconate (Gco)
Myo−inositol (Ino)
D−mannitol (Mtl)
D−sorbitol (Stl)
Maltose (Mal)
Maltotriose (MOS)
Panose (IMO)
Isomaltotriose (IMO)
Melezitose (Mlz)
Cellobiose (BglOS)
Gentiobiose (BglOS)
Laminaritriose (BglOS)
Sophorose (BglOS)
Melibiose (Mel)
Raffinose (RFO)
Lactose (Lac)
Sucrose (Scr)
Fructooligosaccharide DP=3 (scFOS)
Chicory fructooligosaccharides (lcFOS)
Mannotriose (bMnOS)
Xylotriose (XOS)
Arabinotriose (aAOS)
Lacto−N−tetraose (LNT)
Lacto−N−neotetraose (LNnT)
2'−fucosyllactose (2'FL)
3'−sialyllactose (SHMO)
6'−sialyllactose (SHMO)
Potato soluble starch (ST)
Pullulan (PUL)
Gum arabic from acacia tree (GA)
Konjac glucomannan (bMAN)
Wheat flour arabinoxylan (AX)
Sugar beet arabinan (AR)
Tamarind xyloglucan (XGL)
OD600
0.0
0.5
1.0
1.5
2.0
0
10
20
30
40
Time (h)

##### 11.29 *B. pseudocatenulatum* M26A__2F1

Growth curves of *B. pseudocatenulatum* M26A__2F1 in MRS-AC supplemented with various (n = 44) carbon sources.

Unlike all other tested strains, *B. pseudocatenulatum* M26A__2F1 exhibited weak growth in MRS-AC without added carbon source (OD600 in range 0.2-0.25). To account for that, we increased the growth/no growth OD600 threshold for this strain.

```
# read files
strain_name <- "Bifidobacterium pseudocatenulatum M26A_2F1"
growth_data <- "data/growth/formatted/Bpseudocatenulatum_M26A_2F1.txt"
growth_output_file <- "results/growth/growth_curves/Bpseudocatenulatum_M26A_2F1.pdf"
trim_at_time <- 40 # measurements until [input] hours
growth_annotation <- "data/growth/formatted/plate_layout.txt"
blank_wells <- c("blank_1","blank_2","blank_3") # specify blank wells
blank_wells_ST <- c("blank_ST1","blank_ST2","blank_ST3") # specify blank wells for MRS-AC-ST
# read the file with measurements
d <- read_tsv(growth_data) %>%
  filter(time <= trim_at_time)
# calculate mean blank measurements (MRS-AC-Lac without added cells)
blank_d <- d %>%
  dplyr::select(all_of(blank_wells)) %>%
  rowMeans()
# calculate mean blank measurements for ST blank (MRS-AC-ST without added cells)
blank_ST <- d %>%
  dplyr::select(all_of(blank_wells_ST)) %>%
  rowMeans()
# subtract blank measurements and round OD600 values to 3 digits
d_bl <- d %>%
  mutate(across(-c(time, blank_ST1, blank_ST2, blank_ST3, ST1, ST2, ST3), ~ .x - blank_d)) %>%
  mutate(across(c(blank_ST1, blank_ST2, blank_ST3, ST1, ST2, ST3), ~ .x - blank_ST)) %>%
  mutate(across(2:last_col(), ~ round(.x, 3)))

# read the file with plate annotation
ann <- read_tsv(growth_annotation)
d_bl_long <- d_bl %>%
  gather(., well, od, -time)
# create annotated table
d_bl_long_ann <- left_join(d_bl_long, ann, by="well") %>%
  filter(carbon_source != "blank") %>%
  filter(carbon_source != "blank_ST") %>%
  mutate(across(carbon_source, factor, levels = c("No substrate",
                                                  "D-glucose (Glc)",
                                                  "D-galactose (Gal)",
                                                  "D-fructose (Fru)",
                                                  "D-mannose (Man)",
                                                  "D-xylose (Xyl)",
                                                  "L-arabinose (Ara)",
                                                  "D-ribose (Rib)",
                                                  "N-acetyl-D-glucosamine (GlcNAc)",
                                                  "N-acetyl-D-galactosamine (GalNAc)",
                                                  "N-acetyl-D-mannosamine (ManNAc)",
                                                  "N-acetylneuraminic acid (Neu5Ac)",
                                                  "D-glucuronate (GlcA)",
```

```
                                                    "D-galacturonate (GalA)",
                                                    "D-gluconate (Gco)",
                                                    "Myo-inositol (Ino)",
                                                    "D-mannitol (Mtl)",
                                                    "D-sorbitol (Stl)",
                                                    "Maltose (Mal)",
                                                    "Maltotriose (MOS)",
                                                    "Panose (IMO)",
                                                    "Isomaltotriose (IMO)",
                                                    "Melezitose (Mlz)",
                                                    "Cellobiose (BglOS)",
                                                    "Gentiobiose (BglOS)",
                                                    "Laminaritriose (BglOS)",
                                                    "Sophorose (BglOS)",
                                                    "Melibiose (Mel)",
                                                    "Raffinose (RFO)",
                                                    "Lactose (Lac)",
                                                    "Sucrose (Scr)",
                                                    "Fructooligosaccharide DP=3 (scFOS)",
                                                    "Chicory fructooligosaccharides (lcFOS)",
                                                    "Mannotriose (bMnOS)",
                                                    "Xylotriose (XOS)",
                                                    "Arabinotriose (aAOS)",
                                                    "Lacto-N-tetraose (LNT)",
                                                    "Lacto-N-neotetraose (LNnT)",
                                                    "2'-fucosyllactose (2'FL)",
                                                    "3'-sialyllactose (SHMO)",
                                                    "6'-sialyllactose (SHMO)",
                                                    "Potato soluble starch (ST)",
                                                    "Pullulan (PUL)",
                                                    "Gum arabic from acacia tree (GA)",
                                                    "Konjac glucomannan (bMAN)",
                                                    "Wheat flour arabinoxylan (AX)",
                                                    "Sugar beet arabinan (AR)",
                                                    "Tamarind xyloglucan (XGL)")))

# calculate the mean OD600_max value and set thresholds based on this value
mean_max_OD <- mean(head(sort(d_bl_long_ann$od,decreasing=TRUE), n=30))
# threshold for no growth (10% of OD600_max)
growth_limit <- 0.22 * mean_max_OD
# threshold for weak growth (25% of OD600_max)
weak_growth_limit <- 0.55 * mean_max_OD

# calculate mean OD and standard deviation for each carbon source and time
growth_summary <- d_bl_long_ann %>%
  group_by(carbon_source, time) %>%
  summarise(
    mean_od = mean(od),
    sd_od = sd(od),
    n = n()
  )
# calculate lower and upper confidence intervals
confidence_level <- 0.95  # set the desired confidence level (e.g., 95%)
```

```r
z_value <- qnorm((1 + confidence_level) / 2)  # calculate the z-value based on the confidence level
growth_summary <- growth_summary %>%
  mutate(
    lower_ci = mean_od - z_value * (sd_od / sqrt(n)),
    upper_ci = mean_od + z_value * (sd_od / sqrt(n))
  )

# plot growth curves
filtered_summary <- growth_summary[!is.na(growth_summary$mean_od), ]
curves <- ggplot(filtered_summary, aes(x = time, y = mean_od)) +
  # add line
  geom_line() +
  # add confidence interval
  geom_ribbon(aes(ymin = lower_ci, ymax = upper_ci), alpha = 0.2) +
  scale_x_continuous(limits=c(0, trim_at_time),
                     breaks=c(0, trim_at_time*0.25, trim_at_time*0.5, trim_at_time*0.75, trim_at_time))
  # add threshold for no growth
  geom_hline(yintercept=growth_limit , linetype="dashed", color = "red", linewidth=0.2) +
  # add threshold for weak growth
  geom_hline(yintercept=weak_growth_limit, linetype="dashed", color = "#F48326", linewidth=0.2) +
  ggtitle(strain_name) +
  xlab("Time (h)") +
  ylab("OD600") +
  facet_wrap(~ carbon_source, scales="fixed", ncol = 6) +
  theme_bw() +
  theme(
    strip.text = element_text(size = 5)
  )
# save pdf
ggsave(filename = growth_output_file, plot = curves, width = 210, height = 297, units = "mm", dpi = 300
print(curves)
```

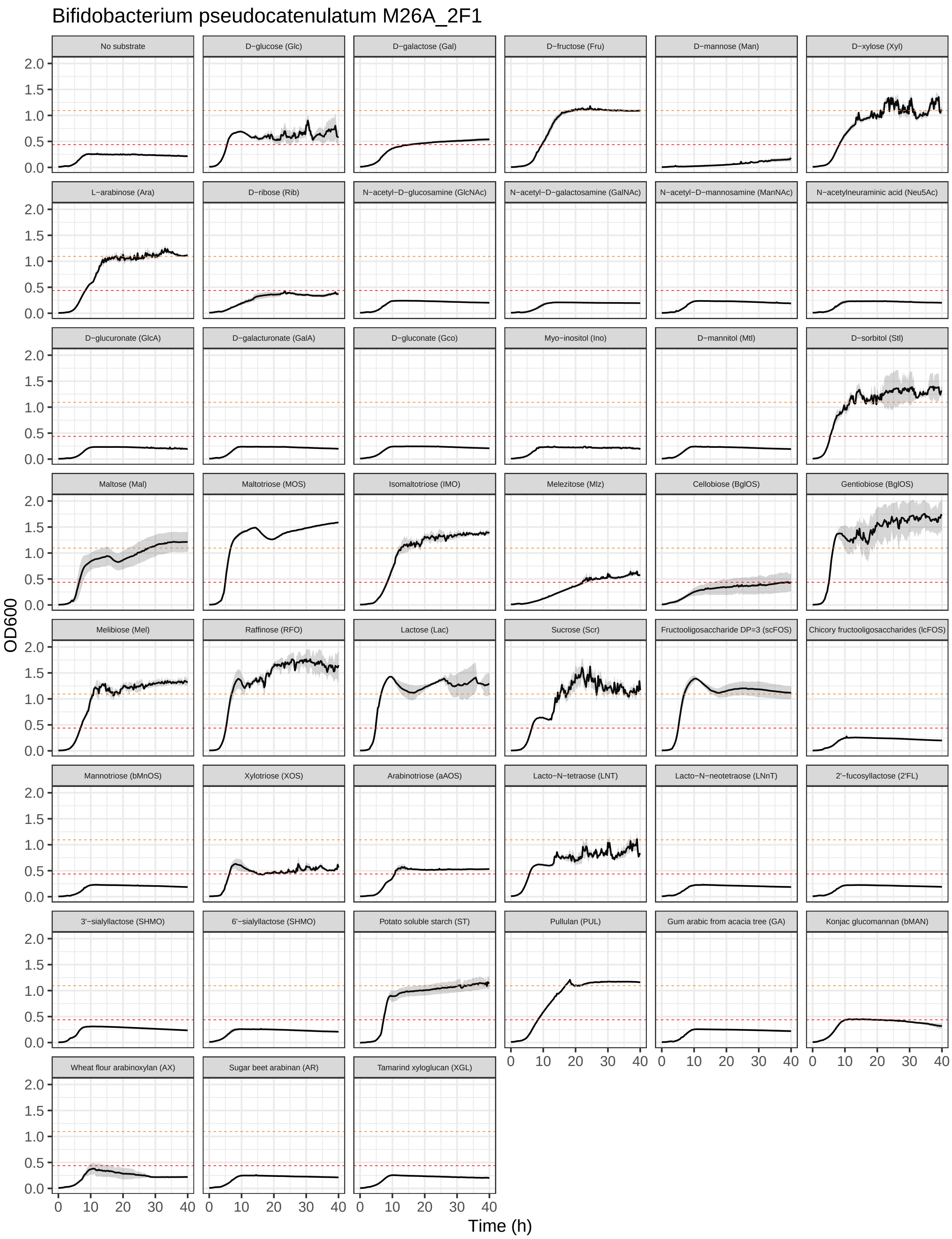
Bifidobacterium pseudocatenulatum M26A_2F1
No substrate
D−glucose (Glc)
D−galactose (Gal)
D−fructose (Fru)
D−mannose (Man)
D−xylose (Xyl)
L−arabinose (Ara)
D−ribose (Rib)
N−acetyl−D−glucosamine (GlcNAc)
N−acetyl−D−galactosamine (GalNAc)
N−acetyl−D−mannosamine (ManNAc)
N−acetylneuraminic acid (Neu5Ac)
D−glucuronate (GlcA)
D−galacturonate (GalA)
D−gluconate (Gco)
Myo−inositol (Ino)
D−mannitol (Mtl)
D−sorbitol (Stl)
Maltose (Mal)
Maltotriose (MOS)
Isomaltotriose (IMO)
Melezitose (Mlz)
Cellobiose (BglOS)
Gentiobiose (BglOS)
Melibiose (Mel)
Raffinose (RFO)
Lactose (Lac)
Sucrose (Scr)
Fructooligosaccharide DP=3 (scFOS)
Chicory fructooligosaccharides (lcFOS)
Mannotriose (bMnOS)
Xylotriose (XOS)
Arabinotriose (aAOS)
Lacto−N−tetraose (LNT)
Lacto−N−neotetraose (LNnT)
2'−fucosyllactose (2'FL)
3'−sialyllactose (SHMO)
6'−sialyllactose (SHMO)
Potato soluble starch (ST)
Pullulan (PUL)
Gum arabic from acacia tree (GA)
Konjac glucomannan (bMAN)
Wheat flour arabinoxylan (AX)
Sugar beet arabinan (AR)
Tamarind xyloglucan (XGL)
OD600
Time (h)
0.0
0.5
1.0
1.5
2.0
0
10
20
30
40

#### 11.30 *B. scardovii* JCM 12489 = DSM 13734

Growth curves of *B. scardovii* JCM 12489 = DSM 13734 in MRS-C supplemented with various (n = 43) carbon sources.

```
growth_curves("Bifidobacterium scardovii JCM 12489 = DSM 13734",
              "data/growth/formatted/Bscardovii_JCM12489.txt",
              "results/growth/growth_curves/Bscardovii_JCM12489.pdf",
              40)
```

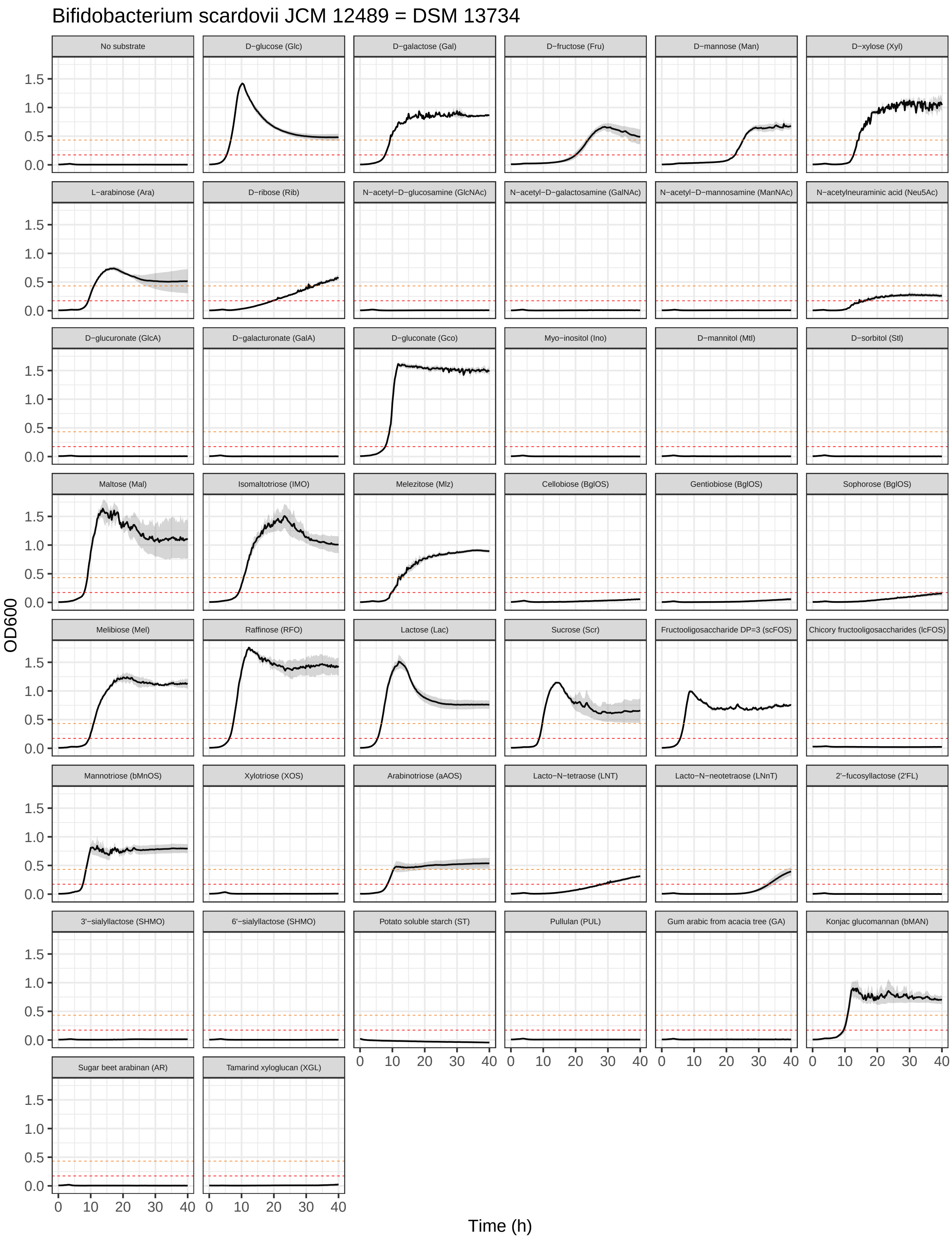
Bifidobacterium scardovii JCM 12489 = DSM 13734
No substrate
D−glucose (Glc)
D−galactose (Gal)
D−fructose (Fru)
D−mannose (Man)
D−xylose (Xyl)
L−arabinose (Ara)
D−ribose (Rib)
N−acetyl−D−glucosamine (GlcNAc)
N−acetyl−D−galactosamine (GalNAc)
N−acetyl−D−mannosamine (ManNAc)
N−acetylneuraminic acid (Neu5Ac)
D−glucuronate (GlcA)
D−galacturonate (GalA)
D−gluconate (Gco)
Myo−inositol (Ino)
D−mannitol (Mtl)
D−sorbitol (Stl)
Maltose (Mal)
Isomaltotriose (IMO)
Melezitose (Mlz)
Cellobiose (BglOS)
Gentiobiose (BglOS)
Sophorose (BglOS)
Melibiose (Mel)
Raffinose (RFO)
Lactose (Lac)
Sucrose (Scr)
Fructooligosaccharide DP=3 (scFOS)
Chicory fructooligosaccharides (lcFOS)
Mannotriose (bMnOS)
Xylotriose (XOS)
Arabinotriose (aAOS)
Lacto−N−tetraose (LNT)
Lacto−N−neotetraose (LNnT)
2'−fucosyllactose (2'FL)
3'−sialyllactose (SHMO)
6'−sialyllactose (SHMO)
Potato soluble starch (ST)
Pullulan (PUL)
Gum arabic from acacia tree (GA)
Konjac glucomannan (bMAN)
Sugar beet arabinan (AR)
Tamarind xyloglucan (XGL)
OD600
0.0
0.5
1.0
1.5
0
10
20
30
40
Time (h)

#### 11.31 Selected growth curves

##### 11.31.1 Growth in MRS-AC supplemented with 0.5% tamarind xyloglucan

```
XGL_output_file <- "results/growth/XGL.pdf"
XGL_data <- "data/growth/formatted_selected/XGL/growth_xyloglucan.txt"
XGL_annotation <- "data/growth/formatted_selected/XGL/growth_xyloglucan_ann.txt"
XGL_trim_at_time <- 40 # measurements until [input] hours
XGL_blank_wells <- c("blank_1","blank_2","blank_3") # specify blank wells

# read a file with measurements
XGL_d <- read_tsv(XGL_data) %>%
  filter(time <= XGL_trim_at_time)
# calculate mean blank measurements (MRS-AC-Lac without added cells)
XGL_blank <- XGL_d %>%
  select(all_of(XGL_blank_wells)) %>%
  rowMeans()
# subtract blank measurements and round OD600 values to 3 digits
XGL_d_bl <- XGL_d %>%
  mutate(across(2:last_col(), ~ .x - XGL_blank)) %>%
  mutate(across(2:last_col(), ~ round(.x, 3)))
# read files with plate annotations
XGL_ann <- read_tsv(XGL_annotation)
XGL_d_bl_long <- XGL_d_bl %>%
  gather(., well, od, -time)
# create annotated tables
XGL_d_bl_long_ann <- left_join(XGL_d_bl_long, XGL_ann, by="well") %>%
  filter(carbon_source != "blank")
# plot growth curves
ggplot(XGL_d_bl_long_ann, aes(time, od, color = strain, fill = strain)) +
  #geom_point() +
  # add line connecting means from three replicates
  stat_summary(
    fun = mean,
    geom='line',
    size=0.5) +
  # add errorbars (standard deviation)
  stat_summary(
    fun.data=mean_sd,
    geom='errorbar',
    size=0.2,
    width=0.2,
    alpha=1) +
  scale_color_manual(name = "Taxonomy",
                     breaks=c("B. adolescentis LFYP80",
                              "B. catenulatum ssp. kashiwanohense Bg42221_1E1",
                              "B. catenulatum subsp. kashiwanohense_A Bg42221_1D3",
                              "B. longum ssp. longum Bg115.S08_3A11",
                              "B. pseudocatenulatum LFYP29"),
                     values=c("black", "tomato2","#00a2ff", "#51796f", "#e68607")) +
  scale_x_continuous(limits=c(0, XGL_trim_at_time),
                     breaks=c(0, XGL_trim_at_time*0.25, XGL_trim_at_time*0.5,
                              XGL_trim_at_time*0.75, XGL_trim_at_time)) +
```

```
  xlab("Time (h)") +
  ylab("OD600") +
  theme_bw() +
  theme(axis.text = element_text(color = "black"))
```

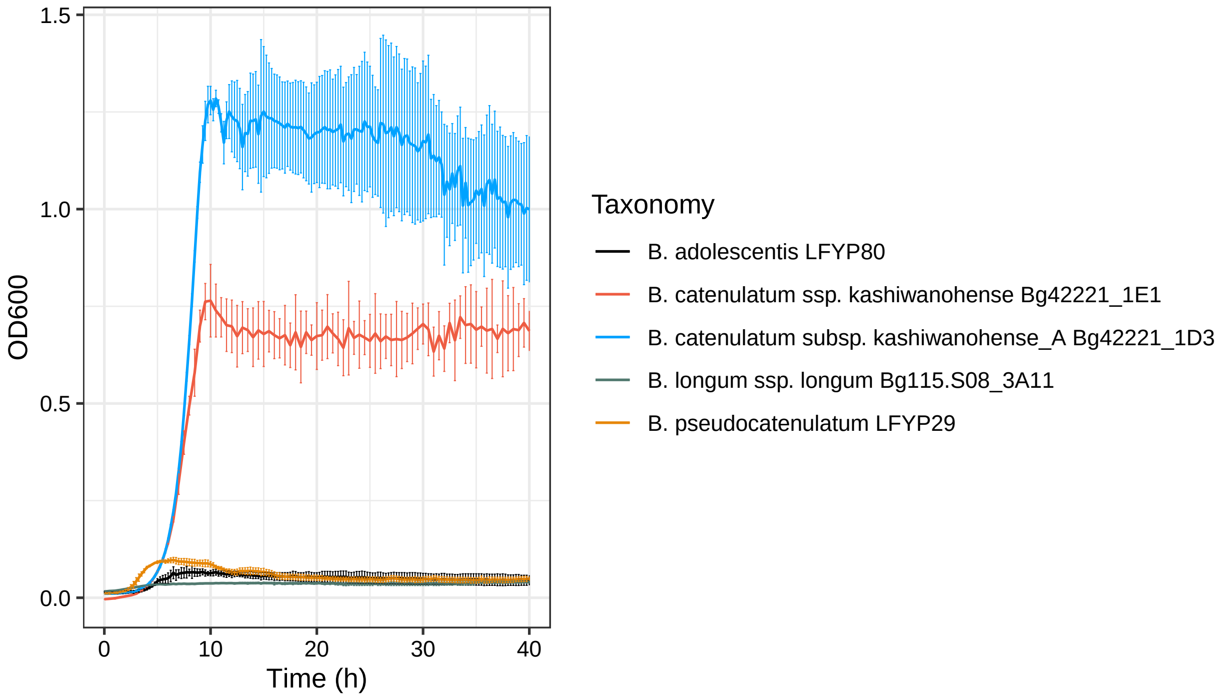

```
# save the plot as pdf
ggsave(filename = XGL_output_file, width = 7, height = 4, units = "in", dpi = 300)
```

##### 11.31.2 Growth in MRS-AC supplemented with 1% N-acetylneuraminic acid

```
# set input files
NANA_output_file <- "results/growth/Neu5Ac.pdf"
NANA_data <- "data/growth/formatted_selected/Neu5Ac/growth_sialic_acid.txt"
NANA_annotation <- "data/growth/formatted_selected/Neu5Ac/growth_sialic_acid_ann.txt"
NANA_trim_at_time <- 40 # measurements until [input] hours
NANA_blank_wells <- c("blank_1","blank_2","blank_3") # specify blank wells

# read a file with measurements
NANA_d <- read_tsv(NANA_data) %>%
  filter(time <= NANA_trim_at_time)
# calculate mean blank measurements (MRS-AC-Lac without added cells)
NANA_blank <- NANA_d %>%
  select(all_of(NANA_blank_wells)) %>%
  rowMeans()
# subtract blank measurements and round OD600 values to 3 digits
NANA_d_bl <- NANA_d %>%
```

```r
  mutate(across(2:last_col(), ~ .x - NANA_blank)) %>%
  mutate(across(2:last_col(), ~ round(.x, 3)))
# read files with plate annotations
NANA_ann <- read_tsv(NANA_annotation)
NANA_d_bl_long <- NANA_d_bl %>%
  gather(., well, od, -time)
# create annotated tables
NANA_d_bl_long_ann <- left_join(NANA_d_bl_long, NANA_ann, by="well") %>%
  filter(carbon_source != "blank")
# plot growth curves
ggplot(NANA_d_bl_long_ann, aes(time, od, color = strain, fill = strain)) +
  # add line connecting means from three replicates
  stat_summary(
    fun = mean,
    geom='line',
    size=0.5) +
  # add errorbars (standard deviation)
  stat_summary(
    fun.data=mean_sd,
    geom='errorbar',
    size=0.2,
    width=0.2,
    alpha=1) +
  scale_color_manual(name = "Taxonomy",
                     breaks=c("B. breve Bg41721_1C11", "B. breve Bg155.S08_4F7",
                             "B. breve JG_Bg463", "B. breve LFYP24",
                             "B. breve Bg131.S11_D6"),
                     values=c("#51796f", "#ffa600", "#00a2ff", "#00b400",
                              "black")) +
  scale_x_continuous(limits=c(0, NANA_trim_at_time),
                   breaks=c(0, NANA_trim_at_time*0.25, NANA_trim_at_time*0.5,
                           NANA_trim_at_time*0.75, NANA_trim_at_time)) +
  xlab("Time (h)") +
  ylab("OD600") +
  theme_bw() +
  theme(axis.text = element_text(color = "black"))
```

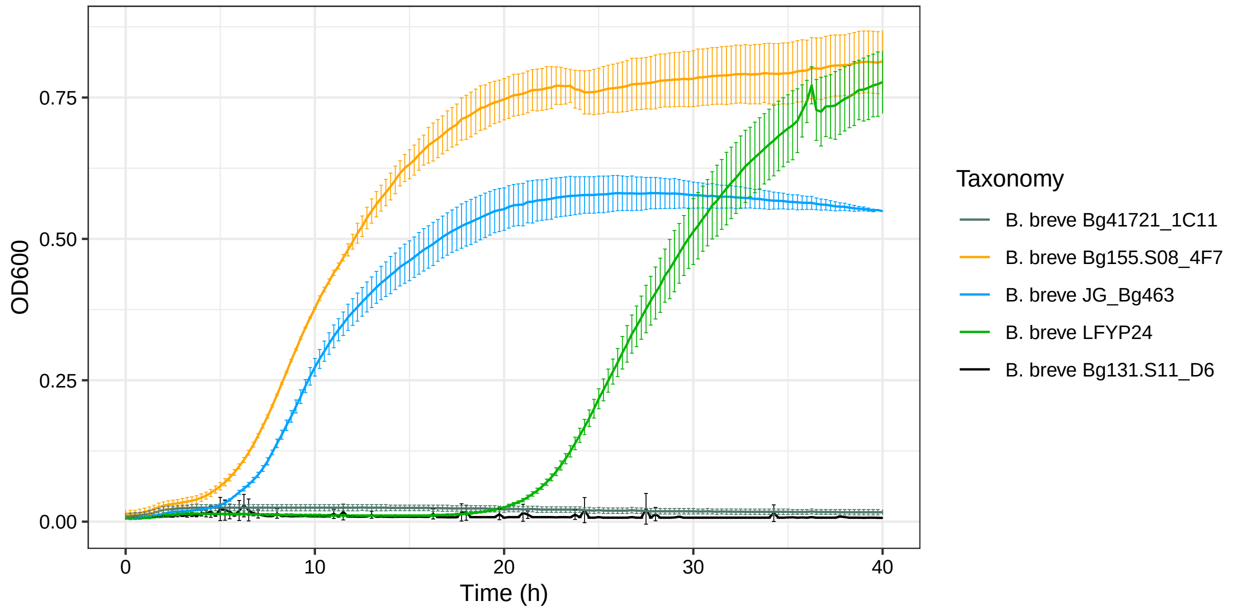

```
# save the plot as pdf
ggsave(filename = NANA_output_file, width = 8, height = 4, units = "in", dpi = 300)
```

###### 11.31.3 Growth in MRS-AC supplemented with 0.5% mannotriose or 0.5% konjac glucomannan

```
bMAN_output_file <- "results/growth/bMAN.pdf"
bMAN_data <- "data/growth/formatted_selected/bMAN/growth_bMAN.txt"
bMAN_annotation <- "data/growth/formatted_selected/bMAN/growth_bMAN_ann.txt"
bMAN_trim_at_time <- 40 # measurements until [input] hours
bMAN_blank_wells <- c("blank_1","blank_2","blank_3") # specify blank wells

# read a file with measurements
bMAN_d <- read_tsv(bMAN_data) %>%
  filter(time <= bMAN_trim_at_time)
# calculate mean blank measurements (MRS-AC-Lac without added cells)
bMAN_blank <- bMAN_d %>%
  select(all_of(bMAN_blank_wells)) %>%
  rowMeans()
# subtract blank measurements and round OD600 values to 3 digits
bMAN_d_bl <- bMAN_d %>%
  mutate(across(2:last_col(), ~ .x - bMAN_blank)) %>%
  mutate(across(2:last_col(), ~ round(.x, 3)))
# read files with plate annotations
bMAN_ann <- read_tsv(bMAN_annotation)
bMAN_d_bl_long <- bMAN_d_bl %>%
  gather(., well, od, -time)
# create annotated tables
bMAN_d_bl_long_ann <- left_join(bMAN_d_bl_long, bMAN_ann, by="well") %>%
  filter(carbon_source != "blank") %>%
  mutate(carbon_source = factor(carbon_source, levels = c("Mannotriose (bMnOS)",
```

```
                                                            "Konjac glucomannan (bMAN)")))
# plot growth curves
ggplot(bMAN_d_bl_long_ann, aes(time, od, color = strain, fill = strain)) +
  # add line connecting means from three replicates
  stat_summary(
    fun = mean,
    geom='line',
    size=0.5) +
  # add errorbars (standard deviation)
  stat_summary(
    fun.data=mean_sd,
    geom='errorbar',
    size=0.2,
    width=0.2,
    alpha=1) +
  scale_color_manual(name = "Taxonomy",
                     breaks=c("B. breve Bg41721_1C11", "B. breve Bg155.S08_4F7",
                              "B. breve LFYP81", "B. dentium LFYP24",
                              "B. scardovii JCM 12489"),
                     values=c("#51796f", "#ffa600", "#00b400", "#8e063c", "grey")) +
  scale_x_continuous(limits=c(0, bMAN_trim_at_time),
                     breaks=c(0, bMAN_trim_at_time*0.25, bMAN_trim_at_time*0.5,
                              bMAN_trim_at_time*0.75, bMAN_trim_at_time)) +
  xlab("Time (h)") +
  ylab("OD600") +
  facet_wrap(~carbon_source) +
  theme_bw() +
  theme(axis.text = element_text(color = "black"))
```

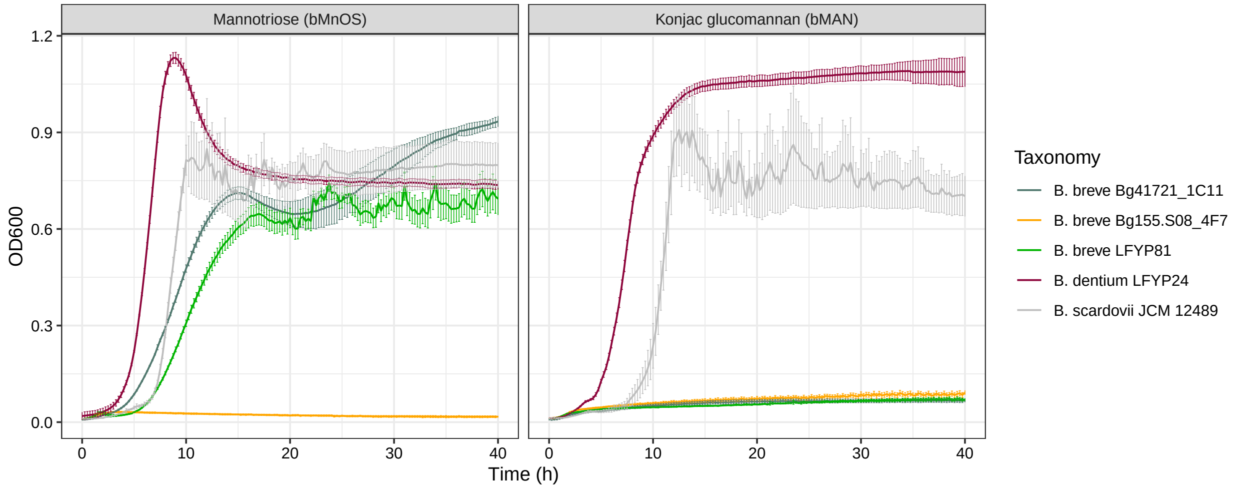

```
# save the plot as pdf
ggsave(filename = bMAN_output_file, width = 10, height = 4, units = "in", dpi = 300)
```

##### 11.31.4 Growth in MRS-AC supplemented with 1% D-glucuronic acid sodium salt

```
GlcA_output_file <- "results/growth/GlcA.pdf"
GlcA_data <- "data/growth/formatted_selected/GlcA/growth_GlcA.txt"
GlcA_annotation <- "data/growth/formatted_selected/GlcA/growth_GlcA_ann.txt"
GlcA_trim_at_time <- 40 # measurements until [input] hours
GlcA_blank_wells <- c("blank_1","blank_2","blank_3") # specify blank wells

# read a file with measurements
GlcA_d <- read_tsv(GlcA_data) %>%
  filter(time <= GlcA_trim_at_time)
# calculate mean blank measurements (MRS-AC-Lac without added cells)
GlcA_blank <- GlcA_d %>%
  select(all_of(GlcA_blank_wells)) %>%
  rowMeans()
# subtract blank measurements and round OD600 values to 3 digits
GlcA_d_bl <- GlcA_d %>%
  mutate(across(2:last_col(), ~ .x - GlcA_blank)) %>%
  mutate(across(2:last_col(), ~ round(.x, 3)))
# read files with plate annotations
GlcA_ann <- read_tsv(GlcA_annotation)
GlcA_d_bl_long <- GlcA_d_bl %>%
  gather(., well, od, -time)
# create annotated tables
GlcA_d_bl_long_ann <- left_join(GlcA_d_bl_long, GlcA_ann, by="well") %>%
  filter(carbon_source != "blank")
# plot growth curves
ggplot(GlcA_d_bl_long_ann, aes(time, od, color = strain, fill = strain)) +
  # add line connecting means from three replicates
  stat_summary(
    fun = mean,
    geom='line',
    size=0.5) +
  # add errorbars (standard deviation)
  stat_summary(
    fun.data=mean_sd,
    geom='errorbar',
    size=0.2,
    width=0.2,
    alpha=1) +
  scale_color_manual(name = "Taxonomy",
                     breaks=c("B. breve Bg41721_1C11",
                              "B. longum ssp. infantis ATCC15697",
                              "B. longum ssp. infantis JG_Bg463",
                              "B. longum ssp. infantis Bg40721_2D9"),
                     values=c("#51796f", "#2032ab", "#8e063c", "black")) +
  scale_x_continuous(limits=c(0, GlcA_trim_at_time),
                     breaks=c(0, GlcA_trim_at_time*0.25, GlcA_trim_at_time*0.5,
                              GlcA_trim_at_time*0.75, GlcA_trim_at_time)) +
  xlab("Time (h)") +
  ylab("OD600") +
  theme_bw() +
  theme(axis.text = element_text(color = "black"))
```

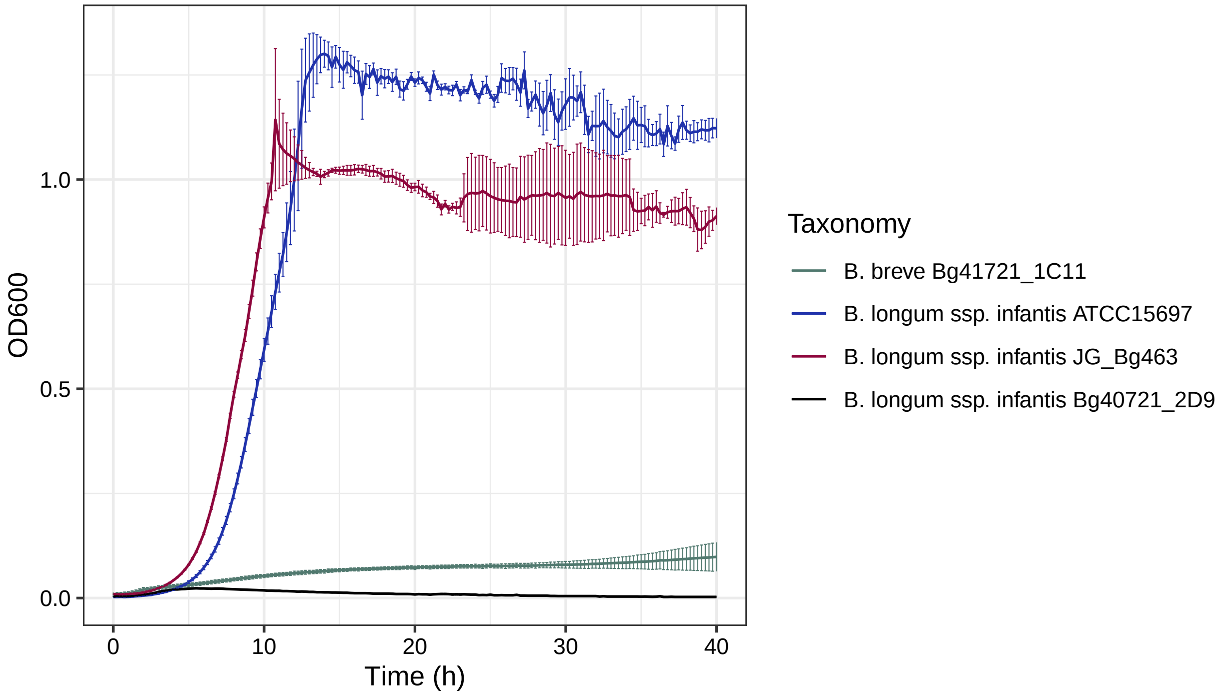

```
# save the plot as pdf
ggsave(filename = GlcA_output_file, width = 7, height = 4, units = "in", dpi = 300)
```

###### 11.31.5 Growth in MRS-AC supplemented with 1% HMOs from pooled human milk (pHMOs)

```
# load libraries
library(tidyverse)
library(ggpubr)
# read_data
pHMO_output_file <- "results/growth/pHMO_24h.pdf"
pHMO_data <- "data/growth/formatted_selected/pHMO/growth_pHMO_24h.txt"
pHMO_annotation <- "data/growth/formatted_selected/pHMO/growth_pHMO_24h_ann.txt"
trim_at_time <- 24 # measurements until [input] hours
blank_wells_HMO <- c("blank_1","blank_2","blank_3") # specify blank wells

# read a file with measurements
d_HMO <- read_tsv(pHMO_data) %>%
  filter(time <= trim_at_time)
# calculate mean blank measurements (MRS-AC-Lac without added cells)
blank_HMO <- d_HMO %>%
  dplyr::select(all_of(blank_wells_HMO)) %>%
  rowMeans()
# subtract blank measurements and round OD600 values to 3 digits
d_bl_HMO <- d_HMO %>%
  mutate(across(2:last_col(), ~ .x - blank_HMO)) %>%
  mutate(across(2:last_col(), ~ round(.x, 3)))
# read files with plate annotations
```

```r
ann <- read_tsv(pHMO_annotation)
d_bl_long_HMO <- d_bl_HMO %>%
  gather(., well, od, -time)
# create annotated tables
d_bl_long_HMO_ann <- left_join(d_bl_long_HMO, ann, by="well") %>%
  filter(carbon_source != "blank")

# plot growth curves
ggplot(d_bl_long_HMO_ann, aes(time, od, color = strain, fill = strain)) +
  #geom_point() +
  # add line connecting means from three replicates
  stat_summary(
    fun = mean,
    geom='line',
    size=0.5) +
  # add errorbars (standard deviation)
  stat_summary(
    fun.data=mean_sd,
    geom='errorbar',
    size=0.2,
    width=0.2,
    alpha=1) +
  scale_color_manual(name = "Taxonomy",
                     breaks=c("B. breve Bg155.S08_4F7",
                              "B. catenulatum ssp. kashiwanohense Bg42221_1E1",
                              "B. catenulatum ssp. kashiwanohense_A Bg42221_1D3",
                              "B. longum ssp. infantis ATCC 15697",
                              "B. longum ssp. infantis Bg40721_2D9",
                              "B. longum ssp. longum Bg115.S08_3A11",
                              "B. longum ssp. suis Bg131.S11_17.F6",
                              "B. longum ssp. suis Bg41121_2E1",
                              "B. longum (ssp. nov.) LFYP82",
                              "B. pseudocatenulatum LFYP29"),
                     values=c("#00a2ff", "#52ab35", "#ffa600",
                              "#52ab35", "#52ab35", "#00a2ff",
                              "#52ab35", "#ffa600", "black",
                              "black")) +
scale_x_continuous(limits=c(0, trim_at_time),
                   breaks=c(0, trim_at_time*0.25, trim_at_time*0.5,
                            trim_at_time*0.75, trim_at_time)) +
  xlab("Time (h)") +
  ylab("OD600") +
  theme_bw() +
  theme(axis.text = element_text(color = "black"))
```

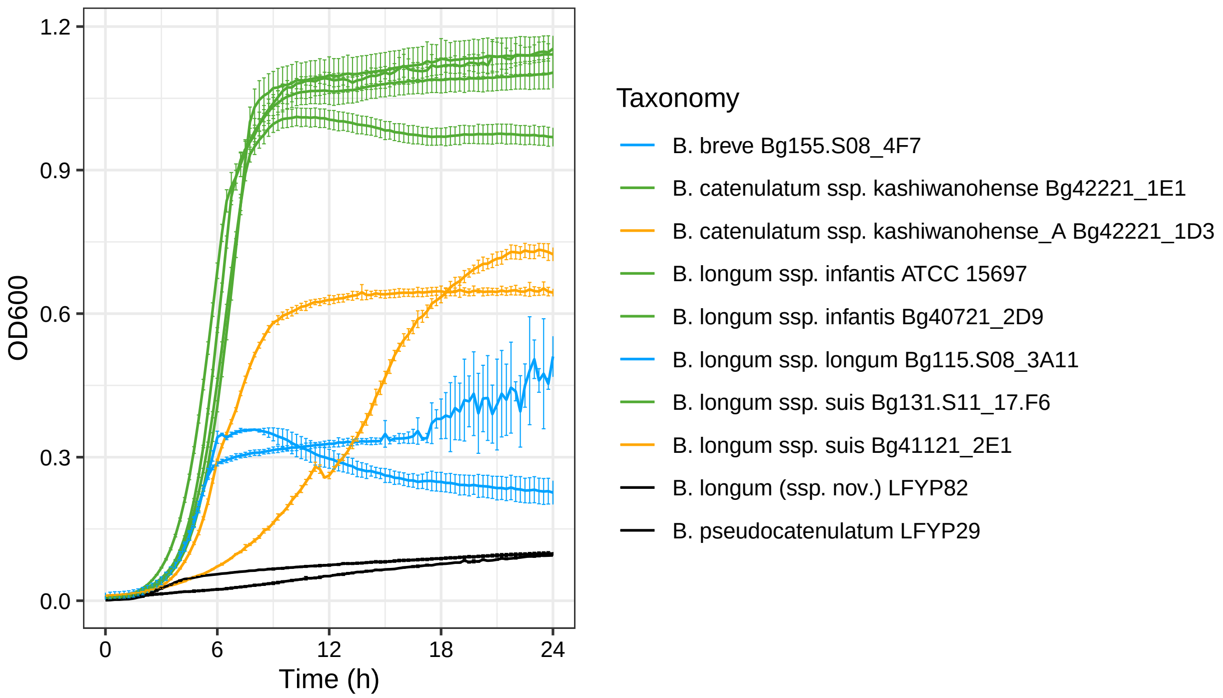

```
# save the plot as pdf
ggsave(filename = pHMO_output_file, width = 7, height = 4, units = "in", dpi = 300)
```

#### 11.32 Summary of growth data

The code chunks below describe the comparison of predicted binary carbohydrate utilization (CU) phenotypes with *in vitro* growth data obtained in this work or described previously in the literature (Bottacini et al., 2014, Arboleya et al., 2018, and Orihara et al. 2023).

Here we compare predicted binary CU phenotypes (**1** or **0**) with growth phenotypes (**+**/**w** and **-**) and calculate the number of true positives (TP), true negatives (TN), false positives (FP), and false negatives (FN). We then calculate various metrics (precision, recall, specificity, accuracy, F-score, and Matthew's correlation coefficient) based on TP, TN, FP, and FN counts.

```
# define a function that will calculate various metrics based on TP, TN, FP, and FN counts
compute_metrics <- function(df) {
  df <- df %>%
    mutate(
      Precision = round(TruePositive / (TruePositive + FalsePositive), 2),
      Recall = round(TruePositive / (TruePositive + FalseNegative), 2),
      Specificity = round(TrueNegative / (TrueNegative + FalsePositive), 2),
      Accuracy = round((TruePositive + TrueNegative) /
                       (TruePositive + TrueNegative + FalsePositive + FalseNegative), 2),
      F_score = round(2 * (Precision * Recall) / (Precision + Recall), 2),
      MCC = round((TruePositive * TrueNegative - FalsePositive * FalseNegative) /
        sqrt((TruePositive + FalsePositive) * (TruePositive + FalseNegative) *
              (TrueNegative + FalsePositive) * (TrueNegative + FalseNegative)), 2)
    )

  return(df)
```

```
}

# read files
# BPM and the summary of growth data for 30 strains from this work
# all strains
bpm_arz <- read_tsv("data/growth/summary/bif_30_strains_BPM.txt") %>%
  select(-country,-dataset)
growth_arz <- read_tsv("data/growth/summary/bif_30_strains_growth_data.txt") %>%
  select(-country,-dataset)
# only reference strains
bpm_arz_ref <- read_tsv("data/growth/summary/bif_30_strains_BPM.txt") %>%
  filter(dataset == "ref") %>%
  select(-country,-dataset)
growth_arz_ref <- read_tsv("data/growth/summary/bif_30_strains_growth_data.txt") %>%
  filter(dataset == "ref") %>%
  select(-country,-dataset)
# only new strains
bpm_arz_ext <- read_tsv("data/growth/summary/bif_30_strains_BPM.txt") %>%
  filter(dataset == "ext") %>%
  select(-country,-dataset)
growth_arz_ext <- read_tsv("data/growth/summary/bif_30_strains_growth_data.txt") %>%
  filter(dataset == "ext") %>%
  select(-country,-dataset)

# BPM and the summary of growth data for 6 strains from Bottacini et al., 2014
bpm_bot <- read_tsv("data/growth/summary/Bottacini_2014_BPM.txt")
growth_bot <- read_tsv("data/growth/summary/Bottacini_2014_growth_data.txt")

# BPM and the summary of growth data for 19 strains from Arboleya et al., 2018
bpm_arb <- read_tsv("data/growth/summary/Arboleya_2018_BPM.txt")
growth_arb <- read_tsv("data/growth/summary/Arboleya_2018_growth_data.txt")

# BPM and the summary of growth data for 8 strains from Orihara et al. 2023
bpm_ori <- read_tsv("data/growth/summary/Orihara_2023_BPM.txt")
growth_ori <- read_tsv("data/growth/summary/Orihara_2023_growth_data.txt")

# phenotype metadata
phenotype_metadata_gr <- read_tsv("data/tables/phenotype_metadata_carbs.txt") %>%
  dplyr::select(phenotype, type_group)

# calculate the number TP, TN, FP, and FN counts
# growth data from this work
results_arz <- calculate_metrics_within_groups(bpm_arz, growth_arz, phenotype_metadata_gr)
results_arz_df <- bind_rows(results_arz, .id = "Group")
results_arz_ref <- calculate_metrics_within_groups(bpm_arz_ref, growth_arz_ref, phenotype_metadata_gr)
results_arz_ref_df <- bind_rows(results_arz_ref, .id = "Group")
results_arz_ext <- calculate_metrics_within_groups(bpm_arz_ext, growth_arz_ext, phenotype_metadata_gr)
results_arz_ext_df <- bind_rows(results_arz_ext, .id = "Group")
# growth data from Bottacini et al., 2014
results_bot <- calculate_metrics_within_groups(bpm_bot, growth_bot, phenotype_metadata_gr)
results_bot_df <- bind_rows(results_bot, .id = "Group")
```

| Group | Tested | TruePositive | TrueNegative | FalsePositive | FalseNegative |
|---|---|---|---|---|---|
| monosaccharides_and_derivatives | 237 | 123 | 93 | 17 | 4 |
| di_and_oligosaccharides | 252 | 152 | 86 | 13 | 1 |
| polysaccharides | 207 | 81 | 121 | 5 | 0 |
| AllGroups | 696 | 356 | 300 | 35 | 5 |

```
# growth data from Arboleya et al., 2018
results_arb <- calculate_metrics_within_groups(bpm_arb, growth_arb, phenotype_metadata_gr)
results_arb_df <- bind_rows(results_arb, .id = "Group")
# growth data from Orihara et al., 2023
results_ori <- calculate_metrics_within_groups(bpm_ori, growth_ori, phenotype_metadata_gr)
results_ori_df <- bind_rows(results_ori, .id = "Group")
# combine TP, TN, FP, and FN count data: Bottacini et al., 2014 + Arboleya et al., 2018
# + Orihara et al., 2023
desired_order <- c(
  "monosaccharides_and_derivatives",
  "di_and_oligosaccharides",
  "polysaccharides",
  "AllGroups"
)
results_papers_df <- bind_rows(results_bot_df, results_arb_df, results_ori_df) %>%
  group_by(Group) %>%
  summarise(
    Tested = sum(Tested),
    TruePositive = sum(TruePositive),
    TrueNegative = sum(TrueNegative),
    FalsePositive = sum(FalsePositive),
    FalseNegative = sum(FalseNegative)
  ) %>%
  arrange(match(Group, desired_order))
```

Summary for data from Bottacini et al., 2014, Arboleya et al., 2018, and Orihara et al. 2023:

```
# calculate the metrics for TP, TN, FP, and FN counts
results_papers_df_m <- compute_metrics(results_papers_df)
# use the gt package to produce the table
gt(results_papers_df_m  ) %>%
  cols_align(
    align = "left",
    columns = everything()
  )
```

Summary for data obtained in this work for 20 reference strains:

```
# calculate the metrics for TP, TN, FP, and FN counts
results_arz_ref_df_m <- compute_metrics(results_arz_ref_df)
# use the gt package to produce the table
gt(results_arz_ref_df_m) %>%
  cols_align(
    align = "left",
```

| Group | Tested | TruePositive | TrueNegative | FalsePositive | FalseNegative |
|---|---|---|---|---|---|
| monosaccharides_and_derivatives | 260 | 119 | 121 | 10 | 10 |
| di_and_oligosaccharides | 357 | 229 | 114 | 6 | 8 |
| polysaccharides | 139 | 21 | 115 | 3 | 0 |
| AllGroups | 756 | 369 | 350 | 19 | 18 |

| Group | Tested | TruePositive | TrueNegative | FalsePositive | FalseNegative |
|---|---|---|---|---|---|
| monosaccharides_and_derivatives | 130 | 52 | 70 | 4 | 4 |
| di_and_oligosaccharides | 177 | 101 | 65 | 6 | 5 |
| polysaccharides | 69 | 7 | 60 | 2 | 0 |
| AllGroups | 376 | 160 | 195 | 12 | 9 |

| Group | Tested | TruePositive | TrueNegative | FalsePositive | FalseNegative |
|---|---|---|---|---|---|
| monosaccharides_and_derivatives | 390 | 171 | 191 | 14 | 14 |
| di_and_oligosaccharides | 534 | 330 | 179 | 12 | 13 |
| polysaccharides | 208 | 28 | 175 | 5 | 0 |
| AllGroups | 1132 | 529 | 545 | 31 | 27 |

```
    columns = everything()
  )
```

Summary for data obtained in this work for 10 additional strains:

```
# calculate the metrics for TP, TN, FP, and FN counts
results_arz_ext_df_m <- compute_metrics(results_arz_ext_df)
# use the gt package to produce the table
gt(results_arz_ext_df_m) %>%
  cols_align(
    align = "left",
    columns = everything()
  )
```

Summary for data obtained in this work for all 30 strains:

```
# calculate the metrics for TP, TN, FP, and FN counts
results_arz_df_m <- compute_metrics(results_arz_df)
# use the gt package to produce the table
gt(results_arz_df_m) %>%
  cols_align(
    align = "left",
    columns = everything()
  )
```

Plot the heatmap with growth results:

```r
# read full growth data
growth_data <- read_tsv("data/growth/summary/Bif_30_strains_growth_data_full.txt")
# extract the binary matrix
growth_mat <- as.matrix((growth_data[, 2:44]))
# add rownames to the matrix
growth_mat_ID <- growth_data$genome_ID
rownames(growth_mat) <- growth_mat_ID

# create annotation vectors
genome_group <- growth_data$dataset
genome_country <- growth_data$country

# create a coloring function
col_fun <- structure(c("#949599", "white", "#d7f2a2", "#5e9155"), names = c("NT","-", "w", "+"))
# create two row annotations specifying taxonomy
ha_row <- HeatmapAnnotation(
  which = c("row"),
  country = genome_country,
  col = list(country = c("US" = "#bdbad9",
                         "Bangladesh" = "#51796f",
                         "Malawi" = "#ffa600",
                         "Europe" = "black")),
             show_annotation_name = FALSE,
             simple_anno_size = unit(3, "mm"))

# plot the heatmap
pdf("results/growth/growth_summary.pdf", width=20, height=20)
ht_growth_data <- ComplexHeatmap::Heatmap(growth_mat,
                              name = "Summary of growth data",
                              right_annotation = ha_row,
                              col = col_fun,
                              cluster_rows = F,
                              cluster_columns = F,
                              rect_gp = gpar(col = "grey", lwd = 0.05),
                              show_row_names = TRUE,
                              row_names_gp = gpar(fontsize = 3),
                              column_names_gp = gpar(fontsize = 5),
                              column_names_rot = 60,
                              column_names_side = "top",
                              width = unit(350, "mm"),
                              height = unit(200, "mm"))
draw(ht_growth_data)
# save the heatmap to a file
dev.off()
```

```
## pdf
##   2
```

```r
# draw the heatmap again to show it in the compiled markdown file
draw(ht_growth_data)
```

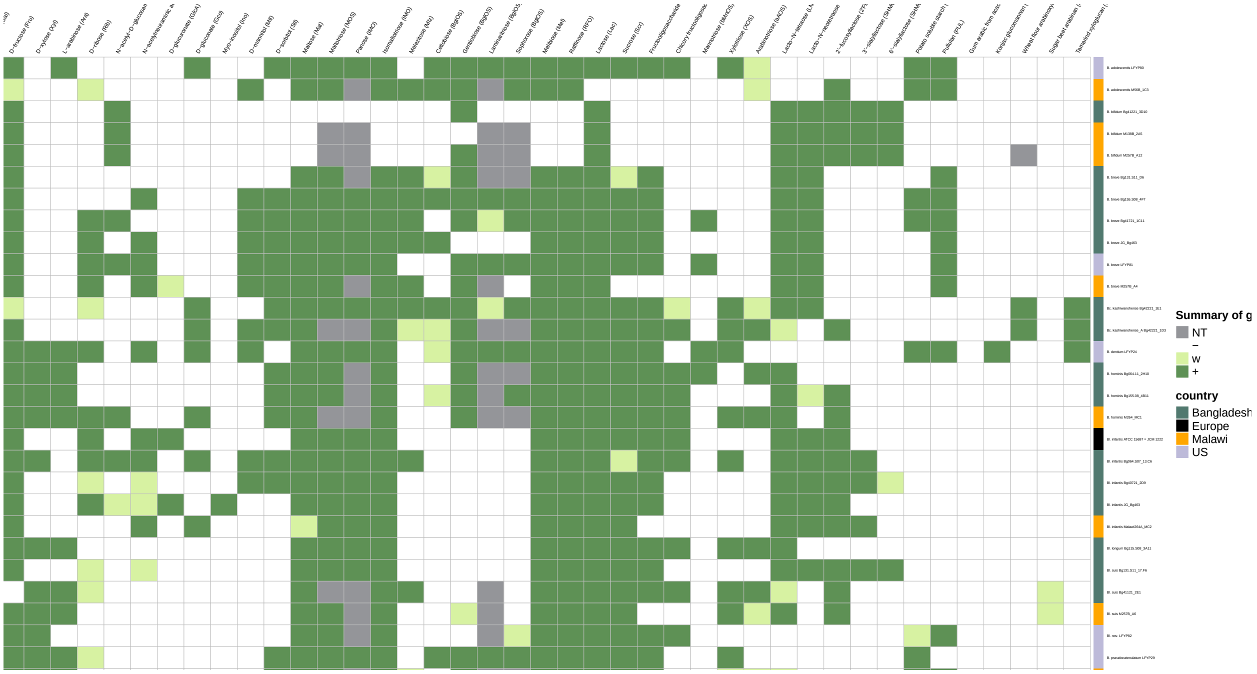

Plot the heatmap with comparison results:

```
# assuming bpm_arz, growth_arz, phenotype_metadata_gr are already defined
# modify comparison table creation to handle NT values
comparison_table <- bpm_arz %>%
  mutate(across(-genome_ID, ~case_when(
    growth_arz[[cur_column()]] == "NT" ~ NA_character_,  # Handle NT first
    . == 1 & growth_arz[[cur_column()]] %in% c("+", "w") ~ "TP",
    . == 0 & growth_arz[[cur_column()]] %in% c("-") ~ "TN",
    . == 1 & growth_arz[[cur_column()]] %in% c("-") ~ "FP",
    . == 0 & growth_arz[[cur_column()]] %in% c("+", "w") ~ "FN",
    TRUE ~ NA_character_  # Ensure all other cases are NA
  )))

# extract the binary matrix
comparison_table_mat <- as.matrix(comparison_table[, 2:ncol(comparison_table)])

# add rownames to the matrix
rownames(comparison_table_mat) <- comparison_table$genome_ID

# extract vectors containing data about glycan type and origin
matrix_column_names <- colnames(comparison_table_mat)
matching_indices <- match(matrix_column_names, phenotype_metadata_gr$phenotype)
glycan_type_vector <- phenotype_metadata_gr$type_group[matching_indices]

# update coloring function to include white for NA (NT cases)
comp_col_fun <- structure(
  c("#C5DEC6", "#C5DEC6", "#fffdbc", "#fffdbc", "#FFFFFF"),
  names = c("TP", "TN", "FP", "FN", NA_character_)
)

# create a column annotation specifying glycan type
```

```r
comp_ha_ann <- HeatmapAnnotation(
  type = glycan_type_vector,
  col = list(type = c("monosaccharides_and_derivatives" = "#FFFFFF",
                      "di_and_oligosaccharides" = "#E6E7E8",
                      "polysaccharides" = "#BCBEC0")),
  show_annotation_name = FALSE,
  simple_anno_size = unit(3, "mm")
)

# plot the heatmap
pdf("results/growth/predictions_vs_growth.pdf", width=15, height=8)

ht_growth <- ComplexHeatmap::Heatmap(comparison_table_mat,
                              name = "Legend",
                              heatmap_legend_param = list(title_position = "topcenter"),
                              bottom_annotation = comp_ha_ann,
                              col = comp_col_fun,
                              cluster_rows = F,
                              cluster_columns = F,
                              rect_gp = gpar(col = "white", lwd = 0.5),
                              show_row_names = TRUE,
                              row_names_gp = gpar(fontsize = 8),
                              column_names_gp = gpar(fontsize = 8),
                              column_names_rot = 45,
                              width = unit(175, "mm"),
                              height = unit(100, "mm"),
                              cell_fun = function(j, i, x, y, width, height, fill) {
                                cell_value <- comparison_table_mat[i, j]
                                if (is.na(cell_value)) {
                                  return()  # Do nothing for NT cases (NA)
                                } else if (cell_value == "FP") {
                                  grid.text(cell_value, x, y, gp = gpar(fontsize = 8, col = "#2c64a3"))
                                } else if (cell_value == "FN") {
                                  grid.text(cell_value, x, y, gp = gpar(fontsize = 8, col = "#be5d20"))
                                } else {
                                  grid.text(cell_value, x, y, gp = gpar(fontsize = 8))
                                }
                              })

draw(ht_growth, heatmap_legend_side = "top", annotation_legend_side = "top")
dev.off()
```

```
## pdf
##   2
```

```r
draw(ht_growth)
```

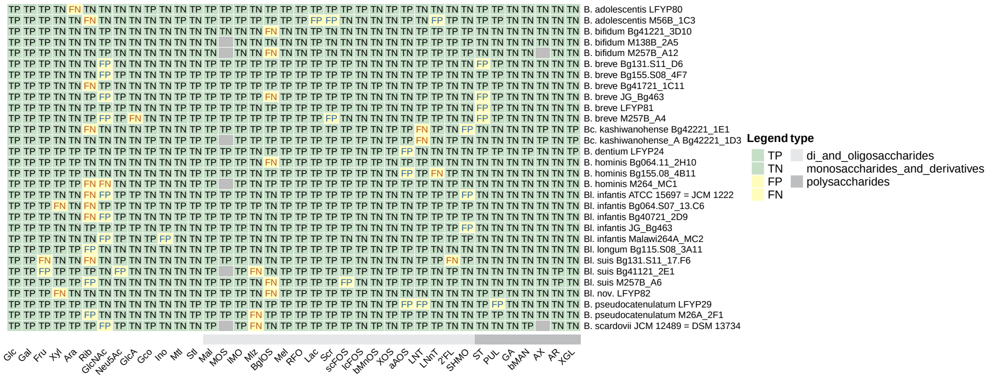

### 12 Analysis of human milk oligosaccharide (HMO) consumption data

#### 12.1 Introduction

This block describes the analysis of tables with information about the concentration of individual HMOs (nmol/mL) in culture supernatants of select *Bifidobacterium* strains grown in MRS-AC-pHMO for (a) 8 hours or (b) 24 hours. The supernatants of two different timepoints were collected in four separate experiments

#### 12.2 Load data

```
# read data
pHMO_data_8h <- "data/hmo_consumption/HMO_consumption_8h.txt"
pHMO_data_24h <- "data/hmo_consumption/HMO_consumption_24h.txt"
```

##### 12.2.1 HMO consumption after 24h

```
# read files with measurements
HMO_24h <- read_tsv(pHMO_data_24h) %>%
  mutate(across(3:last_col(), ~ round(.x, 1)))
# calculate average values
HMO_24h_avg <- HMO_24h %>%
  group_by(strain) %>%
```

```r
  summarize(across(where(is.numeric), mean, na.rm = TRUE)) %>%
  mutate(across(2:last_col(), ~ round(.x, 1)))
# calculate the percent of utilized HMOs
group1_strains <- c("B. catenulatum ssp. kashiwanohense Bg42221_1E1",
                    "B. longum ssp. infantis ATCC 15697",
                    "B. longum ssp. infantis Bg40721_2D9",
                    "B. longum ssp. longum Bg115.S08_3A11",
                    "B. pseudocatenulatum LFYP29")
group2_strains <- c("B. breve Bg155.S08_4F7",
                    "B. catenulatum ssp. kashiwanohense_A Bg42221_1D3",
                    "B. longum (ssp. nov.) LFYP82",
                    "B. longum ssp. suis Bg131.S11_17.F6",
                    "B. longum ssp. suis Bg41121_2E1")
HMO_24h_ut <- HMO_24h_avg %>%
  mutate(across(-strain, ~ case_when(
    strain %in% group1_strains ~ 1 - (. / HMO_24h_avg %>% filter(strain == "medium_batch1")
                                  %>% pull(cur_column())),
    strain %in% group2_strains ~ 1 - (. / HMO_24h_avg %>% filter(strain == "medium_batch2")
                                  %>% pull(cur_column())),
    TRUE ~ .  # leave others unchanged
  ))) %>%
  mutate(across(2:last_col(), ~ round(.x, 3))) %>%
  filter(!strain %in% c("medium_batch1", "medium_batch2"))
  #select(-c(total, total_sial, total_fuc))

# extract matrices
hmo_mat_24h <- as.matrix(HMO_24h_ut[,2:ncol(HMO_24h_ut)])
rownames(hmo_mat_24h) <- pull(HMO_24h_ut, strain)
desired_order <- c("LNT", "LNnT", "LNH", "2'FL", "3FL", "LDFT",
                   "LNFP I", "LNFP II", "LNFP III", "DFLNT", "FLNH",
                   "DFLNH", "3'SL", "6'SL", "LSTb", "LSTc", "DSLNT",
                   "DSLNH", "FDSLNH", "total", "total_fuc", "total_sial")
hmo_mat_24h <- hmo_mat_24h[, desired_order]

# add a coloring function
# backup legend (1, 0.875, 0.75, 0.625, 0.5, 0.375, 0.25, 0.125, 0)
col_fun <- colorRamp2(c(0, 0.125, 0.25, 0.375, 0.5, 0.625, 0.75, 0.875, 1),
                      c("#ffffff", "#DEEBF7", "#C6DBEF", "#9ECAE1", "#6BAED6",
                        "#4292C6", "#2171B5", "#08519C", "#08306B"))

lgd <- Legend(col_fun = col_fun,
              title = "% utilized",
              direction = "horizontal",
              title_position = "lefttop")
# plot the heatmap
pdf("results/HMO_utilization/heatmap_24h.pdf", width=15, height=4)
ht_hmo_24h <- ComplexHeatmap::Heatmap(hmo_mat_24h,
                                     name = "HMO utilization at 24h",
                                     col = col_fun,
                                     cluster_rows = TRUE,
                                     cluster_columns = FALSE,
                                     rect_gp = gpar(col = "grey", lwd = 0.05),
                                     heatmap_legend_param = list(
```

```
                                              col_fun = col_fun,
                                              at = c(0, 0.25, 0.5, 0.75, 1),
                                              title = "% of utilized HMOs",
                                              direction = "horizontal",
                                              title_position = "topcenter",
                                              border = "black",
                                              legend_width = unit(40, "mm")))
draw(ht_hmo_24h)
# save the heatmap to a file
dev.off()
```

```
## pdf
##   2
```

```
# draw the heatmap again to show it in the compiled markdown file
draw(ht_hmo_24h)
```

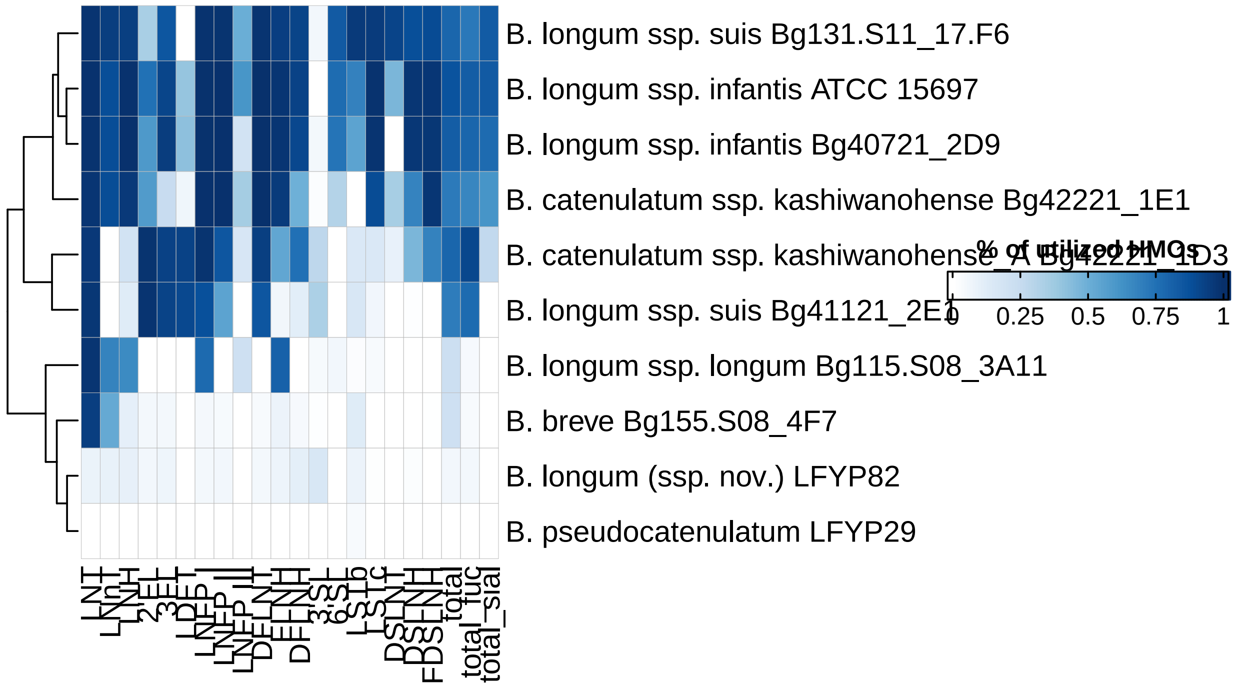

##### 12.2.2 HMO consumption after 8h

```
# read files with measurements
HMO_8h <- read_tsv(pHMO_data_8h) %>%
  mutate(across(3:last_col(), ~ round(.x, 1)))
# calculate average values
HMO_8h_avg <- HMO_8h %>%
  group_by(strain) %>%
  summarize(across(where(is.numeric), mean, na.rm = TRUE)) %>%
```

```r
  mutate(across(2:last_col(), ~ round(.x, 1)))
# calculate percent of utilized HMOs
group1_strains_8h <- c("B. breve Bg155.S08_4F7",
                       "B. catenulatum ssp. kashiwanohense Bg42221_1E1",
                       "B. longum ssp. infantis ATCC 15697",
                       "B. longum ssp. infantis Bg40721_2D9",
                       "B. longum ssp. longum Bg115.S08_3A11",
                       "B. pseudocatenulatum LFYP29")
group2_strains_8h <- c("B. catenulatum ssp. kashiwanohense_A Bg42221_1D3",
                       "B. longum (ssp. nov.) LFYP82",
                       "B. longum ssp. suis Bg131.S11_17.F6",
                       "B. longum ssp. suis Bg41121_2E1")
custom_order <- c("B. longum ssp. suis Bg131.S11_17.F6",
                  "B. longum ssp. infantis ATCC 15697",
                  "B. longum ssp. infantis Bg40721_2D9",
                  "B. catenulatum ssp. kashiwanohense Bg42221_1E1",
                  "B. catenulatum ssp. kashiwanohense_A Bg42221_1D3",
                  "B. longum ssp. suis Bg41121_2E1",
                  "B. longum ssp. longum Bg115.S08_3A11",
                  "B. breve Bg155.S08_4F7",
                  "B. longum (ssp. nov.) LFYP82",
                  "B. pseudocatenulatum LFYP29")
# calculate the percent of utilized HMOs
HMO_8h_ut <- HMO_8h_avg %>%
  mutate(across(-strain, ~ case_when(
    strain %in% group1_strains_8h ~ 1 - (. / HMO_8h_avg %>%
                                          filter(strain == "medium_batch1")
                                     %>% pull(cur_column())),
    strain %in% group2_strains_8h ~ 1 - (. / HMO_8h_avg %>%
                                          filter(strain == "medium_batch2")
                                     %>% pull(cur_column())),
    TRUE ~ .  # leave others unchanged
  ))) %>%
  mutate(across(2:last_col(), ~ round(.x, 3))) %>%
  filter(!strain %in% c("medium_batch1", "medium_batch2")) %>%
  mutate(strain = factor(strain, levels = custom_order)) %>%
  arrange(strain)
# extract matrices

hmo_mat_8h <- as.matrix(HMO_8h_ut[,2:ncol(HMO_8h_ut)])
rownames(hmo_mat_8h) <- pull(HMO_8h_ut, strain)
desired_order <- c("LNT", "LNnT", "LNH", "2'FL", "3FL", "LDFT",
                   "LNFP I", "LNFP II", "LNFP III", "DFLNT", "FLNH",
                   "DFLNH", "3'SL", "6'SL", "LSTb", "LSTc", "DSLNT",
                   "DSLNH", "FDSLNH", "total", "total_fuc", "total_sial")
hmo_mat_8h <- hmo_mat_8h[, desired_order]

# add a coloring function
col_fun <- colorRamp2(c(0, 0.125, 0.25, 0.375, 0.5, 0.625, 0.75, 0.875, 1),
                      c("#ffffff", "#DEEBF7", "#C6DBEF", "#9ECAE1", "#6BAED6",
                        "#4292C6", "#2171B5", "#08519C", "#08306B"))
```

```r
lgd <- Legend(col_fun = col_fun,
              title = "% of predicted utilizers",
              direction = "horizontal",
              title_position = "lefttop")

# specify the name of the output file
pdf("results/HMO_utilization/heatmap_8h.pdf", width=15, height=4)
ht_hmo_8h <- ComplexHeatmap::Heatmap(hmo_mat_8h,
                        name = "HMO consumption at 8h",
                        col = col_fun,
                        cluster_rows = FALSE,
                        cluster_columns = FALSE,
                        rect_gp = gpar(col = "grey", lwd = 0.05),
                        heatmap_legend_param = list(
                          col_fun = col_fun,
                          at = c(0, 0.25, 0.5, 0.75, 1),
                          title = "% of utilized HMOs",
                          direction = "horizontal",
                          title_position = "topcenter",
                          border = "black",
                          legend_width = unit(40, "mm"))
                        )
draw(ht_hmo_8h)
# save the heatmap to a file
dev.off()
```

```
## pdf
##   2
```

```r
# draw the heatmap again to show it in the compiled markdown file
draw(ht_hmo_8h)
```

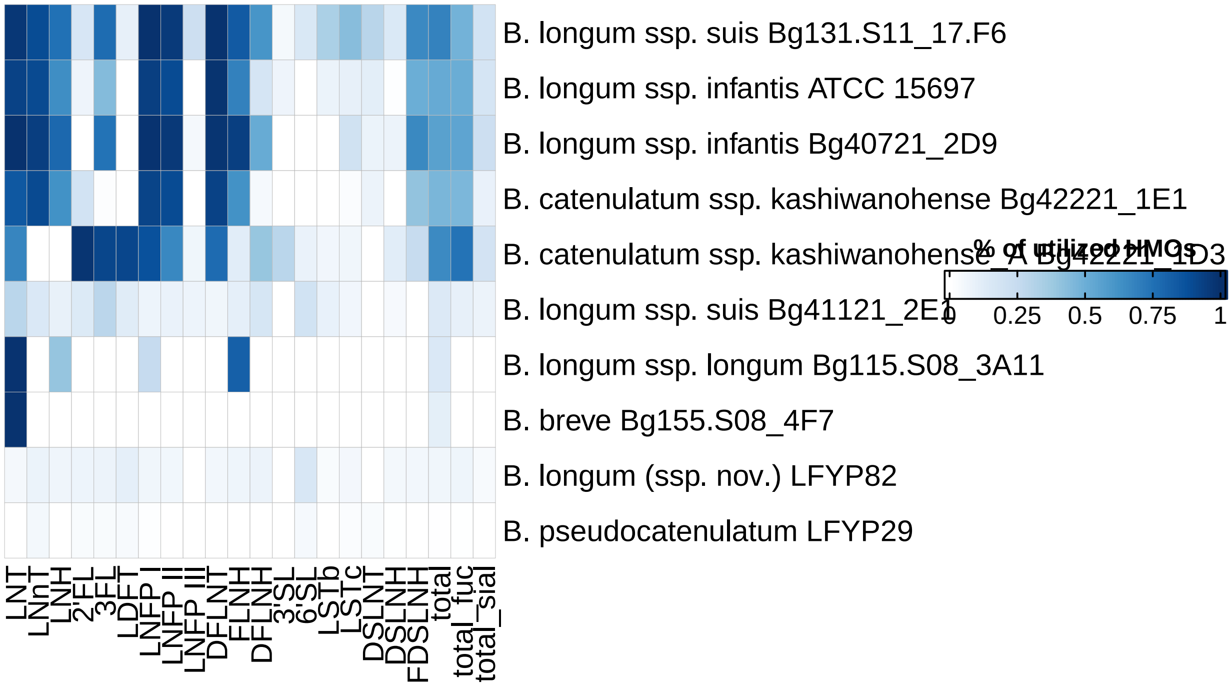

##### 12.2.3 Integration of growth and HMO consumption data

We first use the gcplyr package to calculate growth parameters (maximum OD600, Area Under the Curve (AUC), and growth rate).

```
# read_data
pHMO_data <- "data/growth/formatted_selected/pHMO/growth_pHMO_24h.txt"
pHMO_annotation <- "data/growth/formatted_selected/pHMO/growth_pHMO_24h_ann.txt"
trim_at_time <- 24 # measurements until [input] hours
blank_wells_HMO <- c("blank_1","blank_2","blank_3") # specify blank wells

# read a file with measurements
d_HMO <- read_tsv(pHMO_data) %>%
  filter(time <= trim_at_time)
# calculate mean blank measurements (MRS-AC-Lac without added cells)
blank_HMO <- d_HMO %>%
  dplyr::select(all_of(blank_wells_HMO)) %>%
  rowMeans()
# subtract blank measurements and round OD600 values to 3 digits
d_bl_HMO <- d_HMO %>%
  mutate(across(2:last_col(), ~ .x - blank_HMO)) %>%
  mutate(across(2:last_col(), ~ round(.x, 3)))
# read files with plate annotations
ann <- read_tsv(pHMO_annotation)
d_bl_long_HMO <- d_bl_HMO %>%
  gather(., well, od, -time)
# create annotated tables
d_bl_long_HMO_ann <- left_join(d_bl_long_HMO, ann, by="well") %>%
  filter(carbon_source != "blank")
```

```r
# derivatives and doubling time
ex_dat_mrg <- mutate(group_by(d_bl_long_HMO_ann, well, strain),
                     deriv_percap5 = calc_deriv(x = time, y = od,
                                                percapita = TRUE, blank = 0,
                                                window_width_n = 5, trans_y = "log"),
                     doub_time = doubling_time(y = deriv_percap5)) %>%
              mutate(deriv_percap5_filtered = ifelse(od > 0.05, deriv_percap5, NA))

# calculate maximum density, AUC, and growth rate
ex_dat_mrg_sum <- summarize(group_by(ex_dat_mrg, strain, well),
            max_dens = max_gc(od, na.rm = TRUE),
            auc = auc(x = time, y = od),
            max_growth_rate = max(deriv_percap5_filtered, na.rm = TRUE))
```

PCA of growth and HMO consumption data.

```r
# read data frames with measuerements
auc <- ex_dat_mrg_sum
hmo <- HMO_24h_ut

# aggregate growth data per strain (mean across replicates)
auc_summary <- auc %>%
  group_by(strain) %>%
  summarize(
    auc = mean(auc, na.rm = TRUE),
    max_dens = mean(max_dens, na.rm = TRUE),
    max_growth_rate = mean(max_growth_rate, na.rm = TRUE)
  )

# merge growth and HMO data based on strain
merged_data <- inner_join(auc_summary, hmo, by = "strain")

# remove strain column for PCA
pca_data <- merged_data %>% select(-strain)
strain  <- merged_data$strain
pca.res <- prcomp(pca_data, scale.=T, retx=T)

# plot the results
autoplot(pca.res, merged_data, color = "strain",
         size = 6) +
  theme_bw() +
  coord_fixed() +
  theme(plot.title = element_text(face = "bold"))
```

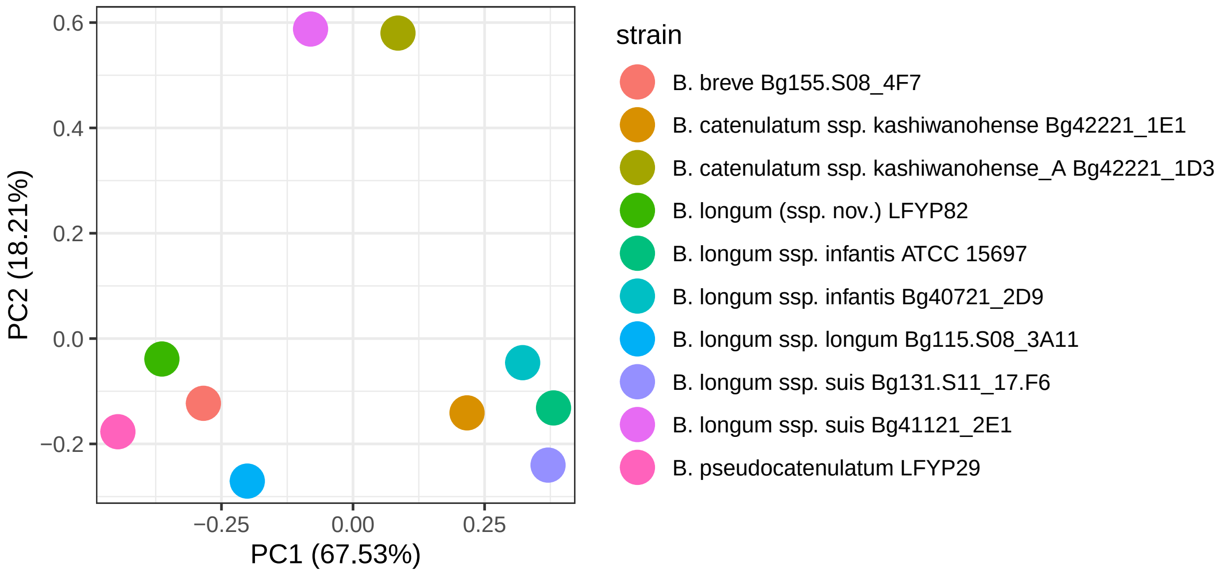

```
# save the plot
ggsave("results/HMO_utilization/PCA.pdf", device = "pdf", width = 10, height = 10)
```

---

### 13 Analysis of RNA-seq data

#### 13.1 Processing of raw FASTQ files and read mapping

The code chuck below describes the processing of raw FASTQ files and mapping reads to the *Bifidobacterium catenulatum* subsp. *kashiwanohense* Bg42221_1E1 transcriptome. The following software is required (can be installed in one conda/mamba environment):

1. FastQC (v0.11.9)
2. Cutadapt (v4.1)
3. Bowtie2 (v 2.4.5)
4. Kallisto (v0.48)
5. MultiQC (v1.13)
6. Parallel (v20220722)

**Note**: due to size limitation, raw FASTQ files could not be stored in the GitHub repo. Thus, you will need to download the FASTQ files from the Gene Expression Omnibus, under **accession**. Put downloaded fastq.gz files to `data/rnaseq/fastq/`.

The reference FASTA files used for building Bowtie2 and Kallisto indeces can be found in `data/rnaseq/refs/`.

**Note**: exact file names (without the .fastq.qz extension) should be entered into `data/rnaseq/runids.txt`.

Alternatively, you can run `code/qc_readmapping.sh` instead of the code chunk below.

Summary of the script:

1. Quality control of raw reads was carried out using FastQC
2. Quality trimming and removal of Illumina sequencing adapters and short reads (< 20 bp) were performed via Cutadapt
3. Reads were mapped against rRNA and tRNA gene sequences extracted from the *Bifidobacterium catenulatum* subsp. *kashiwanohense* Bg42221_1E1 genome (GenBank accession no. ) using Bowtie2. Unmapped (filtered) reads were saved and used further
4. Filtered reads were mapped to the *Bifidobacterium catenulatum* subsp. *kashiwanohense* Bg_1E1 transcriptome using Kallisto
5. The quality of raw/filtered reads, as well as the results of Bowtie2/Kallisto mapping were summarized in `data/rnaseq/multiqc_report.html` generated via MultiQC

```
source ~/.bash_profile
echo $BASH_VERSION
#set -ex

####################
## SOFTWARE SETUP ##
####################
# required tools: fastqc (v0.11.9), cutadapt (v4.1), bowtie2 (v2.4.5), kallisto (v0.48)
# multiqc (v1.13), and parallel (v20220722)
# set the name of the environment with the tools
environment_name="transcriptomics"
# activate conda environment with the required tools
eval "$(command conda 'shell.bash' 'hook' 2> /dev/null)" # initializes conda in sub-shell
conda activate ${environment_name}
conda info|egrep "conda version|active environment"

################
## USER INPUT ##
################
# fastq files should be in data/rnaseq/fastq/
# txt file containing names of fastq files (without the fastq.gz extension)
sample_names="data/rnaseq/runids.txt"
# fasta file containing sequences of rRNA and tRNA genes
rRNA_tRNA_fasta="data/rnaseq/refs/Bc_kashiwanohense_Bg42221_1E1_RNA.fasta"
# fasta file containing the whole transcriptome
transcriptome_fasta="data/rnaseq/refs/Bc_kashiwanohense_Bg42221_1E1.fasta"
# name of the bowtie2 index
bowtie2_index_name="data/rnaseq/refs/Bc_kashiwanohense_Bg42221_1E1_rRNA_tRNA"
# name of the kallisto index
kallisto_index_name="data/rnaseq/refs/Bc_kashiwanohense_Bg42221_1E1_transcriptome.index"

# create directories
echo "Creating directories"
mkdir -p data/rnaseq/qc1 # qc results for raw reads
mkdir -p data/rnaseq/qc2 # qc results for filtered reads
mkdir -p data/rnaseq/fq_trim # trimmed reads
mkdir -p data/rnaseq/fq_filt # filtered reads
mkdir -p data/rnaseq/sam # sam files produced during bowtie2 alignment; will be deleted
mkdir -p data/rnaseq/kallisto # kallisto mapping results

# run fastqc on raw reads
echo "Running FastQC on raw reads"
cat ${sample_names} | parallel "fastqc \
```

```
data/rnaseq/fastq/{}.fastq.gz \
--outdir data/rnaseq/qc1"

# trim adapters with cutadapt
echo "Trimming adapters with Cutadapt"
cat ${sample_names} | parallel "cutadapt \
--nextseq-trim=20 \
-m 20 \
-a AGATCGGAAGAGCACACGTCTGAACTCCAGTC \
-o data/rnaseq/fq_trim/{}.fastq.gz data/rnaseq/fastq/{}.fastq.gz \
&> data/rnaseq/fq_trim/{}.fastq.qz.log"

# filter reads mapped to rRNA and tRNA
echo "Filtering reads that map to rRNA and tRNA using Bowtie2"
## build bowtie2 index
bowtie2-build ${rRNA_tRNA_fasta} \
${bowtie2_index_name}
## align reads via bowtie2; save ones that did not align to a separate file
cat ${sample_names} | \
parallel "bowtie2 -x ${bowtie2_index_name} \
-U data/rnaseq/fq_trim/{}.fastq.gz \
-S data/rnaseq/sam/{}.sam \
--un data/rnaseq/fq_filt/{}.fastq \
&> data/rnaseq/fq_filt/{}_bowtie2.log"
cat ${sample_names} | parallel "gzip data/rnaseq/fq_filt/{}.fastq"

# run fastqc on filtered reads
echo "Running FastQC on filtered reads"
cat ${sample_names} | parallel "fastqc \
data/rnaseq/fq_filt/{}.fastq.gz \
--outdir data/rnaseq/qc2"

# pseudolalign reads to transcriptome
echo "Mapping reads to the transcriptome via Kallisto"
## build kallisto index
kallisto index -i ${kallisto_index_name} \
${transcriptome_fasta}
## map reads to indexed reference via kallisto
cat ${sample_names} | parallel "kallisto quant \
-i ${kallisto_index_name} \
-o data/rnaseq/kallisto/{} \
--single \
--rf-stranded \
-l 150 \
-s 20 \
data/rnaseq/fq_filt/{}.fastq.gz \
&> data/rnaseq/kallisto/{}_kallisto.log"

# run multiqc
echo "Running MultiQC"
export LC_ALL=en_US.utf-8
export LANG=en_US.utf-8
multiqc -d data/rnaseq -o data/rnaseq
```

```
# remove directories with intermediate files
echo "Removing directories"
rm -rf data/rnaseq/qc1
rm -rf data/rnaseq/qc2
rm -rf data/rnaseq/fq_trim
rm -rf data/rnaseq/fq_filt
rm -rf data/rnaseq/sam
```

#### 13.2 Importing count data into R

TxImport was used to read Kallisto output into the R environment.

*Note*: before running the code, double-check that file names in the file_names column in `data/rnaseq/studydesign.txt` are identical to file names in `data/rnaseq/runids.txt`.

```
# read the study design file
targets <- read_tsv("data/rnaseq/studydesign.txt")
# set file paths to Kallisto output folders with quantification data
files <- file.path("data/rnaseq/kallisto", targets$file_name, "abundance.tsv")
# check that all output files are present
all(file.exists(files))
```

```
## [1] TRUE
```

```
# use 'tximport' to import Kallisto output into R
txi_kallisto <- tximport(files,
                         type = "kallisto",
                         txOut = TRUE, # import at transcript level
                         countsFromAbundance = "lengthScaledTPM")

# capture variables of interest from the study design
condition <- as.factor(targets$condition)
condition <- factor(condition, levels = c("Glc", "Lac", "LNT", "XGL"))
batch <- as.factor(targets$batch)
# capture sample labels for later use
sampleLabels <- targets$sample

# create a table with raw counts for GEO submission
raw_counts <- as_tibble(txi_kallisto$counts, rownames = "locus_tag")
colnames(raw_counts) <- c("geneID", sampleLabels)
write_tsv(raw_counts, "results/rnaseq/tables/Arzamasov_raw_count_matrix.txt")

# use the gt package to produce the study design table
gt(targets) %>%
  cols_align(
    align = "left",
    columns = everything()
  )
```

| sample | file_name | condition | batch |
|---|---|---|---|
| Glc1 | Glc1 | Glc | 1 |
| Glc2 | Glc2 | Glc | 1 |
| Glc3 | Glc3 | Glc | 1 |
| Lac1 | Lac1 | Lac | 2 |
| Lac2 | Lac2 | Lac | 2 |
| Lac3 | Lac3 | Lac | 2 |
| LNT1 | LNT1 | LNT | 2 |
| LNT2 | LNT2 | LNT | 2 |
| LNT3 | LNT3 | LNT | 2 |
| XGL1 | XGL1 | XGL | 1 |
| XGL2 | XGL2 | XGL | 1 |
| XGL3 | XGL3 | XGL | 1 |

##### 13.3 Filtering and normalization

```r
# extract counts
myDGEList <- DGEList(txi_kallisto$counts)
# plot unfiltered, non-normalized CPM
p1 <- profile(myDGEList, sampleLabels, "Unfiltered, non-normalized")
# filter counts
cpm <- cpm(myDGEList)
keepers <- rowSums(cpm>1)>=3 # only keep genes that have cpm>1 (== not zeroes)
# in more than 3 samples (minimal group size)
myDGEList.filtered <- myDGEList[keepers,]
# plot filtered, non-normalized CPM
p2 <- profile(myDGEList.filtered, sampleLabels, "Filtered, non-normalized")
# normalize counts via the TMM method implemented in edgeR
myDGEList.filtered.norm <- calcNormFactors(myDGEList.filtered, method = "TMM")
# plot filtered, normalized CPM
p3 <- profile(myDGEList.filtered.norm, sampleLabels, "Filtered, TMM normalized")
# compare distributions of the CPM values
plot_grid(p1, p2, p3, labels = c('A', 'B', 'C'), label_size = 12)
```

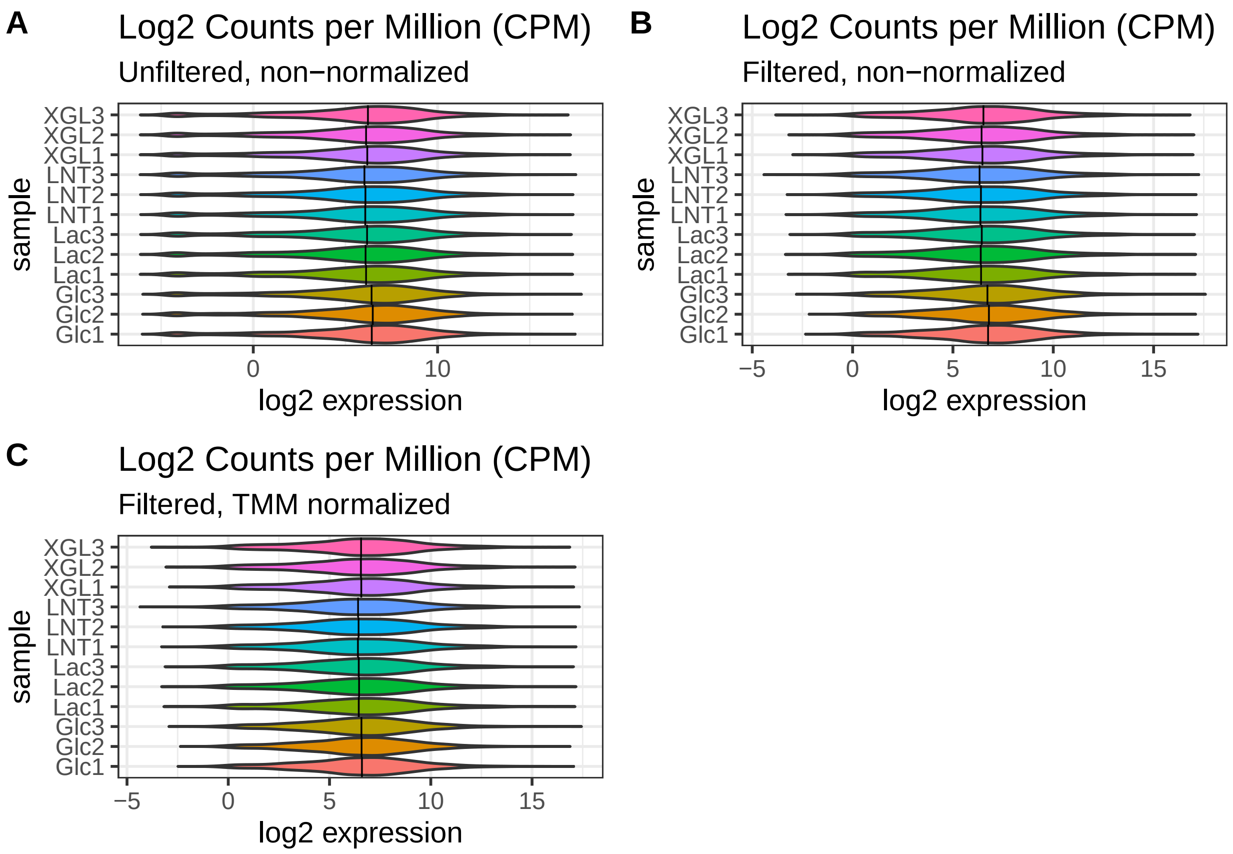

Filtering was carried out to remove lowly expressed genes. Genes with less than 1 count per million (CPM) in at least 3 or more samples were filtered out. This procedure reduced the number of genes from **2099** to **1927**. In addition, the TMM method was used for between-sample normalization.

#### 13.4 PCA plot

Principal Component Analysis (PCA) plots reduce complex datasets to a 2D representation where each axis represents a source of variance (known or unknown) in the dataset. Principal Component 1 (PC1; X-axis) accounted for >43% of the variance in the data. PC2 (Y-axis) accounted for >36% of the variance in the data.

```
# running PCA
log2.cpm.filtered.norm <- cpm(myDGEList.filtered.norm, log=TRUE)
pca.res <- prcomp(t(log2.cpm.filtered.norm), scale.=F, retx=T)
pc.var <- pca.res$sdev^2 # sdev^2 captures eigenvalues from the PCA result
pc.per <- round(pc.var/sum(pc.var)*100, 1) # calculate percentage of the total variation
# explained by each eigenvalue
# converting PCA result into a tibble for plotting
pca.res.df <- as_tibble(pca.res$x)

# plotting PCA
ggplot(pca.res.df) +
  aes(x=PC1, y=PC2, label=sampleLabels, fill = condition) +
  geom_point(size=4, shape = 21) +
  scale_fill_manual(name = "Carbon source",
```

```
                      breaks=c("Glc","Lac", "LNT", "XGL"),
                      values=c("#ed5564", "#ffce54", "#a0d568", "#4fc1e8"),
                      labels=c("Glucose","Lactose", "Lacto-N-tetraose", "Xyloglucan")) +
  guides(fill = guide_legend(override.aes=list(shape=21))) +
  xlab(paste0("PC1 (",pc.per[1],"%",")")) +
  ylab(paste0("PC2 (",pc.per[2],"%",")")) +
  labs(title= "PCA of Bc. kashiwanohense Bg42221_1E1") +
  coord_fixed() +
  theme_bw() +
  theme(plot.title = element_text(face="bold"))
```

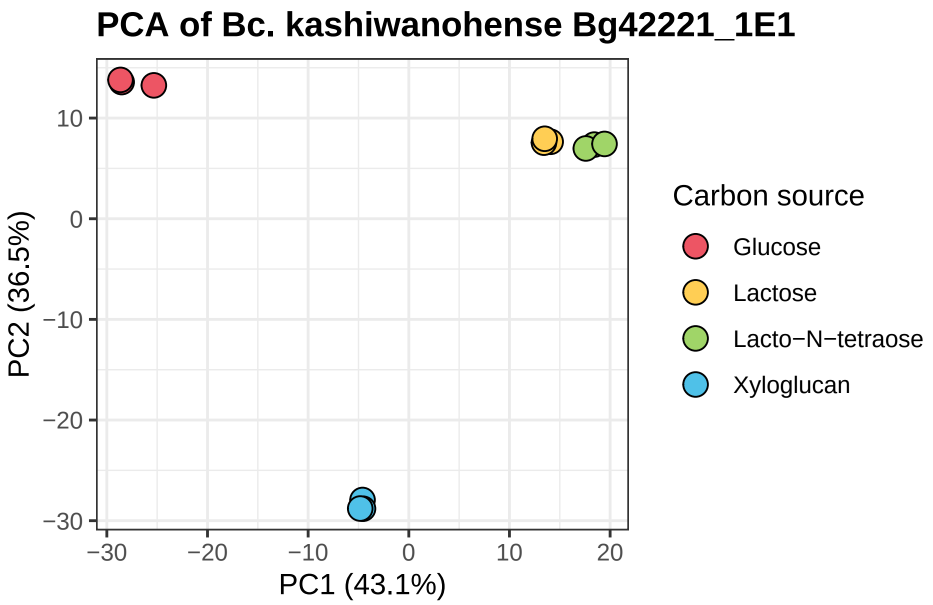

```
# save the figure
ggsave("results/rnaseq/figures/PCA.pdf", device = "pdf", width = 5, height = 5)
```

```
# PCA 'small multiples' chart
# view PCA loadings to understand impact of each sample on each principal component
pca.res.df <- pca.res$x[,1:4] %>%
  as_tibble() %>%
  add_column(sample = sampleLabels,
             group = condition)
```

```r
pca.pivot <- pivot_longer(pca.res.df, # data frame to be pivoted
                          cols = PC1:PC4, # column names to be stored as a SINGLE variable
                          names_to = "PC", # name of that new variable (column)
                          values_to = "loadings") # name of new variable (column) storing all the value

# plot the chart
ggplot(pca.pivot) +
  aes(x=sample, y=loadings, fill=group) +
  geom_bar(stat="identity") +
  facet_wrap(~PC) +
  scale_fill_manual(name = "Carbon source",
                    breaks=c("Glc","Lac", "LNT", "XGL"),
                    values=c("#ed5564", "#ffce54", "#a0d568", "#4fc1e8"),
                    labels=c("Glucose","Lactose", "Lacto-N-tetraose", "Xyloglucan")) +
  labs(title="PCA 'small multiples' plot") +
  theme_bw() +
  coord_flip()
```

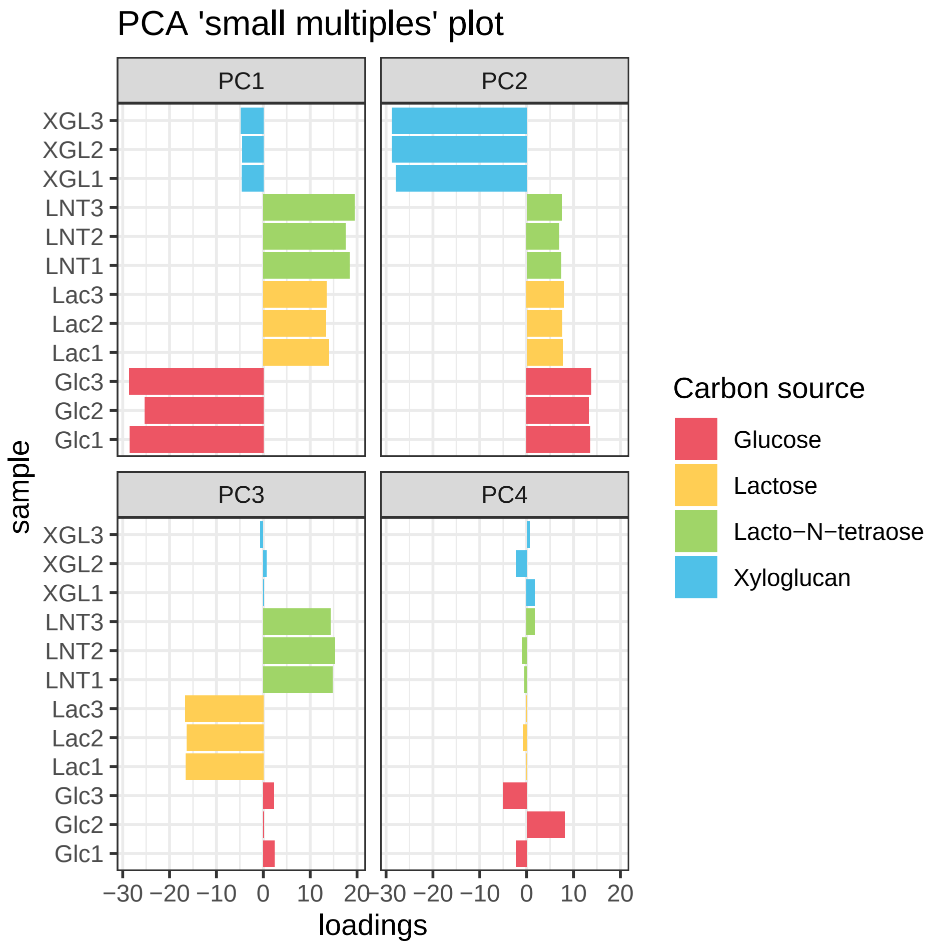

```r
# save the figure
ggsave("results/rnaseq/figures/PCA_small_multiples.pdf", device = "pdf", width = 8, height = 5)
```

##### 13.5 Differentially expressed genes

To identify differentially expressed genes (DEGs), precision weights were first applied to each gene based on its mean-variance relationship using VOOM. Linear modeling and bayesian stats were employed using Limma to find genes that were up- or down-regulated more than 4-fold at false-discovery rate (FDR) of 0.01.

```
# setting up model matrix without intercept
design <- model.matrix(~0 + condition)
colnames(design) <- levels(condition)
# using VOOM function from Limma package to apply precision weights to each gene
v.DEGList.filtered.norm <- voom(myDGEList.filtered.norm, design, plot = TRUE)
```

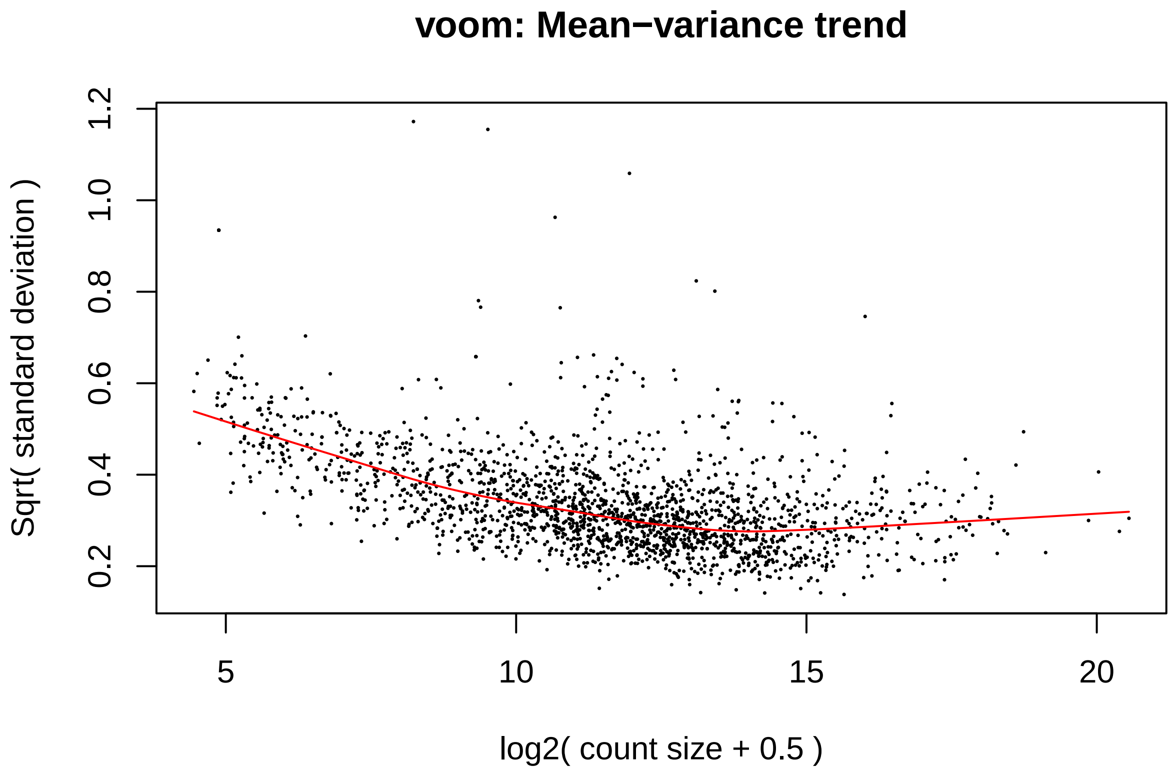

```
fit <- lmFit(v.DEGList.filtered.norm, design)
# setting up contrast matrix for pairwise comparisons of interest
contrast.matrix <- makeContrasts(XGL = XGL - Lac,
                                 LNT = LNT - Lac,
                                 levels=design)
fits <- contrasts.fit(fit, contrast.matrix)
# extracting stats
ebFit <- eBayes(fits)
```

DEGs were annotated based on a RAST-annotated version of the *Bifidobacterium catenulatum* subsp. *kashiwanohense* Bg42221_1E1 genome, which was additionally subjected to extensive manual curation in mcSEED, a private clone of the publicly available SEED platform. The manual curation focused on annotating genes encoding functional roles (transporters, glycoside hydrolases, downstream catabolic enzymes, transcriptional regulators) involved in carbohydrate metabolism.

DEGs: *Bifidobacterium catenulatum* subsp. *kashiwanohense* Bg42221_1E1 grown in MRS-AC-XGL vs. MRS-AC-Lac

```
# create a master annotation table
seed.ann <- read_tsv('data/rnaseq/annotation/mcSEED_annotations.txt')
corr <- read_tsv('data/rnaseq/annotation/locus_tag_comparison.txt')
final.ann <- right_join(seed.ann, corr, by = c('seed_id' = 'seed_id')) %>%
  dplyr::select(seed_id, locus_tag, annotation)

# annotate DEGs
# xyloglucan vs lactose
myTopHits.XGLvsLac <- topTable(ebFit, adjust ="BH", coef=1, number=2600, sort.by="logFC")
deg_list(myTopHits.XGLvsLac, -2, 2, 0.01, "results/rnaseq/tables/table_S15A.txt")
```

DEGs: *Bifidobacterium catenulatum* subsp. *kashiwanohense* Bg42221_1E1 grown in MRS-AC-LNT vs. MRS-AC-Lac

```
# annotate DEGs
# lacto-N-tetraose vs lactose
myTopHits.LNTvsLac <- topTable(ebFit, adjust ="BH", coef=2, number=2600, sort.by="logFC")
deg_list(myTopHits.LNTvsLac, -2, 2, 0.01, "results/rnaseq/tables/table_S15B.txt")
```

#### 13.6 Volcano plot: *Bifidobacterium catenulatum* subsp. *kashiwanohense* Bg42221_1E1 grown in MRS-AC-XGL vs MRS-AC-Lac

Volcano plots are convenient ways to represent gene expression data because they combine magnitude of change (X-axis) with significance (Y-axis). Since the Y-axis is the inverse log10 of the adjusted Pvalue, higher points are more significant. In the case of this particular plot, there are many genes in the upper right of the plot, which represent genes that are significantly **upregulated** in when the bacterium is grown in MRS-AC-XGL, compared MRS-AC-Lac.

```
# list stats for all genes in the dataset to be used for making the volcano plot
myTopHits <- topTable(ebFit, adjust ="BH", coef=1, number=3000, sort.by="logFC")
myTopHits.df <- myTopHits %>%
  as_tibble(rownames = "geneID")
# select only genes with significant logFC and adj.P.Val for the heatmap
myTopHits.df.de <- subset(myTopHits.df, (logFC > 2 | logFC < 2) & adj.P.Val < 0.01)
# create vectors containing locus_tags of genes constituting reconstructed regulons
targets.XglR <- c("BcK1E1_01925", "BcK1E1_01926", "BcK1E1_01927", "BcK1E1_01944",
                  "BcK1E1_01943", "BcK1E1_01942", "BcK1E1_01941", "BcK1E1_01940",
                  "BcK1E1_01939", "BcK1E1_01950", "BcK1E1_01949", "BcK1E1_01948",
                  "BcK1E1_01947", "BcK1E1_01946", "BcK1E1_01945", "BcK1E1_01951")
targets.XylR <- c("BcK1E1_00526", "BcK1E1_00535", "BcK1E1_00536")
targets.XosR <- c("BcK1E1_00357", "BcK1E1_00358", "BcK1E1_00359", "BcK1E1_00360",
                  "BcK1E1_00361", "BcK1E1_00362", "BcK1E1_00363", "BcK1E1_00364",
                  "BcK1E1_00365", "BcK1E1_00366", "BcK1E1_00367", "BcK1E1_00528",
                  "BcK1E1_00529", "BcK1E1_00530", "BcK1E1_00531", "BcK1E1_00532", "BcK1E1_00533")
targets.XosR2 <- c("BcK1E1_01192", "BcK1E1_01191", "BcK1E1_01190", "BcK1E1_01189",
                   "BcK1E1_01188", "BcK1E1_01187", "BcK1E1_01186", "BcK1E1_01185")
# subset data based on targets.XglR/xylR/xosR/xosR2
myTopHits.XglR <- subset(myTopHits.df, geneID %in% targets.XglR)
myTopHits.XylR <- subset(myTopHits.df, geneID %in% targets.XylR)
```

```
myTopHits.XosR <- subset(myTopHits.df, geneID %in% targets.XosR)
myTopHits.XosR2 <- subset(myTopHits.df, geneID %in% targets.XosR2)
# subset data labels (genes in regulons)
## XglR
myTopHits.df$XglR <- myTopHits.df$geneID
myTopHits.XglR_selected <- myTopHits.df$XglR %in% myTopHits.XglR$geneID
myTopHits.df$XglR[!myTopHits.XglR_selected] <- NA
## XylR
myTopHits.df$XylR <- myTopHits.df$geneID
myTopHits.XylR_selected <- myTopHits.df$XylR %in% myTopHits.XylR$geneID
myTopHits.df$XylR[!myTopHits.XylR_selected] <- NA
## XosR
myTopHits.df$XosR <- myTopHits.df$geneID
myTopHits.XosR_selected <- myTopHits.df$XosR %in% myTopHits.XosR$geneID
myTopHits.df$XosR[!myTopHits.XosR_selected] <- NA
## XosR
myTopHits.df$XosR2 <- myTopHits.df$geneID
myTopHits.XosR2_selected <- myTopHits.df$XosR2 %in% myTopHits.XosR2$geneID
myTopHits.df$XosR2[!myTopHits.XosR2_selected] <- NA

# create the volcano plot
ggplot(myTopHits.df) +
  aes(y=-log10(adj.P.Val), x=logFC, text = paste("Symbol:", geneID)) +
  geom_point(size=3, shape = 16, color="black", alpha=.3) +
  geom_point(mapping=NULL, myTopHits.XglR, size = 3, shape = 16, color= "#ee7942",
             inherit.aes = TRUE) +
  geom_point(mapping=NULL, myTopHits.XylR, size = 3, shape = 16, color= "#00a69c",
             inherit.aes = TRUE) +
  geom_point(mapping=NULL, myTopHits.XosR, size = 3, shape = 16, color= "#f8b4ff",
             inherit.aes = TRUE) +
  geom_point(mapping=NULL, myTopHits.XosR2, size = 3, shape = 16, color= "#9ac946",
             inherit.aes = TRUE) +
  geom_text_repel(aes(label = XglR),
                  size = 1, fontface=2, color="black", min.segment.length = 0,
                  seed = 42, box.padding = 0.5, max.overlaps = 100) +
  geom_text_repel(aes(label = XylR),
                  size = 1, fontface=2, color="black", min.segment.length = 0,
                  seed = 42, box.padding = 0.5, max.overlaps = 100) +
  geom_text_repel(aes(label = XosR),
                  size = 1, fontface=2, color="black", min.segment.length = 0,
                  seed = 42, box.padding = 0.5, max.overlaps = 100) +
  geom_text_repel(aes(label = XosR2),
                  size = 1, fontface=2, color="black", min.segment.length = 0,
                  seed = 42, box.padding = 0.5, max.overlaps = 100) +
  geom_hline(yintercept = -log10(0.01), linetype="longdash", colour="grey", size=0.6) +
  geom_vline(xintercept = 2, linetype="longdash", colour="grey", size=0.6) +
  geom_vline(xintercept = -2, linetype="longdash", colour="grey", size=0.6) +
  annotate("text", x=-6, y=-log10(0.01)+0.3,
           label=paste("Padj<0.01"), size=5, fontface="bold") +
  scale_x_continuous(limits=c(-5,10), breaks = -5:10) +
  labs(title="Volcano plot",
       subtitle = "B. catenulatum subsp. kashiwanohense Bg42221_1E1 grown in MRS-AC-XGL vs. MRS-AC-Lac"
  theme(plot.title = element_text(face="bold")) +
```

```r
  theme_cowplot()
```

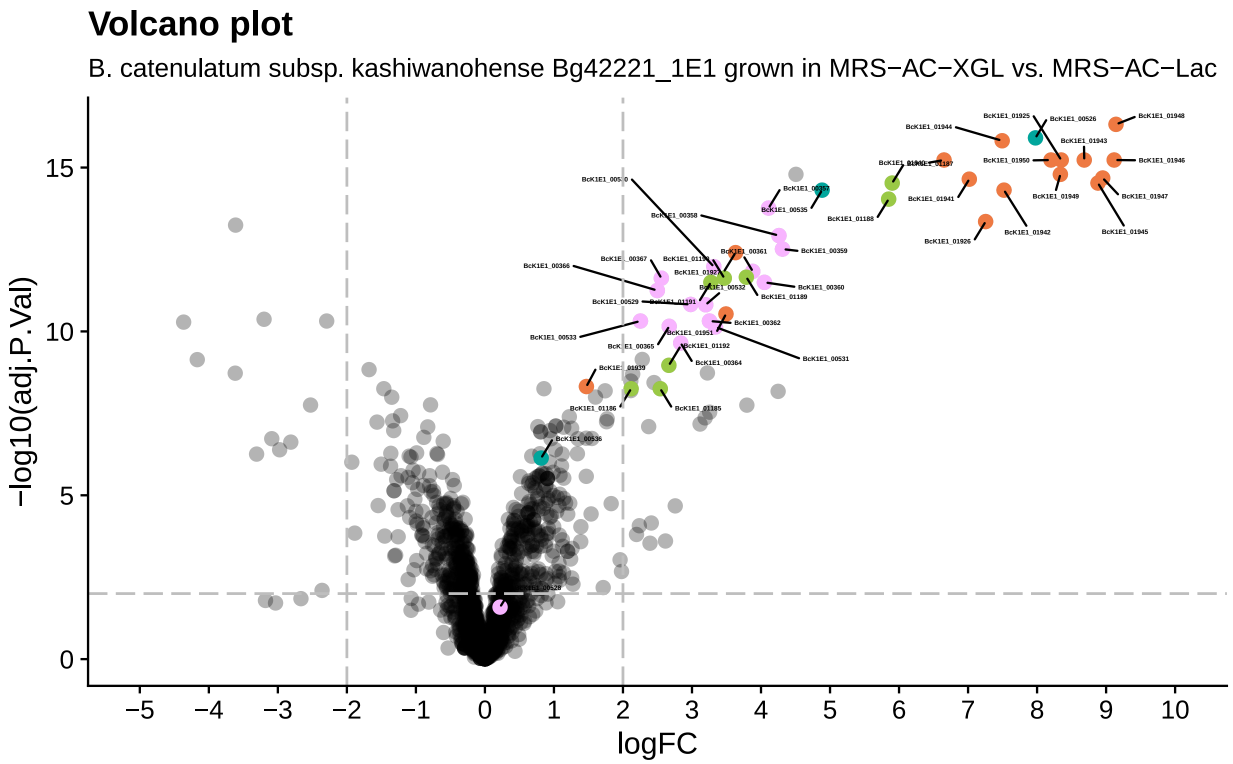

```r
# save the figure
ggsave("results/rnaseq/figures/XGL_vs_Lac.pdf", device = "pdf", width = 8, height = 5)
```

#### 13.7 Volcano plot: *Bifidobacterium catenulatum* subsp. *kashiwanohense* Bg42221_1E1 grown in MRS-AC-LNT vs MRS-AC-Lac

```r
# listing stats for all genes in the dataset to be used for making volcano plot
myTopHits2 <- topTable(ebFit, adjust ="BH", coef=2, number=3000, sort.by="logFC")
myTopHits.df2 <- myTopHits2 %>%
  as_tibble(rownames = "geneID")
# select only genes with significant logFC and adj.P.Val for the heatmap
myTopHits.df2.de <- subset(myTopHits.df2, (logFC > 2 | logFC < 2) & adj.P.Val < 0.01)
# create a vector containing locus_tags of genes predicted to be in the HMO cluster
targets.NagR <- c("BcK1E1_00572", "BcK1E1_01910", "BcK1E1_01909", "BcK1E1_01908",
                  "BcK1E1_01907", "BcK1E1_01911", "BcK1E1_01907", "BcK1E1_02020",
                  "BcK1E1_02019", "BcK1E1_02018", "BcK1E1_02017", "BcK1E1_02016",
                  "BcK1E1_02034", "BcK1E1_02033", "BcK1E1_02032", "BcK1E1_02031",
                  "BcK1E1_02030", "BcK1E1_02029", "BcK1E1_02028", "BcK1E1_02027",
                  "BcK1E1_02026", "BcK1E1_02025", "BcK1E1_02024", "BcK1E1_02023",
                  "BcK1E1_02022", "BcK1E1_02021", "BcK1E1_02035", "BcK1E1_02036",
                  "BcK1E1_02037", "BcK1E1_02038", "BcK1E1_02039")
# subset data based on targets.nagR
myTopHits.NagR <- subset(myTopHits.df2, geneID %in% targets.NagR)
```

```r
# subset data labels (genes in regulons)
myTopHits.df2$NagR <- myTopHits.df2$geneID
myTopHits.NagR_selected <- myTopHits.df2$NagR %in% myTopHits.NagR$geneID
myTopHits.df2$NagR[!myTopHits.NagR_selected] <- NA

# create the volcano plot
ggplot(myTopHits.df2) +
  aes(y=-log10(adj.P.Val), x=logFC, text = paste("Symbol:", geneID)) +
  geom_point(size=3, shape = 16, color="black", alpha=.3) +
  geom_point(mapping=NULL, myTopHits.NagR, size = 3, shape = 16, color= "#cbbedd",
           inherit.aes = TRUE) +
  geom_text_repel(aes(label = NagR),
                  size = 1, fontface=2, color="black", min.segment.length = 0,
                  seed = 42, box.padding = 0.5, max.overlaps = 100) +
  geom_hline(yintercept = -log10(0.01), linetype="longdash", colour="grey", size=0.6) +
  geom_vline(xintercept = 2, linetype="longdash", colour="grey", size=0.6) +
  geom_vline(xintercept = -2, linetype="longdash", colour="grey", size=0.6) +
  annotate("text", x=-6, y=-log10(0.01)+0.3,
          label=paste("Padj<0.01"), size=5, fontface="bold") +
  scale_x_continuous(limits=c(-4,7), breaks = -4:7) +
  labs(title="Volcano plot",
      subtitle = "B. catenulatum subsp. kashiwanohense Bg42221_1E1 grown in MRS-AC-LNT vs. MRS-AC-Lac")
  theme(plot.title = element_text(face="bold")) +
  theme_cowplot()
```

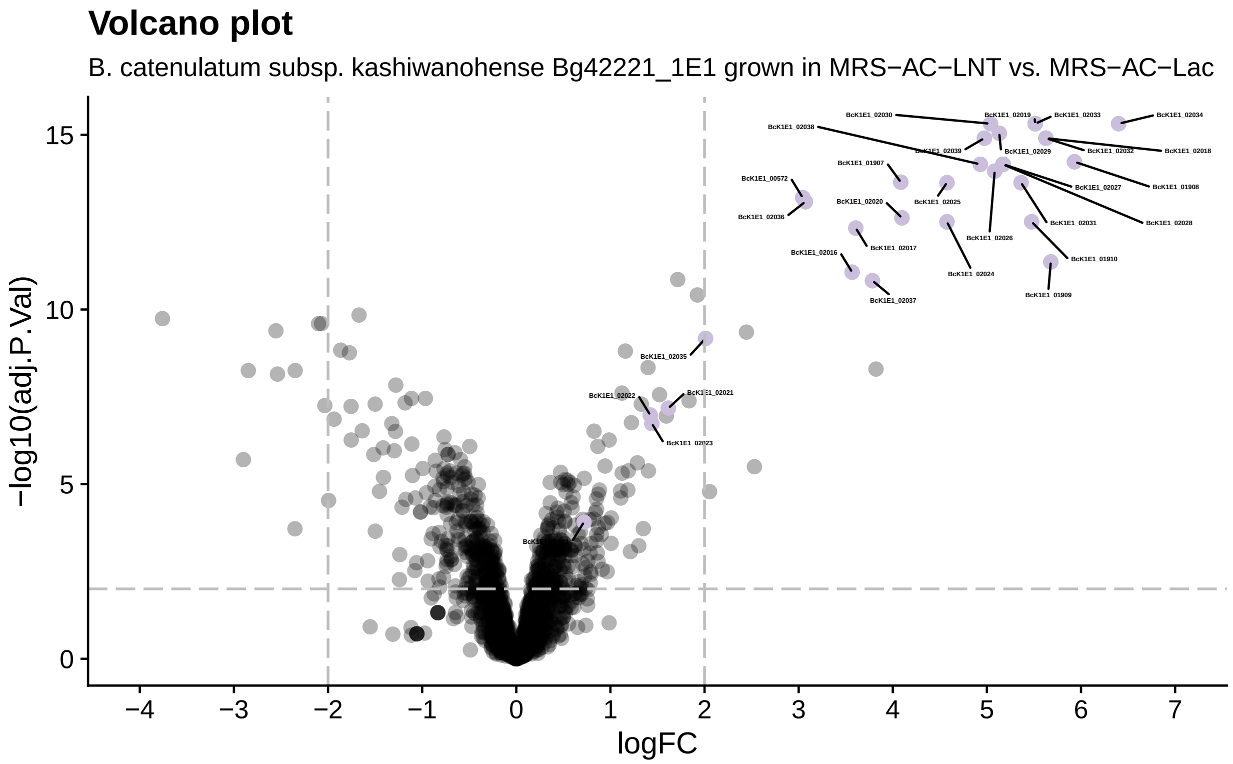

```r
# save the figure
ggsave("results/rnaseq/figures/LNT_vs_Lac.pdf", device = "pdf", width = 8, height = 5)
```

### 14 Session info

The output from running 'sessionInfo' is shown below and details all packages necessary to reproduce the results in this report.

```
sessionInfo()
```

```
## R version 4.3.2 (2023-10-31)
## Platform: aarch64-apple-darwin20 (64-bit)
## Running under: macOS 15.3.1
##
## Matrix products: default
## BLAS:   /Library/Frameworks/R.framework/Versions/4.3-arm64/Resources/lib/libRblas.0.dylib
## LAPACK: /Library/Frameworks/R.framework/Versions/4.3-arm64/Resources/lib/libRlapack.dylib;  LAPACK ve
##
## locale:
## [1] en_US.UTF-8/en_US.UTF-8/en_US.UTF-8/C/en_US.UTF-8/en_US.UTF-8
##
## time zone: America/Los_Angeles
## tzcode source: internal
##
## attached base packages:
## [1] grid      stats     graphics  grDevices utils     datasets  methods
## [8] base
##
## other attached packages:
##  [1] ggordiplots_0.4.3     glue_1.8.0            ggfortify_0.4.17
##  [4] dendextend_1.19.0     phytools_2.4-4        maps_3.4.2.1
##  [7] ape_5.8-1             gcplyr_1.11.0         emmeans_1.10.6
## [10] AER_1.2-14            survival_3.8-3        sandwich_3.1-1
## [13] lmtest_0.9-40         zoo_1.8-12            car_3.1-3
## [16] carData_3.0-5         vegan_2.6-8           lattice_0.22-6
## [19] permute_0.9-7         cowplot_1.1.3         edgeR_4.0.16
## [22] limma_3.58.1          gt_0.11.1             rhdf5_2.46.1
## [25] tximport_1.30.0       ggpubr_0.6.0          circlize_0.4.16
## [28] ggrepel_0.9.6         ggbeeswarm_0.7.2      ComplexHeatmap_2.18.0
## [31] patchwork_1.3.0       readxl_1.4.3          lubridate_1.9.4
## [34] forcats_1.0.0         stringr_1.5.1         dplyr_1.1.4
## [37] purrr_1.0.2           readr_2.1.5           tidyr_1.3.1
## [40] tibble_3.2.1          ggplot2_3.5.1         tidyverse_2.0.0
## [43] pacman_0.5.1          knitr_1.49            tinytex_0.54
## [46] rmarkdown_2.29
##
## loaded via a namespace (and not attached):
##  [1] RColorBrewer_1.1-3      rstudioapi_0.17.1       shape_1.4.6.1
##  [4] magrittr_2.0.3          TH.data_1.1-2           estimability_1.5.1
##  [7] farver_2.1.2            ragg_1.3.3              GlobalOptions_0.1.2
## [10] vctrs_0.6.5             rstatix_0.7.2           htmltools_0.5.8.1
## [13] DEoptim_2.2-8           broom_1.0.7             cellranger_1.1.0
## [16] Rhdf5lib_1.24.2         Formula_1.2-5           igraph_2.1.2
```

```
## [19] lifecycle_1.0.4         iterators_1.0.14        pkgconfig_2.0.3
## [22] Matrix_1.6-4            R6_2.5.1                fastmap_1.2.0
## [25] clue_0.3-66             digest_0.6.37           numDeriv_2016.8-1.1
## [28] colorspace_2.1-1        S4Vectors_0.40.2        textshaping_0.4.1
## [31] labeling_0.4.3          clusterGeneration_1.3.8 timechange_0.3.0
## [34] abind_1.4-8             mgcv_1.9-1              compiler_4.3.2
## [37] bit64_4.6.0-1           withr_3.0.2             doParallel_1.0.17
## [40] backports_1.5.0         optimParallel_1.0-2     viridis_0.6.5
## [43] ggsignif_0.6.4          MASS_7.3-60             rjson_0.2.23
## [46] scatterplot3d_0.3-44    tools_4.3.2             vipor_0.4.7
## [49] beeswarm_0.4.0          quadprog_1.5-8          nlme_3.1-166
## [52] rhdf5filters_1.14.1     cluster_2.1.8           generics_0.1.3
## [55] gtable_0.3.6            tzdb_0.4.0              hms_1.1.3
## [58] xml2_1.3.6              BiocGenerics_0.48.1     foreach_1.5.2
## [61] pillar_1.10.1           vroom_1.6.5             splines_4.3.2
## [64] bit_4.5.0.1             tidyselect_1.2.1        locfit_1.5-9.10
## [67] gridExtra_2.3           IRanges_2.36.0          stats4_4.3.2
## [70] xfun_0.50               expm_1.0-0              statmod_1.5.0
## [73] matrixStats_1.5.0       stringi_1.8.4           yaml_2.3.10
## [76] evaluate_1.0.3          codetools_0.2-20        cli_3.6.3
## [79] systemfonts_1.1.0       xtable_1.8-4            munsell_0.5.1
## [82] Rcpp_1.0.14             coda_0.19-4.1           png_0.1-8
## [85] parallel_4.3.2          phangorn_2.12.1         viridisLite_0.4.2
## [88] mvtnorm_1.3-3           scales_1.3.0            crayon_1.5.3
## [91] combinat_0.0-8          GetoptLong_1.0.5        rlang_1.1.4
## [94] fastmatch_1.1-6         multcomp_1.4-26         mnormt_2.1.1
```
